# An endogenous glucocorticoid-cytokine signaling circuit promotes CD8^+^ T cell dysfunction in the tumor microenvironment

**DOI:** 10.1101/799759

**Authors:** Nandini Acharya, Asaf Madi, Huiyuan Zhang, Max Klapholz, Elena Christian, Karen O. Dixon, Geoffrey Fell, Katherine Tooley, Davide Mangani, Junrong Xia, Meromit Singer, Donna Neuberg, Orit Rozenblatt-Rosen, Aviv Regev, Vijay K. Kuchroo, Ana C. Anderson

**Author notes:** Co-first author (Nandini Acharya, Asaf Madi). Correspondence (V.K.K.), (A.R.), (A.C.A).

## Abstract

Identifying signals in the tumor microenvironment (TME) that promote CD8^+^ T cell dysfunction can inform improved therapeutic approaches for cancer. Here, we identify that *Nr3c1*, the gene encoding the glucocorticoid receptor (GR), is highly expressed in dysfunctional CD8^+^ tumor-infiltrating lymphocytes (TILs). The GR transactivates expression of multiple checkpoint receptors and loss of GR in CD8^+^ T cells limits dysfunctional phenotype in CD8^+^ TILs resulting in improved tumor growth control. We show that glucocorticoids can be produced in the TME and that they co-operate with the immunosuppressive cytokine IL-27 to promote the dysfunction gene program in CD8^+^ T cells. The presence of the glucocorticoid + IL27 signature in CD8^+^ TILs correlates with failure to respond to checkpoint blockade in melanoma patients, highlighting the relevance of this immunoregulatory glucocorticoid-cytokine circuit in tumor tissue.

## Introduction

Although the immune system has the capacity to fight cancer, signals present within the tumor microenvironment (TME) actively suppress anti-tumor immune responses. In particular, CD8^+^ T cells, key mediators of anti-tumor immunity, develop a dysfunctional or “exhausted” state in the highly immunosuppressive TME (Wherry and Kurachi, 2015), but the key mechanisms and factors in the TME that promote T cell dysfunction have not been identified. Dysfunctional CD8^+^ T cells exhibit defective cytolytic activity, loss of pro-inflammatory cytokines, and induction of the immunosuppressive cytokine IL-10 (Jin et al., 2010). Accordingly, dysfunctional CD8^+^ T cells are not only poor mediators of tumor clearance but also contribute to the suppressive microenvironment within tumor tissue. Therefore, understanding the signals, both T cell intrinsic and extrinsic, that contribute to the development of T cell dysfunction is of key importance in devising effective therapies to improve anti-tumor CD8^+^ T cell responses.

We previously defined a gene signature for dysfunctional CD8^+^ tumor-infiltrating lymphocytes (TILs) based on the differential gene expression of CD8^+^ TIL populations that exhibit distinct effector capacities(Sakuishi et al., 2010; Singer et al., 2016). Specifically, the expression of the checkpoint receptors Tim-3 and PD-1 distinguishes CD8^+^ TILs subsets with different degrees of function: Tim-3^+^PD-1^+^ CD8^+^ TILs are severely dysfunctional, Tim-3^−^PD-1^+^ CD8^+^ TILs are partially dysfunctional with intermediate effector function, and Tim-3^−^PD-1^−^ CD8^+^ TILs exhibit strong effector function (Fourcade et al., 2010; Sakuishi et al., 2010), with each of these populations exhibiting distinct transcriptional profiles(Singer et al., 2016). From the transcriptome data of these subsets of CD8^+^ TILs, we identified *Nr3c1*, the gene encoding the glucocorticoid receptor (GR), as being most highly expressed in severely dysfunctional Tim-3^+^PD-1^+^ CD8^+^ TILs. Glucocorticoids (GC), steroid hormones derived from the metabolic breakdown of cholesterol, bind to the GR, which resides in the cytosol in its inactive state and translocates to the nucleus upon ligand binding. In the nucleus, the GR can regulate gene expression either directly by binding to the promoter of a given target gene or indirectly by affecting the binding of other transcription factors (TFs) to the promoter regions of their respective targets (Oakley and Cidlowski, 2013). Both natural and synthetic glucocorticoids suppress a number of inflammatory indices and have been used clinically since the 1950s for treating excessive inflammation in patients with asthma and autoimmune diseases. Currently, glucocorticoids are routinely used to manage excessive inflammation in cancer patients treated with checkpoint blockade (Kumar et al., 2017).

Despite their widespread use, surprisingly little is known regarding the molecular circuitry by which glucocorticoids suppress immune responses (Cain and Cidlowski, 2017; Munck et al., 1984). The prevailing dogma attributes the anti-inflammatory effects of glucocorticoids to transrepression, whereby the GR inhibits the function of TFs that have key roles in driving pro-inflammatory responses. The GR binds to and directly interferes with AP-1 (Jonat et al., 1990; Yang-Yen et al., 1990). The GR can also interfere with NF-κB either directly or indirectly by modulating I*κ*B*α* (Auphan et al., 1995; Rhen and Cidlowski, 2005; Scheinman et al., 1995; Smoak and Cidlowski, 2004). However, glucocorticoids have also been associated with enhanced expression of anti-inflammatory cytokines, such as IL10 (Barrat et al., 2002), raising the possibility that in addition to actively repressing pro-inflammatory gene expression, they may also promote suppression via transactivation of immune-suppressive genes.

We show that activation of GR signaling in CD8^+^ T cells promotes T cell dysfunction in the TME and that the GR transactivates the expression of the checkpoint receptors Tim-3, PD-1, and Lag-3, and of the anti-inflammatory cytokine IL-10. Accordingly, loss of GR in CD8^+^ T cells results in tumor growth inhibition. We further demonstrate that monocyte/macrophage lineage cells are a chief source of glucocorticoid in the TME, that reduced steroidogenesis is correlated with better survival in cancer patients, and that glucocorticoid signaling co-operates with IL-27 signaling to form an immunoregulatory circuit that promotes T cell dysfunction in the TME. High expression of the glucocorticoid + IL-27 signature correlates with failure to respond to checkpoint blockade in melanoma patients, highlighting the relevance of this immunoregulatory circuit in human disease.

## Results

### Glucocorticoid signaling is active in dysfunctional CD8^+^ TILs

Analysis of transcriptional profiles (Singer et al., 2016), showed that *Nr3c1,* the gene encoding the glucocorticoid receptor (GR) is highly expressed in the PD-1^+^ CD8^+^ and Tim-3^+^PD-1^+^ CD8^+^TIL subsets that exhibit intermediate and severe dysfunctional phenotype, respectively (**Figure S1**). Examination of GR protein showed that it is most highly expressed in severely dysfunctional Tim3^+^PD1^+^ CD8^+^ TILs in two different tumor models, MC38-Ova colon carcinoma and B16F10 melanoma (**Figure 1a**), indicating that dysfunctional CD8^+^ T cells may have increased sensitivity to glucocorticoid signaling. Consistent with the expression pattern on murine CD8^+^ TILs subsets, the GR was also most highly expressed in Tim-3^+^PD-1^+^ CD8^+^ TILs from human colon carcinoma tumors (**Figure 1b**). Thus, we hypothesized that glucocorticoid signaling may be associated with the development of CD8^+^ T cell dysfunction in both murine and human tumors.

**Figure 1:**
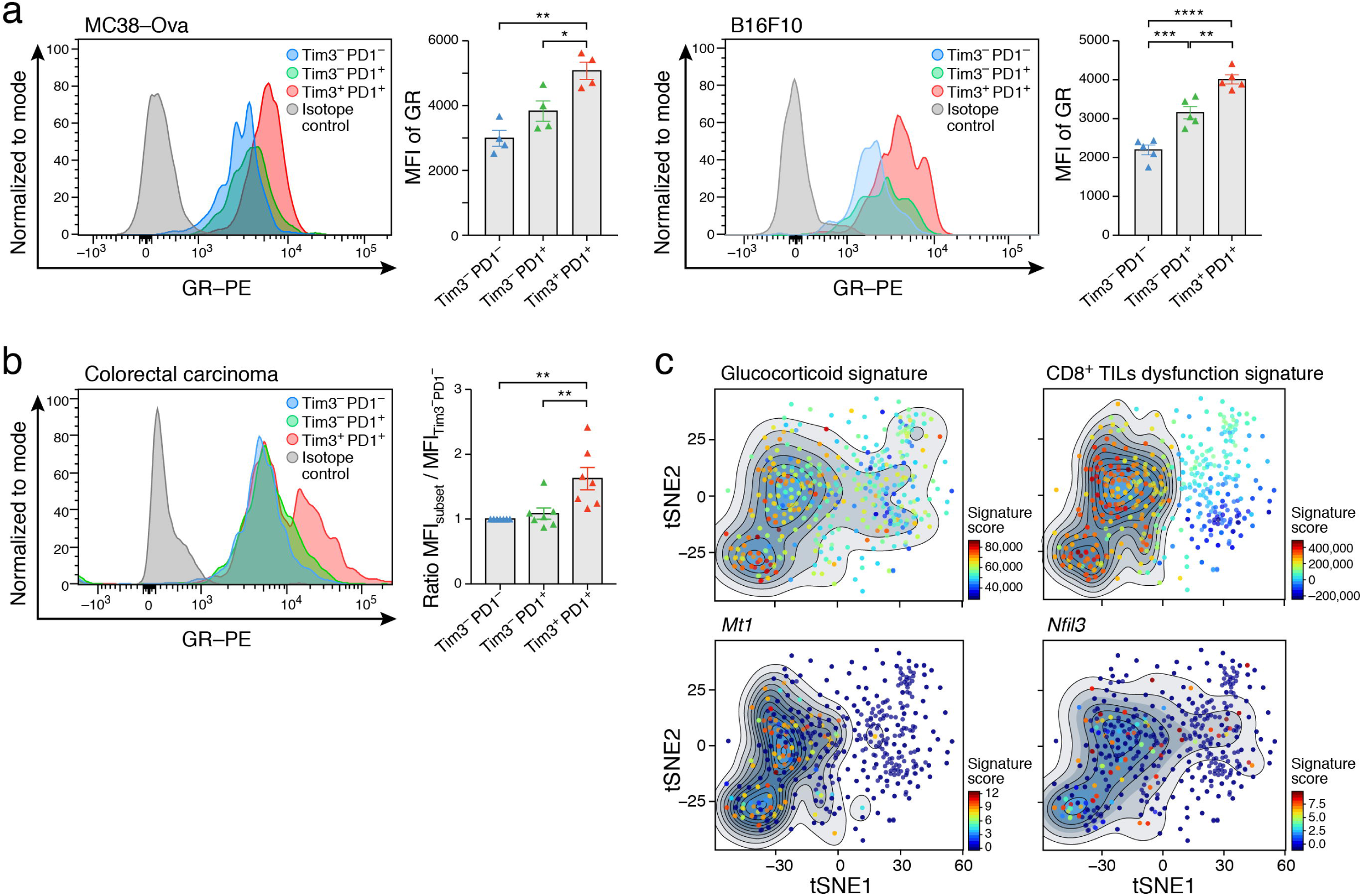
Glucocorticoid signaling is active in dysfunctional CD8^+^ TILs. TILs were harvested from (a) mice bearing MC38-Ova colon carcinoma (n=4) or B16F10 melanoma (n=5) or (b) from human colorectal carcinomas (n=7) for the examination of glucocorticoid receptor (GR) expression by intracellular staining. Representative histograms show GR expression in the indicated CD8^+^ TILs populations. Summary plots show the mean fluorescence intensity (MFI) of GR expression in the indicated populations. For human colorectal carcinoma TILs, data are normalized to the expression level in Tim-3-PD-1-CD8^+^ TILs. *p < 0.05, **p < 0.01, ***p < 0.001, ****p < 0.0001. Ordinary one-way ANOVA (Tukey’s multiple comparisons test). Mean ± SEM is shown. c) tSNE plot showing projection of a glucocorticoid signature (top left), a CD8^+^ T cell dysfunction signature (top right), and *Mt1* (bottom left) and *Nfil3* (bottom right) gene expression onto the single-cell RNA profiles of CD8^+^ TILs(Singer et al., 2016). The contour marks cells showing highest expression and the color scale indicates low (dark blue) to high (red) expressing cells.

To further test the possible association of glucocorticoid signaling with CD8^+^ T cell dysfunction, we scored the expression of a previously established glucocorticoid signature (Phuc Le et al., 2005) (**Methods**) in the single-cell RNA-Seq (scRNA-Seq) profiles of CD8^+^ TILs (Singer et al., 2017) from B16F10 melanoma (**Figure 1c**). Cells expressing the glucocorticoid signature and known GR target genes, such as *Mt1* (Karin and Herschman, 1979) and *Nfil3* (Carey et al., 2013), also scored highly for expression of the T cell dysfunction or “exhaustion” signature (**Methods**), indicating that glucocorticoid signaling was active in CD8^+^ TILs that exhibit dysfunctional phenotype.

### Glucocorticoid signaling promotes features of dysfunctional phenotype in CD8^+^ T cells

Accordingly, we hypothesized that glucocorticoid signaling might promote T cell dysfunction in murine and human cells. We tested the effect of repeated activation of CD8^+^ T cells in the presence of exogenous glucocorticoid (dexamethasone; Dex) *in vitro*. In line with observations in acutely activated cells (Barrat et al., 2002; Brattsand and Linden, 1996; Rhen and Cidlowski, 2005), we found that repeated activation in the presence of glucocorticoid profoundly suppressed the production of the pro-inflammatory cytokines IFN-γ, IL-2, and TNF-*α*, and up-regulated the immune-suppressive cytokine IL-10 (**Figure 2a**), a phenotype consistent with dysfunctional T cells. Indeed, we found that glucocorticoid treatment dramatically upregulated checkpoint receptors associated with dysfunctional phenotype including PD-1, Tim-3, and Lag-3, but not Tigit (**Figure 2b**). Notably, the glucocorticoid-mediated induction of checkpoint receptor expression was conserved in human CD8^+^ T cells (**Figure 2c**). Additionally, we observed that glucocorticoid increased the frequency of Tim-3^+^PD-1^+^ CD8^+^ T cells in both murine and human samples (**Figure S2a,b**). The observed effects of glucocorticoid were not due to reduced T cell survival or altered proliferation (**Figure S2c,d**). We further tested the effect of a natural glucocorticoid, corticosterone, on the expression of checkpoint receptors and found that it recapitulated the effects of Dex (**Figure S2e**).

**Figure 2:**
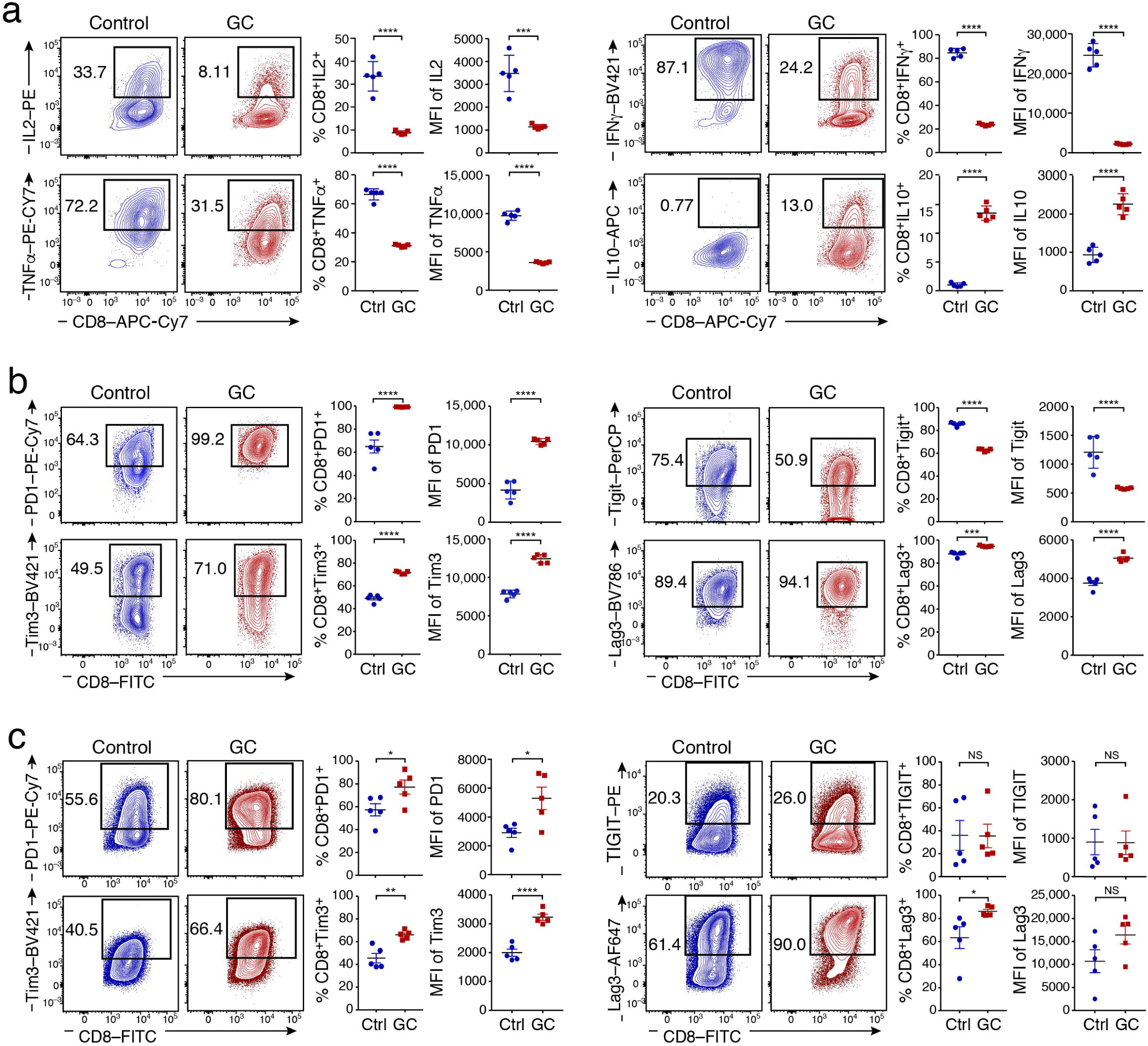
Glucocorticoid signaling promotes checkpoint receptor expression and dampens CD8^+^ T cell effector functions. Naïve CD8^+^ T cells from wild type mice (n= 5) were activated in the presence or absence of Dex (GC) and harvested on Day 9. **a)** Cells were stimulated with PMA/ionomycin for 4 hrs followed by intracellular staining for IL-2, TNF-*α*, IFN-*γ* and IL10. **b)** Expression of Tim-3, PD-1, Lag3, and Tigit was examined by flow cytometry. Data shown are representative of 3 independent experiments. **c)** Human CD8^+^ T cells were activated in the presence or absence of Dex (GC) and expression of Tim-3, PD-1, Lag-3, and TIGIT was examined by flow cytometry on Day 9. Data shown are representative of 2 independent experiments. *p<0.05, **p <0.01, ***p < 0.001, ****p<0.0001, two-tailed Student’s t test. Mean ± SEM is shown.

The observed effects of glucocorticoid on CD8^+^ T cells depended on *Nr3c1*. We examined expression of *Nr3c2*, which encodes the mineralocorticoid receptor (MR) that shares high structural homology with GR and can bind glucocorticoids with high affinity (Arriza et al., 1987). We found that *Nr3c2* is not expressed by wild type CD4^+^ and CD8^+^ T cells or in CD8^+^ T cells from mice that lack *Nr3c1* expression specifically in mature CD8^+^ T cells (E8i-Cre x Nr3c1^fl/fl^) (**Figure S3a**). Further, comparison of the RNA profiles of wild type and E8i-Cre x Nr3c1^fl/fl^ CD8^+^ T cells stimulated with or without glucocorticoid showed distinct glucocorticoid-induced changes in wild type but not E8i-Cre x Nr3c1^fl/fl^ CD8^+^ T cells, indicating that glucocorticoid-induced transcription in CD8^+^ T cells was *Nr3c1* dependent (**Figure S3b**). Thus, repeated stimulation in the presence of active glucocorticoid signaling dramatically influenced the effector differentiation of CD8^+^ T cells, resulting in cells that exhibited features shared with dysfunctional T cells, including up-regulation of multiple checkpoint receptors, dampened pro-inflammatory cytokine production, and increased IL-10 production.

### Glucocorticoid signaling in CD8+ TILs promotes tumor progression

We next tested whether glucocorticoid signaling impacts the functional state of CD8^+^ TILs *in vivo.* For this, we employed E8i-Cre x *Nr3c1*^fl/fl^ mice. Examination of T cell development and the steady state peripheral immune compartment of these mice showed no gross differences compared to wild type (*Nr3c1*^fl/fl^) littermate controls (**Figure S4a-d**). Further, we confirmed that the deletion of *N3rc1* was specific to CD8^+^ T cells (**Figure S4e**). We implanted either ovalbumin expressing MC38 colon carcinoma (MC38-Ova) or B16F10 melanoma cells into wild type and E8i-Cre x *Nr3c1*^fl/fl^ mice and found that E8i-Cre x *Nr3c1*^fl/fl^ mice exhibited improved tumor growth control in both models (**Fig. 3a** and **Figure S5a**), indicating that the effect of glucocorticoid signaling in CD8^+^ T cells was conserved across tumor types.

**Figure 3:**
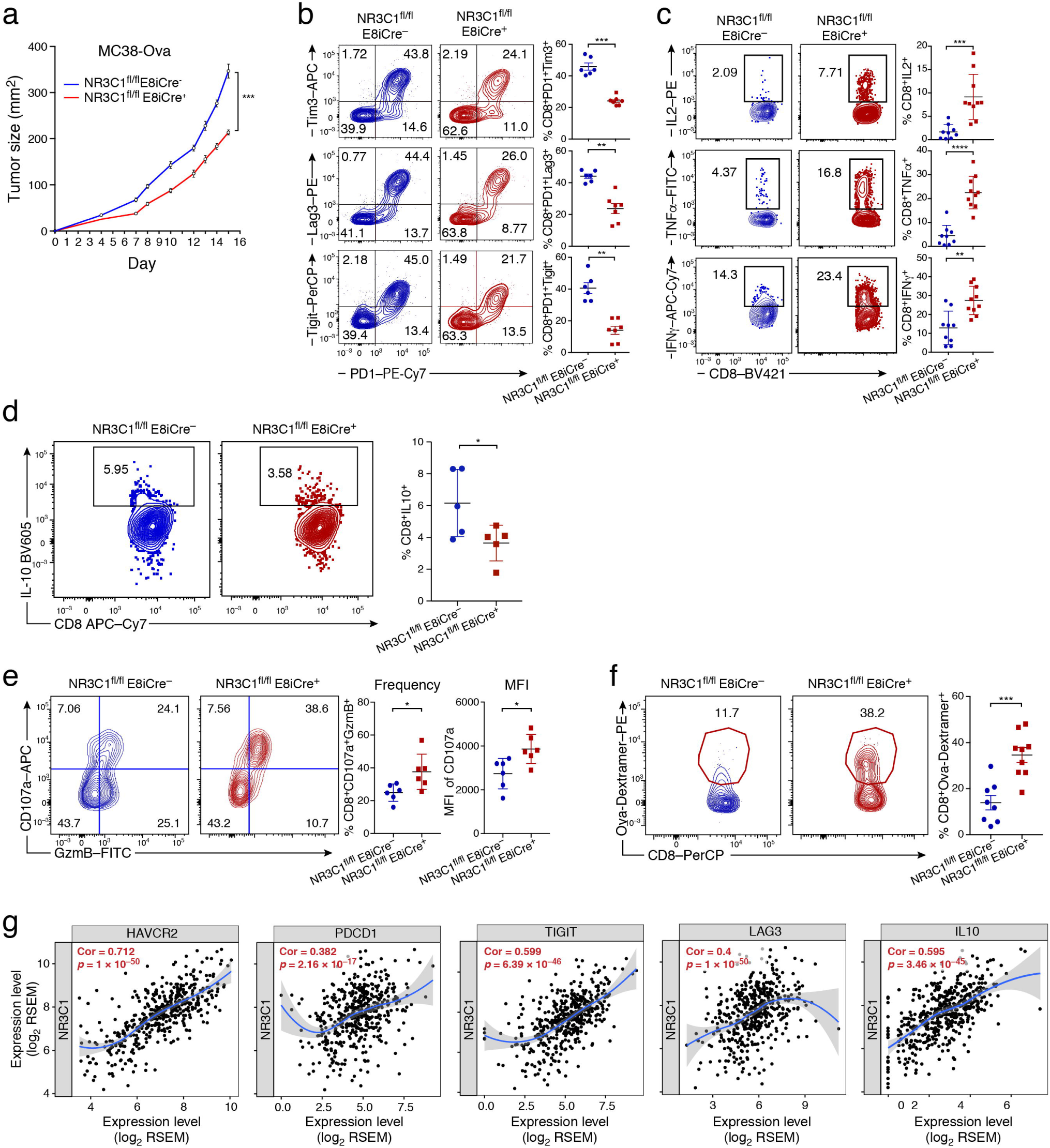
Glucocorticoid signaling dampens CD8^+^ TILs effector functions. **a)** MC38-Ova was implanted into wild type (E8i-Cre^−^*Nr3c1*^fl/fl^) and E8i-Cre^+^*Nr3c1*^fl/fl^ mice (n=8-9). Mean tumor growth is shown, ***p< 0.001, linear mixed model. Data are representative of 3 independent experiments. **b)** TILs were harvested on Day 13 post tumor implantation and the expression of checkpoint receptors was analyzed by flow cytometry. Representative flow cytometry data are shown. Scatter plots show summary data (n=6-7). Data are representative of 3 independent experiments. **c,d,e)** TILs were harvested and activated with 5 μg/ml OVA_257-264_ (SIINFEKL). **c)** Representative flow cytometry data and summary scatter plots showing the frequency of IL-2, TNF-α, and IFN-γ-producing CD8^+^ T cells (n=9-10). Data are pooled from 2 independent experiments. **d)** Representative flow cytometry data and summary scatter plots showing the frequency of IL10 producing CD8^+^ T cells (n=5). **e)** Representative flow and summary scatter plots show the frequency of CD107a and Granzyme B expression (n=6). Data are representative of 2 independent experiments. **f)** TILs were stained with H-2Kb/ OVA257-264 dextramer, scatter plot shows the frequency of tumor antigen-specific CD8^+^ T cells (n=8-9). Data are representative of 2 independent experiments. *p < 0.05, **P < 0.01, ***p < 0.001, ****p<0.0001, two-tailed Student’s t-test. Mean ± SEM are shown. **g)** Correlation of *NR3C1* mRNA with checkpoint receptor and IL10 mRNA in colon adenocarcinoma patients using TIMER.

Glucocorticoid signaling impacted the functional state of CD8^+^ TILs *in vivo*, based on the difference in several key parameters between E8i-Cre x *Nr3c1*^fl/fl^ and wild type MC38-Ova-bearing mice. First, not only was there a dramatic reduction in the frequency of CD8^+^ TILs co-expressing PD-1, Tim-3, Lag-3, and Tigit in CD8^+^ TILs from E8i-Cre x *Nr3c1*^fl/fl^ mice (**Figure 3b**), but also the expression level of each of these checkpoint receptors was significantly reduced (**Figure S5b**). Of note, Tigit expression was down-regulated in CD8^+^ TILs from E8i-Cre x Nr3c1^fl/fl^ mice (**Figure 3b and Figure S5b**), in contrast to our *in vitro* observations where Tigit expression was not induced by GR stimulation (**Figure 2b**). Second, CD8^+^ TILs from E8i-Cre x *Nr3c1*^fl/fl^ mice had enhanced responses to tumor-antigen (OVA_257-264_) as well as polyclonal stimulation, producing more IL-2, TNF-*α*, and IFN-*γ* (**Figure 3c** and **Figure S5c**). Indeed, the CD8^+^ TILs from E8i-Cre x *Nr3c1*^fl/fl^ mice were more polyfunctional in terms of pro-inflammatory cytokine production (**Figure S5d**). Furthermore, the few Tim-3^+^PD-1^+^CD8^+^ TILs in E8i-Cre x *Nr3c1*^fl/fl^ mice exhibited increased pro-inflammatory cytokine production in response to OVA_257-264_ stimulation (**Figure S5e**), in contrast to their typical severe dysfunctional phenotype observed in wild type mice. Third, the CD8^+^ TILs from E8i-Cre x *Nr3c1*^fl/fl^ mice produced lower amounts of the immune-suppressive cytokine IL10 (**Figure 3d** and **Figure S5f**). Fourth, CD8^+^ TILs from E8i-Cre x *Nr3c1*^fl/fl^ mice had higher cytotoxic capacity, as shown by the increased frequency of Granzyme B^+^CD107a^+^ cells upon OVA_257-264_ stimulation (**Figure 3e**). Finally, E8i-Cre x *Nr3c1*^fl/fl^ mice harbored more H-2K^b^/OVA_257-264_ dextramer^+^ CD8^+^ TILs (**Figure 3f**). We analyzed the proliferation status and the absolute number of the CD8^+^ TILs in wild-type and E8i-Cre x *Nr3c1*^fl/fl^ mice and observed no significant differences (**Figure S5g,h**). Notably, checkpoint receptor expression on CD4^+^ TILs in E8i-Cre x *Nr3c1*^fl/fl^ mice was not significantly different from that of wild type CD4^+^ TILs, indicating that the regulation of checkpoint receptors in CD8^+^ TILs was cell-intrinsic and not due to a secondary effect of loss of GR in CD8^+^ TILs (**Figure S5i,j**). Finally, in human colon adenocarcinoma from TCGA (http://cancergenome.nih.gov/), we found (using TIMER (Li et al., 2016)) that *NR3C1* mRNA levels positively correlated with *HAVCR2* (Tim-3), *PDCD1* (PD-1), *LAG3*, *TIGIT* and *IL10* mRNA levels (**Figure 3g**). Collectively, these data supported that glucocorticoid signaling is active in the TME of both murine and human tumors and functions to promote checkpoint receptor expression and dampen the effector function of CD8^+^ TILs.

### The glucocorticoid receptor transactivates the expression of checkpoint receptors and IL-10

We next tested if the GR directly regulates the expression of checkpoint receptor genes and IL-10. First, we analyzed GR-binding peaks in the loci of *Havcr2* (Tim-3), *Pdcd1* (PD-1), *Lag3*, *Tigit,* and *IL10* in publicly available ChIP-seq data(Oh et al., 2017) from bone marrow-derived macrophages (BMDMs) (**Figure S6**). We found GR-binding peaks in the loci of *Havcr2, Lag3*, and *IL10* but not *Pdcd1* or *Tigit*, reflecting the lack of PD-1 and Tigit expression in BMDMs. We further found that some of the GR binding peaks in the *Havcr2, Lag3*, and *IL10* loci overlapped regions of accessible chromatin (based on ATAC-seq) in IL-27-stimulated T cells (Karwacz et al., 2017), which are known to express high levels of these molecules (Chihara et al., 2018) (**Figure S6**). We therefore tested the effect of GR binding to the *cis*-regulatory elements in these peaks in the *Havcr2*, *Pdcd1*, *Lag3*, and *Tigit* loci using luciferase reporter assays. For *IL10*, we utilized luciferase reporters of a previously established enhancer element of *Il10* – HSS^+^2.98 as well as the proximal promoter (−1.5kb)(Karwacz et al., 2017). We transfected the different luciferase reporter constructs along with a *Nr3c1* expressing vector or empty vector into 293T cells and treated the cells with glucocorticoid to assay the transactivation capability of the GR. In line with our observations in glucocorticoid treated CD8^+^ T cells (**Figure 2**), the GR potently transactivated Tim-3, PD-1, Lag-3, and IL-10 expression (**Figure 4**). Tigit was also induced but to a much lower degree (**Figure 4d**).

**Figure 4:**
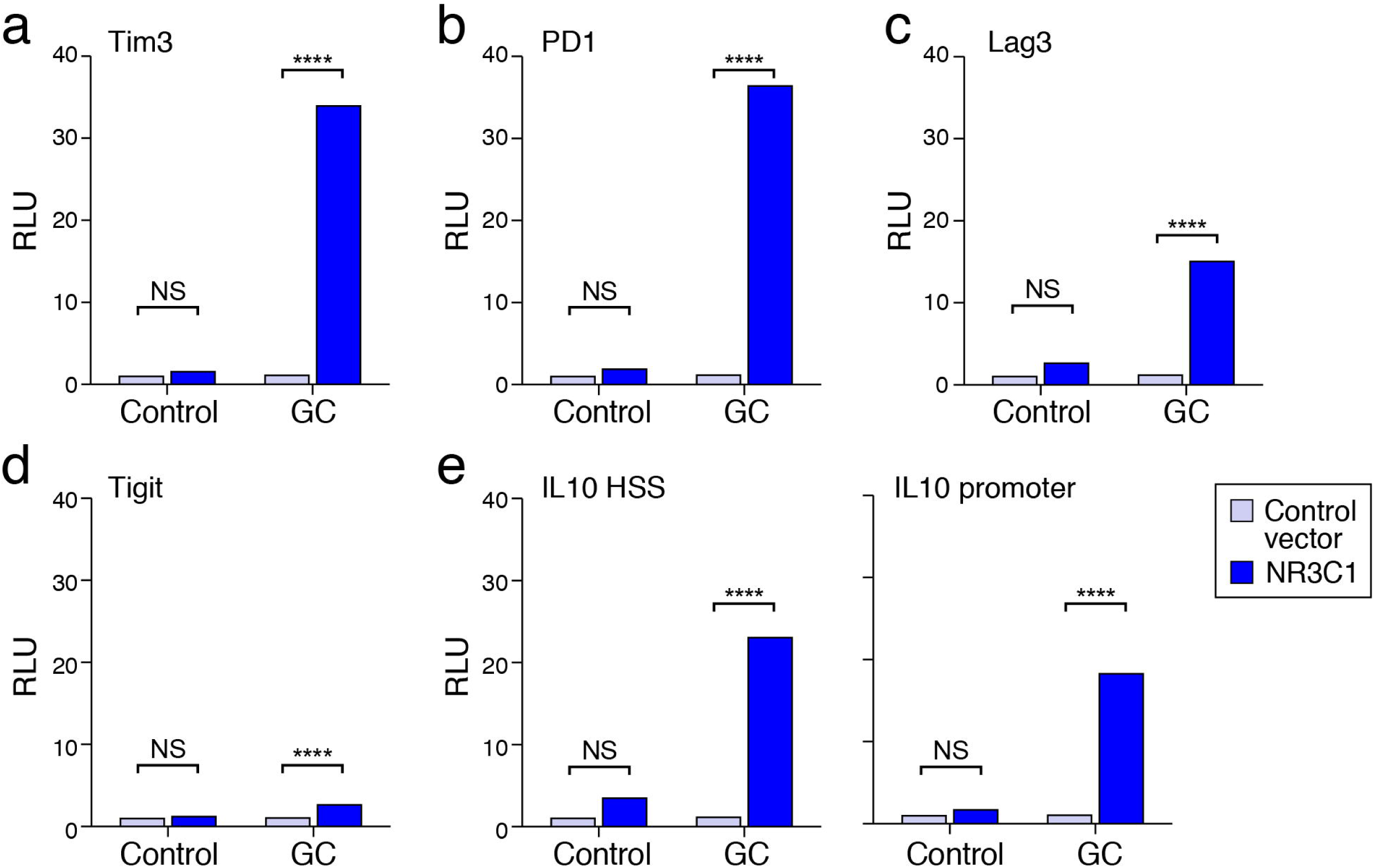
Transactivation of checkpoint receptors and IL-10 by GR. Luciferase activity in 293T cells transfected with pGL4.23 or pGL4.10 luciferase reporters for the loci of a) *Havcr2* (Tim3), b) *Pdcd1* (PD1), c) Lag3 d) Tigit, and e) IL10 together with either empty vector (control) or constructs encoding *Nr3c1*. Cells were treated with GC after 24h. Firefly luciferase activity was measured 48 h after transfection and is presented relative to constitutive Renilla luciferase activity. NS is not significant, ****p<0.0001, two-way ANOVA (Tukey’s multiple comparisons test). Data are mean ± S.E.M. The data are representative of 2 independent experiments.

### Glucocorticoid and IL-27 signaling co-operate to promote CD8^+^ T cell dysfunction

The transactivation of multiple checkpoint receptors and IL-10 by GR and the effect of glucocorticoid signaling on the effector responses of CD8^+^ TILs were reminiscent of our recent observation that IL-27 regulates a gene module that includes checkpoint receptors (Tim-3, Lag3, Tigit) and IL-10 and suppresses the responses of CD8^+^ TILs(Chihara et al., 2018; Zhu et al., 2015). Interestingly, glucocorticoids have been shown to work in concert with TFs such as the STAT family (Petta et al., 2016) and STAT1 and STAT3 are downstream of IL-27. These observations prompted us to address the relationship of the glucocorticoid and IL-27 pathways. We collected and analyzed the RNA-Seq profiles from cells treated with glucocorticoid, IL-27, or both. Unsupervised principle component analysis (PCA) showed that glucocorticoid and IL-27 each induced a distinct transcriptional profile with glucocorticoid + IL-27 treatment inducing the largest transcriptional change relative to control (Figure 5a and b and **Figure S7a**). Examination of differentially expressed genes across all three conditions relative to control showed some common as well as some distinct genes (**Figure S7b**). 6,812 genes were differentially expressed (DE) between glucocorticoid + IL-27 compared to control out of which 3,417 (50%) showed non-additive regulation (**Figure 5c** and **Table S1**). Among the genes that were highly up-regulated by glucocorticoid + IL-27 compared to glucocorticoid or IL-27 alone were *Prdm1* and *Nfil3*, which encode TFs with known roles in promoting or maintaining T cell dysfunction (Chihara et al., 2018; Rutishauser et al., 2009; Shin et al., 2009; Zhu et al., 2015). In contrast, *Tcf7*, which encodes TCF-1, a TF important for maintaining stem-like T cells critical for the response to checkpoint blockade and which shows antagonism with Tim-3 expression (Im et al., 2016; Kurtulus et al., 2019; Miller et al., 2019; Siddiqui et al., 2019), was down-regulated by glucocorticoid + IL-27. Using qPCR, we confirmed that *Prdm1* and *Nfil3* expression was most highly up-regulated by treatment with glucocorticoid + IL-27 compared to glucocorticoid or IL-27 alone (**Figure S7c**). *Tcf7* was dramatically reduced by both glucocorticoid or IL27 alone and treatment with glucocorticoid + IL-27 showed a trend of further reduction (**Figure S7c**). These observations indicated that glucocorticoid + IL-27 signaling may co-operate to promote gene programs associated with T cell dysfunction in CD8^+^ T cells. Accordingly, we tested all of the differentially expressed genes induced by glucocorticoid + IL-27 for overlap with the T cell dysfunction signature. 1,022 out of 6,812 DE genes overlapped with the dysfunction signature (**Table S2**). The genes down-regulated by glucocorticoid + IL-27 significantly overlapped with genes expressed in CD8^+^TIM-3^−^PD-1^−^ TILs (p=2.1×10^−10^, Mean-rank Gene Set Test) and the genes up-regulated by glucocorticoid + IL-27 showed significant overlap with the genes expressed in severely dysfunctional CD8^+^Tim-3^+^PD-1^+^ TILs (p=4.3×10^−5^, Mean-rank Gene Set Test) (**Figure 5d** and **Figure S7d**).

**Figure 5:**
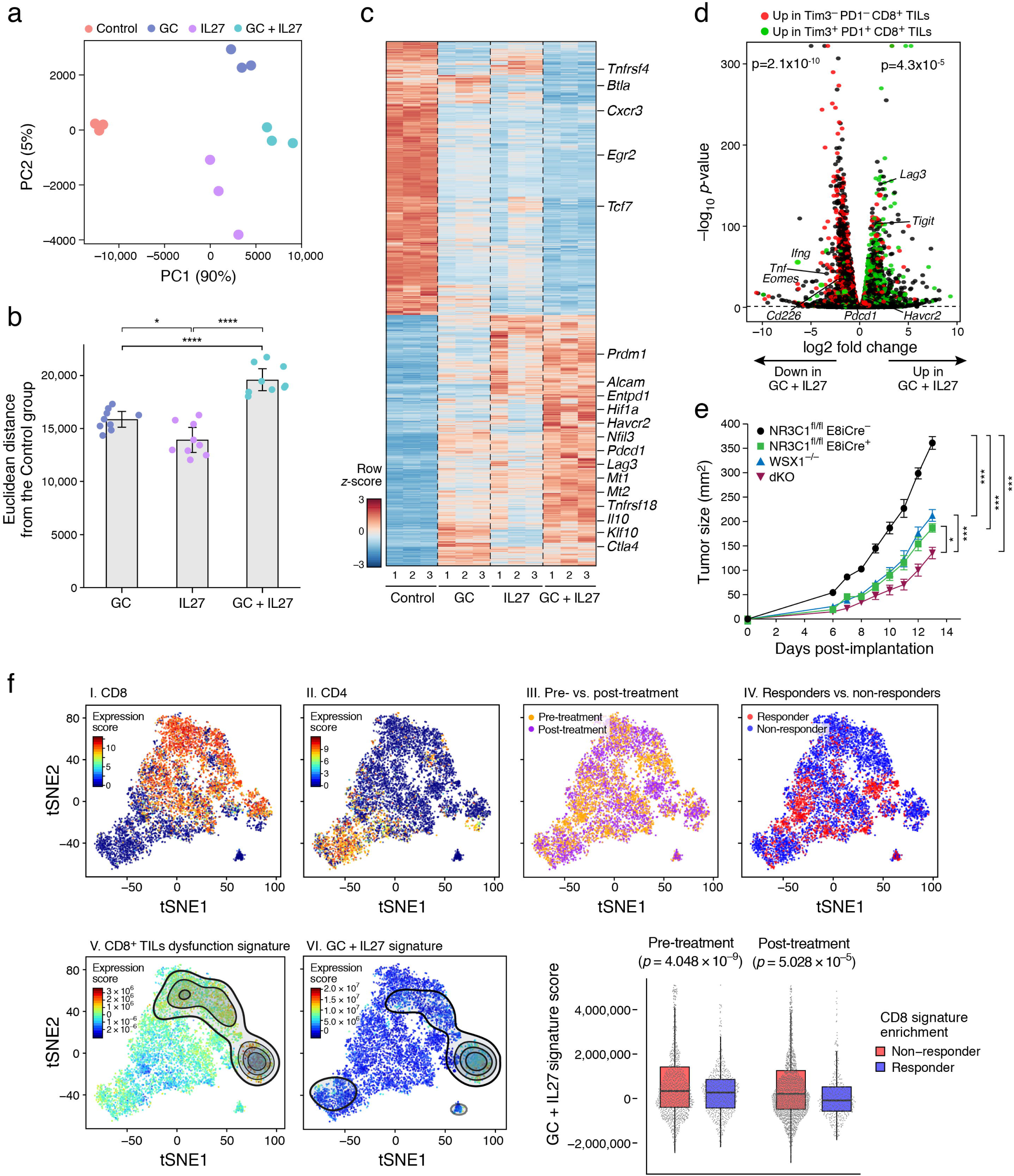
The glucocorticoid and IL-27 pathways co-operate to promote dysfunction in CD8^+^ T cells. **a-c)** Naïve CD8^+^ T cells were cultured *in vitro* with anti CD3/28 in the presence of Dex (GC), IL-27, or GC+IL-27. Cells were harvested on Day 9 and gene expression analyzed by RNA sequencing. **a)** Principle component analysis (PCA) of Ctrl, GC, IL-27, and GC+IL-27 treated CD8^+^ T, the percentage of explained variance for each principal component is indicated. **b)** Bar graph shows the mean delta Euclidean distance between the GC, IL-27, or GC+IL-27 treated groups to the control group, adjusted p-values were calculated using one-way ANOVA (p-value=9.89e-09), followed by Tukey HSD, *p < 0.05, ****p < 0.001. **c)** Heatmap of DE genes between Ctrl and GC + IL-27 treatment. Tick marks indicate selected genes associated with T cell dysfunction. **d)** Volcano plot showing overlap of genes down-regulated by IL-27 + GC with genes expressed in Tim-3^−^PD-1^−^ CD8^+^ TILs (p= 2.1×10^−10^, Mean-rank Gene Set Test), and genes up-regulated by IL-27+GC with Tim-3^+^PD-1^+^ CD8^+^ TILs (p=4.3×10^−5^, Mean-rank Gene Set Test). **e)** CD8 T cells from either WT (E8i-Cre^−^*Nr3c1*^fl/fl^), E8i-Cre^+^*Nr3c1*^fl/fl^, WSX1^−/−^ or and E8i-Cre^+^*Nr3c1*^fl/fl^ WSX1^−/−^ (DKO) mice and CD4^+^ T cells from WT mice were transferred to Rag^−/−^ mice (n=5-6/group) that were implanted with MC38-Ova cells two days later. Mean tumor growth is shown. *p<0.05, ***p< 0.001, linear mixed model. Data are representative of 2 independent experiments. **f)** tSNE plot of single-cell RNA profiles of TILs from melanoma patients. I) CD8 expression, II) CD4 expression, III) pre- (orange) versus post-(purple) treatment samples, IV) Responder (red) versus non-responder (blue), V) Projection of CD8^+^ TILs dysfunction signature, VI) Projection of the GC + IL-27 signature. Box plots show the GC + IL-27 signature score in responder versus non-responders in pre- and post-treatment samples, p = 4.048 x 10^−9^ and p = 5.028 x 10^−5^, respectively (Welch Two Sample t-test). The lower and upper hinges correspond to the first and third quartiles. The upper and lower whiskers extend from the hinge to the largest and smallest value no further than 1.5 times the distance between the first and third quartiles, respectively. Data beyond the end of the whiskers are outlying points and are not plotted individually.

To determine the functional consequences of the glucocorticoid + IL-27 signaling pathways on T cell dysfunction *in vivo*, we crossed E8i-Cre^+^*Nr3c1*^fl/fl^ mice with WSX1^−/−^ (*IL27ra*^−/−^) mice to generate mice that can be used as a source of double knock-out (DKO) CD8^+^ T cells, lacking both the glucocorticoid and IL-27 signaling pathways. We isolated CD8^+^ T cells from wild type, E8i-Cre^+^Nr3c1^fl/fl^, WSX-1^−/−^, or DKO mice and transferred them along with wild type CD4^+^ T cells into Rag-1^−/−^ mice followed by implant of MC38-Ova colon carcinoma cells. In line with our previous findings (**Figure 3a** and **Figure S5a**) (Zhu et al., 2015), absence of either glucocorticoid or IL-27 signaling alone individually conferred tumor growth control; however, absence of both of pathways together led to significantly greater tumor growth inhibition (**Figure 5e**).

To examine the relevance of glucocorticoid + IL-27 signaling in human disease, we scored the glucocorticoid + IL-27 signature in the single-cell data of TILs from melanoma patients pre- and post-treatment with checkpoint blockade (Sade-Feldman et al., 2018). We found that the glucocorticoid + IL-27 signature scored highly in CD8^+^ TILs that also scored highly for the T cell dysfunction signature (**Figure 5f, panels V** and **VI**). Most importantly, we found that high expression of the glucocorticoid + IL-27 signature correlated with non-responsiveness to checkpoint inhibitor in CD8^+^ TILs in both pre- (p= 4.048 x 10^−9^) and post- (p=5.028x 10^−05^) treatment samples (**Figure 5f**). Altogether, our data indicated that glucocorticoid and IL-27 signaling combined in the TME to promote CD8^+^T cell dysfunction and dampen anti-tumor immunity.

### Myeloid cells are the primary sources of glucocorticoid and IL-27 in the TME

We next asked whether local sources in the TME provided endogenous glucocorticoid and IL-27 signals. Although steroids are mainly synthesized in the adrenal cortex, it has been suggested that tumor cells are capable of extra-adrenal steroidogenesis (Sidler et al., 2011). To test whether glucocorticoid was indeed produced in tumor tissue, we measured corticosterone production in the spleen and tumor tissue from MC38-Ova tumor-bearing mice. We found that corticosterone was produced at high levels in the tumor tissue relative to the spleen from tumor-bearing mice while the levels of corticosterone in the spleen of tumor-bearing and non-tumor bearing mice were similar (**Figure 6a**). Further, we cultured tumor explants in the presence or absence of Metyrapone, an inhibitor of glucocorticoid synthesis, and observed reduced levels of corticosterone in the presence of the inhibitor (**Figure S8a**). Together these data indicated local glucocorticoid production in the TME. To test the impact of this on tumor growth, we administered Metyrapone to MC38-Ova tumor-bearing mice by intra-tumoral injection and observed dramatic tumor growth inhibition in the treated mice (**Figure 6b**).

**Figure 6:**
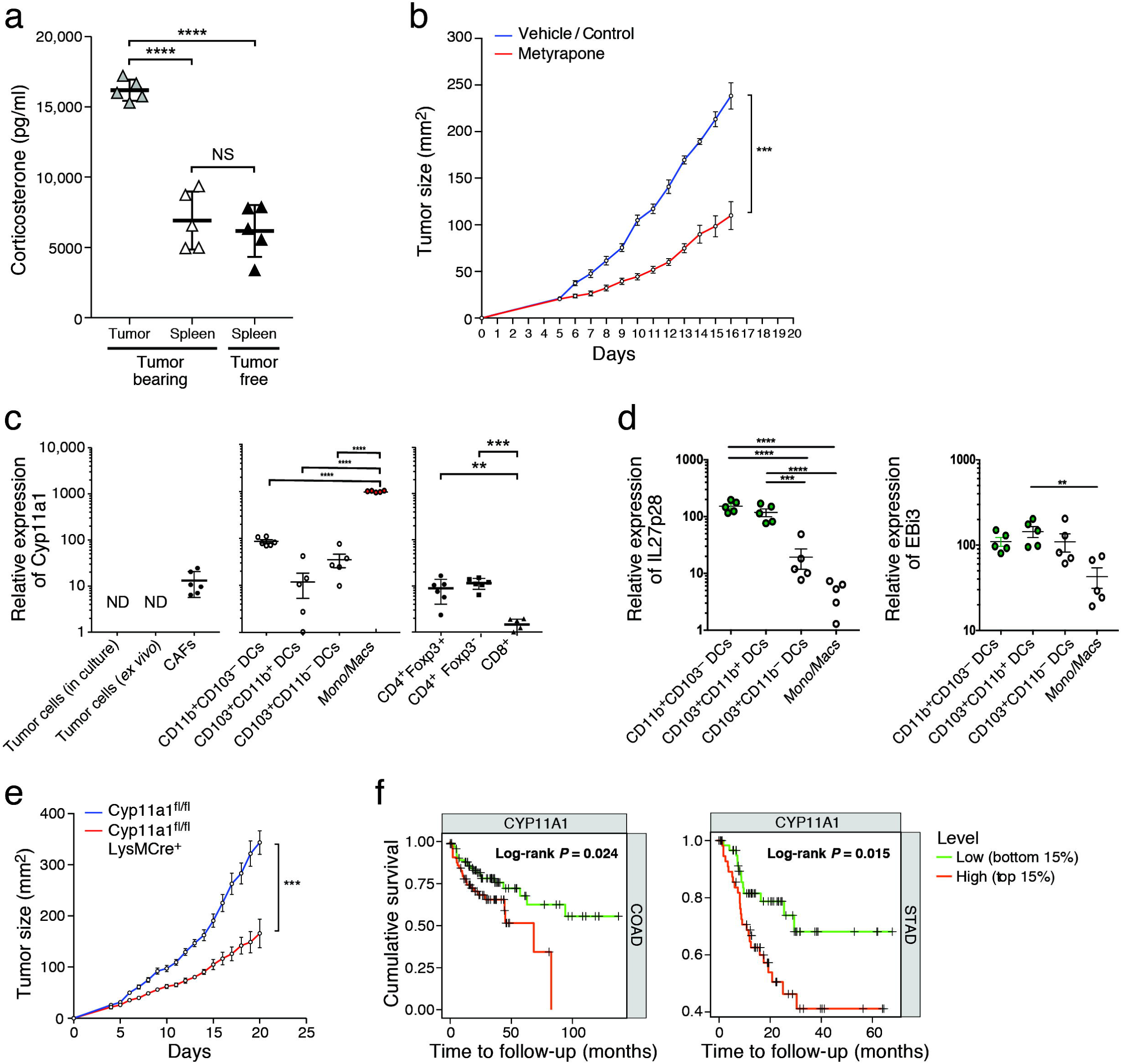
Glucocorticoid and IL-27 are synthesized by different cells in the TME. **a)** Corticosterone levels in the indicated tissues were quantified by ELISA. (n=5). b) MC38-Ova was implanted in wild-type mice. Metyrapone or vehicle control was administered intra-tumorally on Days 5,6,7 and 9 post tumor implantation (n=5/group). Mean tumor growth is shown ***p< 0.001, linear mixed model. Data are representative of 2 independent experiments. Quantitative RT-PCR analysis of (c) Cyp11a1 and (d) IL-27 (p28 and Ebi3) mRNA expression in the indicated cells. Data are pooled from 2 independent experiments. **p < 0.01, ***p < 0.001, ****p<0.0001. Ordinary-one way ANOVA (Tukey’s multiple comparisons test). Data are mean ± S.E.M. e) MC38-Ova was implanted in Cyp11a1^fl/fl^ and Cyp11a1^fl/fl^LysMCre^+^ mice and tumor progression was studied (n=5). Mean tumor growth is shown, ***p< 0.001, linear mixed model. Data are representative of 2 independent experiments. f) Correlation of Cyp11a1 mRNA expression level with survival in patients with colon adenocarcinoma (COAD) and stomach adenocarcinoma (STAD) using TIMER.

Steroids are produced by the enzymatic breakdown of cholesterol, where cytochrome P450 cholesterol side-chain cleavage enzyme (Cyp11a1 or P450scc) catalyzes the first and the rate-limiting step that catalyzes the breakdown of cholesterol to pregnenolone, the precursor of all steroid hormones (Payne and Hales, 2004). Thus, to identify which cell types were responsible for steroid production in the TME, we first examined the expression of Cyp11a1. We examined the MC38-Ova tumor cell line and found that *in vitro* cultured MC38-Ova cells did not express Cyp11a1 (**Figure 6c**). To test if factors present in the TME could induce expression of Cyp11a1 in MC38-Ova tumor cells, we implanted MC38-Ova-GFP cells in mice and examined Cyp11a1 expression in tumor cells (CD45^−^GFP^+^). We did not detect Cyp11a1 expression in the isolated tumor cells (**Figure 6c**). Examination of other cells in the TME showed that cancer-associated fibroblasts (CAFs) (CD45^−^GFP^−^PDGFRa^+^), tumor-associated dendritic cells (TADCs), and T cells (mostly CD4^+^ T cells) expressed Cyp11a1 but at much lower levels compared to tumor-associated monocyte/macrophages (**Figure 6c**). We further examined the expression of the other enzymes involved in glucocorticoid biosynthesis (StAR,Cyp21a1,Cyp17a1,Cyp11b1,Hsd3b1) in monocytes/macrophages from MC38-Ova tumors and found that they expressed all enzymes to varying degrees with the exception of Hsd3b1, which was difficult to detect (**Figure S8b**). To confirm that tumor-associated monocyte/macrophages could indeed produce glucocorticoid, we measured their corticosterone production after culture in the presence or absence of Metyrapone. We found that the cells produced corticosterone and this was dramatically reduced by the addition of Metyrapone (**Figure S8c**). Together these data indicated that monocyte/macrophages, which are present from early time points during tumor progression and comprise more than 50% of CD45^+^ cells in MC38-Ova tumors, are a major source of glucocorticoid in the TME. Conversely, we found that TADCs were the main source of IL-27, as they expressed high levels of both p28 and EBi3 (**Figure 6d**). Therefore, different cell types within the TME produced glucocorticoid and IL-27.

To study the relevance of steroid production from monocyte/macrophages on tumor growth, we implanted tumors in LysMCre^+^ x Cyp11a1^fl/fl^ mice and observed significant tumor growth control (**Figure 6e**). Lastly, we examined the relevance of steroid abundance in the TME in human cancers. Using TIMER, we found that low Cyp11a1 mRNA levels were associated with a substantial survival benefit in patients with colon adenocarcinoma and stomach adenocarcinoma (**Figure 6f**). Collectively, our data demonstrate that endogenous glucocorticoid signaling co-operates with IL-27 signaling to form an immunoregulatory circuit that dampens effective anti-tumor immunity by promoting T cell dysfunction in the TME.

## Discussion

Glucocorticoids, steroid hormones, have been shown to suppress immune responses by interfering with AP-1- and NF-κB-mediated induction of pro-inflammatory cytokines (Auphan et al., 1995; Jonat et al., 1990; Rhen and Cidlowski, 2005; Scheinman et al., 1995; Smoak and Cidlowski, 2004; Yang-Yen et al., 1990). Here, we identify a novel molecular mechanism by which glucocorticoids suppress effector T cell responses through the transactivation of multiple checkpoint receptors (Tim-3, PD-1, and Lag3) together with IL-10. Glucocorticoids can be produced in the TME and combine with the immune suppressive cytokine IL-27 to form an immunoregulatory circuit that promotes gene programs associated with CD8^+^ T cell dysfunction. The relevance of glucocorticoid signaling pathways to human cancer is supported by our observation that increased steroidogenic capacity in the TME correlated with decreased survival and that high expression of the glucocorticoid + IL-27 signature in the CD8^+^ TILs of melanoma patients correlated with failure to respond to checkpoint blockade therapy (Sade-Feldman et al., 2018).

The observation that *Prdm*1 and *Nfil3,* TFs known to be involved in T cell dysfunction (Chihara et al., 2018; Rutishauser et al., 2009; Shin et al., 2009; Zhu et al., 2015), are most highly induced by the combination of glucocorticoid and IL-27 provides a potential mechanism for the co-operativity of these two pathways in the promotion of T cell dysfunction. Moreover, glucocorticoid + IL-27 strongly reduced the expression of *Tcf7*, which encodes TCF-1, a TF involved in maintenance of stem-like memory precursor CD8^+^ TILs that are required for the success of checkpoint blockade therapy (Im et al., 2016; Kurtulus et al., 2019; Miller et al., 2019; Siddiqui et al., 2019). Thus, glucocorticoid + IL-27 signaling may subvert anti-tumor immunity by two mechanisms, promoting T cell dysfunction and decreasing stemness and memory potential in CD8^+^ TILs.

The GR has also been shown to act via chromatin remodeling (Jubb et al., 2017). Therefore, it is possible that glucocorticoid signaling may drive epigenetic changes that together with IL-27 set the stage for CD8^+^ T cell dysfunction. The contributions of glucocorticoid and IL-27 signaling to the distinct epigenetic changes that have been described in dysfunctional CD8^+^ T cells (Pauken et al., 2016; Philip et al., 2017; Sen et al., 2016) remains to be determined.

Our data demonstrate that activation of glucocorticoid and IL-27 signaling promotes the dysfunction gene program in CD8^+^ TILs. However, a previous study implicated the GR in the maintenance of memory-precursor CD8^+^ T cells in the context of bacterial infection (Yu et al., 2017). Thus, the effect of glucocorticoid signaling favoring differentiation into memory or dysfunctional CD8^+^ T cells may be context dependent. It is also possible that the GR may partner with other signaling pathways and TFs depending on the tissue environment to achieve different effects. Further molecular investigation will help delineate the mechanisms operative in different contexts.

Using conditional knockout mice and *in vitro* studies we provide insight into the CD8^+^ T cell intrinsic effects of glucocorticoid signaling. However, glucocorticoid signaling may also act intrinsically in other immune and non-immune cell populations in the TME. Indeed, glucocorticoids have been implicated in increasing the frequency of T_reg_ cells in both humans and mice (Chen et al., 2006; Hu et al., 2012; Ling et al., 2007; Suarez et al., 2006). In myeloid cells, glucocorticoids have been implicated in modulating both antigen presentation and inflammatory cytokine production by DCs (Piemonti et al., 1999) with consequences for the efficacy of anticancer therapies (Yang et al., 2019). Further, in line with our data, a recent study suggested that endogenous glucocorticoids regulate the expression of PD-1 on NK cells in the context of viral infection (Quatrini et al., 2018). Whether these effects of glucocorticoid are due to cell intrinsic or extrinsic effects will require detailed molecular dissection using cell type specific knockouts.

The observations that extra-adrenal glucocorticoid biogenesis can occur in tumor tissue and that its ablation has important consequences for tumor growth underscore the need to identify the signals that promote steroidogenesis in cells within the TME. Our study highlights monocyte/macrophages lineage cells as one of the major sources of glucocorticoid in colon carcinomas. Whether other cell types are the predominant source of extra-adrenal steroid in other cancer types remains to be determined. Further, inhibitors of the enzymes of steroid biogenesis are used in the clinical management of cancers that are predominantly driven by steroid hormone signaling, such as breast and prostate cancer. Our data indicate the potential of such drugs in a broader spectrum of cancer types.

Our study focuses on the effects of endogenous glucocorticoid in the TME; however, exogenous glucocorticoid is often administered to cancer patients. In patients with glioblastoma, dexamethasone is given to prevent cerebral edema. Whether this negatively impacts on the ability of these patients to respond to checkpoint blockade immunotherapy is not known. Glucocorticoids are also used as first-line agents for managing immune-related adverse events (IRAEs) (Kumar et al., 2017) associated with checkpoint blockade immunotherapy. Although initial studies indicated that glucocorticoids can be used to treat IRAEs without negative impact on therapeutic outcome (Beck et al., 2006; Downey et al., 2007; Johnson et al., 2015; Weber et al., 2008), a recent study comparing patients who received checkpoint blockade and developed a severe IRAE followed by treatment with either low- or high-dose glucocorticoids showed that patients who received high-dose exogenous glucocorticoid had both reduced survival and time to treatment failure (Faje et al., 2018).

Further, another recent study showed that low-affinity memory T cells are inhibited by corticosterone and that overall survival (OS) was reduced in melanoma patients who received corticosteroids along with ICB (Tokunaga et al., 2019). These observations highlight the need to understand the effects of low-versus high-dose administration of exogenous glucocorticoids and how these relate to the effects of endogenous glucocorticoid. Notwithstanding these considerations, we have observed that patients who fail to respond to checkpoint blockade (Sade-Feldman et al., 2018) have higher expression of the glucocorticoid + IL-27 signature. Thus, having a better understanding of the mechanisms downstream of glucocorticoids and IL-27 could inform the clinical development of more precise therapies for modulating immune responses in the clinic, the relevance of which extends beyond cancer.

## Acknowledgments

We thank Deneen Kozoriz and Rajesh Kumar Krishnan for cell sorting. Sally Kent for advice in human T cell culture. Leslie Gaffney, Amit Bansal and Anna Hupalowska for figure preparation. This work was supported by grants from the National Institutes of Health (R01NS045937 to VKK, P01AI073748 to VKK and ACA, R01CA229400 to ACA), and by the Klarman Cell Observatory at the Broad Institute and HHMI. A.R. is an Investigator of the Howard Hughes Medical Institute.

## Authors Contributions

N.A. designed, performed and analyzed all biological experiments and wrote the manuscript.

A.M. performed computational analyses for RNA-seq and contributed to interpretation of the results.

H.Z. performed some biological experiments.

M.K. performed some biological experiments.

E.C. prepared RNA-seq libraries.

K.O.D helped with some biological experiments.

G.F. helped with statistical analyses.

K.T. provided reagents.

D.M. helped with some biological experiments.

J.X. helped with some biological experiments.

M.S. contributed to computational analysis.

D.N. helped with statistical analyses.

O.R.R. coordinated and supervised RNA Seq.

A.R. advised and provided insight into the analysis of RNA-seq analysis and contributed to interpretation of the results and writing of the manuscript.

V.K.K. supervised the study and contributed to interpretation of the results and writing of the manuscript.

A.C.A. designed, supervised the study and wrote the manuscript.

## Supplementary Figure Legends

**Figure S1:**
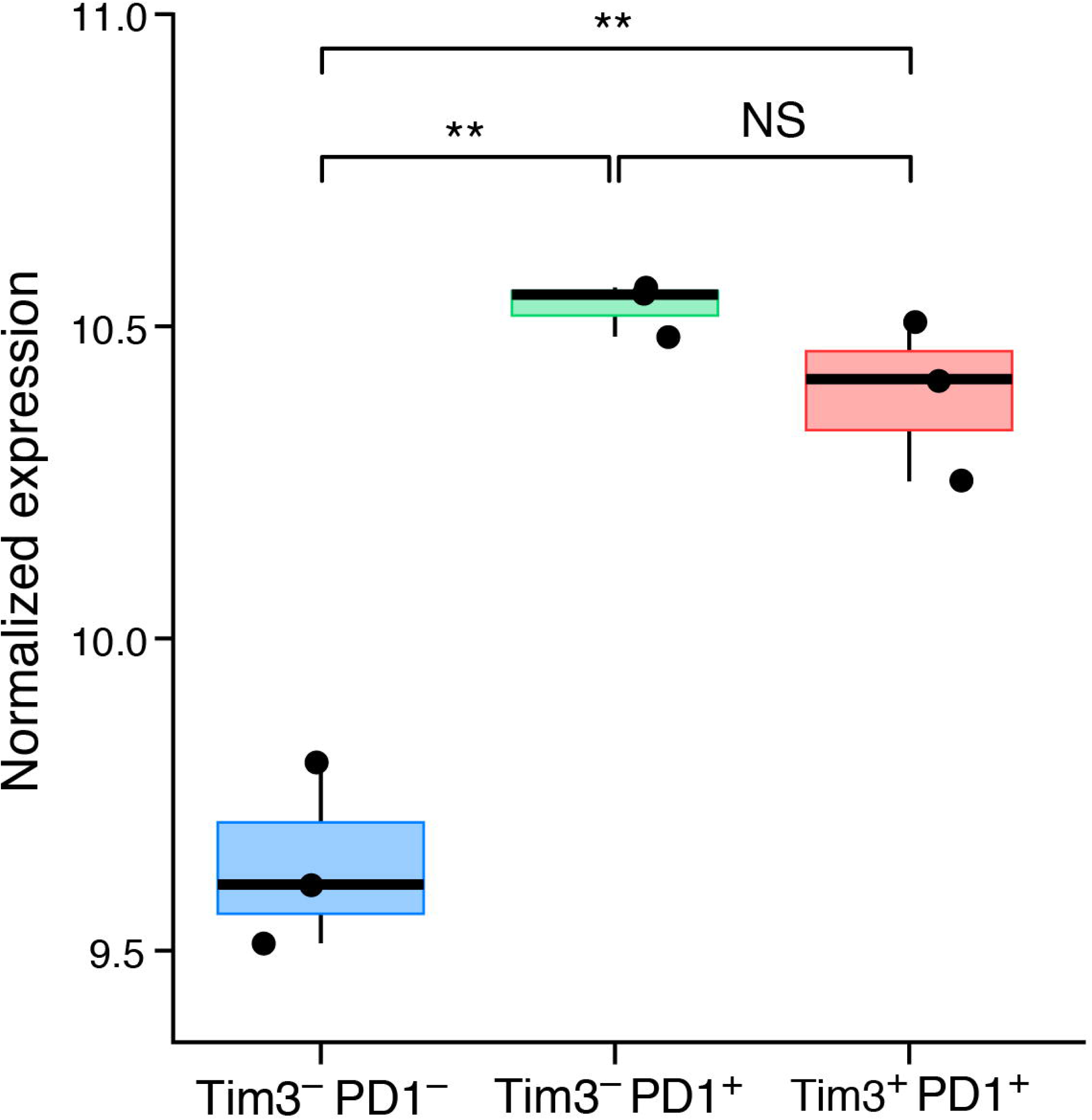
Glucocorticoid receptor expression in CD8^+^ TILs populations. Gene expression value of *Nr3c1* on Tim3^−^PD1^−^, Tim3^−^PD1^+^, and Tim3^+^PD1^+^ CD8^+^ TILs from CT26 colon carcinoma(Singer et al., 2016). NS is not significant, **p < 0.01. One-way ANOVA.

**Figure S2:**
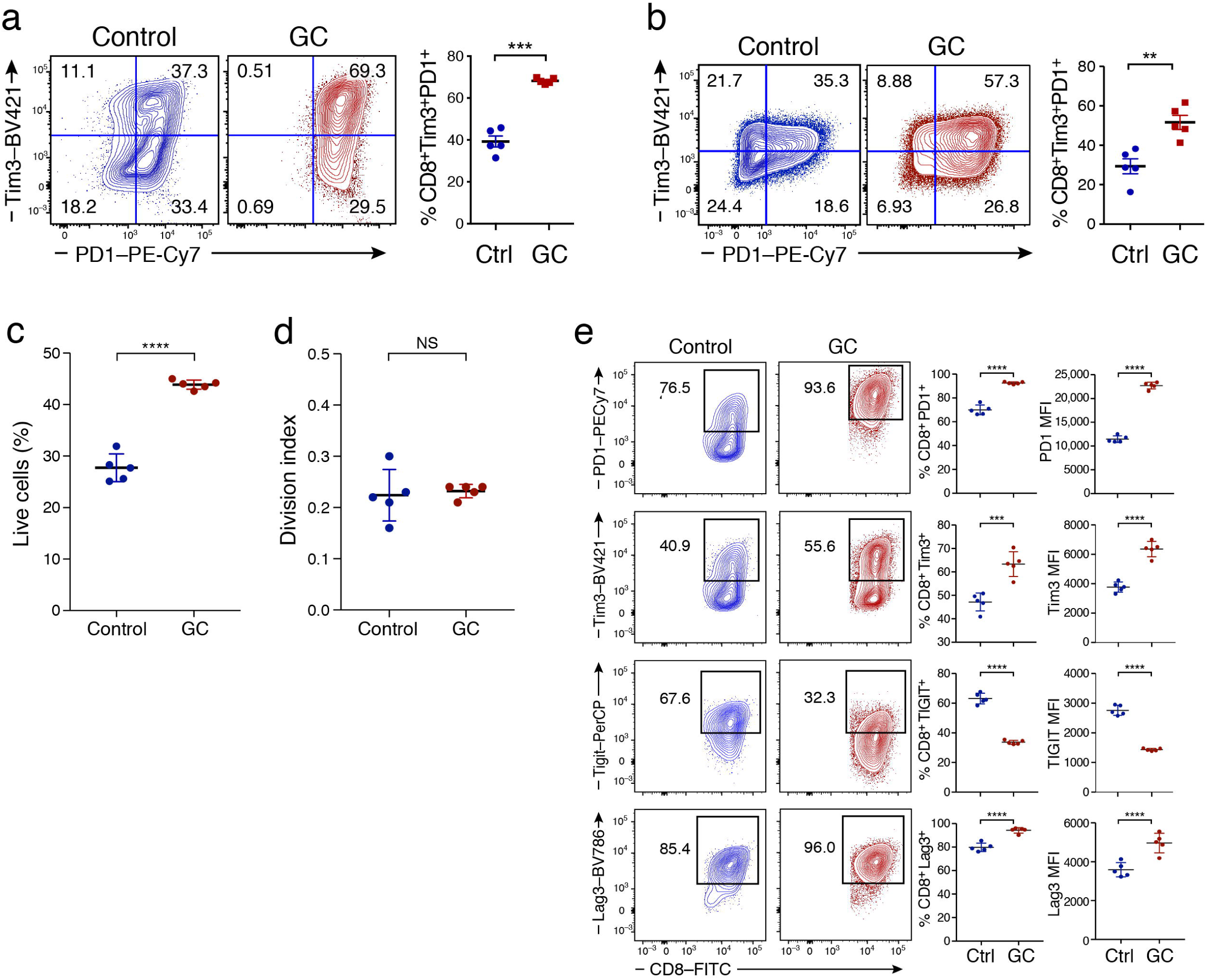
Effect of synthetic and natural Glucocorticoids on CD8^+^ T cells. Naïve CD8^+^ T cells from wild type mice (n= 5) **(a,c,d)** or from human samples (n= 5) **(b)** were activated in the presence or absence of Dex (GC) as in Fig. 2a and harvested on Day9. Representative flow cytometry data show Tim-3 and PD-1 expression**. c)** Frequency of viable CD8^+^ T cells. **d)** Division index of CD8^+^ T cells. **e)** Naïve CD8^+^ T cells from wild type mice (n= 5) were activated in the presence or absence of Corticosterone (GC) as in Fig. 2a and harvested on Day9. Representative flow cytometry data shows the frequency of Tim-3^+^, PD-1^+^, Tigit^+^ and Lag3^+^ cells (n=5). NS is Not Significant **p < 0.01, ***p < 0.001, ****p < 0.0001 two-tailed Student’s t-test. Mean ± SEM are shown.

**Figure S3:**
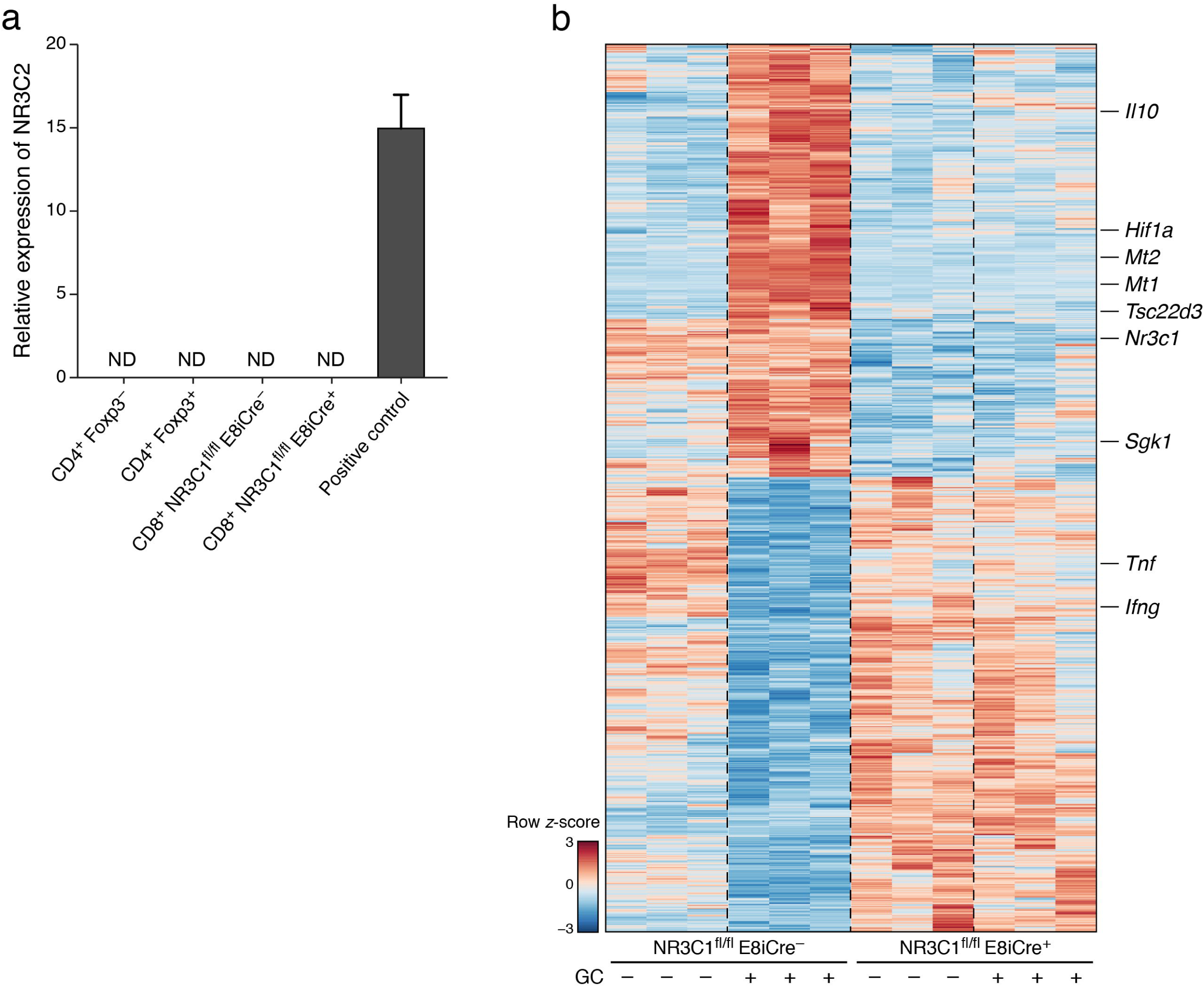
Glucocorticoid-mediated effects on CD8^+^ T cells requires *Nr3c1*. **a)** Expression of *Nr3c2* on T cells was quantified by qPCR. MC38-Ova cell-line was used as the positive control. Data are representative of 2 independent experiments. ND is Not Detected. **b)** Heatmap of differentially expressed genes in wild type (E8i-Cre^−^*Nr3c1*^fl/fl^) or E8i-Cre^+^*Nr3c1*^fl/fl^ CD8^+^ T cells activated in the presence or absence of Dex (GC) for 72hrs. Tick marks indicate selected known GC target genes.

**Figure S4:**
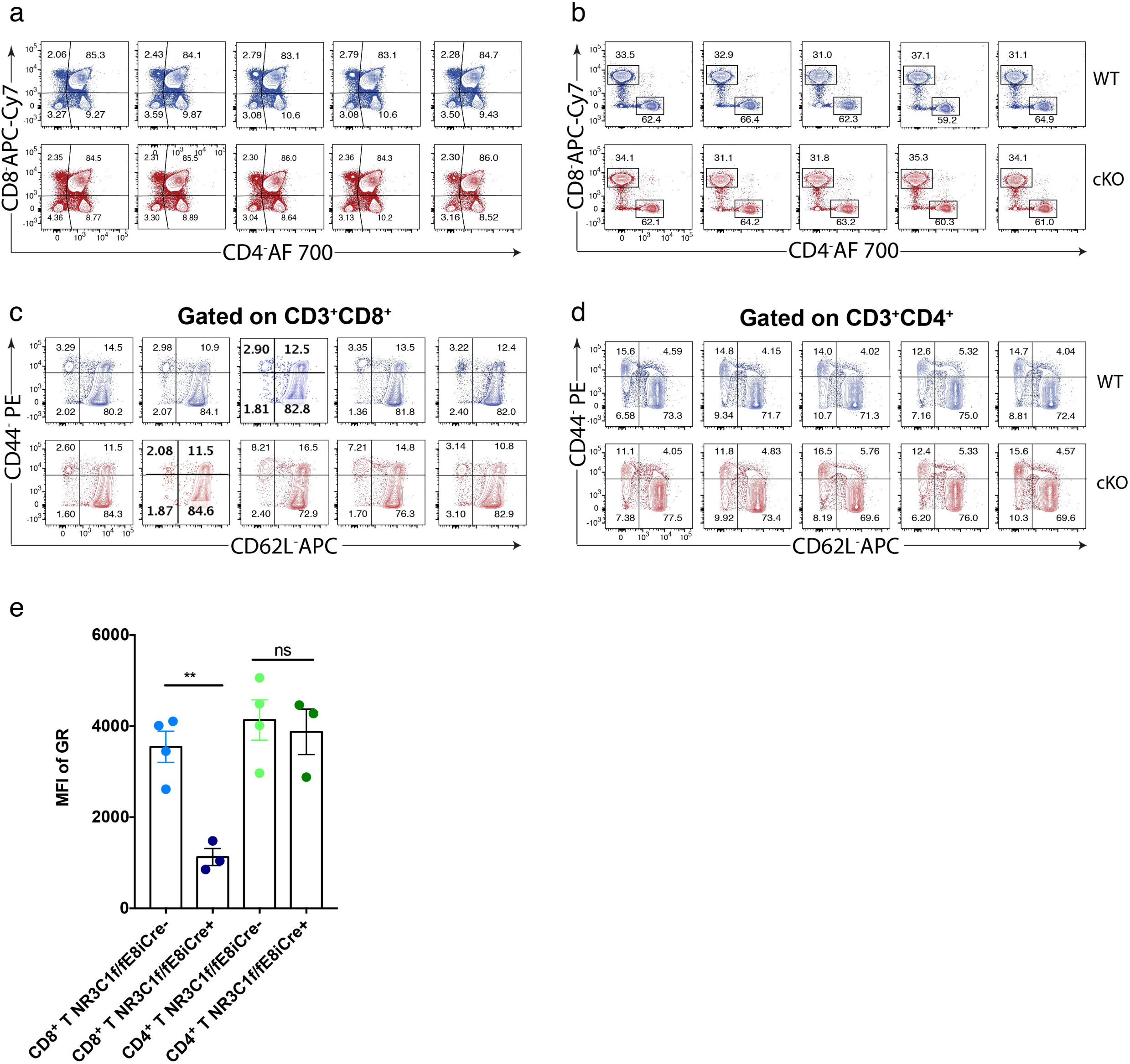
Deletion of Nr3c1 using E8iCre does not affect the development of T cells. Frequency of T cells in the thymus **(a)** and spleen (**b)** of Nr3c1^fl/fl^ (WT) and Nr3c1^fl/fl^E8iCre^+^ (Dodd et al.) mice. **c)** Frequency of naïve and activated CD8^+^ T cells in steady state. **d)** Frequency of naïve and activated CD4^+^ T cells in steady state. **e)** Expression of *Nr3c1* on T cells from Nr3c1^fl/fl^ and Nr3c1^fl/fl^E8iCre^+^ mice was quantified by qPCR. **P < 0.01, Ordinary one-way ANOVA (Tukey’s multiple comparisons test). Mean ± SEM are shown.

**Figure S5:**
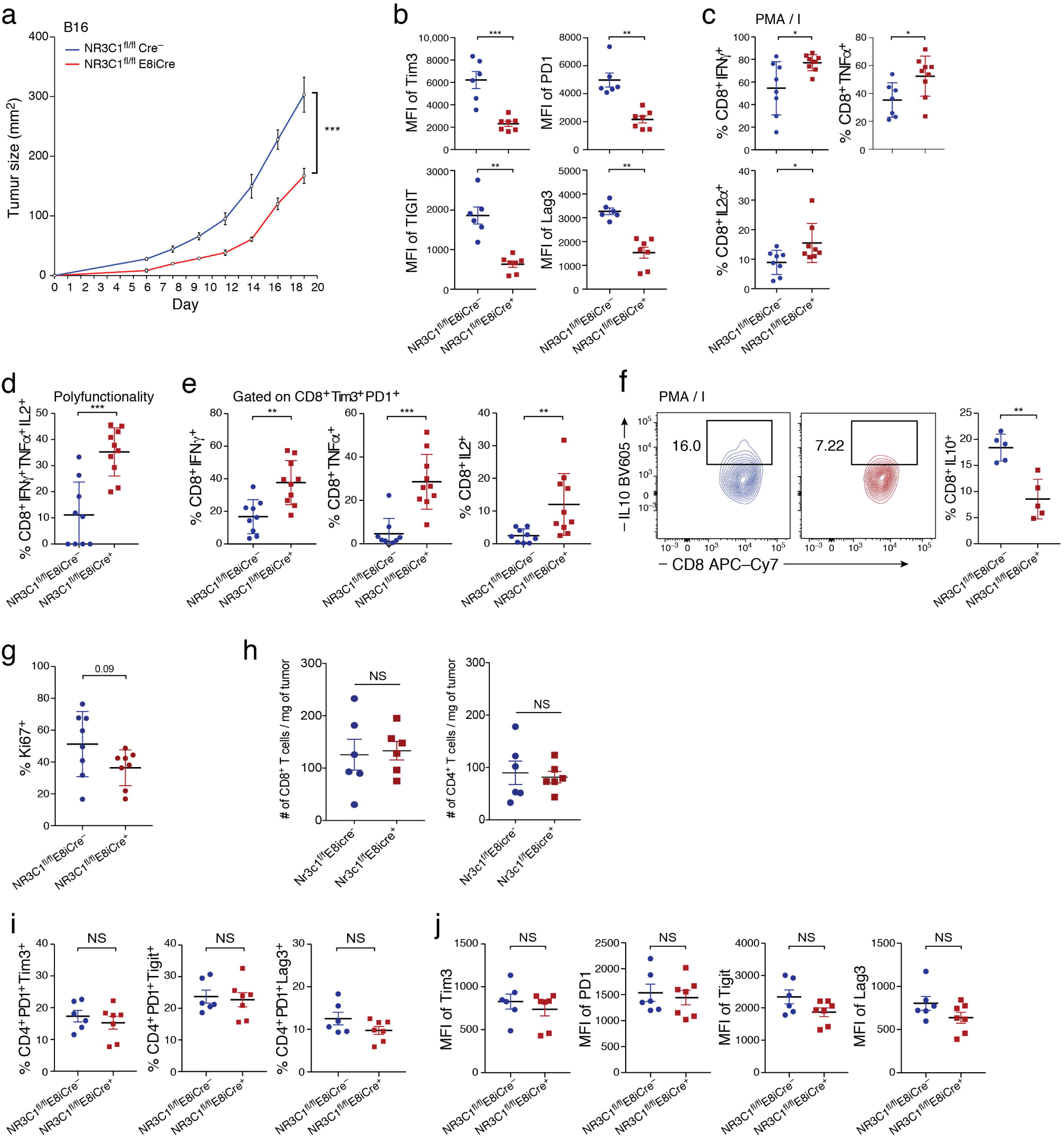
Glucocorticoid signaling dampens effector function of CD8^+^ T cells. **a)** B16F10 was implanted into wild type (E8i-Cre^−^*Nr3c1*^fl/fl^) and E8i-Cre^+^*Nr3c1*^fl/fl^ mice (n=5). Mean tumor growth is shown. ***p< 0.001, linear mixed model. Data are representative of 2 independent experiments. **b-f)** MC38-Ova was implanted into wild type (E8i-Cre^−^*Nr3c1*^fl/fl^) and E8i-Cre^+^*Nr3c1*^fl/fl^ mice. **b)** Expression level of Tim-3, PD-1, Lag-3 and Tigit as indicated by mean fluorescence intensity (MFI) in CD8^+^ T cells from wild type (E8i-Cre^−^*Nr3c1*^fl/fl^) and E8i-Cre^+^*Nr3c1*^fl/fl^ mice (n=6-7). Data are representative of 3 independent experiments. TILs were isolated and activated with PMA/ionomycin **(c,f)** and with OVA_257-264_ (SIINFEKL) **(d,e)** in the presence of Golgi Plug and Golgi Stop for 4 hr prior to extracellular and intracellular staining and analysis by flow cytometry. **c)** Summary plot showing cytokine production in CD8^+^ TILs from wild type (E8i-Cre^−^*Nr3c1*^fl/fl^) and E8i-Cre^+^*Nr3c1*^fl/fl^ mice (n=9-10) following polyclonal activation. Data are pooled from 2 independent experiments. **d)** Summary plot showing poly-functionality of CD8^+^ TILs from wild type (E8i-Cre^−^*Nr3c1*^fl/fl^) and E8i-Cre^+^*Nr3c1*^fl/fl^ mice (n=9-10). Data are pooled from 2 independent experiments. **e)** Summary plots representing cytokine production in Tim3^+^PD1^+^ CD8^+^ TILs from wild type (E8i-Cre^−^ *Nr3c1*^fl/fl^) and E8i-Cre^+^*Nr3c1*^fl/fl^ mice. Data are pooled from 2 independent experiments (n=9-10). **f)** Summary plots representing IL10 production in CD8^+^ TILs from wild type (E8i-Cre^−^ *Nr3c1*^fl/fl^) and E8i-Cre^+^*Nr3c1*^fl/fl^ mice (n=5). **g)** Summary plots representing Ki67^+^ CD8^+^ TILs from wild type (E8i-Cre^−^*Nr3c1*^fl/fl^) and E8i-Cre^+^*Nr3c1*^fl/fl^ mice (n=8). **h)** Summary plots representing absolute number of CD8^+^ TILs from wild type (E8i-Cre^−^*Nr3c1*^fl/fl^) and E8i-Cre^+^*Nr3c1*^fl/fl^ mice (n=6). **i)** Summary plots representing the frequency of CD4^+^ T cells expressing checkpoint receptors in wild type (E8i-Cre^−^*Nr3c1*^fl/fl^) and E8i-Cre^+^*Nr3c1*^fl/fl^ mice (n=6-7). **j)** Summary plots of the MFI of checkpoint receptors on CD4^+^ T cells in WT and E8i-Cre^+^*Nr3c1*^fl/fl^ mice (n=6-7). NS is not significant, *p< 0.05, **p< 0.01, ***p < 0.001, two-tailed Student’s t-test. Mean ± SEM are shown.

**Figure S6:**
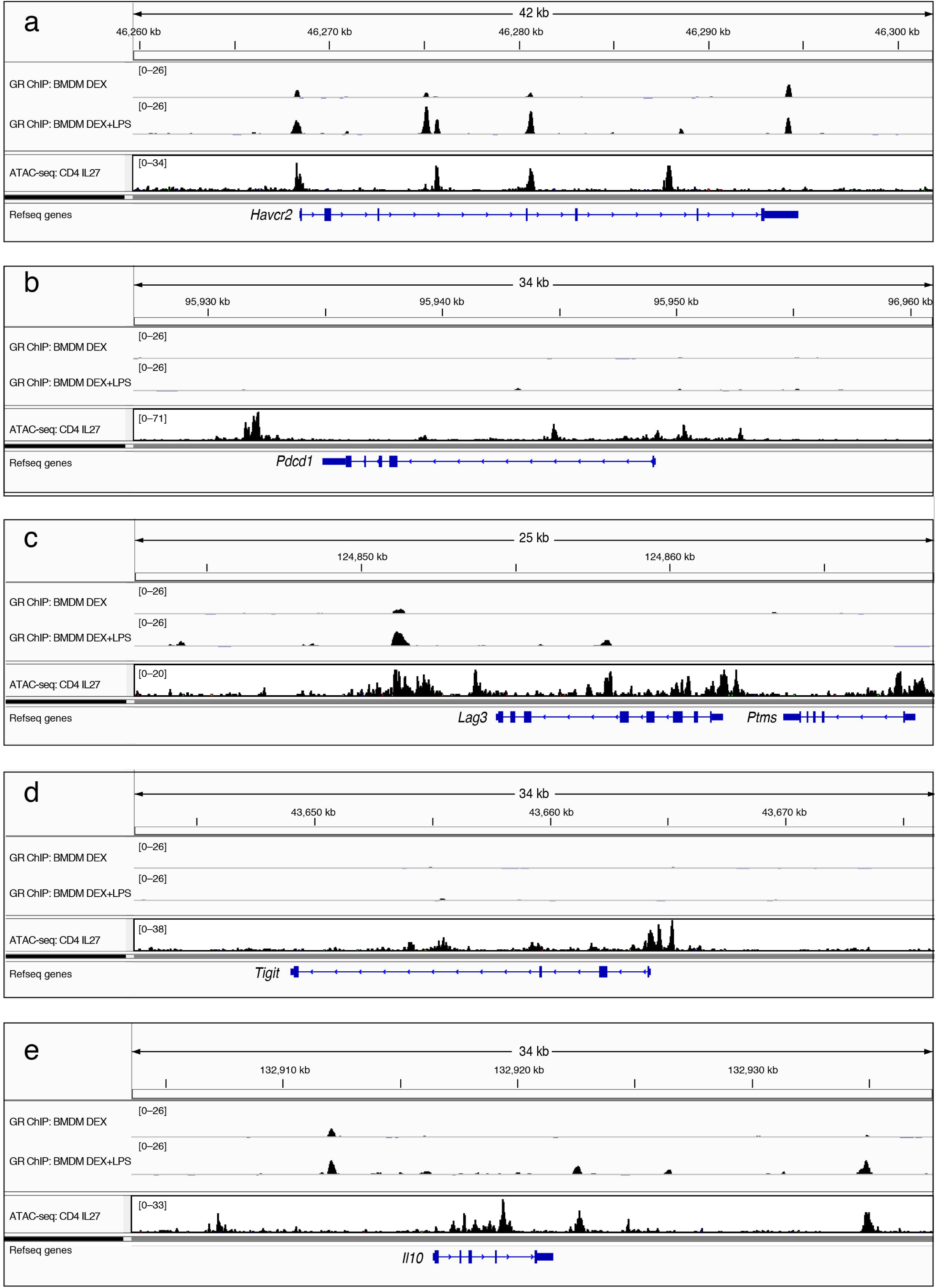
GR binding sites and open chromatin in the loci of checkpoint receptors and IL10. Overlay of ChIP-seq data of GR(Jubb et al., 2017) and ATAC-seq data of naive CD4^+^ cells induced with IL-27(Karwacz et al., 2017) in the loci of **(a)** *Havcr2* (Tim3) **(b)** *Pdcd1* (PD1) **(c)** *Lag3* **(d)** *Tigit* and **(e)** *Il10*.

**Figure S7:**
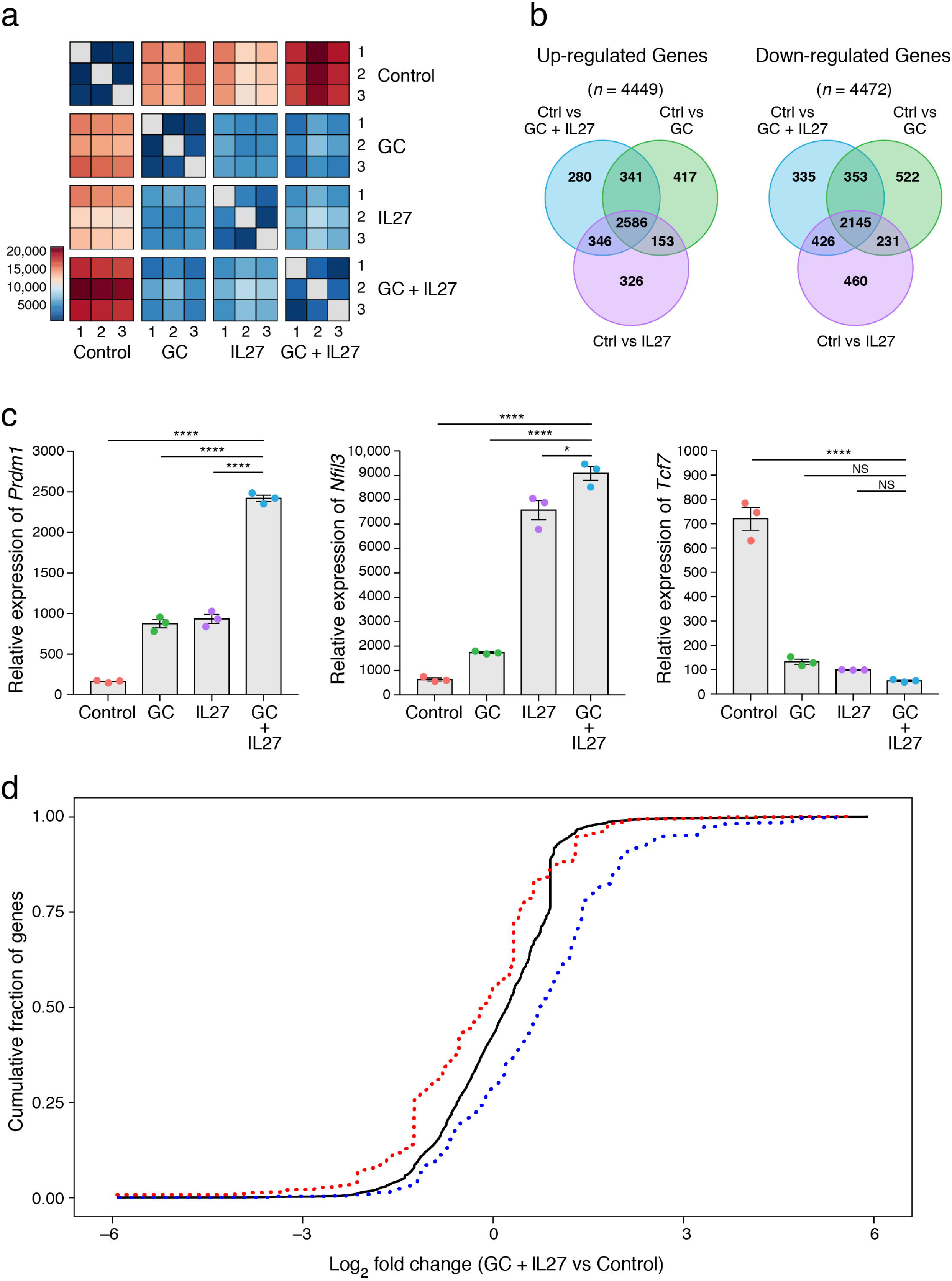
Effects of glucocorticoid and IL-27 in CD8^+^ T cells. Naïve CD8^+^ T cells were cultured *in vitro* with anti CD3/CD28 in the presence of Dex (GC), IL-27, or GC+IL-27. Cells were harvested on Day 9 for analysis. **a)** Heatmap display of the pairwise Euclidean distance between samples calculated for all genes**. b)** Venn plots showing the differentially expressed upregulated (left panel) and downregulated (right panel) genes between GC (Dex), IL-27, or GC+IL-27 treated cells relative to the control. **c)** Quantitative RT-PCR analysis of *Prdm1*, *Nfil3,* and *Tcf7* mRNA expression in the Ctrl, GC, IL-27, or GC+IL-27 treated cells. NS is not significant, *p < 0.05, ****p<0.0001. Ordinary one-way ANOVA (Tukey’s multiple comparisons test). Mean ± SEM are shown. **d)** Kolomogorov Smirnov one-sample curve(Tingey, 1951) showing overlap of genes down-regulated by GC+IL-27 with genes expressed in Tim-3^−^PD-1^−^ CD8+ TILs (p= 5.5×10^−16^), and genes up-regulated by GC+IL-27 with Tim-3^+^PD-1^+^ CD8^+^ TILs (p=7.7x 10^−16^).

**Figure S8:**
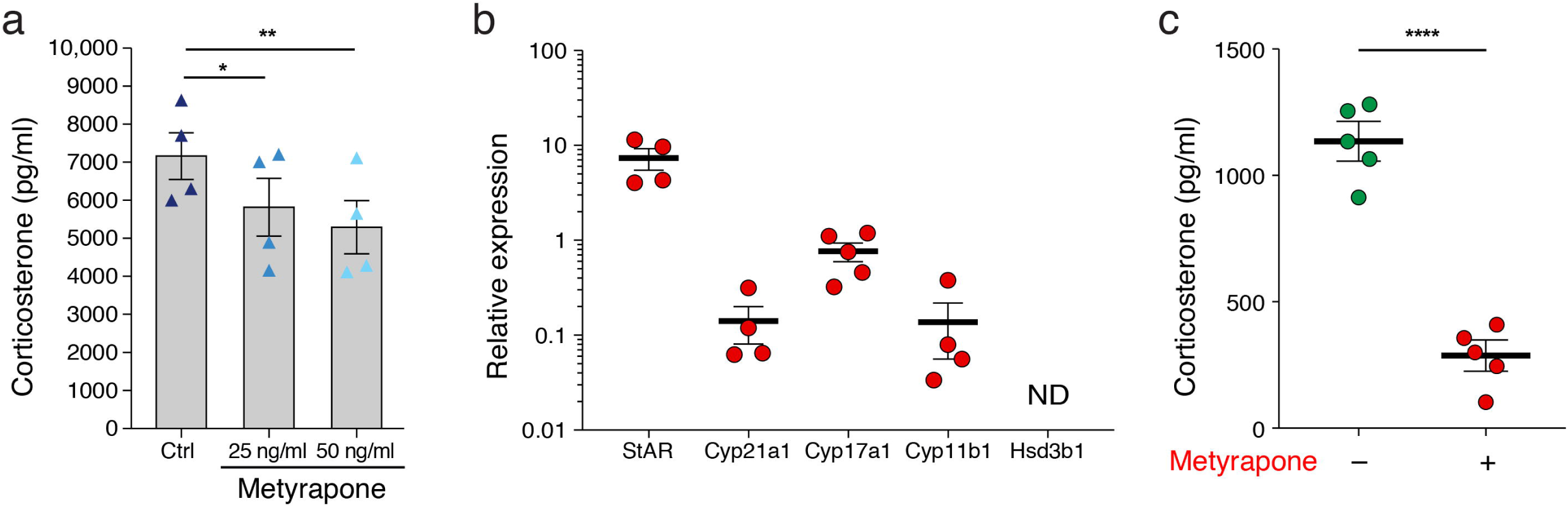
Monocyte/macrophage are the chief source of glucocorticoid in the TME. a) MC38-Ova tumor explants were cultured in 48 well plates in the presence or absence of metyrapone. Supernatants were harvested after 24hrs and the level of corticosterone was evaluated by ELISA. *p < 0.05, **P < 0.01, Ordinary one-way ANOVA (Tukey’s multiple comparisons test). Mean ± SEM are shown. b and c) Lin^−^CD45^+^CD24^−^ monocyte/macrophages were isolated from MC38-Ova tumors and b) RNA extracted for examination of the expression of the enzymes involved in steroid biogenesis by quantitative RT-PCR (c) cultured in the presence or absence of metyrapone. Supernatants were harvested after 24hrs and the level of corticosterone was evaluated by ELISA. ****p<0.0001, two-tailed Student’s t-test. Mean ± SEM are shown.

## Supplementary Tables

**Table S1.** Differentially expressed (DE) genes between control and glucocorticoid + IL-27 treated CD8^+^ T cells.

**Table S2.** Shared genes between the DE genes and T cell dysfunction signatures.

## Methods and Material

### Experimental methods

#### Mice

6–8 week old C57BL/6, *Nr3c1*^fl/fl^, Rag1^−/−^, E8iCre, WSX1^−/−^ and LysM-Cre transgenic mice were purchased from the Jackson Laboratory. NR3C1^fl/fl^ was crossed to E8iCre and/or E8iCre x WSX1^−/−^. Cryopreserved sperm from males bearing a targeted Cyp11a1 allele were obtained from EUCOMM and used to fertilize C57BL/6 oocytes. Heterozygote progeny were confirmed by PCR and bred to mice that express the FlpO recombinase (MMRC, UC Davis) to remove the neomycin resistance cassette followed by breeding with LysM-Cre. All mice were housed under SPF conditions. All experiments involving laboratory animals were performed under protocols approved by the Harvard Medical Area Standing Committee on Animals (Boston, MA).

#### Collection of colorectal carcinoma patient specimens

Primary colorectal carcinoma specimens were obtained under informed consent from untreated patients undergoing surgical resection at the Brigham and Women’s /Dana Farber Cancer Center and Massachusetts General Hospital (IRB protocol 03-189 and 02-240). Freshly resected CRC tumors and adjacent normal colon were recovered in Medium 199 (Thermo Fisher) supplemented with 2% heat-inactivated FCS (Sigma Aldrich) and stored briefly on ice.

#### Cell culture and treatment with glucocorticoid

CD8^+^ T cells from splenocytes and lymph nodes were isolated using CD8 microbeads (Miltenyi). Cells were further stained with antibodies against CD8, CD62L and CD44, and CD8^+^CD62L^hi^CD44^−^ naive cells were sorted by BD FacsAria (BD Biosciences). Sorted cells were cultured for 9 days as described below in DMEM supplemented with 10% (vol/vol) FCS, 50 mM mercaptoethanol, 1 mM sodium pyruvate, nonessential amino acids, L-glutamine, and 100 U/ml penicillin and 100 g/ml streptomycin. Specifically, naive CD8^+^CD62L^hi^CD44^lo^ cells were stimulated with plate bound anti-CD3 (145-2C11, 1 μg/ml) and anti-CD28 (PV-1, 1μg/ml) in the presence of either 10 nM dexamethasone (Sigma), 100 nM Corticosterone (Fisher Scientific), 25 ng/ml IL-27 (R&D), or both dexamethasone and IL27 for 3 days. Cells were then rested in the presence of 5 ng/ml IL2 (Miltenyi) for 3 days followed by re-stimulation with plate bound anti-CD3 (145-2C11, 1 μg/ml) and anti-CD28 (PV-1, 1μg/ml) in the presence of either 10nM dexamethasone (Sigma), 100 nM Corticosterone (Fisher Scientific), 25 ng/ml IL-27 (R&D), or both dexamethasone and IL27 for an additional 3 days.

#### Human CD8^+^ T cell culture

Peripheral blood was procured from healthy volunteers. Mononuclear cells were enriched by density gradient centrifugation on Ficoll-Paque PLUS (GE Healthcare) in SepMate-50 tubes (Stem Cell Technologies). CD8^+^ T cells were isolated from PBMCs using CD8 microbeads (Miltenyi) according to manufacturer protocol. Isolated CD8^+^ T cells were cultured for 9 days in RPMI supplemented with 10% (vol/vol) autologous heat-inactivated serum, 1 mM sodium pyruvate, 1X nonessential amino acids, 2mM L-glutamine, 100 U/ml penicillin, and 100 g/ml streptomycin. CD8^+^cells were stimulated with plate-bound anti-CD3 (Biolegend, clone UCHT1, 1 μg/ml) and anti-CD28 (Biolegend, clone CD28.2, 1μg/ml) in the presence of 10nM dexamethasone (Sigma) or vehicle control for 3 days. Cells were then rested in the presence of 100U/ml IL2 (R&D Systems) for 3 days. Next, the cells were re-stimulated with plate-bound anti-CD3 (1 μg/ml)and anti-CD28 (1μg/ml) in the presence of either 10nM dexamethasone (Sigma) or vehicle control for 3 days.

#### Tumor experiments

MC38-Ova was generously provided by Mark Smyth. B16F10 was purchased from ATCC. MC38-Ova-GFP was generated in our lab as follows, HEK293T cells were transfected with pLenti PGK GFP Puro plasmid. The resulting Lenti virus was then used to infect Mc38Ova cell line to generate GFP expressing cell line. MC38-Ova (0.5 x10^6^) or B16F10 (0.25×10^6^) and MC38-Ova-GFP (0.5 x10^6^) cells were implanted subcutaneously into the right flank of mice. Tumor size was measured in two dimensions by caliper and is expressed as the product of two perpendicular diameters.

#### Isolation of TILs

TILs were isolated by dissociating tumor tissue in the presence of collagenase D (2.5 mg/ml) for 20 minutes prior to centrifugation on a discontinuous Percoll gradient (GE Healthcare). Isolated cells were then used in various assays of T cell function.

#### Flow cytometry

Single cell suspensions were stained with antibodies against surface molecules. For murine samples, antibodies against CD4 (RM4-5), CD8 (53-6.7), CD107a (1D4B), PD-1 (RMP1-30) CD45 (30-F11), CD3 (145-2C11), CD19 (6D5), NK1.1 (V=PK136), Ly-6C (HK1.4), Ly-6G (1A8), CD11b (M1/70), CD11c (N418), CD24 (M1/69), I-A/I-E (M5/114.15.2), F4/80 (BM8) CD103 (2E7) were purchased from BioLegend. Antibodies against LAG-3 (clone C9B7W), Gzmb (NGZB) and Tigit (clone GIGD7) were purchased from eBioscience. Anti-Tim-3 (5D12) antibody was generated in house. Antibody against GR (G5) was purchased from Santa Cruz. Antibody against Siglec-F (E50-2440) was purchased from BD Biosciences. For human samples, antibodies against CD3 (UCHT1), CD8a (RPA-T8), Tim3 (F38-2E2), PD1 (EH12.2H7) and Lag3 (11C3C65) were purchased from Biolegend and antibody against TIGIT (MBSA43) was purchased from Thermo Fisher. Fixable viability dye eF506 (eBioscience) or Zombie UV dye (Biolegend) were used to exclude dead cells. For GR staining, eBioscience Foxp3/transcription factor staining buffer set was used as per manufacturer’s protocol. For intra-cellular cytokine (ICC) staining of CD8^+^ T cells in culture *in vitro*, cells were stimulated with 12-myristate 13-acetate (Coutinho and Chapman) (50ng/ml, Sigma-Aldrich, MO) and ionomycin (1mg/ml, Sigma-Aldrich, MO) in the presence of Golgi Plug (BD Biosciences) and Golgi Stop (BD Biosciences) for four hours prior to cell surface and ICC staining. For intra-cytoplasmic cytokine staining of TILs, cells were stimulated *in vitro* with 5 μg/ml OVA257-264 peptide for 4 hrs in the presence of Golgi stop (BD Biosciences) and Golgi Plug (BD Biosciences) prior to cell surface and ICC staining. Following fixation and permeabilization, staining with antibodies against the following was performed for murine samples: IL-2 (JES6-5H4), TNF-a (MP6-XT22), IFN-g (XMG-1.2), CD107a (1D4B) and Granzyme B (GB11) were purchased from Biolegend. Antigen specific T cells were determined by H-2Kb/ OVA257-264 dextramer staining following the manufacturer’s protocol (Immudex). All data were collected on a BD LsrII (BD Biosciences) or Fortessa (BD Biosciences) and analyzed with FlowJo 10.4.2 software (TreeStar).

#### Adoptive transfers

For adoptive transfer experiments, CD4^+^ (FOXP3^+^ and FOXP3^−^) and CD8^+^ T cells from either WT, *Nr3c1*^fl/fl^ E8iCre, WSX1^−/−^ or *Nr3c1*^fl/fl^ E8iCre^+^WSX1^−/−^ (dKO) mice were isolated by cell sorting using a BD FACSAria. A total of 1.5 × 10^6^ cells at a ratio of 1: 0.5 (CD4/CD8) was mixed in PBS and injected i.v. into Rag^−/−^ mice. Two days later, mice were implanted with MC38-Ova Colon carcinoma cells and followed for tumor growth.

#### In vivo and in vitro blockade of glucocorticoid synthesis

MC38-Ova was implanted in wild-type C57BL/6 mice and either Metyrapone (50mg/kg; Fisher Scientific) or vehicle control PBS (Gibco) was administered intra-tumorally on Day 5,6,7 and 9 post tumor implantation. In some experiments, MC38-Ova tumor explants or sorted lin^−^CD45^+^CD24^−^ cells were cultured in the presence or absence of Metyrapone (25 or 50 ng/ml) for 24hrs. Supernatants were harvested and corticosterone measured by ELISA (Arbor Assays).

#### Measurement of corticosterone in tissue extracts

Organic phase extraction using acetonitrile and hexane (1:2) was employed to extract steroids from tumor and spleen followed by examination of corticosterone by ELISA (Arbor Assays).

#### Luciferase assays

HEK293T cells were transfected with firefly luciferase reporter constructs for IL-10, PD1, Tim3, Lag3 or Tigit, together with Renilla luciferase reporter as internal control and plasmids expressing Nr3c1 or empty control vector. Dex or vehicle control was added to the culture 24hrs after transfection. Cells were analyzed at 48hrs with the dual luciferase assay kit (Promega). Fragments containing the proximal *Il10* promoter (−1.5 kb including the HSS^−^0.12 site), and the HSS^+^2.98 region followed by of the *Il10* minimal promoter were cloned into pGL4.10 Luciferase reporter plasmid (Promega). Fragments containing the *cis*-regulatory elements for the *Havcr2*, *Pdcd1* and *Lag3* loci were cloned into pGL4.23 Luciferase reporter plasmid (Promega).

#### Quantitative PCR

Total RNA was extracted using RNeasy columns (Qiagen). Reverse transcription of mRNA was performed in a thermal cycler (Bio-Rad) using iScript cDNA Synthesis Kit (Bio-Rad). qPCR was performed in the Vii7 Real-Time PCR system (Applied Biosystems) using the primers for Taqman gene expression (Applied Biosystems). Data were normalized to the expression of Actb.

#### RNA-Seq

1,000 cells were sorted into 5 µL of Buffer TCL (Qiagen) supplemented with 1% 2 mercaptoethanol. Plates were thawed on ice for one minute and spun down at 2,000 rpm for one minute. Immediately following, RNA lysate was purified using a 2.2x RNAClean SPRI bead ratio (Beckman Coulter Genomics). The RNA captured beads were processed using a modified SMART-Seq2 protocol(Picelli et al., 2013) entailing RNA secondary structure denaturation (72°C for three minutes), reverse transcription with Maxima Reverse Transcriptase (Life Technologies), and whole-transcription amplification (WTA) with KAPA HiFi HotStart ReadyMix 2X (Kapa Biosystems) for 11 cycles. WTA products were purified with Ampure XP beads (Beckman Coulter), quantified with a Qubit dsDNA HS Assay Kit (ThermoFisher), and quality accessed with a high-sensitivity DNA chip (Agilent). 0.2 ng of purified WTA product was used as input for the Nextera XT DNA Library Preparation Kit (Illumina). Uniquely barcoded libraries were pooled and sequenced with a NextSeq 500 high output V2 75 cycle kit (Illumina) using 38 and 38 paired end reads(Picelli et al., 2013).

### Computational analyses

#### Signature scoring in single cells

CD8^+^ TILs single-cell data were obtained and processed as previously described(Singer et al., 2016). Briefly, Briefly, paired reads were mapped to mouse annotation mm10 using Bowtie (Langmead et al., 2009) (allowing a maximum of one mismatch in seed alignment, and suppressing reads that had more than 10 valid alignments) and TPMs were computed using RSEM(Li and Dewey, 2011), and log2(TPM+1) values were used for subsequent analyses. Next, we filtered out low quality cells and cell doublets, maintaining for subsequent analysis the 588 cells that had (1) 1,000-4,000 detected genes (defined by at least one mapped read), (2) at least 200,000 reads mapped to the transcriptome, and (Cass et al.) at least 50% of the reads mapped to the transcriptome. Here, we restricted the genes considered in subsequent analyses to be the 7,790 genes expressed at log2(TPM+1) R 2 in at least ten percent of the cells. After removal of low-quality cells/genes, the data were normalized using quantile normalization followed by PCA. PCs 1-8 were chosen for subsequent analysis due to a drop in the proportion of variance explained following PC8. We used to visualize single cells in a two-dimensional non-linear embedding. To score each cell for a gene signature, expression data was initially scaled by calculating the z-score across each gene. For each gene signature, a cell-specific signature score was computed by first sorting the normalized scaled gene expression values for each cell followed by summing up the indices (ranks) of the signature genes. For signatures consisting of an upregulated and downregulated set of genes, two ranking scores were obtained separately, and the down-regulated associated signature score was subtracted from the up-regulated generated signature score. A contour plot was added on top of the tSNE space, which takes into account only those cells that have a signature score above the indicated threshold to further emphasize the region of highly scored cells.

#### RNA-Seq data pre-processing

RNA-seq reads were aligned using Tophat(Trapnell et al., 2009) (to mouse genome version mm9), and expression levels were calculated using RSEM(Li and Dewey, 2011) using annotated transcripts (mm9), followed by further processing using the Bioconductor package DESeq in R(Anders and Huber, 2010). The data was normalized using TMM normalization, and differentially expressed genes were defined using the differential expression pipeline on the raw counts with a single call to the function DESeq (FDR-adjusted *P* value <0.05). Heatmap figures were generated using pheatmap package(Kolde and Vilo, 2015) and clustered using Euclidian distance.

#### Analysis of additive and non-additive effects

To test whether the glucocorticoid and IL-27 signaling pathways had additive or non-additive effects on gene expression, we stimulated naïve CD8^+^ T cells in the presence of Dex, IL-27, or Dex+IL-27 *in vitro*. We tested for non-additive effects between IL-27 and glucocorticoid signaling using a negative binomial generalized linear model in order to account for both estimations of the mean and the dispersion across conditions, where dispersion describes the relationship between the mean and variance. The model was applied to the expression data using ANOVA between a model that takes into account the interaction between IL27 and Dex versus no interaction. We found that 1,675 out of 3,496 differentially expressed genes (adjusted P < 0.05, likelihood ratio test and false discovery rate (FDR) correction) between control and Dex+IL-27 stimulated CD8+ cells have non-additive effects.

#### Analysis of human TILs for GC+IL-27 signature

Data was downloaded from(Sade-Feldman et al., 2018) in a in log2(TPM+1) format. PCA was performed after removal of non-expressed genes. PCs 1-8 were chosen for subsequent analysis due to a drop in the proportion of variance explained following PC8. We used tSNE(Maaten, 2008) to visualize single cells in a two-dimensional non-linear embedding. GC+IL27 signature was projected onto single cell RNA profiles of TILs from 48 melanoma patients treated with checkpoint blockade (with 35 anti-PD-1, 11 anti-CTLA4+PD-1, and 2 anti-CTLA4 samples)(Sade-Feldman et al., 2018).

#### Statistical analysis

Significant differences between two groups were analyzed using GraphPad Prism 7 using the two-tailed unpaired Student’s t test or in case of multiple groups one-way or two-way ANOVA with multiple testing (Tukey). Tumor growth curves were analyzed using linear mixed effects models to test the trajectory of growth between various genotypes or treatments over time controlling for mouse. Differentially expressed genes following RNA-seq were defined using the differential expression pipeline on the raw counts with a single call to the function DESeq (FDR-adjusted P value <0.05). Values of *P < 0.05, **P < 0.01, ***P < 0.001 were considered statistically significant.

