## Supplementary material for "An endogenous glucocorticoid-cytokine signaling circuit promotes CD8^+^ T cell dysfunction in the tumor microenvironment": Table S1

| | CD8_1_WT_Ctrl_Day9 | CD8_2_WT_Ctrl_Day9 | CD8_3_WT_Ctrl_Day9 | CD8_1_WT_Dex_Day9 | CD8_2_WT_Dex_Day9 | CD8_3_WT_Dex_Day9 | CD8_1_WT_IL27_Day9 | CD8_2_WT_IL27_Day9 | CD8_3_WT_IL27_Day9 | CD8_1_WT_combo_Day9 | CD8_2_WT_combo_Day9 | CD8_3_WT_combo_Day9 | non-additive interaction |
|---|---|---|---|---|---|---|---|---|---|---|---|---|---|
| Dhx9 | 6.98 | 7.79 | 8.28 | 3.3 | 3.13 | 3.46 | 2.25 | 3.39 | 2.55 | 2.48 | 2.15 | 2.97 | + |
| Ssu72 | 139.67 | 137.69 | 144.5 | 119.1 | 130.19 | 127.37 | 126.32 | 132.54 | 121.93 | 115.52 | 123.3 | 127.64 | + |
| Psma4 | 231.27 | 242.27 | 232.42 | 116.48 | 123.25 | 125.96 | 115.66 | 128.13 | 133.73 | 85.67 | 88.73 | 100.09 | + |
| Bnip3l | 24.95 | 27.57 | 26.73 | 71.26 | 56.93 | 57.08 | 105.17 | 65.33 | 74.78 | 95.88 | 103.18 | 107.99 | + |
| Snip1 | 18.09 | 20.97 | 20.84 | 14.16 | 14.1 | 14.3 | 13.1 | 15.03 | 12.92 | 13.99 | 11.64 | 13.48 | + |
| Ppia | 4507.35 | 4877.36 | 4634.24 | 2576.63 | 2725.51 | 2835.8 | 2841.43 | 2947.02 | 3060.9 | 1986.64 | 2042.19 | 2129.37 | + |
| Rplp2 | 970.55 | 984.22 | 1010.65 | 1437 | 1410.35 | 1474.11 | 1425.85 | 1352.05 | 1346.45 | 1544.44 | 1653.7 | 1583.11 | + |
| Snhg4 | 32.29 | 37.41 | 35.7 | 24.27 | 25.62 | 26.71 | 21.77 | 24.29 | 24.84 | 21.22 | 24.3 | 17.71 | + |
| Psmg2 | 90.94 | 85.54 | 92.36 | 53.94 | 64.96 | 63.41 | 61.15 | 68.38 | 65.42 | 51.97 | 53.95 | 51.44 | + |
| Mettl22 | 21.87 | 23.84 | 25.61 | 21.71 | 26.62 | 25.26 | 20.66 | 30.91 | 26.76 | 16.2 | 16.61 | 16.38 | + |
| Ifi30 | 28.48 | 27.64 | 26.51 | 20.94 | 22.33 | 16.51 | 15.7 | 19.28 | 17.88 | 18.78 | 19.6 | 15.92 | + |
| Hprt | 273.11 | 270.8 | 272.18 | 188.21 | 192.28 | 194.5 | 178.16 | 200.35 | 191.75 | 145.62 | 148.24 | 148.62 | + |
| Chst12 | 35.74 | 32.48 | 33.87 | 55.1 | 52.32 | 51.47 | 53.69 | 52.98 | 48.59 | 56.37 | 54.94 | 65.17 | + |
| Nmt1 | 68.15 | 68.96 | 68.84 | 47.7 | 47.56 | 50.66 | 51.99 | 51.96 | 52.03 | 46.9 | 58.76 | 42.37 | + |
| Pbx4 | 4.55 | 4.32 | 3.81 | 15.58 | 11.07 | 14.11 | 18.85 | 14.08 | 14.54 | 19.17 | 21.69 | 19.75 | + |
| Aig1 | 9.37 | 10.21 | 9.39 | 10.07 | 12.08 | 9.86 | 17.72 | 18.78 | 18.38 | 13.57 | 16.69 | 15.42 | + |
| l7Rn6 | 48.3 | 48.44 | 49.3 | 32.35 | 30.11 | 33.25 | 34.64 | 35.86 | 35.53 | 29.78 | 32.43 | 32.88 | + |
| Dda1 | 61.55 | 63.24 | 60.13 | 38.61 | 42.39 | 41.43 | 43.98 | 45.75 | 48.75 | 37.74 | 44.52 | 42.29 | + |
| Cacfd1 | 8.55 | 9.13 | 8.52 | 17.16 | 16.91 | 15.56 | 15.87 | 12.65 | 15.04 | 17.55 | 13.94 | 18.81 | + |
| Nt5dc1 | 12.88 | 12.66 | 12.42 | 13.87 | 11.24 | 11.51 | 13.65 | 12.67 | 12.74 | 16.66 | 17.22 | 15.67 | + |
| A930016O22Rik | 0.19 | 0.12 | 0.14 | 0.51 | 0.4 | 0.65 | 0.8 | 0.8 | 0.6 | 0.6 | 0.83 | 0.37 | + |
| Dnajc7 | 63.19 | 65.4 | 63.17 | 58.4 | 54.03 | 52.79 | 53.89 | 54.08 | 50 | 53.11 | 53.77 | 53.04 | + |
| Tor3a | 6.81 | 7.34 | 7.78 | 11.93 | 10.44 | 9.99 | 10.43 | 9.31 | 9.01 | 10.01 | 9.67 | 10.83 | + |
| Ccdc134 | 12.57 | 11.67 | 10.83 | 8.59 | 9.2 | 9.39 | 8.99 | 7.21 | 9.16 | 8.44 | 8.02 | 9.7 | + |
| Nup37 | 53.27 | 57.75 | 54.83 | 28.45 | 31.23 | 26.5 | 20 | 25.27 | 23.14 | 17.88 | 19.32 | 17.07 | + |
| Sppl2b | 10.9 | 11.36 | 11.02 | 4.63 | 4.11 | 5.29 | 4.55 | 5.65 | 5.33 | 6.4 | 3.72 | 4.64 | + |
| Aimp1 | 86.51 | 87.59 | 89.68 | 94.13 | 87.4 | 92.76 | 105.71 | 93.02 | 96.09 | 101.56 | 122.27 | 114.36 | + |
| Rpl24 | 1114.43 | 1148.32 | 1121.51 | 1851.8 | 1819.97 | 1960.69 | 1918.74 | 1710.32 | 1796.85 | 2089.35 | 2284.29 | 2148.95 | + |
| Malat1 | 60.16 | 51.62 | 53.19 | 121.37 | 123.81 | 134.75 | 182.32 | 141.28 | 156.2 | 182.17 | 186.83 | 197.01 | + |
| Dock6 | 0.33 | 0.6 | 0.41 | 0.93 | 0.95 | 1.45 | 0.81 | 0.79 | 0.88 | 0.94 | 0.82 | 1.25 | + |
| Timp1 | 34.59 | 24.12 | 36.96 | 30.66 | 51.74 | 41.37 | 17.67 | 43.85 | 35.11 | 12.31 | 7.01 | 7.01 | + |
| Wls | 9.88 | 8.77 | 8.53 | 15.71 | 12.86 | 14.8 | 7.25 | 5.09 | 5.39 | 3.81 | 3.37 | 6.35 | + |
| Wdr3 | 45.08 | 47.05 | 48.46 | 24 | 22.68 | 19.14 | 16.29 | 22.92 | 23.35 | 16.45 | 15.73 | 17.25 | + |
| Ap1s1 | 92.9 | 88.44 | 93.7 | 54.05 | 55.71 | 53.78 | 50.17 | 60.62 | 56.76 | 39.06 | 40.01 | 38.05 | + |
| Ptprs | 10.3 | 10.63 | 10.87 | 13.85 | 13.34 | 13.08 | 14.93 | 18.09 | 17.37 | 15.79 | 16.61 | 16.23 | + |
| Tubb4b | 394.29 | 335.49 | 334.47 | 136.95 | 165.7 | 160.31 | 108.11 | 110.2 | 104.31 | 95.79 | 98.56 | 93.54 | + |
| Lta4h | 83.78 | 84.02 | 81.99 | 58.66 | 57.35 | 59.02 | 77.77 | 75.76 | 73.18 | 53.68 | 61.05 | 59.51 | + |
| Il10 | 0.57 | 1.08 | 0.25 | 81.3 | 76.04 | 69.05 | 4.4 | 5.31 | 8.09 | 65.74 | 51.01 | 50.49 | + |
| Rpph1 | 13.17 | 20.59 | 22.29 | 0.97 | 1.34 | 1.37 | 1.94 | 1.27 | 1.82 | 1.58 | 2.09 | 1.77 | + |
| Caprin1 | 120.3 | 117.15 | 124.3 | 78.8 | 76.41 | 77.82 | 77.97 | 80.62 | 78.01 | 80.31 | 91.24 | 78.73 | + |
| Mdm4 | 14.35 | 15.14 | 14.73 | 23.41 | 23.99 | 22.31 | 20.17 | 22.43 | 22.23 | 22.76 | 24.02 | 20.12 | + |
| Senp3 | 17.89 | 19.13 | 18.18 | 8.03 | 8.85 | 10.28 | 7.7 | 9.62 | 6.84 | 9.17 | 6.3 | 8.49 | + |
| Sytl1 | 17.02 | 18.2 | 18.77 | 49.38 | 40.56 | 44.92 | 31.28 | 21.97 | 22.96 | 40.95 | 44.37 | 45.46 | + |
| Zdhhc16 | 24.88 | 27.47 | 24.59 | 22.76 | 17.91 | 21.61 | 20.6 | 19.8 | 19.31 | 20.12 | 17.44 | 20.32 | + |
| Mrpl1 | 31.39 | 29.54 | 30.49 | 15.25 | 16.68 | 18.14 | 21.73 | 25.05 | 23.19 | 14.36 | 16 | 14.67 | + |
| Eva1b | 14.41 | 17.08 | 14.42 | 67.2 | 47.95 | 48.8 | 46.42 | 33.09 | 40.66 | 52.94 | 47.43 | 51.76 | + |
| Fas | 24.76 | 20.59 | 24.21 | 66.57 | 56.13 | 64.01 | 51.33 | 48.73 | 47.68 | 67.74 | 74.3 | 67.7 | + |
| Ube2m | 10.1 | 8.16 | 9.41 | 5.68 | 5.01 | 4.19 | 4.56 | 4.6 | 4.25 | 3.37 | 2.68 | 3.34 | + |
| Trim24 | 1.25 | 1.27 | 1.03 | 3.18 | 2.8 | 2.52 | 1.7 | 1.47 | 1.53 | 2.34 | 2.47 | 2.21 | + |
| Pla2g12a | 23.34 | 23.6 | 28.81 | 26.66 | 33.86 | 35.48 | 41.81 | 45.24 | 44.6 | 32.82 | 34.31 | 30.56 | + |
| Rbm25 | 33.57 | 35.39 | 34.61 | 27.4 | 27.36 | 26.75 | 24.71 | 22.95 | 26.07 | 28.4 | 28.08 | 27.01 | + |
| Ran | 723.69 | 750.54 | 747.2 | 299.46 | 350.12 | 352.54 | 279.75 | 374.64 | 352.99 | 211.26 | 219.89 | 204.88 | + |
| Cep63 | 7.32 | 6.77 | 7.06 | 3.68 | 5.3 | 5.34 | 5.01 | 5.34 | 4.79 | 3.69 | 4.5 | 4.25 | + |
| Fam173b | 11.95 | 13.62 | 13.73 | 7.37 | 7.23 | 7.22 | 7.29 | 7.4 | 6.58 | 5.74 | 5.71 | 8.45 | + |
| Dmap1 | 25.81 | 25.08 | 24.34 | 20.55 | 19.39 | 19.77 | 18.8 | 18.9 | 20.99 | 17.66 | 19.1 | 19.39 | + |
| Rufy1 | 1.18 | 1.34 | 1.24 | 2.52 | 2.8 | 2.33 | 3.57 | 2.72 | 2.54 | 2.2 | 2.51 | 2.36 | + |
| Paf1 | 31.31 | 31.34 | 30.73 | 22.67 | 19.7 | 21.36 | 17.68 | 18.13 | 16.92 | 20.86 | 20.83 | 18.41 | + |
| Rfx3 | 0.65 | 0.56 | 0.52 | 1.24 | 1.26 | 1.25 | 1.28 | 0.86 | 1 | 1.36 | 1.2 | 1.28 | + |
| Tmco1 | 19 | 20.99 | 18.44 | 15.52 | 15.3 | 14.95 | 14.96 | 16.94 | 15.76 | 15.2 | 15.82 | 16.39 | + |
| Qtrt1 | 26.62 | 27.71 | 26.81 | 19.42 | 17.15 | 19.51 | 20.1 | 19.79 | 17.13 | 19.79 | 17.4 | 14.84 | + |
| Taf6l | 12.48 | 10.89 | 11.97 | 5.47 | 6.37 | 5.76 | 3.38 | 4.9 | 5.48 | 4.17 | 3.81 | 3.35 | + |
| Htt | 3.32 | 2.79 | 3.14 | 3.87 | 4.19 | 4.05 | 3.8 | 4.36 | 4 | 4.09 | 3.76 | 4.04 | + |
| Got1 | 82.22 | 87.84 | 88.02 | 83.06 | 89.87 | 88.52 | 59.7 | 83.29 | 78.92 | 50.84 | 54.9 | 51.17 | + |
| Zfp688 | 8.43 | 9.66 | 8.09 | 17.27 | 14.44 | 16.01 | 14.37 | 16.41 | 14.42 | 16.72 | 17.53 | 20.6 | + |
| Crispld2 | 0.76 | 0.36 | 1.19 | 9.85 | 6.28 | 8.04 | 2.2 | 2.66 | 2.47 | 1.53 | 2.63 | 2.54 | + |
| Ptgir | 0.45 | 0.59 | 0.47 | 0.49 | 0.43 | 0.52 | 0.48 | 0.37 | 0.74 | 0.24 | 0.11 | 0.07 | + |
| Cd97 | 24.97 | 22.06 | 24.72 | 52.02 | 43.85 | 46.04 | 46.24 | 39.64 | 36.14 | 56.56 | 56.46 | 60.03 | + |
| Cysltr2 | 0.72 | 1.03 | 1.83 | 5.22 | 5.15 | 4.17 | 2.27 | 2.15 | 2.37 | 4.36 | 3.78 | 5.19 | + |
| Srsf3 | 320.48 | 344.49 | 332.74 | 131.29 | 141.08 | 140.98 | 89.51 | 125.67 | 117.47 | 81.63 | 86.15 | 85.05 | + |
| Chchd2 | 716.33 | 681.18 | 673.45 | 439.88 | 450.69 | 445.34 | 460.7 | 508.53 | 476.15 | 376.68 | 396.42 | 403.51 | + |
| Hist1h4i | 13.46 | 16.74 | 17.69 | 7.39 | 8.25 | 6.75 | 10.83 | 9.24 | 8.48 | 7.48 | 5.73 | 7.46 | + |
| Otud6b | 32.7 | 32.33 | 33.71 | 14.2 | 19.02 | 18.39 | 12.47 | 17.18 | 16.25 | 12.82 | 12.93 | 11 | + |
| Kbtbd4 | 19.06 | 18.38 | 20.45 | 15.66 | 14.79 | 16.01 | 15.78 | 13.6 | 14.39 | 13.24 | 14.63 | 15.18 | + |
| Gm10094 | 149.89 | 162.73 | 162.24 | 96.68 | 100.76 | 96.98 | 105.68 | 107.97 | 109.56 | 89.74 | 88.37 | 92.45 | + |
| Rab2b | 5.3 | 4.69 | 4.96 | 7.18 | 7.63 | 7.77 | 7.03 | 6.7 | 6.32 | 6.75 | 8.27 | 8.3 | + |
| Abcb6 | 10.67 | 9.71 | 9.68 | 3.9 | 5.09 | 3.61 | 4.92 | 5.67 | 5.4 | 4.19 | 3.53 | 3.18 | + |
| Tbc1d14 | 8.69 | 9.51 | 8.96 | 28.14 | 24.33 | 27.82 | 14.66 | 12.37 | 13.44 | 29 | 25.28 | 27.58 | + |
| Ccnl2 | 27.31 | 34 | 36.11 | 86.67 | 80.19 | 74.95 | 78.2 | 67.26 | 70.5 | 99.65 | 109.39 | 98.55 | + |
| Pbk | 34.26 | 31.04 | 31.86 | 23.79 | 23.78 | 23.28 | 9.18 | 7.92 | 7.92 | 9.43 | 8.31 | 7.11 | + |
| Lig1 | 47.7 | 48.45 | 44.67 | 28.17 | 33.61 | 30.67 | 22.16 | 21.4 | 20.08 | 18.16 | 20.54 | 19.23 | + |
| Haao | 14.37 | 11.98 | 8.44 | 39.35 | 35.85 | 36.53 | 42.15 | 34.32 | 33.8 | 43.18 | 46.07 | 49.5 | + |
| Cux1 | 4.88 | 4.53 | 4.31 | 9.33 | 8 | 8.05 | 9.16 | 8.64 | 8.22 | 10.54 | 10.96 | 10.6 | + |
| Phf11b | 42.4 | 40.62 | 39.94 | 34.48 | 29.28 | 35.08 | 72.28 | 41.22 | 46.38 | 64.51 | 62.86 | 71.08 | + |
| Alg8 | 33.35 | 36.44 | 32.06 | 14.82 | 17.16 | 18.01 | 10.79 | 15.88 | 14.02 | 9.92 | 8.87 | 9.13 | + |
| Cox15 | 5.81 | 4.62 | 5.26 | 3.45 | 2.87 | 3.45 | 2.89 | 3.31 | 3.62 | 2.38 | 2.9 | 2.91 | + |
| Rad54l2 | 4.43 | 4.03 | 4.39 | 2.69 | 3.26 | 3.12 | 3.16 | 3.7 | 3.64 | 2.82 | 3.37 | 2.69 | + |
| Vmn2r-ps129 | 66.81 | 78.38 | 67.72 | 19.52 | 27.4 | 20.42 | 25.83 | 31 | 26.53 | 17.63 | 19.8 | 17.65 | + |
| Dhdds | 12.18 | 12.22 | 13.14 | 10.21 | 11.74 | 10.77 | 10.69 | 10.29 | 9.56 | 9.05 | 10.15 | 10.62 | + |
| Zyg11b | 4.23 | 3.52 | 3.99 | 6.1 | 4.68 | 5.11 | 6.24 | 4.84 | 4.76 | 9.75 | 8.51 | 10.66 | + |
| Samhd1 | 38.97 | 37.06 | 36.31 | 130.31 | 127.11 | 132.15 | 148.24 | 115.88 | 118.36 | 172.06 | 172.79 | 180.91 | + |
| Apoa1bp | 60.37 | 64.87 | 58.96 | 99.1 | 100.01 | 105.14 | 92.2 | 103.25 | 101.68 | 96.51 | 105.35 | 91.4 | + |
| Mthfd1 | 66.67 | 73.12 | 71.37 | 20.44 | 25.67 | 24.77 | 18.65 | 27.95 | 25.58 | 14.85 | 14.02 | 14.27 | + |
| Chd2 | 1.29 | 1.84 | 1.81 | 2.98 | 3.06 | 3.11 | 3.63 | 3.6 | 3.33 | 3.96 | 3.69 | 3.16 | + |
| Noxo1 | 0.28 | 0.21 | 0.27 | 1.38 | 1.03 | 1.03 | 1.21 | 0.5 | 0.86 | 0.92 | 1.02 | 1.31 | + |
| Fanca | 3.94 | 3.92 | 4.41 | 1.72 | 2.23 | 2.68 | 1.51 | 2.01 | 1.67 | 1.3 | 1.46 | 1.36 | + |
| Chmp2a | 82.89 | 88.45 | 85.51 | 107.11 | 116.69 | 110.55 | 131.05 | 116.58 | 116.68 | 111.33 | 108.3 | 112.92 | + |
| Polr2l | 39.78 | 36.97 | 36.04 | 16.73 | 15.38 | 16.95 | 12.85 | 17.58 | 15.01 | 13.65 | 12.61 | 13.59 | + |
| Map4k2 | 20.72 | 21.5 | 20.54 | 31.45 | 29.07 | 28.26 | 34.92 | 31.47 | 31.18 | 35.61 | 33.33 | 36.98 | + |
| Pbdc1 | 36.89 | 38.04 | 37.04 | 18.9 | 20.24 | 19.41 | 16.98 | 19.42 | 19.87 | 14.72 | 15.77 | 14.96 | + |
| Urb1 | 3.02 | 3.78 | 3.76 | 1.72 | 2.01 | 2.2 | 1.33 | 1.7 | 1.44 | 1.17 | 1.71 | 1.78 | + |
| Gtf3c1 | 9.28 | 9.5 | 10.97 | 7.94 | 7.82 | 7.67 | 6.7 | 8.34 | 8.15 | 8.14 | 7.63 | 7.06 | + |
| Thoc7 | 111.55 | 121.81 | 116.92 | 62.73 | 72.41 | 72.96 | 70.1 | 73.41 | 76.27 | 63.08 | 61.08 | 66.47 | + |
| Nom1 | 21.65 | 20.63 | 18.93 | 12.97 | 13.38 | 13.23 | 9.38 | 11.5 | 12.99 | 10.79 | 9.68 | 12.4 | + |
| Lman2 | 67.31 | 73.4 | 70.99 | 47.73 | 48.57 | 48.93 | 51.63 | 54.09 | 52.32 | 46.96 | 44.59 | 47.34 | + |
| Tsen54 | 13.78 | 13.21 | 12.88 | 10.61 | 9.55 | 10.8 | 10.9 | 11.03 | 10.12 | 9.08 | 9.21 | 9.16 | + |
| BC017643 | 23 | 20.79 | 22.61 | 36.69 | 40.73 | 36.79 | 42.92 | 32.32 | 38.82 | 46.53 | 45.3 | 47.7 | + |
| Taf12 | 56.59 | 59.62 | 58.15 | 42.32 | 47.53 | 47.99 | 44.11 | 45.8 | 43.2 | 41.52 | 44.32 | 45.93 | + |
| Ept1 | 9.52 | 9.68 | 9.93 | 6.83 | 5.61 | 5.92 | 5.16 | 6.7 | 6.96 | 5.87 | 6.6 | 6.07 | + |
| Ints8 | 3.02 | 4.4 | 4.29 | 2.77 | 2.26 | 2.61 | 2.52 | 2.09 | 2.29 | 2.17 | 2.39 | 1.82 | + |
| Luc7l3 | 20.61 | 23.73 | 23.19 | 21.87 | 23.84 | 25.12 | 24.16 | 20.92 | 24.78 | 29.4 | 29.98 | 27.06 | + |
| 2900056M20Rik | 3.62 | 2.84 | 3.3 | 5.14 | 5.69 | 5.37 | 5.74 | 5.8 | 7.25 | 5.42 | 6.3 | 6.3 | + |
| Pus7l | 6.61 | 7.76 | 6.51 | 1.8 | 2.43 | 2.42 | 1.88 | 2.95 | 2.58 | 1.67 | 1.41 | 1.07 | + |
| Rps27a | 1786.09 | 1854.06 | 1844.8 | 2284.48 | 2272.36 | 2361.59 | 2313.52 | 2276.8 | 2261.18 | 2198.38 | 2515.94 | 2205 | + |
| Rpf1 | 67.9 | 60.71 | 63.88 | 37.77 | 43.44 | 40.17 | 36.06 | 47.6 | 47.43 | 34.9 | 41.15 | 33.01 | + |
| Ndrg4 | 3.24 | 2.61 | 2.38 | 6.56 | 5.31 | 5.78 | 7.65 | 6.54 | 6.38 | 7.76 | 9.35 | 7.4 | + |
| Lamtor3 | 39.61 | 46.82 | 40.97 | 26.09 | 27.53 | 24.93 | 23.93 | 27.54 | 23.77 | 23.04 | 22.54 | 22.65 | + |
| Man1c1 | 0.05 | 0.06 | 0.12 | 0.44 | 0.13 | 0.38 | 0.66 | 0.89 | 1.01 | 0.36 | 0.39 | 0.53 | + |
| Ankrd27 | 20.48 | 21.56 | 21.33 | 14.48 | 15.33 | 16.09 | 13.99 | 14.12 | 14.88 | 13.61 | 13.74 | 14.68 | + |
| Cdc123 | 89.9 | 90.79 | 88.81 | 55.05 | 57.68 | 55.93 | 52.9 | 60.43 | 52.19 | 47.38 | 46.27 | 45.35 | + |
| Rangap1 | 79.11 | 74.53 | 75.07 | 61.66 | 65.92 | 64 | 53.86 | 57.97 | 53.22 | 49.64 | 50.76 | 46.61 | + |
| Mmadhc | 70.29 | 69.94 | 67.82 | 45.87 | 46.94 | 42.17 | 47.18 | 52.78 | 55.26 | 40.6 | 38.77 | 40.9 | + |
| Mybbp1a | 117.79 | 121.6 | 130.5 | 44.73 | 49.91 | 47.08 | 31.94 | 53.65 | 49 | 29.53 | 24.64 | 25.99 | + |
| Cndp2 | 159.28 | 171.78 | 158.05 | 202.98 | 204.3 | 207.88 | 122.63 | 156.36 | 157.47 | 77.85 | 96.74 | 78.72 | + |
| Pcna | 223.46 | 243.51 | 227.91 | 86.59 | 92.87 | 89.18 | 72.07 | 82.28 | 78.84 | 46.26 | 51.01 | 53.44 | + |
| Lias | 35.87 | 36.25 | 35.68 | 29.54 | 29.71 | 31.14 | 25.67 | 28.59 | 27.6 | 24.68 | 25.09 | 25.16 | + |
| Ipo13 | 12.23 | 12.16 | 12.34 | 8.1 | 7.34 | 10.22 | 6.1 | 8.38 | 9.61 | 8.93 | 7.86 | 7.34 | + |
| 2210404J11Rik | 8.12 | 7.73 | 8.79 | 8.58 | 8.43 | 8.66 | 7.56 | 7.82 | 8.49 | 6.11 | 6.05 | 5.46 | + |
| Gm13212 | 5.88 | 7.93 | 6 | 10.29 | 12.29 | 11.12 | 12.44 | 13.51 | 10.85 | 15.36 | 16.07 | 13.95 | + |
| Smarca2 | 2.78 | 2.73 | 2.56 | 7.05 | 4.83 | 4.86 | 6.61 | 4.5 | 6.22 | 6.87 | 6.79 | 6.23 | + |
| Ppp2r1a | 181.82 | 179.98 | 184.15 | 153.44 | 157.88 | 160.81 | 151.33 | 153.97 | 155.69 | 147.35 | 148.48 | 152.73 | + |
| Wdr34 | 13.83 | 13.86 | 15.87 | 10.22 | 11.38 | 11.41 | 10.24 | 9.6 | 10.38 | 10.86 | 11.31 | 10.36 | + |
| Pdcd10 | 73.95 | 76.01 | 80.28 | 55.33 | 55.45 | 52.94 | 59.53 | 58.09 | 59.69 | 54.67 | 51.12 | 53.77 | + |
| Gsg2 | 9.12 | 7.31 | 8.31 | 5.46 | 6.57 | 6.4 | 4.97 | 3.91 | 4.04 | 4.36 | 4.36 | 3.59 | + |
| 1810009A15Rik | 141 | 136.15 | 143.65 | 77.28 | 85.75 | 91.46 | 68.12 | 83.85 | 80.94 | 61.49 | 60.93 | 68.28 | + |
| Cdipt | 51.39 | 54.24 | 52.29 | 36.83 | 39.67 | 36.92 | 42.41 | 45.05 | 48.43 | 39.92 | 39.12 | 44.95 | + |
| Isoc2b | 3.65 | 3.55 | 3.91 | 10.09 | 10.88 | 6.53 | 13.81 | 7.89 | 8.46 | 12.78 | 11.86 | 13.42 | + |
| Aplp1 | 0.93 | 0.44 | 0.66 | 2.54 | 1.96 | 1.63 | 2.88 | 1.87 | 3.43 | 3.18 | 3.38 | 3.83 | + |
| Usp3 | 30.09 | 29.23 | 27.5 | 83.45 | 63.93 | 72.35 | 75.93 | 56.03 | 60.38 | 81.07 | 91.95 | 87.5 | + |
| Amz2 | 8.8 | 7.15 | 8.39 | 23.02 | 22.36 | 24.16 | 20.05 | 13.72 | 16.94 | 17.82 | 21.05 | 20.41 | + |
| Rlf | 1.53 | 1.85 | 1.6 | 3.42 | 3.71 | 3.47 | 2.54 | 2.97 | 2.44 | 4 | 3.94 | 3.59 | + |
| Cct8 | 476.85 | 479.12 | 469.85 | 220.94 | 238.81 | 245.6 | 210.75 | 270.62 | 247.34 | 179.57 | 205.08 | 183.8 | + |
| Cyp4f13 | 6.95 | 5.07 | 5.47 | 14.18 | 15.61 | 16.25 | 12.93 | 14.07 | 11.63 | 11.94 | 16.18 | 16.87 | + |
| Pa2g4 | 253.43 | 263.74 | 267.85 | 121.16 | 128.6 | 129.85 | 108.2 | 140.99 | 125.36 | 89.52 | 93.95 | 88.02 | + |
| Sh2b3 | 1.38 | 1.99 | 3.16 | 3.71 | 3.9 | 3.03 | 1.48 | 1.66 | 2.49 | 1.04 | 1.22 | 0.72 | + |
| Pigq | 25.7 | 27.59 | 25.99 | 18.69 | 18.12 | 19.15 | 15.44 | 17.48 | 17.5 | 17.1 | 13.25 | 15.24 | + |
| Cox6c | 421.2 | 410.78 | 400.72 | 328.82 | 354.17 | 341.81 | 377.35 | 366.4 | 325.85 | 294.78 | 327.88 | 329.01 | + |
| Eif3b | 23.75 | 24.97 | 24.34 | 11.94 | 11.39 | 12.41 | 10.87 | 12.83 | 13.11 | 10.61 | 11 | 10.53 | + |
| Rnf167 | 12.17 | 14.39 | 16.99 | 39.99 | 33.46 | 33.38 | 55.64 | 39.21 | 49.78 | 57.77 | 60.34 | 63.85 | + |
| Kdelr3 | 1.9 | 2.18 | 2.49 | 5.01 | 7.2 | 5.01 | 10.57 | 11.55 | 9.33 | 8.46 | 11.94 | 10.07 | + |
| Cdv3 | 105.17 | 95.92 | 85.83 | 63.81 | 74.44 | 58.73 | 63.87 | 66.97 | 68.2 | 54.84 | 70.89 | 56.73 | + |

| | | | | | | | | | | | | | |
|---|---|---|---|---|---|---|---|---|---|---|---|---|---|
| Gstt1 | 17.39 | 17.67 | 18.87 | 17.86 | 23.3 | 19.49 | 41.39 | 47.52 | 41.75 | 26.64 | 32.75 | 27.74 | + |
| Limk1 | 0.99 | 0.98 | 1.17 | 2.13 | 2.29 | 2.21 | 1.85 | 1.5 | 2.29 | 2.39 | 1.71 | 2.38 | + |
| Bend3 | 1.5 | 1.8 | 2 | 0.67 | 0.88 | 0.38 | 0.3 | 0.45 | 0.37 | 0.6 | 0.25 | 0.27 | + |
| Prkar2a | 6.47 | 5.95 | 6.28 | 3.68 | 3.55 | 3.87 | 2.97 | 3.56 | 3.69 | 3.53 | 2.72 | 2.91 | + |
| Hnrnpa2b1 | 394.31 | 431.61 | 418.19 | 184.52 | 198.35 | 188.93 | 140.81 | 184.56 | 165.02 | 137.78 | 126.68 | 134.03 | + |
| Kif22 | 41.26 | 38.54 | 35.99 | 30.32 | 36.96 | 32.78 | 19.32 | 18.41 | 16.52 | 16.29 | 21.9 | 17.12 | + |
| Adap1 | 7.09 | 8.76 | 7.08 | 2.32 | 2.06 | 2.05 | 7.67 | 5.21 | 7.38 | 3.51 | 3.44 | 3.7 | + |
| P4ha2 | 9.39 | 6.2 | 8.01 | 17.04 | 14.21 | 16.95 | 46.82 | 21.45 | 34.59 | 34.39 | 33.86 | 36.57 | + |
| Snrpa1 | 113.22 | 116.7 | 114.97 | 64.15 | 75.71 | 78.36 | 70.57 | 91.32 | 82.06 | 57.85 | 73.91 | 61.87 | + |
| Fth1 | 950.09 | 994.32 | 998.17 | 2276.95 | 2350.76 | 2420.82 | 2178.71 | 1449.21 | 1711.04 | 2135.87 | 2278.17 | 2344.37 | + |
| Cmpk1 | 73.49 | 73.6 | 70.26 | 48.71 | 48.04 | 48.42 | 61.33 | 62.55 | 60.82 | 50.63 | 53.55 | 53.96 | + |
| B630005N14Rik | 1.13 | 1.16 | 0.9 | 2.11 | 2.11 | 2.25 | 2.61 | 2.02 | 2.07 | 2.21 | 3.24 | 3.08 | + |
| Ccar1 | 28.04 | 29.13 | 28.37 | 16.17 | 17.58 | 17.65 | 13.24 | 16.85 | 15.53 | 16.18 | 18.41 | 13.37 | + |
| Lrp12 | 2.44 | 2.98 | 2.18 | 4.51 | 3.87 | 3.78 | 4.19 | 4.57 | 5.81 | 4.34 | 3.11 | 3.79 | + |
| Rnf34 | 20.16 | 19.74 | 19.59 | 14.69 | 14.12 | 15.31 | 12.82 | 15.03 | 14.55 | 13.64 | 14.13 | 13.81 | + |
| Lysmd4 | 2.2 | 1.79 | 1.91 | 3.3 | 2.87 | 3.62 | 3.81 | 3.07 | 3.52 | 2.84 | 2.98 | 3.96 | + |
| Rtca | 77.06 | 77.7 | 74.13 | 38.24 | 42.66 | 47.5 | 46.27 | 51.02 | 49.63 | 40.64 | 46.7 | 40.29 | + |
| Dnaja2 | 173.33 | 184.74 | 168.19 | 118.67 | 119.82 | 120.32 | 124.58 | 136.67 | 130.9 | 112.51 | 123.78 | 120.62 | + |
| Lsm6 | 19.5 | 21.24 | 19.06 | 13.84 | 13.44 | 13.67 | 12.09 | 14.22 | 11.81 | 11.01 | 10.53 | 11.74 | + |
| Hnrnpa0 | 5.71 | 6.67 | 6.38 | 4.11 | 2.61 | 2.86 | 2.88 | 2.5 | 3.19 | 3.59 | 2.42 | 3.96 | + |
| Irgq | 0.95 | 0.75 | 0.77 | 1.95 | 1.75 | 1.8 | 1.78 | 1.25 | 1.59 | 2.12 | 2.11 | 1.71 | + |
| Vdac1 | 120 | 124.08 | 114.82 | 90.86 | 84.69 | 91.59 | 102.62 | 96.49 | 95.86 | 72.43 | 91.09 | 77.86 | + |
| Sipa1l2 | 0.13 | 0.22 | 0.16 | 0.13 | 0.18 | 0.15 | 0.45 | 0.32 | 0.35 | 0.67 | 0.45 | 0.87 | + |
| Ccdc109b | 17.43 | 16.63 | 17.76 | 31.71 | 34.57 | 35.67 | 33.92 | 35.2 | 28.29 | 30.29 | 35.14 | 30.35 | + |
| Hspd1 | 552.3 | 575.38 | 596.05 | 174.57 | 218.48 | 209.11 | 159.12 | 258.54 | 232.09 | 122.82 | 143.97 | 119.83 | + |
| Ppif | 72.97 | 78.17 | 73 | 31.09 | 31.43 | 30.7 | 26.52 | 35.91 | 31.21 | 22.59 | 22.26 | 21.65 | + |
| Ufl1 | 9.89 | 10.38 | 10.68 | 13.11 | 13.06 | 12.81 | 15.15 | 14.75 | 14.6 | 13.49 | 14.89 | 13.31 | + |
| Oxa1l | 27.53 | 26.34 | 26.2 | 38.46 | 35.84 | 34.5 | 39.48 | 34.4 | 34.51 | 38.22 | 42.37 | 36.83 | + |
| Snhg11 | 2.21 | 2.38 | 2.04 | 1.19 | 1.08 | 0.97 | 1.18 | 1.14 | 1.17 | 1.11 | 0.92 | 1.03 | + |
| Nup107 | 51.46 | 48.01 | 47 | 20.95 | 26.79 | 24.85 | 15.59 | 20.65 | 20.27 | 13.12 | 15.03 | 12.7 | + |
| Stag1 | 5.46 | 5.22 | 5.17 | 3.65 | 3.66 | 4.15 | 3.39 | 3.3 | 4.27 | 3.65 | 3.97 | 3.58 | + |
| Kars | 197.66 | 186.96 | 192.44 | 114.59 | 122.89 | 126.15 | 123.25 | 138.36 | 132.29 | 102.36 | 116.37 | 102.57 | + |
| Sema3a | 0.11 | 0.09 | 0.11 | 8.77 | 9.27 | 8.3 | 0.13 | 0.02 | 0.11 | 1.82 | 1.67 | 1.41 | + |
| Atp5c1 | 413.54 | 419.56 | 409.56 | 340.84 | 331.21 | 349.27 | 394.58 | 346.32 | 349.16 | 353.03 | 384.54 | 354.3 | + |
| Mtmr2 | 26.09 | 25.44 | 27.07 | 20.38 | 22.12 | 21.29 | 17 | 20.02 | 21 | 18.26 | 19.2 | 17.52 | + |
| Iws1 | 6.22 | 7.01 | 7.13 | 5.68 | 5.81 | 6.2 | 5.08 | 5.42 | 5.51 | 5 | 5.21 | 5.24 | + |
| Gm6793 | 267.55 | 296.44 | 283.44 | 181.14 | 192.09 | 179.07 | 138.07 | 157.65 | 152.57 | 125.88 | 125.14 | 124.11 | + |
| Mrpl12 | 147.22 | 166.02 | 160.56 | 70.59 | 77.1 | 85.07 | 78.72 | 87.04 | 87.08 | 60.89 | 58.57 | 66.39 | + |
| S1pr3 | 1.2 | 1.78 | 2.17 | 2.31 | 2.97 | 3.02 | 1.28 | 1.74 | 1.65 | 0.56 | 0.49 | 0.27 | + |
| Crat | 4.3 | 3.59 | 3.54 | 5.38 | 6.14 | 4.82 | 5.47 | 6.08 | 5.48 | 5.33 | 5.5 | 5.63 | + |
| Uba6 | 4.9 | 4.99 | 5.25 | 2.76 | 4.42 | 3.52 | 3.36 | 3.25 | 3.63 | 3.77 | 3.78 | 3.17 | + |
| Asb6 | 23.56 | 25.5 | 22.21 | 17.71 | 17.37 | 16.93 | 15.26 | 19.01 | 14.55 | 18.03 | 16.6 | 18.22 | + |
| Ptpn18 | 29.98 | 35.11 | 30.32 | 57.3 | 53.41 | 53.35 | 67.22 | 48.21 | 61.6 | 62.32 | 59.24 | 68.89 | + |
| D030028A08Rik | 4.8 | 4.75 | 6.49 | 2.2 | 2.55 | 1.97 | 1.65 | 3.04 | 2.34 | 2.23 | 1.49 | 1.66 | + |
| Txn2 | 175.44 | 153.96 | 176.23 | 114.4 | 127.24 | 124.49 | 116.94 | 148.24 | 139.35 | 111.36 | 115.81 | 114.81 | + |
| Fam98a | 19.4 | 20.62 | 18.86 | 9.96 | 11.41 | 12.02 | 8.79 | 12.76 | 10.34 | 9.2 | 8.44 | 10.29 | + |
| Cpsf6 | 33.34 | 35.59 | 35.29 | 18.39 | 20.22 | 19.79 | 17.15 | 18.96 | 18.89 | 15.58 | 15.94 | 15.69 | + |
| Tfip11 | 13.49 | 13.57 | 13.97 | 8.7 | 9.31 | 9.17 | 8.94 | 7.95 | 8.14 | 10 | 8.22 | 10.3 | + |
| D2Wsu81e | 29.62 | 30.89 | 32.33 | 21.16 | 20.2 | 17.85 | 16.42 | 18.68 | 19.43 | 16.44 | 14.24 | 13.61 | + |
| Rftn1 | 8.27 | 7.79 | 10.84 | 8.91 | 10.14 | 7.19 | 14.36 | 10.22 | 10.59 | 20.78 | 13.94 | 20.23 | + |
| Ifng | 1007.59 | 774.16 | 713.34 | 4.82 | 20.58 | 12.2 | 48.8 | 159.36 | 164.54 | 11.9 | 4.95 | 13.42 | + |
| Cops7a | 70.38 | 71.53 | 72.12 | 39.36 | 40.26 | 38.66 | 32.14 | 44.73 | 41.21 | 29.03 | 25.41 | 26.72 | + |
| Dcp1a | 7.36 | 6.75 | 8.09 | 5.56 | 4.9 | 5.44 | 6.26 | 5.23 | 5.53 | 5.44 | 5.66 | 4.52 | + |
| Gnb1 | 84.02 | 87.52 | 88.45 | 50.58 | 50.9 | 52.65 | 44.89 | 54.05 | 50.7 | 43.18 | 43.07 | 45.16 | + |
| Ndufs6 | 140 | 141.9 | 137.44 | 71.56 | 84.98 | 86.04 | 81.9 | 97.94 | 77.42 | 60.05 | 55.95 | 57.89 | + |
| Ino80e | 49.62 | 50.26 | 53.17 | 26.62 | 27.7 | 27.81 | 18.12 | 27.39 | 24.99 | 20.31 | 21.86 | 20.56 | + |
| Psme3 | 17.89 | 18.9 | 17.73 | 8.06 | 10.13 | 8.84 | 6.64 | 9.11 | 9.1 | 7.89 | 6.19 | 6.12 | + |
| Prpf4b | 15.09 | 14.19 | 15.65 | 10.93 | 11.95 | 10.43 | 9.27 | 12.21 | 11.51 | 12.02 | 12.24 | 11.74 | + |
| Tagap | 76.6 | 66.14 | 80.13 | 36.51 | 39.1 | 43.22 | 55.05 | 72.82 | 64.34 | 45.25 | 43.9 | 44.35 | + |
| Mis12 | 18.7 | 18.87 | 17.98 | 13.58 | 14.67 | 13.25 | 9.7 | 11.08 | 9.41 | 10.53 | 8.82 | 10.71 | + |
| Armcx4 | 0.57 | 0.74 | 0.61 | 0.54 | 0.47 | 0.52 | 0.46 | 0.46 | 0.51 | 0.2 | 0.26 | 0.21 | + |
| Gpr125 | 0.17 | 0.09 | 0.11 | 0.59 | 0.73 | 0.73 | 0.36 | 0.16 | 0.49 | 0.5 | 0.45 | 0.71 | + |
| Trib3 | 22.35 | 25.73 | 23.15 | 28.26 | 37.22 | 36.66 | 33.43 | 52.03 | 53.36 | 14.31 | 14.6 | 15.91 | + |
| Pcbp1 | 62.57 | 65.93 | 68.49 | 40.9 | 40.74 | 40.48 | 36.55 | 44.38 | 33.37 | 37.09 | 38.59 | 37.95 | + |
| Thoc3 | 30.49 | 30.87 | 31.23 | 14.18 | 16.37 | 15.5 | 13.24 | 15.6 | 14.43 | 13.15 | 12.85 | 13.13 | + |
| Pycr1 | 10.19 | 9.73 | 10.07 | 9.36 | 12.57 | 14.03 | 11.79 | 19.53 | 15.71 | 6.34 | 7.82 | 6.47 | + |
| Polr2a | 18.9 | 18.8 | 20.25 | 12.9 | 13.24 | 11.81 | 12.65 | 12.98 | 13.89 | 11.98 | 12.35 | 11.47 | + |
| Rprd1b | 16.71 | 15.17 | 14 | 9.16 | 10.57 | 10.89 | 11.42 | 12.99 | 10.33 | 10.73 | 10.62 | 10.84 | + |
| L3mbtl2 | 24.08 | 22.9 | 21.94 | 10.5 | 12.65 | 12.66 | 11.34 | 14.14 | 11.89 | 12.27 | 11.63 | 9.5 | + |
| Strap | 151.76 | 158.91 | 153.57 | 79.57 | 82.45 | 80.61 | 70.87 | 95.1 | 88.76 | 66.68 | 72.05 | 71.4 | + |
| Crbn | 14.5 | 16.2 | 13.92 | 30.37 | 24.81 | 29.1 | 33.63 | 24.93 | 26.14 | 36.94 | 39.92 | 39.01 | + |
| Usp11 | 7.56 | 7.27 | 8.83 | 16.48 | 19.07 | 18.47 | 15.26 | 17.6 | 16.33 | 14.73 | 16.82 | 17.27 | + |
| 4-Sep | 0.66 | 0.25 | 0.17 | 1.82 | 1.64 | 2.26 | 2.32 | 1.26 | 1.74 | 1.71 | 2.03 | 2.64 | + |
| Haus7 | 22.31 | 25.37 | 24.26 | 16.51 | 14.24 | 14.17 | 12.59 | 13.22 | 13.2 | 10.6 | 10.86 | 10.29 | + |
| Nutf2 | 5.42 | 5.87 | 5.51 | 3.37 | 2.4 | 3.34 | 3.23 | 4.32 | 3.24 | 2.69 | 2.58 | 2.97 | + |
| Cox10 | 15.42 | 15.22 | 15.99 | 8.03 | 8.83 | 8.19 | 6.55 | 9.09 | 8.91 | 5.86 | 6.9 | 5.82 | + |
| Pfdn2 | 78.58 | 85.31 | 88.92 | 41.56 | 49.48 | 45.83 | 39.82 | 51.68 | 51.84 | 38.35 | 36.46 | 35.03 | + |
| Synpo | 0.17 | 0.34 | 0.19 | 0.8 | 0.91 | 0.88 | 0.41 | 0.67 | 0.78 | 0.66 | 0.51 | 0.79 | + |
| Rnh1 | 236.06 | 212.64 | 234.19 | 329.33 | 338.37 | 341 | 355.92 | 377.88 | 359.63 | 413.38 | 417.62 | 426.61 | + |
| Mrps31 | 18.88 | 20.17 | 22.49 | 13.62 | 16.95 | 14.99 | 11.22 | 16.36 | 12.51 | 11.28 | 14.2 | 14.3 | + |
| Ap2m1 | 169.84 | 164.1 | 167.94 | 109.63 | 122.65 | 117.1 | 96.35 | 127.52 | 107.25 | 82.1 | 91.99 | 86.19 | + |
| Supt16 | 40.16 | 42.29 | 40.87 | 23.37 | 24.31 | 26.19 | 18.79 | 22.73 | 22.31 | 18.83 | 21.31 | 18.99 | + |
| Pes1 | 73.39 | 76.06 | 70.6 | 39.95 | 43.32 | 41.81 | 38.77 | 48.14 | 42.89 | 34.71 | 36.06 | 36.54 | + |
| Il4ra | 62.92 | 60.9 | 63.74 | 98.02 | 90.66 | 85.94 | 73.64 | 75.13 | 76.2 | 72.29 | 68.16 | 74.92 | + |
| Skiv2l | 21.13 | 21.49 | 22.85 | 14.39 | 15.18 | 15.64 | 11.94 | 12.86 | 11.81 | 12.37 | 11.05 | 14.78 | + |
| Klf9 | 0.45 | 0.52 | 0.41 | 2.56 | 2.39 | 1.86 | 0.86 | 0.74 | 0.69 | 2.02 | 2.3 | 2.12 | + |
| Ppib | 321.19 | 331.01 | 309.52 | 228.92 | 211.15 | 221.61 | 230.65 | 222.31 | 225.45 | 204.22 | 204.03 | 214.74 | + |
| Airn | 2.7 | 2.53 | 1.71 | 2.77 | 4.12 | 3.12 | 3.16 | 2.1 | 2.28 | 5.25 | 5.71 | 5.46 | + |
| Nat10 | 21.42 | 21.41 | 22.35 | 14.6 | 14.52 | 15.46 | 14.01 | 17.67 | 17.43 | 17.07 | 16.42 | 14.68 | + |
| Gzmb | 7872.71 | 8076.32 | 7098.8 | 2856.59 | 3082.15 | 2987.16 | 4122.96 | 4644.32 | 4348.33 | 3341.06 | 2857.62 | 3551.25 | + |
| Idua | 4.98 | 6.17 | 5.38 | 12.48 | 11.9 | 11.51 | 11.43 | 9.12 | 10.54 | 12 | 11.98 | 11.48 | + |
| Gle1 | 29.35 | 28.91 | 31.81 | 19.83 | 19.74 | 19.35 | 16.03 | 17.27 | 18.16 | 15.76 | 14.48 | 13.05 | + |
| Prmt1 | 72.58 | 80.79 | 84.96 | 30.12 | 33.94 | 28.64 | 24.15 | 37.02 | 31.57 | 17.4 | 18.49 | 16.28 | + |
| Grwd1 | 54.37 | 49.81 | 52.59 | 16.35 | 17.56 | 19.98 | 12.02 | 14.3 | 15.18 | 8.81 | 7.91 | 9.52 | + |
| Hmgn5 | 15.74 | 17.73 | 15.62 | 11.77 | 13.94 | 11.44 | 11.48 | 11.77 | 12.8 | 11.88 | 11.21 | 11.12 | + |
| Clasp1 | 4.75 | 4.93 | 5.01 | 3.57 | 3.61 | 3.91 | 3.03 | 3.61 | 4.03 | 3.23 | 3.44 | 4.06 | + |
| Mapk3 | 7.35 | 5.85 | 6.16 | 9.67 | 10.29 | 11.16 | 11.16 | 11.17 | 10.78 | 11.17 | 12.83 | 12.39 | + |
| Hsp90b1 | 479.6 | 485 | 470.94 | 214.35 | 233.44 | 225.06 | 235.77 | 283.6 | 268.83 | 188.21 | 214.83 | 186.41 | + |
| Paxip1 | 4.85 | 6.18 | 5.3 | 3.18 | 4 | 3.39 | 2.18 | 2.93 | 2.61 | 3.48 | 2.61 | 2.35 | + |
| Nnt | 7.37 | 8.38 | 6.88 | 3.44 | 4.95 | 4.05 | 3.99 | 4.23 | 3.79 | 3.29 | 4.13 | 3.5 | + |
| Emc9 | 9.86 | 9.25 | 8.4 | 18.06 | 13.88 | 17.27 | 17.13 | 14.08 | 13.33 | 17.2 | 19.6 | 15.75 | + |
| Phf5a | 57.91 | 61.35 | 58.88 | 32.56 | 32.35 | 33.86 | 33.34 | 35.57 | 39.62 | 32.13 | 34.59 | 30.99 | + |
| Ddx5 | 345.73 | 352.27 | 349.25 | 293.34 | 297.96 | 291.03 | 255.88 | 284.51 | 272.83 | 291.46 | 288.09 | 297.46 | + |
| Tnfrsf11b | 2.82 | 4.13 | 3.28 | 10.23 | 7.97 | 9.01 | 1.39 | 1.44 | 2.7 | 0.49 | 0.74 | 0.66 | + |
| Rplp0 | 2738.31 | 2657.94 | 2725.53 | 3997 | 3890.73 | 4067.24 | 3815.12 | 3632.91 | 3618.97 | 4466.87 | 4701.83 | 4488.37 | + |
| Calcrl | 3.8 | 2.94 | 3.72 | 3.15 | 3.59 | 3.18 | 3.42 | 4.2 | 3.71 | 5.72 | 4.44 | 4.63 | + |
| 2210404O07Rik | 2.78 | 3.04 | 2.54 | 8.57 | 6.73 | 9.32 | 19.09 | 12.16 | 13.9 | 16.81 | 19.51 | 22.76 | + |
| Rabgef1 | 16.31 | 13.26 | 14.74 | 8.67 | 9.91 | 9.83 | 7.35 | 11.21 | 9.45 | 8.3 | 9.02 | 8.43 | + |
| Rpl39 | 1906.47 | 1937.74 | 1934.45 | 3211.25 | 3311.15 | 3444.47 | 3022.97 | 2809.25 | 2749.62 | 3369.75 | 3739.6 | 3451.41 | + |
| Mrps7 | 68.69 | 67.63 | 64.93 | 36.07 | 35.27 | 33.52 | 36.68 | 39.02 | 38.45 | 29.6 | 27.23 | 36.67 | + |
| Vim | 1068.82 | 1107.35 | 1081.18 | 2366.77 | 2147.57 | 2323.87 | 2376.65 | 1665.52 | 2027.98 | 1825.19 | 2067.61 | 1807.63 | + |
| Casc5 | 3.53 | 2.97 | 2.35 | 1.97 | 2.74 | 2.01 | 1 | 1.07 | 1.08 | 1.54 | 1.43 | 1.19 | + |
| Flot2 | 20.71 | 20.6 | 21.74 | 31.94 | 32.12 | 32.1 | 27.63 | 27.5 | 30.18 | 27.53 | 29.12 | 30.31 | + |
| Fignl1 | 38.93 | 37.94 | 37.33 | 20.05 | 23.79 | 21.66 | 13.72 | 14.29 | 14.03 | 11.29 | 11.01 | 10.22 | + |
| Tgfb3 | 0.1 | 0.12 | 0.03 | 8.53 | 5.51 | 5.9 | 3.6 | 1.14 | 2.06 | 12.08 | 11.67 | 16.33 | + |
| 1110038B12Rik | 82.8 | 78.32 | 81.44 | 192.07 | 188.39 | 190.74 | 158.27 | 172.33 | 141.38 | 215.38 | 248.35 | 195.86 | + |
| Arhgap27 | 9.47 | 9.73 | 7.75 | 21.28 | 17.91 | 17.41 | 18.1 | 17.16 | 16.25 | 23.99 | 20.54 | 26.34 | + |
| Hyou1 | 13.49 | 12.41 | 15.26 | 7.16 | 7.52 | 6.86 | 6.09 | 8.88 | 8.73 | 5.92 | 6.83 | 5.67 | + |
| Psmb5 | 318.94 | 321.89 | 312.26 | 158.63 | 172.98 | 172.87 | 145.02 | 187.76 | 168.73 | 131.05 | 126.3 | 140.57 | + |
| Gps2 | 15.25 | 14.4 | 14.74 | 15.75 | 16.89 | 15.21 | 15.21 | 15.17 | 14.25 | 21.34 | 18.13 | 21.91 | + |
| Arhgef7 | 7.63 | 6.78 | 6.93 | 5.33 | 4.93 | 4.78 | 5.47 | 5.89 | 4.69 | 5.06 | 4.77 | 5.96 | + |
| 2310008H09Rik | 16.34 | 16.1 | 16.24 | 5.9 | 6.46 | 5.63 | 4.88 | 7.23 | 5.86 | 4.86 | 3.57 | 4.86 | + |
| Lsm2 | 90.99 | 86.78 | 87.42 | 41.53 | 47.67 | 51.29 | 37.76 | 42.98 | 41.86 | 29.82 | 32.1 | 29.18 | + |
| Arfip1 | 17.95 | 20.07 | 19.37 | 15.45 | 14.41 | 17.1 | 16.14 | 17.36 | 15.88 | 14.12 | 15.44 | 13.56 | + |
| Ebna1bp2 | 54.61 | 59.37 | 59.32 | 18.55 | 21.07 | 19.17 | 16.83 | 26.94 | 23.51 | 11.77 | 12.62 | 13.98 | + |
| Xbp1 | 83.2 | 98.05 | 105.51 | 124.86 | 130.42 | 122.86 | 85.53 | 105.99 | 105.02 | 76.5 | 81.8 | 77.37 | + |
| Sertad3 | 6.54 | 6.8 | 6.71 | 13.88 | 14.9 | 16.49 | 15.59 | 11.4 | 12.29 | 17.66 | 17.99 | 20.04 | + |
| Dpp9 | 25.25 | 27.24 | 25.59 | 16.95 | 15.46 | 17.54 | 11.64 | 15.52 | 14.98 | 13.16 | 12.89 | 12.23 | + |
| Birc2 | 13.77 | 13.23 | 13.44 | 27.13 | 25.36 | 26.65 | 35.54 | 28.93 | 29.96 | 35.08 | 35.75 | 36.29 | + |
| Snrnp25 | 52.35 | 58.75 | 61.37 | 34.03 | 39.99 | 33.8 | 40.27 | 39.96 | 36.76 | 31.83 | 43.48 | 31.8 | + |
| Ssb | 140.01 | 146.2 | 139.75 | 83.64 | 89.23 | 90.45 | 81.06 | 86 | 84.24 | 77.45 | 71.79 | 78.81 | + |
| Snhg7 | 9.39 | 10.02 | 9.91 | 18.83 | 19.35 | 15.38 | 14.95 | 18.84 | 19.45 | 23.21 | 21.45 | 22.35 | + |
| Ube2v2 | 26.56 | 26.67 | 28.06 | 12.67 | 13.81 | 12.44 | 9.31 | 12.2 | 13.49 | 8.95 | 8.68 | 8.94 | + |
| Arhgef4 | 3.13 | 2.62 | 3.53 | 8.06 | 6.97 | 6.81 | 8.44 | 6.73 | 7.33 | 8.4 | 9.46 | 10.79 | + |
| Tmem70 | 84.95 | 84.69 | 83.51 | 52 | 51.24 | 54.92 | 41.74 | 55.54 | 53.5 | 38.5 | 45.32 | 39.24 | + |
| Ppp4r1l-ps | 0.13 | 0.25 | 0.28 | 0.52 | 0.81 | 0.67 | 0.98 | 0.62 | 0.72 | 1.02 | 1.07 | 0.84 | + |
| Zbed6 | 2.42 | 2.65 | 2.66 | 5.91 | 4.61 | 4.26 | 4.2 | 4.55 | 4.24 | 5.81 | 5.85 | 5.29 | + |
| Atp2b4 | 3.33 | 2.91 | 2.61 | 10.11 | 9.76 | 9.91 | 6.82 | 4.68 | 4.98 | 8.45 | 8.09 | 8.38 | + |
| Zfp583 | 1.59 | 2.06 | 2.05 | 3.15 | 3.05 | 3.55 | 3.55 | 2.87 | 3.96 | 3.17 | 3.1 | 2.89 | + |
| 1700084E18Rik | 0.53 | 0.27 | 0.25 | 2.62 | 3.74 | 1.91 | 2.38 | 1.08 | 1.98 | 3.03 | 2.41 | 1.99 | + |
| Hif1an | 3.38 | 3.59 | 3.35 | 3.75 | 4.35 | 4.17 | 3.57 | 3.62 | 3.69 | 4.55 | 5.07 | 4.32 | + |
| Sumo2 | 601.85 | 609.89 | 599.3 | 424.48 | 434.99 | 435.69 | 407.54 | 426.1 | 412.86 | 393.56 | 429.17 | 390.04 | + |
| Vstm5 | 0.89 | 0.79 | 0.65 | 1.75 | 2.01 | 1.57 | 2.19 | 1.51 | 1.92 | 1.73 | 2.08 | 1.42 | + |
| Lrrc61 | 4.67 | 3.72 | 5.12 | 8.32 | 6.73 | 8.75 | 12.18 | 9.39 | 8.44 | 9.55 | 10.14 | 10.97 | + |
| Ccnd3 | 329.83 | 358.01 | 351.93 | 182.14 | 176.74 | 159.45 | 433.2 | 367.54 | 445.22 | 142.55 | 144.16 | 150.21 | + |
| Zfp292 | 1.42 | 1.42 | 1.3 | 3.15 | 3.03 | 2.83 | 2.76 | 2.94 | 2.9 | 3.7 | 4.07 | 4.37 | + |
| U2af1 | 221.25 | 232.41 | 225.99 | 123.2 | 128.63 | 134.28 | 114.6 | 145.99 | 130.31 | 92.23 | 96.18 | 91.9 | + |
| Azin1 | 26.8 | 25.58 | 26.14 | 16.43 | 15.18 | 14.65 | 14.73 | 16.26 | 14.27 | 15.17 | 17.18 | 13.65 | + |
| Golga4 | 2.82 | 2.27 | 2.37 | 2.52 | 2.69 | 2.74 | 3.65 | 3.06 | 3.17 | 3.82 | 4.28 | 3.98 | + |
| Mettl10 | 33.82 | 31.06 | 34.21 | 23.84 | 25.61 | 25.76 | 20.7 | 25.63 | 24.27 | 18.75 | 21.52 | 21.1 | + |
| Cers2 | 93.31 | 91.22 | 92.07 | 76.94 | 75.12 | 72.97 | 73.38 | 78.51 | 73.57 | 72.57 | 76.91 | 80.04 | + |

| | | | | | | | | | | | | | |
|---|---|---|---|---|---|---|---|---|---|---|---|---|---|
| Rbm15b | 2.9 | 3.57 | 3.23 | 2.49 | 2.46 | 1.88 | 2.17 | 1.84 | 2.03 | 1.78 | 1.92 | 2.41 | + |
| Psap | 35.54 | 30.21 | 26.5 | 69.71 | 61.92 | 60.81 | 54.3 | 48.97 | 48.56 | 53.15 | 67.9 | 55.37 | + |
| Prr13 | 107.89 | 102.91 | 105.58 | 123.26 | 114.43 | 127.38 | 153.49 | 144.41 | 153.18 | 134.37 | 127.31 | 140.82 | + |
| Tra2b | 175.51 | 198.93 | 188.81 | 75.6 | 83.93 | 78.8 | 60.38 | 77.24 | 71.28 | 56.54 | 50.29 | 48.4 | + |
| Bcl3 | 4.32 | 4.06 | 4.01 | 7.89 | 8.47 | 8.69 | 8.6 | 7.84 | 7.66 | 8.88 | 8.9 | 9.9 | + |
| Trdmt1 | 3.42 | 3.96 | 3.92 | 1.49 | 1.3 | 1.81 | 1.61 | 1.9 | 1.63 | 1.26 | 0.91 | 1.01 | + |
| Hist1h2bj | 5.05 | 2.6 | 3.59 | 0.6 | 1.52 | 0.82 | 0.36 | 0.82 | 1.1 | 1.38 | 0.17 | 0.64 | + |
| Zfp446 | 3.68 | 3.73 | 4.01 | 2.1 | 2.31 | 2.68 | 1.85 | 1.84 | 1.37 | 1.32 | 1.41 | 1.84 | + |
| Tmem8b | 0.11 | 0.23 | 0.07 | 0.45 | 0.42 | 0.46 | 0.77 | 0.51 | 0.41 | 0.97 | 0.54 | 0.63 | + |
| Gltp | 40.55 | 46.76 | 42.25 | 62.18 | 57.11 | 57.31 | 81.55 | 72.87 | 73.65 | 74.45 | 76.68 | 66.73 | + |
| Fkbp3 | 273.22 | 285.83 | 268.75 | 154.43 | 169.67 | 177.17 | 176.54 | 199.43 | 178.4 | 134.52 | 138.75 | 139.56 | + |
| Rpl18 | 1145.86 | 1190.55 | 1178.68 | 1563.35 | 1537.61 | 1594.58 | 1474.51 | 1430.89 | 1474.98 | 1554.08 | 1636.47 | 1577.08 | + |
| Rfc5 | 64.55 | 65.92 | 64.32 | 25.28 | 26.81 | 30.29 | 23.32 | 24.72 | 20.93 | 17.72 | 14.43 | 15.93 | + |
| Slc25a44 | 6.34 | 5.89 | 4.78 | 3.68 | 3.5 | 3.72 | 3.45 | 3.31 | 4.15 | 3.61 | 3.94 | 4.06 | + |
| Gm9855 | 31.06 | 37.24 | 35.55 | 13.41 | 13.12 | 13.14 | 11.17 | 15.63 | 16.97 | 11.27 | 11.07 | 10.79 | + |
| Pafah1b2 | 54.17 | 51.25 | 54.85 | 30.81 | 34.27 | 32.61 | 27.07 | 30.8 | 32.06 | 25.05 | 27.89 | 23.03 | + |
| B4galt7 | 12.27 | 11.03 | 13.01 | 25.03 | 17.44 | 23.97 | 21.53 | 19.95 | 24.34 | 23.13 | 24.13 | 25.35 | + |
| Mboat2 | 0.44 | 0.28 | 0.64 | 2.95 | 2.79 | 2.09 | 4.57 | 2.41 | 2.74 | 3.41 | 4.26 | 4.63 | + |
| Deb1 | 53.11 | 49.59 | 50.21 | 29.71 | 35.86 | 39.93 | 36.23 | 37.49 | 33.21 | 31.19 | 27.05 | 29.1 | + |
| Ppp2r2a | 41.47 | 42.55 | 42.75 | 36.93 | 37.85 | 35.73 | 32.93 | 35.62 | 32.62 | 30.62 | 34.97 | 33.09 | + |
| 9330151L19Rik | 0.45 | 0.9 | 0.76 | 1.82 | 2.35 | 1.44 | 2.68 | 2.54 | 3.03 | 2.43 | 3.08 | 2.09 | + |
| Stac2 | 0.07 | 0.06 | 0.04 | 2.22 | 1.54 | 1.8 | 5.32 | 2.39 | 3.24 | 3.75 | 4.65 | 4.9 | + |
| Xrcc2 | 6.33 | 5.94 | 5.59 | 2.95 | 4.1 | 4.08 | 2.62 | 2.55 | 2.5 | 2.45 | 2.97 | 2.51 | + |
| Uckl1 | 23.57 | 22.52 | 23.04 | 33.12 | 34.02 | 31.59 | 37.07 | 37.89 | 34.74 | 42.28 | 35.77 | 39.37 | + |
| Dclre1c | 17.15 | 19.05 | 18.84 | 8.32 | 10.6 | 10.36 | 9.42 | 9.88 | 10.53 | 10.21 | 7.49 | 8.97 | + |
| Atp6v1c1 | 44.55 | 44.18 | 43.49 | 24.31 | 23.48 | 27.96 | 27.42 | 30.35 | 30.12 | 23.34 | 27.91 | 24.95 | + |
| Plekha7 | 0.21 | 0.39 | 0.18 | 1.13 | 1.01 | 0.9 | 1.09 | 0.84 | 0.81 | 1.51 | 1.3 | 1.02 | + |
| Wdr18 | 45.65 | 48.38 | 51.42 | 17.44 | 20.35 | 20.64 | 18.75 | 23.74 | 20.87 | 10.61 | 13.08 | 13.05 | + |
| Bzw1 | 170.14 | 182.41 | 172.34 | 77.6 | 84.02 | 79.1 | 62.5 | 75.67 | 75.55 | 78.99 | 76.88 | 74.6 | + |
| 5330426P16Rik | 8.68 | 9.07 | 8.55 | 15.94 | 13.83 | 13.89 | 12.85 | 12.59 | 13.8 | 12.04 | 11.82 | 13.21 | + |
| Mrpl15 | 43.07 | 45.54 | 46.8 | 30.34 | 32.59 | 32.39 | 28.44 | 33.05 | 32.6 | 25.94 | 26.87 | 26.66 | + |
| Akap9 | 2.53 | 2.67 | 2.6 | 4.81 | 4.6 | 5 | 5.22 | 4.98 | 5.24 | 6.01 | 6.62 | 5.47 | + |
| Ccl4 | 542.03 | 585.58 | 491.44 | 56.03 | 118.65 | 100.01 | 80.92 | 216.52 | 228.17 | 142.05 | 83.02 | 114.08 | + |
| Bcas2 | 129.8 | 137.2 | 132.63 | 94.59 | 93.77 | 93.47 | 99.19 | 105.48 | 96.33 | 91.96 | 99.11 | 94.39 | + |
| Zscan12 | 3.67 | 3.44 | 4.25 | 7.77 | 7.52 | 8.27 | 3.95 | 5.6 | 5.26 | 4.8 | 6.07 | 5.13 | + |
| Tpm4 | 228.43 | 226.71 | 223.57 | 192.45 | 187.38 | 184.74 | 189.65 | 181.46 | 187.76 | 183.98 | 159.22 | 192.06 | + |
| H2-Q4 | 15.37 | 15.46 | 16.44 | 34.3 | 34.78 | 32.87 | 35.6 | 29.9 | 30.83 | 40.62 | 47.85 | 44.52 | + |
| Kctd20 | 15.85 | 17.15 | 13.29 | 11.72 | 9.65 | 10.68 | 10.36 | 9.64 | 9.86 | 8.58 | 7.6 | 8.86 | + |
| Adrm1 | 97.84 | 106.35 | 102.46 | 59.89 | 65.09 | 58.77 | 50.03 | 62.76 | 57.07 | 53.24 | 56.41 | 53.69 | + |
| Pcca | 19.81 | 21.84 | 20.52 | 16.12 | 19.28 | 16 | 14.87 | 20.15 | 17.42 | 9.12 | 10.08 | 9.65 | + |
| Frrs1 | 14.62 | 15.41 | 15.89 | 10.82 | 9.82 | 11.25 | 10.74 | 13.19 | 11.05 | 10.69 | 9.91 | 10.19 | + |
| Isg20l2 | 8.32 | 9.67 | 10.47 | 4.63 | 5.29 | 4.19 | 4.05 | 4.16 | 5.12 | 5.64 | 4.25 | 4.44 | + |
| 1600014C10Rik | 15.11 | 15.23 | 14.72 | 34.3 | 29.73 | 31.78 | 34.4 | 24.54 | 27.78 | 40.69 | 33.45 | 40.18 | + |
| Fam214a | 2.18 | 2.11 | 2.35 | 5.28 | 3.39 | 3.86 | 3.97 | 4.14 | 4.13 | 4.07 | 4.55 | 5.56 | + |
| Slc35b4 | 15.72 | 15.64 | 14.48 | 11.17 | 10.94 | 11.68 | 10.49 | 10.44 | 11.26 | 9.87 | 10.34 | 9.71 | + |
| 5430416N02Rik | 14.14 | 16.82 | 18.31 | 46.57 | 42.88 | 46.43 | 40.21 | 52.61 | 46.71 | 54.19 | 73.44 | 43.65 | + |
| Ampd2 | 35.16 | 35.93 | 35.43 | 22.08 | 21.22 | 20.73 | 19.22 | 24.93 | 24.91 | 16.16 | 17.32 | 16.96 | + |
| Gm20300 | 4.61 | 4.85 | 5.25 | 8.22 | 8.21 | 8.22 | 8.33 | 6.99 | 6.99 | 9.34 | 9.11 | 9.83 | + |
| Mvb12a | 97.94 | 91.99 | 88.08 | 101.77 | 101.75 | 105.5 | 125.32 | 125.14 | 122.3 | 108.38 | 112.76 | 114.3 | + |
| Slc25a36 | 7.8 | 8.5 | 9.31 | 13.82 | 15.13 | 14.17 | 16.13 | 13.01 | 16.76 | 17.76 | 16.65 | 16.98 | + |
| Cox6b2 | 18.33 | 10.75 | 18.37 | 26.54 | 33.28 | 38.44 | 32.82 | 30.8 | 25.83 | 25.41 | 33.59 | 30.02 | + |
| Dcaf13 | 106.63 | 105.86 | 104.42 | 55.53 | 57.79 | 55.69 | 53.25 | 69.48 | 65.25 | 46.44 | 51.63 | 56.97 | + |
| Gm17644 | 10.47 | 9.92 | 10.2 | 17.48 | 17.27 | 15.63 | 15.28 | 16.26 | 13.56 | 19.92 | 16.91 | 18.04 | + |
| Sptlc1 | 21.32 | 19.85 | 17.73 | 14.2 | 15.63 | 13.43 | 14.11 | 13.31 | 14.41 | 16.11 | 12.99 | 16.54 | + |
| Mroh1 | 4.49 | 4.62 | 4.53 | 9.03 | 8.38 | 8.82 | 8.11 | 8.38 | 7.27 | 9.5 | 8.16 | 8.27 | + |
| Zgpat | 15.42 | 16.95 | 15.01 | 18.92 | 15.9 | 14.57 | 17.56 | 16.33 | 19.43 | 25.32 | 20.66 | 25.02 | + |
| Rqcd1 | 56.94 | 58.89 | 57.12 | 33.77 | 37.59 | 34.85 | 34.27 | 36.39 | 35.28 | 30.34 | 32.36 | 33.93 | + |
| Gimap3 | 108.61 | 107.61 | 92.49 | 355.26 | 296.58 | 308.35 | 254.79 | 233.36 | 252.12 | 326.62 | 279.34 | 330.62 | + |
| Rbx1 | 108.55 | 109.38 | 108.73 | 83.29 | 84.87 | 95.21 | 94.58 | 95.45 | 96.43 | 80.79 | 86.65 | 86.65 | + |
| Brix1 | 62.77 | 64.55 | 63.94 | 29.72 | 31.47 | 29.25 | 24.86 | 35.55 | 32.13 | 22.45 | 22.55 | 21.46 | + |
| Taf9 | 229.99 | 220.98 | 217.73 | 109.57 | 121.87 | 122.86 | 98.46 | 127.61 | 115.25 | 96.63 | 104.62 | 95.03 | + |
| Cyb561d2 | 19.73 | 18.78 | 18.63 | 11.99 | 11.59 | 11.63 | 12.64 | 12.33 | 13.19 | 8.95 | 8.37 | 9.68 | + |
| Prrg4 | 9.97 | 9.53 | 8.99 | 4.44 | 6.04 | 6.39 | 5.14 | 7.89 | 7.64 | 5.37 | 6.93 | 6.12 | + |
| Polr3f | 6.3 | 7.45 | 6.6 | 4.62 | 3.46 | 3.61 | 2.87 | 3.63 | 3.77 | 4.49 | 3.43 | 3 | + |
| Gpd1l | 8.02 | 8.76 | 8.21 | 5.21 | 5.44 | 4.99 | 4.64 | 5.12 | 4.96 | 4.62 | 5.09 | 5.17 | + |
| Mcm4 | 87.14 | 91.44 | 84.13 | 46.09 | 50.89 | 48.5 | 30.89 | 34.96 | 34.29 | 29 | 28.4 | 28.95 | + |
| Glipr2 | 187 | 184.27 | 183.43 | 293.35 | 293.78 | 305.02 | 249.61 | 241.32 | 265.3 | 227.96 | 240.72 | 237.17 | + |
| Eif4h | 232.5 | 255.04 | 252.19 | 177.12 | 176.76 | 178.88 | 191.08 | 188.93 | 175.13 | 176.12 | 186.54 | 179.2 | + |
| Trip6 | 0.97 | 0.69 | 0.92 | 0.47 | 0.67 | 0.76 | 0.95 | 0.97 | 0.7 | 2.01 | 1.75 | 2 | + |
| Siva1 | 115.71 | 121.48 | 123.62 | 64.58 | 68 | 68.95 | 66.66 | 64.87 | 65.23 | 51.25 | 56.16 | 59.42 | + |
| Klhdc4 | 40.84 | 39.42 | 39.22 | 19.78 | 19.24 | 20.7 | 16.66 | 20.07 | 18 | 16.18 | 15.46 | 13.31 | + |
| Gmeb1 | 6.67 | 7.89 | 6.94 | 4.31 | 5.38 | 4.94 | 5.04 | 5.16 | 5.72 | 4.85 | 4.43 | 5.3 | + |
| Gpr160 | 4.38 | 4.55 | 5.13 | 4.43 | 3.29 | 4.25 | 17.33 | 15.08 | 13.6 | 15.17 | 17.28 | 16.37 | + |
| Bckdha | 10.05 | 7.91 | 9.82 | 20.28 | 17.15 | 21.19 | 22.41 | 23.92 | 20.91 | 26.84 | 24.19 | 26.17 | + |
| Faim | 41.7 | 46.96 | 37.26 | 28.02 | 27.38 | 26.16 | 28.7 | 34.37 | 30.7 | 25.88 | 25.27 | 31.17 | + |
| Rps24 | 1145.96 | 1149.35 | 1158.88 | 1756.2 | 1715.85 | 1837.61 | 1723.42 | 1649.89 | 1660.97 | 1939.22 | 2096.33 | 1958.42 | + |
| Sdhaf1 | 22.97 | 26.6 | 27.71 | 44.97 | 46.61 | 43.39 | 29.59 | 36.81 | 36.72 | 42.48 | 47.93 | 36.36 | + |
| Mrpl20 | 209.38 | 225.64 | 226.45 | 85.62 | 87.31 | 86.47 | 82.09 | 102.01 | 91.08 | 79.81 | 69.41 | 76.36 | + |
| Hnrnpr | 45.83 | 44.81 | 42.56 | 32.75 | 36.4 | 33.87 | 28.85 | 31.59 | 30.15 | 28.49 | 30.04 | 29.48 | + |
| Zbtb4 | 0.34 | 0.32 | 0.28 | 1.33 | 1.01 | 1.03 | 1.01 | 0.62 | 0.73 | 1.02 | 0.98 | 1.03 | + |
| Serpine2 | 148.88 | 116.7 | 128.62 | 42.2 | 52.89 | 58.74 | 66.76 | 90.51 | 72.89 | 56.64 | 62.76 | 63.95 | + |
| Tnfrsf21 | 2.9 | 2.89 | 2.34 | 0.8 | 0.9 | 0.93 | 1.62 | 1.98 | 1.95 | 0.96 | 1.16 | 1.02 | + |
| Slmo2 | 128 | 137.56 | 148.75 | 71.15 | 73.72 | 75.69 | 65.14 | 82.92 | 79.82 | 63.3 | 55.2 | 50.68 | + |
| Exoc3l | 3.22 | 3.59 | 3.2 | 0.73 | 0.92 | 1.07 | 1.27 | 0.87 | 0.97 | 1.03 | 1.01 | 1.22 | + |
| Armc6 | 18.09 | 16.85 | 17.45 | 6.8 | 7.69 | 8.82 | 6.59 | 10.87 | 8.89 | 4.68 | 5.6 | 5.85 | + |
| Dtx3 | 6.67 | 6.5 | 5.4 | 19.15 | 18.27 | 16.44 | 22.58 | 14.32 | 15.16 | 24.25 | 21.82 | 24.28 | + |
| Tfdp1 | 79.35 | 78.2 | 78.71 | 36.07 | 45.38 | 42.88 | 35.93 | 42.74 | 40.26 | 25.81 | 32.88 | 26.96 | + |
| Rpa2 | 61.86 | 66.09 | 58.76 | 36.89 | 38.11 | 39.93 | 30.93 | 28.87 | 30.4 | 30.46 | 28.08 | 30.72 | + |
| Tmem50b | 10.64 | 12.33 | 11.56 | 25.17 | 23.91 | 25.02 | 15.85 | 14.91 | 17.12 | 14.8 | 20.13 | 18.47 | + |
| Nop58 | 180.72 | 195.84 | 187.78 | 84.34 | 108.35 | 101.76 | 55.71 | 89.43 | 81.04 | 57.56 | 51.77 | 55.44 | + |
| Brms1 | 34.64 | 37.27 | 29.7 | 23 | 26.69 | 22.49 | 20.54 | 20.63 | 20.73 | 21.85 | 21.42 | 23.87 | + |
| Mybl2 | 5.67 | 6.04 | 5.62 | 2.07 | 3.46 | 2.63 | 1.75 | 2.68 | 1.9 | 1.82 | 1.45 | 1.25 | + |
| Med29 | 14.15 | 13.56 | 13.85 | 8.13 | 8.92 | 7.61 | 6.25 | 7.17 | 7.39 | 6.06 | 6.47 | 7.41 | + |
| Limd1 | 13.74 | 14.63 | 13.29 | 16.7 | 13.3 | 14.61 | 16.78 | 15.55 | 15.71 | 20.06 | 18.77 | 20.5 | + |
| I730030J21Rik | 7.97 | 9.09 | 9.8 | 25.52 | 21.32 | 24.34 | 16.53 | 10.88 | 12.91 | 23.26 | 19.94 | 24.55 | + |
| Prps2 | 28.83 | 28.64 | 27.62 | 24.19 | 24.79 | 25.02 | 19.98 | 24.56 | 21.97 | 22.1 | 24.68 | 21.53 | + |
| Wbp1 | 6.11 | 7.68 | 9.35 | 19.9 | 17.38 | 16.35 | 22.24 | 17.78 | 16.08 | 23.87 | 20.77 | 19.81 | + |
| Nsun5 | 19.18 | 19.17 | 20.18 | 12.3 | 13.07 | 12.5 | 8.24 | 11.32 | 10.41 | 9.09 | 8.22 | 9.62 | + |
| Tcp11l2 | 2.63 | 2.96 | 3.14 | 17.55 | 13.13 | 15.24 | 23.51 | 13.82 | 16.66 | 31.7 | 29.92 | 38.25 | + |
| Tcof1 | 32.03 | 34.17 | 37.03 | 20.16 | 21.1 | 20.48 | 15.82 | 22.17 | 21.21 | 18.37 | 16.11 | 16.15 | + |
| Rasal1 | 2.26 | 1.92 | 2.44 | 1 | 0.82 | 0.78 | 1.68 | 1.98 | 1.79 | 1.14 | 1.23 | 1.4 | + |
| Arsb | 9.03 | 12.41 | 8.76 | 10.55 | 11.04 | 9.29 | 11.17 | 7.37 | 8.53 | 14.37 | 13.34 | 15.64 | + |
| 4933426M11Rik | 1.96 | 2.21 | 1.9 | 2.9 | 2.63 | 2.98 | 2.92 | 3.46 | 3.82 | 3.56 | 3.18 | 3.32 | + |
| Slc25a22 | 15.09 | 13.99 | 15.52 | 9.12 | 9.55 | 11.11 | 8.95 | 8.66 | 9.24 | 7.83 | 8.65 | 9.86 | + |
| Dnajb6 | 108.36 | 113.83 | 105.11 | 96.1 | 90.85 | 89.41 | 116.62 | 112.77 | 105.79 | 127.9 | 134.83 | 129.46 | + |
| Tbc1d23 | 1.6 | 2.39 | 1.97 | 4.66 | 3.87 | 4.23 | 5.1 | 3.94 | 5.88 | 6.35 | 5.84 | 3.79 | + |
| Poli | 9.56 | 9.66 | 9.8 | 14.69 | 14.29 | 14.31 | 13.35 | 11.61 | 13.97 | 13.14 | 12.85 | 15.37 | + |
| Pisd-ps2 | 12.69 | 8.72 | 11.05 | 19.51 | 19.91 | 24.18 | 27.09 | 22.41 | 25.23 | 20.75 | 23.61 | 24.69 | + |
| Rps18 | 2484.5 | 2562.64 | 2609.01 | 3859.2 | 3873.7 | 3989.53 | 3865.8 | 3707.15 | 3715.16 | 4042.88 | 4463.16 | 3965.04 | + |
| Adat3 | 5.26 | 5.73 | 6.63 | 2.96 | 3.67 | 3.07 | 3.58 | 2.37 | 3.38 | 4.01 | 2.75 | 2.86 | + |
| Malt1 | 4.75 | 6.05 | 6.16 | 33.68 | 28.17 | 26.05 | 14.09 | 11.11 | 12.76 | 38.07 | 38.07 | 32.05 | + |
| Dhcr24 | 4.59 | 5.16 | 5.48 | 2.9 | 4.04 | 3.16 | 4.05 | 4.1 | 3.53 | 3.04 | 4.12 | 3.09 | + |
| Mphosph6 | 95.43 | 90.89 | 86.94 | 35.02 | 40.21 | 39.21 | 46.33 | 56.76 | 57.29 | 31.55 | 38.52 | 28.92 | + |
| Atp5a1 | 520.3 | 516.3 | 526.62 | 326.4 | 333.54 | 339.59 | 307.3 | 364.56 | 348.63 | 267.8 | 284.99 | 268.11 | + |
| Pik3r2 | 11.73 | 10.92 | 11.6 | 11.22 | 11.58 | 12.73 | 9.9 | 14.24 | 11.91 | 8.08 | 9.93 | 8.41 | + |
| Nat9 | 13.84 | 13.18 | 13.88 | 19.46 | 19.78 | 21.67 | 21 | 21.47 | 19.23 | 19.09 | 20.18 | 19.49 | + |
| Enpp2 | 0.19 | 0.13 | 0.13 | 5.75 | 6.55 | 6.39 | 1.03 | 1.54 | 1.72 | 4.11 | 2.53 | 3.8 | + |
| Rsph3b | 16.67 | 15.51 | 12.8 | 24.31 | 22.9 | 29.36 | 32.16 | 27.08 | 26.27 | 30.51 | 32.14 | 31.66 | + |
| Nsmaf | 8.4 | 8.93 | 7.72 | 9.53 | 7.96 | 8 | 9.21 | 8.29 | 9.49 | 11.16 | 12.14 | 13.29 | + |
| 2810408A11Rik | 1.36 | 0.49 | 0.52 | 2.34 | 2.76 | 2.93 | 7.11 | 2.49 | 4.97 | 3.62 | 3.31 | 3.48 | + |
| 4931406H21Rik | 0.73 | 0.65 | 0.84 | 2.39 | 1.73 | 1.83 | 1.91 | 1.54 | 1.71 | 2.48 | 2.17 | 2.2 | + |
| Tipin | 194.31 | 206.66 | 201.81 | 89.15 | 98.24 | 95.26 | 73.16 | 91.57 | 86.31 | 59.47 | 59.02 | 61.68 | + |
| Mrpl51 | 34.77 | 33.61 | 35.95 | 21.38 | 20.18 | 21.96 | 15.59 | 21.15 | 18.02 | 16.69 | 18.45 | 16.72 | + |
| Commd4 | 81.6 | 79.05 | 80.25 | 71.87 | 74.46 | 76.31 | 95.88 | 81.97 | 85.15 | 97.47 | 97.23 | 96.33 | + |
| Cul9 | 1.35 | 0.99 | 1 | 3.13 | 3 | 2.64 | 2.92 | 2.67 | 2.75 | 2.99 | 3.72 | 3.01 | + |
| Cdca3 | 62.11 | 58.78 | 57.66 | 37.5 | 50.06 | 44.57 | 28.23 | 31.54 | 27.11 | 28.1 | 35.21 | 23.86 | + |
| Arpc1a | 64.35 | 71.42 | 67.89 | 56.12 | 53.59 | 56.63 | 41.36 | 46.71 | 46.92 | 46.15 | 52.28 | 43.78 | + |
| Dram2 | 15.4 | 13.92 | 14.11 | 19.49 | 20.81 | 20.9 | 17.43 | 19.03 | 18.39 | 16.72 | 19.71 | 20.56 | + |
| Hat1 | 98.73 | 104.82 | 95.92 | 47.24 | 48.8 | 51.63 | 42.14 | 42.52 | 42.11 | 30.35 | 34.92 | 33.69 | + |
| Cenpn | 10.67 | 10.4 | 9.11 | 6.07 | 5.39 | 5.81 | 4.58 | 4.15 | 3.61 | 3.92 | 3.63 | 2.67 | + |
| Lmnb2 | 13.85 | 14.21 | 13.7 | 7.46 | 8.77 | 8.09 | 7.38 | 8.48 | 7.42 | 5.89 | 6.69 | 5.93 | + |
| Daglb | 6.04 | 7.56 | 5.45 | 4.82 | 4.24 | 4.84 | 4.45 | 4.84 | 5.26 | 4.27 | 4.39 | 4.85 | + |
| Cdc27 | 10.17 | 9.19 | 10.59 | 5.7 | 7.16 | 6.52 | 6.22 | 7.13 | 6.56 | 6.79 | 6.89 | 5.34 | + |
| Prss16 | 3.05 | 3.27 | 2.54 | 12.48 | 8.9 | 9.96 | 7.42 | 4.91 | 5.76 | 6.19 | 6.42 | 6.16 | + |
| Uqcr11 | 258.87 | 251.14 | 238.72 | 122.78 | 131.7 | 129.88 | 118.22 | 130.12 | 126.38 | 78.46 | 74.01 | 90.43 | + |
| Anapc10 | 11.22 | 10.07 | 9.69 | 7.24 | 7.13 | 6.44 | 6.93 | 6.78 | 6.76 | 7.13 | 6.55 | 7.63 | + |
| Psmf1 | 21.39 | 23.76 | 22.12 | 18.42 | 17.76 | 17.64 | 17.36 | 18.49 | 15.93 | 15.26 | 16.44 | 16.45 | + |
| Dnph1 | 41.64 | 38.88 | 37.01 | 10.75 | 13.54 | 9.75 | 10.47 | 15.54 | 15.24 | 4.74 | 6.25 | 6.12 | + |
| Ing4 | 18.04 | 16.63 | 15.25 | 50.65 | 39.55 | 45.2 | 57.64 | 44.83 | 46.15 | 68.39 | 71.92 | 73.33 | + |
| Chd6 | 1.09 | 1.01 | 1.12 | 1.95 | 2.14 | 2.1 | 1.61 | 1.73 | 1.74 | 1.98 | 2.27 | 1.78 | + |
| Ints5 | 19.71 | 21.29 | 18.48 | 11.42 | 11.23 | 10.75 | 10.5 | 10.7 | 12.12 | 10.01 | 8.21 | 12.09 | + |
| Adam17 | 17.41 | 15.71 | 14.75 | 14.25 | 12.02 | 13.28 | 10.33 | 12.98 | 11.61 | 13 | 12.7 | 11.16 | + |
| Rarg | 2.05 | 1.69 | 1.88 | 6.75 | 7.39 | 6.24 | 4.91 | 5.57 | 4.9 | 6.11 | 7.65 | 6.98 | + |
| Hnrnpk | 607.87 | 632.91 | 613.17 | 422.22 | 436.31 | 439.43 | 387.27 | 418.51 | 403.65 | 396.01 | 394.81 | 410.31 | + |
| Ust | 0.09 | 0.11 | 0.04 | 1.56 | 1.17 | 1.37 | 0.63 | 0.16 | 0.23 | 1.63 | 1.04 | 1.31 | + |
| Slc35b1 | 59.43 | 58.01 | 56.52 | 21.73 | 22.46 | 19.2 | 24.78 | 30.2 | 27.36 | 20.84 | 16.24 | 16.96 | + |
| Pop1 | 14.91 | 16.63 | 15.12 | 4.15 | 4.95 | 5.6 | 3.11 | 5.62 | 4.21 | 3.67 | 3.56 | 2.72 | + |
| Polr2h | 74.06 | 77.72 | 73.51 | 55.49 | 64.95 | 62.23 | 64.26 | 66.11 | 63.47 | 53.17 | 63.82 | 58.23 | + |
| C3 | 0.1 | 0.19 | 0.15 | 1.18 | 1.39 | 1.04 | 4.88 | 1.2 | 2.61 | 3.19 | 3.73 | 3.22 | + |
| Letm2 | 1.55 | 1.88 | 2.09 | 4.79 | 4.71 | 5.41 | 5.39 | 3.75 | 4.16 | 3.87 | 3.99 | 4.28 | + |
| Klc3 | 7.29 | 8.39 | 7.75 | 10.09 | 8.6 | 10.22 | 16.43 | 15.66 | 15.18 | 9.51 | 11.43 | 12.83 | + |

| | | | | | | | | | | | | | |
|---|---|---|---|---|---|---|---|---|---|---|---|---|---|
| Psmd2 | 176.97 | 172.08 | 170 | 104.96 | 107.94 | 108.02 | 103.14 | 108.39 | 108.44 | 95.67 | 107.32 | 99.26 | + |
| Drosha | 4.94 | 5.63 | 5.91 | 3.52 | 3.4 | 3.3 | 3.37 | 3.06 | 2.93 | 2.4 | 2.32 | 3.21 | + |
| C230052I12Rik | 28.16 | 26.95 | 28.54 | 11.83 | 13.86 | 13.13 | 7.81 | 12.47 | 9.36 | 7.7 | 6.45 | 6.64 | + |
| Hmgb3 | 39.19 | 36.94 | 38.24 | 32.04 | 39.54 | 35.47 | 18.81 | 22.8 | 19.33 | 22.87 | 23.29 | 24.54 | + |
| Klc4 | 3.2 | 4.03 | 3.19 | 11.27 | 9.03 | 9.67 | 10.91 | 9.81 | 10.81 | 11.14 | 12.92 | 12.98 | + |
| Wrap53 | 13.97 | 12.77 | 12.75 | 9.67 | 8.76 | 10.7 | 8.82 | 8.83 | 8.89 | 8.66 | 10.01 | 9.88 | + |
| Lgals4 | 8.08 | 6.99 | 6.58 | 25.29 | 21.21 | 24.74 | 32.42 | 24.29 | 29.99 | 43.83 | 50.76 | 43.26 | + |
| Zbtb45 | 3.17 | 3.53 | 3.76 | 2.07 | 2.16 | 1.64 | 0.91 | 1.37 | 1.45 | 1.6 | 1.12 | 2.1 | + |
| Stk4 | 9.1 | 9.46 | 9.16 | 9.85 | 8.7 | 9.46 | 11.53 | 10.2 | 9.68 | 13.65 | 13.64 | 13.6 | + |
| Aaas | 84.11 | 72.98 | 76.09 | 65.75 | 78.1 | 73.95 | 61.45 | 66.14 | 56.46 | 59.38 | 70.97 | 60.19 | + |
| Mir703 | 16.67 | 20.76 | 21.36 | 24.87 | 36.93 | 42.2 | 49.23 | 34.27 | 45.79 | 37.6 | 47.68 | 37.74 | + |
| Tbc1d17 | 11.08 | 9.84 | 11.7 | 19.76 | 17.03 | 16.69 | 19.15 | 19.89 | 19.88 | 21.82 | 25.79 | 24.68 | + |
| Gtf3c2 | 24.22 | 22.02 | 22.24 | 19.3 | 18.87 | 16.24 | 13.98 | 15.18 | 14.36 | 15.07 | 16.14 | 13.46 | + |
| Fchsd1 | 2.49 | 1.58 | 1.63 | 3.99 | 3.9 | 3.32 | 4.43 | 3.74 | 4.08 | 4.31 | 5.57 | 3.89 | + |
| Cisd2 | 27.2 | 25.26 | 26.96 | 18.85 | 21.93 | 20.46 | 22.17 | 23.71 | 22.64 | 20.68 | 19.76 | 20.78 | + |
| Polr1e | 20.17 | 18.45 | 18.89 | 9.92 | 9.99 | 9.88 | 8.18 | 9.18 | 10.56 | 6.59 | 6.28 | 7.34 | + |
| Nup133 | 13.26 | 13.42 | 14.66 | 8.55 | 10.26 | 9.79 | 7.34 | 10.07 | 7.45 | 6.55 | 7.58 | 6.5 | + |
| H2afz | 980.34 | 1090.68 | 977.45 | 690.21 | 729.38 | 735.63 | 476.87 | 536.31 | 517.47 | 449.95 | 476.44 | 474.43 | + |
| Zer1 | 0.35 | 0.64 | 0.36 | 1.65 | 1.51 | 1.66 | 1.66 | 1.13 | 1.32 | 2.05 | 2.03 | 1.98 | + |
| Gm5122 | 0.45 | 0.36 | 0.07 | 10.85 | 15.83 | 12.56 | 0.35 | 0.29 | 0.52 | 4.89 | 6.85 | 5.07 | + |
| 2610301B20Rik | 19.7 | 17.39 | 19.36 | 29.89 | 27.91 | 30.02 | 25.04 | 28.81 | 29.61 | 27.06 | 30.5 | 31.57 | + |
| Idh2 | 41.11 | 37.49 | 39.86 | 39.53 | 37.21 | 39.66 | 44.05 | 36.95 | 39.55 | 26.89 | 32.43 | 28.69 | + |
| Katnal1 | 1.51 | 1.35 | 1.51 | 2.04 | 1.94 | 1.93 | 2.55 | 2.25 | 2.41 | 2.18 | 2.02 | 2.12 | + |
| Rpn1 | 136.43 | 138.62 | 132.04 | 74.44 | 79.46 | 75.84 | 63.97 | 92.02 | 85.77 | 52.5 | 61.47 | 54.22 | + |
| Atp2a1 | 0.05 | 0.05 | 0.03 | 0.44 | 0.25 | 0.44 | 1.25 | 0.61 | 0.51 | 0.47 | 0.74 | 0.74 | + |
| Lepr | 1.53 | 1.38 | 1.36 | 0.55 | 0.7 | 0.47 | 0.7 | 0.65 | 0.69 | 0.61 | 0.67 | 0.52 | + |
| Tmem101 | 13.55 | 13.25 | 15.03 | 11.46 | 10.22 | 10.48 | 10.79 | 11.89 | 11.88 | 10.96 | 10.4 | 9.33 | + |
| Ctsz | 236.14 | 205.2 | 230.49 | 149.36 | 154.02 | 164.7 | 107.05 | 144.46 | 127.26 | 102.79 | 116.67 | 99.6 | + |
| Ppp1cb | 78.32 | 63.56 | 73.14 | 14.51 | 19.74 | 15.27 | 17.43 | 33.32 | 31.03 | 16.24 | 14.55 | 15.45 | + |
| Mrpl44 | 49.27 | 48.49 | 46.13 | 34.66 | 31.65 | 35.77 | 33.7 | 36.38 | 35.8 | 37.6 | 33.49 | 37.57 | + |
| Asap3 | 0.07 | 0.04 | 0.06 | 0.19 | 0.24 | 0.22 | 0.21 | 0.29 | 0.2 | 0.26 | 0.23 | 0.39 | + |
| Aprt | 296.98 | 280.13 | 284.08 | 164.87 | 168.89 | 168.63 | 201.2 | 228.68 | 213.18 | 166.6 | 180.27 | 164.21 | + |
| Cyc1 | 258.41 | 260.67 | 250.31 | 140.77 | 158.52 | 150.71 | 133.54 | 160.57 | 154.08 | 125.05 | 112.53 | 114.46 | + |
| Il1r2 | 17.05 | 9.36 | 9.78 | 58.83 | 73.07 | 79.34 | 48.35 | 33.33 | 37.11 | 38.8 | 54.61 | 52.75 | + |
| Cstf2 | 23.23 | 22.07 | 24.68 | 13.32 | 11.96 | 13.26 | 9.48 | 12.85 | 11.98 | 8.87 | 8.62 | 10.4 | + |
| Ncapd3 | 9.06 | 9.45 | 10.1 | 7.35 | 9.32 | 6.69 | 4.71 | 5.67 | 5.5 | 5.12 | 6.22 | 5.23 | + |
| Rpl5 | 1541.89 | 1599.04 | 1600.2 | 2023.58 | 2075.11 | 2114.96 | 1917.45 | 1959.24 | 1902.99 | 1913.28 | 2177.92 | 1916.59 | + |
| Nfia | 0.06 | 0.11 | 0.08 | 0.47 | 0.34 | 0.39 | 0.3 | 0.18 | 0.23 | 0.52 | 0.68 | 0.57 | + |
| Vdr | 1.47 | 1 | 1.49 | 4.29 | 6.14 | 5.84 | 2.42 | 2.1 | 2.09 | 5.26 | 4.6 | 3.67 | + |
| 9330179D12Rik | 0.43 | 0.79 | 2.01 | 3.69 | 4.83 | 5.51 | 3.01 | 4.59 | 3.62 | 2.65 | 5.42 | 4.81 | + |
| Actg1 | 3944.49 | 3910.57 | 3939.64 | 2234.98 | 2143.54 | 2200.82 | 3470.09 | 3069.97 | 3229.78 | 2559.97 | 2347.99 | 2587.7 | + |
| Gemin4 | 31.06 | 30.38 | 29.03 | 13.56 | 13.11 | 13.82 | 8.82 | 12.08 | 10.85 | 8.38 | 6.47 | 6.1 | + |
| Tep1 | 6.85 | 5.59 | 5.63 | 9.79 | 8.47 | 9.89 | 10.06 | 7.93 | 9.84 | 10.19 | 10.25 | 10.19 | + |
| Acad9 | 20.13 | 20.72 | 19.68 | 16 | 16.56 | 16.83 | 11.52 | 14.76 | 13.62 | 13.39 | 13.28 | 13.29 | + |
| Rrp1 | 178.51 | 168.1 | 174.94 | 124.07 | 120.72 | 124.27 | 115.85 | 116.83 | 120.23 | 111.11 | 111.32 | 112.79 | + |
| Mtap | 56.98 | 53.61 | 57.1 | 22.19 | 23.47 | 22.23 | 22.1 | 33.13 | 29.6 | 15.72 | 15.56 | 17.01 | + |
| Nup214 | 8.85 | 9.34 | 9.29 | 6.13 | 6.57 | 7.08 | 5.92 | 6.72 | 5.3 | 6.67 | 7.78 | 6.69 | + |
| Haus6 | 15.23 | 14.68 | 16.15 | 8.67 | 11.22 | 8.25 | 7.88 | 7.39 | 7.2 | 5.77 | 8.49 | 6.22 | + |
| Sgta | 79.64 | 76.82 | 75.24 | 55.9 | 58.67 | 59.09 | 52.09 | 59.22 | 52.46 | 51.36 | 51.94 | 53.29 | + |
| Cks2 | 228.79 | 211.7 | 210.68 | 129.92 | 165.55 | 158.97 | 111.95 | 118.4 | 119.47 | 115.32 | 114.47 | 109.39 | + |
| AI480653 | 2.44 | 2.28 | 2.5 | 9.2 | 9.19 | 8.22 | 1.2 | 1.7 | 1.4 | 1.62 | 1.39 | 0.63 | + |
| Cep97 | 9.09 | 9.91 | 10.21 | 15.08 | 15.7 | 15.6 | 17.61 | 17.32 | 15.7 | 18.96 | 18.07 | 18.99 | + |
| Zcrb1 | 84.74 | 89.31 | 83.72 | 49.28 | 50.47 | 58.91 | 50.29 | 54.61 | 56.44 | 47.03 | 42.96 | 42.56 | + |
| Sbno2 | 8.03 | 6.96 | 8.58 | 15.8 | 14.46 | 13.87 | 12.74 | 12.52 | 12.37 | 13.39 | 12.47 | 14.37 | + |
| Fam171b | 0.67 | 0.5 | 0.56 | 2.7 | 3.29 | 3.1 | 1.52 | 3.77 | 2.28 | 2.01 | 2.12 | 2.28 | + |
| Dhrs7 | 37.43 | 34.54 | 38.56 | 90.01 | 61.93 | 63.82 | 43.42 | 46.55 | 39.41 | 56.77 | 54.78 | 54.99 | + |
| Lsm7 | 171.26 | 168.72 | 173.64 | 71.54 | 68.16 | 67.6 | 63.6 | 95.63 | 82.65 | 51.46 | 52.28 | 52.22 | + |
| Lama5 | 3.95 | 4.01 | 3.63 | 22.11 | 21.43 | 20.65 | 6.45 | 8.13 | 7.89 | 6.76 | 9.1 | 7.1 | + |
| 2210016F16Rik | 37.82 | 35.2 | 36.17 | 22.26 | 24.02 | 24.32 | 15.5 | 19.88 | 18.82 | 15.17 | 12.51 | 15.85 | + |
| Rtn3 | 113.06 | 114.42 | 116.34 | 91.06 | 90.14 | 94.72 | 98.81 | 108.19 | 104.17 | 87.52 | 89 | 86.58 | + |
| U2af2 | 19.26 | 26.47 | 24.69 | 10.57 | 11.58 | 10.67 | 9.38 | 11.89 | 10.74 | 8.79 | 10.7 | 9.3 | + |
| Pgam5 | 62.33 | 60.56 | 63.61 | 50.17 | 51.25 | 54.18 | 49.34 | 52.61 | 49.36 | 45.44 | 49.07 | 50.73 | + |
| Gramd1a | 23.17 | 22.7 | 23.07 | 53.96 | 48.31 | 46.38 | 51.99 | 43.29 | 47.48 | 51.04 | 49.93 | 50.2 | + |
| Shcbp1 | 33.57 | 29.39 | 26.65 | 13.52 | 18.71 | 16.06 | 8.14 | 9.06 | 10.19 | 8.61 | 7.83 | 6.68 | + |
| Cbx3 | 334.31 | 341.84 | 338.34 | 206.91 | 216.86 | 221.02 | 176.11 | 185.26 | 186.32 | 177.76 | 182.85 | 182.16 | + |
| Irf2bp1 | 2.88 | 2.87 | 2.13 | 0.87 | 1.78 | 1.37 | 1.17 | 1.04 | 1.67 | 1.23 | 1.32 | 1.39 | + |
| Timm8b | 125.01 | 118.75 | 116.83 | 80.83 | 88 | 78.89 | 98.45 | 95.28 | 94.17 | 74.22 | 81.04 | 87.72 | + |
| Ubn2 | 2.35 | 2.13 | 2.04 | 3.88 | 3.96 | 3.93 | 3.87 | 3.73 | 3.3 | 4.95 | 5.3 | 5.01 | + |
| Bola3 | 79.71 | 80.15 | 82.63 | 41.26 | 37.81 | 40.92 | 30.83 | 42 | 39.58 | 25.47 | 25.82 | 27.08 | + |
| Mrpl28 | 114.39 | 111.46 | 115.77 | 61.54 | 61.49 | 61.86 | 62.54 | 66.49 | 69.62 | 44.05 | 44.44 | 47.74 | + |
| Zc3h4 | 5.69 | 5.52 | 5.41 | 6.05 | 5.16 | 5.34 | 5.61 | 5.08 | 5.27 | 7.45 | 6.3 | 7.36 | + |
| Rpl37a | 645.21 | 659.07 | 638.54 | 1078.73 | 1043.88 | 1109.84 | 1120.73 | 962.84 | 971.61 | 1284.4 | 1404.82 | 1309.62 | + |
| Tsg101 | 56.4 | 62.04 | 62.18 | 48.09 | 48.09 | 46.4 | 49.44 | 53.45 | 52.38 | 43.77 | 44.98 | 42.22 | + |
| Fbxo30 | 6.05 | 5.26 | 5.58 | 2.34 | 3.71 | 3.41 | 1.8 | 2.67 | 2.74 | 2.45 | 3.21 | 2.58 | + |
| Gm12505 | 10.83 | 10.84 | 10.8 | 15.7 | 20.86 | 18.15 | 17.36 | 19.56 | 18.36 | 20.36 | 22.48 | 20.31 | + |
| Cops5 | 93.99 | 90.71 | 92.62 | 53.61 | 57.03 | 55.22 | 53.99 | 55.52 | 53.73 | 47.58 | 52.98 | 43.79 | + |
| Drg1 | 136.11 | 131.47 | 131.91 | 85.65 | 88.39 | 88.88 | 72.66 | 76.04 | 78 | 63.81 | 73.68 | 67.97 | + |
| Golga2 | 10.7 | 11.13 | 10.46 | 17.26 | 14.63 | 15.46 | 18.94 | 17.35 | 17.22 | 18.07 | 20.77 | 18.66 | + |
| Pmpca | 86.84 | 83.14 | 87.04 | 43.2 | 44.71 | 47.73 | 37.18 | 44.7 | 45.44 | 31.02 | 27.5 | 32.23 | + |
| Snhg1 | 77.77 | 85.69 | 86.21 | 192.61 | 188.91 | 231.78 | 210.48 | 230.25 | 212.59 | 324.64 | 376.01 | 299.44 | + |
| Inpp5b | 48.24 | 44.84 | 49.4 | 21.23 | 25.64 | 27.19 | 27.24 | 33.42 | 32.27 | 20.14 | 20.41 | 19.99 | + |
| Cflar | 16.49 | 16.26 | 17.21 | 11.58 | 12.21 | 12.55 | 14.13 | 13.73 | 15.8 | 14.15 | 13.54 | 13.68 | + |
| Srgn | 788.25 | 691.04 | 748.43 | 1002.43 | 1056.73 | 1102.95 | 813.11 | 858.75 | 842.32 | 841.66 | 898.28 | 852.42 | + |
| Tjp3 | 2.26 | 1.69 | 1.66 | 4.82 | 5.31 | 4.73 | 5.82 | 5.17 | 5.03 | 8.02 | 6.69 | 7.8 | + |
| Cmah | 13.15 | 13.99 | 14.84 | 6.8 | 7.79 | 6.9 | 6.31 | 6.26 | 6.31 | 4.9 | 4.76 | 4.13 | + |
| Tmem219 | 25.76 | 29.59 | 28.64 | 48.68 | 42.72 | 45.8 | 46.26 | 47.22 | 47.74 | 58.24 | 56.6 | 57.83 | + |
| Pet112 | 11.88 | 10.54 | 11.27 | 4.22 | 5.82 | 4.4 | 6.3 | 6.57 | 6.37 | 3.79 | 4.57 | 4.05 | + |
| Wbp1l | 7.91 | 7.94 | 7.53 | 8.21 | 8.16 | 8.36 | 8.36 | 8.02 | 7.65 | 12.4 | 10.56 | 11.6 | + |
| Slc37a1 | 6.53 | 6.21 | 8.02 | 9.34 | 8.89 | 7.72 | 11.27 | 10.62 | 10.35 | 9.84 | 8.73 | 9.27 | + |
| Exosc4 | 38.77 | 37.75 | 38.62 | 28.32 | 27.15 | 28.67 | 25.85 | 26.94 | 25.76 | 25.84 | 27.36 | 26.62 | + |
| Dnajb1 | 36.66 | 32.36 | 39.43 | 23.99 | 26.86 | 26.84 | 25.2 | 32.04 | 28.95 | 25.8 | 25.23 | 28.74 | + |
| Mmgt2 | 13.28 | 14.77 | 13.33 | 14.37 | 13.66 | 12.66 | 17.67 | 17.03 | 15.63 | 16.44 | 18.03 | 19.19 | + |
| Npc1 | 5.73 | 6.54 | 6.32 | 9.99 | 9.74 | 10.25 | 9.6 | 9.9 | 9.66 | 10.76 | 9.59 | 10.96 | + |
| Psmc3 | 137.81 | 147.27 | 134.93 | 76.62 | 82.35 | 84.2 | 67.02 | 80.43 | 72.19 | 63.71 | 62.61 | 64.68 | + |
| Cct3 | 456.69 | 455.14 | 456.81 | 185.22 | 215.9 | 204.85 | 149.28 | 231.96 | 198.94 | 127.89 | 134.99 | 134.49 | + |
| Sqstm1 | 130.24 | 125.2 | 134.95 | 232.18 | 238.83 | 251.09 | 241.63 | 272.42 | 240.83 | 270.96 | 275.53 | 274.71 | + |
| Chchd5 | 19.82 | 18.19 | 20.39 | 32.43 | 32.75 | 34.48 | 34.38 | 31.84 | 29.75 | 31.37 | 35.93 | 29.54 | + |
| Pacsin3 | 1.23 | 2.19 | 1.57 | 5.07 | 5.57 | 5.69 | 5.41 | 4.8 | 5.66 | 6.15 | 5.9 | 5.02 | + |
| Txnrd1 | 19.81 | 21.92 | 23.03 | 10.75 | 12.59 | 10.18 | 9.95 | 12.52 | 10.9 | 9.18 | 10.19 | 7.5 | + |
| Dclk1 | 0.62 | 0.7 | 0.77 | 0.12 | 0.11 | 0.07 | 0.07 | 0.17 | 0.33 | 0.09 | 0.03 | 0.23 | + |
| Ankrd52 | 6.84 | 8.01 | 8.53 | 5.97 | 6.28 | 6.01 | 4.37 | 5.05 | 5.41 | 5.08 | 5.48 | 4.72 | + |
| Banp | 4.89 | 4.25 | 5.15 | 6.47 | 6.86 | 6.88 | 7.16 | 8.1 | 7.07 | 7.11 | 8.71 | 7.66 | + |
| Vcpip1 | 6.52 | 6 | 6.16 | 6.36 | 6.23 | 6.17 | 6.7 | 6.06 | 7.46 | 7.54 | 9.45 | 8.08 | + |
| Pdcd4 | 15.02 | 14.77 | 13.54 | 113.82 | 100.67 | 108.97 | 27.75 | 22.95 | 25.03 | 57.25 | 52.66 | 55.19 | + |
| Slc25a32 | 21.58 | 22.48 | 20.8 | 9.16 | 9.35 | 10.61 | 6.77 | 11.09 | 9.95 | 6.55 | 5.15 | 5.58 | + |
| Tbp | 27.49 | 28.49 | 28.09 | 17.84 | 19.01 | 19.25 | 14.88 | 16.23 | 16.28 | 17.52 | 14.89 | 15.4 | + |
| Lbh | 5.03 | 5.73 | 4.74 | 11.86 | 9.93 | 11.28 | 18.07 | 11.25 | 13.39 | 10.82 | 12.37 | 13.89 | + |
| Alg5 | 59.81 | 61.24 | 58.13 | 43.81 | 45.02 | 46.82 | 48.36 | 48.95 | 45.66 | 48.07 | 47.14 | 47.17 | + |
| Gpr18 | 37.04 | 37.37 | 40.54 | 147.68 | 131.16 | 133.08 | 70.3 | 69.63 | 69.21 | 117.52 | 112.06 | 127.65 | + |
| Mrps23 | 64.93 | 63.19 | 62.04 | 54.39 | 48.08 | 51.09 | 55.48 | 52.78 | 52.83 | 53.92 | 53.72 | 51.89 | + |
| Ndufa6 | 150.42 | 140.09 | 146.89 | 269.39 | 253.69 | 271.11 | 277.5 | 229.8 | 246.83 | 297.22 | 330.48 | 311.89 | + |
| Cox7a2l | 200.45 | 177.47 | 184.33 | 370.18 | 348.43 | 363.08 | 397.97 | 348.15 | 355.1 | 405.49 | 442.38 | 406.46 | + |
| 1110038F14Rik | 24.36 | 24.32 | 24.63 | 20.74 | 20.08 | 20.75 | 19.49 | 20.67 | 18.03 | 17.77 | 16.6 | 20.02 | + |
| Tmem199 | 46.21 | 47.14 | 51.41 | 23.76 | 24.42 | 25.1 | 21.78 | 27.03 | 27.54 | 19.6 | 20.08 | 18.32 | + |
| Ndufc2 | 163.81 | 172 | 173.05 | 125.11 | 137.31 | 138.15 | 147.22 | 147.32 | 149.21 | 115.57 | 140.73 | 120.01 | + |
| Kpna2 | 380.91 | 278.33 | 242.14 | 179.79 | 229.77 | 209.35 | 112.63 | 122.46 | 122.37 | 119.45 | 123.35 | 100.88 | + |
| Impad1 | 2.52 | 2.43 | 2.27 | 1.89 | 1.97 | 1.82 | 1.33 | 1.65 | 1.56 | 1.41 | 1.75 | 1.63 | + |
| Ttll12 | 20.49 | 19.58 | 20.61 | 10.85 | 10.96 | 11.16 | 14.56 | 13.55 | 13.43 | 11.15 | 9.65 | 8.7 | + |
| Ormdl2 | 37 | 34.79 | 32.44 | 21.6 | 24.07 | 24.05 | 17.42 | 23.52 | 22.17 | 19.03 | 20.44 | 20.24 | + |
| Nefh | 8.55 | 8.99 | 7.96 | 2.26 | 2.24 | 2.49 | 2.81 | 3.63 | 3.25 | 1.07 | 1.61 | 1 | + |
| Ech1 | 145.33 | 134.98 | 130.86 | 190.45 | 192.75 | 193.48 | 269.65 | 240.96 | 245.66 | 225.31 | 233.77 | 251.73 | + |
| Elp6 | 11.26 | 10.91 | 12.35 | 8.21 | 8.17 | 7.48 | 8.58 | 8.2 | 7.75 | 6.93 | 7.2 | 8.18 | + |
| Rnmt | 19.67 | 20.51 | 20.88 | 16.09 | 15.2 | 16.13 | 14.5 | 16.13 | 17.2 | 13.51 | 15.1 | 13.63 | + |
| Esco2 | 11.56 | 10.47 | 9.49 | 5.04 | 7.3 | 7.28 | 2.48 | 3.88 | 3.04 | 3.65 | 3.98 | 2.9 | + |
| Exosc9 | 59.04 | 60.27 | 56.7 | 30.08 | 37.58 | 37.15 | 35.28 | 39.67 | 35.58 | 30.4 | 34.54 | 26.24 | + |
| Mto1 | 13.98 | 11.27 | 12.95 | 8.56 | 8.73 | 10.55 | 10.52 | 11.23 | 9.62 | 9.17 | 9.07 | 9.24 | + |
| Asb2 | 0.57 | 0.54 | 0.62 | 1.79 | 1.67 | 1.14 | 3.61 | 1.53 | 3.11 | 3.04 | 1.63 | 2.57 | + |
| Snrpd3 | 125.61 | 131.41 | 139.17 | 61.84 | 70.17 | 61.84 | 65.35 | 72.45 | 67.51 | 43 | 44.9 | 41.13 | + |
| Parp2 | 38.15 | 39.14 | 41.79 | 32.03 | 30.44 | 30.51 | 27 | 31.24 | 29.62 | 24.67 | 30.68 | 27.31 | + |
| Rpf2 | 100.67 | 106.25 | 101.15 | 39.18 | 47.24 | 43.27 | 35.16 | 50.15 | 46.93 | 28.85 | 30.82 | 28.09 | + |
| Dhx8 | 12.41 | 12.08 | 13.81 | 9.76 | 8.41 | 10.26 | 9.3 | 10.22 | 11.31 | 10.24 | 9.78 | 8.98 | + |
| Myo6 | 0.31 | 0.21 | 0.16 | 0.62 | 0.45 | 0.56 | 0.73 | 0.57 | 0.55 | 0.53 | 0.48 | 0.64 | + |
| 2610507B11Rik | 9.74 | 9.51 | 10.56 | 7.59 | 7.02 | 8.24 | 6.11 | 7.26 | 7.26 | 6.46 | 6.38 | 6.57 | + |
| Ppp1r3b | 8.34 | 9.13 | 8.78 | 15.9 | 16.91 | 14.64 | 23.09 | 21.76 | 23.93 | 10.99 | 14.52 | 14.61 | + |
| Zfp280b | 7.1 | 7.66 | 7.28 | 5.49 | 5.73 | 5.17 | 5.83 | 4.49 | 4.84 | 4.58 | 5.01 | 4.3 | + |
| Cpsf1 | 32.92 | 32.81 | 31.84 | 27.03 | 28.77 | 28.66 | 22.73 | 26.49 | 26 | 25.62 | 23.44 | 25.86 | + |
| Jup | 0.18 | 0.21 | 0.12 | 0.57 | 0.72 | 0.66 | 3.69 | 2.02 | 2.15 | 2.27 | 2.53 | 2.5 | + |
| Eno3 | 18.22 | 17.31 | 18.66 | 9.35 | 11.93 | 9.6 | 9.46 | 10.72 | 9.22 | 7.81 | 5.95 | 7.37 | + |
| Rbm38 | 95.67 | 104.04 | 103.59 | 83.16 | 76.22 | 77.25 | 80.27 | 89.8 | 90.69 | 78.83 | 88.06 | 74.38 | + |
| Smarca5 | 10.65 | 12.29 | 11.46 | 6.56 | 7.45 | 7.03 | 7.59 | 7.77 | 6.93 | 8.35 | 7.34 | 6.32 | + |
| Parp16 | 3.91 | 3.66 | 3.92 | 12.4 | 13.58 | 13.51 | 22.31 | 17.07 | 16.52 | 21.84 | 23.98 | 19.36 | + |
| Msto1 | 42.84 | 41.94 | 44.15 | 23.68 | 25.3 | 21.94 | 25.86 | 26.98 | 24.67 | 22.3 | 18.82 | 20.13 | + |
| Pla2g6 | 6.16 | 5.06 | 5.67 | 8.77 | 7.51 | 8.14 | 7.56 | 7.73 | 7.17 | 8.85 | 7.4 | 6.95 | + |
| Mvp | 21.07 | 24.08 | 23.24 | 31.24 | 28.26 | 30.23 | 30.25 | 32.84 | 31.08 | 28.91 | 35.79 | 27.51 | + |
| Cpsf2 | 22.79 | 24.57 | 22.17 | 11.47 | 14.81 | 13.65 | 13.49 | 15.61 | 16.2 | 13.27 | 12.11 | 12.8 | + |
| Hnrnph2 | 45.98 | 48.27 | 45.28 | 32.91 | 34.17 | 33.13 | 28.83 | 31.13 | 30.01 | 33.12 | 28.13 | 30.92 | + |
| Ero1l | 77.36 | 87.01 | 99.35 | 104.43 | 120.87 | 110.54 | 316.03 | 224.49 | 284.34 | 123.1 | 150.64 | 143.26 | + |
| Ccdc59 | 32.2 | 32.6 | 33.05 | 22.51 | 23.51 | 24.43 | 22.34 | 23.76 | 23.34 | 22.5 | 22.75 | 22.73 | + |
| Slc5a6 | 5.06 | 4.3 | 4.4 | 2.85 | 2.91 | 2.1 | 2.55 | 2.59 | 2.39 | 2.61 | 2.44 | 2.77 | + |
| 1500012F01Rik | 82.48 | 79 | 77.07 | 291.27 | 268.3 | 293.2 | 279.54 | 269.32 | 256.91 | 365.71 | 422.48 | 361.79 | + |
| Tmem41b | 20.04 | 20.08 | 20.04 | 13.07 | 11.97 | 12.9 | 11.98 | 11.83 | 11.99 | 11.19 | 9.87 | 9.91 | + |

| | | | | | | | | | | | | | |
|---|---|---|---|---|---|---|---|---|---|---|---|---|---|
| Bcl2a1b | 108.19 | 103.95 | 92.37 | 90.68 | 94.82 | 98.43 | 123.53 | 96.63 | 106.05 | 191.18 | 170.56 | 197.69 | + |
| Cops8 | 62.39 | 62.16 | 62.69 | 38.07 | 39.19 | 39.51 | 36.28 | 40.54 | 40.31 | 34.36 | 35.15 | 38.39 | + |
| Rpl23 | 1242.34 | 1234.39 | 1222.64 | 1871.36 | 1864.83 | 1938.2 | 1719.12 | 1676.62 | 1649.05 | 1857.98 | 2027.07 | 1884.55 | + |
| Dennd4a | 9.56 | 9.78 | 9.03 | 16.13 | 14.04 | 14.46 | 13.28 | 12.21 | 11.11 | 12.89 | 13.96 | 14.03 | + |
| Sdf2l1 | 122.58 | 142.72 | 142.86 | 41.35 | 42.14 | 42.47 | 44.81 | 62.28 | 53.41 | 28.45 | 32.65 | 25.54 | + |
| Pwp1 | 59.21 | 61.8 | 60.89 | 34.46 | 37.47 | 37.02 | 25.47 | 39.16 | 33.98 | 28.72 | 32.36 | 26.26 | + |
| Pitrm1 | 37.36 | 36.4 | 35.62 | 15.49 | 19.1 | 15.91 | 12.2 | 17.5 | 14.54 | 9.19 | 10.83 | 10.56 | + |
| Smg9 | 10.59 | 8.71 | 9.18 | 14.13 | 15.35 | 13.14 | 14.57 | 12.87 | 13.24 | 14.06 | 15.63 | 14.28 | + |
| Sepn1 | 1.72 | 1.86 | 2.02 | 2.48 | 3.13 | 2.94 | 3.12 | 3.21 | 3.6 | 2.94 | 3.65 | 2.83 | + |
| Nolc1 | 77.53 | 83.57 | 81.03 | 24.45 | 26.56 | 29.99 | 16.75 | 30.85 | 28.79 | 16.88 | 15.99 | 14.46 | + |
| Tmem14c | 133.89 | 143.25 | 135.49 | 90.19 | 95.62 | 92.26 | 99.54 | 107.31 | 97.52 | 78.18 | 94.04 | 90.95 | + |
| Atad2 | 7.5 | 8.42 | 7.98 | 4.49 | 5.78 | 5.77 | 3.95 | 4.78 | 5.21 | 3.99 | 5.11 | 3.84 | + |
| Cmss1 | 14.92 | 15.3 | 11.87 | 8.58 | 7.94 | 8.71 | 7.9 | 8.58 | 9.12 | 6.94 | 6.05 | 6.35 | + |
| Ddx1 | 146.39 | 148.02 | 143.21 | 78.77 | 85.54 | 85.91 | 70.64 | 87.75 | 83.34 | 60.08 | 66.77 | 62.41 | + |
| Trmt44 | 3.98 | 3.58 | 3.94 | 1.9 | 3.1 | 2.06 | 1.32 | 2.62 | 1.91 | 2.59 | 2.13 | 2.32 | + |
| E430025E21Rik | 16.69 | 17.09 | 16.47 | 12.63 | 10.69 | 12.55 | 14.2 | 14.57 | 14.93 | 12.33 | 13.48 | 12.88 | + |
| Vps25 | 102.03 | 104.71 | 101.65 | 87.58 | 82.9 | 78.49 | 82.19 | 83.6 | 81.2 | 76.72 | 67.29 | 73.31 | + |
| Dync1li1 | 39.81 | 36.49 | 38.95 | 26.96 | 27.79 | 27.31 | 28.53 | 32.28 | 31.37 | 25.84 | 26.22 | 26.97 | + |
| Eif3f | 115.54 | 171.87 | 205.95 | 392.01 | 363.62 | 322.56 | 357.41 | 358.51 | 362.13 | 484.6 | 461.4 | 365.67 | + |
| Nap1l1 | 376.96 | 389.9 | 393.84 | 262.65 | 285.22 | 285.78 | 255.78 | 273.21 | 278.28 | 221.11 | 263.64 | 235.54 | + |
| Sgsm2 | 1.42 | 1.42 | 1.47 | 2.86 | 3.36 | 2.75 | 2.99 | 3.29 | 3.01 | 3.28 | 3.97 | 2.63 | + |
| Gadd45gip1 | 51.94 | 56.46 | 57.94 | 41.54 | 43.09 | 44.04 | 54.45 | 50.79 | 48.37 | 43.66 | 45.55 | 40.58 | + |
| Kcnab3 | 0.2 | 0.14 | 0.11 | 0.82 | 0.39 | 0.77 | 0.91 | 0.23 | 0.7 | 0.47 | 1.04 | 0.7 | + |
| Grpel1 | 43.89 | 44.03 | 41.03 | 28.63 | 25.24 | 30.39 | 28.89 | 28.58 | 31.03 | 23.27 | 23.74 | 21.86 | + |
| Tctn1 | 1.97 | 1.52 | 1.68 | 3.04 | 2.6 | 2.95 | 3.12 | 3.07 | 2.88 | 3.4 | 2.77 | 3.62 | + |
| Fbxl12 | 2.83 | 3.49 | 3.23 | 13.03 | 9.84 | 9.09 | 10.9 | 10.33 | 10.79 | 13.31 | 13.25 | 12.29 | + |
| Senp2 | 6.22 | 6.83 | 7.24 | 4.39 | 4.95 | 4.74 | 4.57 | 4.61 | 4.27 | 4.93 | 4.16 | 3.82 | + |
| Zmym6 | 1.64 | 1.15 | 1.7 | 3.67 | 3.13 | 3.43 | 4.34 | 3.33 | 3.1 | 3.6 | 3.18 | 4.55 | + |
| Cast | 45.82 | 40.47 | 41.84 | 65.59 | 64.43 | 61.72 | 66.38 | 62.1 | 64.32 | 75.47 | 78.02 | 78.48 | + |
| Gm20139 | 0.02 | 0.01 | 0.08 | 0.33 | 0.36 | 0.44 | 0.62 | 0.9 | 0.62 | 0.96 | 0.43 | 0.46 | + |
| Macrod1 | 0.18 | 0.23 | 0.3 | 0.64 | 1.03 | 0.45 | 0.77 | 1.38 | 1.35 | 1.44 | 0.77 | 0.95 | + |
| Mtmr1 | 2.94 | 2.9 | 2.81 | 3.64 | 3.42 | 3.7 | 2.27 | 3.09 | 3.23 | 5.33 | 5.25 | 5.23 | + |
| Abcb1b | 14.87 | 13.79 | 16.07 | 7.45 | 6.55 | 5.36 | 5.89 | 6.82 | 4.95 | 6.44 | 5.38 | 5.02 | + |
| Ssna1 | 74.66 | 67.56 | 75.57 | 47.4 | 61.24 | 60.38 | 62.13 | 63.9 | 57.81 | 59.92 | 59.42 | 56.44 | + |
| Zmynd19 | 5.37 | 6.33 | 6.02 | 3.49 | 4.26 | 3.21 | 2.66 | 2.82 | 3.4 | 2.23 | 2.75 | 3.79 | + |
| Gpr137b-ps | 1.03 | 1.27 | 0.85 | 2.53 | 2.09 | 1.97 | 5.62 | 3.1 | 3.45 | 3.37 | 3.2 | 4.97 | + |
| Krr1 | 19.3 | 19.76 | 19.84 | 11.9 | 13.22 | 10.9 | 13.52 | 15.77 | 15.23 | 13.1 | 14.55 | 13.13 | + |
| Ypel3 | 1.32 | 1.16 | 1.82 | 7.96 | 6.35 | 8.97 | 9.95 | 9.55 | 6.59 | 11.95 | 12.29 | 12.28 | + |
| Aldh16a1 | 18.03 | 17.31 | 16.99 | 11.12 | 11.13 | 10.17 | 10.5 | 9.86 | 10.31 | 8.29 | 8.36 | 9.82 | + |
| Birc5 | 148.89 | 138.65 | 124.49 | 76.57 | 100.06 | 92.12 | 61.1 | 59.96 | 50.77 | 54 | 56.68 | 44.51 | + |
| Car9 | 1.69 | 1.25 | 1.52 | 3.29 | 2.01 | 2.7 | 5.92 | 4.62 | 6.19 | 1.96 | 3.11 | 4.33 | + |
| Mllt3 | 2.62 | 2.73 | 2.79 | 6.84 | 5.89 | 6.16 | 5.54 | 4.92 | 4.58 | 8.27 | 8.11 | 8.29 | + |
| Zc3h12d | 15.48 | 13.56 | 17.53 | 22.27 | 24.84 | 24.35 | 30.44 | 27.72 | 25.73 | 26.94 | 31.77 | 29.42 | + |
| Ptrh2 | 19.12 | 17.64 | 19.04 | 9.54 | 11.55 | 11.99 | 11.58 | 13.68 | 12.03 | 10.86 | 10.54 | 11.87 | + |
| Lin9 | 5.23 | 5.96 | 4.94 | 3.36 | 4.23 | 4.3 | 2.4 | 3.19 | 3.38 | 3.32 | 3.26 | 2.46 | + |
| Mad2l1 | 71.1 | 70.11 | 67.15 | 44.73 | 52.3 | 50.14 | 35.85 | 37.64 | 38.45 | 33.85 | 38.96 | 36.09 | + |
| Gimap6 | 88.46 | 86.2 | 80.19 | 237.52 | 195.67 | 210.54 | 277.3 | 186.73 | 206.91 | 341.47 | 309.8 | 356.96 | + |
| Magohb | 47.7 | 46.57 | 42.54 | 22.94 | 20.94 | 21.31 | 22.66 | 27.99 | 28.72 | 19.54 | 21 | 19.04 | + |
| Aar2 | 12.01 | 12.23 | 12.02 | 10.54 | 10.25 | 9.1 | 7.68 | 8.23 | 9 | 9.79 | 7.33 | 9.95 | + |
| Sugt1 | 82.43 | 82.33 | 82.75 | 89.9 | 87.69 | 91.2 | 96.14 | 92.24 | 91.21 | 100.13 | 109.72 | 103.49 | + |
| Dennd5b | 0.55 | 0.55 | 0.7 | 0.19 | 0.43 | 0.24 | 0.28 | 0.37 | 0.29 | 0.23 | 0.35 | 0.26 | + |
| Dlat | 33.9 | 35.4 | 34.92 | 18.98 | 19.99 | 19.78 | 17.82 | 23.65 | 20.02 | 14.2 | 15.47 | 14.91 | + |
| Nmrk1 | 56.03 | 50.14 | 47.82 | 39.8 | 52.2 | 47.53 | 28.75 | 34.41 | 34.88 | 38.99 | 36.28 | 38.23 | + |
| Rpl23a | 2161.41 | 2264.65 | 2215.53 | 4118.65 | 4138.27 | 4301.16 | 3462.94 | 3246.04 | 3344.24 | 4139.28 | 4475.15 | 4234.77 | + |
| Cd24a | 1.01 | 0.32 | 0.45 | 3.34 | 3.85 | 2.98 | 5.19 | 3.01 | 3.6 | 7.66 | 10.11 | 8.82 | + |
| Mtx1 | 41.11 | 42.63 | 40.57 | 24.92 | 25.18 | 27.06 | 22.13 | 24.51 | 27.01 | 17.97 | 19.32 | 19.77 | + |
| Rfc2 | 99.05 | 103.1 | 104.5 | 70.7 | 72.16 | 69.75 | 62.2 | 62.45 | 60.95 | 45.33 | 48.55 | 47.69 | + |
| Snrpe | 512.96 | 569.42 | 530.63 | 331.92 | 327.14 | 337.06 | 349.81 | 336.6 | 339.46 | 325.94 | 343.84 | 329.57 | + |
| Sh3bp5l | 8.52 | 7.63 | 7.67 | 5.69 | 4.52 | 6.67 | 6.91 | 6.51 | 5.84 | 5.72 | 6.08 | 5.78 | + |
| Esf1 | 21.27 | 23.45 | 21.06 | 9.03 | 11.3 | 11.17 | 10.18 | 13.84 | 12.59 | 8.86 | 8.56 | 7.82 | + |
| Map3k2 | 0.59 | 0.54 | 0.53 | 1.05 | 0.96 | 0.86 | 0.94 | 0.79 | 1.06 | 0.86 | 1.19 | 1.12 | + |
| Samsn1 | 46.44 | 43.49 | 43.05 | 46.21 | 52.55 | 56.17 | 78.33 | 69.61 | 77.93 | 52.36 | 63.37 | 62.28 | + |
| Smarce1 | 41.26 | 44.31 | 43.15 | 35.41 | 36.9 | 35.88 | 32.04 | 35.24 | 35.25 | 30.92 | 33.35 | 32.19 | + |
| Slfn5 | 8.66 | 9.53 | 8.65 | 3.57 | 4.47 | 4.12 | 4.03 | 4.67 | 3.83 | 3.26 | 4.36 | 3.24 | + |
| Trip13 | 45.4 | 36.76 | 41.34 | 17.44 | 20.49 | 20 | 9.18 | 15.05 | 12.69 | 10.12 | 9.46 | 9.1 | + |
| Nme1 | 364.71 | 366.49 | 345.63 | 166.53 | 177.41 | 178.2 | 148.98 | 207.94 | 194.19 | 110.66 | 124.01 | 118.28 | + |
| Stradb | 4.9 | 4.65 | 5.34 | 8.54 | 9.11 | 9.19 | 10.27 | 9.22 | 8.41 | 8.57 | 8.94 | 10.37 | + |
| Srsf6 | 77.07 | 82.33 | 79.2 | 28.71 | 31.07 | 28.19 | 23.17 | 28.27 | 26.58 | 19.84 | 16.91 | 18.55 | + |
| Mrps12 | 74.19 | 70.27 | 70.98 | 40.47 | 37.23 | 42.42 | 32.68 | 37.1 | 37.49 | 31.7 | 35.68 | 32.36 | + |
| Cenpe | 5.47 | 4.71 | 4.23 | 3.57 | 4.85 | 4.62 | 2.88 | 2.72 | 3.01 | 3.63 | 3.5 | 2.94 | + |
| Rnf145 | 8.98 | 8.57 | 10.58 | 9.17 | 10.24 | 8.92 | 12.8 | 11.93 | 12.87 | 21.85 | 17.25 | 15.86 | + |
| Plcg2 | 8.53 | 9.92 | 9.47 | 19.26 | 15.31 | 17.82 | 25.08 | 19.12 | 22.01 | 20.62 | 19.7 | 24.29 | + |
| Nupl2 | 8.34 | 8.9 | 8.38 | 4.54 | 4.96 | 4.63 | 3.95 | 4.6 | 4.06 | 4.58 | 3.03 | 3.69 | + |
| Blvra | 57.53 | 55.55 | 54.61 | 31.01 | 36.58 | 33.03 | 32.86 | 40.21 | 34.12 | 25.96 | 27.65 | 30.94 | + |
| Erp29 | 71.73 | 72.18 | 73.21 | 59.44 | 61.22 | 59.15 | 62.69 | 63.6 | 63.84 | 51.18 | 61.68 | 57.6 | + |
| Rpl14 | 930.38 | 950.51 | 958.6 | 1268.01 | 1242.55 | 1302.12 | 1344.08 | 1235.19 | 1281.79 | 1324.67 | 1495.59 | 1314.84 | + |
| Ppm1j | 16.16 | 17.15 | 12.51 | 33.73 | 30.85 | 29.07 | 46.67 | 42 | 46.24 | 29.01 | 27.91 | 29.56 | + |
| Hsdl1 | 16.28 | 16.46 | 18.72 | 34.4 | 34.4 | 31.8 | 29.77 | 30.3 | 30.3 | 31.32 | 35.8 | 35.87 | + |
| Mbd3 | 12.29 | 13.73 | 15.11 | 7.04 | 8.01 | 7.91 | 7.05 | 7.28 | 7.33 | 5.76 | 6.55 | 5.22 | + |
| Trappc6a | 71.21 | 61.21 | 63.43 | 114.9 | 107.86 | 104.85 | 163.98 | 131.67 | 134.66 | 132.62 | 134.21 | 150.88 | + |
| Slco3a1 | 4.21 | 3.61 | 4.35 | 1.76 | 1.89 | 2.3 | 1.47 | 2.1 | 1.47 | 2.11 | 2.11 | 2.53 | + |
| Rchy1 | 28.99 | 31.56 | 32.8 | 44.47 | 44.12 | 48.39 | 51.16 | 53.64 | 50.92 | 47.26 | 59.88 | 50.99 | + |
| Slc39a1 | 47.56 | 49.99 | 51.2 | 45.35 | 43.83 | 46.19 | 35.2 | 35.24 | 33.33 | 36.51 | 35.74 | 39.62 | + |
| Hmbs | 62.92 | 64.2 | 61.21 | 37.26 | 41.73 | 43.29 | 38.56 | 42.76 | 42.71 | 29.14 | 29.27 | 28.5 | + |
| Bcl2 | 57.87 | 63.34 | 59.97 | 135.36 | 120.71 | 116.57 | 64.11 | 82.57 | 93.5 | 32.01 | 32.21 | 32.05 | + |
| Oaz1 | 465.53 | 460.24 | 449.24 | 374.68 | 375.2 | 365.02 | 354.8 | 379.29 | 382.59 | 335.08 | 345.53 | 355.76 | + |
| Gm6644 | 152.93 | 166.36 | 158.92 | 86.59 | 80.95 | 88.21 | 107.46 | 91.66 | 84.21 | 91.99 | 79.58 | 92.67 | + |
| Cars2 | 11.17 | 13.97 | 15.29 | 7.38 | 7.86 | 7.14 | 8.64 | 9.27 | 8.21 | 8.05 | 7.86 | 8.27 | + |
| Fam3a | 7.12 | 7.06 | 6.59 | 12.43 | 9.44 | 10.37 | 12.24 | 10.75 | 12.2 | 12.19 | 13.16 | 12.02 | + |
| Ncdn | 11.83 | 11.41 | 13.34 | 7.8 | 7.32 | 9.27 | 6.13 | 8.27 | 7.6 | 7.96 | 6.42 | 6.78 | + |
| Tspan32 | 17.05 | 16.16 | 20.59 | 31.93 | 31.93 | 31.76 | 30.25 | 26.9 | 25.79 | 30.91 | 29.94 | 26.36 | + |
| Nras | 70.77 | 75.39 | 74.82 | 53.93 | 60.48 | 61.72 | 56.93 | 63.92 | 65.8 | 57.47 | 58.91 | 55.51 | + |
| Hells | 39.55 | 39.93 | 45.65 | 24.08 | 29.55 | 28.69 | 18.77 | 18.81 | 20.58 | 21.03 | 21.48 | 21.61 | + |
| Zfp692 | 14.05 | 13.15 | 13.57 | 17.84 | 17.55 | 18.11 | 22.91 | 19.88 | 20.24 | 22 | 23.36 | 20.97 | + |
| Slc17a9 | 16.29 | 14.36 | 14.49 | 18.35 | 19.58 | 20.78 | 20.27 | 29.78 | 25.35 | 20.75 | 22.96 | 19.49 | + |
| Gfra1 | 0.05 | 0.13 | 0.12 | 0.35 | 0.81 | 0.26 | 0.87 | 2.53 | 1.31 | 0.75 | 1.01 | 0.61 | + |
| Dok1 | 19.36 | 18.06 | 19.23 | 44.75 | 43.5 | 44.76 | 33.27 | 34.98 | 33.62 | 31.06 | 36.33 | 28.33 | + |
| 4833439L19Rik | 57.17 | 53.07 | 53.23 | 29.51 | 29.38 | 29.01 | 28.95 | 36.24 | 38.2 | 27.81 | 26.85 | 27.73 | + |
| Rasgrp4 | 0.86 | 1.01 | 0.9 | 3 | 2.29 | 2.03 | 3.71 | 2.83 | 3.07 | 2.92 | 2.79 | 3.78 | + |
| Cenpw | 28.88 | 27.09 | 29.22 | 19.1 | 21.48 | 20.41 | 15.43 | 15.89 | 14.44 | 14.26 | 15.73 | 13.34 | + |
| 2410004N09Rik | 38.78 | 35.35 | 35.36 | 94.23 | 97.12 | 98.63 | 74.48 | 64.96 | 68.55 | 92.05 | 117.32 | 93.02 | + |
| Rnps1 | 152.29 | 142.94 | 150.26 | 113.28 | 115.29 | 117.5 | 98.63 | 112.88 | 108.16 | 96.5 | 97.15 | 98.66 | + |
| Psmd7 | 166.87 | 168.48 | 173.02 | 79.01 | 81.26 | 79.07 | 62.51 | 89.57 | 80.91 | 58.08 | 54.43 | 53.31 | + |
| Actb | 4433.72 | 4688.6 | 4295.84 | 1966.53 | 2002.43 | 2000.56 | 2545.49 | 2515.51 | 2651.86 | 1762.14 | 1482.72 | 1869.08 | + |
| Eno1 | 1742.64 | 1788.08 | 1828.97 | 1224.54 | 1196.52 | 1175 | 1663.36 | 1533.92 | 1657.05 | 1155.66 | 1172.08 | 1366.04 | + |
| Actr2 | 105.85 | 106.96 | 96.11 | 79.61 | 85.3 | 78.49 | 76.27 | 77.56 | 81.08 | 73.58 | 78.48 | 71.96 | + |
| Cox7a1 | 0.78 | 3.82 | 3.1 | 4.32 | 4.09 | 4.18 | 10.21 | 9.95 | 10.41 | 10.11 | 8.06 | 8.82 | + |
| Snrnp48 | 14.45 | 14.72 | 15.08 | 36.35 | 33.74 | 36.29 | 35.01 | 30.78 | 29.87 | 39.8 | 51.7 | 42.73 | + |
| Plcxd1 | 1.6 | 1.85 | 1.6 | 3.82 | 3.53 | 3.98 | 11.97 | 5.58 | 7.63 | 5.56 | 8.32 | 9.27 | + |
| Fam114a1 | 0.6 | 0.69 | 0.28 | 3.04 | 2.82 | 4.93 | 5.07 | 4.3 | 5.24 | 3.94 | 4.44 | 4.14 | + |
| Gcsh | 61.3 | 65.68 | 64.74 | 23.49 | 26.99 | 27.62 | 24.14 | 32.36 | 29.71 | 18.23 | 19.83 | 21.89 | + |
| Sass6 | 4.79 | 4.37 | 5.26 | 2.85 | 3.28 | 3.06 | 2.68 | 2.58 | 2.39 | 2.72 | 2.71 | 3.1 | + |
| Bdh1 | 17.67 | 17.16 | 17.35 | 9.17 | 8.99 | 10.59 | 10.33 | 12.04 | 11.34 | 7.3 | 7.39 | 7.83 | + |
| Noc4l | 97.71 | 98.11 | 98.88 | 39.09 | 42.67 | 37.29 | 33.71 | 44.65 | 37.86 | 27.44 | 28.62 | 29.34 | + |
| Dcaf6 | 1.57 | 1.36 | 1.31 | 3.46 | 3.51 | 3.01 | 3.6 | 3.13 | 2.48 | 4.05 | 4.55 | 3.58 | + |
| Ndufa4 | 372.95 | 395.46 | 385.06 | 164.09 | 185.86 | 194.52 | 155.95 | 202.36 | 195.07 | 118.17 | 137.17 | 125.48 | + |
| Ddr1 | 0.39 | 0.38 | 0.43 | 2.05 | 1.81 | 1.81 | 1.73 | 0.97 | 1.14 | 2.08 | 2.37 | 1.78 | + |
| Ccl9 | 25.69 | 17.27 | 17.01 | 2.84 | 3.5 | 4.18 | 14.71 | 35.46 | 21.68 | 12.67 | 12.83 | 13.69 | + |
| Wbscr22 | 31.75 | 33.91 | 35.15 | 18.17 | 19.97 | 17.81 | 17.56 | 19.76 | 17.68 | 14.03 | 13.71 | 16 | + |
| Acad11 | 5.34 | 4.41 | 4.62 | 8.25 | 7.46 | 8.55 | 7.96 | 9.03 | 6.97 | 6.23 | 9.28 | 6.85 | + |
| Usp15 | 51.37 | 46.45 | 46.88 | 32.24 | 31.94 | 34.54 | 32.12 | 30.66 | 34.31 | 27.67 | 26.93 | 32.31 | + |
| Cd151 | 1.42 | 1.61 | 1.49 | 3.77 | 2.86 | 3.52 | 3.71 | 2.5 | 3.53 | 3.19 | 4.32 | 4.55 | + |
| Laptm5 | 396.55 | 377.56 | 374.53 | 754.84 | 686.52 | 706.86 | 768.44 | 636.56 | 671.15 | 922.26 | 925.62 | 959.84 | + |
| Zwilch | 14.54 | 14.46 | 15.62 | 7.29 | 8.88 | 8.2 | 5.08 | 5.86 | 4.95 | 4.73 | 6.22 | 4.06 | + |
| Wdr8 | 30.94 | 30.04 | 25.82 | 22.44 | 20.29 | 21.08 | 21.48 | 22.06 | 21.83 | 21.18 | 21.8 | 22.88 | + |
| Spcs2 | 116.14 | 129.32 | 119.32 | 79.59 | 79.78 | 85.66 | 84.39 | 101.92 | 92.66 | 65.66 | 68.96 | 69.84 | + |
| Dus3l | 19.95 | 16.74 | 18.92 | 14.57 | 14.96 | 12.73 | 14.64 | 14.91 | 13.46 | 13.83 | 12.71 | 13.26 | + |
| Tmem147 | 111.45 | 105.97 | 98.42 | 69.39 | 67.32 | 69.28 | 72.11 | 81 | 84.69 | 63.4 | 58.51 | 64.98 | + |
| Eif2b1 | 59.71 | 58.43 | 60.81 | 27.43 | 28.8 | 30.34 | 28.35 | 30.51 | 29.58 | 30.2 | 26.7 | 26.09 | + |
| Atp6v1a | 55.93 | 46.36 | 49.08 | 24.33 | 27.64 | 27.62 | 28.77 | 32.07 | 31.49 | 21.03 | 20.34 | 18.9 | + |
| Ctsl | 7.04 | 6.79 | 7.79 | 17.57 | 16.77 | 17.37 | 14.17 | 13.56 | 12.19 | 18.69 | 19.95 | 18.16 | + |
| Minpp1 | 34.86 | 37.51 | 37.1 | 25 | 23.95 | 25.54 | 22.55 | 26.35 | 25.59 | 24.4 | 20.24 | 24.31 | + |
| Ccdc124 | 61.54 | 70.45 | 69.2 | 36.77 | 42.82 | 37.67 | 41.76 | 46.03 | 41.58 | 34.41 | 34.43 | 35.56 | + |
| Bin2 | 81.36 | 82.05 | 79.18 | 151.57 | 138.38 | 143.69 | 136.35 | 133.13 | 130.61 | 144.21 | 154.32 | 156.35 | + |
| Spp1 | 261.25 | 261.26 | 148.96 | 13.84 | 23.97 | 20.41 | 231.34 | 343.23 | 280.52 | 56.65 | 50.31 | 90.42 | + |
| Btla | 12.59 | 10.1 | 12.11 | 11.1 | 11.96 | 12.72 | 5.35 | 7.23 | 6.94 | 2.88 | 3.53 | 3.48 | + |
| Akap13 | 10.97 | 11.79 | 11.95 | 23.49 | 22.46 | 22.45 | 17.96 | 15.98 | 15.74 | 20.53 | 18.37 | 20.36 | + |
| Snrpb2 | 75.6 | 76.51 | 79.46 | 52.67 | 48.11 | 52.88 | 54.52 | 54.53 | 54.29 | 50.37 | 56.17 | 52.49 | + |
| Nup85 | 74.89 | 71.28 | 69.37 | 26.68 | 30.78 | 30.91 | 19.21 | 28.78 | 23.01 | 14.41 | 13.05 | 13.94 | + |
| Rbpms2 | 3.18 | 2.71 | 3.01 | 0.94 | 1.24 | 0.46 | 1.56 | 3.03 | 1.97 | 1.11 | 1.35 | 1.15 | + |
| Fkbp4 | 202.47 | 201.65 | 212.18 | 83.98 | 98.17 | 94.08 | 68.79 | 107.2 | 97.86 | 59.03 | 63.93 | 64.05 | + |
| Arrdc2 | 4.16 | 4.34 | 4.86 | 9.29 | 9.34 | 8.02 | 9.94 | 7.93 | 8.03 | 12.75 | 12.19 | 9.04 | + |
| Aptx | 3.73 | 3.61 | 3.41 | 3.27 | 3.17 | 3.19 | 3.34 | 3.31 | 2.91 | 2.15 | 2.38 | 2.13 | + |
| Rabac1 | 101.06 | 91.24 | 91.14 | 202.76 | 170.85 | 202.9 | 261.12 | 203.86 | 205.35 | 254.13 | 279.46 | 281.17 | + |
| Xpnpep1 | 57.07 | 54.16 | 56.11 | 36.28 | 36.82 | 34.66 | 40.03 | 37.72 | 39.75 | 29.55 | 30.3 | 34.91 | + |
| Wdr13 | 2.88 | 2.73 | 3.28 | 7.25 | 5.58 | 6.16 | 7.86 | 4.91 | 6.82 | 6 | 7.03 | 6.45 | + |
| Rpl35a | 1381.29 | 1404.95 | 1409.39 | 1854.87 | 1934.8 | 1996.96 | 1769.83 | 1749.42 | 1683.72 | 1839.29 | 2088.67 | 1855.24 | + |
| Ddx46 | 26.12 | 28.08 | 25.2 | 15.44 | 15.71 | 15.91 | 16.29 | 16.81 | 16.74 | 18.65 | 16.39 | 16.1 | + |
| Pcnxl4 | 6.45 | 5.25 | 5.7 | 4.66 | 3.81 | 3.38 | 2.56 | 3.35 | 2.97 | 2.52 | 2.47 | 3.34 | + |
| Ranbp3 | 35.02 | 36.8 | 35.2 | 25.83 | 24.56 | 24.23 | 26.76 | 25.68 | 22.54 | 24.92 | 23.14 | 26.61 | + |
| Irs2 | 1.14 | 0.86 | 1.08 | 4.07 | 4.8 | 4.07 | 1.42 | 2.44 | 1.77 | 2.02 | 2.6 | 2.84 | + |

| | | | | | | | | | | | | |
|---|---|---|---|---|---|---|---|---|---|---|---|---|
| Zc3h12a | 6.6 | 7.68 | 6.46 | 4.46 | 4.4 | 3.47 | 4.74 | 4.46 | 4.23 | 4.38 | 3.55 | 4.27 | + |
| Jag2 | 0.51 | 0.47 | 0.52 | 2.03 | 0.89 | 1.38 | 1.88 | 1.56 | 1.86 | 1.68 | 1.46 | 1.87 | + |
| Polr1a | 12.64 | 14.09 | 13.77 | 7.02 | 7.54 | 8.77 | 5.82 | 8.73 | 7.24 | 5.57 | 4.89 | 5.5 | + |
| Skp2 | 5.1 | 5.11 | 4.8 | 2.33 | 2.62 | 2.83 | 1.71 | 1.98 | 1.46 | 1.34 | 2.61 | 1.85 | + |
| Eif5a2 | 6.66 | 6.67 | 6.56 | 2.44 | 4.14 | 3.65 | 3.09 | 3.28 | 3.85 | 3.07 | 2.65 | 2.9 | + |
| Nap1l4 | 97.19 | 90.27 | 94.73 | 76.42 | 74.86 | 73.7 | 81.62 | 77.06 | 75.26 | 74.16 | 71.41 | 75.03 | + |
| Atp5f1 | 420.29 | 432.13 | 399.67 | 302.95 | 299.94 | 309.13 | 311.97 | 359.71 | 332.71 | 261.49 | 278.43 | 264.76 | + |
| Itpa | 65.08 | 68.23 | 71.85 | 42.75 | 47.99 | 44.29 | 39.57 | 50.02 | 47.12 | 43.09 | 51.84 | 45.56 | + |
| Acap1 | 65.37 | 63.97 | 65.63 | 162.36 | 148.47 | 161.63 | 255.39 | 176.81 | 195.67 | 217.82 | 230.81 | 235.48 | + |
| Ikzf3 | 5.57 | 5.91 | 4.88 | 8.61 | 7.91 | 7.11 | 13.25 | 11.41 | 12.15 | 11.82 | 13.58 | 11.79 | + |
| Acp1 | 62.33 | 58.62 | 60.33 | 50.04 | 53.26 | 56.53 | 51.42 | 46.71 | 45.87 | 50.26 | 55.03 | 49.98 | + |
| Rps15 | 1485.3 | 1520.05 | 1531.19 | 2401.14 | 2403.77 | 2513.85 | 2303.68 | 2182.45 | 2263.39 | 2378.9 | 2648.13 | 2453.58 | + |
| Vamp4 | 3.38 | 3.95 | 3.99 | 6.74 | 6.8 | 6.1 | 8.72 | 6.5 | 6.93 | 7.52 | 8.25 | 9.77 | + |
| Ppp6r2 | 0.74 | 1.49 | 1.27 | 3.22 | 2.18 | 3.22 | 3.46 | 3.75 | 3.41 | 3.67 | 4.18 | 2.19 | + |
| Ddx11 | 6.21 | 5.5 | 5.78 | 3.98 | 5.05 | 4.92 | 3.95 | 3.82 | 4.01 | 4.52 | 3.66 | 4.83 | + |
| Pomgnt1 | 4.81 | 4.3 | 6 | 1.79 | 1.88 | 2.6 | 1.48 | 2.76 | 1.87 | 1.69 | 1.63 | 1.74 | + |
| Tmem209 | 21.2 | 24.27 | 24.42 | 24.61 | 25.79 | 26.8 | 19.94 | 23.41 | 22 | 18.62 | 17.17 | 17.26 | + |
| Plrg1 | 53.35 | 60.39 | 60.18 | 43.87 | 49.59 | 43.28 | 42.28 | 40.86 | 40.5 | 37.19 | 41.71 | 36.52 | + |
| Hnrnpl | 8.98 | 9.1 | 9.52 | 5.26 | 6.5 | 5.96 | 4.62 | 6.11 | 5.54 | 6.54 | 6.65 | 6.62 | + |
| 2610008E11Rik | 1.54 | 1.8 | 1.59 | 3.08 | 3.24 | 2.74 | 2.85 | 2.56 | 2.74 | 3.4 | 2.62 | 2.52 | + |
| Arl4c | 2.99 | 3.79 | 3.15 | 12.14 | 9.88 | 9.33 | 10.76 | 8.43 | 10.96 | 9.77 | 10.78 | 11.31 | + |
| Trmt6 | 29.64 | 30.64 | 28.72 | 13.3 | 12.91 | 13.59 | 9.26 | 11.83 | 13.51 | 9.74 | 7.69 | 8.48 | + |
| Galk2 | 30.92 | 27.62 | 27.48 | 15.37 | 17.49 | 15.82 | 19.59 | 20.6 | 21.08 | 16.96 | 17.77 | 19.9 | + |
| Tmbim4 | 96.21 | 95.93 | 90.27 | 91.6 | 88.77 | 96.53 | 104.76 | 97.19 | 94.88 | 115.75 | 113.84 | 133.15 | + |
| Golgb1 | 1.75 | 1.51 | 1.61 | 3.06 | 3.1 | 3.79 | 3.84 | 2.79 | 3.24 | 3.46 | 4.3 | 3.25 | + |
| Zbtb25 | 8.37 | 7.94 | 9.8 | 13.6 | 11.83 | 13.57 | 19.05 | 16.88 | 14.84 | 17.36 | 18.92 | 19.25 | + |
| Nudt16 | 17.49 | 18.37 | 19.09 | 27.81 | 32.81 | 29.1 | 34.1 | 32.97 | 31.4 | 36.65 | 32.08 | 36.03 | + |
| Ndufaf1 | 31.71 | 30.52 | 31.5 | 21.52 | 18.87 | 21.45 | 18.61 | 23.56 | 20.74 | 16.52 | 19.11 | 18.26 | + |
| Ccdc116 | 1.28 | 1.35 | 1.61 | 0.35 | 0.22 | 0.3 | 0.15 | 0.2 | 0.4 | 0.18 | 0.27 | 0.2 | + |
| BC029214 | 10.07 | 11.88 | 8.46 | 21.73 | 19.93 | 20.94 | 23.18 | 21.55 | 19.21 | 24.07 | 31.97 | 26.43 | + |
| Ftsj3 | 58.84 | 53.7 | 61.56 | 18.66 | 22.94 | 23.73 | 14.68 | 24.68 | 21.07 | 12.82 | 11.94 | 13.49 | + |
| Epcam | 0.8 | 0.88 | 0.63 | 2.05 | 2.3 | 2.14 | 4.24 | 3.62 | 3.44 | 1.89 | 1.85 | 1.29 | + |
| Acss1 | 2.21 | 2.82 | 2.74 | 14.25 | 12.09 | 10.46 | 15.07 | 8.33 | 9.23 | 25.5 | 22.98 | 25.59 | + |
| Ttc13 | 16.22 | 16.35 | 16.18 | 10.12 | 12.58 | 10.35 | 12.48 | 11.62 | 13.2 | 9.1 | 10.39 | 10.3 | + |
| Map2k3 | 88.35 | 80.91 | 83.37 | 57.05 | 59.83 | 62.54 | 46.32 | 44.77 | 45.1 | 53.43 | 54.64 | 53.67 | + |
| Fbxo10 | 0.15 | 0.18 | 0.13 | 0.86 | 0.75 | 0.61 | 1 | 0.72 | 0.54 | 0.65 | 0.88 | 1.06 | + |
| Rbks | 5.57 | 6.64 | 6.8 | 2.16 | 2.31 | 2.55 | 2.92 | 2.41 | 2.72 | 2.01 | 1.98 | 2.31 | + |
| Grhpr | 38.75 | 39.63 | 35.57 | 24.41 | 23.01 | 27.26 | 28.5 | 20.17 | 19.91 | 19.22 | 23.28 | 26.66 | + |
| Slc10a3 | 13.4 | 12.52 | 12.21 | 6.66 | 7.45 | 7.57 | 5.51 | 6.34 | 5.95 | 6.6 | 4.25 | 6.94 | + |
| Hiat1 | 21.22 | 22.86 | 22.64 | 21.08 | 18.91 | 19.78 | 17.49 | 17.09 | 16.98 | 15.95 | 16.32 | 18.96 | + |
| Tmem171 | 0.79 | 1.81 | 0.42 | 0.6 | 0.86 | 0.39 | 1.64 | 1.32 | 1.95 | 3.89 | 1.85 | 3.64 | + |
| Ap3d1 | 16.69 | 17.84 | 15.47 | 12.12 | 11.28 | 11.33 | 12.42 | 12.17 | 11.69 | 12.03 | 11.82 | 12.05 | + |
| Krtcap2 | 201.48 | 219.95 | 192.94 | 255.44 | 252.48 | 269.67 | 332.56 | 307.33 | 318.27 | 271.27 | 284.26 | 283.29 | + |
| Gm9199 | 0.89 | 0.99 | 0.72 | 3.68 | 2.99 | 2.75 | 2.73 | 1.97 | 2.02 | 3.29 | 3.48 | 3.82 | + |
| Pfas | 39.42 | 39.59 | 38.89 | 13.05 | 14.06 | 13.06 | 8.76 | 12.53 | 11.37 | 7.24 | 6.93 | 6.87 | + |
| Dnajb11 | 49.53 | 69.08 | 71.84 | 30.75 | 38.25 | 30.99 | 36.91 | 47.19 | 39.22 | 39.4 | 33.05 | 30.09 | + |
| Prdx5 | 147.62 | 129.15 | 130.3 | 162.15 | 162.86 | 166.4 | 271.45 | 212.74 | 216.57 | 188.17 | 210.15 | 193.71 | + |
| A330032B11Rik | 4.87 | 6.92 | 6.2 | 3.29 | 3.35 | 3.39 | 3.35 | 3.39 | 3.82 | 3.45 | 3.4 | 2.88 | + |
| Gm12191 | 905.85 | 908.94 | 907.46 | 1432.59 | 1417.22 | 1520.97 | 1423.29 | 1393.82 | 1406.34 | 1515.69 | 1653.64 | 1508.9 | + |
| Pgd | 85.56 | 91.14 | 88.15 | 46.49 | 44.52 | 40.32 | 40.15 | 40.29 | 44.31 | 35.52 | 37.33 | 42.15 | + |
| Ier5l | 0.15 | 0.07 | 0.21 | 1.78 | 1.16 | 1.43 | 1.36 | 1.54 | 1.24 | 2.68 | 2.13 | 3.62 | + |
| Scap | 13.4 | 15.32 | 12.55 | 7.22 | 8.25 | 7.25 | 7.16 | 7.87 | 7.88 | 5.37 | 4.79 | 5.68 | + |
| Wdr46 | 25.99 | 24.85 | 27.21 | 14.35 | 14.51 | 14.86 | 11.32 | 12.75 | 12.2 | 8.65 | 9.43 | 9.23 | + |
| Fam65a | 4.34 | 4.15 | 4.38 | 7.25 | 7.28 | 8.01 | 9.41 | 6.89 | 6.57 | 9.43 | 8.34 | 9.36 | + |
| Ercc3 | 18.43 | 17.62 | 19.01 | 10.88 | 11.7 | 11.14 | 10.04 | 12.61 | 12.2 | 9.7 | 9.8 | 10.5 | + |
| Sf3a3 | 101.62 | 104.32 | 111.97 | 58.47 | 60.73 | 65.8 | 46.9 | 56.28 | 54.41 | 42.61 | 45.69 | 48.67 | + |
| Hnrnpab | 31.62 | 30.15 | 30.26 | 8.28 | 9.78 | 11.07 | 7.64 | 11.35 | 10.38 | 6.1 | 4.76 | 5.54 | + |
| Dalrd3 | 17.47 | 16.75 | 17.63 | 11.6 | 15.03 | 13.09 | 10.43 | 12.21 | 9.96 | 13.05 | 13.46 | 13.43 | + |
| Eme1 | 7.94 | 6.82 | 7.6 | 4.02 | 3.67 | 3.16 | 1.87 | 1.98 | 2.83 | 2.53 | 1.95 | 1.77 | + |
| G3bp2 | 59.21 | 66.26 | 63.63 | 36.87 | 41.27 | 38.5 | 34.12 | 34.34 | 36.43 | 34.43 | 33.12 | 32.61 | + |
| Ddb1 | 127.65 | 119.06 | 125.4 | 86.81 | 95.51 | 91.94 | 89.59 | 101.95 | 93.75 | 79.62 | 88.74 | 80.45 | + |
| Nfkbil1 | 9.12 | 9.26 | 10.15 | 19.91 | 17.88 | 17.31 | 27.44 | 19.9 | 19.23 | 24.59 | 31.99 | 28.74 | + |
| Hspbp1 | 66.83 | 61.39 | 64.77 | 26.46 | 29.11 | 27.56 | 22.86 | 34.38 | 32.56 | 19.03 | 14.33 | 19.25 | + |
| Aim1 | 11.5 | 11.81 | 10.85 | 29.08 | 26.66 | 26.74 | 13.42 | 11.55 | 12.49 | 21.43 | 19.75 | 21.6 | + |
| Pole | 6.68 | 6.52 | 5.9 | 5.12 | 5.24 | 5.27 | 3.32 | 3.84 | 3.79 | 2.23 | 2.26 | 1.93 | + |
| B3gnt8 | 0.28 | 0.37 | 0.67 | 1.49 | 2.34 | 2.19 | 1.6 | 2.23 | 1.87 | 1.59 | 1.42 | 2.5 | + |
| Loxl2 | 4.27 | 5.09 | 4.41 | 1.42 | 2.14 | 1.89 | 2.75 | 2.33 | 2.67 | 1.66 | 1.44 | 1.65 | + |
| Sc5d | 58.13 | 55.41 | 53.07 | 49.11 | 45.85 | 49.49 | 47.55 | 45.83 | 47.83 | 45.27 | 41.58 | 46.12 | + |
| Mettl17 | 12.68 | 11.62 | 13.59 | 8.61 | 9.03 | 7.66 | 8.54 | 7.66 | 8.89 | 10.13 | 7.1 | 9.73 | + |
| Nup160 | 11.77 | 11.86 | 12.11 | 5.55 | 6.47 | 5.79 | 3.74 | 5.55 | 5.48 | 3.74 | 3.85 | 2.75 | + |
| Gramd4 | 3.15 | 2.96 | 3.44 | 10.53 | 9.58 | 9.9 | 6.92 | 4.72 | 4.65 | 11.73 | 9.82 | 10.71 | + |
| Rft1 | 7.18 | 7 | 6.66 | 5.62 | 5.24 | 5 | 4.46 | 4.89 | 5.42 | 4.59 | 4.37 | 4.53 | + |
| Dnajc3 | 16.77 | 19.68 | 17.07 | 14.91 | 15.08 | 13.96 | 12.84 | 12.69 | 12.23 | 12.62 | 16.04 | 14.1 | + |
| Casp4 | 17.1 | 15.32 | 16.57 | 36.64 | 35.4 | 37.5 | 24.93 | 29.16 | 22.1 | 33.08 | 28.84 | 28.28 | + |
| Lrrc47 | 8.26 | 9.13 | 9.71 | 5.26 | 6.77 | 7.59 | 5.61 | 6.48 | 6.23 | 6.87 | 6.84 | 6.54 | + |
| Wnk1 | 20.6 | 22 | 20.78 | 38.63 | 38.33 | 39.27 | 31.12 | 30.64 | 34.21 | 28.09 | 27.82 | 27.93 | + |
| Timm44 | 64.78 | 60.84 | 65.74 | 44.58 | 42.39 | 43.98 | 47.98 | 46.61 | 43.69 | 42.34 | 40.74 | 45.16 | + |
| Trim16 | 6.9 | 6.25 | 5.9 | 3.88 | 3.44 | 4.01 | 8.78 | 7.07 | 7.59 | 2.85 | 2.66 | 2.4 | + |
| Rnf126 | 5.35 | 5.78 | 5.85 | 3.71 | 3.47 | 3.27 | 4.6 | 4.15 | 4.22 | 3.54 | 3.31 | 3.42 | + |
| Snx21 | 1.05 | 1.53 | 1.24 | 5.53 | 4.64 | 3.9 | 6.05 | 3.98 | 4.34 | 5.27 | 5.23 | 6.33 | + |
| Dlst | 73.43 | 67.15 | 74.42 | 49.27 | 54.08 | 47.74 | 36.09 | 49.84 | 45.96 | 37.62 | 41.81 | 37.62 | + |
| Mak16 | 106.94 | 103.43 | 102.19 | 63.33 | 61.08 | 64.74 | 70.56 | 79.81 | 79.99 | 62.41 | 66.51 | 59.36 | + |
| Wdr91 | 8.38 | 6.49 | 6.89 | 10.79 | 12.11 | 13.37 | 12.75 | 12.72 | 13.49 | 14.49 | 15.81 | 16.66 | + |
| Sumo3 | 56.29 | 56.74 | 51.02 | 36.38 | 43.36 | 36.82 | 32.44 | 40.55 | 38.86 | 33.08 | 36.42 | 29.83 | + |
| Haghl | 9.1 | 9.21 | 9.05 | 17.3 | 18.58 | 18.12 | 18.91 | 15.37 | 17.34 | 17.09 | 14.77 | 18.59 | + |
| Elovl1 | 106.22 | 109.49 | 105.3 | 73.3 | 71.58 | 68.65 | 72.22 | 81.59 | 82.23 | 70.1 | 69.57 | 67.85 | + |
| Gtf2h3 | 19.26 | 19.96 | 17.56 | 13.29 | 11.76 | 12.85 | 11.04 | 11.09 | 11.78 | 10.29 | 9.16 | 9.46 | + |
| Neu3 | 17.52 | 15.26 | 15.67 | 9.05 | 9.33 | 7.28 | 9.37 | 12.22 | 11.93 | 8.79 | 7.22 | 7.99 | + |
| Trpm4 | 2.58 | 2.08 | 1.95 | 3.2 | 2.76 | 3.24 | 5.9 | 5.11 | 5.26 | 4.07 | 4.65 | 5.2 | + |
| Pdia6 | 236.68 | 221.37 | 225.78 | 90.3 | 102.35 | 112.66 | 94.25 | 130.32 | 114.97 | 68.48 | 83.03 | 73.64 | + |
| Ddx55 | 9.08 | 11.23 | 9.99 | 5.85 | 6.73 | 6.86 | 6.92 | 6.07 | 7.51 | 6.63 | 6.66 | 6.62 | + |
| Zmpste24 | 27.16 | 28.96 | 27.35 | 15.42 | 17.82 | 16.82 | 14.67 | 15.99 | 16.85 | 13.93 | 14.44 | 15.41 | + |
| Ccdc17 | 1.37 | 0.93 | 1.11 | 2.24 | 3.24 | 2.54 | 1.95 | 2.85 | 2.17 | 2.86 | 2.15 | 2.56 | + |
| Cops7b | 16.33 | 17.81 | 16.1 | 13.25 | 11.89 | 13.81 | 13.93 | 14.16 | 13.51 | 11.1 | 13.84 | 12.35 | + |
| H2-T10 | 5.99 | 6.52 | 4.81 | 9.38 | 10.13 | 9.86 | 10.17 | 9.26 | 9.58 | 11.9 | 13.97 | 12.45 | + |
| Cep170b | 2.8 | 2.75 | 2.91 | 4.57 | 4.91 | 4.51 | 5.06 | 3.95 | 4.33 | 4.68 | 5.63 | 5.53 | + |
| Brwd1 | 2.86 | 3.51 | 3.54 | 6.6 | 6.62 | 6.54 | 6.15 | 5.08 | 5.88 | 6.99 | 8.72 | 6.86 | + |
| Akap8l | 13.72 | 13.48 | 12.34 | 28.98 | 28.65 | 28.65 | 29.46 | 27.16 | 26.63 | 35.48 | 38.43 | 35.69 | + |
| Tnfrsf18 | 253.75 | 260.35 | 263.38 | 720.3 | 685.7 | 687.65 | 486.6 | 474.49 | 503.16 | 384.77 | 445.12 | 474.82 | + |
| Gnl3 | 85.42 | 86.14 | 89.26 | 40.95 | 46.29 | 45.93 | 28.46 | 50.04 | 48.19 | 35.92 | 35.1 | 27.66 | + |
| Ubtf | 20.73 | 25.56 | 24.66 | 13.11 | 13.38 | 12.02 | 9.21 | 11.28 | 10.17 | 10.54 | 8.63 | 9.39 | + |
| Rps15a-ps6 | 175.33 | 161.99 | 158.79 | 295.88 | 284.73 | 295.38 | 342.18 | 286.52 | 286.46 | 388.28 | 374.4 | 369.48 | + |
| Ramp3 | 3.69 | 1.43 | 1.95 | 40.17 | 52.32 | 52.75 | 4.57 | 5.58 | 3.87 | 22.02 | 21.6 | 23.22 | + |
| Rab8a | 107.07 | 103.7 | 97.95 | 105.2 | 103.49 | 100.06 | 141.25 | 103.19 | 106.87 | 150.39 | 131.16 | 156.39 | + |
| Ythdf2 | 50.73 | 49.29 | 49.68 | 41.3 | 39.97 | 40.5 | 30.65 | 37.91 | 36.32 | 34.64 | 37.1 | 33.16 | + |
| Gspt1 | 6.84 | 7.01 | 7.1 | 4.4 | 4.8 | 5.61 | 3.74 | 5.41 | 4.01 | 4.99 | 5.57 | 4.55 | + |
| Prmt5 | 63.94 | 65.55 | 65.95 | 33.96 | 32.55 | 32.11 | 29.92 | 35.52 | 33.68 | 23.24 | 28 | 29.28 | + |
| Bloc1s3 | 1.64 | 1.54 | 1.74 | 3.15 | 2.85 | 2.58 | 2.92 | 2.58 | 3.17 | 3.32 | 3.28 | 2.92 | + |
| Qdpr | 101.54 | 102.65 | 104.55 | 78.63 | 72.42 | 83.4 | 82.98 | 84.01 | 82.63 | 67.55 | 72.36 | 70.41 | + |
| Bms1 | 31.61 | 31.31 | 36.72 | 16.64 | 18.28 | 18.17 | 14.79 | 18 | 15.73 | 13.85 | 14.43 | 13.81 | + |
| Pbld1 | 28.09 | 31.58 | 30.37 | 8.6 | 10.53 | 9.95 | 11.62 | 13.09 | 12.18 | 10.65 | 10.13 | 8.67 | + |
| Rnf31 | 15.59 | 14.15 | 13.77 | 25.48 | 25.85 | 26.52 | 25.21 | 23.85 | 23.87 | 30.15 | 30.11 | 31.86 | + |
| Tmem97 | 120.85 | 116.42 | 120.02 | 49.59 | 56.94 | 57.07 | 49.45 | 60.45 | 55.38 | 36.24 | 40.55 | 34.85 | + |
| Atl2 | 9.01 | 8.65 | 9.49 | 6.95 | 7.24 | 6.94 | 4.05 | 5.06 | 4.39 | 5.82 | 7.03 | 4.87 | + |
| Lrrk2 | 0.25 | 0.18 | 0.2 | 0.09 | 0.08 | 0.09 | 0.52 | 0.54 | 0.22 | 0.67 | 0.52 | 0.48 | + |
| Tmed5 | 41.7 | 45.85 | 40.56 | 27.2 | 31.75 | 30.52 | 24.91 | 28.72 | 29.21 | 28.85 | 25.66 | 28.31 | + |
| Tuba1c | 143.91 | 105.77 | 108.23 | 52.93 | 75.01 | 69.79 | 41.24 | 42.47 | 43.16 | 30.96 | 32.78 | 33.11 | + |
| Ubap2 | 33.18 | 29.83 | 31.4 | 22 | 22.31 | 22.68 | 18.21 | 20.5 | 21.5 | 17.87 | 18.28 | 18.8 | + |
| Ube2i | 133.69 | 133.77 | 133.66 | 97.14 | 99.63 | 102.05 | 103.15 | 103.54 | 102.72 | 100.75 | 94.74 | 100.67 | + |
| Ube2l3 | 106.41 | 103.37 | 104.69 | 63.51 | 67.4 | 67.66 | 66.57 | 70.88 | 70.81 | 55.54 | 59.67 | 56.06 | + |
| Arrdc4 | 0.89 | 1.02 | 1.9 | 9.62 | 10.89 | 11.95 | 11.84 | 12.19 | 11.16 | 22.39 | 17.5 | 11.99 | + |
| Sar1b | 93.31 | 96.04 | 94.72 | 73.31 | 70.79 | 72.7 | 68.01 | 78.47 | 75.02 | 69.71 | 75.56 | 70.16 | + |
| Mthfs | 19.9 | 15.51 | 19.51 | 38.51 | 36.07 | 40.35 | 51.37 | 35.34 | 38.67 | 54.88 | 53.26 | 56.61 | + |
| Mcm3 | 167.15 | 180.06 | 168.8 | 90.49 | 103.54 | 97.24 | 66.13 | 77.38 | 76.44 | 55.91 | 48.22 | 57.69 | + |
| Acsl4 | 40.31 | 37.08 | 43.23 | 21.92 | 25.72 | 24.4 | 22.55 | 26.34 | 25.98 | 20.2 | 17.68 | 17.9 | + |
| Tom1l2 | 2.69 | 2.35 | 2.31 | 5.81 | 4.67 | 5.64 | 4.44 | 3.79 | 4.13 | 4.72 | 4.14 | 4.63 | + |
| Odc1 | 37.96 | 39.43 | 38.19 | 19.94 | 21.79 | 20.35 | 16.79 | 17.53 | 17.59 | 15.33 | 17.12 | 14.42 | + |
| 4930539E08Rik | 2.67 | 2.55 | 2.09 | 4.08 | 4.67 | 3.72 | 5.66 | 6.67 | 5.36 | 2.85 | 4.83 | 4.49 | + |
| Mss51 | 0.41 | 0.29 | 0.18 | 0.55 | 0.65 | 1.26 | 1.51 | 0.87 | 1.06 | 1.22 | 1.21 | 1.29 | + |
| BC030867 | 11.81 | 9.24 | 9.95 | 6.21 | 7.63 | 7.1 | 4.36 | 3.9 | 3.78 | 3.73 | 4.29 | 3.35 | + |
| Riok3 | 40.04 | 37.99 | 37.29 | 48.33 | 52.41 | 49.98 | 50.75 | 52.94 | 52.12 | 45.7 | 56.67 | 47.39 | + |
| Snapc3 | 34.34 | 32.49 | 31.85 | 25.74 | 24.69 | 25.55 | 24.6 | 27.61 | 25.49 | 21.94 | 20.96 | 20.94 | + |
| Prps1l3 | 29.06 | 31.04 | 30.85 | 15.63 | 16.48 | 16.2 | 11.43 | 15.86 | 15.92 | 12.34 | 10.94 | 12.95 | + |
| Cish | 223.23 | 221.22 | 220.99 | 241.73 | 245.16 | 244.84 | 124.42 | 181.5 | 184.04 | 89.45 | 104.41 | 89.56 | + |
| Ccp110 | 5.54 | 5.64 | 5.69 | 3.1 | 2.82 | 3.03 | 2.44 | 3.11 | 3.02 | 2.48 | 2.98 | 2.58 | + |
| Rac3 | 2.81 | 1.78 | 2.21 | 4.4 | 4.34 | 4.73 | 5.81 | 7.3 | 7.86 | 6.69 | 6.23 | 6.84 | + |
| Nsa2 | 69.88 | 66.85 | 66.12 | 92.7 | 95.63 | 93.85 | 85.77 | 83.33 | 83.76 | 96.82 | 104.33 | 97.17 | + |
| Rnaseh2b | 37.46 | 37.18 | 37.55 | 21.34 | 28.45 | 28.95 | 16.68 | 21.85 | 21.69 | 16.41 | 19.8 | 17.88 | + |
| Eef2 | 1787.93 | 1727.18 | 1799.42 | 2542.63 | 2548.36 | 2606.62 | 2749 | 2721.58 | 2674.46 | 2540.47 | 2871.76 | 2492.22 | + |
| Ssr3 | 132.16 | 129.14 | 132.94 | 146.37 | 142.78 | 147.09 | 163.25 | 160.04 | 155.04 | 176.12 | 200.43 | 170.53 | + |
| Tnfrsf26 | 20.38 | 15.5 | 17.87 | 39.93 | 34.8 | 36.45 | 28.24 | 29.11 | 30.36 | 33.66 | 30.25 | 34.55 | + |
| Szrd1 | 14.91 | 14.61 | 13.51 | 9.47 | 8.63 | 8.49 | 8.27 | 9.49 | 8.06 | 6.7 | 9.28 | 7.78 | + |
| Atp6ap2 | 47.52 | 48.01 | 49.82 | 31.9 | 34.9 | 34.64 | 28.91 | 30.81 | 32.16 | 29.45 | 30.75 | 32.86 | + |
| Cryl1 | 7.41 | 6.97 | 6.02 | 5.2 | 4.52 | 4.61 | 4.24 | 4.21 | 3.96 | 3.65 | 4.06 | 3.19 | + |
| Srsf9 | 7.62 | 6.76 | 6.28 | 3.75 | 4.46 | 3.95 | 2.42 | 3.23 | 3.09 | 2.51 | 2.76 | 2.83 | + |
| Eif2s3x | 192.95 | 188.24 | 204.47 | 124.6 | 131.57 | 130.45 | 122.37 | 144.82 | 143.61 | 104.36 | 131.06 | 111.52 | + |
| Sfr1 | 112.6 | 115.39 | 112.18 | 121.24 | 121.86 | 126.19 | 124.35 | 122.54 | 121.73 | 129.77 | 148.34 | 139.72 | + |
| Cpsf3l | 65.53 | 57.14 | 62.64 | 44.63 | 49.42 | 43.57 | 43.33 | 45.88 | 41.23 | 35.46 | 36.3 | 42 | + |
| Plekhb2 | 38.95 | 37.11 | 39.26 | 73.18 | 72.42 | 68.41 | 84.26 | 79.1 | 86.41 | 81.01 | 97.65 | 84.82 | + |
| Creld2 | 98.78 | 90.34 | 98.82 | 25.9 | 28.79 | 27.96 | 23.42 | 36.39 | 28.4 | 18.75 | 18.03 | 17.37 | + |
| Mboat1 | 5.77 | 6.6 | 6.19 | 2.48 | 2.71 | 2.39 | 3.15 | 2.67 | 3.35 | 2.88 | 1.71 | 2.42 | + |

| | | | | | | | | | | | | |
|---|---|---|---|---|---|---|---|---|---|---|---|---|
| Ccr2 | 7.12 | 9.95 | 5.21 | 3.53 | 1.67 | 2.13 | 52.88 | 26.55 | 32.1 | 30.45 | 17.27 | 36.68 | + |
| Crtam | 67.48 | 54.38 | 59.61 | 19.52 | 18.26 | 19.7 | 32.13 | 48.13 | 41.15 | 22.05 | 25.75 | 20.24 | + |
| Lars2 | 159.7 | 168.4 | 177.51 | 255.42 | 251.46 | 265.36 | 259.6 | 273.28 | 267.32 | 298.78 | 289.53 | 269.86 | + |
| Sdf4 | 79.59 | 78.22 | 84.19 | 197.56 | 196.62 | 192.64 | 173.03 | 137.85 | 160.89 | 195.87 | 202.03 | 207.21 | + |
| Ccl3 | 839.16 | 721.18 | 717.97 | 32.48 | 72.3 | 45.12 | 76.74 | 186.87 | 197.2 | 64.16 | 39.38 | 61.1 | + |
| Igfbp4 | 25.53 | 29.34 | 24.64 | 38.12 | 32.8 | 37.84 | 33.65 | 24.82 | 34.85 | 9.66 | 9.78 | 10.27 | + |
| Exosc10 | 41.16 | 41.24 | 41.88 | 27.42 | 30.84 | 31.55 | 28.8 | 28.24 | 31.56 | 24.63 | 27.78 | 27.59 | + |
| Serpinb6a | 133.12 | 124.94 | 125.38 | 113.22 | 101.4 | 110.17 | 118.07 | 105.7 | 108.21 | 105.9 | 104.49 | 109.9 | + |
| Banf1 | 317.27 | 319.22 | 328.18 | 158.08 | 182.65 | 173.87 | 183.51 | 215.16 | 213.4 | 117.67 | 135.06 | 127.37 | + |
| Mrps27 | 50.82 | 47.26 | 45.25 | 28.87 | 29.45 | 31.99 | 30.47 | 31.11 | 33.53 | 20.49 | 25.82 | 24.8 | + |
| Lpcat3 | 50.21 | 51.67 | 49.56 | 41.04 | 39.37 | 41.32 | 32.46 | 29.88 | 32.14 | 31.59 | 28.36 | 31.9 | + |
| Rer1 | 128.96 | 130 | 130.82 | 103.16 | 103.78 | 113.75 | 113.19 | 113.14 | 112.04 | 103.71 | 107.65 | 105.57 | + |
| Nmd3 | 104.85 | 108.92 | 101.65 | 71.39 | 74.51 | 72.14 | 64.75 | 77.61 | 78.04 | 61.1 | 66.52 | 60.82 | + |
| Mtmr3 | 5.07 | 6.07 | 5.38 | 13.14 | 8.97 | 9.81 | 9.7 | 9.23 | 9.34 | 8.21 | 8.75 | 7.14 | + |
| Rb1 | 2.65 | 2.77 | 2.59 | 3.69 | 3.85 | 3.89 | 5.01 | 4.51 | 4.65 | 4.22 | 4.88 | 4.55 | + |
| Wrb | 12.55 | 13.87 | 13.15 | 5.12 | 5.43 | 5.24 | 4.42 | 6.65 | 5.12 | 5.32 | 4.03 | 5.13 | + |
| Gadd45g | 101.02 | 115.06 | 107.84 | 92.59 | 109.8 | 93.83 | 80.43 | 100.83 | 103.29 | 32.21 | 37.15 | 33.36 | + |
| Naa38 | 120.82 | 121.78 | 119.2 | 79.9 | 74.1 | 76.13 | 80.13 | 81.27 | 72.68 | 63.39 | 56.35 | 61.49 | + |
| Xlr3a | 0.5 | 0.48 | 1.12 | 1.86 | 2.27 | 1.75 | 2.05 | 1.81 | 2.35 | 1.82 | 1.67 | 2.1 | + |
| Mapre3 | 2.12 | 1.13 | 2.04 | 6.76 | 4.06 | 5.35 | 6.21 | 4.99 | 5.35 | 8.22 | 6.82 | 6.17 | + |
| Emc8 | 12.46 | 13.52 | 13.36 | 4.2 | 4.91 | 4.64 | 3.82 | 4.8 | 4.5 | 3.4 | 2.86 | 3.53 | + |
| 1600020E01Rik | 6.27 | 5.29 | 7.14 | 14.64 | 12.42 | 12.25 | 15.38 | 12.39 | 11.6 | 19.72 | 16.6 | 17.9 | + |
| Mkks | 8.49 | 8.29 | 7.88 | 6.13 | 6.96 | 6.29 | 6.09 | 6.33 | 5.98 | 5.23 | 5.15 | 5.83 | + |
| Taf13 | 19.44 | 22.39 | 21.18 | 14.96 | 15 | 13.36 | 12.48 | 17.45 | 16.44 | 12.91 | 14.3 | 11.99 | + |
| Imp3 | 81.42 | 78.19 | 82.41 | 57.3 | 64.32 | 64.91 | 67.88 | 75 | 74.51 | 66.37 | 69.45 | 64.5 | + |
| Gpx4 | 268.45 | 259.43 | 269.03 | 351.23 | 362.17 | 365.94 | 330.56 | 331.29 | 309.57 | 319.6 | 350.16 | 344.87 | + |
| Phyh | 16.58 | 16.03 | 16.43 | 28.29 | 28.5 | 27.04 | 27.48 | 24.98 | 27.33 | 24.9 | 27.3 | 30.49 | + |
| Srp19 | 131.36 | 142.31 | 147.57 | 85.45 | 91.38 | 92.67 | 89.03 | 104.25 | 103.13 | 76.02 | 71.96 | 71.38 | + |
| Ubxn6 | 27.19 | 27.25 | 28.02 | 41.17 | 43.08 | 41.13 | 58.45 | 49.44 | 48.4 | 50.88 | 52.08 | 51 | + |
| Gm2a | 55.64 | 53.25 | 54.04 | 88.14 | 84.56 | 88.4 | 146.27 | 112.12 | 121.57 | 132.84 | 135.08 | 133.98 | + |
| Vdac2 | 238.8 | 241.04 | 234.9 | 176.63 | 177.33 | 181.76 | 202.48 | 215.18 | 208.92 | 176.21 | 193.31 | 179.31 | + |
| Prmt6 | 3.96 | 4.12 | 3.83 | 2.51 | 2.17 | 2.46 | 1.92 | 1.94 | 2.13 | 1.68 | 2.31 | 1.26 | + |
| Nup153 | 7.04 | 6.89 | 6.41 | 4.7 | 4.83 | 4.94 | 4.87 | 4.94 | 5.51 | 4.47 | 4.91 | 3.66 | + |
| Gmppb | 38.25 | 36.08 | 40.23 | 17.71 | 18.61 | 19.56 | 16.45 | 20.21 | 19.46 | 12.31 | 10.42 | 12.03 | + |
| Anapc4 | 47.55 | 44.84 | 46.98 | 32.78 | 36.33 | 37.44 | 32.65 | 34.87 | 30.5 | 31.87 | 30.72 | 25.85 | + |
| Eif6 | 189.84 | 193.17 | 187.66 | 97.92 | 106.27 | 108.84 | 83 | 117.16 | 105.16 | 72.44 | 80.51 | 79.9 | + |
| Bphl | 9.26 | 8.23 | 8.05 | 14.53 | 11.98 | 16.98 | 17.36 | 13.6 | 15.17 | 14.43 | 12.34 | 12.25 | + |
| Hist1h2ao | 414.31 | 415.22 | 385.61 | 202.84 | 288.52 | 279.6 | 162.94 | 176.57 | 150.93 | 135.27 | 159.43 | 119.86 | + |
| Atp10d | 0.83 | 0.77 | 0.57 | 3.86 | 5.3 | 4.48 | 1.57 | 1.49 | 2.08 | 3.71 | 4.39 | 2.98 | + |
| Ss18l1 | 0.43 | 0.28 | 0.61 | 1.42 | 1.37 | 1.76 | 1.66 | 1.91 | 1.03 | 1.47 | 1.64 | 1.56 | + |
| Nudt19 | 40.36 | 40.75 | 39.97 | 25.55 | 21.7 | 26.67 | 22.09 | 23.34 | 24.15 | 19.67 | 19.59 | 21.35 | + |
| Dnaja3 | 43.7 | 42.55 | 44.41 | 34.4 | 38.44 | 38.6 | 40.96 | 41.07 | 38.27 | 36.47 | 38.3 | 33.83 | + |
| Mllt4 | 0.58 | 0.42 | 0.58 | 1.04 | 1.06 | 1.02 | 1.26 | 1.05 | 1.34 | 1.02 | 1.29 | 1.05 | + |
| Tmem205 | 17.76 | 20.17 | 17.52 | 29.11 | 32.11 | 29.38 | 36.98 | 31.73 | 38.09 | 23.71 | 24.51 | 27.36 | + |
| Usb1 | 30.91 | 31.1 | 31.48 | 18.38 | 19.38 | 20.64 | 24.69 | 24 | 26.66 | 19.19 | 17.89 | 20.85 | + |
| Fam193b | 2.01 | 2.21 | 2.03 | 5.52 | 5.45 | 5.23 | 5.67 | 4.56 | 5.17 | 6.54 | 7.54 | 6.71 | + |
| Nup205 | 12.71 | 12.57 | 12.93 | 7.63 | 8.16 | 7.01 | 6.31 | 7.33 | 7.29 | 4.9 | 6.44 | 4.91 | + |
| Cct7 | 479.94 | 471.54 | 489.28 | 262.15 | 291.78 | 276.2 | 265.58 | 336.93 | 314.39 | 213.44 | 247.2 | 216.72 | + |
| Klhl24 | 1.17 | 0.75 | 1.13 | 7.63 | 6.08 | 7.49 | 8.86 | 8.18 | 8.7 | 11.9 | 13.85 | 11.08 | + |
| Mrpl13 | 72.91 | 71.36 | 69.33 | 42.6 | 51.22 | 48.16 | 48.59 | 56.79 | 49.45 | 38.29 | 38.47 | 39.96 | + |
| Sf3b4 | 57.09 | 64.26 | 63.56 | 29.68 | 28.48 | 28.18 | 21.33 | 25.27 | 23.65 | 19.96 | 19.39 | 21.37 | + |
| Zfp828 | 14.7 | 17.47 | 16.71 | 8.8 | 9.84 | 9.56 | 6.55 | 8.94 | 9.24 | 8.18 | 7.03 | 7.4 | + |
| Fyttd1 | 18.1 | 17.54 | 18.29 | 13.3 | 12.37 | 14.9 | 12.92 | 13.47 | 13.99 | 11.31 | 12.82 | 11.44 | + |
| Nhp2 | 277.2 | 279.4 | 285.32 | 120.36 | 133.17 | 136.8 | 116.3 | 154.62 | 138.65 | 86.03 | 89.87 | 83.82 | + |
| Clpx | 28.33 | 29.34 | 26.85 | 22.74 | 26.52 | 23.52 | 25.14 | 25.26 | 24.02 | 23.28 | 24.57 | 21.06 | + |
| Parp4 | 1.85 | 2.09 | 1.95 | 3.35 | 3.63 | 3.68 | 3.5 | 4.44 | 4 | 4.76 | 4.89 | 4.09 | + |
| Arl6ip4 | 64.15 | 61.09 | 65.73 | 46.15 | 45.21 | 46.51 | 44.63 | 41.99 | 41.89 | 45.4 | 40.72 | 42.67 | + |
| Lgr4 | 0.42 | 0.31 | 0.29 | 0.99 | 0.83 | 0.95 | 0.65 | 0.71 | 0.43 | 0.47 | 1 | 0.98 | + |
| Rpl9 | 1997.08 | 2042.85 | 2041.13 | 3143.54 | 3094.21 | 3251.62 | 3098.59 | 2876.92 | 2919.07 | 3322.8 | 3624.6 | 3298.1 | + |
| 2310057M21Rik | 12.99 | 13.85 | 13.42 | 6.39 | 5.81 | 7.37 | 5.58 | 5.73 | 6.25 | 4.86 | 5.29 | 4.79 | + |
| Selk | 110.53 | 109.66 | 108.08 | 76.87 | 78.37 | 84.57 | 95 | 88.85 | 87.17 | 91.03 | 88.94 | 90.61 | + |
| Rere | 0.68 | 0.75 | 0.6 | 1.67 | 1.24 | 1.82 | 1.09 | 0.83 | 1.1 | 1.58 | 1.21 | 1.35 | + |
| Adh5 | 105.13 | 109.32 | 112.3 | 85.3 | 85.71 | 85.15 | 90.94 | 94.98 | 93.25 | 77.58 | 80.24 | 77.53 | + |
| Metap2 | 99.16 | 105.16 | 97.77 | 54.14 | 60.21 | 54.12 | 49.06 | 56.94 | 55.26 | 42.88 | 45.13 | 43.75 | + |
| Sh2d1a | 80.51 | 81.78 | 85.27 | 57.45 | 54.46 | 55.6 | 54.73 | 46.33 | 46.22 | 44.98 | 42.22 | 48.21 | + |
| Cluh | 15.13 | 16.33 | 18.87 | 7.71 | 8.57 | 9.3 | 6.28 | 8.76 | 8.82 | 7.48 | 6.28 | 5.71 | + |
| Ftsj1 | 9.91 | 10.17 | 9.4 | 6.62 | 6.08 | 6.59 | 4.91 | 5.53 | 6.28 | 6.43 | 5.4 | 5.68 | + |
| H2afv | 21.67 | 21.48 | 20.66 | 41.81 | 43.42 | 46.43 | 42.04 | 33.78 | 31.81 | 42.48 | 50.55 | 40.66 | + |
| Tfg | 81.99 | 84.42 | 86.56 | 66.05 | 69.87 | 71.75 | 59.21 | 73.68 | 66.85 | 58.82 | 70.01 | 60.57 | + |
| Psma3 | 425.86 | 436.49 | 443.07 | 242.34 | 261.99 | 265.77 | 218.89 | 279.35 | 251.06 | 203.55 | 218.85 | 200.28 | + |
| Cep57 | 27.6 | 31.31 | 26.17 | 25.33 | 25.11 | 22.64 | 20.99 | 22.23 | 18.4 | 22.25 | 24.48 | 22.61 | + |
| Eif2b3 | 63.05 | 62.88 | 62.24 | 32.48 | 32.57 | 34.57 | 37.86 | 59.47 | 51.91 | 34.14 | 33.45 | 35.05 | + |
| Lsm3 | 140.96 | 143.04 | 138.52 | 62.5 | 70.93 | 77.62 | 58.84 | 66.1 | 61.98 | 40.8 | 52.25 | 47.88 | + |
| Mecp2 | 1.68 | 1.91 | 1.87 | 2.65 | 2.73 | 2.96 | 3.04 | 2.88 | 2.56 | 2.63 | 3.29 | 2.73 | + |
| Rpl36 | 743.35 | 778.05 | 742.07 | 1245.59 | 1222.4 | 1279.73 | 1314.76 | 1173.22 | 1202.34 | 1333.81 | 1412.98 | 1367.65 | + |
| H2-Oa | 14.1 | 10.32 | 12.94 | 31.87 | 25.23 | 29.48 | 30.73 | 19.02 | 20.76 | 19.8 | 20.15 | 23.57 | + |
| Lman1 | 52.49 | 50.33 | 50.53 | 30.67 | 35.55 | 33.2 | 33.65 | 37.07 | 36.7 | 25.4 | 30.58 | 27.67 | + |
| Itpripl1 | 42.72 | 44.08 | 44.24 | 33.45 | 37.74 | 37.4 | 23.84 | 28.69 | 28.23 | 29.96 | 24.8 | 29.47 | + |
| Cdkn1b | 3.8 | 3.61 | 3.87 | 11.17 | 10.9 | 11.88 | 12.55 | 9.41 | 10.35 | 15.86 | 16.63 | 18.23 | + |
| Mrfap1 | 133.62 | 151.03 | 153.83 | 107.8 | 113.52 | 114.55 | 100.94 | 115.58 | 114.15 | 101.32 | 98.61 | 106.59 | + |
| Nme2 | 603.74 | 588.25 | 582.02 | 464.34 | 487.02 | 498.38 | 520.61 | 534.98 | 529.58 | 474.54 | 519.54 | 440.2 | + |
| Wasf2 | 12.46 | 11.84 | 11.88 | 17.46 | 17.74 | 17.39 | 19.06 | 16.86 | 18.27 | 21 | 19.57 | 19.54 | + |
| Fam173a | 48.99 | 46.69 | 45.63 | 62.43 | 66.5 | 64.41 | 62.7 | 62.68 | 64.62 | 65.77 | 65.48 | 63.47 | + |
| Secisbp2 | 10.07 | 10.02 | 9.84 | 7.07 | 6.24 | 6.41 | 7.77 | 7.96 | 6.87 | 7.38 | 6.95 | 8.23 | + |
| Mrpl9 | 76.04 | 76.28 | 74.92 | 77.04 | 68.65 | 75.25 | 89.27 | 73.87 | 72.38 | 93.61 | 97.82 | 105.17 | + |
| Ip6k1 | 7.91 | 7.69 | 8.62 | 32.83 | 28.91 | 32.06 | 33.85 | 24.28 | 27.72 | 42.87 | 53 | 45.62 | + |
| Cmip | 3.47 | 2.89 | 2.66 | 6.76 | 5.82 | 6.81 | 5.51 | 4.66 | 5.01 | 6.22 | 5.96 | 6.09 | + |
| Rpa3 | 93.69 | 95.15 | 95.51 | 42.99 | 46.76 | 52.48 | 37.56 | 45.11 | 45.91 | 33.53 | 37.77 | 35.92 | + |
| Srgap2 | 3.93 | 3.52 | 3.41 | 4.86 | 4.98 | 4.64 | 5.62 | 5.1 | 5.11 | 4.89 | 5.82 | 5.27 | + |
| Arhgdia | 280.23 | 291.04 | 285.08 | 194.68 | 192.56 | 190.85 | 182.13 | 201.31 | 192.1 | 168.94 | 159.5 | 171.1 | + |
| Doc2g | 1.76 | 1.67 | 1.16 | 3.75 | 3.26 | 4.06 | 3.19 | 3.32 | 3.46 | 4.42 | 4.72 | 4.08 | + |
| Afap1 | 0.68 | 0.27 | 0.69 | 1.17 | 1.25 | 1.2 | 1.23 | 1.53 | 1.62 | 1.08 | 0.82 | 1.42 | + |
| Ifitm3 | 212.61 | 231.22 | 176.74 | 75.9 | 79.75 | 71.55 | 86.11 | 74.65 | 73.09 | 66.45 | 57.98 | 81.54 | + |
| Bub3 | 140.24 | 118.82 | 118.31 | 89.23 | 109.5 | 98.84 | 82.41 | 92.92 | 89.68 | 83.07 | 82.68 | 83.31 | + |
| Zdhhc12 | 25.92 | 25.66 | 23.96 | 14.37 | 14.43 | 16.66 | 15.36 | 20.12 | 18.57 | 15.35 | 13.39 | 17.06 | + |
| Zfp768 | 0.69 | 0.67 | 0.67 | 1.58 | 1.66 | 1.53 | 1.04 | 1.46 | 2.05 | 1.17 | 2.22 | 1.46 | + |
| Rbbp7 | 243.89 | 245.86 | 244.88 | 133.63 | 160.83 | 157.87 | 117.69 | 140.29 | 127.68 | 100.96 | 110.4 | 103.82 | + |
| Bud31 | 86.2 | 83.97 | 75.12 | 37.09 | 38.27 | 45.13 | 40.06 | 46.55 | 42.98 | 36.73 | 37.25 | 38.56 | + |
| Ptprv | 1.35 | 1.35 | 1.56 | 1.98 | 2.01 | 1.96 | 3.13 | 2.44 | 2.56 | 2.53 | 2.89 | 2.27 | + |
| Zmat2 | 39.05 | 35.09 | 35.43 | 26.63 | 27.96 | 26.39 | 25.6 | 26.36 | 27.55 | 25.55 | 25.79 | 27.23 | + |
| Serpinf1 | 2.1 | 1.95 | 3.15 | 6.64 | 8.31 | 8.11 | 49.15 | 55.03 | 51.39 | 34.5 | 41.75 | 35.03 | + |
| Rhot2 | 12.54 | 13.7 | 12.9 | 6.69 | 5.91 | 6.61 | 6.06 | 6.1 | 5.04 | 5.46 | 5.42 | 4.33 | + |
| Cirh1a | 88.52 | 89.04 | 88.92 | 43.23 | 45.6 | 46.42 | 39.68 | 47.54 | 40.66 | 35.38 | 34.06 | 35.84 | + |
| 2210408F21Rik | 1.25 | 0.88 | 1.11 | 7.28 | 5.56 | 6.12 | 3.15 | 1.37 | 2.57 | 4.4 | 4.56 | 5.45 | + |
| Hmg20a | 5.33 | 4.52 | 4.63 | 9.4 | 8.92 | 9.87 | 8.94 | 7.73 | 8.73 | 8.82 | 10.77 | 9.43 | + |
| Hivep3 | 1.53 | 1.6 | 1.77 | 1.13 | 0.95 | 0.98 | 0.97 | 1.13 | 1 | 0.82 | 1.07 | 0.81 | + |
| Mov10 | 20.66 | 22.09 | 19.92 | 31.49 | 31.42 | 31.62 | 12.93 | 15.34 | 15.85 | 8.5 | 7.53 | 7.54 | + |
| Psmd6 | 186.62 | 185.24 | 180.05 | 69.88 | 80.97 | 75.24 | 70.46 | 86.15 | 79.6 | 54.73 | 52.31 | 63.2 | + |
| Nubp2 | 118.25 | 115.05 | 113.07 | 56.12 | 60.68 | 58.38 | 47.48 | 60.54 | 59.78 | 55.81 | 51.38 | 48.74 | + |
| Cnbd2 | 0.14 | 0.19 | 0.05 | 0.63 | 0.74 | 0.61 | 0.85 | 0.56 | 0.84 | 1.03 | 0.67 | 0.41 | + |
| Snrnp70 | 56.19 | 60.25 | 63.47 | 39.19 | 40.16 | 42.92 | 43.17 | 46.66 | 44.81 | 34.76 | 40.99 | 38.55 | + |
| Oas1a | 6.35 | 6.86 | 5.15 | 3.18 | 5.06 | 4.81 | 1.55 | 1.34 | 0.97 | 2.39 | 1.14 | 1.34 | + |
| Slc25a26 | 4.7 | 4.42 | 5.77 | 2.27 | 2.05 | 2.07 | 3.74 | 2.89 | 3.15 | 2.53 | 1.86 | 1.89 | + |
| Gnpda2 | 20.84 | 21.46 | 25.56 | 33.46 | 40.31 | 39.8 | 24.01 | 28.34 | 28.66 | 26.77 | 30.25 | 31.61 | + |
| Ptges3 | 176.44 | 203.02 | 216.89 | 111.37 | 125.9 | 121.15 | 94.07 | 119.29 | 122.59 | 87.48 | 92.99 | 90.5 | + |
| Mbtps1 | 16.29 | 16.3 | 15.7 | 14.21 | 13.34 | 11.26 | 11.37 | 10.79 | 10.59 | 12.43 | 10.23 | 12.12 | + |
| Tmem69 | 23.94 | 25.56 | 24.12 | 9.33 | 7.74 | 8.37 | 9.89 | 10.02 | 9.11 | 8.32 | 9.56 | 9.12 | + |
| Ndfip2 | 11.03 | 12.01 | 12.7 | 14.57 | 13.91 | 15.24 | 30.38 | 24.98 | 27.6 | 20.21 | 17.05 | 22.44 | + |
| Acvrl1 | 0.38 | 0.22 | 0.25 | 2.7 | 1.46 | 1.51 | 7.05 | 1.43 | 4.4 | 3.49 | 4.66 | 6.28 | + |
| Timm13 | 120.93 | 120.6 | 124.42 | 68.41 | 75.57 | 74.65 | 68.21 | 86.11 | 81.9 | 55.49 | 56.85 | 58.63 | + |
| Sec23a | 11.58 | 11.97 | 12.34 | 18.22 | 18.08 | 18.15 | 17.04 | 21.02 | 20.81 | 16.08 | 18.89 | 17.86 | + |
| Polr2d | 47.83 | 50.49 | 47.93 | 31.47 | 28.86 | 27.49 | 30.79 | 30.5 | 29.93 | 28.88 | 33.89 | 32.28 | + |
| Ints7 | 17.11 | 17.69 | 15.66 | 12.46 | 9.58 | 9.98 | 9.87 | 9.79 | 9.44 | 9.37 | 7.9 | 9.2 | + |
| Snx3 | 186.85 | 190.28 | 198.43 | 165.52 | 161.2 | 170.76 | 151.35 | 164.19 | 161.57 | 142.42 | 151.5 | 144.22 | + |
| Arhgap33 | 0.95 | 0.95 | 1.05 | 1.98 | 1.7 | 1.7 | 2.04 | 1.43 | 1.45 | 1.5 | 2.03 | 1.72 | + |
| Cpped1 | 9.24 | 7.76 | 8.81 | 4.79 | 4.31 | 4.55 | 4.81 | 5.96 | 5.52 | 3.89 | 4.05 | 5.08 | + |
| Ccdc115 | 34.56 | 34.65 | 32.91 | 24.84 | 21.89 | 26.25 | 28.75 | 24.87 | 26.62 | 22.55 | 26.1 | 27.39 | + |
| Psmd13 | 151.87 | 148.8 | 145.04 | 74.36 | 70.05 | 79.57 | 71.94 | 83.87 | 75.48 | 64.61 | 64.82 | 73.06 | + |
| Smpd5 | 3.54 | 4.47 | 3.03 | 6.8 | 7.5 | 7.96 | 7.82 | 7.2 | 5.56 | 6.37 | 5.41 | 7.99 | + |
| Lcp2 | 74.08 | 78.31 | 71.87 | 42.75 | 40.16 | 38.76 | 48.54 | 46.54 | 52.23 | 38.77 | 30.78 | 40.4 | + |
| Trmt2a | 29.84 | 32.23 | 30.2 | 19.17 | 18.93 | 18.12 | 11.2 | 16.2 | 14.01 | 13.19 | 13.02 | 12.99 | + |
| Stard4 | 17.6 | 17.1 | 17.73 | 10.95 | 12.42 | 12.73 | 12.39 | 14.5 | 13.25 | 10.47 | 10.61 | 10.66 | + |
| Gnl3l | 18.07 | 17.89 | 18.5 | 12.08 | 11.56 | 11.82 | 11.39 | 14.44 | 13.2 | 10.67 | 10.07 | 9.16 | + |
| Arfgap2 | 36.43 | 34.07 | 36.67 | 30.67 | 27.87 | 30.27 | 28.84 | 27.2 | 27.06 | 31.24 | 26.31 | 29.19 | + |
| Fanci | 5.58 | 5.4 | 7.09 | 3.75 | 4.03 | 4.71 | 2.23 | 2.34 | 2.55 | 2.5 | 2.49 | 2.36 | + |
| Gipr | 0.1 | 0.06 | 0.18 | 2.12 | 1.36 | 1.71 | 7.66 | 1.15 | 3.98 | 3.86 | 3.82 | 5.05 | + |
| Tssc4 | 22.05 | 23.32 | 22.98 | 17.68 | 15.08 | 15.69 | 12.49 | 16.04 | 11.9 | 15.92 | 12.44 | 15.53 | + |
| Hs3st3b1 | 1.13 | 1.33 | 1.26 | 0.56 | 0.55 | 0.53 | 0.79 | 0.79 | 1.16 | 0.63 | 0.73 | 0.85 | + |
| 5-Mar | 10.04 | 9.68 | 9.86 | 5.1 | 6.87 | 6.81 | 6.51 | 8.34 | 6.93 | 5.85 | 7.6 | 5.86 | + |
| Lmna | 17.32 | 16.48 | 18.04 | 9.82 | 15.79 | 14.41 | 5.08 | 8.87 | 8 | 11.58 | 9.81 | 8.84 | + |
| A730043L09Rik | 0.09 | 0.2 | 0.18 | 0.44 | 0.41 | 1.27 | 2.86 | 1.22 | 1.74 | 1.52 | 2.05 | 1.72 | + |
| Chmp2b | 29.13 | 27.73 | 30.18 | 41.48 | 37.36 | 39.65 | 40.99 | 38.18 | 33.13 | 38.53 | 43.56 | 40.07 | + |
| Nxt1 | 72.51 | 76.59 | 73.67 | 52.78 | 62.54 | 64.55 | 51.99 | 53.38 | 54.47 | 55.27 | 51.53 | 49.49 | + |
| Timm50 | 104.49 | 104.29 | 109.01 | 55.06 | 57.7 | 58.2 | 46.35 | 60.68 | 55.98 | 41.97 | 43.7 | 38.77 | + |
| St6gal1 | 2.54 | 2.07 | 1.56 | 6.19 | 3.43 | 4.65 | 1.02 | 0.33 | 0.49 | 0.65 | 0.4 | 0.43 | + |
| Orc6 | 69.48 | 80.22 | 73.85 | 29.47 | 31.72 | 33.81 | 21.31 | 31.23 | 28.77 | 18.06 | 19.53 | 18.72 | + |
| Madd | 6.96 | 7.68 | 6.64 | 9.01 | 8.72 | 8.3 | 13.2 | 10.78 | 10.57 | 9.95 | 8.33 | 9.84 | + |
| Spata6 | 13.71 | 10.57 | 12.37 | 23.25 | 21.51 | 23.25 | 28.05 | 19.92 | 20.53 | 20.41 | 20.55 | 23.46 | + |
| Zfc3h1 | 4.13 | 4.11 | 4.2 | 6.56 | 7.4 | 7.41 | 6.6 | 7.1 | 6.4 | 8.46 | 8.42 | 7.59 | + |
| Selenbp1 | 6.33 | 5.48 | 4.65 | 21.38 | 22.91 | 21 | 91.51 | 25.77 | 49.59 | 55.58 | 65.13 | 78.16 | + |
| Zfp706 | 40.55 | 44.8 | 46.1 | 33.05 | 31.13 | 31.88 | 33.74 | 33.64 | 35.21 | 34.24 | 33.35 | 33.25 | + |

| | | | | | | | | | | | | |
|---|---|---|---|---|---|---|---|---|---|---|---|---|
| Rbm10 | 39.3 | 41.09 | 38.62 | 26.6 | 28.24 | 28.02 | 24.66 | 27.89 | 23.68 | 24.87 | 24.15 | 25.24 | + |
| Pwwp2b | 0.91 | 1.08 | 0.8 | 2.13 | 2.7 | 2.54 | 2.55 | 2.78 | 2.11 | 1.8 | 1.75 | 1.88 | + |
| Pacs1 | 2.68 | 2.98 | 3.32 | 4.77 | 5.49 | 5.98 | 8.38 | 5.95 | 6.89 | 7.85 | 8.53 | 7.63 | + |
| Fam107b | 161.43 | 165.98 | 169.18 | 158.63 | 154.32 | 143.66 | 193.84 | 177.92 | 178.64 | 218.07 | 212.76 | 235.01 | + |
| Tsta3 | 62.01 | 61.91 | 59.63 | 49.39 | 49.39 | 49.43 | 50.45 | 50.75 | 51.16 | 43.21 | 46.21 | 49.38 | + |
| Rps12 | 1605.23 | 1610.48 | 1640.84 | 2201.39 | 2166.69 | 2255.88 | 2104.88 | 2071.1 | 2052.29 | 2310.46 | 2520.16 | 2308.35 | + |
| Ppil3 | 90.94 | 78.23 | 76.03 | 60.98 | 61.54 | 59.49 | 56.97 | 60.66 | 54.4 | 56.46 | 49.97 | 56.86 | + |
| Med13l | 2.79 | 2.82 | 3.24 | 4.87 | 4.23 | 4.71 | 5.14 | 5.38 | 5.82 | 4.75 | 4.65 | 4.74 | + |
| Calm1 | 241.21 | 255.47 | 241.85 | 121.17 | 123.67 | 119.36 | 119.78 | 127.82 | 122.35 | 105.99 | 90.39 | 108.48 | + |
| Ddx21 | 72.66 | 73.36 | 78.51 | 40.65 | 41.94 | 40.63 | 34.44 | 47.68 | 46.54 | 35.29 | 33.2 | 35.03 | + |
| Pafah1b3 | 26.64 | 26.89 | 27.03 | 52.41 | 44.97 | 46.47 | 63.81 | 52.46 | 50.88 | 51.16 | 49.52 | 54.52 | + |
| Cd48 | 150.17 | 165.34 | 148.86 | 51.8 | 54.22 | 49.92 | 52.89 | 52.15 | 51.96 | 51.09 | 37.88 | 44.4 | + |
| Ppa1 | 217.88 | 222.62 | 234.47 | 81.69 | 92.47 | 88.85 | 64.45 | 95.09 | 83.55 | 51.36 | 56.29 | 57.95 | + |
| Ttc17 | 2.11 | 2.47 | 2.11 | 5.08 | 5.43 | 4.5 | 4.14 | 4.55 | 3.69 | 5.55 | 6.09 | 6.97 | + |
| Xrcc1 | 29.01 | 28.73 | 27.37 | 26.13 | 26.38 | 27.7 | 19.65 | 18.56 | 19.92 | 17.92 | 21.07 | 18.59 | + |
| Uqcrfs1 | 203.4 | 206.12 | 206.8 | 122.76 | 112.7 | 111.09 | 139.5 | 132.22 | 138.24 | 117.12 | 111.53 | 116.52 | + |
| Psmc2 | 103.5 | 108.92 | 103.68 | 58.54 | 64.67 | 65.62 | 53.4 | 66.55 | 61.24 | 44.86 | 51.74 | 47 | + |
| Dhx29 | 12.38 | 13.92 | 12.98 | 9.56 | 10.08 | 9.07 | 8.68 | 10.75 | 9.83 | 8.18 | 9.82 | 8.57 | + |
| Srek1 | 13.56 | 14.85 | 15.13 | 10.9 | 11.1 | 10.57 | 8.69 | 10.34 | 10.29 | 8.46 | 10.33 | 8.34 | + |
| Vill | 3.18 | 2.59 | 1.72 | 10.01 | 9.12 | 8.74 | 8.74 | 6.94 | 7.29 | 7.58 | 7.54 | 11.73 | + |
| Tpra1 | 9.83 | 11.26 | 10.25 | 21.67 | 20.27 | 19.94 | 22.06 | 21.74 | 19.45 | 25.79 | 21.7 | 29.91 | + |
| Gm12504 | 3.4 | 3.21 | 2.61 | 1.8 | 1.91 | 1.51 | 1.72 | 1.88 | 1.88 | 1.41 | 1.62 | 1.03 | + |
| Tcta | 7.08 | 6.5 | 6.47 | 11.95 | 9.89 | 11.59 | 12.42 | 17.13 | 13.4 | 14.36 | 14.79 | 13.34 | + |
| Slc20a1 | 33.11 | 39.06 | 31.61 | 22.08 | 20.06 | 19.31 | 24.6 | 22.08 | 23.34 | 21.26 | 20.68 | 19.5 | + |
| 2500004C02Rik | 5.64 | 6.13 | 5.52 | 3.2 | 2.62 | 3.59 | 3.04 | 4.13 | 3.59 | 3.44 | 3.34 | 3.36 | + |
| Fasl | 35.52 | 39.06 | 38.69 | 25.71 | 19.79 | 23.05 | 14.41 | 18.09 | 19.81 | 17.32 | 12.45 | 15.25 | + |
| Ptcd3 | 56.53 | 59.22 | 56.26 | 21.97 | 25.83 | 24.51 | 17.22 | 24.79 | 21.42 | 16.61 | 15.69 | 15.53 | + |
| B4galt1 | 18.03 | 18.61 | 15.42 | 38.03 | 35.8 | 37.53 | 23.19 | 19 | 20.12 | 32.86 | 32.45 | 32.84 | + |
| Ddx50 | 34.65 | 33.28 | 32.91 | 65.69 | 66.84 | 71.2 | 72.21 | 70.89 | 71.54 | 79.41 | 93.85 | 76.6 | + |
| B4galnt1 | 96.77 | 106.25 | 90.86 | 133.13 | 118.8 | 131.39 | 103.99 | 82.7 | 90.92 | 157.12 | 127.76 | 173.49 | + |
| Larp4 | 6.86 | 7.91 | 7.2 | 4.29 | 3.93 | 4.99 | 2.92 | 3.95 | 3.91 | 3.03 | 3.08 | 2.89 | + |
| Skiv2l2 | 51.43 | 51.35 | 49.9 | 30 | 32.9 | 31.97 | 25.6 | 30.58 | 29.23 | 24.63 | 22.59 | 21.79 | + |
| Dpagt1 | 27.96 | 24.75 | 26.97 | 18.25 | 18.97 | 19.32 | 19.19 | 19.08 | 16.22 | 18.92 | 14.95 | 21 | + |
| Eif2b4 | 40.35 | 42.23 | 41.11 | 25.05 | 24.39 | 23.43 | 23.65 | 24.56 | 22.27 | 23.77 | 22.61 | 26.99 | + |
| Ccnb1 | 148.47 | 112.51 | 109.35 | 74.5 | 100.98 | 85.91 | 42.88 | 46.21 | 42.58 | 51.26 | 48.48 | 37.08 | + |
| Aco1 | 12.22 | 12.68 | 13.37 | 8.71 | 10.1 | 9.73 | 8.66 | 9.95 | 9.65 | 8.8 | 10.49 | 9.53 | + |
| Hspa4 | 115.53 | 123.08 | 125 | 72.81 | 76.51 | 79.75 | 68.93 | 84.78 | 79.96 | 62.72 | 72.69 | 63.49 | + |
| Nek8 | 4.95 | 4.84 | 4.73 | 11.06 | 12.06 | 9.09 | 5.93 | 5.6 | 6.61 | 6.52 | 7.46 | 7.82 | + |
| Top1 | 30.71 | 30.1 | 30.5 | 16.4 | 17.65 | 16.58 | 13.59 | 17.86 | 13.75 | 14.29 | 14.77 | 13.35 | + |
| Dgat1 | 29.76 | 35.2 | 41.65 | 69.3 | 66.35 | 62.68 | 99.95 | 97.09 | 87.18 | 109.07 | 89.75 | 107.91 | + |
| Amica1 | 7.71 | 11.24 | 5.7 | 80.91 | 58.94 | 61.63 | 7.46 | 4.56 | 6.31 | 28.29 | 25.17 | 25.95 | + |
| Fam133b | 15.5 | 17.82 | 16.04 | 13.37 | 11.64 | 12.44 | 10.68 | 10.36 | 10.55 | 10.56 | 10.55 | 10.37 | + |
| Efcab4b | 1.04 | 1.76 | 1.58 | 1.96 | 1.08 | 1.87 | 2.26 | 1.83 | 1.32 | 3.95 | 2.82 | 3.58 | + |
| Ash2l | 6.8 | 6.41 | 6.42 | 4.47 | 4.53 | 4.19 | 4.82 | 5.35 | 4.23 | 3.59 | 4.34 | 4.15 | + |
| Pgls | 121.16 | 120.28 | 122.28 | 139.15 | 145.13 | 143.78 | 156.23 | 152.19 | 149.89 | 140.52 | 144.92 | 156.74 | + |
| Srsf2 | 63.16 | 71.7 | 73.29 | 83.23 | 76.05 | 77.43 | 91.28 | 71.8 | 81.02 | 112.76 | 109.53 | 105.8 | + |
| Ndufaf4 | 15.14 | 15.2 | 16.58 | 7.6 | 7.25 | 7.34 | 4.68 | 7.94 | 8.97 | 5.32 | 5.53 | 5.22 | + |
| Timm22 | 11.41 | 12.04 | 11.35 | 8.53 | 8.16 | 7.74 | 8.26 | 9.32 | 8.1 | 7.29 | 6.6 | 7.72 | + |
| Rhov | 0.1 | 0.28 | 0.39 | 1.34 | 1.24 | 1.04 | 0.83 | 0.96 | 0.65 | 1.35 | 1.08 | 1.77 | + |
| Myadm | 4.4 | 3.82 | 4.24 | 7.92 | 6.01 | 7.84 | 11.05 | 7.42 | 9.21 | 10.58 | 7.1 | 8.93 | + |
| Xpo5 | 18.16 | 17.79 | 17.76 | 9.3 | 9.48 | 7.48 | 5.19 | 8.3 | 7.59 | 5.52 | 4.89 | 5.98 | + |
| Ung | 16.8 | 22.57 | 23.84 | 8.58 | 7.81 | 8.16 | 7.65 | 9.57 | 7.78 | 5.47 | 5.92 | 5.29 | + |
| Churc1 | 363.31 | 398.8 | 368.58 | 223.57 | 246.03 | 206.51 | 257.14 | 268.17 | 235.16 | 224.33 | 241.37 | 235.4 | + |
| Ciao1 | 13.7 | 12.77 | 12.73 | 10.26 | 10.9 | 10.93 | 10.07 | 9.66 | 10.13 | 8.28 | 10.12 | 11.43 | + |
| Zcchc10 | 29.61 | 27.26 | 27.96 | 9.41 | 10.53 | 9.79 | 10.65 | 13.69 | 12.59 | 7.28 | 7.06 | 6.14 | + |
| Ap3m2 | 1.98 | 2.31 | 2.16 | 4.58 | 3.98 | 4.46 | 4.59 | 3.57 | 3.28 | 4.57 | 5.83 | 4.25 | + |
| Tmem48 | 18.42 | 19.86 | 18.89 | 10.67 | 9.8 | 10.67 | 6.45 | 8.05 | 6.44 | 4.71 | 4.74 | 4.03 | + |
| Pdlim1 | 47.53 | 55.56 | 54.36 | 58.5 | 52.84 | 55.88 | 45.83 | 44.2 | 48.2 | 33.9 | 37.97 | 39.3 | + |
| Csnk1d | 28.95 | 27.13 | 29.21 | 48.5 | 44.91 | 47.52 | 48.13 | 42.5 | 43.42 | 53.89 | 60.93 | 56.72 | + |
| Pum2 | 10.99 | 10.87 | 11.55 | 9.91 | 9.21 | 8.45 | 7.77 | 8.64 | 8.61 | 9.77 | 8.26 | 8.01 | + |
| A130077B15Rik | 76.65 | 77.79 | 81.33 | 131.47 | 121.26 | 118.75 | 114.31 | 113.07 | 100.48 | 141.98 | 138.07 | 133.73 | + |
| 3110001I22Rik | 6.13 | 4.41 | 4.63 | 3.14 | 2.87 | 2.1 | 1.96 | 1.82 | 2.03 | 2.56 | 1.8 | 2.29 | + |
| Ldha | 3162.33 | 3160.43 | 3208.14 | 1753.16 | 1709.44 | 1732.71 | 2412.21 | 2116.46 | 2283.17 | 1678.03 | 1757.39 | 1945.95 | + |
| Eif4a1 | 340.8 | 381.33 | 412.98 | 194.91 | 201.89 | 201.95 | 209.59 | 224.95 | 215.51 | 182.47 | 179.19 | 198.25 | + |
| Rpl32 | 1892.9 | 1999.02 | 1959.36 | 3266.2 | 3167.95 | 3356.27 | 3395.41 | 3029.36 | 3181.04 | 3530.67 | 3905.65 | 3590.58 | + |
| Cenpt | 18.16 | 16.99 | 17.17 | 16.36 | 20.45 | 18.88 | 10.99 | 15.36 | 13.74 | 8.97 | 8.81 | 9.59 | + |
| Nipal3 | 1.12 | 1.11 | 1.07 | 3.04 | 2.66 | 3.48 | 4.45 | 3.44 | 4.01 | 4.35 | 3.57 | 3.5 | + |
| Unc93b1 | 28.09 | 30.96 | 26.37 | 42.23 | 37.41 | 40.44 | 42.81 | 40.07 | 36.36 | 35.95 | 32.14 | 41.69 | + |
| Ctns | 1.69 | 1.91 | 1.6 | 3.44 | 2.61 | 2.39 | 4.75 | 3.88 | 3.71 | 3.67 | 3.52 | 4.33 | + |
| Sec61a2 | 9.13 | 7.59 | 8.7 | 5.36 | 6.06 | 5.6 | 4.07 | 5.6 | 4.8 | 4.56 | 5.52 | 3.6 | + |
| Mrpl50 | 36.08 | 32.81 | 36.47 | 19.63 | 21.33 | 22.71 | 17.93 | 21 | 21.49 | 19.31 | 20.72 | 16.94 | + |
| Evi5l | 3.52 | 1.85 | 2.81 | 5.04 | 4.98 | 5 | 8.53 | 6.25 | 8.21 | 4.3 | 6.31 | 6.09 | + |
| Hilpda | 77.84 | 69.54 | 75.7 | 85.98 | 114.83 | 96.92 | 133.26 | 127.31 | 131.1 | 80.44 | 109.18 | 122.95 | + |
| C330018D20Rik | 8.71 | 7.55 | 7.64 | 4.11 | 5.2 | 5.37 | 3.67 | 4.67 | 4.35 | 3.84 | 3.73 | 2.91 | + |
| Gm11110 | 7.34 | 7.54 | 6.48 | 24.21 | 21.96 | 20.51 | 28.23 | 17.69 | 20.08 | 31.81 | 30.64 | 36.07 | + |
| Pitpnb | 66.32 | 67.21 | 71.36 | 39.81 | 42.3 | 39.2 | 33.74 | 39.21 | 39.11 | 30.7 | 33.31 | 33.1 | + |
| Wdr12 | 53.08 | 48.73 | 52.33 | 25.53 | 29.54 | 30.91 | 19.76 | 26.07 | 26.29 | 24.93 | 22.17 | 23.46 | + |
| Elac2 | 20.96 | 25.15 | 32.54 | 14.47 | 12.3 | 13.19 | 12.3 | 16.6 | 16.26 | 10.67 | 12.59 | 10.43 | + |
| Mbp | 5.95 | 5.57 | 5.92 | 8.76 | 7.71 | 7.41 | 8.13 | 6.88 | 7.19 | 7.59 | 8.24 | 7.34 | + |
| Galnt1 | 7.03 | 8.49 | 8.37 | 4.22 | 4.04 | 4.22 | 4.24 | 4.54 | 4.22 | 3.92 | 3.11 | 3.63 | + |
| Mrm1 | 8.48 | 8.86 | 9.5 | 16.89 | 16.58 | 14.03 | 14.38 | 13.33 | 15.8 | 18.5 | 17.79 | 16.57 | + |
| Cyb5b | 67.93 | 66.86 | 67.13 | 36.38 | 39.61 | 42.44 | 32.93 | 44.3 | 41.58 | 32.15 | 28.3 | 28.68 | + |
| Ddx41 | 43.18 | 45.55 | 42.62 | 30.67 | 29.24 | 29.7 | 32.34 | 31.8 | 30.56 | 29.09 | 27.14 | 27.79 | + |
| Gfpt2 | 1.49 | 1.07 | 1.67 | 0.95 | 1.42 | 1.6 | 1.08 | 1.93 | 1.89 | 0.61 | 0.5 | 0.64 | + |
| Fam134a | 3.84 | 3.89 | 3.78 | 6.87 | 5.37 | 7.66 | 6.97 | 7.05 | 5.75 | 6.89 | 7.47 | 8.36 | + |
| Ndufv3 | 138.51 | 142.91 | 138.92 | 247.27 | 217.31 | 240.94 | 336.83 | 240.88 | 284.66 | 259.88 | 289.29 | 287.57 | + |
| Ppp1r7 | 19.91 | 20.44 | 21.19 | 16.97 | 15.58 | 15.59 | 13.74 | 13.9 | 13.35 | 12.93 | 11.88 | 12.18 | + |
| Gm15772 | 1681.28 | 1652.58 | 1693.55 | 2642.1 | 2579.89 | 2763.84 | 2542.48 | 2500.79 | 2516.72 | 2745.29 | 2999.35 | 2722.37 | + |
| 1810022K09Rik | 239.32 | 220.94 | 219.41 | 154.31 | 150.01 | 150.76 | 139.73 | 142.66 | 146.64 | 127.73 | 124.49 | 125.88 | + |
| H3f3b | 364.13 | 386.87 | 354.71 | 221.46 | 239.12 | 237.37 | 199.18 | 199.8 | 206.21 | 204.89 | 187.91 | 190.84 | + |
| Rps23 | 2057.17 | 2083.08 | 2079.68 | 3102.04 | 3147.49 | 3300.27 | 3122.9 | 3001.88 | 3109.2 | 3340.78 | 3650.22 | 3434.3 | + |
| Bend4 | 6.23 | 5.89 | 6.19 | 9.12 | 10.86 | 10.41 | 9.63 | 11.7 | 11.21 | 12.14 | 11.27 | 12.24 | + |
| Naprt1 | 1.75 | 1.85 | 1.57 | 6.55 | 8.88 | 7.15 | 4.77 | 4.94 | 4.88 | 8.81 | 8.28 | 8.69 | + |
| Cdkn2d | 72.64 | 65.89 | 65.65 | 124.47 | 111.29 | 123.34 | 132.31 | 110.86 | 119 | 113.82 | 133.89 | 122.68 | + |
| Tyr | 10.02 | 10.57 | 11.88 | 5.04 | 5.69 | 5.31 | 5.78 | 6.05 | 5.66 | 6.32 | 5.9 | 4.87 | + |
| Wdr43 | 96.56 | 101.83 | 101.02 | 52.92 | 59.28 | 57.54 | 43.21 | 63.7 | 58.33 | 44.56 | 42.66 | 42.9 | + |
| Rps10 | 1505.83 | 1489.93 | 1472.11 | 2045.43 | 2058.96 | 2123.27 | 2041.14 | 1971.18 | 2003.98 | 2143.16 | 2329.83 | 2158.65 | + |
| Dctpp1 | 92.28 | 85.48 | 79.6 | 41.27 | 47.49 | 50.22 | 49.8 | 62.71 | 51.78 | 36.9 | 41.9 | 40.38 | + |
| Fam111a | 20.56 | 21.12 | 19.86 | 9.92 | 10.14 | 10.09 | 7.56 | 7.4 | 8.34 | 8.1 | 5.57 | 7.5 | + |
| Slc35e4 | 3.42 | 2.72 | 3.38 | 1.69 | 1.61 | 1.42 | 2.18 | 2.67 | 1.44 | 2.08 | 1.53 | 1.58 | + |
| Sfxn3 | 7.62 | 7.07 | 6.83 | 20.54 | 17.7 | 20.63 | 28.25 | 21.66 | 24.64 | 26.64 | 26.38 | 30.98 | + |
| Haus1 | 28.54 | 29.91 | 29.73 | 19.33 | 19.52 | 22.8 | 18.09 | 17.67 | 16.33 | 14.59 | 13.5 | 16.62 | + |
| Rsl1d1 | 220.89 | 232.48 | 231.44 | 127.45 | 130.61 | 135.52 | 118.25 | 141.92 | 139.33 | 110.76 | 118.75 | 113.19 | + |
| Ranbp9 | 4.03 | 4.66 | 4.24 | 6.97 | 7.21 | 7.53 | 6.42 | 7.48 | 5.91 | 6.71 | 7.29 | 6.94 | + |
| Ndufb9 | 238.09 | 238.93 | 244.78 | 183.3 | 203 | 210.71 | 205.85 | 208.79 | 200.76 | 175.4 | 182.71 | 190.2 | + |
| Ltb4r1 | 34.86 | 36.45 | 30.1 | 56.08 | 50.45 | 47.22 | 116.17 | 83.29 | 95.36 | 74.1 | 60.17 | 79.36 | + |
| Nde1 | 49.08 | 49.71 | 43.35 | 36.43 | 44.31 | 43.95 | 33.94 | 34.14 | 30.81 | 35.61 | 31.82 | 35.3 | + |
| Foxk2 | 8.37 | 8.13 | 8.59 | 5.46 | 6.5 | 6.53 | 4.67 | 6.56 | 5.18 | 4.98 | 5.1 | 4.48 | + |
| Bpnt1 | 12.08 | 9.44 | 9.51 | 5.96 | 6.2 | 5.17 | 6.66 | 5.71 | 6.74 | 5.87 | 5.39 | 4.53 | + |
| E2f4 | 18.19 | 21.34 | 20.65 | 16.3 | 14.41 | 15.2 | 14.73 | 15.44 | 14.27 | 14.5 | 15.34 | 15.17 | + |
| Slc23a2 | 3.04 | 2.76 | 2.47 | 2.5 | 2.31 | 2.37 | 1.95 | 1.86 | 1.66 | 1.7 | 2.04 | 1.73 | + |
| Btaf1 | 2.5 | 2.64 | 2.93 | 1.73 | 1.72 | 1.62 | 1.63 | 1.83 | 1.83 | 1.43 | 1.78 | 1.61 | + |
| Cpne1 | 20.36 | 19.95 | 17.23 | 27.78 | 28.3 | 25.35 | 28.97 | 28.93 | 30.46 | 31.75 | 31.32 | 32.98 | + |
| Dguok | 23.31 | 23.29 | 21.26 | 48.5 | 50.53 | 48.18 | 50.32 | 47.03 | 49.59 | 51.13 | 48.92 | 52.87 | + |
| Med6 | 23.09 | 24.36 | 24.07 | 17.62 | 21.01 | 18.04 | 15.15 | 17.37 | 16.56 | 15.67 | 17.62 | 13.89 | + |
| Vps36 | 66.21 | 61.06 | 59.57 | 52.92 | 50.15 | 53.19 | 49.72 | 52.71 | 52.68 | 43.36 | 48.41 | 45.77 | + |
| Zfp317 | 4.86 | 5.03 | 5.36 | 9.92 | 9.63 | 9.06 | 8.32 | 8.3 | 8.05 | 11.49 | 10.89 | 10.06 | + |
| Gyg | 61.68 | 66.06 | 63.75 | 124.48 | 104.74 | 112.02 | 168.56 | 126.35 | 148.47 | 172.31 | 180.86 | 178.07 | + |
| Arhgef10 | 0.93 | 0.89 | 0.63 | 7.02 | 6.79 | 8.04 | 1.53 | 1.71 | 1.87 | 2.91 | 3.46 | 2.86 | + |
| Pycr2 | 111.86 | 93.7 | 102.84 | 54.75 | 53.6 | 55.4 | 57.83 | 66.45 | 67.85 | 53.38 | 43.13 | 48.49 | + |
| Ivns1abp | 81.47 | 79.95 | 84.08 | 67.64 | 66.67 | 72 | 60.29 | 64.75 | 64.55 | 72.1 | 71.95 | 62.15 | + |
| Tmem91 | 0.91 | 1 | 0.99 | 2.24 | 1.57 | 2.62 | 5.97 | 4.36 | 4.47 | 2.87 | 5.21 | 4.28 | + |
| Osgin2 | 25.91 | 23.01 | 24.6 | 13.12 | 16.55 | 13.81 | 10.46 | 16.13 | 15.63 | 12.34 | 11.87 | 10.63 | + |
| Ncoa1 | 1.49 | 1.78 | 1.6 | 2.53 | 2.87 | 2.38 | 2.73 | 2.81 | 2.39 | 2.94 | 2.96 | 3.04 | + |
| Cks1b | 218.1 | 218.6 | 214.07 | 114.71 | 141.7 | 129.32 | 94.07 | 106.56 | 91.32 | 65.78 | 73.92 | 63.53 | + |
| Neat1 | 27.09 | 28.5 | 33.84 | 44.77 | 39.64 | 42.37 | 79.57 | 63.6 | 63.24 | 58.08 | 49.97 | 61.51 | + |
| Sirt7 | 48.77 | 45.43 | 46.41 | 45.59 | 42.48 | 44.43 | 52.71 | 47.6 | 46.93 | 57.92 | 52.97 | 55.49 | + |
| Wdr5 | 40.12 | 41.97 | 37.06 | 24.69 | 28.93 | 27.66 | 22.71 | 26.41 | 25.77 | 25.78 | 25.84 | 23.69 | + |
| Lonp1 | 25.43 | 24.43 | 27.94 | 35.66 | 40.42 | 37.9 | 41.78 | 49.62 | 48.77 | 35.75 | 34.55 | 30.54 | + |
| Srp54c | 83.3 | 78.09 | 81.25 | 56.29 | 58.37 | 61.64 | 49.55 | 58.22 | 50.44 | 46.39 | 43.27 | 46.95 | + |
| Ttk | 9.74 | 7.83 | 8.27 | 4.11 | 7.14 | 5.14 | 3.59 | 3.31 | 3.78 | 3.76 | 3.98 | 2.68 | + |
| Cdh23 | 0.21 | 0.18 | 0.22 | 0.09 | 0.03 | 0.05 | 0.13 | 0.15 | 0.12 | 0.04 | 0.08 | 0.09 | + |
| Ccdc82 | 39.82 | 39.09 | 44.8 | 35.61 | 35.43 | 33.06 | 29.58 | 30.42 | 30.96 | 35.64 | 35.52 | 34.77 | + |
| Arhgap18 | 8.84 | 8.28 | 6.92 | 11.03 | 11.09 | 12.82 | 19.02 | 15.4 | 15.65 | 12.21 | 12.24 | 13.01 | + |
| Lrdd | 10.39 | 9.39 | 10.7 | 5.47 | 6.69 | 5.87 | 5.63 | 6.14 | 5.72 | 3.84 | 5.44 | 4.43 | + |
| Tceb1 | 123.11 | 129.11 | 133.76 | 81.11 | 80.73 | 84.94 | 72.53 | 82.63 | 77.41 | 60.25 | 73.61 | 63.8 | + |
| Car12 | 11.99 | 11.26 | 11.11 | 3.83 | 4.84 | 4.08 | 4.14 | 6.95 | 5.85 | 3.33 | 3.25 | 2.47 | + |
| Rdh13 | 17.67 | 17.47 | 15.65 | 12.08 | 13.29 | 13.42 | 10.79 | 13.12 | 13.68 | 13.21 | 11.78 | 11.65 | + |
| Zbtb8os | 81.83 | 82.17 | 76.79 | 36.86 | 43.22 | 44.16 | 33.52 | 44.05 | 33.06 | 31.37 | 26.5 | 33.28 | + |
| Slc33a1 | 17.53 | 17.49 | 19.71 | 8.32 | 10.25 | 10.84 | 10.01 | 10.93 | 9.97 | 10.13 | 9.52 | 10.14 | + |
| Mphosph10 | 36.32 | 37.13 | 38.17 | 21.74 | 23.83 | 22.7 | 18.37 | 25.89 | 22.51 | 18.62 | 22.17 | 17.34 | + |
| Slc7a3 | 29.4 | 28.34 | 27.45 | 12.02 | 19.18 | 13.76 | 16.89 | 31.8 | 28.08 | 6.36 | 8.45 | 4.97 | + |
| Usp10 | 51.93 | 56.19 | 59.29 | 32.02 | 35.44 | 34.32 | 26.37 | 35.28 | 30.31 | 22.43 | 25.5 | 22.59 | + |
| Plekhg5 | 0.62 | 0.38 | 0.45 | 1.24 | 1.21 | 1.12 | 0.88 | 1.06 | 0.95 | 1.47 | 0.97 | 0.71 | + |
| Agpat6 | 31.25 | 32.75 | 29.14 | 31.3 | 28.46 | 28.78 | 21.56 | 24.78 | 23.04 | 24.41 | 24.08 | 22.99 | + |
| Jagn1 | 63.48 | 62.43 | 54.84 | 33.72 | 35.66 | 36.43 | 30.37 | 39.67 | 31.78 | 26.11 | 25.15 | 28.26 | + |
| Ikbkap | 12.25 | 12.02 | 13.88 | 6.86 | 6.61 | 6.77 | 4.75 | 7.33 | 6.4 | 4.47 | 4.61 | 4.65 | + |
| Vma21 | 14.21 | 15.58 | 14.81 | 8.13 | 8.21 | 7.87 | 5.97 | 8.18 | 8.34 | 6.35 | 6.57 | 6.41 | + |
| Atf1 | 21.92 | 27.78 | 25.4 | 19.19 | 21.78 | 20.95 | 17.35 | 19 | 19.46 | 20.24 | 20.75 | 19.05 | + |
| Dhx35 | 12.92 | 13.61 | 13.13 | 18.63 | 18.46 | 18.08 | 15.74 | 15.31 | 16.86 | 16.77 | 17.69 | 16.49 | + |
| 4930579G24Rik | 18.68 | 17.96 | 17.25 | 10.23 | 12.13 | 11.37 | 8.91 | 8.17 | 8.27 | 7.58 | 5.84 | 7 | + |

| | | | | | | | | | | | | | |
|---|---|---|---|---|---|---|---|---|---|---|---|---|---|
| Dnajc19 | 73.09 | 83.08 | 79.19 | 49.87 | 61.89 | 54.17 | 51.77 | 55.93 | 53.3 | 39.76 | 47.88 | 47.44 | + |
| Hcls1 | 106.52 | 112.95 | 107.64 | 100.72 | 93.82 | 93.57 | 86.97 | 79.97 | 85.61 | 98.39 | 84.6 | 96.63 | + |
| Comt | 57.03 | 61.89 | 60.18 | 35.22 | 36.32 | 33.76 | 30.98 | 34.37 | 31.66 | 23.57 | 26.84 | 27.38 | + |
| Elp4 | 7.42 | 6.75 | 6.34 | 4.16 | 4.8 | 4.43 | 3.55 | 4.09 | 3.01 | 4.08 | 2.59 | 2.98 | + |
| Rpp40 | 7.7 | 7.73 | 7 | 3.89 | 4.42 | 3.75 | 3.2 | 4.87 | 3.95 | 2.58 | 2.74 | 2.89 | + |
| Rps4x | 3111.6 | 3131.97 | 3101.67 | 4947.99 | 4894.71 | 5121.89 | 4633.64 | 4342.55 | 4419.5 | 5021.04 | 5517.28 | 5012.14 | + |
| Zcwpw1 | 3.62 | 3.34 | 3.23 | 9.75 | 9.34 | 7.02 | 3.58 | 4.04 | 3.61 | 4.95 | 4.72 | 6.73 | + |
| 4632428N05Rik | 16.11 | 14.48 | 11.58 | 25.96 | 21.87 | 21.53 | 43.22 | 26.45 | 36.12 | 31.83 | 22.8 | 33.92 | + |
| Zfp61 | 4.82 | 5.33 | 5.26 | 4.2 | 4.12 | 6.52 | 5.97 | 6.56 | 5.44 | 6.78 | 8.72 | 7.63 | + |
| Lmf1 | 10.92 | 9.97 | 9.99 | 14.74 | 14.73 | 16.93 | 14.61 | 15.31 | 15.42 | 17.06 | 13.51 | 16.14 | + |
| Snrnp40 | 103.82 | 107.4 | 102.65 | 54.6 | 63.96 | 58.91 | 46.8 | 52.41 | 55.04 | 41.76 | 46.73 | 43.21 | + |
| Chac1 | 11.28 | 10.05 | 10.72 | 8.29 | 12.77 | 15.2 | 12.82 | 21.6 | 16.99 | 6.05 | 5.38 | 6.28 | + |
| Lgals3bp | 124.21 | 126.09 | 109.76 | 100.26 | 95.05 | 101.34 | 45.99 | 46.3 | 45.93 | 72.88 | 52.33 | 78.54 | + |
| Armc1 | 25.32 | 26.13 | 22.74 | 16.8 | 17.04 | 17.41 | 14.8 | 16.6 | 15.82 | 15.36 | 14.3 | 14.91 | + |
| Pinx1 | 17.16 | 14.95 | 15.69 | 8.37 | 8.64 | 8.17 | 5.66 | 8.48 | 10.13 | 7.07 | 7.94 | 6.83 | + |
| Gtf2f1 | 93.06 | 91.39 | 87.11 | 57.58 | 59.66 | 58.17 | 53.12 | 59.09 | 51.46 | 52.83 | 50.56 | 48.06 | + |
| Cars | 52.33 | 49.11 | 50.23 | 41.05 | 51.55 | 52.2 | 45.14 | 59 | 52.69 | 35.12 | 39.84 | 30.46 | + |
| G3bp1 | 290.47 | 304.26 | 298.02 | 182.19 | 191.6 | 190.41 | 149.38 | 181.45 | 168.44 | 140.67 | 145.85 | 138.06 | + |
| Apip | 26.86 | 31.05 | 26.86 | 14.84 | 19.66 | 18.08 | 13.81 | 15.64 | 16.72 | 12.01 | 12.01 | 17.53 | + |
| Arhgef3 | 6.08 | 7.06 | 5.58 | 9.33 | 8.55 | 10.32 | 15.09 | 9.98 | 10.81 | 9.46 | 7.02 | 10.13 | + |
| Plxna3 | 1.29 | 1.3 | 1.38 | 1.14 | 1.11 | 1.19 | 1.39 | 1.73 | 1.07 | 0.77 | 0.71 | 0.76 | + |
| H2afj | 35.88 | 34.79 | 34.52 | 39.98 | 39.92 | 38.55 | 52.49 | 58.38 | 52 | 48.49 | 55.83 | 48.89 | + |
| Tmem71 | 2.92 | 3.22 | 2.32 | 6.93 | 5.37 | 5.72 | 14.84 | 8.62 | 11.08 | 13.08 | 12.13 | 16.56 | + |
| Mlf1 | 5.58 | 4.69 | 4.6 | 2.1 | 1.62 | 2.29 | 0.67 | 0.84 | 0.62 | 0.97 | 0.22 | 1.14 | + |
| Rbm12 | 13.97 | 14.44 | 13.94 | 7.22 | 7.32 | 7.43 | 6.09 | 7.01 | 5.87 | 5.26 | 5.69 | 5.46 | + |
| Satb1 | 15.85 | 17.77 | 18.55 | 36.38 | 37.73 | 37.32 | 30.48 | 26.4 | 26.97 | 43.99 | 37.56 | 45.54 | + |
| Oxct1 | 49.46 | 47.78 | 49.17 | 40.24 | 40.8 | 41.65 | 33.35 | 36.02 | 33.93 | 31.76 | 34.41 | 34 | + |
| Mars2 | 13.23 | 14.21 | 13.81 | 7.5 | 9.27 | 8.89 | 7.52 | 7.46 | 8.28 | 8.58 | 7.68 | 7.44 | + |
| Sorbs1 | 1.18 | 1.03 | 1.02 | 2.34 | 1.59 | 1.57 | 2.06 | 1.72 | 2.02 | 2.01 | 1.66 | 2.15 | + |
| Gltpd1 | 13.94 | 12 | 12.57 | 7.33 | 8.44 | 6.08 | 5.76 | 6.96 | 4.57 | 4.06 | 4.17 | 4.37 | + |
| Adsl | 86.22 | 90.51 | 91.95 | 41.25 | 43.44 | 44.37 | 28.28 | 42.66 | 39.19 | 25.09 | 22.46 | 25.27 | + |
| Wdr55 | 58.65 | 55.3 | 58.04 | 31.35 | 34.05 | 36.14 | 28 | 34.39 | 34.35 | 24.78 | 26.9 | 25.24 | + |
| Sssca1 | 39.69 | 46.3 | 40.97 | 22.08 | 21.99 | 23.2 | 16.49 | 20.17 | 20.37 | 15.34 | 18.68 | 17.21 | + |
| Tma7 | 214.62 | 239.2 | 225.86 | 140.15 | 146.11 | 152.87 | 162.83 | 156 | 162.38 | 143.78 | 145.32 | 143.61 | + |
| Cysltr1 | 5.66 | 5.43 | 5.91 | 5.12 | 6.96 | 5.7 | 7.93 | 6.29 | 9.97 | 2.18 | 2.04 | 1.78 | + |
| Gab2 | 1.44 | 0.82 | 0.77 | 6.68 | 5.65 | 5.68 | 4.98 | 4.22 | 5.66 | 5.82 | 5.06 | 4.49 | + |
| Vimp | 80.55 | 73.47 | 69.14 | 41.68 | 43.71 | 42.03 | 41.98 | 55.95 | 46.3 | 39.19 | 40.45 | 33.33 | + |
| Selrc1 | 7.07 | 6.8 | 6.19 | 3.3 | 3.86 | 4.12 | 3.8 | 3.8 | 3.38 | 3.64 | 2.76 | 2.79 | + |
| Ttc3 | 5.47 | 4.57 | 5.22 | 8.64 | 8.09 | 9.7 | 8.57 | 7.89 | 7.82 | 8.67 | 11.39 | 7.38 | + |
| Cenpa | 105.17 | 106.03 | 100.43 | 80.41 | 94.47 | 93.29 | 64.47 | 71.6 | 63.03 | 68.55 | 75.18 | 62.95 | + |
| Slc25a30 | 7.09 | 8.7 | 9.16 | 19.22 | 17.64 | 18.84 | 15.19 | 15.04 | 13.7 | 18.03 | 21.1 | 14.26 | + |
| Mrpl22 | 81.95 | 75.55 | 86.85 | 44 | 51.68 | 46.19 | 45.34 | 51.63 | 56.31 | 45.75 | 45.31 | 43.7 | + |
| Rpl11 | 1336.24 | 1429.82 | 1420.1 | 2064.73 | 1944.26 | 2028.17 | 2121.67 | 1946.41 | 2047.48 | 2210.68 | 2353.06 | 2202.68 | + |
| Snrpa | 142.21 | 152.98 | 153.85 | 94.29 | 99.22 | 104.43 | 88.3 | 100.88 | 98.33 | 84.65 | 83.69 | 74.51 | + |
| Ctc1 | 12.38 | 12.04 | 11.97 | 17.87 | 18.43 | 18.22 | 16.31 | 17.27 | 16.16 | 14.91 | 21.11 | 14.92 | + |
| Mapk11 | 8.07 | 9.56 | 9.39 | 6.82 | 7.13 | 6.87 | 8.31 | 8.47 | 8.96 | 5.41 | 4.12 | 4.3 | + |
| Pus10 | 26.08 | 23.67 | 24.53 | 14.4 | 14.93 | 17.71 | 11.67 | 17.26 | 14.81 | 13.79 | 14.46 | 10.78 | + |
| Syt12 | 0.14 | 0.38 | 0.08 | 0.38 | 1.12 | 0.86 | 0.71 | 0.83 | 0.63 | 0.87 | 0.68 | 0.5 | + |
| Polr2k | 121.4 | 120.92 | 117.75 | 63.85 | 63.45 | 68.06 | 53.91 | 68.97 | 66.68 | 46.63 | 48.25 | 45.36 | + |
| Ccdc28a | 2.64 | 3.29 | 2.83 | 7.37 | 5.67 | 7.16 | 7.2 | 8.46 | 6.95 | 6.92 | 7.66 | 8.06 | + |
| Itsn2 | 2.58 | 2.8 | 3.49 | 4.43 | 3.64 | 3.73 | 4.84 | 4.83 | 5.04 | 4.83 | 4.25 | 3.61 | + |
| Nup155 | 10.06 | 11.03 | 10.03 | 5.37 | 6.23 | 5.49 | 3.75 | 4.71 | 4.01 | 2.68 | 2.53 | 3.29 | + |
| Kat2b | 1.04 | 1.15 | 1.25 | 2.22 | 2.27 | 2.04 | 2.14 | 1.85 | 2.45 | 2.83 | 2.87 | 2.45 | + |
| Lat | 187.18 | 208.55 | 182.44 | 125.54 | 100.41 | 102.94 | 144.55 | 136.46 | 133.09 | 109.08 | 95.89 | 118.09 | + |
| Cers5 | 55.95 | 56.97 | 50.88 | 40.12 | 36.09 | 37.97 | 48.03 | 42.52 | 45.59 | 37.74 | 37.16 | 36.87 | + |
| Mfsd1 | 30.69 | 30.52 | 30.32 | 21.93 | 20.77 | 22.79 | 27.79 | 28.73 | 26.55 | 24.56 | 24.93 | 27.39 | + |
| Rmnd5b | 34.73 | 31.13 | 27.59 | 21.05 | 20.05 | 21.55 | 24.07 | 23.63 | 24.73 | 23.58 | 18.18 | 21.3 | + |
| Cxcr3 | 7.81 | 6.81 | 6.72 | 3.73 | 4.11 | 4.01 | 2.72 | 1.42 | 2.94 | 5.18 | 2.49 | 3.48 | + |
| Mrps22 | 61.71 | 60.2 | 63.01 | 31.78 | 36.17 | 35.06 | 29.42 | 37.37 | 35.84 | 26.18 | 23.28 | 21.82 | + |
| Tmed10 | 65.04 | 62.18 | 62.49 | 47.47 | 51 | 49.58 | 46.47 | 51.9 | 51.2 | 43.35 | 44.66 | 44.2 | + |
| Slc35g1 | 4.26 | 4.22 | 4.51 | 2.39 | 2.26 | 2.33 | 2.27 | 2.38 | 2.49 | 1.39 | 1.5 | 2.33 | + |
| Zbed4 | 4.37 | 4.15 | 4.08 | 2.72 | 3.28 | 3.16 | 2.67 | 3.33 | 2.57 | 3.41 | 2.84 | 2.66 | + |
| Mbnl1 | 162.25 | 170.3 | 183.47 | 112.42 | 108.05 | 97.49 | 107.89 | 111.12 | 122.78 | 120.41 | 121.23 | 105.95 | + |
| Parp1 | 51.34 | 53.26 | 55.55 | 42.41 | 43.18 | 41.2 | 36.2 | 41.94 | 37.76 | 36.47 | 37.97 | 36.69 | + |
| Ndufa1 | 195.82 | 184.6 | 173.2 | 139.48 | 154.48 | 147.34 | 156.66 | 161.67 | 156.24 | 133.71 | 137.01 | 135.63 | + |
| Kpna1 | 12.98 | 15.56 | 14.2 | 10.83 | 11.78 | 10.69 | 11.16 | 11.41 | 11.96 | 11.45 | 11.71 | 10.39 | + |
| Mrto4 | 117.14 | 121.71 | 117.45 | 40.9 | 40.08 | 42.03 | 32.17 | 54.11 | 51.03 | 27.45 | 30.23 | 23.03 | + |
| Slc39a13 | 3.32 | 2.87 | 2.62 | 7.07 | 6.78 | 6.73 | 5.61 | 5.87 | 5.66 | 8.03 | 8.76 | 7.23 | + |
| Mrps30 | 71.95 | 71.69 | 77.43 | 62.54 | 61.01 | 66.23 | 55.91 | 62.03 | 58.1 | 56.3 | 63.1 | 59.85 | + |
| Tmem218 | 3.47 | 3.94 | 3.55 | 7.37 | 8.38 | 6.62 | 7.38 | 5.01 | 6.21 | 7.52 | 7.85 | 8.42 | + |
| Nrp1 | 1.65 | 1.87 | 2 | 4.84 | 4.48 | 4.09 | 0.42 | 0.74 | 0.54 | 0.6 | 0.42 | 0.85 | + |
| Lypla1 | 73.1 | 75.5 | 81.03 | 59.52 | 59.67 | 59.47 | 57.36 | 57.18 | 62.79 | 58.27 | 56.58 | 56.7 | + |
| Vwa9 | 22.25 | 23.89 | 23.28 | 13.75 | 12.97 | 13.92 | 14.16 | 13.24 | 14.39 | 11.23 | 12.86 | 13.22 | + |
| Cct2 | 405.77 | 406.08 | 421.54 | 240.67 | 253.37 | 259.25 | 280.01 | 321.66 | 310.03 | 236.23 | 263.15 | 233.6 | + |
| Gtf2b | 62.44 | 65.13 | 62.06 | 38.09 | 35.94 | 38.46 | 35.39 | 38.56 | 35.82 | 33.17 | 29.34 | 34.02 | + |
| Exosc1 | 39.83 | 44.34 | 44.38 | 25.68 | 25.05 | 24.64 | 17.04 | 24.4 | 24.08 | 17.88 | 14.84 | 19.9 | + |
| Orc3 | 25.01 | 25.15 | 24.24 | 15.27 | 15.01 | 15.97 | 16.65 | 16.63 | 16.36 | 13.89 | 14.3 | 14.31 | + |
| Zcchc24 | 0.59 | 0.31 | 0.17 | 2.15 | 2.1 | 1.98 | 2.13 | 1.36 | 1.59 | 1.94 | 2.31 | 2.45 | + |
| Idi1 | 168.16 | 155.67 | 171.21 | 65.91 | 77.17 | 76.68 | 56.1 | 73.9 | 66.28 | 45.16 | 49.63 | 45.07 | + |
| Nmi | 25.48 | 27.96 | 26.32 | 16.19 | 18.58 | 16.34 | 16.61 | 17.33 | 17.63 | 15.98 | 14.18 | 19.49 | + |
| Mdp1 | 35.92 | 33.4 | 35.12 | 55.01 | 56.77 | 52.28 | 50.43 | 58.47 | 53.48 | 59.4 | 58.87 | 64.38 | + |
| Dach2 | 0.39 | 0.32 | 0.46 | 2.83 | 2.65 | 2.41 | 0.66 | 0.76 | 0.63 | 1.29 | 1.37 | 1.98 | + |
| Srfbp1 | 34.26 | 35.55 | 36.76 | 22.11 | 20.59 | 25.08 | 21.69 | 24.53 | 23 | 19.66 | 22.79 | 20.4 | + |
| Ilkap | 50.16 | 47.62 | 51.45 | 26.98 | 29.52 | 28.9 | 24.24 | 31.05 | 30.2 | 23.97 | 20.99 | 23.43 | + |
| Dbr1 | 28.99 | 27.06 | 27.72 | 15.87 | 17.22 | 16.56 | 13.23 | 14.91 | 13.98 | 12.84 | 12.31 | 14.41 | + |
| Sephs1 | 18.98 | 19.72 | 17.61 | 10.65 | 13.11 | 11.74 | 8.88 | 10.24 | 11.05 | 9.25 | 8.78 | 9.52 | + |
| Nudc | 221.24 | 222.29 | 228.91 | 88.49 | 105.8 | 105.74 | 77.56 | 105.48 | 95.96 | 53.77 | 68.54 | 57.6 | + |
| Sf3a2 | 3.89 | 4.38 | 6.56 | 2 | 2.57 | 2.48 | 2.75 | 2.95 | 2.18 | 1.85 | 2.55 | 2.48 | + |
| Acss2 | 10.25 | 9.43 | 10.88 | 15.68 | 16.31 | 12.61 | 19.33 | 18.18 | 16.1 | 13.46 | 16.51 | 14.85 | + |
| B3galt4 | 3.08 | 2.66 | 2.67 | 6.21 | 4.24 | 5.14 | 5.43 | 5.92 | 7.45 | 6.6 | 5.24 | 7.46 | + |
| Tpm3 | 424.5 | 437.25 | 426.65 | 278.66 | 293.97 | 301.35 | 269.39 | 318.78 | 314.04 | 245.65 | 233.36 | 246.32 | + |
| Tmem229b | 8.03 | 9.46 | 5.81 | 3.92 | 3.56 | 3.84 | 8.08 | 5.96 | 4.91 | 5.9 | 3.47 | 5.33 | + |
| Srp14 | 204.93 | 195.15 | 198.15 | 159.9 | 156.61 | 153.73 | 178.03 | 192.7 | 179.55 | 154.44 | 157.34 | 154.24 | + |
| Gcat | 42.79 | 43.98 | 45.84 | 14.98 | 17.08 | 17.4 | 18.42 | 22.89 | 20.87 | 12.46 | 10.11 | 10.83 | + |
| Trp53 | 127.54 | 139.62 | 125.47 | 73.02 | 86.13 | 80.78 | 74.14 | 92.13 | 82.26 | 72.13 | 69.09 | 67.24 | + |
| Eef1a1 | 5363.7 | 5544.66 | 5514.62 | 9961.68 | 9871.72 | 10343.76 | 10038.67 | 9257.04 | 9650.95 | 10307.58 | 11468.54 | 10127.76 | + |
| Angptl6 | 7.99 | 6.75 | 8.21 | 27.58 | 32.35 | 29.52 | 24.01 | 32.6 | 33.18 | 15.98 | 18.72 | 15.85 | + |
| 4933421O10Rik | 1.22 | 1.22 | 1.13 | 1.78 | 1.73 | 2.07 | 1.99 | 1.76 | 2.15 | 2.25 | 2.17 | 2.02 | + |
| Hmgcr | 73.12 | 79.52 | 76.31 | 60 | 58.73 | 61.92 | 48.59 | 54.42 | 54.26 | 46.34 | 49.65 | 44.27 | + |
| Smim14 | 18.08 | 16.56 | 17.41 | 28.87 | 28.02 | 29.71 | 39.52 | 34.63 | 37.35 | 35.62 | 33.98 | 35.25 | + |
| Ppid | 138.52 | 141.95 | 144.9 | 37.93 | 43.89 | 40.82 | 29.06 | 49.15 | 44.12 | 25.79 | 28.1 | 22.28 | + |
| Prdm1 | 1.62 | 2.09 | 2.13 | 2.26 | 2.31 | 1.55 | 3.6 | 2.91 | 2.23 | 4.48 | 6.17 | 6.68 | + |
| Tasp1 | 41.18 | 30.96 | 38.9 | 18.31 | 25.41 | 25.26 | 13.07 | 22.33 | 18.76 | 21.04 | 22.21 | 21.33 | + |
| Tbl3 | 50.63 | 49.69 | 49.78 | 18.75 | 20.05 | 19.97 | 14.69 | 23.12 | 19.88 | 15.04 | 13.51 | 17.19 | + |
| Gpatch11 | 5.78 | 5.76 | 5.63 | 4.5 | 4.35 | 5.31 | 3.73 | 4.04 | 4.18 | 3.97 | 4.56 | 3.44 | + |
| Pkn1 | 6.67 | 7.1 | 6.9 | 14.93 | 12.1 | 12.38 | 15.55 | 12.26 | 14.74 | 16.38 | 15.83 | 17.96 | + |
| Sh3bgrl | 54.62 | 54.27 | 53.03 | 43.05 | 45.36 | 45.43 | 44.36 | 47.46 | 48.12 | 38.61 | 44.92 | 41.4 | + |
| Vcp | 206.92 | 204.42 | 209.49 | 122.85 | 126.03 | 127.78 | 113.23 | 145.94 | 128.39 | 111.03 | 122.57 | 112.33 | + |
| Eif4g2 | 77.07 | 90.48 | 89.96 | 40.99 | 45 | 41.49 | 36.66 | 45.48 | 41.32 | 35.96 | 38.31 | 33.83 | + |
| 1700001O22Rik | 1.03 | 1.41 | 1.24 | 3.69 | 4.12 | 3.81 | 3.31 | 2.61 | 3.06 | 4.05 | 4.67 | 4.28 | + |
| Ahdc1 | 0.73 | 0.98 | 0.97 | 2.13 | 1.33 | 1.55 | 1.56 | 1.3 | 1.38 | 1.27 | 1.34 | 1.75 | + |
| Olfr613 | 3.73 | 4.5 | 4.02 | 7.27 | 7.26 | 6.24 | 6.43 | 5.88 | 5.41 | 7.68 | 7.07 | 7.42 | + |
| Nanog | 2.78 | 3.53 | 2.9 | 0.87 | 1.37 | 0.73 | 1.22 | 1.28 | 0.86 | 0.8 | 1.82 | 1.07 | + |
| Polr3g | 14.4 | 12.87 | 15.1 | 9.79 | 10.59 | 10.05 | 8.32 | 11.37 | 9.68 | 7.48 | 11.37 | 8.35 | + |
| Ccser2 | 3.07 | 2.8 | 2.62 | 4.74 | 4.26 | 5.57 | 6.47 | 6.13 | 5.67 | 6.99 | 7.89 | 7.36 | + |
| Uchl3 | 167.97 | 168.95 | 165.69 | 71.6 | 74.78 | 76.28 | 71.46 | 94.16 | 91.03 | 51.36 | 58.58 | 50 | + |
| Ormdl3 | 11.68 | 10.44 | 11.84 | 24.41 | 24.03 | 20.08 | 22.21 | 20.76 | 22.09 | 24.68 | 26.62 | 21.73 | + |
| Zfp956 | 3.88 | 4.39 | 3.18 | 7.79 | 7.7 | 7.59 | 5.47 | 6.92 | 6.4 | 7.06 | 7.9 | 8.38 | + |
| St6galnac4 | 30.19 | 35.69 | 30.53 | 16.62 | 19.03 | 18.41 | 14.13 | 18.04 | 17.4 | 14.96 | 13.56 | 14.67 | + |
| Filip1 | 4.4 | 3.13 | 3.75 | 0.72 | 1.13 | 1.11 | 12.72 | 11.77 | 12.72 | 11.84 | 11.33 | 13.94 | + |
| Tmem66 | 110.43 | 106.12 | 100.45 | 226.49 | 176.95 | 198.83 | 242.72 | 226.04 | 214.68 | 251.17 | 246.07 | 255.4 | + |
| Xrcc6 | 66.32 | 55.84 | 61.41 | 35.8 | 38.05 | 37.12 | 32.47 | 41.77 | 38.82 | 30.22 | 30.01 | 29.38 | + |
| Tsr2 | 7.74 | 6.91 | 7.43 | 4.61 | 4.11 | 4.5 | 3.09 | 4.27 | 4.38 | 3.67 | 3.5 | 3.07 | + |
| Zcchc17 | 35.18 | 36.42 | 30 | 19.8 | 21.19 | 19.85 | 15.19 | 17.98 | 18.29 | 16.87 | 14.98 | 18.47 | + |
| Col18a1 | 0.37 | 0.55 | 0.55 | 1.39 | 1.51 | 1.43 | 4.62 | 3.12 | 3.34 | 3.49 | 4.07 | 4.65 | + |
| Rsf1 | 1.01 | 1.23 | 0.96 | 1.93 | 1.7 | 1.53 | 1.73 | 1.81 | 1.64 | 2.14 | 2.02 | 1.5 | + |
| Cacybp | 224.39 | 224.96 | 232.84 | 93.68 | 106.39 | 101.12 | 71.68 | 111.2 | 101.33 | 66.81 | 66.84 | 59.08 | + |
| Rpp25l | 30.42 | 31 | 29 | 18.08 | 20.98 | 25.62 | 21.71 | 20.72 | 20.65 | 21.48 | 19.69 | 23.36 | + |
| Calcoco1 | 1.41 | 1.65 | 1.4 | 8.42 | 6.81 | 6.01 | 9.48 | 7.47 | 5.55 | 8.81 | 9.01 | 10.7 | + |
| Gprasp1 | 1.75 | 2.13 | 1.93 | 3.81 | 3.61 | 3.9 | 3.33 | 3.19 | 3.21 | 2.75 | 3.85 | 3.07 | + |
| Hn1 | 264.53 | 245.08 | 230.24 | 197.41 | 211.84 | 210.92 | 175.67 | 168.42 | 168.28 | 187.79 | 193.7 | 198.59 | + |
| Vps33a | 9.78 | 9.36 | 9.04 | 6.9 | 7.27 | 5.71 | 6.26 | 6.15 | 7.7 | 6.46 | 6.43 | 6.94 | + |
| Tnip2 | 20.88 | 19.98 | 23.62 | 49.12 | 52.17 | 53.75 | 34.02 | 37.19 | 33.29 | 48.05 | 63.58 | 47.04 | + |
| Agl | 1.25 | 1.36 | 1.12 | 1.7 | 1.8 | 1.95 | 3.08 | 2.28 | 2.45 | 2.16 | 2.35 | 1.85 | + |
| Fbxo42 | 8.66 | 9.44 | 8.71 | 6.3 | 7.27 | 7.06 | 6.13 | 5.94 | 6.45 | 6.58 | 6.64 | 6.3 | + |
| Camk2d | 4.13 | 3.53 | 4.39 | 10.63 | 9.24 | 10.08 | 7.14 | 6.33 | 8.11 | 9.27 | 10.71 | 8.4 | + |
| H2-DMb1 | 5.19 | 6.48 | 5.13 | 9.36 | 8.21 | 8.77 | 18.5 | 16.08 | 17.15 | 12.89 | 9.6 | 11.38 | + |
| Tarbp2 | 47.44 | 42.67 | 43.52 | 33.12 | 36.74 | 37.83 | 35.72 | 36.37 | 36.24 | 31.34 | 36.77 | 33.41 | + |
| Cebpz | 23.15 | 25.41 | 26.2 | 16.75 | 17.11 | 16.82 | 10.64 | 16.59 | 17.19 | 13.84 | 16.18 | 13.05 | + |
| Paxbp1 | 8.17 | 10.3 | 9.98 | 18.24 | 18.13 | 17.3 | 11.22 | 12.79 | 12.18 | 13.31 | 14.73 | 14.35 | + |
| Pwp2 | 23.19 | 23.66 | 22.14 | 10.78 | 12.16 | 9.79 | 7.39 | 10.76 | 10.66 | 6.04 | 5.84 | 7.09 | + |
| Ube3c | 6.78 | 5.63 | 7.05 | 4.71 | 5.03 | 4.76 | 3.4 | 3.42 | 3.59 | 4.05 | 3.29 | 2.84 | + |
| Hpse | 1.09 | 1.72 | 1.75 | 3.49 | 2.94 | 2.96 | 8.63 | 5.16 | 6.32 | 4.25 | 4.72 | 3.85 | + |
| Eef1e1 | 204.34 | 210.24 | 209.11 | 83.06 | 95.84 | 98 | 83.98 | 115.07 | 115.21 | 61.94 | 64.18 | 61.73 | + |
| Nipbl | 3.69 | 3.81 | 5.04 | 9.88 | 9.01 | 9.19 | 7.2 | 6.97 | 7.27 | 11.38 | 11.47 | 10.7 | + |
| Gltscr1l | 1.18 | 1.29 | 1.37 | 2.51 | 2.68 | 2.84 | 2.68 | 2.46 | 2.25 | 3.47 | 3.66 | 2.55 | + |
| Iigp1 | 4.37 | 7.39 | 4.75 | 3.78 | 1.91 | 1.84 | 0.93 | 0.32 | 0.38 | 2.06 | 0.41 | 1.61 | + |
| Wbp11 | 22.12 | 22.89 | 26.93 | 17.53 | 18.78 | 17.06 | 17.56 | 18.71 | 16.12 | 16.14 | 17.32 | 15.82 | + |
| 2610035D17Rik | 0.22 | 0.94 | 0.32 | 2.98 | 2.67 | 2.8 | 3.23 | 1.72 | 2.29 | 4.24 | 3.23 | 4.71 | + |
| Adora2a | 5.05 | 4.13 | 4.86 | 5.19 | 6.7 | 6.47 | 6.3 | 6.35 | 6.18 | 12.96 | 11.26 | 13.03 | + |
| Zbtb7a | 2.17 | 2.51 | 3.28 | 4.86 | 4.62 | 4.43 | 4.98 | 5.37 | 5.07 | 6.06 | 6.03 | 5.32 | + |

| | | | | | | | | | | | | | |
|---|---|---|---|---|---|---|---|---|---|---|---|---|---|
| Ankrd32 | 3.98 | 2.83 | 3.9 | 1.67 | 2.15 | 1.23 | 1.47 | 1.3 | 1.15 | 1.21 | 1.14 | 1.14 | + |
| Calm2 | 307.82 | 261.97 | 271.52 | 247.46 | 272.01 | 278.01 | 222.11 | 220.2 | 210.11 | 236.67 | 229.23 | 231.65 | + |
| Eif3l | 263.7 | 263.16 | 248.63 | 161.86 | 162.47 | 169.03 | 182.3 | 196.06 | 184.68 | 155.7 | 175.15 | 155.07 | + |
| Kdm4b | 1.71 | 2.16 | 1.82 | 5.74 | 5.2 | 5.05 | 8.04 | 4.54 | 5.8 | 7.33 | 8.34 | 7.45 | + |
| Dffb | 6.55 | 7.9 | 4.74 | 3.3 | 4.33 | 3.99 | 2.65 | 4.12 | 3.82 | 4.47 | 3.7 | 2.76 | + |
| Als2cl | 8.18 | 8.81 | 9.75 | 15.44 | 14.44 | 16.13 | 16.04 | 16.64 | 15.67 | 17.2 | 17.19 | 18.62 | + |
| Gngt2 | 10.84 | 12.61 | 10.32 | 33.84 | 33.9 | 32.77 | 46.6 | 34.27 | 37.92 | 50.18 | 44.57 | 48.77 | + |
| Slc27a1 | 3.07 | 3.2 | 3.35 | 9.03 | 7.95 | 5.25 | 11.12 | 7.86 | 8.23 | 10.18 | 10.07 | 12.79 | + |
| 2810474O19Rik | 6.53 | 5.97 | 5.61 | 13.2 | 11.72 | 12.38 | 9.79 | 9.68 | 9.6 | 9.72 | 9.67 | 11.75 | + |
| Slc25a5 | 1107.99 | 1170.7 | 1079.62 | 739.69 | 805.36 | 775.89 | 768.91 | 712.36 | 708.7 | 664.05 | 730.08 | 649.35 | + |
| Agbl5 | 1.8 | 1.33 | 1.45 | 3.48 | 3.06 | 3.81 | 4.64 | 3.25 | 3.59 | 4.19 | 4.59 | 4.3 | + |
| Ift27 | 26.97 | 22.84 | 23.53 | 8.66 | 9.11 | 9.37 | 10.82 | 13.22 | 9.63 | 6.21 | 6.56 | 10.06 | + |
| Dna2 | 5.62 | 6.41 | 5.4 | 1.55 | 1.91 | 1.62 | 1.13 | 1.68 | 1.34 | 0.86 | 0.7 | 0.81 | + |
| Slc35a1 | 22.37 | 22.55 | 22.65 | 14.46 | 15.41 | 16.46 | 12.22 | 13.08 | 13.71 | 11 | 11.46 | 10.29 | + |
| G6pdx | 35.77 | 37.59 | 39.12 | 19.64 | 17.99 | 18.19 | 15.22 | 19.15 | 18.27 | 15.5 | 15.15 | 16.22 | + |
| Ing5 | 11.16 | 10.4 | 10.76 | 6.08 | 6.25 | 6.68 | 6.12 | 7.13 | 6.14 | 6.37 | 6.41 | 4.5 | + |
| Folr4 | 4.54 | 3.82 | 4.25 | 11.88 | 14.97 | 15.83 | 9.44 | 8.08 | 7.99 | 12.06 | 15.96 | 14 | + |
| Psrc1 | 29.2 | 23.14 | 26.38 | 51.58 | 62.28 | 63.81 | 57.5 | 54 | 54.24 | 55.82 | 74.32 | 60.09 | + |
| Hmgb2 | 175.63 | 193.3 | 164.49 | 81.64 | 96.82 | 91.62 | 68.3 | 67.73 | 66.18 | 55.38 | 55.29 | 55.75 | + |
| Hsp90ab1 | 1781.49 | 1815.45 | 1902.05 | 1239.63 | 1306.25 | 1280.52 | 1460.01 | 1484.25 | 1473.74 | 1173.95 | 1357.37 | 1161.22 | + |
| Acer3 | 6.61 | 7.32 | 7.68 | 3.46 | 4.8 | 4.61 | 4.43 | 5.1 | 4.48 | 3.52 | 3.45 | 4.38 | + |
| Phf1 | 1.27 | 1.01 | 1.45 | 5.31 | 4.96 | 4.73 | 4.99 | 4.04 | 4.46 | 7.08 | 5.51 | 6.38 | + |
| Kidins220 | 6.5 | 6.22 | 6.27 | 10.84 | 9.59 | 9.88 | 10.44 | 8.92 | 9.61 | 10.68 | 10.82 | 10.87 | + |
| Rbm4 | 14.64 | 14.85 | 13.7 | 9.94 | 9.84 | 10.91 | 8.53 | 11.46 | 9.47 | 9.87 | 7.65 | 9.69 | + |
| Rdx | 40.78 | 45.04 | 48.61 | 32.9 | 38.02 | 34.42 | 31.71 | 35.18 | 34.58 | 31.73 | 36.43 | 30.24 | + |
| Gm20594 | 2425.89 | 2446.4 | 2374.6 | 2904.86 | 2950.24 | 3048.62 | 2799.65 | 2399.81 | 2397.54 | 3220.02 | 3288.05 | 3444.08 | + |
| Yipf2 | 7.54 | 8.12 | 9.53 | 12.31 | 13.45 | 12.17 | 13.85 | 14.13 | 13.29 | 12.4 | 12.14 | 13.71 | + |
| AI662270 | 54.65 | 65.87 | 66.51 | 32.54 | 35.91 | 30.12 | 34.12 | 39.92 | 32.54 | 29.86 | 30.03 | 34.01 | + |
| Utp3 | 61.02 | 64.51 | 61.7 | 40.42 | 40.96 | 44.71 | 44.08 | 44.72 | 43.36 | 44.91 | 39 | 45.19 | + |
| Rest | 2.1 | 2.45 | 2.62 | 3.33 | 3.51 | 3.69 | 4.54 | 4.67 | 3.75 | 4.41 | 4.41 | 4.19 | + |
| Qsox1 | 6.65 | 7.53 | 7.01 | 19.58 | 16.56 | 17.23 | 19.42 | 14.83 | 13.85 | 19.29 | 20.72 | 17.81 | + |
| Gtf2h2 | 34.71 | 32.38 | 35.29 | 19.05 | 19.78 | 22.45 | 19.96 | 20.68 | 21.88 | 16.29 | 19.93 | 16.88 | + |
| Ddx39b | 172.52 | 182.78 | 182.38 | 107.81 | 105.32 | 107.61 | 79.99 | 97.12 | 84.12 | 74.96 | 75.97 | 80.32 | + |
| Gtf3c4 | 1.83 | 2.5 | 1.66 | 1.4 | 1.29 | 1.2 | 1.16 | 0.84 | 0.88 | 1.07 | 0.97 | 1.14 | + |
| Snrpf | 174 | 190.01 | 173.23 | 108.73 | 114.88 | 110.19 | 106.97 | 122.58 | 116.49 | 91.5 | 96.73 | 94.76 | + |
| Rpn2 | 198.68 | 201.02 | 193.5 | 144.63 | 143.2 | 151.07 | 141.32 | 153 | 147.27 | 105.59 | 128.11 | 109.24 | + |
| Sf3b5 | 142.26 | 151.29 | 143.22 | 89.55 | 94.76 | 92.09 | 88.95 | 94.81 | 93.7 | 68.89 | 72.97 | 76.79 | + |
| Chek1 | 11.86 | 13.12 | 12.23 | 5.88 | 8.29 | 7.1 | 2.83 | 4.29 | 3.77 | 3.1 | 3.14 | 3.01 | + |
| Skap1 | 55.46 | 57.28 | 54.27 | 83.52 | 72.13 | 73.84 | 130.82 | 94.34 | 100.53 | 98.16 | 101.67 | 109.28 | + |
| Gabarap | 114.03 | 107.1 | 100.71 | 177.61 | 166.73 | 179.9 | 200.39 | 171.37 | 175.93 | 209.59 | 235.43 | 218.69 | + |
| Uqcr10 | 246.06 | 238.7 | 245.15 | 139.98 | 147.92 | 176.41 | 148.8 | 178.16 | 161.79 | 118.05 | 141.44 | 124.26 | + |
| Rfwd3 | 26.32 | 25.13 | 24.96 | 15.22 | 17.3 | 16.49 | 11.78 | 13.67 | 13.95 | 12.98 | 14.29 | 12.35 | + |
| Rpl7a | 1337.97 | 1377.63 | 1393.33 | 1812.52 | 1779.9 | 1881.89 | 1999.33 | 1839.03 | 1929.06 | 1937.63 | 2184.79 | 1977.13 | + |
| H2-T23 | 247.21 | 234.09 | 227.62 | 218.67 | 210.22 | 225.6 | 244.16 | 240.17 | 230.52 | 263.74 | 277.56 | 281.97 | + |
| Plek | 62.49 | 48.51 | 60.44 | 24.94 | 31.25 | 32.34 | 33.42 | 52.32 | 42.28 | 39.74 | 31.67 | 34.13 | + |
| Hars | 115.97 | 114.5 | 116.31 | 52.15 | 51.7 | 57.73 | 44.2 | 61.65 | 58.02 | 35.2 | 31.26 | 29.88 | + |
| Ankrd13d | 4.8 | 3.64 | 5.4 | 6.7 | 7.09 | 8.31 | 12.05 | 8.72 | 10.15 | 9.4 | 9.93 | 10.8 | + |
| Klf11 | 0.97 | 1.29 | 1.37 | 1.91 | 1.86 | 2.36 | 2.53 | 2.2 | 1.77 | 2.28 | 2.35 | 2.56 | + |
| Smug1 | 2.59 | 2.27 | 2.48 | 3.55 | 4.06 | 3.83 | 4.47 | 3.81 | 3.95 | 4.02 | 4.11 | 3.64 | + |
| Gmfg | 98.74 | 96.87 | 98.03 | 136.05 | 126.7 | 122.12 | 219.38 | 185.57 | 188.02 | 162.71 | 161.44 | 185.94 | + |
| Nop14 | 45.16 | 44.18 | 45.58 | 29.3 | 29.71 | 32.35 | 23.82 | 32.48 | 29.33 | 30.13 | 29.79 | 25.93 | + |
| Coq4 | 10.06 | 8.11 | 10.18 | 2.79 | 3.39 | 3.88 | 2.49 | 3.22 | 2.92 | 2.33 | 1.09 | 1.91 | + |
| Sfpq | 13.77 | 14.83 | 14.96 | 8.76 | 9.02 | 8.27 | 5.7 | 6.71 | 6.96 | 6.82 | 6.54 | 6.38 | + |
| Mycbp | 26.1 | 30.13 | 27.23 | 15.1 | 12.99 | 14.04 | 12.85 | 13.53 | 13.25 | 11.59 | 9.88 | 12.64 | + |
| Rad52 | 10.37 | 8.86 | 9.76 | 20.38 | 20.43 | 20.24 | 26.38 | 22.55 | 23.2 | 32.08 | 31.21 | 34 | + |
| Gpx8 | 0.73 | 1.43 | 0.32 | 4.91 | 3.56 | 3.57 | 5.88 | 4.22 | 5.02 | 5.74 | 5.58 | 5.9 | + |
| Sarnp | 190.19 | 208.58 | 196.02 | 122.31 | 128.61 | 132.83 | 112.28 | 140.02 | 123.37 | 97.54 | 104.86 | 100.79 | + |
| Rpl18a | 1520.76 | 1544.2 | 1540.04 | 2822.15 | 2719.54 | 2776.07 | 2591.72 | 2302.32 | 2384.97 | 2968.5 | 3123.82 | 3026.02 | + |
| Mrpl16 | 48.89 | 44.26 | 45.37 | 23.56 | 27.6 | 28.43 | 26.28 | 33.04 | 28.64 | 18.27 | 21.44 | 22.87 | + |
| Rnf157 | 4.51 | 3.51 | 3.83 | 7.52 | 8.48 | 8.89 | 5.83 | 6.7 | 6.68 | 7.83 | 6.62 | 7.8 | + |
| Jhdm1d | 0.39 | 0.41 | 0.54 | 1.76 | 1.45 | 1.11 | 1.64 | 1.23 | 1.17 | 1.59 | 1.72 | 1.77 | + |
| Ints4 | 22.17 | 19.93 | 21.75 | 13.94 | 14.32 | 13.64 | 12.31 | 14.14 | 14.9 | 10.99 | 12.58 | 9.74 | + |
| Mrpl27 | 60.12 | 57.26 | 63.32 | 34.89 | 34.03 | 37.61 | 38.75 | 39.59 | 41.17 | 32.17 | 35.7 | 36.4 | + |
| Rpp30 | 53.92 | 58.24 | 55.33 | 30.34 | 34.91 | 35.14 | 22.73 | 31.5 | 27.75 | 28.26 | 27.56 | 24.93 | + |
| Psmd14 | 214.55 | 206.64 | 208.33 | 119.22 | 127.87 | 125.59 | 97.84 | 135.84 | 129.79 | 97.78 | 96.99 | 90.69 | + |
| Rps6ka1 | 40.69 | 39.28 | 36.37 | 30.41 | 25.72 | 27.96 | 34.8 | 33.21 | 28.98 | 32.07 | 27.53 | 32.87 | + |
| Mrpl3 | 54.55 | 64.07 | 61.1 | 31.13 | 34.24 | 28.77 | 27.02 | 35.95 | 35.08 | 20.22 | 20.81 | 25.87 | + |
| Zc3h18 | 16.27 | 16.67 | 17.05 | 10.31 | 10.89 | 10.13 | 9.01 | 9.9 | 9.97 | 8.52 | 7.95 | 9.6 | + |
| Tsn | 54.2 | 55.37 | 54.44 | 38.03 | 37.77 | 36.89 | 35.38 | 37.06 | 36.14 | 32.68 | 33.67 | 32.05 | + |
| Kpnb1 | 24.04 | 23.35 | 24.92 | 16.93 | 16.02 | 17.06 | 15.74 | 16.18 | 15.4 | 17.98 | 17.75 | 14.63 | + |
| Ahsa1 | 155.26 | 152.83 | 159.98 | 59.09 | 64.59 | 64.54 | 48.24 | 70.25 | 59.55 | 48.32 | 45.34 | 49.52 | + |
| Rps3a1 | 1978.72 | 2050.19 | 1986.26 | 2321.25 | 2330.34 | 2467.82 | 2462.97 | 2466.29 | 2460.9 | 2238.96 | 2557.72 | 2225.79 | + |
| Gtf3a | 95.04 | 97.73 | 96.36 | 67.29 | 74.98 | 71.26 | 73.63 | 85.85 | 79.9 | 68.32 | 78.74 | 68.79 | + |
| Adam9 | 7 | 6.76 | 7.69 | 35.62 | 31.34 | 31.1 | 7.33 | 7.68 | 9.04 | 22.23 | 21.26 | 19.31 | + |
| Nfu1 | 69.43 | 71.41 | 66.63 | 62.29 | 66.13 | 61.82 | 65.4 | 76.01 | 62.63 | 51.73 | 50.31 | 48.56 | + |
| Tpd52 | 20.65 | 20.12 | 20.53 | 38.42 | 33.46 | 33.68 | 44.28 | 36.79 | 43.93 | 38.84 | 42.46 | 42.96 | + |
| Gins4 | 38.17 | 40.39 | 38.75 | 27.94 | 29.68 | 30.85 | 23.9 | 24.88 | 23.99 | 22.69 | 23.48 | 19.4 | + |
| Psma7 | 401.77 | 424.97 | 389.58 | 241.18 | 242.39 | 244.83 | 244.5 | 252.86 | 255.94 | 220.51 | 248.27 | 226.7 | + |
| Sod1 | 703.41 | 715.02 | 688.28 | 687.23 | 651.67 | 719.75 | 841.1 | 663.62 | 705.8 | 793.96 | 917.61 | 810.13 | + |
| Magt1 | 29.77 | 32.23 | 30.67 | 18.64 | 19.5 | 20.31 | 17.61 | 22.07 | 21.73 | 16.05 | 17.14 | 17.06 | + |
| Tgs1 | 17.44 | 15.85 | 15.88 | 10.71 | 11.69 | 10.7 | 9.54 | 9.61 | 10.64 | 8.23 | 10.35 | 9.3 | + |
| Zfr2 | 0.03 | 0.05 | 0.15 | 1.72 | 1.27 | 1.09 | 1.87 | 1.51 | 1.59 | 1.83 | 1.73 | 1.64 | + |
| Utp23 | 16.81 | 16.68 | 15.71 | 10.9 | 12.48 | 12.13 | 9.29 | 10.72 | 11.24 | 9.99 | 10.27 | 10.57 | + |
| Nop10 | 256.95 | 254.49 | 247.08 | 150.51 | 152.24 | 153.69 | 141.72 | 175.52 | 161.88 | 137.38 | 146.5 | 135.11 | + |
| Kti12 | 52.19 | 53.79 | 48.31 | 24.62 | 22.14 | 25.49 | 23.9 | 27.48 | 24.15 | 22.54 | 18.34 | 20.63 | + |
| Lgals8 | 5.99 | 5.31 | 5.1 | 13.35 | 11.17 | 11.92 | 15.87 | 9.51 | 10.39 | 15.24 | 15.71 | 17.33 | + |
| Cd300a | 3.86 | 5.38 | 4.8 | 2.98 | 2.51 | 2.61 | 2.73 | 3.07 | 2.92 | 2.88 | 2.68 | 2.66 | + |
| Rad51ap1 | 23.46 | 20.64 | 20.36 | 11.02 | 13.34 | 13.43 | 6.65 | 7.84 | 5.76 | 4.89 | 5.78 | 4.65 | + |
| Slc7a6 | 22.93 | 24.79 | 23.35 | 20.52 | 21.71 | 21.16 | 17.24 | 22.16 | 19.4 | 18.72 | 20.37 | 19.47 | + |
| Trmt11 | 19.68 | 16.35 | 18.47 | 11.6 | 12.68 | 11.68 | 7.6 | 11.02 | 11.47 | 8.8 | 10.55 | 8.43 | + |
| Pak1ip1 | 57.94 | 53.08 | 53.17 | 41.61 | 32.8 | 35.58 | 36.12 | 42.45 | 36.3 | 31.61 | 32.29 | 29 | + |
| Znhit3 | 54.73 | 50.86 | 54.9 | 27.94 | 28.98 | 25.88 | 24.26 | 29.25 | 23.09 | 20.57 | 17.26 | 18.65 | + |
| Ralgps2 | 1.92 | 2.01 | 1.93 | 3.19 | 2.59 | 3.34 | 4.33 | 3.25 | 3.67 | 3.06 | 4.5 | 3.13 | + |
| Relb | 16.02 | 17.88 | 16.64 | 32.2 | 33.42 | 29.73 | 26.93 | 30.71 | 25.96 | 25.91 | 33.76 | 29.42 | + |
| Prnp | 8.43 | 9.26 | 7.54 | 85.76 | 57.11 | 59.16 | 16.35 | 13.76 | 13.57 | 29.46 | 27.3 | 29.86 | + |
| Taz | 20.13 | 17.53 | 17.14 | 34.73 | 37.72 | 37.25 | 37.12 | 31.8 | 33.2 | 36.93 | 41.49 | 38.31 | + |
| Cox6b1 | 434.67 | 437.01 | 418.01 | 311.4 | 312.12 | 331.09 | 365.39 | 329.49 | 335.68 | 305.09 | 324.02 | 326.68 | + |
| Usp7 | 7.95 | 8.37 | 7.68 | 6.97 | 6.84 | 6.61 | 6.42 | 6.77 | 6.13 | 5.84 | 6.63 | 6.13 | + |
| Prrc2c | 16.88 | 17.47 | 18.09 | 19.21 | 18.07 | 18.96 | 17.12 | 16.96 | 17.81 | 21.93 | 23.61 | 21.26 | + |
| Igf2r | 5.68 | 5.7 | 5.03 | 7.5 | 6.2 | 6.95 | 6.71 | 5.78 | 6.14 | 9.12 | 9.39 | 9 | + |
| Gzf1 | 4.24 | 3.49 | 4.41 | 15.07 | 15.73 | 16.9 | 6.14 | 6.51 | 5.47 | 14.28 | 17.57 | 15.68 | + |
| Bola2 | 74.54 | 89.44 | 85.67 | 65.26 | 63.16 | 61.23 | 73.49 | 70.59 | 67.02 | 63.32 | 59.07 | 61.92 | + |
| E4f1 | 8.45 | 8.31 | 7.69 | 13.72 | 13.56 | 15.46 | 18.78 | 17.67 | 16.49 | 19.5 | 21.31 | 15.08 | + |
| Slc19a1 | 55.28 | 51.48 | 54.42 | 22.93 | 22.14 | 21.99 | 21.47 | 28.97 | 23.7 | 17.62 | 17.6 | 15.3 | + |
| Phyhd1 | 4.99 | 4.06 | 3.98 | 22.82 | 25.94 | 25.81 | 7.4 | 13.74 | 11.51 | 7.37 | 10.3 | 7.34 | + |
| Nup88 | 89.6 | 91.1 | 87.2 | 52.92 | 56.43 | 59.19 | 47.93 | 56.35 | 55.55 | 47.18 | 44.32 | 39.06 | + |
| Slc25a4 | 326.84 | 315.99 | 324.7 | 260.39 | 253.51 | 260.72 | 306.22 | 266.69 | 285.62 | 268.5 | 279.67 | 281.52 | + |
| Acot13 | 45.52 | 39.61 | 41.85 | 64.86 | 65.06 | 61.08 | 82.26 | 69.67 | 67.19 | 65.96 | 62.71 | 73.02 | + |
| Tmem223 | 40.15 | 46.52 | 43.08 | 32.19 | 31.63 | 32.82 | 30.47 | 36.6 | 34.99 | 30.73 | 29.99 | 31.52 | + |
| Rpl4 | 2605.69 | 2567.06 | 2661.16 | 3526.62 | 3618.24 | 3713.98 | 3350.04 | 3408.02 | 3364.38 | 3422.64 | 3807.72 | 3368.21 | + |
| Ube2n | 63.29 | 66.71 | 62.4 | 35.04 | 36.36 | 34.45 | 32.47 | 36.6 | 36.71 | 29.65 | 32.52 | 31.75 | + |
| Nsfl1c | 82.87 | 82.09 | 82.53 | 49.74 | 50.09 | 50.3 | 41.34 | 59.61 | 54.1 | 39.65 | 42.95 | 40.76 | + |
| Cstf3 | 25.46 | 25.31 | 25.19 | 13.6 | 13.88 | 17.11 | 16.77 | 16.49 | 18.07 | 14.02 | 20.28 | 15.69 | + |
| Pkp2 | 0.39 | 0.28 | 0.32 | 1.22 | 1.4 | 0.62 | 5.24 | 2.68 | 3.3 | 3.96 | 4.56 | 4.89 | + |
| 2700094K13Rik | 183.92 | 202.29 | 185.14 | 147.93 | 147.79 | 139.56 | 135.31 | 132.55 | 121.79 | 131.48 | 144.85 | 120 | + |
| Bccip | 75.62 | 82.86 | 70.07 | 43.56 | 45.38 | 45.38 | 39.53 | 47.42 | 46 | 31.34 | 35.43 | 32.41 | + |
| Nelfcd | 29.96 | 32.04 | 28.68 | 27.62 | 26.02 | 25.07 | 23.75 | 23.77 | 23.16 | 21.49 | 22.39 | 21.9 | + |
| Supt4a | 133.97 | 124.67 | 124.22 | 130.09 | 124.45 | 126.98 | 195.25 | 142.73 | 139.59 | 229.98 | 233.04 | 241.54 | + |
| Zfp383 | 0.28 | 0.22 | 0.39 | 0.77 | 0.76 | 0.8 | 0.79 | 1.28 | 0.72 | 0.85 | 0.89 | 1.21 | + |
| Mmp11 | 0.61 | 0.73 | 0.79 | 3.49 | 3.14 | 2.16 | 2.94 | 2.43 | 2.96 | 2.72 | 4.17 | 3.67 | + |
| Tor2a | 39.84 | 38.87 | 40.36 | 49.81 | 52.72 | 51.99 | 51.71 | 54.22 | 53.11 | 53.64 | 55.13 | 49.23 | + |
| Tmem248 | 7.2 | 7.73 | 8.25 | 5.25 | 5.62 | 5.21 | 3.66 | 5.49 | 5.65 | 5.67 | 6.2 | 4.02 | + |
| Npnt | 1.85 | 1.09 | 1.56 | 0.4 | 1.22 | 0.45 | 0.39 | 1.01 | 0.66 | 0.8 | 0.87 | 0.75 | + |
| Dnpep | 41.26 | 40.6 | 41.76 | 55.68 | 50.14 | 56.29 | 64.34 | 57.79 | 55.58 | 58.71 | 65.35 | 57.31 | + |
| Hypk | 143.18 | 142.47 | 134.36 | 90.2 | 91.27 | 88.16 | 79.16 | 95.2 | 89 | 64.5 | 69.39 | 60.51 | + |
| Iffo1 | 3.25 | 3.22 | 2.67 | 4.71 | 4.98 | 6.24 | 4.38 | 4.54 | 4.07 | 4.47 | 5.61 | 5.35 | + |
| Cpeb2 | 0.21 | 0.11 | 0.14 | 0.59 | 0.71 | 0.62 | 0.37 | 0.52 | 0.36 | 0.8 | 0.87 | 0.61 | + |
| Aup1 | 110.44 | 110.79 | 108.49 | 147.28 | 133.85 | 143.16 | 144.28 | 141.64 | 139.54 | 153.58 | 151.09 | 144.04 | + |
| Ltbp4 | 0.36 | 0.26 | 0.36 | 0.79 | 0.72 | 0.76 | 0.81 | 0.78 | 0.65 | 0.76 | 0.8 | 0.76 | + |
| Chchd10 | 159.47 | 158.49 | 174.18 | 186.81 | 228.56 | 219.64 | 166.7 | 242.25 | 212.45 | 109.05 | 122.55 | 115.47 | + |
| Atpaf2 | 30.11 | 28.78 | 29.64 | 18 | 15.98 | 16.7 | 16.82 | 16.99 | 16.35 | 14.78 | 13.07 | 15.48 | + |
| Ewsr1 | 220.06 | 236.74 | 240.78 | 158.36 | 163.21 | 161.03 | 132.96 | 161.91 | 156.15 | 130.41 | 125.88 | 131.26 | + |
| Itgae | 0.13 | 0.11 | 0.16 | 0.58 | 0.65 | 0.55 | 0.57 | 0.4 | 0.48 | 0.3 | 0.6 | 0.48 | + |
| Igflr1 | 7.89 | 8.1 | 8.16 | 32.61 | 22.63 | 20.65 | 17.64 | 11.98 | 17.82 | 25.52 | 16.2 | 27.7 | + |
| Kif20a | 35.1 | 33.6 | 31.13 | 18.81 | 28.35 | 26.28 | 13.39 | 11.87 | 10.99 | 16.11 | 14 | 10.21 | + |
| Dynll1 | 75.49 | 82.84 | 79.83 | 31.7 | 33.32 | 36.63 | 32.02 | 35.69 | 37.78 | 25.15 | 24.02 | 25.52 | + |
| Snrpd2 | 194.2 | 204.49 | 192.56 | 131.34 | 130.09 | 129.24 | 155.32 | 142.38 | 149.1 | 135.29 | 160.24 | 136.67 | + |
| Cep170 | 4.02 | 3.47 | 4.28 | 5.11 | 5.77 | 5.18 | 7.73 | 6.73 | 8.28 | 7.19 | 7.48 | 6.77 | + |
| Gjc3 | 1.96 | 2.17 | 2.5 | 3.29 | 3.67 | 3.32 | 3.18 | 3.11 | 3.31 | 3.62 | 3.69 | 3.7 | + |
| Scaper | 1.7 | 2.26 | 2.05 | 4.67 | 4.63 | 4.06 | 4.09 | 3.16 | 3.38 | 4.18 | 3.44 | 3.47 | + |
| Pmpcb | 77.44 | 73.76 | 75.53 | 56.63 | 57.06 | 57.42 | 48.2 | 59.43 | 52.65 | 50.04 | 52.97 | 43.75 | + |
| Atp1b3 | 144.1 | 147.86 | 146.84 | 165.95 | 160.1 | 159.87 | 123.88 | 113.39 | 110.68 | 197.43 | 177.99 | 197.41 | + |
| Psme1 | 342.38 | 351.04 | 341.36 | 288.38 | 284.57 | 293.57 | 287.21 | 278.22 | 288.08 | 289.11 | 275.4 | 313.99 | + |
| Mapk8ip3 | 10.53 | 9.27 | 10.42 | 22.14 | 22.41 | 23.33 | 22.24 | 19.61 | 20.74 | 27 | 27.89 | 25.49 | + |
| Hnrnpm | 151.01 | 163.84 | 161.02 | 64.22 | 67.06 | 63.43 | 53.66 | 62.11 | 57.32 | 46.59 | 41.95 | 51.15 | + |
| Gpn1 | 43.1 | 44.05 | 42.27 | 26.36 | 25.58 | 24.67 | 22.37 | 26.93 | 24.69 | 21.27 | 19.09 | 18.51 | + |
| Cox19 | 44.61 | 42.75 | 43.48 | 28.99 | 26.61 | 28.6 | 32.77 | 30.02 | 30.95 | 26.16 | 26.64 | 27.72 | + |
| Cetn3 | 97.07 | 101.09 | 87.37 | 46.09 | 44.42 | 52.49 | 54.12 | 57.92 | 58.25 | 46.45 | 46.35 | 40.17 | + |
| Stub1 | 64.6 | 71.2 | 72.31 | 61.5 | 57.07 | 60.76 | 47.27 | 47.99 | 49.45 | 47.38 | 43.11 | 51.3 | + |
| Nol12 | 23.63 | 22.64 | 21.8 | 13.61 | 15.44 | 15.95 | 14.59 | 16.62 | 15.76 | 15.12 | 12.85 | 14.64 | + |
| Ppp3r1 | 32.08 | 34.66 | 33.87 | 23.02 | 26.03 | 23.78 | 22 | 27.88 | 25.84 | 21.86 | 21.74 | 20.93 | + |

| | | | | | | | | | | | | | |
|---|---|---|---|---|---|---|---|---|---|---|---|---|---|
| Snapc5 | 30.89 | 31.92 | 31.23 | 15.2 | 15.59 | 15.59 | 14.42 | 19.53 | 18.89 | 16.54 | 14.25 | 19.24 | + |
| Mfsd11 | 18.62 | 20.41 | 20.62 | 11.76 | 11.64 | 11.29 | 18.5 | 13.15 | 12.46 | 14.15 | 15.01 | 14.95 | + |
| C1d | 26.03 | 26.76 | 24.68 | 16.2 | 16.36 | 16.58 | 16.87 | 16.84 | 17.75 | 15.44 | 14.63 | 15.58 | + |
| Mta3 | 18.19 | 19.93 | 20.75 | 31.62 | 29.08 | 30.14 | 34.57 | 27.22 | 26.36 | 30.73 | 31.76 | 34.89 | + |
| Siah1b | 6.47 | 8.39 | 6.17 | 3.88 | 3.2 | 2.46 | 1.45 | 1.96 | 2.81 | 2.18 | 1.82 | 2.06 | + |
| Idh3a | 132.23 | 128.44 | 136.04 | 54.55 | 57.69 | 62.53 | 49.36 | 65.77 | 60.66 | 35.91 | 35.63 | 38.13 | + |
| Glrx5 | 9.88 | 10.7 | 13.08 | 6.18 | 5.84 | 5.76 | 6.55 | 7.26 | 7.38 | 5.6 | 4.87 | 4.83 | + |
| Kcnk7 | 3.12 | 2.4 | 2.66 | 4.75 | 3.13 | 3.77 | 11.16 | 6.05 | 10.57 | 6.5 | 7.78 | 6.5 | + |
| Timmdc1 | 16.07 | 16.08 | 16.51 | 11.8 | 11.01 | 12.87 | 9.37 | 11.62 | 12.2 | 9.19 | 9.44 | 10.38 | + |
| Ecm1 | 1.47 | 0.58 | 0.86 | 26.59 | 20.46 | 23.06 | 8.79 | 7.8 | 8.29 | 19.39 | 22.55 | 23.01 | + |
| Extl3 | 5.9 | 6.7 | 6.39 | 4.24 | 3.73 | 4.98 | 3.74 | 4.16 | 4.17 | 3.92 | 3.52 | 3.62 | + |
| Slc25a23 | 0.76 | 0.4 | 0.35 | 1.26 | 1.69 | 1.32 | 0.89 | 1.29 | 1.85 | 1.7 | 1.46 | + |
| Itga4 | 7.44 | 6.61 | 8.04 | 6.39 | 5.41 | 5.75 | 16.07 | 13.6 | 14.49 | 15.47 | 14.9 | 15.55 | + |
| E130309D02Rik | 6.09 | 7.15 | 6.46 | 3.56 | 4.26 | 4.3 | 3.29 | 5.39 | 4.14 | 4.2 | 4.18 | 3.05 | + |
| Snx17 | 49.89 | 54.05 | 52.51 | 47.18 | 45.28 | 43.94 | 43.09 | 41.39 | 43.14 | 40.05 | 38.52 | 46.03 | + |
| Tspan9 | 1.32 | 1.08 | 0.92 | 6.05 | 4.78 | 4.97 | 7.14 | 2.93 | 7.09 | 10.46 | 8.29 | 12.17 | + |
| Fam217b | 0.08 | 0.15 | 0.12 | 0.59 | 0.85 | 0.51 | 0.8 | 0.52 | 0.47 | 1.19 | 1.14 | 1.15 | + |
| Tmie | 0.96 | 0.93 | 1.2 | 3.61 | 4.71 | 3.32 | 4.69 | 2.58 | 2.69 | 4.51 | 5.72 | 4.94 | + |
| Rbmx2 | 17.49 | 20.19 | 19.82 | 12.47 | 10.92 | 11.09 | 9.2 | 9.84 | 9.54 | 6.66 | 7.53 | 5.75 | + |
| Sema4f | 2.57 | 1.72 | 2.92 | 6.47 | 6.17 | 6.37 | 14.35 | 8.85 | 11.87 | 14.48 | 14.23 | 16.93 | + |
| Elovl5 | 50.29 | 54.38 | 48.73 | 88.61 | 76.54 | 83.65 | 73.67 | 65.83 | 68.45 | 85.21 | 83.18 | 91.84 | + |
| S100a10 | 912.18 | 789.18 | 814.71 | 908.77 | 975.86 | 1011.06 | 1319.43 | 1244.74 | 1218.76 | 1139.65 | 1207.5 | 1164.58 | + |
| Tmem167b | 10.17 | 8.33 | 7.88 | 16.11 | 15.39 | 16.85 | 17.56 | 15.98 | 16.9 | 17.71 | 24.84 | 23.19 | + |
| Pmaip1 | 8.35 | 8.49 | 8.37 | 11.49 | 10.84 | 11.96 | 13.25 | 10.56 | 10.16 | 10.8 | 12.14 | 11.77 | + |
| Tpgs2 | 4.68 | 5.19 | 4.51 | 2.67 | 2.44 | 2.89 | 2.45 | 2.68 | 3.02 | 2.73 | 2.58 | 2.86 | + |
| Dhx16 | 22.73 | 25.5 | 24.12 | 13.63 | 12.67 | 13.62 | 17.22 | 15.04 | 17.34 | 17.38 | 14.27 | 20.08 | + |
| Wbp5 | 30.56 | 37.19 | 29.8 | 52.79 | 47.07 | 49.43 | 56 | 49.27 | 54.76 | 54.28 | 60.96 | 56.96 | + |
| Psmb4 | 279.4 | 273.66 | 272.85 | 138.73 | 155.57 | 154.73 | 144.33 | 178.25 | 154.79 | 132.65 | 125.02 | 130.53 | + |
| Casc4 | 23.32 | 25.84 | 24.3 | 8.96 | 11.13 | 9.39 | 9.59 | 12.21 | 10.46 | 8.05 | 9.4 | 8.16 | + |
| Ebf1 | 4.42 | 3.14 | 3.37 | 3.6 | 5.76 | 5.03 | 8.65 | 12 | 11.93 | 7.95 | 8.24 | 8.17 | + |
| Tfrc | 140.15 | 143.86 | 160.48 | 28.08 | 34.58 | 32.24 | 15.41 | 26.89 | 22.38 | 12.58 | 12.44 | 11.69 | + |
| Ddx54 | 34.46 | 34.81 | 34.9 | 27.62 | 25.15 | 21.61 | 28.82 | 28.55 | 30.33 | 24.32 | 26.39 | 26.82 | + |
| Psmd1 | 105.57 | 107.91 | 103.5 | 35.32 | 42.66 | 40.13 | 28.97 | 40.47 | 35.08 | 28.07 | 29.85 | 25.79 | + |
| Wipi1 | 1.1 | 0.67 | 0.51 | 4 | 3.16 | 2.91 | 5.66 | 4.04 | 3.25 | 5.54 | 7.56 | 7.77 | + |
| Mrpl55 | 45.3 | 45.1 | 44.78 | 26.67 | 25.45 | 25.67 | 23.87 | 28.09 | 26.76 | 20.39 | 16.28 | 18.09 | + |
| Plcd1 | 0.62 | 0.46 | 0.24 | 9.04 | 8.39 | 7.8 | 8.48 | 2.9 | 5.14 | 15.2 | 13.72 | 14.25 | + |
| Thrap3 | 36.68 | 38.19 | 37.09 | 25.85 | 24.87 | 25.01 | 23.28 | 25.94 | 23.83 | 23.77 | 21.68 | 21.67 | + |
| Hvcn1 | 29.78 | 27.15 | 26.93 | 25.93 | 23.14 | 25.48 | 27.73 | 23.95 | 27.04 | 34.74 | 40.99 | 42.2 | + |
| Dnajc2 | 81.61 | 83.95 | 81.9 | 57.85 | 61.02 | 61.17 | 51.29 | 66.08 | 60.46 | 46.39 | 60.03 | 46.69 | + |
| Hdac1 | 137.12 | 150.58 | 148.38 | 75.19 | 77.56 | 74.63 | 63.91 | 74.81 | 68.91 | 60.08 | 55.64 | 62.29 | + |
| Rpl22l1 | 484.54 | 491.58 | 487.66 | 872.94 | 946.5 | 949.3 | 870.42 | 822.1 | 820.74 | 938.4 | 1040.3 | 947.34 | + |
| Pde4b | 5.66 | 5.14 | 6.66 | 34.11 | 32.07 | 32.27 | 18.73 | 16.9 | 15.59 | 32.24 | 34.47 | 33.5 | + |
| Dhx15 | 109.36 | 106.97 | 114.29 | 72.55 | 74.59 | 73.46 | 64.2 | 73.6 | 71.89 | 65.16 | 76.63 | 60.48 | + |
| Thoc5 | 27.84 | 26.61 | 25.5 | 21.27 | 20.75 | 22.8 | 19.59 | 20.5 | 20.47 | 19.67 | 19 | 15.58 | + |
| Cox14 | 73.51 | 72.51 | 72.84 | 92.49 | 86.6 | 104.78 | 97.52 | 107.48 | 110.34 | 105.23 | 108.86 | 94.4 | + |
| Vps52 | 25.99 | 25.67 | 22.56 | 20.29 | 17.08 | 19.64 | 16.91 | 16.22 | 16.93 | 17.39 | 17.71 | 17.58 | + |
| Mvd | 50.48 | 56.31 | 51.74 | 74.05 | 77.4 | 79.27 | 80.92 | 65.93 | 70.07 | 61.37 | 72.54 | 61.48 | + |
| Zfp239 | 8.32 | 7.75 | 7.91 | 4.57 | 3.6 | 3.3 | 3.12 | 3.63 | 3.74 | 2.67 | 2.4 | 2.84 | + |
| Aldh6a1 | 0.82 | 0.78 | 1.13 | 5.16 | 3.15 | 3.93 | 3.18 | 1.77 | 1.99 | 3.3 | 4 | 3.26 | + |
| Zfp280c | 7.66 | 6.89 | 7.28 | 5.53 | 4.85 | 5.45 | 4.05 | 4.71 | 4.38 | 4.6 | 5.13 | 4.05 | + |
| Emg1 | 184.65 | 182.88 | 184.32 | 112.09 | 117.81 | 115.89 | 119.91 | 111.18 | 119.18 | 108.84 | 115.87 | 107.36 | + |
| Rps21 | 456.42 | 503.71 | 473.79 | 730.81 | 687.15 | 720.17 | 757.31 | 654.72 | 670.15 | 800.48 | 840.11 | 805.31 | + |
| Txn1 | 535.27 | 551.36 | 548.81 | 265.26 | 288.63 | 292.58 | 250.8 | 302.9 | 272.55 | 228.5 | 232.98 | 222.58 | + |
| Prf1 | 394.81 | 331.03 | 309.66 | 106.21 | 124.73 | 121.47 | 155.65 | 179.56 | 165.18 | 224.23 | 165.94 | 259.33 | + |
| Yme1l1 | 44.48 | 42.21 | 45.68 | 37.28 | 38.38 | 38.54 | 33.83 | 38.51 | 36.3 | 36.1 | 37.31 | 33.31 | + |
| Sptssa | 62.8 | 67.14 | 64.26 | 47.81 | 48.09 | 50.25 | 50.18 | 48.69 | 49.8 | 46.88 | 50.25 | 48.85 | + |
| 2810407C02Rik | 7.05 | 8.93 | 7.29 | 5.95 | 5.38 | 5.11 | 4.8 | 4.42 | 5.09 | 5.36 | 4.64 | 5.87 | + |
| Eny2 | 25.49 | 29.32 | 25.81 | 21.36 | 21.31 | 23.8 | 18.33 | 17.19 | 18.28 | 18.72 | 19.43 | 17.82 | + |
| Slc41a2 | 0.27 | 0.04 | 0.26 | 0.59 | 0.88 | 0.79 | 1.82 | 0.96 | 1.79 | 2.22 | 1.81 | 2.23 | + |
| Sdha | 164.89 | 166.92 | 167.05 | 152.17 | 150.59 | 145.28 | 140.31 | 134.9 | 131.53 | 129.8 | 124.32 | 126.03 | + |
| Lgals3 | 201.88 | 166.68 | 157.96 | 102.13 | 85.19 | 90.55 | 455.98 | 369.18 | 386.58 | 257.96 | 264.85 | 296.95 | + |
| Cycs | 160.35 | 160.28 | 154.27 | 56.97 | 63.62 | 65.02 | 74.18 | 95.82 | 89.15 | 51.04 | 48.93 | 51.31 | + |
| Use1 | 47.39 | 48.83 | 49.76 | 102.84 | 97.56 | 94.76 | 86.79 | 85.28 | 90.23 | 111.8 | 103.86 | 120.3 | + |
| 2410006H16Rik | 236.44 | 218.75 | 249.23 | 1296.39 | 1248.84 | 1343.56 | 1030.71 | 1050.85 | 1018.02 | 1471.83 | 1733.77 | 1476.58 | + |
| Cnot6 | 4.52 | 4.51 | 4.39 | 2.75 | 3.59 | 3.49 | 3.22 | 3.41 | 3.15 | 3.04 | 3.1 | 2.88 | + |
| Ptpn2 | 87.93 | 88.02 | 90.84 | 120.62 | 133.54 | 134.65 | 123.53 | 118.08 | 124.14 | 134.87 | 140.36 | 133.33 | + |
| Mogs | 9.23 | 10.94 | 9.54 | 5.25 | 4.44 | 5.38 | 3.98 | 4.86 | 4.16 | 4.79 | 3.16 | 4 | + |
| Gdpd5 | 3.15 | 2.74 | 1.93 | 6.3 | 4.7 | 5.39 | 7.15 | 5.48 | 5.23 | 5.99 | 6.5 | 6.44 | + |
| Casp8ap2 | 4.04 | 4.68 | 4.54 | 3.44 | 2.97 | 3.48 | 2.55 | 2.47 | 2.73 | 3 | 3.11 | 3.15 | + |
| Pdxk | 11.01 | 10.85 | 11.2 | 6.94 | 7.54 | 7.69 | 7.62 | 10.23 | 8.34 | 7.67 | 8.45 | 8.27 | + |
| Pdcd11 | 10.97 | 11.13 | 11.95 | 5.32 | 4.8 | 5 | 3.54 | 5.27 | 5.35 | 3.77 | 3.35 | 3.17 | + |
| Ccdc50 | 24.55 | 21.8 | 23.05 | 11.4 | 13.16 | 13.8 | 10.42 | 14.21 | 12.85 | 9.91 | 9.6 | 9.26 | + |
| Ttc38 | 1.86 | 1.32 | 1.95 | 3.23 | 4.11 | 3.53 | 5.4 | 3.75 | 3.36 | 4.73 | 4.18 | 3.34 | + |
| Myh10 | 0.85 | 0.5 | 0.76 | 0.15 | 0.2 | 0.15 | 0.21 | 0.18 | 0.21 | 0.07 | 0.13 | 0.07 | + |
| Gtf2h5 | 51.42 | 49.74 | 51.34 | 30.33 | 33.37 | 33.66 | 29.64 | 31.1 | 34.33 | 28.29 | 27.67 | 24.91 | + |
| Cd74 | 6.14 | 5.69 | 5.38 | 2.96 | 4.09 | 3.25 | 5.58 | 7.79 | 7.9 | 2.13 | 1.54 | 1.4 | + |
| Lpgat1 | 2.62 | 2.68 | 3.24 | 1.92 | 1.88 | 1.4 | 2.19 | 2.03 | 1.73 | 1.8 | 1.85 | 1.71 | + |
| Mllt11 | 7.55 | 9.19 | 9.89 | 6.33 | 5.22 | 5.72 | 5.8 | 6.74 | 6.81 | 5.68 | 6.07 | 6.13 | + |
| Apex2 | 14.05 | 14.08 | 11.36 | 8.32 | 6.92 | 6.26 | 4.79 | 6.04 | 6.63 | 6.6 | 5.13 | 7 | + |
| Abcg3 | 3.26 | 2.82 | 3.53 | 5.5 | 7.16 | 7.48 | 10.23 | 6.99 | 7.77 | 6.43 | 6.52 | 7.28 | + |
| Zkscan17 | 10.29 | 9.95 | 9.69 | 18 | 17.34 | 17.25 | 21.59 | 15.98 | 18.57 | 21.12 | 18.61 | 20.2 | + |
| Tmem167 | 15.53 | 15.14 | 15.03 | 10.46 | 10.61 | 11.71 | 11.91 | 12.11 | 12.66 | 10.33 | 10.58 | 10.5 | + |
| Etf1 | 62.75 | 72.3 | 72.99 | 38.6 | 44.88 | 45.65 | 38.55 | 49.31 | 44.37 | 36.9 | 44.05 | 34.98 | + |
| Sec24a | 7.26 | 7.34 | 7.53 | 13.17 | 12.02 | 13.13 | 12.84 | 10.85 | 12.67 | 13.92 | 15.82 | 14.85 | + |
| Socs1 | 50.76 | 82.31 | 81.68 | 136 | 127.84 | 122.36 | 51.6 | 67.17 | 73.26 | 36.98 | 37.59 | 35.39 | + |
| Utp6 | 23.25 | 24.16 | 24 | 12.82 | 13.56 | 11.91 | 8.74 | 12.04 | 12.43 | 7.11 | 8.27 | 7.36 | + |
| Eif1ax | 187.25 | 201.7 | 195.24 | 134.05 | 136.17 | 135.37 | 118.79 | 135.19 | 129.06 | 102.04 | 129.43 | 107.94 | + |
| Fscn1 | 0.43 | 0.37 | 0.53 | 0.62 | 0.8 | 0.69 | 2.54 | 1.23 | 1.99 | 1.23 | 1.18 | 1.62 | + |
| Psenen | 72.2 | 62.75 | 68.28 | 102.48 | 95.8 | 103.51 | 96.64 | 95.33 | 95.33 | 104.28 | 114.82 | 119.07 | + |
| Gltscr2 | 151.84 | 158.87 | 150.69 | 326.6 | 291.15 | 314.64 | 334.97 | 271.18 | 301.02 | 370.12 | 376.7 | 386.84 | + |
| Dpy30 | 124.51 | 132.04 | 134.79 | 61.12 | 66.72 | 72.17 | 68.87 | 75.14 | 74.96 | 58.97 | 58.78 | 60.74 | + |
| Polr2g | 125.87 | 121.85 | 127.28 | 84.98 | 86.8 | 100.96 | 93.15 | 105.22 | 108.78 | 93.72 | 97.98 | 94.46 | + |
| 1110004F10Rik | 157.62 | 162.55 | 152.29 | 105.91 | 115.44 | 113.42 | 92.39 | 115.44 | 108.03 | 93.08 | 92.59 | 99.81 | + |
| Tex10 | 28.47 | 26.99 | 26.29 | 23.21 | 21.46 | 25.01 | 22.79 | 20.89 | 19.25 | 23.12 | 22.28 | 23.18 | + |
| Cdk2 | 31.53 | 31.67 | 34.02 | 20.3 | 24.53 | 22.7 | 16.38 | 18.76 | 18.69 | 16.36 | 16.62 | 16.8 | + |
| Derl2 | 17.24 | 17.72 | 17.14 | 14.32 | 13.04 | 13.25 | 14.36 | 14.01 | 13.79 | 11.95 | 12.93 | 13.92 | + |
| Tlr1 | 2 | 3.05 | 1.58 | 2.19 | 2.47 | 2.64 | 1.32 | 2.25 | 1.73 | 0.81 | 0.93 | 0.8 | + |
| Med15 | 25.23 | 23.41 | 26.11 | 36.68 | 36.22 | 35.61 | 37.35 | 42.79 | 36.73 | 39.05 | 48.92 | 38.87 | + |
| 2310039H08Rik | 15.95 | 13.55 | 14.41 | 20.85 | 24.31 | 21.78 | 27.13 | 28.34 | 26.17 | 19.98 | 25.83 | 18.12 | + |
| Nhp2l1 | 324.07 | 326.22 | 320.2 | 151 | 150.8 | 152.24 | 131.58 | 173.15 | 159.49 | 113.18 | 113.6 | 113 | + |
| Rpl15 | 845.76 | 869.69 | 858.24 | 1062.06 | 1097.77 | 1133.21 | 981.16 | 1029.46 | 996.54 | 988.05 | 1094.79 | 976.11 | + |
| Suv39h1 | 31.66 | 31.02 | 30.41 | 20.63 | 23.92 | 23.71 | 17.74 | 18.55 | 18.4 | 16.17 | 16.32 | 15.56 | + |
| Slc4a7 | 9.14 | 6.72 | 9.2 | 4.05 | 4.06 | 3.88 | 6.5 | 6.81 | 7.48 | 6.09 | 4.94 | 4.84 | + |
| Gigyf1 | 0.65 | 0.77 | 0.85 | 1.91 | 2 | 2.13 | 2.16 | 1.87 | 1.75 | 3.24 | 3.05 | 3.13 | + |
| Gm8773 | 4.86 | 5.69 | 6.26 | 3.89 | 3.69 | 4.73 | 3.26 | 3.1 | 2.79 | 0.54 | 0.87 | 1.27 | + |
| Rap1b | 24.61 | 24.72 | 25.07 | 22.03 | 18.88 | 18.4 | 15.49 | 17.26 | 15.83 | 19.26 | 16.32 | 17.97 | + |
| Camkk1 | 1.9 | 1.65 | 1.69 | 4.89 | 3.87 | 3.95 | 5 | 3.52 | 4.04 | 4.03 | 5.8 | 4.63 | + |
| Elmsan1 | 2.22 | 1.9 | 2.25 | 3.93 | 3 | 3.47 | 2.75 | 2.69 | 2.75 | 2.74 | 2.76 | 3.27 | + |
| Gm10845 | 13.49 | 14.89 | 14.77 | 26.87 | 25.63 | 27.2 | 33.16 | 28.99 | 27.79 | 37.74 | 41.21 | 36.41 | + |
| Ccdc43 | 23.21 | 21.92 | 20.57 | 15.29 | 17.75 | 15.89 | 15.02 | 17.41 | 16.62 | 15.11 | 17.34 | 15.98 | + |
| Tyw1 | 19.06 | 20.38 | 18.03 | 14.36 | 15.8 | 15.47 | 15.6 | 16.31 | 17.8 | 13.39 | 17.09 | 14.7 | + |
| Serf2 | 60.32 | 61.61 | 60.43 | 80.44 | 75.44 | 84.34 | 90.77 | 83.79 | 85.61 | 82.6 | 85.85 | 89.59 | + |
| Rmi2 | 6.19 | 6.73 | 6.58 | 4.61 | 4.91 | 4.44 | 5.05 | 4.91 | 4.64 | 3.96 | 4.31 | 4.17 | + |
| Slc7a14 | 4.19 | 4.37 | 4.28 | 1.69 | 1.91 | 2.16 | 2.06 | 2.1 | 2.08 | 1.99 | 1.9 | 1.77 | + |
| Exoc2 | 23.44 | 24.38 | 23.93 | 29.06 | 29.42 | 28.39 | 36.96 | 37.6 | 36.43 | 32.77 | 39.3 | 34.04 | + |
| BC055324 | 10 | 8.24 | 7.82 | 5.2 | 5.62 | 5.72 | 5.16 | 5.73 | 4.14 | 4.08 | 4.65 | 3.37 | + |
| Ndufa10 | 159.86 | 161.43 | 152.47 | 128.13 | 127.81 | 128.49 | 138.17 | 139.32 | 146.35 | 121.18 | 121.68 | 115.95 | + |
| Rabgap1l | 70.79 | 53.79 | 66.72 | 71.14 | 86.83 | 86.27 | 80.34 | 99.72 | 94.6 | 76.31 | 85.24 | 79.36 | + |
| Usp18 | 12.25 | 13.07 | 12.85 | 7.95 | 9.94 | 9.54 | 2.27 | 3.49 | 3.7 | 6.27 | 5.71 | 8.66 | + |
| Cml1 | 1.02 | 1.55 | 1.18 | 2.88 | 2.53 | 3.34 | 3.87 | 2.56 | 2.87 | 3.56 | 4.08 | 4.45 | + |
| Txnl4a | 184.85 | 183.61 | 182.7 | 113.1 | 116.19 | 112.95 | 98.73 | 124.29 | 114.48 | 99.72 | 89.6 | 96.45 | + |
| Isy1 | 65.22 | 67.74 | 65.45 | 35.66 | 38.59 | 38.83 | 51.68 | 49.89 | 50.31 | 49.96 | 41.64 | 47.62 | + |
| Tmem108 | 2.81 | 3.23 | 3.05 | 2.93 | 3.21 | 3.28 | 9.79 | 7.37 | 8.73 | 6.1 | 5.76 | 6.32 | + |
| Pold4 | 24.45 | 19.69 | 23.2 | 79.47 | 75.75 | 72.78 | 95.61 | 82.47 | 79.56 | 93.68 | 111.79 | 108.95 | + |
| Cox5b | 387.48 | 395.6 | 371.73 | 254.26 | 252.16 | 254.74 | 261.7 | 297.08 | 298.75 | 231.33 | 232.87 | 236.04 | + |
| Xist | 35.22 | 29.44 | 30.79 | 64.49 | 67.35 | 66.48 | 53.51 | 49.29 | 50.68 | 67.38 | 70.28 | 61.01 | + |
| Dtd2 | 8.64 | 7.35 | 7.09 | 5.15 | 5.45 | 6.79 | 4.87 | 6.16 | 5.02 | 4.8 | 5.07 | 5.76 | + |
| Zbtb32 | 270.61 | 247.91 | 246.38 | 135.06 | 152.91 | 163.34 | 157.56 | 210.34 | 192.07 | 146.2 | 138.03 | 137.83 | + |
| Ddx47 | 74.86 | 76.55 | 77.2 | 59.37 | 53.08 | 56.85 | 56.27 | 57.56 | 61.03 | 59.27 | 64.71 | 57.01 | + |
| Tspan13 | 130.92 | 122.27 | 140.35 | 250.67 | 248.74 | 259.27 | 201.77 | 222.89 | 217.06 | 201.81 | 223.88 | 212.57 | + |
| BC068157 | 0.77 | 0.84 | 0.78 | 1.82 | 1.33 | 1.48 | 3.24 | 1.74 | 1.57 | 1.8 | 2.43 | 1.72 | + |
| Pds5a | 5.93 | 7.41 | 7.55 | 5 | 5.44 | 5.26 | 4.38 | 5.23 | 5.23 | 4.9 | 5.35 | 4.3 | + |
| Gp49a | 45.21 | 48.41 | 51.97 | 88.57 | 83.66 | 76.21 | 46.62 | 53.83 | 50.55 | 20.24 | 20.37 | 20.63 | + |
| Uba52 | 1428.23 | 1517.25 | 1476.68 | 2431.57 | 2270.04 | 2375.44 | 2457.79 | 2083.74 | 2260.27 | 2694.62 | 2815.54 | 2770.93 | + |
| Fastkd3 | 21.92 | 21.64 | 22.95 | 10.22 | 11.07 | 11.09 | 8.96 | 9.52 | 9.96 | 7.41 | 7.72 | 6.78 | + |
| Ccdc86 | 20.66 | 23.52 | 23.21 | 8.56 | 9.17 | 8.9 | 6.17 | 8.2 | 7.62 | 5.48 | 4.99 | 4.93 | + |
| Arhgef9 | 1.3 | 1.02 | 1.2 | 2.48 | 2.98 | 2.86 | 4.7 | 3.45 | 3.45 | 2.92 | 3.48 | 3.12 | + |
| Yy2 | 3.68 | 3.78 | 3.38 | 1.63 | 1.48 | 2 | 1.66 | 1.82 | 1.98 | 1.02 | 1.07 | 1.39 | + |
| Bcs1l | 12.2 | 9.65 | 9.6 | 18.86 | 17.5 | 18.52 | 16.88 | 15.41 | 18.03 | 18.9 | 19.72 | 18.71 | + |
| Ptp4a2 | 83.54 | 83.94 | 84.47 | 84.06 | 79.19 | 84.42 | 104.54 | 97.93 | 97.16 | 100.12 | 112.54 | 101.65 | + |
| Ost4 | 177.47 | 170.57 | 165.12 | 105.19 | 101.46 | 106.26 | 114.2 | 116 | 112.75 | 88.31 | 103.82 | 102.27 | + |
| Phb2 | 232.13 | 249.38 | 241.65 | 161.6 | 158.8 | 154.32 | 156.16 | 171.43 | 170.97 | 146.38 | 151.49 | 147.26 | + |
| Iars2 | 10.3 | 10.68 | 12.34 | 8.21 | 7.55 | 7.48 | 7.06 | 8.28 | 8.32 | 7.45 | 6.05 | 6.98 | + |
| Nrn1 | 32.32 | 27.22 | 31.25 | 113.68 | 90.91 | 93.4 | 60.13 | 43.45 | 42.38 | 50.14 | 49.15 | 55.25 | + |
| Ube2d3 | 152.8 | 156.36 | 167.64 | 166.59 | 165.53 | 162.37 | 178.83 | 164.43 | 164.91 | 204.73 | 202.18 | 198.23 | + |
| Gyk | 11.47 | 9.32 | 9.49 | 3.79 | 4.3 | 5.65 | 4.26 | 5.02 | 4.64 | 3.15 | 3.15 | 3.63 | + |
| Arpp19 | 22.3 | 25.03 | 23.91 | 14.11 | 14.57 | 14.01 | 13.25 | 14.37 | 15.19 | 10.3 | 10.67 | 12.93 | + |
| Hsf1 | 25.94 | 25.46 | 27.09 | 20.2 | 20.49 | 18.74 | 19.71 | 18.14 | 19.82 | 16.5 | 18.02 | 18.17 | + |
| Cbl | 5.32 | 5.35 | 5.99 | 7.57 | 6.63 | 7.28 | 8.11 | 7.68 | 7.91 | 7.87 | 7.26 | 6.97 | + |
| Srrt | 48.26 | 51.8 | 49.93 | 18.53 | 18.88 | 20.21 | 11.44 | 14.78 | 13.61 | 9.15 | 10.41 | 10.49 | + |
| Usp14 | 52.95 | 53.24 | 51.93 | 30.63 | 31.92 | 30.93 | 29.26 | 31.61 | 33.44 | 25.4 | 26.94 | 24.85 | + |

| | | | | | | | | | | | | | |
|---|---|---|---|---|---|---|---|---|---|---|---|---|---|
| 6430531B16Rik | 0.2 | 0.2 | 0.37 | 1.03 | 1.61 | 0.71 | 2.13 | 1.62 | 1.62 | 1.58 | 2.34 | 1.69 | + |
| Rpl35 | 1554.08 | 1585.05 | 1577.57 | 2250.69 | 2331.87 | 2424.77 | 2163.6 | 2162.33 | 2092.72 | 2312.25 | 2537.51 | 2330.34 | + |
| Uspl1 | 22.82 | 21.35 | 23.84 | 15.1 | 18.38 | 17.06 | 13.68 | 15.13 | 16.28 | 15.29 | 13.77 | 13.79 | + |
| Slc35e1 | 4.15 | 3.91 | 4.4 | 2.62 | 3.36 | 2.99 | 3.21 | 2.72 | 2.69 | 2.95 | 2.64 | 2.72 | + |
| Slbp | 96.07 | 114.41 | 102.56 | 63.74 | 66.52 | 62.69 | 46.79 | 45.73 | 52.59 | 51.92 | 43.79 | 49.7 | + |
| Ccdc61 | 5.79 | 6.2 | 7.32 | 10.69 | 11.99 | 11.96 | 9.76 | 10.11 | 10.33 | 10.36 | 8.71 | 9.06 | + |
| 2310001H17Rik | 6.48 | 6.61 | 7.26 | 12.61 | 10.98 | 12.81 | 14.88 | 13.18 | 11.74 | 15.79 | 16.65 | 16.05 | + |
| Nrf1 | 19.76 | 20.56 | 21.23 | 13.31 | 13.93 | 13.04 | 11.32 | 11.89 | 11.55 | 12.94 | 11.88 | 14.95 | + |
| Mrps5 | 57.2 | 57.38 | 51.69 | 32.09 | 34.04 | 36.37 | 28.73 | 35.27 | 32.92 | 24.33 | 25.96 | 23.47 | + |
| Leprotl1 | 55.3 | 55.11 | 56.21 | 79.2 | 74.55 | 73.28 | 92.93 | 89.78 | 95.91 | 81.34 | 83.45 | 84 | + |
| 9-Sep | 142.68 | 152.25 | 158.4 | 100.44 | 98.82 | 99.34 | 121.3 | 136.38 | 133.64 | 107.43 | 97.77 | 105.6 | + |
| Psmg3 | 53.01 | 47.23 | 53.29 | 27.72 | 32.44 | 26.83 | 26.23 | 29.82 | 29.48 | 21 | 19.54 | 20.82 | + |
| Vmac | 1.46 | 2.57 | 1.94 | 4.75 | 4.48 | 4.64 | 6.66 | 6.55 | 5.09 | 7.59 | 5.57 | 6.31 | + |
| Ruvbl2 | 198.15 | 187.77 | 194.61 | 69.99 | 83.7 | 85.27 | 67.28 | 94.75 | 91.96 | 50.48 | 47.06 | 48.23 | + |
| Trmt5 | 18.09 | 15.86 | 17.2 | 10.79 | 11.78 | 11.54 | 11.4 | 11.89 | 11.5 | 9.6 | 9.49 | 8.46 | + |
| Atp13a1 | 32.24 | 29.52 | 30.05 | 24.14 | 25.55 | 24 | 20.89 | 21.25 | 21.1 | 22.86 | 23.93 | 23.01 | + |
| Gpn3 | 36.77 | 34.86 | 37.91 | 23.3 | 25.69 | 28.61 | 23.32 | 27.31 | 26.4 | 21.83 | 27.39 | 24.96 | + |
| Actn4 | 92.92 | 94.63 | 95.19 | 64.28 | 63.77 | 61.06 | 56.22 | 63.13 | 55.97 | 54.03 | 50.03 | 54.18 | + |
| Med4 | 36.38 | 32.89 | 31.91 | 22.44 | 25.36 | 24.79 | 26.66 | 24.95 | 20.65 | 24.13 | 24.29 | 23.56 | + |
| Neil1 | 6.39 | 7.21 | 6.42 | 9.51 | 8.32 | 8.18 | 13.6 | 12.9 | 11.33 | 11.15 | 7.85 | 11.97 | + |
| C1qtnf4 | 0.71 | 0.43 | 0.28 | 1.16 | 1.42 | 1.82 | 2.99 | 2.02 | 2.28 | 2.05 | 2.17 | 2.24 | + |
| Def8 | 7.65 | 7.96 | 7.42 | 6.27 | 4.74 | 4.71 | 5.13 | 6.4 | 5.13 | 5 | 5.54 | 6.77 | + |
| Dnajc11 | 59.86 | 54.72 | 54.86 | 22.5 | 21.82 | 23.26 | 17.64 | 24.78 | 21.11 | 14.83 | 11.79 | 14.45 | + |
| Bbx | 3.27 | 3.03 | 3.11 | 6.86 | 6.58 | 6.92 | 5.93 | 5.79 | 5.94 | 6.87 | 7.56 | 5.89 | + |
| Mad1l1 | 12.07 | 11.06 | 12.06 | 6.18 | 6.41 | 7.01 | 5.35 | 5.83 | 4.84 | 3.64 | 4.44 | 4.67 | + |
| Tnpo1 | 25.72 | 22.76 | 22.74 | 14.69 | 15.84 | 15.91 | 12.78 | 16.81 | 16.21 | 11.79 | 14.52 | 10.56 | + |
| Asap1 | 1.11 | 1.63 | 1.74 | 3.98 | 2.95 | 2.55 | 3.23 | 2.2 | 2.33 | 3.11 | 2.85 | 3 | + |
| Sorl1 | 0.63 | 0.96 | 0.57 | 2.23 | 1.51 | 2.16 | 2.69 | 1.54 | 1.66 | 2.58 | 2.52 | 2.5 | + |
| Rpl37 | 509.77 | 535.45 | 539.81 | 782.59 | 805.63 | 821.4 | 819.64 | 742.71 | 717.52 | 813.03 | 897.73 | 804.82 | + |
| Ptger3 | 2.4 | 3.37 | 3.46 | 1.6 | 1.39 | 1.17 | 0.63 | 0.81 | 0.51 | 0.57 | 0.73 | 0.6 | + |
| Taf5l | 10.48 | 12.16 | 11.76 | 6.24 | 6.92 | 6.01 | 7.8 | 8.95 | 7.3 | 6.68 | 6.43 | 6.89 | + |
| Pfkl | 76.53 | 76.16 | 82.21 | 51.34 | 50.67 | 47.86 | 59.15 | 52.71 | 54.59 | 42.27 | 39.68 | 55.8 | + |
| Cinp | 39.31 | 35.5 | 37.12 | 13.1 | 13.46 | 14.84 | 11.91 | 14.84 | 14.86 | 9.1 | 8.2 | 7.87 | + |
| Nln | 12.98 | 12.12 | 12.86 | 7.94 | 7.45 | 6.95 | 8.98 | 8.92 | 7.82 | 7.49 | 8.17 | 5.93 | + |
| Gnaq | 1.82 | 2.16 | 1.88 | 5.13 | 4.71 | 4.94 | 4.66 | 4.47 | 4.63 | 5.72 | 6.09 | 5.66 | + |
| Larp7 | 21.16 | 24.5 | 22.49 | 11.71 | 12.7 | 10.91 | 10.32 | 13.34 | 12.18 | 8.47 | 11.39 | 10.6 | + |
| Baz1a | 5.75 | 6.5 | 5.36 | 3.82 | 5 | 4.08 | 3.79 | 4.2 | 4.45 | 4.78 | 4.19 | 4.59 | + |
| Zfp677 | 4.67 | 3.8 | 3.88 | 7.49 | 5.44 | 8.54 | 8.06 | 7.79 | 7.24 | 8.29 | 6.95 | 6.57 | + |
| C1qbp | 362.26 | 382.95 | 370.98 | 169.31 | 189.48 | 188.08 | 171.34 | 226.48 | 208.32 | 131.96 | 135.99 | 118.45 | + |
| Xcl1 | 924.37 | 751.99 | 763.77 | 55.4 | 61.72 | 99.34 | 53.46 | 131.59 | 108.69 | 26.44 | 20.75 | 29.51 | + |
| Mpv17l | 6.79 | 6.78 | 7.18 | 2.26 | 2.8 | 2.11 | 3.65 | 3.83 | 3.27 | 1.92 | 3.12 | 2.4 | + |
| Eef1d | 189.9 | 195.56 | 187.01 | 130.5 | 131.41 | 138.06 | 133.41 | 131.81 | 131.55 | 145.86 | 151.48 | 138.8 | + |
| Llph | 132.02 | 126.17 | 123.58 | 79.85 | 81.91 | 84.41 | 83.38 | 95.69 | 92.3 | 86.22 | 94.23 | 82.57 | + |
| Lamc2 | 1.05 | 0.55 | 1.03 | 7.22 | 9.92 | 9.29 | 5.9 | 9.07 | 6.77 | 13.69 | 16.16 | 11.57 | + |
| Zbtb38 | 1.2 | 1.04 | 1.36 | 1.88 | 1.91 | 2 | 2.73 | 2.63 | 2.54 | 2.52 | 2.4 | 2.32 | + |
| Tacc3 | 81.97 | 70.97 | 73.86 | 50.3 | 58.9 | 60.44 | 32.22 | 34.1 | 31.09 | 30.51 | 35.78 | 27.18 | + |
| Tomm40 | 139.31 | 154.51 | 165.5 | 48.32 | 51.73 | 53.89 | 35.71 | 56.52 | 51.98 | 30.05 | 26.46 | 31.78 | + |
| Mrps33 | 115.33 | 121.19 | 114.55 | 77.04 | 88.48 | 79.83 | 87.77 | 94.45 | 95.73 | 75.83 | 75.08 | 66.31 | + |
| Snhg5 | 62.01 | 66.6 | 59.14 | 131.58 | 130.09 | 132.44 | 122.79 | 118.33 | 107.06 | 170.69 | 202.91 | 179.99 | + |
| Cox7b | 228.35 | 225.21 | 218.54 | 159.81 | 162.14 | 164.97 | 168.32 | 183.19 | 170.21 | 150.08 | 156.14 | 149.61 | + |
| Tecpr1 | 8.68 | 9.65 | 8.47 | 34.9 | 27.94 | 27.07 | 35.72 | 27.19 | 27.47 | 36.58 | 33.51 | 37.34 | + |
| Gck | 0.13 | 0.29 | 0.31 | 1.46 | 0.86 | 1.31 | 1.79 | 1.33 | 1.31 | 1.61 | 2.17 | 3.08 | + |
| Calr | 283.55 | 267.31 | 274.51 | 103.83 | 110.92 | 108.4 | 100.15 | 130.49 | 122.95 | 84.73 | 94.21 | 76.27 | + |
| Ddah2 | 3.09 | 1.68 | 2.68 | 13.74 | 13.04 | 15.49 | 19.41 | 10.83 | 14.23 | 17.16 | 23.88 | 21.14 | + |
| Dpy19l1 | 4.16 | 3.87 | 4.1 | 2.62 | 2.15 | 1.62 | 2.41 | 1.96 | 2.55 | 2.78 | 2.46 | 1.97 | + |
| Nop56 | 271.86 | 277.62 | 279.59 | 99.69 | 113.51 | 110.49 | 66.91 | 99.92 | 97.79 | 58.35 | 52.23 | 55.97 | + |
| Runx2 | 9.32 | 8.2 | 10.17 | 6.55 | 6.88 | 6.56 | 9.72 | 9.99 | 10.17 | 11.06 | 13.01 | 10.68 | + |
| Suclg1 | 112.38 | 115.5 | 113.69 | 74.55 | 77.72 | 79.53 | 72.45 | 89.88 | 86.73 | 62.22 | 63.4 | 65.35 | + |
| Rpl34-ps1 | 1369.59 | 1419.84 | 1379.38 | 2012.6 | 2004.09 | 2115.14 | 1994.43 | 1895.97 | 1938.43 | 2202.86 | 2379.9 | 2247.43 | + |
| Heca | 1.57 | 1.1 | 1.28 | 3.64 | 3.51 | 3.01 | 4.65 | 3.27 | 3.13 | 4.66 | 5.5 | 4.97 | + |
| 2810008D09Rik | 21.6 | 21.53 | 23.07 | 45.3 | 37.97 | 44.75 | 39.96 | 35.57 | 37.65 | 56.44 | 49.92 | 47.08 | + |
| Ostc | 189.28 | 194.23 | 181.52 | 105.4 | 109.47 | 117.09 | 121.05 | 161.05 | 145.16 | 95.67 | 103.93 | 105.49 | + |
| Lrrc32 | 7.85 | 12.35 | 5.19 | 16.78 | 12.36 | 11.78 | 5.44 | 3.22 | 7.87 | 2.79 | 3.47 | 2.13 | + |
| 5430427O19Rik | 6.4 | 5.89 | 7.15 | 2.49 | 3.3 | 1.98 | 3.21 | 3.75 | 3.36 | 2.57 | 2.19 | 2.86 | + |
| Zfp28 | 2.33 | 1.25 | 1.8 | 2.32 | 3.31 | 3.32 | 3.49 | 2.97 | 3.98 | 2.98 | 3.11 | 3.23 | + |
| Marf1 | 3.22 | 2.62 | 3.29 | 5.81 | 5.26 | 5.47 | 5.22 | 5.19 | 5.1 | 6.09 | 6.14 | 6.78 | + |
| Atat1 | 3.78 | 2.88 | 2.74 | 7.38 | 6.92 | 8.56 | 7.25 | 5.41 | 6.67 | 4.93 | 7.08 | 7.54 | + |
| 4931406C07Rik | 1.25 | 1.03 | 1.18 | 2.88 | 3.48 | 2.12 | 4.07 | 2.88 | 2.31 | 2.52 | 3.93 | 2.93 | + |
| Fmr1 | 6.03 | 6.12 | 6.71 | 4.41 | 4.09 | 4.12 | 3.87 | 3.58 | 3.19 | 3.49 | 3.44 | 3.91 | + |
| Mrps26 | 69.73 | 72.83 | 68.79 | 46.06 | 40.34 | 44.08 | 43.95 | 43.66 | 43.04 | 41.45 | 33.02 | 43.86 | + |
| Polr2c | 145.08 | 137.01 | 147.71 | 79.32 | 82.85 | 82.76 | 74.96 | 96.51 | 85.54 | 61.01 | 68.46 | 67.31 | + |
| Srprb | 29.74 | 29.55 | 28.63 | 22.63 | 23.34 | 22.13 | 18.9 | 23.54 | 23.21 | 17.49 | 21.43 | 19.77 | + |
| Agrn | 0.97 | 0.79 | 0.68 | 2.35 | 2.19 | 2.4 | 1.44 | 0.88 | 1.34 | 1.12 | 1.71 | 1.66 | + |
| Spry2 | 12.23 | 12.34 | 12.07 | 4.42 | 5.71 | 6.02 | 8.7 | 10.13 | 9.9 | 6.5 | 6.2 | 6.71 | + |
| Eif3g | 188.59 | 203.31 | 199.86 | 153.21 | 161.42 | 166 | 136.72 | 155.03 | 147.37 | 135.51 | 140.11 | 135.09 | + |
| Chaf1b | 31.87 | 37.35 | 29.68 | 13.46 | 16.22 | 15.1 | 14.99 | 14.49 | 14.28 | 8.34 | 7.44 | 9.25 | + |
| Msh6 | 36.35 | 34.56 | 35.29 | 31.67 | 34.49 | 34.25 | 27.47 | 25.7 | 26.21 | 30.63 | 30.61 | 27.3 | + |
| Tomm22 | 187.12 | 188.02 | 178.15 | 98.7 | 108.02 | 105.63 | 123.11 | 112.82 | 122.17 | 106.46 | 106.05 | 111.46 | + |
| H2-Ob | 2.35 | 2.27 | 2.57 | 5.4 | 4.08 | 5.14 | 12.59 | 6.75 | 6.53 | 6.26 | 5.44 | 8.72 | + |
| Smyd3 | 5.68 | 5.01 | 5.28 | 13.34 | 14.6 | 13.32 | 13.81 | 12.38 | 14.94 | 10.73 | 12.6 | 9.93 | + |
| Hist1h2ae | 2.49 | 4.85 | 3.9 | 1.28 | 0.97 | 0.69 | 1.72 | 0.85 | 0.93 | 0.65 | 1.34 | 0.87 | + |
| Heatr3 | 28.27 | 31.4 | 32.94 | 8.98 | 10.37 | 11.77 | 6.94 | 9.1 | 7.89 | 5.08 | 5.36 | 5.55 | + |
| Sp4 | 1.25 | 1.62 | 1.81 | 4.13 | 3.1 | 3.06 | 3.33 | 2.61 | 3.05 | 4.2 | 4.55 | 3.81 | + |
| Snhg12 | 28.92 | 30.73 | 29.89 | 164.68 | 149.24 | 166.48 | 116.73 | 154.98 | 132.78 | 165.59 | 209.03 | 159.22 | + |
| Nol10 | 10.79 | 11.22 | 10.61 | 5.8 | 6.46 | 5.05 | 5.68 | 7.07 | 6.17 | 6.11 | 5.48 | 4.62 | + |
| Fam3c | 27.75 | 28.87 | 33.56 | 14.12 | 12.44 | 12.74 | 23.18 | 30.6 | 25.93 | 17.36 | 17.28 | 14.77 | + |
| Smu1 | 78.96 | 78.77 | 75.36 | 49.76 | 47.74 | 53.64 | 46.32 | 53.26 | 50.19 | 45.09 | 43.43 | 47.97 | + |
| Cdc20 | 116.6 | 108.48 | 110.07 | 76.91 | 91.36 | 86.76 | 45.23 | 44.42 | 40.72 | 43.42 | 52.51 | 37.38 | + |
| Tbc1d22a | 11.99 | 9.84 | 10.26 | 16.58 | 14.85 | 14.93 | 21.26 | 19.35 | 19.44 | 19.35 | 21.11 | 18.08 | + |
| Slco4a1 | 40.22 | 38.34 | 37.77 | 17.2 | 17.88 | 18.03 | 29.39 | 26.52 | 27.43 | 14.4 | 15.83 | 15.1 | + |
| Galnt7 | 6.1 | 7.86 | 7.36 | 3.83 | 4.47 | 4.61 | 3.02 | 3.4 | 2.58 | 2.39 | 1.93 | 2.63 | + |
| Dpp3 | 56.15 | 56.34 | 53.43 | 27.88 | 24.4 | 25.48 | 24.61 | 28.72 | 28.08 | 24.73 | 19.88 | 23.55 | + |
| Tspan31 | 84.01 | 74.27 | 78.79 | 75.68 | 73.33 | 84.2 | 89.43 | 77.67 | 79.85 | 89.89 | 95.07 | 101.03 | + |
| Shpk | 0.53 | 1.21 | 1.09 | 4.55 | 2.96 | 2.68 | 3.72 | 3.97 | 2.91 | 4.93 | 5.36 | 5.41 | + |
| Bclaf1 | 41.83 | 42.45 | 41.7 | 20.27 | 20.94 | 20.03 | 15.84 | 19.86 | 17.99 | 19.94 | 18.22 | 17.97 | + |
| Mfsd3 | 4.33 | 4.58 | 5.13 | 3.56 | 2.55 | 2.32 | 1.8 | 2.74 | 2.31 | 2.34 | 2.52 | 2.47 | + |
| Wdhd1 | 14.12 | 15.64 | 15.95 | 6.92 | 8.37 | 7.49 | 5.3 | 5.4 | 6.08 | 4.77 | 4.71 | 5.17 | + |
| Htatip2 | 30.61 | 28.45 | 25.83 | 30.43 | 38.21 | 35.09 | 38.91 | 39.44 | 41.35 | 34.22 | 34.64 | 37.96 | + |
| Lilrb4 | 87.12 | 92.82 | 96.51 | 179.28 | 174.41 | 160.09 | 73.62 | 94.25 | 105.87 | 31.55 | 29.82 | 27.03 | + |
| Abtb1 | 6.41 | 5.57 | 4.46 | 15.06 | 14.58 | 14.93 | 34.45 | 19.18 | 24 | 30.92 | 32.3 | 29.93 | + |
| Cyfip2 | 41.4 | 40.76 | 43.85 | 63.23 | 57.17 | 60.83 | 75.15 | 73.51 | 76.5 | 76.86 | 89.03 | 71.82 | + |
| Sltm | 29.27 | 29.48 | 30.47 | 18.31 | 20.87 | 19.29 | 16.76 | 19.77 | 17.79 | 16.5 | 16 | 16.8 | + |
| Poldip2 | 47.14 | 46.8 | 44.22 | 37.71 | 36.37 | 38.14 | 29.87 | 34.86 | 33.91 | 27.42 | 30.99 | 29.9 | + |
| Anks3 | 7.56 | 6.26 | 7.29 | 7.61 | 7.79 | 7.35 | 7.8 | 8.14 | 6.53 | 9.12 | 10.9 | 9.38 | + |
| Aifm1 | 36.1 | 35.38 | 36.28 | 18.6 | 19.17 | 19.5 | 15.35 | 19.91 | 17.5 | 14.9 | 16.84 | 15.34 | + |
| Ccnc | 37.34 | 37.03 | 40.88 | 24.42 | 27.35 | 27.11 | 20.8 | 24.76 | 25.38 | 22.9 | 22.67 | 24.05 | + |
| Tcf19 | 27.71 | 29.56 | 25.08 | 18 | 17.4 | 15.99 | 14.78 | 15.08 | 13.51 | 13.97 | 11.3 | 14.62 | + |
| 4632415K11Rik | 3.37 | 2.98 | 3.68 | 5.01 | 4.9 | 5.75 | 5.8 | 5.92 | 4.84 | 6.45 | 5.75 | 5.69 | + |
| Dtd1 | 49.63 | 48.75 | 47.1 | 30.2 | 34.54 | 30.37 | 35.11 | 42.12 | 36.3 | 25.24 | 30.36 | 31 | + |
| Wdr92 | 23.96 | 21.99 | 23.68 | 32.27 | 34.9 | 35.52 | 25.69 | 30.88 | 29.46 | 31.68 | 32.3 | 30.54 | + |
| 2310033P09Rik | 9.84 | 10.74 | 10.24 | 6.06 | 7.14 | 5.34 | 4.63 | 4.45 | 3.98 | 4.56 | 5.02 | 5.25 | + |
| Wdr36 | 65.01 | 67.23 | 63.57 | 39.11 | 39.18 | 40.11 | 37.27 | 49.15 | 41.87 | 31.06 | 31.13 | 28.84 | + |
| Hnrnpf | 578.21 | 616.39 | 626.48 | 405.3 | 404.65 | 392.63 | 360.15 | 425.88 | 410.35 | 367.72 | 356.88 | 374.08 | + |
| Txnl1 | 84.66 | 84.25 | 86.51 | 65.76 | 67.57 | 71.52 | 71.73 | 72.73 | 72.95 | 62.6 | 75.15 | 64.75 | + |
| Fads2 | 28.17 | 31.33 | 33.97 | 12.74 | 15.56 | 13.49 | 13.45 | 14.1 | 13.16 | 7.7 | 7.44 | 9.59 | + |
| Piezo1 | 7.02 | 6.11 | 6.65 | 4.97 | 4.69 | 4.52 | 4.46 | 4.71 | 4.5 | 4.8 | 4.28 | 4.82 | + |
| Cdc45 | 63.16 | 58.56 | 59.67 | 23.91 | 30.33 | 30.36 | 17.48 | 25.52 | 18.56 | 11.84 | 14.85 | 12.9 | + |
| Pphln1 | 26.41 | 27.07 | 25.31 | 12.82 | 16.02 | 13 | 9.78 | 14.23 | 12.41 | 9.88 | 10.28 | 9.61 | + |
| Eya3 | 13.83 | 11.79 | 12.47 | 8.28 | 8.14 | 9.35 | 9.09 | 9.48 | 9.6 | 7.79 | 8.82 | 8.18 | + |
| Nr4a2 | 9.23 | 10.73 | 12.61 | 3.84 | 6.22 | 7.04 | 5.24 | 7.74 | 6.04 | 6.74 | 6.86 | 8.29 | + |
| Guca1a | 4.45 | 2.8 | 2.54 | 7.67 | 7.51 | 5.62 | 16.19 | 12.58 | 14.75 | 7.6 | 11.44 | 8.58 | + |
| Ncbp1 | 60.48 | 59.45 | 61.91 | 29.09 | 29.6 | 33.25 | 26 | 33.95 | 32.77 | 24.92 | 23.5 | 23.55 | + |
| Stard5 | 6.64 | 8.16 | 7.09 | 10.24 | 11.79 | 12.31 | 9.24 | 11.29 | 9.65 | 9.65 | 9.76 | 10.48 | + |
| Kbtbd11 | 1.48 | 2.12 | 2.17 | 15.43 | 13.14 | 13.64 | 3.97 | 2.85 | 3.1 | 6.33 | 5.2 | 6.82 | + |
| Tyw3 | 1.56 | 1.52 | 1.95 | 0.55 | 0.48 | 0.76 | 0.69 | 0.86 | 0.47 | 0.61 | 0.61 | 0.79 | + |
| Ndrg2 | 22.87 | 22.81 | 28.72 | 31.88 | 33.34 | 33.48 | 40.63 | 42.11 | 43.4 | 30.3 | 34.94 | 32.74 | + |
| Hspa5 | 555.82 | 554.12 | 547.6 | 230.82 | 274.58 | 280.02 | 213.98 | 354.98 | 285.05 | 189.93 | 226.01 | 180.71 | + |
| Cyb5rl | 2.77 | 2.36 | 2.05 | 0.83 | 1.4 | 1.23 | 0.72 | 0.51 | 1.07 | 0.82 | 0.65 | 0.54 | + |
| Frg1 | 60.56 | 54.8 | 46.22 | 72.67 | 74.35 | 74.31 | 86.35 | 78.74 | 80.67 | 83.88 | 85.76 | 85.72 | + |
| Dnajb13 | 4.25 | 4.53 | 3.78 | 7.8 | 7.32 | 6.53 | 11.97 | 6.99 | 7.18 | 7.96 | 7.36 | 9.48 | + |
| Tigit | 93.67 | 80.96 | 88.87 | 122.48 | 128.27 | 128.91 | 130.51 | 129.47 | 124.43 | 316.6 | 261.5 | 298.35 | + |
| Tardbp | 30.54 | 32.66 | 33.3 | 19.03 | 19.13 | 20.05 | 14.7 | 19.48 | 17.91 | 17.08 | 17.51 | 16.38 | + |
| Riok2 | 16.42 | 16.4 | 15.02 | 10.7 | 9.78 | 8.7 | 8.11 | 9.79 | 9.47 | 7.05 | 7.71 | 7.76 | + |
| D10Wsu52e | 150.24 | 159.57 | 154.55 | 134.37 | 129.18 | 140.98 | 122.2 | 133.22 | 135.61 | 129.36 | 139.19 | 123.26 | + |
| Ppa2 | 56.9 | 56.41 | 54.78 | 33.71 | 38.53 | 38.82 | 30.42 | 37.63 | 26.63 | 25.95 | 24.63 | 26.32 | + |
| Lepre1 | 6.98 | 7.7 | 6.6 | 4.23 | 4.61 | 4.48 | 3.64 | 5.76 | 4.57 | 4.51 | 4.13 | 4.4 | + |
| Bcap31 | 133.08 | 139.43 | 131.71 | 96.99 | 98.56 | 103.85 | 107.49 | 108.32 | 102.44 | 104.37 | 106.74 | 105.71 | + |
| Tollip | 23.42 | 23.85 | 22.82 | 17.35 | 15.85 | 17.63 | 17.18 | 15.71 | 16.47 | 18.07 | 15.14 | 16.28 | + |
| Cox7a2 | 313.94 | 317.98 | 306.19 | 208.3 | 218.18 | 226.44 | 242.36 | 229.96 | 245.5 | 200.39 | 211.68 | 192.59 | + |
| Supt6 | 15.36 | 13.21 | 16.12 | 12.24 | 12.59 | 12.31 | 10.6 | 11.65 | 11.47 | 10.96 | 12.01 | 10.43 | + |
| Crot | 17.18 | 14.76 | 16.08 | 23.92 | 23.74 | 22.98 | 31.27 | 26.2 | 24.84 | 30.18 | 27.84 | 29.72 | + |
| Acads | 25.89 | 27.2 | 25.62 | 32.38 | 29.51 | 30.62 | 38.11 | 34.71 | 35.03 | 30.96 | 37.73 | 34.16 | + |
| Gpn2 | 33.81 | 28.79 | 33.93 | 17.82 | 19.43 | 16.7 | 14.15 | 18.54 | 19.51 | 13.56 | 13.95 | 12.37 | + |
| Rwdd1 | 157.71 | 159.64 | 156.11 | 107.6 | 119.02 | 125.66 | 124.68 | 139.56 | 139.63 | 113.31 | 123.51 | 107.99 | + |
| Upf3a | 3.34 | 4.53 | 4.51 | 6.27 | 6.46 | 7.15 | 7.05 | 7.63 | 8.58 | 7.1 | 8.23 | 5.51 | + |
| Hsd17b10 | 178.78 | 158.51 | 170.78 | 207.07 | 220.91 | 221.19 | 273.25 | 292.76 | 281.77 | 223.99 | 261.31 | 246.56 | + |
| Trappc4 | 103.59 | 96.42 | 97.49 | 63.96 | 62.42 | 64.76 | 72.92 | 79.37 | 71.28 | 61.12 | 63.25 | 65.61 | + |
| Tmem123 | 42.59 | 49.12 | 53.91 | 78.98 | 72.62 | 71.94 | 107.57 | 94.67 | 99.52 | 110.64 | 97.71 | 108.41 | + |
| Sqle | 74.21 | 81.01 | 75.36 | 62.24 | 68.19 | 64.94 | 54.79 | 62 | 56.18 | 56.28 | 59 | 53.83 | + |
| 4930581F22Rik | 0.36 | 0.61 | 0.18 | 0.98 | 1.12 | 0.85 | 1.22 | 0.83 | 1.05 | 0.83 | 1.39 | 1.06 | + |
| Ankrd12 | 2.01 | 1.91 | 2.01 | 3.49 | 3.24 | 3.28 | 3.63 | 3.39 | 3.54 | 4.18 | 4.29 | 4.99 | + |

| | | | | | | | | | | | | | |
|---|---|---|---|---|---|---|---|---|---|---|---|---|---|
| Sumf2 | 20.7 | 20.8 | 22.64 | 14.61 | 14.72 | 16.34 | 12.34 | 14.44 | 13.09 | 11.96 | 10.89 | 12.29 | + |
| Recql | 17.51 | 17.53 | 17.63 | 10.11 | 10.51 | 10.36 | 10.07 | 9.9 | 8.8 | 8.89 | 8.06 | 8.76 | + |
| Rabif | 30.46 | 26.76 | 29.09 | 19.03 | 21.23 | 20.5 | 23.29 | 24.25 | 21.2 | 21.55 | 20.42 | 20.97 | + |
| 2310014L17Rik | 0.41 | 0.54 | 0.55 | 0.06 | 0.15 | 0.16 | 0.09 | 0.07 | 0.1 | 0.12 | 0.17 | 0.09 | + |
| Khnyn | 4.24 | 4.52 | 4.51 | 13.71 | 11.85 | 11.37 | 19.24 | 11.51 | 16.84 | 16.89 | 20.75 | 18.93 | + |
| Abhd14a | 26.88 | 29.36 | 27.03 | 18.63 | 22.94 | 22.99 | 19.09 | 23.69 | 23.45 | 23.17 | 21.04 | 18.72 | + |
| Abcb8 | 14.29 | 15.18 | 13.9 | 10.21 | 11.08 | 11.06 | 11.45 | 10.83 | 10.37 | 9.13 | 8.26 | 10.78 | + |
| Glrx3 | 166.09 | 177.25 | 158.77 | 135.85 | 125.91 | 139.96 | 136.56 | 141.15 | 146.05 | 129.21 | 146.08 | 127.79 | + |
| Taok2 | 7.97 | 7.72 | 6.7 | 8.39 | 7.45 | 7.42 | 7.87 | 6.29 | 7.27 | 10.31 | 8.34 | 10.33 | + |
| Hagh | 29.82 | 23.26 | 26.33 | 25.63 | 26.63 | 26.77 | 28.85 | 36.4 | 28.24 | 20.03 | 21.99 | 15.58 | + |
| Rps7 | 754.1 | 797.02 | 766.49 | 1245.61 | 1223.64 | 1304.24 | 1230.6 | 1140.34 | 1164.89 | 1316.07 | 1445.47 | 1316.16 | + |
| Sh3glb2 | 18 | 18.65 | 17.55 | 14.51 | 14.11 | 12.91 | 14.62 | 13.72 | 14.12 | 14.39 | 12.04 | 13.63 | + |
| Arrb1 | 2.33 | 2.41 | 2.64 | 10.02 | 7.52 | 8.09 | 12.77 | 7.67 | 9.48 | 10.12 | 11.78 | 14.37 | + |
| Gnl2 | 55.95 | 57 | 62.41 | 43.81 | 43.06 | 48.14 | 38.93 | 44.03 | 39.48 | 34.37 | 35.69 | 35.54 | + |
| Acox3 | 6.58 | 6.07 | 5.45 | 12 | 12.22 | 12.85 | 12.49 | 10.76 | 8.87 | 12.67 | 11.54 | 13.4 | + |
| Zc3h6 | 0.33 | 0.58 | 0.46 | 1.22 | 1.73 | 1.49 | 1.08 | 1.12 | 1.1 | 1 | 1.29 | 1.75 | + |
| Rwdd2a | 1.94 | 2.25 | 1.66 | 6.72 | 5.29 | 5.15 | 4.92 | 4.17 | 4.89 | 7.81 | 5.82 | 7.91 | + |
| Prdx3 | 182.6 | 174.3 | 168.7 | 111.37 | 114.13 | 110.99 | 110.28 | 131.69 | 123.66 | 93.99 | 94.46 | 98.59 | + |
| Ccne2 | 21.84 | 20.63 | 22.56 | 5.15 | 5.97 | 6.06 | 4.93 | 4.53 | 4.43 | 4 | 3.34 | 4.38 | + |
| Arap2 | 1.8 | 1.9 | 2.24 | 7.08 | 6.65 | 6.14 | 2.55 | 2.94 | 2.63 | 4.55 | 4.43 | 4.46 | + |
| Lancl1 | 11.9 | 11.93 | 12.55 | 15.78 | 17.62 | 17.18 | 36.73 | 24.91 | 30.99 | 26.54 | 24.88 | 28.17 | + |
| Ntng2 | 1.63 | 1.51 | 1.26 | 2.65 | 3.37 | 3 | 4.51 | 4.89 | 4.26 | 6.7 | 5.65 | 5.59 | + |
| Nupr1 | 12.77 | 13.57 | 13.47 | 11.81 | 14.26 | 21.7 | 11.93 | 21.86 | 19.79 | 3.56 | 7.19 | 3.68 | + |
| Pbxip1 | 7.66 | 7.25 | 7.41 | 19.11 | 15.94 | 16.71 | 25.9 | 16.36 | 18.79 | 25.93 | 25.21 | 27.63 | + |
| Rab33b | 13.24 | 15.05 | 13.91 | 9.55 | 9.68 | 8.82 | 10.46 | 9.74 | 9.69 | 9.54 | 9.87 | 9.47 | + |
| Adat1 | 2.53 | 2.23 | 2.89 | 1.62 | 1.34 | 1.8 | 1.4 | 1.49 | 1.48 | 1.7 | 1.15 | 1.33 | + |
| Cops3 | 99.24 | 104.87 | 101.69 | 59.55 | 66.91 | 66.26 | 53.13 | 56.47 | 57.07 | 47.2 | 47.81 | 41.96 | + |
| Rcc1 | 66.64 | 61.93 | 69.69 | 24.75 | 34.29 | 30.86 | 19.47 | 28.14 | 27.55 | 17.41 | 16.26 | 14.83 | + |
| Asf1b | 55.73 | 59.61 | 54.44 | 28.42 | 29.47 | 29.63 | 18.34 | 18.73 | 18.99 | 16.81 | 13.63 | 12.8 | + |
| Dscc1 | 6.67 | 8.58 | 8.72 | 2.27 | 2.96 | 2.44 | 2.11 | 1.44 | 1.64 | 1.31 | 0.99 | 1.5 | + |
| Ndufb7 | 165.91 | 173.21 | 171.23 | 115.5 | 118.64 | 118.33 | 122.47 | 128.93 | 136.28 | 92.09 | 98.86 | 100.25 | + |
| Rnf169 | 2.15 | 1.85 | 1.75 | 3.26 | 3.58 | 3.4 | 2.89 | 3.19 | 3.01 | 3.94 | 3.28 | 3.58 | + |
| Fam183b | 0.72 | 0.94 | 0.29 | 10.25 | 6.64 | 7.09 | 17.64 | 4.87 | 9.09 | 12.68 | 14.21 | 20.36 | + |
| Tomm34 | 69.47 | 70.88 | 72.23 | 48.65 | 54.5 | 56.12 | 49.92 | 51.56 | 51.9 | 48.16 | 45.48 | 53.06 | + |
| AB124611 | 11.92 | 13 | 10.25 | 29.99 | 23.77 | 28.2 | 24.09 | 16.37 | 18.16 | 24.11 | 25.12 | 22.89 | + |
| Rpl29 | 1217.13 | 1234.91 | 1284.51 | 1564.2 | 1551.22 | 1615.37 | 1596.87 | 1602.12 | 1591.01 | 1629.59 | 1780.46 | 1586.92 | + |
| Parl | 56.83 | 59.95 | 56.61 | 36.14 | 36.07 | 36.27 | 31.74 | 40.15 | 39.65 | 32.5 | 26.71 | 29.56 | + |
| 4930579K19Rik | 0.25 | 0.15 | 0.42 | 0.82 | 0.7 | 1.61 | 1.35 | 0.82 | 1.35 | 1.44 | 1.07 | 1.3 | + |
| Msn | 189.1 | 190.03 | 196.38 | 198.82 | 192.83 | 191.32 | 207.45 | 191.96 | 203.82 | 228.42 | 216.13 | 224.83 | + |
| Psmc5 | 181.21 | 170.6 | 166.64 | 71.46 | 88.86 | 84.65 | 63.17 | 91.16 | 77.59 | 60.28 | 58.59 | 56.7 | + |
| Coq7 | 44.57 | 40.9 | 46.68 | 11.24 | 15.84 | 19.84 | 10.29 | 15.85 | 15.25 | 7.75 | 7.36 | 7.21 | + |
| Ilf2 | 67.93 | 66.04 | 73.86 | 42.73 | 45.69 | 48.76 | 33.4 | 40.94 | 36.5 | 31.66 | 38.58 | 33.06 | + |
| Sesn3 | 2.33 | 1.62 | 2.66 | 5.13 | 5.03 | 4.32 | 5.78 | 4.6 | 5.72 | 5.46 | 6 | 4.89 | + |
| 1600016N20Rik | 1.14 | 0.85 | 0.98 | 2.04 | 2.9 | 2.6 | 3.93 | 3.1 | 2.7 | 1.9 | 2.17 | 4.06 | + |
| Mcm7 | 119.82 | 135.45 | 134.47 | 66.66 | 74.39 | 71.41 | 54.16 | 51.73 | 50.63 | 41.19 | 44.55 | 44.87 | + |
| Nup54 | 39.13 | 42.46 | 43.9 | 30.25 | 30.02 | 31.7 | 27.7 | 34.95 | 30.93 | 25.39 | 29.81 | 29.58 | + |
| Isyna1 | 89.1 | 83.69 | 79.72 | 47.05 | 46.83 | 45.52 | 44.24 | 51.57 | 49.97 | 38.08 | 40.44 | 38.7 | + |
| AA467197 | 282.87 | 296.49 | 255.31 | 400.99 | 449.17 | 433.96 | 407.69 | 463.41 | 440.1 | 363.02 | 391.47 | 375.97 | + |
| Thoc1 | 25.19 | 23.68 | 25.59 | 15.07 | 15.34 | 17.03 | 12.86 | 16.12 | 13.8 | 12.18 | 11.88 | 12.34 | + |
| Hmgn2 | 166.99 | 213.08 | 165.21 | 47.78 | 54.81 | 50.36 | 36.98 | 36.21 | 32.13 | 26.89 | 30.23 | 29.74 | + |
| Chrna1 | 1.54 | 1.71 | 1.61 | 0.79 | 0.69 | 0.65 | 0.59 | 0.95 | 0.78 | 0.87 | 0.88 | 0.67 | + |
| Mrpl53 | 57.27 | 54.06 | 53.28 | 38.33 | 40.68 | 38.9 | 44.4 | 42.16 | 44.74 | 35.62 | 39.26 | 33.47 | + |
| Rps16 | 1798.89 | 1818.82 | 1839.16 | 3101.49 | 2966.41 | 3163.72 | 2910.9 | 2611.44 | 2724.16 | 3110.76 | 3365.75 | 3138.21 | + |
| Polk | 7.08 | 6.95 | 7.92 | 9.84 | 10.5 | 10.94 | 11.33 | 10.66 | 12.14 | 11.99 | 12.7 | 11.21 | + |
| Gusb | 53.53 | 52.24 | 53.18 | 20.82 | 24.53 | 25.13 | 17.17 | 21.12 | 22.34 | 18.03 | 15.92 | 16.48 | + |
| Fam169b | 25.06 | 30 | 26.91 | 51.19 | 46.77 | 44.98 | 28.79 | 29.53 | 30.31 | 33.9 | 34.3 | 35.06 | + |
| Hcfc1r1 | 21.61 | 18.44 | 21.3 | 29.66 | 31.49 | 30.05 | 42.62 | 32.21 | 37.14 | 40.15 | 39.46 | 40.41 | + |
| Trmt10c | 37.78 | 39.53 | 39.45 | 27.09 | 27.8 | 27.26 | 21.95 | 26.75 | 27.09 | 25.91 | 21.37 | 24.51 | + |
| Ecd | 47.53 | 44.56 | 46.9 | 27.55 | 31.86 | 27.02 | 22.99 | 30.01 | 26.08 | 20.28 | 19.52 | 21.5 | + |
| Shq1 | 16.55 | 17.81 | 16.25 | 6.8 | 7.43 | 9.07 | 5.2 | 6.16 | 6.09 | 5 | 4.12 | 4.37 | + |
| Tdg | 5.74 | 5.6 | 4.69 | 2.95 | 3.8 | 3.21 | 2.73 | 3.47 | 2.09 | 2.81 | 3.28 | 2.25 | + |
| Fen1 | 72.15 | 73.16 | 73.63 | 25.42 | 28.72 | 31.66 | 21.64 | 23.03 | 25.82 | 15.66 | 12.48 | 14.97 | + |
| Far1 | 32.82 | 29.7 | 33.48 | 16.58 | 19.72 | 19.74 | 17.04 | 17.4 | 19.08 | 17.5 | 15.31 | 14.22 | + |
| Serinc5 | 0.86 | 0.69 | 1.15 | 2.06 | 1.75 | 1.79 | 2.28 | 2.53 | 2.28 | 2.58 | 2.18 | 2.24 | + |
| Rps26 | 1806.37 | 1826.48 | 1834.91 | 2372.69 | 2440.83 | 2542.97 | 2329.71 | 2279.45 | 2235.68 | 2188.99 | 2455.54 | 2168.33 | + |
| Eaf1 | 22.06 | 19.55 | 21.82 | 10.19 | 12.23 | 9.62 | 11.06 | 14.53 | 15.76 | 9.43 | 11.26 | 9.64 | + |
| Tubg1 | 83.33 | 81.49 | 80.26 | 30.54 | 35.58 | 32.35 | 28.75 | 32.32 | 27.37 | 23.3 | 25.54 | 22.04 | + |
| Ptpn1 | 77.16 | 79.6 | 82.17 | 139.66 | 133.79 | 130.78 | 47.25 | 62.92 | 55.43 | 45.22 | 47.81 | 49.88 | + |
| Dolk | 6.55 | 7.72 | 6.92 | 3.33 | 3.85 | 4.44 | 4.24 | 4.59 | 4.11 | 3.44 | 3.51 | 4.12 | + |
| Sec61a1 | 124.27 | 131.06 | 129.72 | 96.12 | 94.39 | 98.68 | 104.12 | 108.34 | 109.17 | 98.78 | 106.14 | 99.89 | + |
| Fam160a2 | 1.73 | 1.69 | 1.93 | 3.1 | 3.27 | 3.84 | 3.89 | 2.87 | 3.44 | 3.37 | 3.2 | 3.8 | + |
| Rbm34 | 16.28 | 16.58 | 15.65 | 11.26 | 9.81 | 10.12 | 8.52 | 11.05 | 9.89 | 8.4 | 8.14 | 7.16 | + |
| Nudt18 | 2.73 | 4.37 | 5.33 | 8.67 | 8.39 | 7.64 | 11.93 | 9.14 | 9.91 | 9.57 | 9.94 | 10.6 | + |
| Txlna | 30.96 | 31.14 | 30.69 | 19.64 | 21.89 | 21.68 | 18.46 | 19.82 | 20.34 | 15.56 | 18.47 | 17.05 | + |
| Anxa3 | 6.86 | 5.03 | 8.5 | 3.44 | 3.91 | 2.6 | 2.77 | 8.67 | 5.21 | 1 | 1.27 | 0.54 | + |
| Rsl24d1 | 84.1 | 90.03 | 82.58 | 45.43 | 45.38 | 36.79 | 45.63 | 50.81 | 49.28 | 42.37 | 44.83 | 43.67 | + |
| Trap1 | 77.98 | 78.58 | 77.28 | 49.25 | 52.86 | 50.45 | 47.21 | 54.46 | 52.53 | 42.56 | 40.22 | 41.13 | + |
| Cnnm2 | 0.28 | 0.46 | 0.51 | 1.1 | 0.8 | 0.47 | 1.71 | 1.55 | 1.42 | 1.31 | 1.02 | 1.18 | + |
| Fbxo22 | 22.13 | 25.4 | 23.02 | 14.09 | 13.66 | 17.27 | 16.45 | 15.7 | 16.12 | 15.4 | 14.96 | 17.47 | + |
| Igbp1 | 60.16 | 58.38 | 57.49 | 117.25 | 101.12 | 110.95 | 123.5 | 105.44 | 110.03 | 133.91 | 138.14 | 138.92 | + |
| Psma2 | 563.74 | 575.15 | 557.01 | 336.75 | 351.11 | 359 | 362.62 | 396.56 | 385.11 | 319.16 | 333.3 | 313.03 | + |
| Wdr4 | 20.14 | 21.14 | 20.78 | 7.8 | 7.48 | 7.64 | 6.18 | 9.77 | 8.11 | 5.72 | 3.55 | 5.96 | + |
| A430005L14Rik | 25.43 | 22.78 | 22.2 | 14.22 | 15.59 | 15.28 | 16.61 | 12.14 | 12.61 | 12.87 | 9.3 | 13.03 | + |
| Ggct | 11.48 | 13.12 | 11.59 | 5.99 | 5.82 | 6.74 | 5.11 | 7.55 | 7.33 | 4.33 | 4.93 | 6.52 | + |
| Ndrg1 | 10.71 | 10.81 | 11.08 | 77.33 | 72.71 | 84.45 | 66.41 | 36.48 | 45.9 | 104.96 | 105.57 | 114.36 | + |
| Pop5 | 66.38 | 69.81 | 67.68 | 37.59 | 42.54 | 42.11 | 48.17 | 52.43 | 51.21 | 42.23 | 50.16 | 42.78 | + |
| Inpp4a | 3.92 | 3.14 | 3.82 | 7.57 | 6.97 | 6.9 | 7.11 | 6.86 | 6.53 | 8.04 | 7.95 | 7.89 | + |
| Trim26 | 10.25 | 11.86 | 10.93 | 11.96 | 13.8 | 12.75 | 12.99 | 11.76 | 10.82 | 16.72 | 15.2 | 16.13 | + |
| Lrp1 | 2.04 | 1.7 | 2.25 | 6.07 | 8.06 | 8.45 | 1.68 | 1.37 | 1.6 | 3.07 | 3.19 | 3.12 | + |
| Dnttip2 | 58.34 | 63.2 | 59.43 | 37.43 | 37.87 | 38.67 | 31.41 | 36.37 | 30.6 | 28.96 | 29.78 | 33 | + |
| Filip1l | 2.25 | 3.28 | 2.87 | 6.68 | 5.39 | 4.57 | 6.16 | 3.86 | 6.45 | 5.81 | 3.92 | 5.08 | + |
| Nkiras1 | 2.95 | 3.26 | 3.45 | 2.01 | 1.86 | 1.81 | 1.45 | 2.08 | 1.75 | 1.92 | 1.42 | 1.45 | + |
| Slc16a13 | 7.95 | 7.34 | 10.17 | 4.13 | 3.56 | 2.8 | 4 | 4.65 | 4.2 | 2.49 | 2.24 | 3.53 | + |
| Cfl1 | 895.23 | 945.72 | 927.91 | 635.61 | 651.3 | 636.88 | 674.2 | 685.82 | 694.06 | 567.36 | 554.04 | 604.07 | + |
| Rpl7 | 1917.91 | 1900.02 | 1906.53 | 2480.78 | 2436.87 | 2588.95 | 2398.08 | 2292.1 | 2330.43 | 2522.07 | 2759.8 | 2468.27 | + |
| Zbtb20 | 0.77 | 0.84 | 0.81 | 2.82 | 2.92 | 3.89 | 3.17 | 2.19 | 2.41 | 4.45 | 3.38 | 4.61 | + |
| Tmem144 | 1.07 | 0.94 | 1.39 | 0.36 | 0.44 | 0.19 | 0.63 | 0.56 | 0.61 | 0.25 | 0.7 | 0.3 | + |
| Obsl1 | 0.48 | 0.36 | 0.45 | 0.94 | 1.28 | 0.93 | 0.97 | 0.84 | 0.99 | 1 | 1.64 | 1.16 | + |
| Pdzk1ip1 | 20.8 | 10.88 | 11.88 | 1.6 | 0.79 | 1.41 | 1.64 | 2.73 | 2.72 | 0.3 | 0.33 | 0.85 | + |
| Mettl20 | 5.32 | 5.31 | 4.44 | 12.61 | 11.69 | 10.75 | 12.65 | 10.77 | 12.54 | 10.14 | 11.03 | 15.45 | + |
| Map1b | 0.74 | 0.58 | 0.63 | 0.98 | 1.27 | 1.33 | 0.91 | 1.04 | 0.9 | 0.41 | 0.24 | 0.42 | + |
| Mrpl41 | 16.6 | 13.69 | 15.02 | 9.64 | 7.87 | 11.25 | 7.89 | 6.97 | 6.72 | 7.9 | 7.41 | 7.9 | + |
| Prkrip1 | 22.17 | 21.74 | 20.63 | 11.43 | 13.89 | 12.76 | 10.92 | 14.06 | 13.56 | 9.22 | 10.51 | 9.65 | + |
| Mrpl42 | 203.69 | 202.65 | 208.23 | 99.42 | 113.41 | 111.67 | 89.56 | 122.94 | 116.77 | 84.86 | 87.63 | 73.8 | + |
| H2-Ke2 | 53.75 | 56.73 | 53.99 | 34.81 | 34.29 | 35.12 | 37.13 | 35.53 | 35.74 | 29.84 | 30.09 | 29.09 | + |
| Igsf8 | 12.92 | 13.53 | 11.89 | 7.22 | 5.76 | 5 | 6.87 | 7.08 | 6.18 | 4.23 | 5 | 6.25 | + |
| Fosl2 | 3.73 | 3.92 | 3.71 | 7.39 | 7.34 | 8.43 | 4.85 | 6.12 | 6.27 | 5.25 | 4.57 | 6.17 | + |
| Tsfm | 59.15 | 61.84 | 59.57 | 35.93 | 35.39 | 36.05 | 35.65 | 36.14 | 38.19 | 30.35 | 29.46 | 28.63 | + |
| Rilpl2 | 137.68 | 136.04 | 138.52 | 115.68 | 122.67 | 122.94 | 189.77 | 156.11 | 156.46 | 178.37 | 184.79 | 187.29 | + |
| Flnb | 6.4 | 4.68 | 5.68 | 2.92 | 3.74 | 3.92 | 3.11 | 3.64 | 2.99 | 2.71 | 3.62 | 2.47 | + |
| Dhrs4 | 7.15 | 6.77 | 6.25 | 2.18 | 2.86 | 2.63 | 1.9 | 3.75 | 3.78 | 3.09 | 1.66 | 2.9 | + |
| Tamm41 | 32.02 | 28.4 | 28.88 | 19.46 | 18.58 | 17.28 | 16.81 | 17.68 | 15.73 | 14.38 | 15.91 | 15.08 | + |
| 2610301G19Rik | 16.66 | 19.47 | 17.97 | 12.9 | 12.91 | 10.64 | 10.64 | 11.32 | 11.46 | 8.98 | 9.7 | 11.11 | + |
| Dbp | 0.43 | 0.29 | 0.43 | 1.78 | 1.24 | 1.43 | 1.94 | 1.64 | 1.92 | 1.66 | 2.08 | 2.48 | + |
| Ebp | 44.14 | 42.98 | 40.65 | 29.44 | 29.18 | 28.28 | 28.18 | 28.64 | 30.2 | 28.49 | 26.86 | 29.2 | + |
| Tnfaip8l1 | 4.13 | 3.42 | 3.38 | 6.31 | 7.52 | 7.66 | 6.83 | 4.64 | 6.8 | 7.26 | 7.06 | 6.8 | + |
| 9130401M01Rik | 49.49 | 49.46 | 46.54 | 30.42 | 29.09 | 27.91 | 29.94 | 34.33 | 36.53 | 23.24 | 22.63 | 23.64 | + |
| U2surp | 19.98 | 20.79 | 20.21 | 15.76 | 15.77 | 15.41 | 12.64 | 12.26 | 13.8 | 12.08 | 13.55 | 13.31 | + |
| Vav2 | 1.26 | 1.52 | 1.12 | 2.98 | 2.56 | 3.08 | 2.53 | 2.67 | 2.54 | 2.37 | 2.15 | 3.05 | + |
| Nup35 | 25.49 | 23.59 | 21.94 | 9.18 | 12.82 | 12.08 | 8.16 | 9.42 | 7.87 | 7.63 | 7.53 | 8.46 | + |
| Aco2 | 220.06 | 212.85 | 218.79 | 138.34 | 156.03 | 150.07 | 149.51 | 226.46 | 210.21 | 88.43 | 90.07 | 91.81 | + |
| Zbtb7b | 1.61 | 1.97 | 1.71 | 3.97 | 3.42 | 3.39 | 4.11 | 4.54 | 5 | 3.13 | 3.31 | 3.41 | + |
| Rpl8 | 1896.93 | 1942.99 | 1952.71 | 2412.93 | 2450.4 | 2502.28 | 2250.28 | 2258.84 | 2232.63 | 2308.96 | 2501.03 | 2234.26 | + |
| Sap130 | 10.69 | 10.33 | 11.07 | 10.99 | 11.65 | 11.98 | 11.51 | 11.44 | 10.11 | 13.97 | 14.31 | 12.91 | + |
| 4933409K07Rik | 23.43 | 22.88 | 23.44 | 39.01 | 37.85 | 41.24 | 54.21 | 49.43 | 47.04 | 62.34 | 68.19 | 56.58 | + |
| Sft2d3 | 0.33 | 0.33 | 0.33 | 1.06 | 0.76 | 0.5 | 1.02 | 0.92 | 1.32 | 1.2 | 0.9 | 1.01 | + |
| Imp4 | 62.37 | 55.85 | 57.31 | 19.64 | 20.87 | 21.49 | 16.12 | 20.95 | 20.44 | 15 | 11.18 | 12.74 | + |
| Med17 | 17.32 | 18.02 | 19.4 | 14.31 | 12.15 | 13.13 | 12.51 | 11.85 | 12.08 | 10.44 | 9.6 | 12.65 | + |
| Coq10b | 23.17 | 21.17 | 21.29 | 9.12 | 10.31 | 10.05 | 9.85 | 13.99 | 14.7 | 7.69 | 7.82 | 9.62 | + |
| Gtf3c3 | 12.58 | 14.66 | 12.56 | 8.96 | 10.26 | 9.68 | 8.59 | 10.47 | 9.15 | 8.62 | 8.88 | 7.86 | + |
| Stk16 | 18.13 | 20.3 | 18.09 | 12.44 | 13.55 | 12.58 | 15.05 | 12.54 | 13.3 | 13.99 | 13.29 | 16.67 | + |
| Adam19 | 5.78 | 6.23 | 4.58 | 13.88 | 10.18 | 9.92 | 9.49 | 7.55 | 7.96 | 10.4 | 9.72 | 11.28 | + |
| Sh2d2a | 52.05 | 55.58 | 49.74 | 33.34 | 30.51 | 30.62 | 28.98 | 28.33 | 28.59 | 34.77 | 23.86 | 33.54 | + |
| Ccrn4l | 4.26 | 3.68 | 4.39 | 2.68 | 2.12 | 2.39 | 2.35 | 2.55 | 2.69 | 2.91 | 2.29 | 2.07 | + |
| Stip1 | 233.21 | 231.53 | 236.57 | 100.78 | 120.77 | 116.27 | 83.88 | 123.63 | 106.75 | 78.37 | 81.99 | 76.05 | + |
| Hyi | 7.42 | 6.94 | 6.32 | 28.73 | 26.18 | 30.28 | 30.88 | 25.11 | 27.88 | 34.06 | 33.4 | 34.59 | + |
| Zfand6 | 37.22 | 41.48 | 42.11 | 58.13 | 57.13 | 59.73 | 84.54 | 70.74 | 76.57 | 64.45 | 68.28 | 59.57 | + |
| R3hdm2 | 3.06 | 2.42 | 2.7 | 4.29 | 4.07 | 4.88 | 4.51 | 4.42 | 4.51 | 3.41 | 5.05 | 5.2 | + |
| Notch2 | 13.34 | 12.03 | 13.07 | 6.85 | 7.67 | 6.34 | 4.27 | 6.17 | 5.62 | 4.11 | 3.97 | 3.56 | + |
| Hmgb1 | 298.95 | 351.52 | 307.92 | 193.36 | 184.32 | 184.62 | 157.27 | 141.26 | 157.89 | 132.77 | 127.33 | 137.52 | + |
| Glrp1 | 1.87 | 1.93 | 2.27 | 0.81 | 1.45 | 1.66 | 2.08 | 2.1 | 2.06 | 0.53 | 0.47 | 0.88 | + |
| Cltb | 9.63 | 11.33 | 11.49 | 4.05 | 4.79 | 4.41 | 5.94 | 7.23 | 7.35 | 5.08 | 3.87 | 3.04 | + |
| Ctnnbl1 | 33.11 | 31.66 | 32.06 | 21.33 | 23.27 | 23.61 | 21.27 | 20.98 | 21.24 | 18.32 | 19.95 | 19.37 | + |
| Dnajc24 | 24.45 | 23.91 | 22.71 | 18.14 | 18.07 | 18.78 | 13.34 | 15.1 | 17.32 | 14.69 | 13.82 | 15.19 | + |
| Oxsr1 | 5.28 | 5.44 | 6.1 | 5.92 | 6.56 | 6.8 | 4.22 | 4.34 | 4.05 | 7.33 | 8.03 | 6.65 | + |
| Prim1 | 55.26 | 52.83 | 54.81 | 33.27 | 41.74 | 40.35 | 26.69 | 27.24 | 27.26 | 20.08 | 23 | 22.72 | + |
| Rdh14 | 3.57 | 4.82 | 6.67 | 9.79 | 12.64 | 10.87 | 10.74 | 8.98 | 8.35 | 9.44 | 6.91 | 12.69 | + |
| Tmem221 | 2.97 | 3.21 | 3.02 | 8.01 | 6.04 | 6.05 | 9.75 | 7.98 | 9.69 | 10.26 | 6.25 | 8.76 | + |
| Dkc1 | 121.82 | 129.89 | 127.24 | 50.45 | 52.12 | 53.51 | 31.52 | 53.72 | 45.44 | 29.57 | 27.77 | 33.22 | + |
| Gtf2f2 | 51.19 | 55.1 | 46.44 | 33.9 | 36.36 | 37.04 | 44.17 | 39.64 | 38.25 | 31.54 | 35.88 | 34.38 | + |
| Kcnab2 | 34.4 | 32.65 | 36.17 | 26.86 | 25.8 | 24.08 | 30.82 | 34.12 | 32.41 | 25.56 | 27.34 | 28.06 | + |

| | | | | | | | | | | | | | |
|---|---|---|---|---|---|---|---|---|---|---|---|---|---|
| Slc44a2 | 63.27 | 66.31 | 63.86 | 135.69 | 120.38 | 125.99 | 82.13 | 82.96 | 86.07 | 105.97 | 105.64 | 113.74 | + |
| Gins1 | 13.32 | 16.68 | 15.49 | 4.31 | 7.4 | 6.61 | 4.77 | 5.7 | 4.01 | 3.58 | 2.36 | 3.44 | + |
| Ablim2 | 2.03 | 1.33 | 1.91 | 12.01 | 9.84 | 12.27 | 11.45 | 9.25 | 10.56 | 13.99 | 14.14 | 14.95 | + |
| Adamtsl4 | 0.07 | 0.08 | 0.11 | 1.51 | 1.15 | 0.99 | 0.46 | 0.62 | 1 | 1.8 | 1.47 | 1.31 | + |
| Snrpb | 419.84 | 450.14 | 417.74 | 213.91 | 211.86 | 197.54 | 168.39 | 202.7 | 184.73 | 159.95 | 150.67 | 157.6 | + |
| Tmem68 | 18.17 | 19.5 | 20.09 | 12.4 | 11.81 | 12.16 | 12.31 | 12.39 | 13.03 | 10.23 | 11.21 | 9.32 | + |
| Rnf19a | 9.93 | 8.55 | 9.95 | 17.66 | 18.31 | 20.12 | 20.67 | 18.06 | 19.2 | 23.01 | 29.03 | 25.93 | + |
| Cdca4 | 65.67 | 60.23 | 63.23 | 45.04 | 44.15 | 48.14 | 41.41 | 41.49 | 43.34 | 42.19 | 41.63 | 44.18 | + |
| Hras1 | 2.51 | 3.47 | 2.3 | 1.44 | 1.74 | 1.57 | 1.03 | 1.03 | 1.07 | 1.41 | 0.99 | 1.53 | + |
| Dnajc9 | 103.63 | 104.7 | 99.42 | 80.47 | 89.48 | 82.38 | 66.17 | 67.82 | 65.31 | 59.07 | 60.55 | 61.85 | + |
| Rpl22 | 124.19 | 128.78 | 130.52 | 211.84 | 202.95 | 215.44 | 214.23 | 191.85 | 196.91 | 220.99 | 244.68 | 227.01 | + |
| Rpl36a | 980.44 | 966.39 | 978.26 | 1604.16 | 1605.4 | 1670.52 | 1495.49 | 1400.59 | 1463 | 1650.48 | 1786.08 | 1625.66 | + |
| Spcs3 | 105.3 | 100.88 | 105.55 | 67.48 | 64.66 | 66.42 | 69.6 | 82.27 | 78.1 | 58.06 | 53.3 | 54.56 | + |
| Cox17 | 423.97 | 407.21 | 406.25 | 304.96 | 313.97 | 319.44 | 241.7 | 287.45 | 270.78 | 239.06 | 229.7 | 236.94 | + |
| Chchd4 | 88.27 | 82.4 | 99.69 | 34.01 | 38.81 | 35.96 | 28.9 | 44.72 | 36.63 | 24.35 | 26.07 | 22.27 | + |
| Mtch2 | 125.15 | 124 | 119.38 | 81.22 | 82.87 | 85.25 | 78.62 | 94.37 | 86.54 | 70.59 | 70.75 | 69.2 | + |
| Atp5l | 331.63 | 344.33 | 319.47 | 223.42 | 228.74 | 230.6 | 236.47 | 236.69 | 242.42 | 215.51 | 220.81 | 231.53 | + |
| Ddhd2 | 13.03 | 12.29 | 13.24 | 16.76 | 14.57 | 14.46 | 14.59 | 13.56 | 13.64 | 17.76 | 19.64 | 18.89 | + |
| Zfp52 | 22.63 | 18.79 | 21.63 | 29.95 | 33.53 | 30.58 | 28.08 | 32.64 | 30.2 | 31.78 | 38.14 | 32.58 | + |
| Tax1bp3 | 11.32 | 8.54 | 9.77 | 23.1 | 21.32 | 23.64 | 29.48 | 20.31 | 24.12 | 31.33 | 27.1 | 30.35 | + |
| Il21r | 88.83 | 83.74 | 82.94 | 100.54 | 109.29 | 98.81 | 115.79 | 126.25 | 121.34 | 98.26 | 105.23 | 107.73 | + |
| Fndc3a | 6.14 | 6.46 | 7.17 | 18.14 | 16.99 | 17.02 | 13.9 | 10.77 | 13.04 | 14.69 | 15.5 | 14.15 | + |
| Stap1 | 32.06 | 28.33 | 29.08 | 25.43 | 23.12 | 23.24 | 24.24 | 25.69 | 24.8 | 22.86 | 22.35 | 23.34 | + |
| Med21 | 82.33 | 80.72 | 82.81 | 53.97 | 62.98 | 52.04 | 62.11 | 62.27 | 67.6 | 51.63 | 55.39 | 56.02 | + |
| Gata3 | 7.23 | 6.3 | 8.76 | 15.02 | 14.33 | 13.23 | 4.47 | 3.42 | 4.24 | 4.02 | 3.47 | 4.16 | + |
| Wdr76 | 4.3 | 4.98 | 5.05 | 2.84 | 3.02 | 3.43 | 2.22 | 2.24 | 2.59 | 2.42 | 2.4 | 2.55 | + |
| Itprip | 5.08 | 4.73 | 3.83 | 5.09 | 5.45 | 6.06 | 4.33 | 4.06 | 4.15 | 6.61 | 5.73 | 7.03 | + |
| Map4k4 | 2.97 | 2.98 | 3.28 | 5.6 | 5.45 | 6.94 | 4.1 | 4.51 | 3.8 | 4.7 | 5.93 | 5.09 | + |
| Prpf8 | 28.07 | 27.35 | 29.7 | 22.12 | 21.93 | 24.76 | 20.28 | 25.03 | 23.58 | 22 | 24.5 | 20.55 | + |
| Crtap | 28.34 | 27.51 | 27.03 | 28.12 | 30.1 | 28.21 | 27.02 | 27.18 | 26.47 | 20.56 | 22.11 | 21.76 | + |
| Gcn1l1 | 15.12 | 15.5 | 16.45 | 10.21 | 11.26 | 11.42 | 8.92 | 11.54 | 11.41 | 9.49 | 8.46 | 9.08 | + |
| Smtnl2 | 0.22 | 0.15 | 0.42 | 1.91 | 0.87 | 1.26 | 4.28 | 1.71 | 3.45 | 2.58 | 4.2 | 3.69 | + |
| Ppih | 56.44 | 60.65 | 58.89 | 22.99 | 22.06 | 24.11 | 15.63 | 17.45 | 19.46 | 10.18 | 14.84 | 13.15 | + |
| Zfp106 | 11.56 | 11.2 | 11.93 | 7.9 | 9.04 | 8.77 | 7.39 | 8.9 | 7.95 | 6.77 | 7.01 | 6.4 | + |
| Rps3 | 1132.79 | 1112.5 | 1133.01 | 1719.74 | 1688.16 | 1734.32 | 1574.62 | 1540.03 | 1536.01 | 1726.61 | 1815.4 | 1723.05 | + |
| Spred3 | 0.28 | 0.2 | 0.21 | 1.2 | 1.08 | 1.31 | 1.6 | 1.23 | 1.03 | 1.59 | 1.49 | 1.04 | + |
| Mlh1 | 19.07 | 20.44 | 18.69 | 10.9 | 11.99 | 11.93 | 9.74 | 11.57 | 9.5 | 7.31 | 6.7 | 6.35 | + |
| Rbmxl1 | 128.5 | 136.59 | 137.46 | 66.83 | 73.96 | 70.75 | 51.37 | 67.07 | 60.61 | 56.82 | 55.24 | 51.09 | + |
| Kdelr1 | 22.76 | 23.11 | 23.54 | 45.37 | 40.73 | 44.2 | 51.29 | 43.02 | 49.01 | 58.21 | 63 | 64.52 | + |
| Mlx | 54.63 | 50.08 | 47.63 | 33.58 | 32.68 | 33.17 | 31.74 | 34.74 | 35.72 | 28.77 | 34.17 | 32.18 | + |
| Hsph1 | 89.02 | 88.16 | 95.48 | 24.53 | 35.07 | 34.37 | 27.33 | 50.8 | 39.64 | 22.86 | 31.03 | 22.28 | + |
| Myo1h | 0.08 | 0.03 | 0.23 | 0.64 | 0.27 | 0.71 | 0.66 | 0.71 | 0.58 | 0.44 | 0.76 | 0.66 | + |
| Gatsl3 | 18.26 | 19.8 | 20.54 | 48.18 | 47.18 | 44.6 | 47.33 | 35.21 | 39.61 | 28.01 | 29.73 | 32.13 | + |
| Nt5c3 | 41.96 | 43.87 | 41.89 | 48.81 | 42.79 | 40.27 | 46.21 | 42.91 | 43.73 | 51.58 | 48.42 | 52.54 | + |
| Thap4 | 11.17 | 11.83 | 10.59 | 21.48 | 19.19 | 19.78 | 23.73 | 23.25 | 19.39 | 22.67 | 28.47 | 22.39 | + |
| Rab3d | 6.5 | 6.38 | 5.7 | 10.62 | 9.67 | 8.59 | 11.8 | 10.13 | 10.53 | 10.25 | 8.61 | 9.61 | + |
| Clpp | 76.97 | 73.61 | 74.97 | 35.35 | 38.63 | 39.68 | 31.23 | 45.95 | 42.98 | 29.57 | 27.83 | 28.61 | + |
| Dynlt3 | 28.9 | 27.14 | 28.66 | 36.87 | 35.11 | 37.63 | 38.01 | 42.03 | 37.95 | 38.83 | 37.07 | 39.87 | + |
| Kif2c | 14.91 | 13.07 | 12.99 | 10.41 | 11.25 | 11.58 | 7.09 | 6.18 | 5.5 | 6.33 | 6.63 | 6.72 | + |
| Rras2 | 5.09 | 5.96 | 5.32 | 5.05 | 4.2 | 4.32 | 4.7 | 3.43 | 5.6 | 2.16 | 2.41 | 1.9 | + |
| Gtf2ird1 | 1.54 | 1.64 | 2.14 | 3.45 | 2.7 | 2.71 | 2.79 | 3.11 | 2.45 | 2.62 | 2.91 | 2.81 | + |
| Zfp142 | 3.63 | 3.72 | 3.18 | 3.01 | 2.69 | 2.73 | 2.09 | 2.46 | 2.97 | 2.81 | 2.55 | 2.33 | + |
| Lyar | 99.52 | 103.92 | 102.51 | 31.5 | 35.16 | 34.75 | 28.27 | 45.01 | 37.12 | 22.31 | 21.42 | 17.69 | + |
| Tmsb4x | 3520.69 | 3277.74 | 2940.65 | 1926.57 | 1929.9 | 2054.2 | 3823.13 | 3679.82 | 3705.43 | 2529.75 | 2626.56 | 2533.61 | + |
| 3110043O21Rik | 6.67 | 6.24 | 6.66 | 9.63 | 12.37 | 11.1 | 11.93 | 18.42 | 14.19 | 10.8 | 13.38 | 13.25 | + |
| Ncapd2 | 34.05 | 33.72 | 31.62 | 30.39 | 33.66 | 28.04 | 22.93 | 23.04 | 18.22 | 23.95 | 25.4 | 19.63 | + |
| Vopp1 | 11 | 11.23 | 10.12 | 24.66 | 20.93 | 21.68 | 25.72 | 23.89 | 21.54 | 24.93 | 31.97 | 28.41 | + |
| Rpl27a | 864.24 | 876.29 | 872.11 | 1380.91 | 1341.26 | 1423.21 | 1480.09 | 1342.97 | 1384.6 | 1529.58 | 1669.98 | 1529.46 | + |
| Pole2 | 14.62 | 13.24 | 13.1 | 7.2 | 7.4 | 7.15 | 5.22 | 7.57 | 6.71 | 3.71 | 4.29 | 4.73 | + |
| Eif2b5 | 10.94 | 13.18 | 12.86 | 6.65 | 7.82 | 8.46 | 7.89 | 8.78 | 7.96 | 8.12 | 7.62 | 8.25 | + |
| Rrp7a | 96.06 | 99.58 | 96.13 | 58.69 | 59.56 | 59.87 | 50.98 | 58.19 | 58.13 | 49.48 | 48.32 | 50.49 | + |
| Hbp1 | 7.37 | 6.12 | 7.03 | 23.86 | 20.57 | 21.46 | 27.9 | 20.6 | 21.21 | 39.73 | 41.35 | 39.5 | + |
| Rbm28 | 25.1 | 24.01 | 23.49 | 16.25 | 18 | 17.27 | 15.15 | 16.38 | 15.09 | 13.9 | 13.81 | 12.59 | + |
| Atp11b | 17.82 | 20.97 | 25.47 | 42.61 | 41.32 | 38.48 | 28.42 | 26.22 | 32.09 | 31.05 | 32.43 | 28.44 | + |
| Vldlr | 0.54 | 0.36 | 0.45 | 2.45 | 1.93 | 1.83 | 2.17 | 1.86 | 2.47 | 1.76 | 2.36 | 2.68 | + |
| Ptgfrn | 0.61 | 0.65 | 0.84 | 1.35 | 1.28 | 1.72 | 0.28 | 0.3 | 0.42 | 0.36 | 0.23 | 0.27 | + |
| Dctn4 | 27.61 | 25.85 | 27.74 | 51.29 | 49.31 | 48.85 | 43.5 | 44.59 | 41.81 | 51.41 | 59.16 | 49.98 | + |
| Pcmtd1 | 3.81 | 3.43 | 3.81 | 6.33 | 5.22 | 5.89 | 7.51 | 6.05 | 8.4 | 7.59 | 7.22 | 7.13 | + |
| Ahsa2 | 21.59 | 21.34 | 21.72 | 12.77 | 15.84 | 14.82 | 12.63 | 13.86 | 14.5 | 11.64 | 11.48 | 11.2 | + |
| Rmdn2 | 1.55 | 1.19 | 1.23 | 0.21 | 0.48 | 0.18 | 0.22 | 0.26 | 0.57 | 0.35 | 0.34 | 0.16 | + |
| Syngr3 | 7.34 | 6.27 | 6.79 | 14.1 | 10.71 | 14.13 | 24 | 17.24 | 19.42 | 20.44 | 18.25 | 21.89 | + |
| Atp5j2 | 241.25 | 255.99 | 251.47 | 199.96 | 200.8 | 210.97 | 204.83 | 209.14 | 196.72 | 176.78 | 197.98 | 199.16 | + |
| Mospd1 | 7.1 | 6.93 | 6.5 | 7.93 | 5.73 | 7.28 | 8.05 | 7.54 | 7.65 | 13.35 | 9.33 | 11.84 | + |
| Slc7a4 | 1.7 | 1.21 | 0.8 | 3.11 | 2.42 | 1.82 | 3.93 | 3.54 | 3.29 | 2.5 | 2.38 | 2.57 | + |
| Gemin5 | 17.58 | 16.13 | 17.59 | 9.66 | 10.27 | 10.8 | 8.04 | 10.88 | 11.29 | 8.44 | 9.7 | 7.21 | + |
| Smarca4 | 21.84 | 23 | 24.67 | 15.4 | 15.55 | 15.11 | 12.66 | 14.46 | 13.57 | 10.4 | 12.16 | 10.66 | + |
| Lcmt2 | 24.75 | 27.94 | 24.64 | 9.66 | 10.76 | 10.54 | 10.09 | 12.58 | 10.24 | 7.91 | 8.85 | 8.96 | + |
| Cxcr2 | 12.46 | 13.96 | 12.58 | 4.96 | 5.36 | 5.68 | 5.93 | 6.53 | 5.29 | 5.17 | 4.99 | 5.01 | + |
| Gbp8 | 16.71 | 14.39 | 16.2 | 46.2 | 39.54 | 44.19 | 33.82 | 37.36 | 36.95 | 32.63 | 34.51 | 34.52 | + |
| Fbxo36 | 4.19 | 4.14 | 4.09 | 13.9 | 13.94 | 14.81 | 11.36 | 9.2 | 9.75 | 11.76 | 13.97 | 9.48 | + |
| Asns | 220.51 | 212.02 | 204.46 | 139.13 | 181.22 | 179.39 | 163.67 | 265.7 | 234.77 | 94.82 | 109.74 | 88.1 | + |
| Ppfia1 | 3.69 | 3.18 | 3.3 | 2.29 | 2.12 | 2.26 | 1.76 | 2.06 | 2.3 | 1.88 | 2.12 | 2.16 | + |
| Smim13 | 1.63 | 1.66 | 1.48 | 1.16 | 0.86 | 0.84 | 0.95 | 1.34 | 1.02 | 0.84 | 0.84 | 0.86 | + |
| Eif4a2 | 139.61 | 133.01 | 144.53 | 254.4 | 226.31 | 255.75 | 251.64 | 244.96 | 238.65 | 300.05 | 365.91 | 294.4 | + |
| Atp5o | 230.28 | 237.74 | 225.26 | 189.85 | 190.67 | 194.79 | 204.62 | 195.53 | 179.5 | 178.36 | 185.15 | 176.46 | + |
| Uhrf2 | 3.81 | 4.33 | 4.07 | 8.28 | 8.38 | 8.44 | 7.26 | 7.6 | 7.05 | 10.43 | 10.8 | 10.12 | + |
| Pdcd6 | 120.83 | 124.37 | 124.02 | 109.03 | 97.94 | 103.15 | 93.02 | 100.05 | 99.48 | 101.77 | 109.57 | 102.81 | + |
| Rgs10 | 6.72 | 7.85 | 5.62 | 20.44 | 13.7 | 12.54 | 27.06 | 10.87 | 17.23 | 14.52 | 10.42 | 13.31 | + |
| Rab14 | 49.48 | 49.02 | 48.6 | 44.43 | 42.82 | 45.41 | 35.2 | 38.7 | 37.89 | 37.04 | 39.07 | 39.03 | + |
| Ccna2 | 100.9 | 92.61 | 86.27 | 55.76 | 78.05 | 65.6 | 32.47 | 35.61 | 31.14 | 37.01 | 38.16 | 31.47 | + |
| Fbxl20 | 0.87 | 0.88 | 1.17 | 2.93 | 2.68 | 3.29 | 3.41 | 2.94 | 2.73 | 5.05 | 4.1 | 3.43 | + |
| Hcfc1 | 29.74 | 26.5 | 29.81 | 21.47 | 22.65 | 21.42 | 18.32 | 20.1 | 18.83 | 19.06 | 18.6 | 19.62 | + |
| Ino80c | 21.73 | 21.5 | 22.61 | 11.03 | 14.26 | 13.24 | 13.85 | 15.13 | 13.65 | 11.83 | 10.58 | 11.5 | + |
| Me2 | 54.19 | 60.37 | 53.55 | 33.06 | 32.85 | 31.01 | 63.19 | 35.7 | 45.51 | 35.93 | 39.19 | 42.58 | + |
| Gart | 101.88 | 101.68 | 103.73 | 39.01 | 44.67 | 40.07 | 29.37 | 46.6 | 42.28 | 25.33 | 22.93 | 24.77 | + |
| Tbcc | 8.67 | 10.29 | 10.36 | 6.34 | 9.42 | 7.72 | 5.18 | 6.18 | 5.88 | 6.64 | 5.58 | 6.63 | + |
| Itm2c | 24.79 | 23.92 | 24.28 | 61.39 | 57.41 | 54.48 | 57.21 | 53.68 | 51.9 | 54.86 | 58.34 | 58.41 | + |
| Ptpn7 | 58.17 | 57.55 | 53.6 | 33.91 | 37.02 | 36.6 | 39.53 | 38.81 | 35.79 | 32.43 | 33.25 | 36.32 | + |
| Ptrhd1 | 4.21 | 3.58 | 3.79 | 2.2 | 2.47 | 2.41 | 2.54 | 3.12 | 2.9 | 2.04 | 2.24 | 2.17 | + |
| Mtfmt | 21.38 | 19.06 | 21.88 | 16.34 | 17.42 | 13.79 | 13.97 | 15.47 | 15.61 | 14.52 | 14.45 | 13.67 | + |
| Rac2 | 301.93 | 306.63 | 284.33 | 441.43 | 425.46 | 433.19 | 519.31 | 449.62 | 479.76 | 518.08 | 519.83 | 555.1 | + |
| 1600029O15Rik | 4.96 | 5.46 | 5.99 | 9.54 | 7.46 | 8.57 | 9.28 | 7.59 | 9.16 | 8.25 | 8.77 | 9.85 | + |
| Atcay | 1.83 | 1.61 | 1.56 | 1.77 | 2.05 | 1.16 | 1.22 | 2.72 | 2.1 | 0.98 | 0.96 | 1 | + |
| Tfr2 | 0.45 | 0.26 | 0.63 | 0.8 | 1 | 0.83 | 1.63 | 1.91 | 1.12 | 1.49 | 1.26 | 1.64 | + |
| Srsf1 | 17.83 | 16.85 | 17.83 | 8.82 | 8.83 | 8.97 | 7.18 | 7.47 | 9.59 | 8.52 | 8.38 | 7.54 | + |
| Usp35 | 1.05 | 0.92 | 0.43 | 1.87 | 2.18 | 2.63 | 4.4 | 2.83 | 2.87 | 3.1 | 3.25 | 2.56 | + |
| Ppap2c | 16.75 | 12.94 | 13.92 | 20.54 | 21.19 | 23.75 | 32.07 | 33.91 | 34.03 | 24.85 | 23.59 | 28.4 | + |
| Cnot7 | 53.44 | 52.87 | 49.29 | 37.08 | 37.14 | 36.55 | 37.16 | 41.04 | 41.35 | 34.06 | 35.65 | 34.87 | + |
| Pggt1b | 7.89 | 8.17 | 8.66 | 4.25 | 5.25 | 5.41 | 4.58 | 5.78 | 5.37 | 4.66 | 5.7 | 3.96 | + |
| Pycrl | 36.15 | 36.86 | 40.95 | 22.04 | 22.15 | 19.8 | 20.54 | 22.66 | 19.11 | 21.46 | 21.04 | 20.62 | + |
| Sf3b2 | 82.55 | 85.72 | 90.84 | 78.25 | 75.36 | 75.75 | 74.3 | 71.62 | 68.37 | 69.48 | 74.4 | 69.15 | + |
| Haus5 | 28.87 | 22.77 | 24.59 | 11.44 | 15.56 | 14.06 | 6.83 | 9.69 | 9.85 | 6.1 | 6.26 | 7.77 | + |
| Tpp2 | 38.73 | 40.4 | 41.17 | 30.57 | 31.04 | 30.56 | 31.66 | 29.12 | 30.59 | 29.41 | 33.59 | 28.37 | + |
| Traf3ip2 | 12.29 | 11.63 | 12.6 | 16.76 | 18.01 | 18.01 | 19.2 | 19.11 | 19.24 | 18.52 | 17.18 | 22 | + |
| Leng8 | 14.56 | 16.31 | 15.8 | 37.41 | 31.12 | 30.11 | 40.5 | 32.47 | 34.23 | 37.29 | 44.63 | 38.94 | + |
| Ifnar2 | 20.16 | 22.64 | 25.65 | 37.57 | 35.86 | 38.71 | 33.55 | 33.84 | 33.06 | 35.74 | 38.55 | 32.43 | + |
| Anapc1 | 7.78 | 7.37 | 9.11 | 5.83 | 5.15 | 5.73 | 5.73 | 6.07 | 5.67 | 4.65 | 5.85 | 5.03 | + |
| Itga7 | 0.07 | 0.07 | 0.04 | 1.51 | 0.36 | 0.35 | 8.99 | 1.48 | 5.08 | 2.32 | 3.53 | 3.63 | + |
| Ppp1r8 | 14.65 | 16.59 | 14.81 | 8.61 | 8.7 | 10.31 | 7.43 | 10.68 | 7.73 | 7.43 | 7.64 | 6.07 | + |
| Slc30a6 | 11.02 | 11.19 | 11.03 | 8.77 | 7.82 | 7.13 | 6.61 | 7.79 | 7.07 | 5.68 | 7.4 | 7.3 | + |
| Laptm4a | 19.99 | 17.76 | 15.72 | 29.63 | 32.28 | 30.68 | 25.42 | 25.25 | 23.64 | 25.23 | 29.84 | 23.6 | + |
| Eif3d | 206.54 | 208.35 | 206.67 | 163.86 | 174.17 | 175.56 | 159.6 | 174.54 | 172.45 | 153.47 | 158.96 | 156.15 | + |
| Aqr | 13.72 | 14.32 | 15.56 | 10.97 | 10.32 | 10.68 | 8.81 | 10.06 | 10.93 | 10.18 | 10.95 | 9.16 | + |
| Ankfy1 | 16.57 | 14.45 | 16.56 | 13.19 | 14.13 | 14.38 | 13.55 | 13.24 | 13.06 | 13.45 | 13.41 | 13.19 | + |
| N4bp1 | 1.37 | 1.25 | 1.04 | 3.62 | 3.6 | 3.68 | 2.03 | 2.06 | 2.12 | 3.9 | 3.96 | 4.25 | + |
| Psmc1 | 166.65 | 171.4 | 170.24 | 88.28 | 93.54 | 93.48 | 73.95 | 82.72 | 75.01 | 68.1 | 68.98 | 62.72 | + |
| Pdss2 | 9.57 | 9.69 | 8.36 | 4.47 | 5.06 | 4.55 | 6.25 | 6.05 | 6.2 | 2.98 | 4.05 | 4.36 | + |
| Hmg20b | 10.14 | 10.9 | 10.99 | 16.87 | 17.58 | 16.1 | 16.66 | 15.31 | 16.45 | 15.15 | 16.16 | 14.68 | + |
| Dpf2 | 31.62 | 33.13 | 31.82 | 44.39 | 44.74 | 44.88 | 38.73 | 43.05 | 41.01 | 38.15 | 42.04 | 40.97 | + |
| Zfp110 | 11.47 | 13.91 | 11.26 | 18.3 | 18.08 | 16.97 | 15.74 | 16.65 | 15.72 | 18.25 | 19.4 | 16.89 | + |
| Cd93 | 0.47 | 0.61 | 0.46 | 0.41 | 0.92 | 0.96 | 4.47 | 4.63 | 3.92 | 2.58 | 2.17 | 2.77 | + |
| Rnd1 | 3.59 | 2.42 | 3.85 | 2.91 | 3.36 | 2.36 | 2.55 | 3.32 | 4.24 | 1.16 | 1.39 | 1.58 | + |
| Tle2 | 0.47 | 0.46 | 0.58 | 3.53 | 2.54 | 2.51 | 3.89 | 2.41 | 2.37 | 4.08 | 3.58 | 5.13 | + |
| Rps6 | 1560.88 | 1629.82 | 1569.9 | 2427.7 | 2379.54 | 2512.29 | 2290.95 | 2157.14 | 2214.56 | 2455.77 | 2652.7 | 2423.88 | + |
| Zfyve26 | 1.52 | 1.61 | 1.46 | 2.39 | 2.17 | 2.57 | 2.16 | 2.14 | 2.09 | 2.48 | 2.51 | 2.66 | + |
| Adamts10 | 4.34 | 4.67 | 4.75 | 11.61 | 9.01 | 8.59 | 12.68 | 8.32 | 8.84 | 13.8 | 12.95 | 13.37 | + |
| Ppp5c | 99.48 | 98.74 | 97.54 | 48.48 | 46.79 | 49.62 | 46.1 | 57.43 | 51.66 | 40.62 | 42.6 | 41.31 | + |
| Zfp932 | 42.32 | 45.57 | 44.87 | 34.17 | 36.05 | 34.77 | 25.7 | 33.08 | 30.78 | 32.37 | 29.43 | 29.74 | + |
| Ctps | 56.09 | 60.92 | 60.33 | 25.11 | 27.42 | 27.47 | 24.02 | 36.83 | 30.7 | 17.55 | 19.22 | 17.17 | + |
| Slc38a1 | 20.19 | 21.8 | 20.41 | 16.01 | 15.1 | 17.1 | 14.02 | 16.28 | 16.36 | 15.22 | 15.08 | 14.65 | + |
| Ugt1a10 | 13.83 | 6.3 | 9.5 | 0.19 | 0.09 | 0.1 | 0.34 | 2.18 | 1.27 | 0.05 | 0.15 | 0.2 | + |
| Rbm3 | 574.9 | 615.5 | 572.82 | 356.73 | 362.31 | 356.8 | 394.91 | 373.68 | 395.57 | 316.71 | 303.69 | 319.9 | + |
| Camsap1 | 2.25 | 1.82 | 2.11 | 1.09 | 1.13 | 1.29 | 0.89 | 1.14 | 1.56 | 1.3 | 1.2 | 1.14 | + |
| Hdac3 | 56.46 | 58.66 | 56.89 | 39.26 | 39.07 | 36.76 | 32.42 | 34.51 | 32 | 31.81 | 28.37 | 32 | + |
| Tmem33 | 27.01 | 27.39 | 28.48 | 16.2 | 17.84 | 18.43 | 15.59 | 21.04 | 19.59 | 16.14 | 16.97 | 17.03 | + |
| Clpb | 17.65 | 16.19 | 15.29 | 14.25 | 15.58 | 14.96 | 13.83 | 15.25 | 15.55 | 10.06 | 10.99 | 10.66 | + |
| Fbxl8 | 8.38 | 7.81 | 8.42 | 19.55 | 19.21 | 20.43 | 37.15 | 21.41 | 25.6 | 36.62 | 34.79 | 39.72 | + |
| Vps53 | 21.14 | 19.86 | 20.72 | 10.41 | 13.88 | 11.59 | 11.77 | 13.87 | 13.83 | 11.69 | 10.92 | 8.82 | + |
| Ttc39c | 20.68 | 19.05 | 24.03 | 5.91 | 8.79 | 7.49 | 6.53 | 18.1 | 12.33 | 1.49 | 2.22 | 1.57 | + |
| Tubb6 | 187.77 | 155.8 | 153.08 | 68.65 | 77.49 | 79.63 | 84.65 | 72.77 | 80.74 | 55.17 | 59.02 | 66.15 | + |
| D16Ertd472e | 20.4 | 21.16 | 21.19 | 26.61 | 27.03 | 28.66 | 31.11 | 36.2 | 36.63 | 28.92 | 29.57 | 31.34 | + |

| | | | | | | | | | | | | | |
|---|---|---|---|---|---|---|---|---|---|---|---|---|---|
| Metrnl | 0.28 | 0.5 | 0.33 | 1.9 | 1.62 | 1.65 | 0.81 | 0.93 | 0.83 | 1.6 | 1.35 | 1.61 | + |
| Ankrd9 | 3.55 | 3.12 | 2.73 | 2.15 | 1.59 | 2.57 | 2.18 | 1.58 | 1.85 | 1.5 | 1.76 | 2.28 | + |
| Tap2 | 73.44 | 79.85 | 73.55 | 86.91 | 80.3 | 82.72 | 93.45 | 82.36 | 83.28 | 123.88 | 104.63 | 123.65 | + |
| Napb | 0.39 | 0.13 | 0.25 | 0.94 | 0.78 | 0.81 | 0.72 | 0.94 | 0.67 | 0.83 | 1.35 | 0.92 | + |
| Pigt | 8.42 | 9.49 | 7.84 | 5.64 | 5.08 | 6.83 | 6.39 | 6.66 | 6.08 | 5.77 | 5.28 | 5.18 | + |
| Tet2 | 2.37 | 2.17 | 2.31 | 6.93 | 7.46 | 7.53 | 5.76 | 4.92 | 5 | 8.36 | 10.23 | 9.56 | + |
| Eftud2 | 82.84 | 83.47 | 85.62 | 34.72 | 38.84 | 37.3 | 26.32 | 37.85 | 36.08 | 27.98 | 23.97 | 28.13 | + |
| Pdia3 | 325.46 | 328.54 | 306.43 | 210.47 | 211.97 | 216.27 | 177.26 | 193.42 | 183.96 | 190.27 | 202.3 | 191.19 | + |
| Vmp1 | 52.77 | 54.3 | 56.47 | 79.47 | 77.43 | 86.89 | 87.08 | 79.29 | 77.89 | 101.15 | 92.66 | 92.37 | + |
| Sord | 9.5 | 7.81 | 6.63 | 13.09 | 14.58 | 14.26 | 29.66 | 18.15 | 19.47 | 19.14 | 24.61 | 18.1 | + |
| Zfp300 | 1.76 | 1.59 | 1.68 | 2.26 | 2.18 | 2.16 | 1.73 | 2.09 | 1.76 | 2.85 | 2.84 | 2.7 | + |
| Pdcl | 18.89 | 18.08 | 18.28 | 12.91 | 14.2 | 14.01 | 12.89 | 14.51 | 15.41 | 12.11 | 13.37 | 13.13 | + |
| Dab2 | 0.14 | 0.15 | 0.14 | 0.72 | 1.12 | 0.53 | 0.41 | 0.46 | 0.72 | 0.63 | 0.6 | 0.48 | + |
| Zdhhc5 | 1.84 | 2.14 | 2 | 1.28 | 0.91 | 1.09 | 1.19 | 1.45 | 1.16 | 1.07 | 0.99 | 1.18 | + |
| Lmf2 | 28.16 | 30.38 | 27 | 23.8 | 26.62 | 24.71 | 23.43 | 22.6 | 21.06 | 18.48 | 23.58 | 22.45 | + |
| Poldip3 | 37.99 | 36.09 | 36.89 | 41.58 | 39.09 | 38.43 | 36.38 | 32.18 | 34.22 | 40.26 | 48.58 | 42.47 | + |
| Usp46 | 5.77 | 4.76 | 5.96 | 3 | 3.48 | 3.87 | 2.76 | 2.62 | 3.15 | 2.48 | 2.3 | 2.47 | + |
| Mppe1 | 6.4 | 6.98 | 6.95 | 10.37 | 10.31 | 9.49 | 10.89 | 11.97 | 10.93 | 13.02 | 11.47 | 9.61 | + |
| Carns1 | 0.81 | 1.01 | 0.86 | 4.52 | 3.16 | 2.84 | 3.17 | 2.09 | 2.46 | 5.47 | 4.58 | 5.17 | + |
| Pebp1 | 317.08 | 286.35 | 304.95 | 229.49 | 242.46 | 249.44 | 266.41 | 272.8 | 275.22 | 220.33 | 246.25 | 224.71 | + |
| Prpf40a | 47.64 | 47.75 | 49.47 | 36.2 | 38.22 | 43.01 | 32.45 | 37.45 | 36.27 | 31.04 | 35.87 | 30.45 | + |
| Bet1l | 14.75 | 13.31 | 14.74 | 20.44 | 19.7 | 21.08 | 22.73 | 23.77 | 20.74 | 21.38 | 22.71 | 22.21 | + |
| Mtfr1 | 33.46 | 34.5 | 33.88 | 18.54 | 21.2 | 22.94 | 20.66 | 26.47 | 25.21 | 18.09 | 16.94 | 17.78 | + |
| Rpl30 | 162.6 | 153.42 | 129.79 | 52.2 | 57.48 | 57.16 | 39.33 | 77.81 | 58.14 | 36.61 | 29.09 | 30.79 | + |
| Rbm19 | 8.71 | 8.55 | 8.85 | 4.72 | 3.25 | 4.03 | 2.54 | 4.2 | 3.54 | 3.12 | 1.94 | 2.78 | + |
| Smn1 | 20.2 | 24.41 | 25.45 | 10.38 | 13.52 | 13.34 | 11.02 | 15.72 | 12.22 | 9.71 | 12.51 | 7.91 | + |
| Naa20 | 135.17 | 130.49 | 132.04 | 69.87 | 76.84 | 74.56 | 61.63 | 64.02 | 64.41 | 81.83 | 69.8 | 79.18 | + |
| Aldh18a1 | 82.33 | 77.18 | 79.65 | 62.12 | 70.02 | 64.05 | 68.1 | 90.26 | 85.33 | 46.59 | 47.52 | 49.6 | + |
| Slc3a2 | 124.51 | 133.56 | 126.15 | 104.78 | 132.32 | 128.48 | 131.13 | 194.84 | 177.14 | 100.96 | 113.96 | 97.26 | + |
| Ypel5 | 10.57 | 8.43 | 9.27 | 19.87 | 17.42 | 19.05 | 22.17 | 21.26 | 21.84 | 22.8 | 23.77 | 23.55 | + |
| Ccpg1 | 7.59 | 7.46 | 7.18 | 14.78 | 13.24 | 12.74 | 17.1 | 16.24 | 12.83 | 17.54 | 16.83 | 18.32 | + |
| Fkbp2 | 65.01 | 72.52 | 65.68 | 33.74 | 37.47 | 44.32 | 30.81 | 37.76 | 37.56 | 26.84 | 27.74 | 28.23 | + |
| Ap4s1 | 11.9 | 12.6 | 11.9 | 7.21 | 7.52 | 8.59 | 7.64 | 6.6 | 6.22 | 7.81 | 7.36 | 6.37 | + |
| Mrpl37 | 56.89 | 59.48 | 59.2 | 31.48 | 30.75 | 31.5 | 31.54 | 33.84 | 31.64 | 25.85 | 25.77 | 25.89 | + |
| Mocos | 4.39 | 3.83 | 3.3 | 6.81 | 6.15 | 6.48 | 7.26 | 8.12 | 7.7 | 6.37 | 5.85 | 4.91 | + |
| Purb | 3.89 | 4.41 | 4.21 | 3.4 | 2.88 | 3.54 | 2.9 | 2.92 | 3.27 | 3.22 | 3.01 | 3.17 | + |
| Gtf2h1 | 40.31 | 40.04 | 41.4 | 19.1 | 18.44 | 21.33 | 16.48 | 20.24 | 20.83 | 16.92 | 16.35 | 17.89 | + |
| Dusp22 | 24.44 | 23.85 | 26.45 | 49.16 | 60.89 | 58.75 | 49.17 | 52.12 | 49.03 | 61.42 | 81.4 | 66.33 | + |
| Mrps17 | 67.04 | 63.28 | 67.37 | 40.38 | 48.27 | 45.95 | 38.04 | 44.83 | 43.78 | 36.38 | 37.62 | 33.73 | + |
| Cacnb3 | 2.57 | 2.72 | 2.17 | 4.99 | 4.75 | 4.71 | 8.24 | 5.54 | 6.45 | 8.53 | 7.52 | 6.09 | + |
| Mrps6 | 156.99 | 137.33 | 155.47 | 91.95 | 105.23 | 103.81 | 123.81 | 147.05 | 137.69 | 109.12 | 125.15 | 109.95 | + |
| Tango6 | 3.19 | 4.58 | 3.77 | 2.52 | 2.76 | 1.85 | 2.33 | 2.9 | 2.46 | 2.37 | 2.21 | 2.77 | + |
| Kctd11 | 2.83 | 2.59 | 2.09 | 4.76 | 5.41 | 3.82 | 9.67 | 8.42 | 9.72 | 4.48 | 5.5 | 6.18 | + |
| Trmt12 | 2.87 | 2.73 | 2.93 | 0.64 | 1.27 | 1.17 | 0.72 | 0.9 | 1.06 | 1.02 | 0.45 | 0.71 | + |
| Pcnx | 0.89 | 0.69 | 0.76 | 1.42 | 1.56 | 1.7 | 1.29 | 1.43 | 1.48 | 1.51 | 1.7 | 1.51 | + |
| Ikzf1 | 27.9 | 27.25 | 27.5 | 34.85 | 32.85 | 34.58 | 30.4 | 27.98 | 28.55 | 41.37 | 39.37 | 41.48 | + |
| Cep68 | 6.09 | 5.96 | 6.11 | 6.45 | 6.6 | 6.14 | 7.26 | 5.03 | 6.54 | 8.72 | 8.63 | 8.65 | + |
| Poc1a | 15.49 | 17.59 | 16.85 | 8.66 | 10.93 | 10.2 | 5.92 | 8.02 | 6.1 | 5.19 | 6.5 | 4.76 | + |
| Sirt3 | 12.35 | 15.37 | 12.99 | 23.34 | 21.35 | 22.55 | 25.14 | 22.4 | 21.07 | 27.4 | 31.04 | 26.84 | + |
| Anapc15 | 56.99 | 52.95 | 56.01 | 25.79 | 31.72 | 31.53 | 29.92 | 31.78 | 33.42 | 22.69 | 23.91 | 17.03 | + |
| Ftsjd2 | 47.9 | 44.04 | 49.83 | 43.51 | 44.7 | 44.76 | 32.6 | 40.51 | 36.14 | 43.05 | 39.5 | 38.9 | + |
| Hexdc | 1.07 | 0.95 | 0.96 | 4.01 | 2.86 | 3.52 | 3.65 | 1.53 | 3.01 | 2.05 | 1.91 | 3.03 | + |
| Dnajc15 | 191.97 | 195.58 | 177.78 | 105.53 | 107.98 | 116.12 | 166.81 | 150.25 | 144.32 | 133.92 | 111.86 | 125.05 | + |
| Pfdn4 | 33.85 | 38.38 | 35.46 | 14.68 | 16.47 | 14.59 | 18.98 | 21.65 | 26.07 | 14 | 14.59 | 12.41 | + |
| Prrc1 | 13.26 | 14.07 | 13.45 | 19.09 | 20.19 | 20.94 | 22.4 | 25.08 | 23.72 | 20.73 | 27.79 | 21.72 | + |
| Btg1 | 3.52 | 3.63 | 3.84 | 14.7 | 14.95 | 14.96 | 11.96 | 9.42 | 11.68 | 17.82 | 20.01 | 17.09 | + |
| Cyth4 | 30.58 | 29.98 | 29.12 | 31.55 | 25.39 | 26.41 | 37.78 | 30.39 | 33.46 | 41.79 | 36.45 | 38.83 | + |
| Pygl | 4.53 | 4.35 | 3.72 | 23.25 | 15.31 | 15.46 | 30.55 | 13.79 | 22.82 | 24.47 | 17.95 | 21.95 | + |
| BC035044 | 5.44 | 6.1 | 5.53 | 1.09 | 1.16 | 1.27 | 1.98 | 4.7 | 4.09 | 2.56 | 2.41 | 3.16 | + |
| Map6 | 15.46 | 16.84 | 13.29 | 14.64 | 10.21 | 10.75 | 15.92 | 15.06 | 16.55 | 8.7 | 8.73 | 6.59 | + |
| Pold3 | 31.33 | 30.15 | 29.45 | 23.39 | 26.51 | 27.36 | 19.43 | 18.1 | 19.42 | 21.4 | 23.51 | 21.56 | + |
| Cetn2 | 53.47 | 50.29 | 52.65 | 42.24 | 37.85 | 38.5 | 40.22 | 40.96 | 44.38 | 38.88 | 37.1 | 40.87 | + |
| BC003965 | 13.29 | 14.1 | 14.03 | 7.99 | 8.49 | 7.34 | 6.5 | 8.39 | 7.49 | 5.75 | 5.99 | 6.75 | + |
| Dzip1 | 0.26 | 0.48 | 0.69 | 2.16 | 1.46 | 1.51 | 0.87 | 0.85 | 0.95 | 0.83 | 1.06 | 1.48 | + |
| Pdlim5 | 2.12 | 2.43 | 2.35 | 3.34 | 2.35 | 2.41 | 2.44 | 2.75 | 2.27 | 4.07 | 4.29 | 3.73 | + |
| Psmb8 | 765.75 | 781.17 | 744.14 | 796.02 | 777.66 | 808.04 | 846.63 | 804.9 | 794.55 | 866.16 | 876.71 | 932.46 | + |
| Mri1 | 31.67 | 31.52 | 26.69 | 31.5 | 34.6 | 33.98 | 44.26 | 44.09 | 43.27 | 38.68 | 37.84 | 38.51 | + |
| Ctdsp2 | 2.02 | 1.82 | 1.63 | 4.6 | 4.38 | 5.1 | 7.75 | 4.32 | 5.87 | 8.06 | 7.09 | 7.61 | + |
| Gm20098 | 0.1 | 0.32 | 0.16 | 1.24 | 0.8 | 0.81 | 0.91 | 1.02 | 0.74 | 0.83 | 0.6 | 0.69 | + |
| Pak1 | 1.62 | 1.18 | 1.29 | 2.99 | 2.1 | 2.17 | 4.43 | 3.26 | 3.98 | 2.75 | 3.83 | 4.09 | + |
| Tnfsf14 | 10.14 | 10.28 | 13.34 | 19.85 | 16.65 | 19.93 | 4.71 | 9.11 | 7.75 | 4.16 | 5.71 | 5.42 | + |
| D19Bwg1357e | 51.82 | 51.65 | 49.79 | 28.05 | 29.28 | 26.79 | 23.73 | 32.28 | 30.36 | 23.67 | 20.58 | 21.51 | + |
| Aste1 | 6.13 | 5.82 | 7.07 | 5.33 | 5.04 | 4.38 | 2.6 | 4.4 | 3.81 | 4.85 | 4.4 | 4.1 | + |
| G6pc3 | 25.36 | 25.85 | 26.79 | 17.72 | 20.04 | 19.12 | 19.01 | 20.46 | 23.07 | 18.82 | 21.75 | 18.62 | + |
| Senp5 | 6.46 | 6.37 | 6.57 | 4.03 | 3.91 | 4.11 | 3.44 | 4.21 | 3.99 | 3.14 | 3.53 | 2.7 | + |
| Apoo | 53.55 | 53.52 | 51.06 | 28.38 | 27.95 | 33.54 | 30.23 | 29.11 | 25.2 | 23.57 | 23.26 | 20.26 | + |
| Tfdp2 | 1.82 | 1.59 | 2.07 | 4.73 | 4.41 | 4.44 | 4.31 | 3.69 | 4.12 | 5.9 | 4.69 | 4.39 | + |
| Gnb5 | 15.6 | 13.49 | 13.53 | 34.74 | 27.02 | 33.1 | 32.9 | 29.59 | 31.44 | 26.4 | 30.86 | 23.7 | + |
| Snx4 | 46.62 | 45.48 | 47.14 | 39.57 | 39.36 | 38.25 | 40.19 | 35.09 | 36.91 | 35.83 | 35.91 | 39.11 | + |
| Pcyt2 | 76.32 | 76.58 | 72.84 | 67.26 | 63.14 | 68.15 | 61 | 54.16 | 57.86 | 52.96 | 58.04 | 59.81 | + |
| Il24 | 280.07 | 260.93 | 314.42 | 156.6 | 224.35 | 191 | 86.97 | 190.89 | 192.77 | 12.82 | 44.26 | 14.08 | + |
| Bap1 | 31.48 | 32.66 | 33.8 | 24.59 | 25.13 | 26.74 | 21.95 | 22.4 | 22.63 | 22.34 | 20.22 | 23.1 | + |
| B4galt5 | 5.49 | 8.36 | 7.39 | 1.7 | 2.36 | 2.7 | 1.52 | 2.18 | 1.77 | 1.33 | 1.36 | 1.44 | + |
| Mcm6 | 201.49 | 210.62 | 210.63 | 146.07 | 160.4 | 156.79 | 163.99 | 154.42 | 151.98 | 142.69 | 142.94 | 155.48 | + |
| Tubg2 | 2.75 | 2.84 | 2.45 | 4.31 | 4.81 | 3.9 | 8.12 | 5.51 | 7.97 | 5.02 | 5.08 | 7.23 | + |
| Foxred1 | 18.54 | 19.09 | 16.51 | 12.76 | 11.73 | 12.75 | 13.74 | 12.63 | 13.11 | 12.1 | 13.75 | 12.14 | + |
| Gzmc | 211.9 | 257.1 | 85.9 | 30.86 | 17.38 | 15.16 | 162.49 | 71.51 | 108.68 | 28.91 | 34.78 | 36.04 | + |
| Ddb2 | 7.78 | 8.1 | 7.91 | 12.57 | 11.62 | 13.09 | 20.25 | 15.24 | 19.57 | 17.43 | 18.48 | 20.07 | + |
| Tuba1a | 421.31 | 365.58 | 389.69 | 165.76 | 174.43 | 181.26 | 162.52 | 171.14 | 167.54 | 106.68 | 118.57 | 114.83 | + |
| Ogt | 22.75 | 22.28 | 25.03 | 39.27 | 36.96 | 38.9 | 45.14 | 38.07 | 40.94 | 52.37 | 52.35 | 49.84 | + |
| Prcp | 14.15 | 13.68 | 12.15 | 17.59 | 19.59 | 18.33 | 22.74 | 21.72 | 21.31 | 22.15 | 21.82 | 21.49 | + |
| Tk1 | 71.54 | 74.92 | 74.32 | 25.56 | 26.97 | 31.29 | 27.27 | 25.72 | 25.38 | 13.93 | 15.94 | 16.35 | + |
| Idi2 | 27.99 | 13.93 | 21.74 | 5.33 | 7.12 | 5.43 | 1.67 | 2.62 | 1.44 | 2.35 | 2.31 | 1.64 | + |
| Obfc1 | 21.4 | 18.65 | 19.56 | 37.99 | 33.2 | 32.7 | 32.52 | 29.52 | 29.58 | 31.32 | 31.47 | 29.4 | + |
| H2-DMb2 | 8.22 | 6.52 | 5.38 | 13.57 | 11.05 | 11.26 | 18.68 | 12.96 | 15.19 | 13.87 | 11.6 | 12.36 | + |
| Paip2 | 118.07 | 104.12 | 110.04 | 174.56 | 174.29 | 182.76 | 203.7 | 199.29 | 194.25 | 214.24 | 234.3 | 216.27 | + |
| Dgcr2 | 18.27 | 19.11 | 17.12 | 15.58 | 13.34 | 16.21 | 11.63 | 12.83 | 11.94 | 12.78 | 12.71 | 12.68 | + |
| Nip7 | 56.15 | 59.77 | 59.55 | 25.45 | 26.93 | 26.65 | 20.57 | 29.78 | 27.86 | 19.06 | 19.29 | 19.61 | + |
| Hnrnpa3 | 23.02 | 27.2 | 27.03 | 20.44 | 22.27 | 18.38 | 16.64 | 16.04 | 16.71 | 20.2 | 18.24 | 17.12 | + |
| Arfip2 | 14.67 | 14.73 | 15.77 | 11.33 | 12.95 | 12.14 | 12.42 | 12.53 | 12.6 | 10.7 | 12.33 | 11.35 | + |
| Tcp1 | 538.46 | 530.96 | 538.5 | 371 | 388.28 | 392.71 | 440.16 | 449.06 | 436.14 | 360.87 | 425.29 | 352.57 | + |
| Sf1 | 47.58 | 52.62 | 49.99 | 35.61 | 33.11 | 35.27 | 30.86 | 32.05 | 30.62 | 36.79 | 32.38 | 33.27 | + |
| Rars | 110.05 | 111.72 | 112.74 | 53.57 | 59.23 | 57.86 | 48.12 | 68.35 | 65.23 | 42.2 | 39.24 | 40.02 | + |
| Pno1 | 79.53 | 79.58 | 81.11 | 32.11 | 38.06 | 36.95 | 31.43 | 45.66 | 41.88 | 30.05 | 25.94 | 26.15 | + |
| Hacl1 | 7.65 | 4.51 | 4.89 | 9.61 | 10.04 | 10.81 | 9.97 | 8.86 | 9.28 | 8.19 | 8.47 | 8.87 | + |
| Prps1 | 73.33 | 78.5 | 77.57 | 45.33 | 45.08 | 47.09 | 28.1 | 42.54 | 35.87 | 30.52 | 32.91 | 28.84 | + |
| Gm17762 | 0.16 | 0.22 | 0.21 | 0.68 | 0.57 | 0.75 | 0.79 | 0.9 | 0.67 | 0.67 | 0.79 | 0.48 | + |
| Fam168a | 1.93 | 2.37 | 1.96 | 4.25 | 3.83 | 3.54 | 4.43 | 3.22 | 3.37 | 3.76 | 4.47 | 5.1 | + |
| Akna | 34.46 | 34.97 | 37.17 | 36.73 | 33.16 | 32.79 | 39.68 | 33.88 | 38.03 | 48.99 | 43.5 | 53.91 | + |
| Morf4l2 | 130.95 | 133.8 | 128.93 | 98.06 | 102.8 | 99.31 | 84.68 | 95.85 | 100.21 | 89.37 | 93.77 | 82.47 | + |
| Pdap1 | 21.59 | 20.94 | 21.77 | 12.42 | 13.83 | 13.34 | 11.68 | 15.54 | 14.61 | 10.15 | 11.56 | 10.2 | + |
| Thap3 | 10.86 | 9.71 | 11.65 | 23.72 | 19.03 | 19.27 | 27.53 | 19.23 | 20.72 | 24.04 | 27.4 | 27.24 | + |
| Pcgf1 | 11.04 | 9.11 | 9.09 | 14.03 | 13.64 | 13.28 | 19.2 | 15.39 | 19.26 | 17.28 | 18.76 | 17.28 | + |
| BC023829 | 10.82 | 9.8 | 9.54 | 8.39 | 6.43 | 7.53 | 6.41 | 5.43 | 6.56 | 6.76 | 6.82 | 6.87 | + |
| Asnsd1 | 61.92 | 61.03 | 59.15 | 71.49 | 74.35 | 78.29 | 73.02 | 76.04 | 73.88 | 68.61 | 82.74 | 72.29 | + |
| Man2a2 | 3.66 | 3.8 | 3.68 | 9.06 | 7.54 | 8.51 | 3.84 | 4.22 | 4.78 | 4.99 | 5.23 | 5.15 | + |
| Zfp358 | 0.9 | 0.61 | 0.89 | 1.77 | 1.62 | 2.14 | 2.12 | 1.79 | 1.58 | 1.75 | 1.45 | 2.54 | + |
| Cdc37 | 126.57 | 131.18 | 127.97 | 99.41 | 99.25 | 107.19 | 102.43 | 108.52 | 103.63 | 98.66 | 96.24 | 103.22 | + |
| 9430016H08Rik | 28.09 | 28.11 | 29.18 | 18.36 | 19.13 | 20.94 | 18.82 | 21.72 | 20.25 | 15.54 | 17.21 | 14.14 | + |
| Exosc2 | 55.8 | 56.27 | 57.19 | 29.51 | 28.5 | 31.28 | 20.63 | 31.01 | 26.94 | 18.57 | 18.83 | 15.11 | + |
| Tnfrsf23 | 1.11 | 0.98 | 0.97 | 1.9 | 1.82 | 2.11 | 1.7 | 1.57 | 1.97 | 1.78 | 2.03 | 1.67 | + |
| Armc8 | 9.4 | 9.98 | 8.47 | 6.34 | 6.54 | 6.63 | 6.16 | 6.89 | 6.72 | 6.8 | 6.23 | 5.74 | + |
| Mapk14 | 6.66 | 5.77 | 5.13 | 8.01 | 7.17 | 7.02 | 4.95 | 5.75 | 4.9 | 12.87 | 11.07 | 11.21 | + |
| Sucla2 | 62.42 | 64.15 | 59.65 | 37.55 | 42.05 | 40.69 | 37.98 | 40.38 | 39.77 | 33.71 | 33.24 | 35.19 | + |
| Gfm2 | 20.84 | 17.08 | 17.85 | 12.55 | 13.46 | 13.35 | 11.95 | 14.71 | 13.84 | 11.6 | 11.91 | 10.25 | + |
| Irf2 | 6.48 | 6.93 | 7.65 | 13.4 | 12.44 | 11.53 | 20.15 | 14.8 | 15.23 | 19.71 | 21.17 | 20.16 | + |
| Arfgap3 | 9.01 | 8.25 | 7.99 | 19.64 | 16.52 | 20.12 | 24.47 | 18.54 | 18.59 | 26.99 | 30.33 | 24.83 | + |
| Cfdp1 | 115.16 | 109.96 | 112.31 | 79.38 | 89.21 | 89.23 | 81.29 | 93.78 | 82.31 | 79.19 | 80.87 | 68.84 | + |
| Zfp7 | 8.93 | 9.91 | 9.36 | 10.44 | 9.25 | 10.4 | 9.62 | 9.61 | 9.59 | 11.44 | 12.25 | 12.24 | + |
| Rhoq | 3 | 2.78 | 3.2 | 6.52 | 4.61 | 6.01 | 5.43 | 4.5 | 5.51 | 6.22 | 5.99 | 5.45 | + |
| Eif4g1 | 28.34 | 32.09 | 34.72 | 17.84 | 19.26 | 20.98 | 14.39 | 21.34 | 17.61 | 15.45 | 16.26 | 14.66 | + |
| Fbxw4 | 1.29 | 1.14 | 1.47 | 2.53 | 3.53 | 3.04 | 3.78 | 2.08 | 2.95 | 2.15 | 2.91 | 2.7 | + |
| C920025E04Rik | 43.46 | 39.47 | 39.65 | 74.86 | 82.06 | 83.61 | 94.04 | 78.71 | 83.32 | 84.35 | 95.17 | 91.25 | + |
| Avpi1 | 41.53 | 43.34 | 50.46 | 37.62 | 51.11 | 46.78 | 53.7 | 53.76 | 52.94 | 28.31 | 30.38 | 30.07 | + |
| Rpl41 | 3672.53 | 3798.7 | 3751.62 | 5017.53 | 5018.31 | 5334.69 | 5404.08 | 4912.32 | 5034.58 | 5574.46 | 5957.59 | 5486.11 | + |
| Gja1 | 0.97 | 0.88 | 1.17 | 2.12 | 4.46 | 3.74 | 2.13 | 3.31 | 1.92 | 2.1 | 3.12 | 2.32 | + |
| 3110082I17Rik | 26.61 | 25.09 | 27.31 | 6.45 | 8.59 | 8.08 | 4.56 | 9.97 | 6.03 | 4.23 | 3.08 | 3.91 | + |
| Nagpa | 15.56 | 16.11 | 17.37 | 9.08 | 9.49 | 9.79 | 8.53 | 8.58 | 9.95 | 12.33 | 9.49 | 11.6 | + |
| Mrpl17 | 19.98 | 20.27 | 20.73 | 11.78 | 12.93 | 13 | 10.76 | 13.21 | 12.92 | 8.17 | 8.73 | 9.03 | + |
| H2-DMa | 18.66 | 20.88 | 21.39 | 44.03 | 39.08 | 39.81 | 44.36 | 40.83 | 40.64 | 39.96 | 47.48 | 42.97 | + |
| Bnip1 | 25.62 | 24.4 | 22.07 | 13.75 | 15.48 | 19.04 | 17.9 | 21.17 | 18.84 | 14.24 | 15.96 | 14.93 | + |
| Chordc1 | 86.63 | 89.72 | 83 | 53 | 55.54 | 54.46 | 56.68 | 67.32 | 64.99 | 54.77 | 60.43 | 57.25 | + |
| Golt1b | 64.64 | 65.1 | 64.01 | 50.75 | 52.04 | 50.41 | 48.68 | 54.23 | 54.2 | 49.2 | 53.58 | 50 | + |
| Osbpl3 | 9.98 | 8.81 | 7.71 | 7.88 | 8.6 | 8.15 | 7.4 | 8.18 | 7.95 | 3.6 | 4.96 | 3.84 | + |
| Rae1 | 95.24 | 98.08 | 103.05 | 67.04 | 73.42 | 70.62 | 62.32 | 77.92 | 75.76 | 59.77 | 67.5 | 64.23 | + |
| E130102H24Rik | 0.53 | 0.47 | 0.58 | 2.53 | 2.09 | 1.62 | 2.07 | 1.74 | 1.94 | 3.52 | 2.8 | 2.3 | + |
| Mocs1 | 3.62 | 3.42 | 2.93 | 5.99 | 6.03 | 6.58 | 4.64 | 5.57 | 5.68 | 5.25 | 6.17 | 4.5 | + |
| Diexf | 3.05 | 3.09 | 3.64 | 1.29 | 1.75 | 2.35 | 1.04 | 1.78 | 1.7 | 1.9 | 1.74 | 1.49 | + |
| Il18r1 | 34.18 | 29.69 | 30.55 | 54.78 | 49.02 | 52.17 | 45.28 | 46.37 | 43.84 | 43 | 50.32 | 47.27 | + |
| Rnft1 | 14.27 | 13.01 | 14.22 | 10.37 | 9.81 | 8.41 | 9.53 | 8.32 | 8.99 | 8.98 | 10.08 | 8.5 | + |
| Fbxo46 | 5.18 | 5.66 | 5.26 | 3.15 | 4 | 4.46 | 4.67 | 3.53 | 3.19 | 3.11 | 4.06 | 4.13 | + |

| | | | | | | | | | | | | | |
|---|---|---|---|---|---|---|---|---|---|---|---|---|---|
| Arxes2 | 5.59 | 6.4 | 6.74 | 3 | 2.12 | 1.9 | 1.53 | 2.75 | 1.71 | 1.71 | 1.6 | 1.72 | + |
| Rpap1 | 12.65 | 12.41 | 12.52 | 8.6 | 9.19 | 9.57 | 7.68 | 9.49 | 9.85 | 7.42 | 8.31 | 8.65 | + |
| Polr3h | 24.87 | 26.43 | 24.73 | 11.35 | 10.99 | 8.78 | 8.99 | 12.41 | 10.53 | 6.83 | 6.44 | 6.75 | + |
| Gmps | 72.43 | 71.34 | 73.83 | 42.12 | 44.58 | 47.75 | 37.2 | 45.04 | 42 | 39.76 | 41.02 | 39.67 | + |
| Cdc34 | 41.54 | 40.13 | 43.99 | 14.93 | 18.54 | 22.47 | 19.15 | 28.71 | 22.08 | 15 | 14.36 | 15.57 | + |
| Nol8 | 10.13 | 11.57 | 11.2 | 6.92 | 7.32 | 7.72 | 6.73 | 7.78 | 8.08 | 7.38 | 6.82 | 6.72 | + |
| Zfp808 | 10.98 | 10.18 | 11.36 | 6.16 | 8.62 | 9.07 | 5.47 | 5.01 | 5.35 | 5.6 | 5.69 | 5.31 | + |
| Ddit3 | 28.02 | 25.62 | 29.94 | 89.88 | 98.15 | 98.52 | 114.27 | 135.79 | 132.19 | 91.03 | 112.17 | 98.48 | + |
| Dnajc1 | 10.63 | 10.29 | 11.67 | 14.59 | 15.72 | 16.61 | 22.49 | 22.41 | 20.07 | 22.01 | 20.54 | 22.22 | + |
| Glod4 | 42.38 | 46.8 | 42.56 | 53.52 | 55.33 | 57.88 | 61.49 | 61.87 | 60.18 | 56.27 | 59.7 | 57.06 | + |
| Ltv1 | 72.26 | 62.24 | 75.24 | 33.09 | 35.27 | 34.74 | 27.68 | 39.57 | 37.54 | 24.37 | 27.64 | 24.35 | + |
| Cdt1 | 32.38 | 38.43 | 34.37 | 23.1 | 19.7 | 17.18 | 16.26 | 14.73 | 14.23 | 17.02 | 13.49 | 17.14 | + |
| Zcchc11 | 2.54 | 2.1 | 1.95 | 3.85 | 3.75 | 3.9 | 3.97 | 3.2 | 3.6 | 4.28 | 4.17 | 4.74 | + |
| Naa10 | 137.84 | 140.6 | 142.26 | 106.1 | 107.74 | 106.19 | 99.11 | 109.4 | 102 | 92.22 | 89.18 | 92.99 | + |
| Ptcra | 2.44 | 6.56 | 3.84 | 0.8 | 2.53 | 1.25 | 1.58 | 2.29 | 1.18 | 1.07 | 2.97 | 1.63 | + |
| Rps27l | 672.08 | 673.96 | 646.17 | 333.84 | 371.73 | 386.41 | 359.28 | 413.73 | 413.6 | 278.27 | 304.67 | 262.27 | + |
| Ezr | 152.34 | 158.95 | 156.53 | 89.45 | 90.5 | 89.55 | 105.59 | 112.34 | 114.47 | 80.23 | 81.84 | 75.28 | + |
| Socs2 | 52.53 | 58.48 | 53.74 | 76.95 | 81.89 | 73.68 | 25.85 | 31.51 | 33.43 | 17.96 | 18.27 | 17.72 | + |
| Ifngr2 | 4.52 | 6.55 | 6.26 | 16.64 | 13.71 | 14.27 | 3.42 | 3.48 | 4.15 | 2.25 | 2.23 | 2.47 | + |
| Tm9sf3 | 9.7 | 9.14 | 9.24 | 7.48 | 6.81 | 7.19 | 6.9 | 7.91 | 7.18 | 7.92 | 7.48 | 6.82 | + |
| Scand1 | 1.65 | 2.91 | 3.77 | 4.91 | 6 | 4.62 | 7.26 | 6.79 | 7.34 | 7.45 | 5.68 | 5.16 | + |
| Rps13 | 1759.16 | 1854.45 | 1758.9 | 2427.66 | 2366.61 | 2431.4 | 2350.24 | 2249.52 | 2205.17 | 2532.39 | 2700.87 | 2605.27 | + |
| Ikbip | 6.64 | 5.9 | 5.61 | 8.74 | 8.09 | 8.58 | 9.29 | 9.56 | 9.88 | 9.86 | 9.18 | 10.12 | + |
| Tesc | 2.55 | 5 | 5.28 | 12.01 | 9.21 | 10.36 | 24.77 | 18.64 | 21.19 | 31.09 | 24.53 | 19.03 | + |
| Dohh | 88.13 | 93.08 | 89.71 | 42.24 | 38.17 | 43.31 | 46.12 | 51.86 | 48.91 | 41.4 | 42.53 | 41.56 | + |
| Las1l | 43.29 | 42.2 | 44.16 | 27.31 | 27.19 | 25.11 | 21.15 | 26.23 | 24.75 | 20.87 | 20.83 | 20.21 | + |
| Smc2 | 17.48 | 13.96 | 15.92 | 9.16 | 11.61 | 10.1 | 6.67 | 7.52 | 7.33 | 6.42 | 6.74 | 5.12 | + |
| Eif5b | 14.63 | 15.94 | 14.89 | 11.59 | 11.29 | 11.14 | 9.85 | 11.84 | 11.23 | 10.02 | 10.1 | 10.85 | + |
| Gfpt1 | 7.72 | 7.34 | 7.75 | 4.25 | 4.39 | 4.45 | 5 | 5.15 | 5.49 | 3.54 | 4.22 | 3.1 | + |
| Ndufs1 | 56.92 | 60.29 | 54.36 | 35.89 | 35.53 | 35.22 | 29.99 | 37.18 | 33.32 | 29.52 | 28 | 27.27 | + |
| Med8 | 32.18 | 30.29 | 34.15 | 24.04 | 23.85 | 24.01 | 20.55 | 24.47 | 23.56 | 18.31 | 21.41 | 19.36 | + |
| Tsc22d3 | 11.61 | 9.03 | 10.7 | 242.45 | 194.36 | 235.05 | 60.49 | 50.52 | 48.83 | 389.68 | 420.23 | 407.41 | + |
| Kdm8 | 5.11 | 5.14 | 5.38 | 2.41 | 3.45 | 2.4 | 3.01 | 3.1 | 3.09 | 2.25 | 1.89 | 2.05 | + |
| Tmem168 | 14.07 | 15.79 | 14.66 | 8.45 | 8.88 | 6.88 | 9.07 | 9.92 | 8.95 | 8.93 | 8.2 | 7.93 | + |
| Cenpp | 15.44 | 16 | 14.77 | 10.73 | 12.91 | 12.84 | 10.3 | 8.93 | 8.09 | 10.96 | 11.35 | 10.76 | + |
| Fam161a | 0.35 | 0.21 | 0.36 | 1.37 | 1.45 | 1.09 | 1.81 | 1 | 1.22 | 1.27 | 1.24 | 1.52 | + |
| Hsp90aa1 | 438.43 | 423.7 | 437.89 | 134.84 | 165.02 | 153.88 | 124.07 | 192.35 | 164.8 | 96.36 | 96.92 | 90.06 | + |
| Pnp | 27.73 | 25.24 | 23.82 | 14.81 | 14.89 | 15.34 | 13.69 | 14.7 | 14.27 | 12.77 | 12.54 | 10.71 | + |
| Polr3gl | 8.84 | 9.59 | 8.86 | 23.61 | 21.1 | 21.14 | 20.17 | 15.7 | 18.83 | 25.93 | 25.03 | 25.71 | + |
| Pmvk | 59.52 | 60.95 | 59.46 | 54.4 | 52.52 | 55.59 | 80.36 | 69.93 | 69.34 | 71.63 | 68.24 | 78.01 | + |
| Zfp948 | 14.2 | 13.11 | 15.22 | 14.77 | 15.19 | 17.26 | 15.01 | 14.12 | 12.69 | 18.38 | 21.54 | 15.54 | + |
| Thop1 | 54.73 | 46.35 | 52.73 | 15.7 | 18.64 | 16.72 | 14.07 | 19.06 | 17.05 | 9.29 | 8.87 | 10.77 | + |
| Slc12a2 | 1.65 | 1.73 | 1.52 | 1.11 | 1.14 | 1.27 | 0.83 | 0.8 | 0.74 | 0.95 | 0.72 | 0.96 | + |
| Copz2 | 0.31 | 0.44 | 0.58 | 1.56 | 1.88 | 1.51 | 3.15 | 1.71 | 3.23 | 1.79 | 2.79 | 1.8 | + |
| Ddx19a | 7.95 | 8.45 | 8.19 | 6.1 | 6.94 | 5.68 | 5.64 | 6.12 | 5.47 | 5.03 | 5.15 | 4.91 | + |
| Vps26a | 24.57 | 23.81 | 23.19 | 19.62 | 18.93 | 18.37 | 16.48 | 20.6 | 18.37 | 17.55 | 16.69 | 17.62 | + |
| Elavl1 | 28.89 | 36.96 | 38.82 | 22.75 | 24.83 | 24.96 | 20.44 | 23.29 | 22.07 | 19.62 | 20.9 | 18.84 | + |
| Arcn1 | 91.73 | 92.9 | 97.45 | 74.67 | 83.95 | 80.92 | 70.4 | 82.17 | 74.35 | 68.39 | 76.69 | 71.72 | + |
| Mrpl18 | 149.36 | 159.45 | 155.55 | 86 | 83.33 | 89.31 | 101.91 | 86.9 | 90.51 | 80.83 | 80.04 | 80.13 | + |
| Med12 | 6.35 | 5.9 | 6.23 | 10.45 | 10.21 | 10.94 | 10.73 | 9.75 | 9.76 | 11.41 | 11.69 | 10.57 | + |
| Cyb5r1 | 32.07 | 30.14 | 29.23 | 36.72 | 42.33 | 43.68 | 29.93 | 45.73 | 41.36 | 18.47 | 19.82 | 20.44 | + |
| Iqcb1 | 8.24 | 7.21 | 6.92 | 13.91 | 11.24 | 13.17 | 9.73 | 11.75 | 10.56 | 12.64 | 11.63 | 11.14 | + |
| Insr | 0.86 | 0.97 | 0.95 | 1.97 | 1.64 | 1.75 | 1.67 | 1.64 | 1.76 | 2.34 | 1.85 | 2.3 | + |
| Slc16a6 | 10.45 | 9.88 | 8.65 | 4.92 | 5.02 | 4.89 | 5.5 | 6.54 | 4.97 | 4.64 | 4.64 | 4.88 | + |
| Ddx24 | 82.35 | 86.05 | 84.76 | 63.62 | 62.7 | 59.31 | 62.3 | 58.7 | 59.68 | 75.21 | 66.59 | 71.77 | + |
| Gpr108 | 26.33 | 20.59 | 21.76 | 33.59 | 35.06 | 33.71 | 33.64 | 30.69 | 27.73 | 32.26 | 31.58 | 34.06 | + |
| Ppp1r10 | 19.64 | 17.63 | 18.96 | 13.66 | 14.56 | 14.36 | 9.93 | 12.54 | 9.59 | 12.79 | 10.14 | 10.11 | + |
| Klf10 | 7.76 | 8.73 | 8.83 | 28.23 | 26.5 | 28.16 | 28.33 | 20.08 | 20.95 | 42.79 | 43.65 | 42.84 | + |
| Mrps2 | 30.72 | 28.82 | 32.74 | 15.78 | 18.2 | 13.4 | 7.57 | 14.07 | 14.34 | 10.43 | 9.13 | 9.76 | + |
| Prpf4 | 11.84 | 12.23 | 12.74 | 8.86 | 7.53 | 8.61 | 7.54 | 8.1 | 7.8 | 7.35 | 6.68 | 7.14 | + |
| Ubl3 | 8.08 | 7.6 | 7.86 | 17.4 | 16.85 | 16.98 | 11.57 | 14.65 | 14.01 | 14.65 | 14.2 | 16.65 | + |
| Degs1 | 95.93 | 101.95 | 92.84 | 62.42 | 57.72 | 58.25 | 68.33 | 69.87 | 68.06 | 61.29 | 54.34 | 64.27 | + |
| Zfand2b | 18.83 | 16.19 | 15.81 | 21.63 | 23.02 | 25.06 | 31.4 | 26.69 | 27 | 27.61 | 28.21 | 30.99 | + |
| Nbeal1 | 0.8 | 0.8 | 0.81 | 2.04 | 2.03 | 2.02 | 1.9 | 1.98 | 1.85 | 2 | 2.4 | 1.96 | + |
| Ube2k | 22.1 | 24.32 | 23.7 | 15.62 | 16.43 | 17.08 | 14.06 | 17.8 | 15.83 | 14.13 | 15.14 | 13.97 | + |
| 1810043H04Rik | 51.66 | 56.19 | 52.6 | 31.86 | 31.09 | 40.66 | 33.73 | 45.06 | 36.93 | 27.56 | 30.69 | 34.42 | + |
| Rnpep | 53.52 | 55 | 56.43 | 22.23 | 25.33 | 24.02 | 32.45 | 34.89 | 30.7 | 32.81 | 34.74 | 32.88 | + |
| Pin4 | 59.95 | 57.58 | 53.26 | 36.99 | 39.95 | 39.2 | 39.2 | 44.51 | 42.36 | 40.95 | 36.98 | 42.6 | + |
| Ddx39 | 248.01 | 269.34 | 264.14 | 66.18 | 82.93 | 72.15 | 53.73 | 75.48 | 59.97 | 42.01 | 36.35 | 39.92 | + |
| Sdhd | 146.72 | 146.72 | 142.39 | 95.53 | 103.18 | 99.77 | 111.03 | 97.53 | 99.19 | 105.9 | 124.75 | 107.4 | + |
| Mcmbp | 14.21 | 15.33 | 14.13 | 12.24 | 11.35 | 11.64 | 9.67 | 10.03 | 9.83 | 12.69 | 11.31 | 11.26 | + |
| Xlr4b | 36.99 | 38.67 | 39.71 | 22.38 | 24.5 | 27.43 | 23.83 | 27.64 | 27.33 | 20.84 | 21.86 | 21.55 | + |
| Ing3 | 5.63 | 6.57 | 5.87 | 6.1 | 6.38 | 5.81 | 5.88 | 6.05 | 6.19 | 8.01 | 8.09 | 7.75 | + |
| Cdca5 | 30.7 | 25.37 | 25.35 | 18.34 | 21.42 | 20.84 | 14.38 | 12.2 | 12.53 | 14.47 | 12.64 | 10.59 | + |
| Hk2 | 28.74 | 26.25 | 31.87 | 9.13 | 12.17 | 11.5 | 11.68 | 16.7 | 14.95 | 7.31 | 7.4 | 7.64 | + |
| Rpl7l1 | 82.53 | 84.42 | 84.68 | 35.11 | 43 | 42.68 | 27.71 | 34.83 | 33.68 | 23.89 | 22.1 | 27.55 | + |
| Stk38 | 25.13 | 23.11 | 22.66 | 18.01 | 18.24 | 18.97 | 21.65 | 20.67 | 19.01 | 17.66 | 16.63 | 18.07 | + |
| Nrip1 | 0.61 | 0.57 | 0.58 | 1.28 | 0.79 | 0.99 | 1.61 | 0.99 | 0.9 | 1.02 | 1.25 | 1.08 | + |
| Ccdc132 | 9.21 | 7.95 | 8.73 | 5.17 | 5.8 | 4.64 | 6.09 | 4.86 | 4.98 | 5.04 | 4.94 | 5.41 | + |
| Rbbp4 | 47.16 | 47.63 | 44.21 | 39.12 | 40.16 | 38.73 | 28.58 | 30.95 | 29.48 | 28.8 | 32.19 | 28.6 | + |
| Polr2f | 110.46 | 108.04 | 111.26 | 46.06 | 48.5 | 47.46 | 44.98 | 54.56 | 53.33 | 37.75 | 36.39 | 41.25 | + |
| 4932415G12Rik | 0.58 | 0.49 | 0.44 | 1.35 | 1.39 | 1.36 | 0.65 | 0.95 | 0.61 | 0.97 | 1.28 | 1.39 | + |
| Nicn1 | 6.27 | 5.67 | 5.8 | 16.74 | 14.66 | 17.04 | 18.96 | 14.46 | 14.21 | 14.99 | 20.63 | 18.39 | + |
| Slc35a4 | 65.51 | 63.38 | 62.51 | 34.57 | 32.2 | 33.87 | 24.87 | 29.73 | 32.06 | 26.25 | 22.43 | 26.37 | + |
| Ezh1 | 9.24 | 7.78 | 7.93 | 18.55 | 16.35 | 16.36 | 33.23 | 23.49 | 23.24 | 31.97 | 35.77 | 35.12 | + |
| Ints3 | 16.63 | 16.75 | 15.59 | 11.67 | 13.44 | 11.86 | 11.14 | 11.76 | 11.16 | 10.41 | 13.64 | 12.52 | + |
| Haus4 | 41.77 | 40.37 | 41.82 | 29.39 | 31.7 | 32.3 | 30.38 | 27.68 | 28.57 | 27.75 | 28.81 | 24.92 | + |
| Map1lc3a | 2.13 | 1.73 | 2.38 | 7.62 | 6.09 | 5.19 | 6.71 | 7.42 | 6.78 | 9.74 | 8.61 | 9.16 | + |
| Nup93 | 51.47 | 47.5 | 49.78 | 19.65 | 23.54 | 20.01 | 19.04 | 25.14 | 23.16 | 16.58 | 15.15 | 15.18 | + |
| Rps15a-ps4 | 95.4 | 102.57 | 94.39 | 163.55 | 163.39 | 180.03 | 188.56 | 157.45 | 172.65 | 203.15 | 213.77 | 200.66 | + |
| Adam15 | 10.82 | 9.71 | 10.44 | 16.59 | 16.03 | 16.73 | 16.37 | 16.8 | 16.85 | 15.14 | 15.24 | 16.19 | + |
| Ascc1 | 25.64 | 25.03 | 26.34 | 45.54 | 41.47 | 43.93 | 42.07 | 43.78 | 40.29 | 45.31 | 47.61 | 45.89 | + |
| Fam210b | 2.8 | 2.47 | 2.55 | 5.21 | 4.03 | 4.76 | 6.22 | 6.01 | 4.75 | 5.46 | 5.47 | 7.79 | + |
| Mpi | 18.85 | 15.49 | 16.46 | 12.77 | 12.08 | 10.89 | 12.95 | 13.57 | 11.27 | 9.77 | 10.63 | 12.6 | + |
| Zdhhc13 | 21.44 | 19.19 | 19.32 | 15.94 | 15.22 | 15.98 | 11.86 | 13.42 | 11.46 | 11.84 | 10.63 | 10.51 | + |
| Dgcr6 | 7.83 | 10.47 | 10.28 | 16.67 | 16.22 | 16.52 | 16.39 | 14.99 | 13.69 | 14.32 | 14.6 | 12.24 | + |
| Nucks1 | 13.91 | 13.21 | 13.92 | 8.72 | 9.86 | 7.76 | 6.02 | 6.85 | 6.9 | 6.27 | 6.9 | 6.36 | + |
| Arid5a | 33.22 | 34.84 | 33.37 | 77.65 | 68.37 | 71.01 | 47.56 | 44.39 | 47.38 | 51.8 | 42.77 | 52.81 | + |
| Rasa4 | 4.4 | 4.18 | 3.75 | 10.82 | 11.22 | 10.06 | 14.28 | 8.52 | 9.35 | 11.46 | 12.32 | 13 | + |
| Cbx1 | 11.87 | 12.58 | 11.8 | 9.67 | 9.16 | 8.36 | 7.26 | 6.92 | 7.87 | 6.69 | 6.34 | 6.6 | + |
| Memo1 | 84.34 | 84.52 | 84.29 | 54.11 | 61.63 | 62.07 | 55.28 | 61.16 | 63.31 | 50.82 | 54.81 | 55.9 | + |
| Tbc1d15 | 16.69 | 16.43 | 16.84 | 21.93 | 20.71 | 21.69 | 21.76 | 20.3 | 19.86 | 26.34 | 29.85 | 26.16 | + |
| Mafk | 17.3 | 19.59 | 17.74 | 9.37 | 12.44 | 13.12 | 8.4 | 10.04 | 11.81 | 11.04 | 11.34 | 11.51 | + |
| Ap1m1 | 78.56 | 68.79 | 74.26 | 45.46 | 47.77 | 45.23 | 44.48 | 44.88 | 47.21 | 39.03 | 38.19 | 43.88 | + |
| Eml5 | 0.8 | 0.62 | 0.63 | 1.36 | 1.68 | 1.31 | 1.5 | 1.37 | 1.37 | 1.59 | 1.61 | 1.58 | + |
| Snrnp200 | 22.06 | 20.33 | 22.48 | 17.69 | 17.91 | 18.13 | 17.2 | 17.67 | 18.7 | 16.54 | 18.05 | 17.33 | + |
| Rps15a | 139.21 | 140.72 | 143.23 | 241.81 | 234.06 | 246.32 | 264.66 | 238.58 | 248.86 | 300.87 | 309.52 | 308.16 | + |
| Kansl1l | 0.46 | 0.38 | 0.43 | 0.8 | 0.76 | 0.59 | 1.07 | 0.89 | 0.74 | 0.92 | 0.74 | 1.04 | + |
| Pfn1 | 1219.1 | 1291.98 | 1228.17 | 665.44 | 715.69 | 723.92 | 685.76 | 781.32 | 780.63 | 549.63 | 457.53 | 563.14 | + |
| Dapk2 | 1.16 | 0.95 | 0.86 | 10.18 | 3.88 | 3.93 | 18.35 | 8.53 | 11.53 | 17.79 | 13.31 | 19.52 | + |
| Plk1 | 73.56 | 53.83 | 51.85 | 35.28 | 44.13 | 41.65 | 21.39 | 20.2 | 20.26 | 20.49 | 21.66 | 19.76 | + |
| Ipo11 | 21.01 | 20.27 | 20.77 | 10.51 | 11.74 | 9.05 | 8.3 | 9.98 | 8.81 | 6.52 | 7.65 | 7.2 | + |
| Cct5 | 578.6 | 588.6 | 601.74 | 353.23 | 370.68 | 383.93 | 348.88 | 425.43 | 406.19 | 311.99 | 325.79 | 307.75 | + |
| Tmco6 | 3.52 | 5.45 | 6.46 | 7.23 | 7.77 | 6.64 | 10.26 | 10.91 | 9.34 | 8.92 | 7.33 | 8 | + |
| Hip1r | 13.14 | 12.64 | 10.73 | 17.38 | 15.28 | 15.52 | 19.97 | 13.4 | 14.62 | 29.66 | 30.93 | 31.11 | + |
| Dad1 | 200.42 | 204.01 | 211.69 | 146.79 | 151.04 | 162.56 | 165.02 | 151.32 | 150.69 | 153.3 | 163.45 | 175.72 | + |
| Sit1 | 18.25 | 16.72 | 13.15 | 25.1 | 21.67 | 21.24 | 32.34 | 25.87 | 30.34 | 26.34 | 24.43 | 24.86 | + |
| Amigo1 | 0.41 | 0.2 | 0.17 | 0.46 | 0.44 | 0.49 | 1.67 | 1.12 | 1.39 | 1.02 | 0.91 | 1.23 | + |
| Tsnax | 77.33 | 72.03 | 74.41 | 51.16 | 50.02 | 52.09 | 49.21 | 49.6 | 53.29 | 39.62 | 40.62 | 43.08 | + |
| Mllt1 | 5.71 | 5.46 | 5.44 | 3.77 | 3.48 | 3.39 | 3.2 | 3.08 | 2.9 | 3.19 | 3.4 | 2.95 | + |
| Mrpl47 | 31.16 | 33.57 | 34.76 | 22.35 | 20.97 | 21.71 | 20.1 | 22.36 | 23.75 | 14.5 | 22.76 | 16.92 | + |
| Syncrip | 62.67 | 65.52 | 67.32 | 25.47 | 31.17 | 27.64 | 19.67 | 28.16 | 24.33 | 19.97 | 20.54 | 18.72 | + |
| Sox5 | 0.62 | 0.36 | 0.69 | 1.61 | 1.58 | 1.47 | 0.71 | 0.81 | 0.95 | 1.27 | 1.11 | 1.2 | + |
| Abcf2 | 93.61 | 94.64 | 92.96 | 36.74 | 38.34 | 37.48 | 28.69 | 37.46 | 35.22 | 25.33 | 24.58 | 28.36 | + |
| Optn | 1.63 | 1.64 | 1.08 | 3.48 | 3.05 | 3.38 | 4.38 | 4.38 | 4.42 | 3.75 | 2.15 | 3.94 | + |
| Hspa8 | 3501.84 | 3316.47 | 3490.05 | 2035.01 | 2152.47 | 2136.58 | 2193.22 | 2309.12 | 2336.71 | 2051.14 | 2062.58 | 2078.03 | + |
| Tubgcp3 | 17.02 | 18.13 | 16.48 | 13.26 | 13.77 | 14.14 | 11.34 | 13.38 | 12.63 | 12.98 | 11.45 | 12.31 | + |
| 1700017B05Rik | 14.04 | 16.65 | 13.9 | 9.71 | 8.1 | 8.08 | 20.21 | 15.17 | 18.49 | 23.04 | 20.74 | 24.85 | + |
| Tnpo3 | 43.55 | 40.4 | 40.19 | 26.13 | 28.48 | 27.15 | 28.78 | 33.37 | 29.76 | 26.51 | 29.02 | 27.27 | + |
| 9930021J03Rik | 0.73 | 0.69 | 0.82 | 1.28 | 1.18 | 1.52 | 1.47 | 1.41 | 1.22 | 1.58 | 1.72 | 0.88 | + |
| Rbm8a | 61.5 | 62.37 | 60.43 | 34.31 | 32.83 | 35.32 | 27.69 | 39.38 | 34.91 | 22.45 | 24.4 | 22.25 | + |
| Mnat1 | 12.67 | 12.85 | 12.81 | 19.53 | 18.74 | 19.01 | 20.39 | 19.81 | 18.05 | 18.75 | 20.79 | 19.01 | + |
| Fem1a | 5.85 | 5.89 | 5.65 | 3.34 | 2.83 | 3.38 | 1.96 | 2.76 | 2.95 | 3 | 2.2 | 2.7 | + |
| Ctsc | 44.32 | 49.64 | 45.05 | 31.15 | 29.72 | 34 | 21.39 | 31.67 | 28.06 | 36.17 | 28.45 | 34.86 | + |
| Zrsr2 | 4.22 | 4.62 | 5.17 | 2.81 | 2.53 | 3.02 | 2.06 | 2.48 | 3.27 | 2.59 | 2.97 | 3.04 | + |
| Ngly1 | 29.07 | 29.51 | 26.88 | 36.08 | 31.95 | 33.9 | 56.75 | 47.46 | 49.02 | 43.56 | 45.48 | 50.87 | + |
| Dbi | 291.84 | 286.83 | 274.2 | 131.12 | 157.38 | 160.47 | 129.76 | 150.99 | 144.31 | 106.03 | 99.85 | 100.75 | + |
| Ddx18 | 66.53 | 66.01 | 66.93 | 30.41 | 36.33 | 31.56 | 29.95 | 37.58 | 34.38 | 25.79 | 26.91 | 26.5 | + |
| H2-Ke6 | 1.84 | 2.96 | 3.42 | 8.46 | 7.38 | 6.02 | 7.93 | 7.98 | 7.47 | 9.44 | 6.27 | 7.06 | + |
| Psma6 | 286.15 | 294.29 | 287.41 | 164.95 | 164.48 | 161.23 | 148.76 | 173.16 | 160.72 | 135.15 | 135.75 | 152.05 | + |
| 2610002M06Rik | 4.23 | 4.55 | 3.98 | 3 | 3.29 | 2.83 | 3.39 | 3.3 | 2.95 | 2.27 | 3 | 2.55 | + |
| Tfam | 16.06 | 17.37 | 16.36 | 11.68 | 12.57 | 11.57 | 11.35 | 11.49 | 12.08 | 9.62 | 10.57 | 11.23 | + |
| Lsm5 | 77.71 | 93.95 | 84.92 | 63.71 | 60.87 | 64.92 | 48.29 | 53.69 | 58.04 | 54.01 | 62.59 | 56.95 | + |
| Fau | 1179.03 | 1211.72 | 1194.06 | 2426.33 | 2174.88 | 2306.02 | 2514.66 | 2030.87 | 2200.34 | 2877.09 | 3020.6 | 3064.63 | + |
| Ptprc | 133.05 | 138.1 | 133.56 | 140.42 | 118.66 | 127.24 | 135.61 | 121.3 | 134.69 | 165.49 | 150.25 | 149.42 | + |
| Phc1 | 1.02 | 0.96 | 0.95 | 2.04 | 1.73 | 1.81 | 2 | 2.06 | 1.91 | 2.34 | 1.86 | 1.68 | + |
| Twistnb | 35.99 | 41.52 | 34.69 | 23.63 | 25.46 | 22.97 | 18.27 | 21.53 | 23.39 | 19.28 | 16.87 | 18.49 | + |
| Eif4enif1 | 15.63 | 16.41 | 18.4 | 14.13 | 13.52 | 12.31 | 12.22 | 13.34 | 12.95 | 12.82 | 14.78 | 12.08 | + |
| Ltbp3 | 0.94 | 0.91 | 0.66 | 1.44 | 1.83 | 1.48 | 3.24 | 2.45 | 2.93 | 2.63 | 2.47 | 3.47 | + |
| Dbt | 7.73 | 7.98 | 8.6 | 3.76 | 4.95 | 4.81 | 3.83 | 4.23 | 5.55 | 3.83 | 3.09 | 3.65 | + |
| Armc10 | 36.53 | 37.99 | 36.76 | 46.98 | 43.66 | 44.33 | 50.56 | 49.9 | 50.44 | 45.35 | 49.16 | 45.94 | + |

| | | | | | | | | | | | | | |
|---|---|---|---|---|---|---|---|---|---|---|---|---|---|
| Cog1 | 22.41 | 21.42 | 19.92 | 18.44 | 18.3 | 18.77 | 15.16 | 18.2 | 14.88 | 17.04 | 15.89 | 18.24 | + |
| Mrps36 | 110.99 | 113.48 | 108.68 | 82.16 | 86.03 | 90.14 | 83.13 | 84.06 | 80.9 | 72.44 | 80.63 | 74.85 | + |
| Parp9 | 25.62 | 25.07 | 24.78 | 29.83 | 27.02 | 25.33 | 28.16 | 19.76 | 19.66 | 35.92 | 30.45 | 37.58 | + |
| Mrpl2 | 61.78 | 59.3 | 57.01 | 44.01 | 45.75 | 41.84 | 39.18 | 43.41 | 44.64 | 32.79 | 38.21 | 35.43 | + |
| Ssbp1 | 72.11 | 77.28 | 73.51 | 50.07 | 50.52 | 52.75 | 43.98 | 52.02 | 52.61 | 42.31 | 41.33 | 42.23 | + |
| Tomm70a | 14.78 | 14.48 | 14.3 | 9.3 | 9.09 | 9.16 | 8.7 | 9.77 | 9.26 | 7.86 | 7.96 | 8.7 | + |
| Klhl22 | 10.18 | 10.05 | 11.05 | 15.43 | 17.45 | 16.49 | 19.29 | 18.89 | 15 | 18.51 | 20.03 | 21.38 | + |
| Gng2 | 16.47 | 16.59 | 15.95 | 32.29 | 29.28 | 28.26 | 29.85 | 23.29 | 24.43 | 31.8 | 32.53 | 31.37 | + |
| Nop16 | 90.31 | 94.78 | 101.52 | 37.1 | 40.49 | 40.53 | 29.89 | 47.78 | 40.29 | 23.82 | 25.31 | 25.4 | + |
| Snrpd1 | 259.15 | 257.15 | 246.61 | 130.26 | 139.54 | 143.54 | 115.84 | 137.4 | 126.98 | 95.7 | 113.76 | 98.23 | + |
| Dpm2 | 51.19 | 46.66 | 53.62 | 39.35 | 38.86 | 39.11 | 30.83 | 38.95 | 38.51 | 36.94 | 34 | 39.43 | + |
| Atf4 | 180.03 | 183.74 | 177.5 | 201.29 | 220.47 | 218.59 | 257.49 | 291.21 | 281.62 | 207.1 | 262.32 | 206.62 | + |
| Tnfrsf14 | 3.2 | 2.35 | 2.3 | 4.6 | 4.12 | 5.21 | 5.21 | 4.6 | 4.39 | 6.34 | 4.95 | 4.84 | + |
| Hspa14 | 96.38 | 98.82 | 87.6 | 49.9 | 58.82 | 60.89 | 42.67 | 48.59 | 50.54 | 37.94 | 36.82 | 42.13 | + |
| Fancm | 0.94 | 0.93 | 1.05 | 0.37 | 0.51 | 0.72 | 0.52 | 0.51 | 0.51 | 0.44 | 0.37 | 0.48 | + |
| Cdk11b | 37.88 | 36.55 | 34.29 | 31.95 | 29.99 | 29.67 | 27.48 | 27.81 | 26.4 | 26.74 | 29.52 | 25.07 | + |
| Znhit6 | 15.36 | 15.3 | 17.32 | 9.1 | 10.01 | 10.16 | 9.32 | 10.46 | 11.06 | 7.5 | 7.06 | 8.03 | + |
| Rab5b | 15.6 | 14.47 | 15.55 | 24.24 | 23.52 | 21.69 | 25.89 | 25.76 | 24.87 | 23.68 | 28.78 | 25.53 | + |
| Phf21a | 1.42 | 1.24 | 1.48 | 3.21 | 3.09 | 3.22 | 2.88 | 2.81 | 2.74 | 3.18 | 4.17 | 3.27 | + |
| Abcd3 | 35.32 | 29.33 | 35.15 | 15.19 | 19.93 | 18.44 | 8.83 | 14.76 | 14.52 | 11.04 | 11.83 | 11.56 | + |
| Acadl | 133.05 | 141.07 | 137.95 | 157.43 | 161.63 | 156.36 | 334.34 | 334.26 | 338.72 | 297.01 | 281.54 | 326.71 | + |
| Cdk2ap1 | 13.38 | 13.45 | 14.36 | 7.62 | 7.03 | 7.72 | 7.69 | 9.86 | 9.19 | 6.18 | 7.36 | 7.14 | + |
| Prpsap1 | 83.01 | 86.98 | 81.62 | 70.51 | 73.96 | 71.58 | 63.12 | 71.53 | 61.72 | 66.67 | 65.41 | 67.39 | + |
| Ccdc127 | 21.72 | 22.14 | 21.53 | 14.59 | 15.76 | 17.87 | 14.78 | 14.35 | 13.84 | 13.05 | 14.61 | 14.15 | + |
| Timm17a | 208.33 | 198.49 | 200.79 | 71.28 | 77.71 | 78.17 | 70.71 | 93.7 | 88.99 | 56.1 | 57.49 | 61.1 | + |
| Sqrdl | 2.88 | 2.94 | 2.53 | 9.08 | 8.54 | 6.82 | 7.87 | 7.91 | 7.9 | 8.77 | 10 | 9.63 | + |
| Nudt1 | 21.28 | 24.97 | 20.38 | 8.14 | 9.17 | 8.42 | 4.53 | 7.82 | 7.54 | 3.72 | 4.2 | 5.52 | + |
| Vwa5a | 5.84 | 5.82 | 6 | 17.93 | 15.7 | 15.14 | 10.84 | 9.92 | 11.47 | 11.04 | 8.8 | 10.39 | + |
| A730017L22Rik | 6.3 | 5.51 | 5.54 | 9.85 | 9.22 | 10.02 | 9.16 | 8 | 8.41 | 11.12 | 9.13 | 9.42 | + |
| Scoc | 30.87 | 30.98 | 33.48 | 19.8 | 19.92 | 22.51 | 12.19 | 18.93 | 17.68 | 13.94 | 16.04 | 12.55 | + |
| 2010016I18Rik | 11.26 | 11.38 | 12.55 | 38.87 | 36.06 | 34.95 | 40.86 | 30.48 | 35.71 | 40.42 | 39.63 | 44.86 | + |
| Ssr4 | 195.74 | 210.48 | 192.71 | 176.28 | 176.22 | 188.54 | 160.17 | 179.96 | 172.38 | 160.71 | 165.66 | 155.09 | + |
| Xrcc5 | 8.89 | 11.36 | 10.83 | 6.65 | 5.59 | 6.26 | 4.74 | 4.32 | 3.38 | 3.8 | 3.23 | 3.97 | + |
| Cdc42ep4 | 0.56 | 0.69 | 0.42 | 1.82 | 2.29 | 1.93 | 1.72 | 1.26 | 1.59 | 1.77 | 2.03 | 1.27 | + |
| Dip2a | 0.6 | 0.71 | 0.58 | 1.17 | 1.21 | 1.45 | 1.5 | 1.08 | 1.2 | 1.69 | 1.4 | 1.6 | + |
| Pld2 | 0.21 | 0.37 | 0.28 | 0.81 | 0.97 | 0.74 | 0.88 | 0.9 | 0.66 | 0.92 | 0.7 | 0.52 | + |
| Ptma | 1272.28 | 1468.02 | 1345.65 | 719.67 | 722.83 | 744.07 | 749.04 | 677 | 714.21 | 595.04 | 618.36 | 605.07 | + |
| Cdc6 | 19.79 | 21.81 | 20.83 | 6.15 | 6.31 | 5.99 | 3.64 | 4.48 | 3.69 | 1.79 | 1.77 | 1.76 | + |
| Bag2 | 47.55 | 50.42 | 44.85 | 21.8 | 25.93 | 25.83 | 22.54 | 31.39 | 29.24 | 17.02 | 17.05 | 18.61 | + |
| Auh | 21.58 | 23.66 | 23.49 | 35.82 | 38.66 | 38.45 | 42.51 | 40.83 | 42.72 | 39.64 | 47.01 | 45.21 | + |
| Hipk2 | 1.48 | 1.56 | 1.67 | 2.16 | 2.54 | 3.37 | 3.54 | 2.8 | 2.74 | 2.3 | 2.88 | 2.59 | + |
| Anp32e | 92.48 | 91.21 | 92.04 | 46.08 | 54.28 | 52.2 | 42.5 | 49.13 | 41.59 | 31.99 | 36.47 | 34.96 | + |
| Ptrf | 2.08 | 0.96 | 1.9 | 0.1 | 0.19 | 0.16 | 0.64 | 0.46 | 0.92 | 0.5 | 0.52 | 0.36 | + |
| Mbnl2 | 5.7 | 5.82 | 6.3 | 10.23 | 8.93 | 8.77 | 10.19 | 10.39 | 10.58 | 10.82 | 11.58 | 10.65 | + |
| Nomo1 | 29.46 | 26.75 | 29.19 | 7.48 | 9 | 9.25 | 7.84 | 13.57 | 11.65 | 7.48 | 7.09 | 5.93 | + |
| Gtf2i | 14.46 | 15.41 | 15.28 | 20.39 | 20.16 | 17.79 | 25.16 | 16.55 | 20.05 | 39.13 | 31.14 | 37.34 | + |
| Usf1 | 10 | 9.46 | 10.45 | 10.72 | 11.04 | 11.1 | 13.28 | 11.18 | 12.95 | 16.12 | 17.33 | 18.66 | + |
| Myo19 | 6.98 | 5.49 | 6.18 | 1.78 | 1.82 | 2.24 | 1.54 | 2.24 | 1.59 | 1.93 | 1.17 | 2.74 | + |
| Atp5k | 105.21 | 117.56 | 119.43 | 77.27 | 68.62 | 70.04 | 77.14 | 66.78 | 70 | 62.5 | 60.36 | 64.75 | + |
| Fam98c | 4.48 | 7.39 | 9.66 | 16.24 | 14.31 | 14.8 | 19.52 | 14.52 | 16.21 | 17.25 | 16.49 | 17.2 | + |
| Mrps25 | 13.14 | 12.46 | 13.31 | 9.01 | 9.17 | 9.16 | 10.05 | 9.54 | 9.61 | 9.46 | 10.37 | 9.3 | + |
| Ndufa8 | 115.94 | 129.72 | 113.77 | 100.49 | 97.65 | 100.46 | 97.41 | 101.96 | 91.57 | 89.1 | 93.01 | 92.43 | + |
| Puf60 | 119.53 | 117.59 | 125.26 | 88.39 | 84.07 | 86.44 | 84.87 | 80.42 | 77.81 | 84.79 | 79.63 | 81.02 | + |
| Lmo4 | 2.85 | 2.8 | 2.6 | 4.75 | 6.34 | 4.44 | 4.07 | 4.8 | 5.25 | 4.26 | 6.04 | 5.64 | + |
| Fbxl4 | 9.33 | 8.03 | 8.75 | 11.93 | 13.62 | 14.58 | 11.17 | 11.63 | 12.9 | 11.38 | 10.24 | 12.7 | + |
| Emc7 | 100.62 | 105.52 | 107.78 | 69.81 | 74.42 | 77.22 | 72.99 | 75.62 | 72.87 | 67.75 | 78.94 | 68.49 | + |
| 2410004B18Rik | 11.07 | 12.29 | 11.7 | 5.9 | 7.81 | 5.43 | 3.83 | 6.45 | 6.93 | 6.78 | 7.08 | 6.21 | + |
| Mical1 | 25.98 | 27.14 | 26.25 | 35.7 | 35.29 | 32.92 | 47.59 | 38.22 | 44.22 | 41.41 | 36.32 | 41.3 | + |
| Cd3d | 202.81 | 203.07 | 191.29 | 318.71 | 286.58 | 316.52 | 345.15 | 280.35 | 306.69 | 326.18 | 285.36 | 341.75 | + |
| Gpx1 | 217.26 | 221.04 | 217.94 | 131.13 | 128.91 | 132.61 | 138.85 | 155.29 | 154.79 | 121.53 | 125.92 | 117.71 | + |
| Shroom3 | 7.08 | 6.4 | 8.02 | 2.02 | 3.45 | 2.78 | 3.74 | 5.65 | 5.74 | 1.14 | 1.02 | 0.94 | + |
| Stx5a | 16 | 14.19 | 14.29 | 28.35 | 28.85 | 28.28 | 31.77 | 29.56 | 27.58 | 32.35 | 36.44 | 33.75 | + |
| Etfb | 151.57 | 157.41 | 152.17 | 87.94 | 84.04 | 89.2 | 83.91 | 99.14 | 91.63 | 54.32 | 66.2 | 61.28 | + |
| Cd44 | 87.52 | 67.67 | 84.19 | 31.27 | 31.5 | 30.42 | 28.38 | 38.32 | 33.27 | 19.95 | 21.44 | 20.91 | + |
| Tmem141 | 2.8 | 3.2 | 2.97 | 6.05 | 6.12 | 4.88 | 7.15 | 4.78 | 5.52 | 7.12 | 6.93 | 4.69 | + |
| Lnp | 6.72 | 6.99 | 6.08 | 3.28 | 4.16 | 4.01 | 3.22 | 4.01 | 3.69 | 3.97 | 3.75 | 2.94 | + |
| C2cd3 | 2.81 | 3.27 | 3.48 | 3.57 | 3.45 | 3.19 | 3.24 | 2.79 | 3.11 | 4.05 | 4.81 | 4.13 | + |
| Gm15645 | 16.45 | 21.44 | 18.21 | 6.81 | 6.99 | 8.02 | 8.9 | 8.87 | 8.48 | 5.89 | 7.03 | 5.96 | + |
| Sh2b1 | 8.81 | 8.32 | 9.62 | 16.28 | 13.68 | 15.03 | 21.04 | 15.02 | 18.76 | 18.82 | 17.2 | 18.33 | + |
| Nol11 | 46.01 | 47.3 | 47.01 | 24.36 | 28.49 | 25.39 | 17.89 | 25.26 | 23.8 | 17.71 | 19.52 | 17.74 | + |
| Uqcrh | 318.81 | 300.74 | 289.27 | 417.73 | 366.38 | 395.73 | 433.82 | 344.61 | 345.1 | 523.79 | 549.81 | 579.6 | + |
| Uqcrc1 | 326.11 | 324.04 | 316.9 | 214.75 | 218.07 | 220.81 | 204.11 | 226.05 | 218.2 | 178.39 | 183.55 | 179.83 | + |
| Trim62 | 0.12 | 0.29 | 0.17 | 0.95 | 0.8 | 0.54 | 0.41 | 0.31 | 0.65 | 0.89 | 0.6 | 0.49 | + |
| Chaf1a | 18.85 | 24.91 | 24.79 | 12.76 | 11.68 | 12.86 | 9.45 | 10.19 | 9.81 | 7.02 | 7.48 | 6.33 | + |
| Cherp | 19.08 | 19.75 | 21.97 | 15.67 | 13.72 | 14.36 | 12.97 | 13.76 | 11.83 | 14.06 | 13.67 | 13.99 | + |
| Psmb3 | 414.07 | 432.95 | 398.59 | 223.08 | 240.9 | 238.25 | 217.67 | 268.84 | 258.93 | 176.86 | 186.42 | 187.04 | + |
| Pex13 | 7.23 | 8.57 | 7.49 | 12.54 | 11.45 | 13.61 | 14.89 | 12.85 | 12.19 | 15.52 | 17.24 | 13.92 | + |
| Ick | 1.17 | 1.57 | 1.31 | 1.81 | 2.08 | 1.83 | 2.25 | 2.21 | 1.9 | 2.05 | 2.34 | 1.75 | + |
| Nqo2 | 7.23 | 7.05 | 6.8 | 3.92 | 3.42 | 3.73 | 5.37 | 6.43 | 5.89 | 4.41 | 4.06 | 5.44 | + |
| Rab2a | 10.38 | 12.02 | 10.49 | 9.03 | 7.53 | 7.97 | 7.74 | 9.11 | 8.06 | 8.48 | 7.82 | 7.62 | + |
| Tex30 | 84.73 | 82.9 | 79.92 | 36.66 | 41.88 | 46.33 | 27.59 | 34.04 | 34.09 | 28.13 | 26.99 | 23.38 | + |
| Icam2 | 12.15 | 14.24 | 13.35 | 3.76 | 4.48 | 3.85 | 4.53 | 3.34 | 4.6 | 3.21 | 2.48 | 4.59 | + |
| Ssh1 | 2.82 | 3.32 | 2.86 | 6.11 | 5.78 | 5.28 | 7.22 | 5.53 | 7.27 | 5.92 | 6.6 | 6.81 | + |
| Scyl2 | 8.64 | 8.23 | 7.97 | 11.49 | 12.38 | 12.76 | 11.26 | 12.77 | 12.26 | 12.25 | 14.99 | 11.56 | + |
| Rps29 | 953.49 | 1038.92 | 1046.77 | 1413.13 | 1519.27 | 1533.35 | 1426.02 | 1289.74 | 1204.26 | 1449.29 | 1630.19 | 1431.02 | + |
| Dhps | 50.21 | 49.24 | 52.49 | 26.11 | 24.86 | 24.82 | 23.12 | 32.44 | 29.63 | 25.43 | 20.4 | 22.61 | + |
| Ccne1 | 76.65 | 81.2 | 80.17 | 33.35 | 33.44 | 33.19 | 36.19 | 41.77 | 41.36 | 36.41 | 35.26 | 32.56 | + |
| Sephs2 | 65.32 | 67.87 | 61.47 | 38.08 | 40.57 | 37.78 | 34.54 | 40.11 | 42.88 | 42.84 | 32.55 | 36.11 | + |
| Orc2 | 38.74 | 36.01 | 36.75 | 17.32 | 18.67 | 19.01 | 12.69 | 19.71 | 19.24 | 12.39 | 16.09 | 12.26 | + |
| Myo1e | 3.5 | 3.06 | 2.92 | 1.11 | 0.8 | 0.64 | 1.98 | 2.83 | 2.33 | 0.9 | 1.25 | 0.87 | + |
| Serpinc1 | 4.76 | 3.61 | 3.7 | 0.5 | 0.35 | 0.83 | 1.28 | 2.2 | 1.49 | 0.53 | 0.48 | 0.51 | + |
| Tmem39a | 21.48 | 24.29 | 21.62 | 12.68 | 12.36 | 14.14 | 13.15 | 18.65 | 15.72 | 12.74 | 14.37 | 11.1 | + |
| Enoph1 | 27.68 | 31.6 | 29.97 | 17.08 | 15.12 | 16.47 | 14.07 | 16.06 | 15.59 | 12.81 | 13.16 | 12.59 | + |
| Abca7 | 5.12 | 5.13 | 5.16 | 10.46 | 8.78 | 8.5 | 11.72 | 8.46 | 10.19 | 12.68 | 12.51 | 12.71 | + |
| Ikzf5 | 4.36 | 4.19 | 4.15 | 2.94 | 2.77 | 2.67 | 2.25 | 3.1 | 3.52 | 3.27 | 2.88 | 2.65 | + |
| Hnrnpu | 41.7 | 42.24 | 40.39 | 20.29 | 22.2 | 20.5 | 18.8 | 19.42 | 18.99 | 18.08 | 15.38 | 18.5 | + |
| Rrm1 | 125.91 | 125.44 | 120.31 | 47.26 | 56.93 | 55.34 | 34.7 | 41.01 | 37.34 | 28.15 | 29.22 | 29.69 | + |
| Irf7 | 11.33 | 11.17 | 7.76 | 14.89 | 15.12 | 14.81 | 8.51 | 9.22 | 6.92 | 25.27 | 18.08 | 24.91 | + |
| Naga | 9.22 | 8.55 | 10.26 | 21.87 | 17.82 | 19.83 | 20.43 | 17.35 | 18.37 | 19.35 | 23.13 | 22.63 | + |
| Clybl | 23.41 | 20.77 | 23.26 | 43.49 | 45.97 | 41.73 | 61.73 | 50.11 | 51.84 | 46.35 | 47.82 | 58.29 | + |
| Ppp2r4 | 62.28 | 61.08 | 58.39 | 46.89 | 46.77 | 48.85 | 50.8 | 51.1 | 46.94 | 43.95 | 47.69 | 44.79 | + |
| Ptms | 4.5 | 5.29 | 5.06 | 8.54 | 8 | 8.72 | 10.6 | 10.93 | 9.23 | 11.84 | 10.14 | 10.73 | + |
| Tomm5 | 308.81 | 304.94 | 314.74 | 114.62 | 128.48 | 127.08 | 116.93 | 146.17 | 141.05 | 102.4 | 104.27 | 95.45 | + |
| Brcc3 | 72.48 | 74.54 | 72.03 | 47.36 | 44.66 | 52.73 | 47.37 | 49.19 | 49.55 | 40.04 | 44.42 | 44.15 | + |
| Slirp | 156.71 | 148.36 | 143.97 | 104.52 | 108.01 | 120.28 | 120.53 | 108.75 | 103.19 | 91.16 | 100.74 | 94.85 | + |
| Pde4c | 1.56 | 1.28 | 1.37 | 3.57 | 3.93 | 4.37 | 3.58 | 3.45 | 2.67 | 3.97 | 4.11 | 4.36 | + |
| Psmd9 | 19.79 | 18.05 | 16.74 | 9.29 | 11.09 | 10.63 | 8.81 | 11.74 | 10.34 | 8.52 | 8.31 | 9.16 | + |
| Ssrp1 | 91.91 | 101.56 | 90.6 | 52.54 | 54.29 | 49.69 | 49.1 | 56.1 | 53.62 | 39.99 | 40.26 | 44.43 | + |
| Tufm | 131.67 | 131.47 | 128.16 | 74.38 | 76.71 | 79.85 | 85.58 | 85.28 | 89.33 | 75.12 | 74.59 | 73.22 | + |
| Nuf2 | 22.48 | 19.56 | 20.6 | 10.85 | 17.73 | 13.58 | 8.44 | 9.11 | 7.45 | 8.41 | 9.76 | 8.18 | + |
| Rpl12 | 1752.52 | 1817.33 | 1825.64 | 2848.67 | 2818.09 | 2948.09 | 2591.95 | 2658.77 | 2627.13 | 2910.72 | 3094.15 | 2874.2 | + |
| Usp5 | 49.51 | 47.51 | 47.73 | 38.33 | 38.21 | 35.55 | 41.9 | 38.86 | 40.31 | 37.01 | 39.5 | 36.23 | + |
| Mtmr6 | 26.31 | 26.18 | 25.85 | 30.46 | 30.47 | 29.88 | 32.65 | 33.68 | 31.25 | 38.92 | 38.52 | 38.31 | + |
| Lrrc14 | 13.97 | 13.9 | 16.3 | 11.63 | 11.43 | 11.45 | 7.17 | 8.98 | 9.09 | 9.56 | 7.5 | 9.36 | + |
| Mettl25 | 6.08 | 5.88 | 5.2 | 2.91 | 2.39 | 3.22 | 3.33 | 3.06 | 4.03 | 2.22 | 2.42 | 2.54 | + |
| Gm14005 | 2.53 | 3.2 | 2.74 | 0.8 | 1.33 | 1.37 | 1.48 | 2.01 | 1.27 | 1.76 | 1.48 | 1.58 | + |
| Cd3eap | 14.38 | 16.82 | 15.35 | 7.46 | 8.15 | 7.95 | 8.18 | 7.51 | 7.99 | 7.03 | 6.41 | 7.1 | + |
| Mettl13 | 16.58 | 15.02 | 16.87 | 7.59 | 9.17 | 9.42 | 6.69 | 10.08 | 8.23 | 5.82 | 7.07 | 7 | + |
| Cyb5r3 | 43.4 | 41.23 | 41.66 | 34.73 | 36.22 | 36.86 | 34.03 | 36.05 | 34.33 | 30.18 | 34.46 | 37.06 | + |
| Otub1 | 81.77 | 78.7 | 80.41 | 59.48 | 62.65 | 63.43 | 51.26 | 57.37 | 59.34 | 46.67 | 47.46 | 51.45 | + |
| Blvrb | 6.07 | 7.06 | 7.31 | 19.58 | 19.85 | 21.68 | 19.4 | 15.41 | 18.86 | 23.08 | 23.94 | 22.72 | + |
| Rhbdd1 | 8.49 | 8.04 | 9.03 | 15.56 | 16.38 | 15.85 | 17.44 | 21 | 17.52 | 12.82 | 15.39 | 14.97 | + |
| Dhrs7b | 24.39 | 20.57 | 23.86 | 15.77 | 16.98 | 15.91 | 14.6 | 17.14 | 16.14 | 15.04 | 15.97 | 13.85 | + |
| H2-Q5 | 55.69 | 50.31 | 53.61 | 113.36 | 108.13 | 97.58 | 121.07 | 108.21 | 113.38 | 150.33 | 166.22 | 139.52 | + |
| Slc41a3 | 0.94 | 0.26 | 0.57 | 2.58 | 2.18 | 2.24 | 4 | 2.92 | 2.52 | 4.23 | 4.15 | 4.32 | + |
| Lsg1 | 39.18 | 36.78 | 37.92 | 21.09 | 20.52 | 21.89 | 16.35 | 21.01 | 21.7 | 19.46 | 15.64 | 16.63 | + |
| Utp20 | 3.72 | 3.62 | 4.16 | 1.4 | 1.68 | 1.47 | 1.53 | 1.84 | 1.82 | 0.88 | 1.12 | 1.33 | + |
| Ggh | 33.15 | 25.32 | 24.09 | 13.92 | 12.8 | 12.65 | 14.65 | 16.83 | 16.27 | 12.78 | 13.2 | 14.35 | + |
| Rnaseh2c | 37.91 | 39.36 | 42.02 | 22.21 | 21.53 | 28.23 | 22.51 | 20.18 | 17.85 | 20.99 | 22.23 | 24.62 | + |
| Mllt10 | 3.53 | 3.6 | 3.21 | 4.42 | 3.89 | 4.04 | 4.56 | 4.19 | 3.99 | 5.66 | 6.14 | 5.1 | + |
| Etfa | 103.25 | 106.39 | 102.79 | 81.03 | 83.53 | 83.87 | 83.86 | 90.15 | 86.38 | 74.06 | 81.07 | 67.75 | + |
| 6030458C11Rik | 15.49 | 14.93 | 14.68 | 10.38 | 9.91 | 10.07 | 9.29 | 9.7 | 8.61 | 8.43 | 7.78 | 7.44 | + |
| Utrn | 2.22 | 1.96 | 2.48 | 5.55 | 4.18 | 4.39 | 5.49 | 4.37 | 4.41 | 4.34 | 5.37 | 4.3 | + |
| Stt3a | 155.85 | 163.6 | 159.09 | 123.34 | 130.51 | 139.43 | 128.81 | 151.31 | 139.5 | 131.38 | 152.1 | 123.67 | + |
| Pnpt1 | 42.73 | 40.2 | 44.51 | 21.06 | 21.17 | 24.33 | 18.26 | 20.24 | 22.94 | 17.81 | 17.29 | 16.27 | + |
| Pole3 | 25.07 | 27.2 | 28.97 | 12.35 | 14.07 | 14.27 | 13.22 | 11.18 | 12.81 | 9.36 | 9.16 | 9.43 | + |
| Napsa | 11.17 | 10.75 | 9.02 | 81.92 | 32.27 | 51.31 | 6.4 | 6.7 | 6.64 | 14.53 | 12.6 | 18.05 | + |
| Spag9 | 5.04 | 5.1 | 3.91 | 7.05 | 7.87 | 7.43 | 7.14 | 5.92 | 6.35 | 6.28 | 7.82 | 6.02 | + |
| 2810004N23Rik | 44.04 | 40.48 | 44.01 | 23.83 | 22.54 | 22.84 | 19.67 | 22.61 | 25.58 | 16.56 | 16.23 | 17.98 | + |
| F2rl3 | 9.49 | 11.13 | 9.64 | 4.66 | 3.47 | 3.69 | 2.87 | 4.47 | 4.05 | 2.37 | 2.1 | 1.83 | + |
| Fam214b | 2.72 | 3.43 | 3.31 | 3.97 | 5.15 | 5.07 | 5.74 | 5.64 | 4.85 | 6.26 | 4.8 | 5.1 | + |
| Atad5 | 2.15 | 1.95 | 1.88 | 0.9 | 1.09 | 1.39 | 0.94 | 1.11 | 0.91 | 1.07 | 1.04 | 0.79 | + |
| Glyr1 | 47.19 | 43.53 | 46.33 | 40.21 | 40.29 | 37.69 | 37.25 | 34.04 | 36.26 | 37.01 | 35.39 | 35.66 | + |
| Fam203a | 23.75 | 24.61 | 22.65 | 13.65 | 14.58 | 12.7 | 10.51 | 15.12 | 13.73 | 10.38 | 8.77 | 10.45 | + |
| Rpp14 | 26.38 | 26.28 | 27.97 | 13.55 | 14.22 | 13.36 | 10.76 | 15.76 | 13.96 | 11.43 | 8.89 | 10.91 | + |
| Cd40lg | 5.02 | 7.63 | 11.12 | 0.46 | 1.4 | 0.51 | 0.28 | 0.47 | 1.03 | 0.18 | 0.59 | 0.32 | + |
| Lonrf3 | 0.05 | 0.06 | 0.13 | 0.52 | 0.4 | 0.59 | 0.37 | 0.25 | 0.28 | 0.21 | 0.61 | 0.5 | + |
| Fam49b | 26.92 | 31.12 | 32.07 | 52.19 | 51.46 | 45.64 | 46.88 | 45.86 | 47.74 | 55.95 | 57.69 | 45.51 | + |
| Nedd8 | 254.44 | 263.1 | 259.47 | 155.06 | 150.87 | 162.93 | 146.54 | 171.81 | 164.07 | 148.66 | 155.82 | 144.56 | + |
| Nudcd3 | 27.85 | 25.49 | 26.96 | 19.59 | 20.4 | 17.51 | 20 | 22.17 | 20.11 | 22.03 | 19.85 | 19.84 | + |
| Dclre1b | 5.86 | 5.48 | 5.18 | 3.65 | 3.75 | 3.99 | 1.96 | 2.43 | 2.05 | 2.66 | 1.82 | 2.26 | + |

| | | | | | | | | | | | | |
|---|---|---|---|---|---|---|---|---|---|---|---|---|
| Lrrc59 | 32.29 | 36 | 39.86 | 22.61 | 20.75 | 19.45 | 20.44 | 24.74 | 23.81 | 18.63 | 17.76 | 14.83 | + |
| Rn45s | 15125.56 | 15570.73 | 15762.05 | 25424.96 | 24343.03 | 26661.25 | 25913.53 | 23869.62 | 24440.09 | 27950.41 | 30052.88 | 28690.85 | + |
| Pfkfb4 | 1.8 | 2.34 | 2.24 | 3.89 | 4.33 | 4.36 | 4.63 | 3.16 | 4.95 | 4.31 | 3.96 | 4.06 | + |
| Zfp472 | 10.02 | 9.89 | 9.9 | 8.82 | 7.59 | 7.2 | 4.4 | 6.84 | 4.64 | 7.1 | 7.46 | 5.8 | + |
| Fh1 | 91.61 | 89.71 | 92.46 | 40.43 | 42.95 | 43.7 | 39.89 | 45.13 | 40.5 | 29.79 | 30.78 | 29.74 | + |
| Gatsl2 | 1.03 | 1.12 | 1.85 | 4.3 | 3.09 | 3.21 | 3.88 | 3.38 | 3.57 | 4.46 | 4.18 | 3.45 | + |
| Cct6a | 354.56 | 356.58 | 370.04 | 157.59 | 166.91 | 166.98 | 126.27 | 178.75 | 168.03 | 112.1 | 119.74 | 115.53 | + |
| Rnf166 | 6.79 | 5.42 | 4.73 | 20.52 | 13.55 | 14.13 | 20.14 | 13.21 | 14.89 | 20.02 | 22.88 | 29.16 | + |
| Nbr1 | 8.87 | 8.89 | 9.38 | 14.25 | 14.47 | 14.99 | 15.42 | 13.92 | 15.14 | 15.6 | 17.76 | 16.37 | + |
| Trnt1 | 59.06 | 55.01 | 58.4 | 33.89 | 36.87 | 37.29 | 27.95 | 37.96 | 34.43 | 27.62 | 27.73 | 24.57 | + |
| Rab34 | 22.48 | 19.76 | 24.18 | 42.83 | 47.76 | 51.04 | 23.75 | 22.57 | 22.17 | 28.6 | 37.31 | 31.45 | + |
| Plxna1 | 0.71 | 0.44 | 0.65 | 1.41 | 0.96 | 1.31 | 0.87 | 1.32 | 1.37 | 1.21 | 1.47 | 0.75 | + |
| Il1a | 27.43 | 26.85 | 22.4 | 0.22 | 1.26 | 0.3 | 0.2 | 4.57 | 1.4 | 0.29 | 0.42 | 0.39 | + |
| Ttc7b | 2.72 | 2.2 | 2.2 | 1.19 | 0.92 | 0.94 | 1.55 | 1.59 | 1.59 | 1.2 | 1.18 | 0.97 | + |
| Ddx3x | 29.59 | 32.26 | 31.19 | 13.19 | 13.78 | 12.73 | 9.85 | 12.05 | 13.42 | 12.17 | 9.41 | 9.85 | + |
| Uqcrb | 305.06 | 324.92 | 299.66 | 214.11 | 210.8 | 228.46 | 214.09 | 221.71 | 233.48 | 207.84 | 205.48 | 202.61 | + |
| Tnfrsf1b | 13.46 | 14.83 | 12.14 | 12.63 | 13.38 | 13.56 | 15.64 | 13.79 | 14.51 | 19.57 | 16.57 | 19.19 | + |
| Snx18 | 1.29 | 1.35 | 1.26 | 3.45 | 2.39 | 3.51 | 2.67 | 2.21 | 2.38 | 2.69 | 3.22 | 3.1 | + |
| Dlgap4 | 10.8 | 9.73 | 9.57 | 13.48 | 11.96 | 12.09 | 16.9 | 14.29 | 15.82 | 14.81 | 14.49 | 13.56 | + |
| Sh3glb1 | 18.37 | 23.93 | 27.4 | 49.13 | 50.18 | 51.11 | 65.14 | 52.88 | 56.77 | 66.14 | 77.21 | 62.18 | + |
| Pola2 | 38.03 | 38.3 | 38.47 | 20.82 | 21.83 | 22.32 | 16.51 | 18.69 | 19.02 | 16.53 | 17.69 | 18.05 | + |
| Dimt1 | 18.93 | 21.23 | 19.23 | 9.28 | 10.79 | 10.62 | 7.47 | 11.53 | 10.1 | 8.18 | 8.48 | 8.03 | + |
| Smc6 | 14.2 | 15.53 | 15.06 | 11.5 | 13.35 | 12.04 | 11.12 | 11.67 | 12.02 | 11.22 | 10.77 | 11.91 | + |
| Klrk1 | 68.74 | 70.35 | 66.39 | 76.65 | 76.57 | 85.53 | 105.29 | 109.65 | 107.61 | 91.82 | 81.14 | 88.11 | + |
| Ndufs7 | 153.33 | 164.74 | 152.65 | 129.66 | 132.83 | 127.29 | 129.71 | 136.68 | 133.77 | 110.56 | 116.15 | 118.02 | + |
| Mr1 | 5.39 | 4.55 | 5.42 | 8.89 | 8.73 | 10.04 | 10.54 | 10.35 | 8.83 | 9.8 | 10.32 | 9.63 | + |
| Ndufa9 | 121.78 | 123.19 | 120.24 | 79.7 | 79.63 | 83.79 | 82.94 | 94.5 | 93.54 | 66.38 | 69.42 | 65.08 | + |
| Zfp260 | 6.84 | 6.06 | 5.97 | 12.1 | 11.48 | 11.73 | 12.8 | 11.45 | 10.15 | 16.03 | 14.5 | 16.86 | + |
| Isca1 | 34.07 | 33 | 34.47 | 25.78 | 28.46 | 27.91 | 25.11 | 28.92 | 26.17 | 24.14 | 24.91 | 28.81 | + |
| Lrpprc | 30.3 | 32.79 | 33.43 | 14.29 | 14.73 | 12.92 | 10.69 | 16.11 | 15.17 | 9.02 | 8.5 | 10.47 | + |
| Ncaph | 27.32 | 23.48 | 23.76 | 22.31 | 26.6 | 25.51 | 14.78 | 14.56 | 13.58 | 17.03 | 17.89 | 14.74 | + |
| Trim3 | 1.16 | 0.96 | 1.1 | 2.46 | 2.41 | 2.24 | 2.39 | 1.29 | 1.92 | 1.65 | 1.9 | 2.3 | + |
| Fastkd2 | 18.72 | 19.7 | 21.15 | 7.43 | 9 | 7.48 | 5.65 | 8.21 | 6.48 | 5.55 | 5.25 | 4.48 | + |
| 2610005L07Rik | 8.82 | 9.46 | 9.12 | 15.04 | 13.93 | 15.25 | 13.83 | 16.75 | 14.71 | 15.51 | 16.67 | 14.37 | + |
| Tuba1b | 1852.55 | 1808.31 | 1739.27 | 610.11 | 743.22 | 740.01 | 539.56 | 562.46 | 560.78 | 389.17 | 419.73 | 420.69 | + |
| Ahcy | 172.89 | 175.87 | 175.2 | 250.11 | 250.77 | 244.08 | 180.46 | 191.82 | 173.89 | 345.22 | 353.17 | 304.03 | + |
| Sms | 67.88 | 61.04 | 60.63 | 42.41 | 45.99 | 45.99 | 40.98 | 48.96 | 43.5 | 48.94 | 47.19 | 42.62 | + |
| Dhx37 | 10.96 | 11.29 | 11.37 | 5.89 | 5.3 | 5.39 | 4.98 | 6.44 | 5.63 | 3.84 | 5.05 | 3.97 | + |
| Tnp2 | 9.65 | 12.91 | 8.55 | 11.61 | 14.53 | 12.64 | 3.97 | 3.31 | 6.54 | 3.43 | 3.25 | 1.99 | + |
| Rps19 | 1724.26 | 1762.69 | 1719.95 | 2917.53 | 2887.37 | 2999.74 | 2933.14 | 2633.87 | 2700.52 | 3120.77 | 3420.63 | 3159.77 | + |
| Timm9 | 55.38 | 52.72 | 53 | 29.72 | 29.91 | 30.84 | 23.64 | 31.34 | 31.19 | 20.67 | 25.9 | 21.36 | + |
| Nfyc | 32.54 | 33.31 | 28.26 | 14.84 | 14.29 | 12.89 | 9.49 | 11.99 | 11.28 | 9.68 | 8.02 | 10.56 | + |
| Zfp467 | 0.53 | 0.22 | 0.41 | 2.11 | 1.58 | 1.24 | 2.96 | 1.34 | 1.54 | 1.83 | 1.29 | 1.82 | + |
| Arsg | 0.47 | 0.3 | 0.64 | 2.14 | 1.71 | 1.6 | 0.93 | 0.84 | 1.18 | 1.68 | 1.28 | 1.6 | + |
| Psmb10 | 53.09 | 58.46 | 55.82 | 37.72 | 31.6 | 36.66 | 40.86 | 38.76 | 34.68 | 40.49 | 31.69 | 42.16 | + |
| Dnmt1 | 45.56 | 45.18 | 47.24 | 28.51 | 31.75 | 30.42 | 23.28 | 26.78 | 25.1 | 21.02 | 21.29 | 18.51 | + |
| Traf7 | 57.56 | 53.47 | 55.16 | 41.01 | 40.2 | 42.73 | 40.9 | 39.6 | 42.16 | 37.93 | 40.16 | 43.44 | + |
| Pygb | 44.21 | 42.97 | 46.32 | 67.35 | 57.74 | 59.5 | 62.2 | 60.61 | 59.58 | 63.9 | 70.35 | 63.36 | + |
| Gsdmd | 23.43 | 25.52 | 22.8 | 30.5 | 33.97 | 32.23 | 36.56 | 31.56 | 33.53 | 33.79 | 31.85 | 35.54 | + |
| Zcchc6 | 5.51 | 5.3 | 6.01 | 14.83 | 14.08 | 14.12 | 9.96 | 9.39 | 10.99 | 13.45 | 12.08 | 12.22 | + |
| Trib2 | 0.2 | 0.14 | 0.08 | 1.91 | 0.98 | 0.96 | 4.24 | 1.09 | 2.88 | 2.84 | 3.14 | 3.36 | + |
| Gm5531 | 3.67 | 3.16 | 2.95 | 1.75 | 1.61 | 1.52 | 1.66 | 1.66 | 1.61 | 1.38 | 1.18 | 1.28 | + |
| Vat1 | 2.03 | 1.98 | 2.22 | 2.36 | 2.9 | 3.15 | 3.49 | 3.85 | 3.14 | 2.78 | 3.9 | 3.19 | + |
| 2610318N02Rik | 13.84 | 14.75 | 12.41 | 5.59 | 6.66 | 7 | 4.31 | 5.43 | 5.81 | 3.93 | 4.28 | 3.78 | + |
| Trp73 | 2.62 | 2.67 | 2.73 | 1.6 | 1.82 | 1.82 | 1.96 | 2.07 | 1.27 | 1.2 | 1.69 | 1.71 | + |
| Fip1l1 | 45.63 | 50.32 | 48.45 | 52.48 | 51.65 | 54.07 | 50.99 | 52.53 | 50.72 | 57.27 | 59.23 | 56.34 | + |
| Myef2 | 31.18 | 30.83 | 31.73 | 16.83 | 19.95 | 19.27 | 11.24 | 14.61 | 15.38 | 14.38 | 12.85 | 10.98 | + |
| Casp6 | 31.7 | 29.36 | 41.51 | 57.82 | 73.45 | 75.21 | 56.31 | 55.83 | 58.07 | 49.46 | 51.06 | 50.09 | + |
| Ift80 | 5.4 | 4.69 | 6.13 | 5.39 | 6.66 | 6.21 | 4.41 | 4.31 | 3.91 | 2.8 | 2.89 | 3.62 | + |
| Rnf130 | 0.76 | 1.36 | 0.54 | 3.08 | 3.33 | 2.62 | 3.76 | 2.16 | 3.26 | 4.27 | 2.84 | 3.73 | + |
| Klhdc1 | 4.25 | 3.79 | 5.24 | 9.37 | 7.85 | 9.23 | 9.49 | 8.23 | 7.57 | 8.34 | 8.16 | 11 | + |
| 1700011J10Rik | 3.83 | 3.49 | 3.68 | 6.77 | 7.56 | 6.15 | 7.19 | 6.09 | 6.03 | 7.88 | 7.34 | 7.83 | + |
| Mtrr | 10.93 | 11.39 | 10.98 | 7.02 | 7.1 | 9.23 | 5.32 | 5.72 | 6.02 | 6.51 | 5.2 | 5.29 | + |
| Oxld1 | 6.2 | 8.19 | 8.41 | 17.4 | 18.26 | 16.3 | 21 | 21.25 | 22.26 | 17.3 | 19.93 | 19.71 | + |
| Uhrf1 | 84.03 | 87.55 | 85.78 | 43.85 | 46.92 | 49.75 | 35.17 | 38.67 | 36.37 | 25.89 | 28.09 | 28.72 | + |
| Rrp15 | 48.1 | 50.5 | 53.13 | 15.94 | 21.51 | 16.72 | 12.95 | 22.23 | 21 | 11.88 | 11.13 | 11.28 | + |
| Pkp3 | 44.8 | 38.52 | 43.54 | 61.65 | 61 | 67.34 | 63.27 | 60.66 | 59.35 | 60.07 | 52.16 | 62.48 | + |
| Pan2 | 3.66 | 3.08 | 3.86 | 6.92 | 7.36 | 7.38 | 7.48 | 6.66 | 6.74 | 8.48 | 8.44 | 8.48 | + |
| Ccdc90b | 8.12 | 7.8 | 6.64 | 11.6 | 11.38 | 11.27 | 10.3 | 11.37 | 9.85 | 11.68 | 9.46 | 11.7 | + |
| Gps1 | 130.73 | 125.48 | 128.27 | 70.3 | 81.01 | 72.75 | 59.67 | 83 | 69.91 | 53.69 | 56.17 | 44.04 | + |
| Sdc3 | 2.38 | 2.22 | 2.38 | 3.92 | 4.27 | 4.16 | 6.96 | 5.48 | 6.29 | 5.53 | 5.84 | 6.67 | + |
| Prpf38b | 26.98 | 29.33 | 30.97 | 24.67 | 26.03 | 23.7 | 23.5 | 26.37 | 25.47 | 23.16 | 25.34 | 23.32 | + |
| Aatf | 35.43 | 39.4 | 34.54 | 24.19 | 22.19 | 24.16 | 17.65 | 23.55 | 24.21 | 20.42 | 20.82 | 21.4 | + |
| Ptger4 | 16.72 | 16.22 | 17.64 | 13.17 | 17.58 | 17.35 | 7.7 | 9.68 | 12.36 | 5.67 | 5.46 | 5.8 | + |
| Agpat4 | 34.24 | 29.49 | 33.33 | 30.44 | 32.62 | 34.06 | 41.28 | 34.84 | 38.69 | 43.06 | 45.61 | 43.62 | + |
| Tubb4a | 1.33 | 1.12 | 1.64 | 9.39 | 7.95 | 8.64 | 12.03 | 6.17 | 7.01 | 8.73 | 10.79 | 11.54 | + |
| Dnajc4 | 9.38 | 9.14 | 9.78 | 11.73 | 13.82 | 14.4 | 16.56 | 14.89 | 17.56 | 15.87 | 12.55 | 15.23 | + |
| Pdss1 | 18.97 | 15.24 | 15.64 | 4.96 | 6.13 | 5.78 | 3.3 | 4.36 | 4.96 | 2.67 | 2.93 | 2.03 | + |
| Aimp2 | 76.33 | 75.5 | 76.44 | 36.21 | 44.18 | 41.87 | 42.29 | 48.07 | 46.63 | 32.28 | 35.84 | 34.55 | + |
| Noa1 | 36.69 | 38.64 | 40.92 | 20.53 | 20.87 | 24.64 | 23.06 | 27.57 | 22.88 | 22.18 | 23.12 | 21.13 | + |
| Rps20 | 1267.68 | 1302.96 | 1271.25 | 1884.53 | 1913.11 | 1982 | 1914.91 | 1771.26 | 1803.42 | 2015.64 | 2156.32 | 1947.72 | + |
| Adamtsl5 | 1.81 | 1.47 | 1.62 | 4.51 | 6.22 | 5.55 | 7.74 | 7.14 | 7.38 | 7.96 | 9.08 | 10.67 | + |
| Creb1 | 3.4 | 3.41 | 3.33 | 3.24 | 3.29 | 2.78 | 3.9 | 3.14 | 3.62 | 4.29 | 4.12 | 4.4 | + |
| Ulk3 | 7.92 | 7.15 | 6.07 | 11.33 | 9.93 | 10.86 | 9.86 | 11.04 | 10.86 | 11.63 | 9.97 | 12.18 | + |
| 7-Sep | 135.45 | 130.11 | 133.6 | 100.1 | 105.55 | 108.74 | 91.19 | 103.62 | 101.39 | 93.82 | 94.98 | 89.14 | + |
| Vamp7 | 19.33 | 22.64 | 21.44 | 14.87 | 17.69 | 15.02 | 17.78 | 16.13 | 17.28 | 14.58 | 17.05 | 15.49 | + |
| Snhg3 | 184.48 | 180.81 | 202.63 | 89.66 | 89.63 | 92.47 | 85.68 | 96.37 | 77.1 | 81.4 | 94.28 | 77.14 | + |
| Traip | 11.6 | 10.52 | 10.99 | 5.09 | 5.86 | 5.55 | 3.22 | 3.57 | 3.34 | 3.05 | 2.45 | 2.93 | + |
| Nop2 | 93.18 | 92.06 | 95.81 | 38.89 | 40.1 | 41.25 | 31.35 | 48.79 | 45.98 | 26.65 | 26.26 | 27.47 | + |
| BC026585 | 7.31 | 5.1 | 5.75 | 8.43 | 11.04 | 10.19 | 13.33 | 15.08 | 13.11 | 9.42 | 12.14 | 9.35 | + |
| Xpo7 | 3.73 | 3.83 | 3.49 | 2.92 | 2.84 | 2.64 | 2.63 | 2.83 | 2.77 | 2.8 | 2.8 | 2.5 | + |
| Gem | 46.2 | 46.97 | 36.91 | 21.84 | 29.69 | 40.37 | 81.98 | 96.05 | 91.03 | 32.11 | 27.64 | 34.56 | + |
| Inha | 0.27 | 0.44 | 0.22 | 2.29 | 2.09 | 1.96 | 1.73 | 2.25 | 1.43 | 3.28 | 1.6 | 1.97 | + |
| Sgpl1 | 11.37 | 10.91 | 13.91 | 24.11 | 21.38 | 24.81 | 23.15 | 23.44 | 22.6 | 28.4 | 35.37 | 29.47 | + |
| Susd1 | 1.32 | 1.13 | 0.89 | 1.32 | 0.93 | 0.6 | 1.31 | 1.66 | 1.84 | 0.43 | 0.58 | 0.52 | + |
| Bbip1 | 14.91 | 14.6 | 15.66 | 10.45 | 10.42 | 10.71 | 12.42 | 11.26 | 11.81 | 9.36 | 9.84 | 11.7 | + |
| Pdhx | 12.59 | 10.74 | 11.18 | 6.63 | 4.92 | 5.58 | 5.96 | 5.79 | 5.52 | 3.53 | 4.83 | 4.79 | + |
| Nt5e | 3.46 | 3.71 | 3.47 | 23.96 | 14.45 | 14.33 | 8.86 | 6.32 | 6.56 | 7.53 | 7.35 | 9.68 | + |
| Elof1 | 54.38 | 57.61 | 50.06 | 32.9 | 35.86 | 33.67 | 38.42 | 32.69 | 33.42 | 35.59 | 29.26 | 34.21 | + |
| Fam136a | 107.43 | 107.71 | 101.12 | 50.22 | 50.36 | 49.15 | 41.36 | 47.71 | 49.87 | 33.88 | 27.02 | 32.02 | + |
| Thoc6 | 24.38 | 22.93 | 22.49 | 12.13 | 15.57 | 13.02 | 12.45 | 15.81 | 12.24 | 9.15 | 10.73 | 11.22 | + |
| Hook2 | 13.33 | 12.04 | 14.73 | 17.83 | 18.15 | 20.5 | 22.63 | 20 | 20.16 | 17.97 | 22.9 | 19.43 | + |
| Fam71f2 | 1.31 | 0.74 | 1.2 | 4.94 | 4.51 | 5.91 | 7.38 | 6.14 | 7.48 | 7.51 | 9.49 | 11.37 | + |
| Sfxn1 | 80.08 | 90.59 | 83.6 | 47.64 | 50.71 | 48.33 | 40.61 | 51.85 | 48.13 | 33.59 | 37.2 | 33.46 | + |
| Pkib | 6.15 | 5.16 | 5.7 | 3.82 | 5.47 | 5.2 | 0.87 | 1.81 | 1.48 | 2.04 | 2.61 | 1.72 | + |
| Elk3 | 7.95 | 7.87 | 9.09 | 4.96 | 6.34 | 5.47 | 7.68 | 7.8 | 7.62 | 5.51 | 6.51 | 6.66 | + |
| Mrps18b | 148.17 | 151.91 | 146.18 | 76.23 | 86.32 | 87.34 | 78.68 | 106.98 | 93.65 | 70.08 | 64.67 | 62.41 | + |
| Toe1 | 18.61 | 19.21 | 20.86 | 12.65 | 8.96 | 11.35 | 7.32 | 10.11 | 9.29 | 8.73 | 7.42 | 10.01 | + |
| Spr | 15.27 | 13.85 | 16.55 | 10.05 | 8.96 | 8.31 | 9.61 | 10.58 | 11.16 | 8.82 | 7.49 | 7.95 | + |
| Clhc1 | 11.69 | 12.37 | 10.77 | 3.58 | 6.09 | 6.78 | 2.44 | 2.91 | 4.39 | 2.66 | 2.43 | 3.49 | + |
| Pcid2 | 22.9 | 21.84 | 21.07 | 12.76 | 13.44 | 13.61 | 15.06 | 15.57 | 13.38 | 10.86 | 12.16 | 11.32 | + |
| Nutf2-ps1 | 182.39 | 185.93 | 188.77 | 80.06 | 91.43 | 87.53 | 65.64 | 95.21 | 83.58 | 53.11 | 64.78 | 53.56 | + |
| Umps | 49.11 | 49.52 | 49.24 | 22.05 | 19.49 | 23.28 | 17.78 | 23.32 | 22.3 | 13.35 | 16.72 | 16.39 | + |
| Rpl10a | 1323.86 | 1315.62 | 1297.92 | 2236.6 | 2185.49 | 2322.73 | 2207.78 | 2081.16 | 2135.25 | 2447.58 | 2640.71 | 2384.77 | + |
| Qtrtd1 | 5.21 | 4.68 | 6.12 | 2.71 | 3.68 | 3.82 | 2.24 | 2.62 | 2.75 | 2.88 | 2.18 | 2.41 | + |
| Sp140 | 21.27 | 18.89 | 19.23 | 47.77 | 39.51 | 44.63 | 40.63 | 38.87 | 38.32 | 47.34 | 42.64 | 49.89 | + |
| Dap | 64.53 | 67.7 | 57.23 | 145.38 | 122.96 | 135.6 | 217.91 | 141.46 | 158.24 | 222.95 | 202.91 | 221.85 | + |
| Grina | 34.11 | 31.48 | 27.33 | 117.35 | 104.75 | 107.28 | 127.14 | 82.43 | 89.27 | 165.48 | 161.77 | 182.6 | + |
| Pfdn5 | 122.09 | 131.79 | 122.57 | 217.95 | 207.25 | 221.48 | 267.97 | 207.3 | 219.82 | 294.38 | 304.84 | 294.2 | + |
| Pim2 | 20.8 | 17.82 | 19.21 | 41.71 | 39.94 | 38.78 | 19.2 | 19.4 | 20.61 | 15.39 | 14.65 | 14.83 | + |
| Dicer1 | 2.89 | 2.78 | 3.01 | 1.86 | 1.68 | 2.07 | 1.85 | 1.8 | 2.25 | 2.37 | 1.82 | 1.86 | + |
| Srpr | 64.66 | 66.59 | 61.91 | 40.95 | 41.83 | 41.76 | 44.79 | 48.73 | 44 | 39.66 | 51.88 | 39.51 | + |
| Atxn2l | 20.36 | 21.87 | 24.56 | 35.58 | 37.81 | 37.03 | 33.65 | 44.99 | 41.9 | 41.11 | 41.53 | 33.67 | + |
| Ap2s1 | 144 | 136.94 | 144.28 | 111.12 | 115.37 | 116.76 | 117.16 | 111.79 | 119.34 | 104.38 | 110.29 | 115.21 | + |
| Polr2e | 192.22 | 185.3 | 192.46 | 149.54 | 148 | 158.04 | 156.03 | 164.49 | 158.54 | 149.32 | 168.02 | 161.66 | + |
| Spcs1 | 129.4 | 133.27 | 125.12 | 88.83 | 91.03 | 93.94 | 82.43 | 97.43 | 92.48 | 82.11 | 71.77 | 72.88 | + |
| Heatr2 | 2.39 | 2.74 | 2.83 | 4.61 | 4.6 | 5.27 | 7.68 | 5.02 | 7.14 | 6.6 | 4.35 | 5.69 | + |
| Pdia4 | 96.99 | 93.79 | 95.36 | 50.98 | 49.56 | 56.11 | 49.14 | 57.08 | 53.72 | 51.63 | 50.5 | 48.79 | + |
| Ttc1 | 42.78 | 40.19 | 42.6 | 28.16 | 32.91 | 30.82 | 24.84 | 28.1 | 31.17 | 25.13 | 25.64 | 30.17 | + |
| Slc7a11 | 5.1 | 4.76 | 4.58 | 1.47 | 1.97 | 2.18 | 1.66 | 1.85 | 1.74 | 1.07 | 1.38 | 1.14 | + |
| Pias3 | 3.77 | 4.42 | 3.9 | 8.58 | 8.41 | 9.27 | 8.1 | 7.09 | 7.65 | 7.63 | 8.66 | 10.44 | + |
| Gtpbp2 | 17.88 | 17.19 | 16.04 | 36.83 | 40.37 | 39.42 | 33.53 | 37.76 | 37.91 | 29.95 | 35.59 | 30.69 | + |
| Egr1 | 10.46 | 13.49 | 14.56 | 9.83 | 11.32 | 14.2 | 6.87 | 6.12 | 7.49 | 3.78 | 3.51 | 4.03 | + |
| Chd7 | 6.31 | 7.13 | 8.43 | 13.84 | 13.6 | 13.61 | 14.73 | 12.16 | 14.79 | 17.62 | 15.3 | 16.53 | + |
| Dynlt1c | 177.61 | 184.58 | 182.31 | 131.72 | 157.47 | 129.59 | 147.52 | 152.37 | 147.44 | 153.05 | 139.15 | 139.55 | + |
| Rnf141 | 13.97 | 11.85 | 9.88 | 12.68 | 10.43 | 11.72 | 10.03 | 10.06 | 9.35 | 7.03 | 7.56 | 5.63 | + |
| Dck | 26.27 | 28.33 | 27.61 | 15.26 | 15.48 | 15.56 | 17.89 | 13.98 | 18.27 | 15.65 | 17.27 | 17.63 | + |
| Erh | 458.98 | 486.12 | 486.66 | 192.18 | 197.75 | 197.94 | 166.96 | 211.21 | 196.01 | 141.47 | 138.2 | 143.32 | + |
| Zmynd8 | 79.47 | 95.17 | 88.01 | 35.06 | 39.27 | 38.89 | 39.04 | 44.46 | 41.71 | 36.02 | 40.24 | 37.45 | + |
| Snrpc | 71.23 | 78.86 | 75.14 | 35.84 | 43.21 | 34.17 | 33.72 | 42.04 | 40.82 | 29.66 | 29.92 | 28.9 | + |
| Mrpl24 | 61.61 | 59.32 | 56.81 | 77.14 | 73.68 | 80.09 | 77.34 | 79.45 | 76.3 | 73.01 | 82.07 | 78.92 | + |
| Mfn2 | 20.07 | 17.63 | 16.78 | 14.57 | 14.09 | 14.92 | 13.58 | 13.8 | 13.77 | 14.64 | 11.88 | 13.06 | + |
| Igf1r | 0.1 | 0.09 | 0.07 | 0.56 | 0.38 | 0.37 | 0.32 | 0.18 | 0.28 | 0.42 | 0.36 | 0.46 | + |
| Tmx1 | 78.91 | 78.79 | 80.08 | 46.72 | 50.75 | 49.38 | 41.81 | 50.96 | 47.01 | 38.02 | 40.26 | 43.31 | + |
| Kbtbd3 | 1.03 | 1.34 | 1.46 | 2.7 | 2.91 | 3.22 | 2.8 | 2.35 | 2.13 | 2.89 | 3.67 | 2.96 | + |
| Rpl17 | 1553.64 | 1581.64 | 1563.8 | 2579.01 | 2651.23 | 2721.98 | 2525.8 | 2411.04 | 2440.97 | 2706.69 | 2953.52 | 2746.21 | + |
| Uchl5 | 143.76 | 138 | 142.52 | 71.66 | 77.71 | 78.07 | 63.63 | 89.86 | 79.49 | 60.51 | 61.26 | 52.7 | + |
| Adssl1 | 0.89 | 2.11 | 1.06 | 12.03 | 7.6 | 9.65 | 6.48 | 4.92 | 7.28 | 13.7 | 11.77 | 10.14 | + |
| Gm12942 | 8.67 | 7.48 | 8.5 | 17.16 | 14.77 | 16.17 | 22.8 | 22.29 | 18.11 | 16.55 | 13.76 | 21.7 | + |
| Srsf7 | 136.3 | 142.95 | 134.78 | 61.35 | 63.15 | 61.82 | 37.73 | 50.43 | 52.37 | 36.25 | 37.05 | 36.36 | + |
| Ccnl1 | 22.27 | 24.01 | 24.07 | 27.37 | 22.2 | 23.16 | 22.32 | 23.38 | 21.81 | 30.96 | 30.65 | 31.21 | + |

| | | | | | | | | | | | | |
|---|---|---|---|---|---|---|---|---|---|---|---|---|
| Dera | 38.25 | 37.2 | 35.15 | 19.81 | 22.71 | 19.01 | 25.84 | 25.57 | 23.49 | 23.38 | 19.39 | 20.25 | + |
| Cldn25 | 36.99 | 36.26 | 32.69 | 25.31 | 23.26 | 21.6 | 12.19 | 9.22 | 10.83 | 8.74 | 8.9 | 9.64 | + |
| Zfp672 | 24.17 | 22.78 | 28.96 | 23.53 | 19.36 | 19.26 | 15.99 | 13.56 | 15.16 | 17.17 | 14.86 | 16.67 | + |
| Tkt | 243.72 | 240.17 | 243.57 | 181.49 | 172.73 | 179.33 | 197.53 | 184.13 | 189.51 | 177.01 | 185.14 | 179.86 | + |
| Rnf114 | 50.83 | 50.41 | 51.09 | 111.9 | 96.81 | 105.3 | 126.48 | 96.44 | 99.91 | 156.4 | 162.48 | 164.81 | + |
| Uso1 | 4.16 | 5.08 | 4.64 | 2.71 | 2.72 | 3.21 | 3.11 | 3.33 | 3.33 | 3.54 | 2.8 | 2.53 | + |
| 1110007C09Rik | 13.56 | 14.26 | 16.87 | 51.33 | 51.01 | 53.96 | 50.31 | 34.7 | 42.11 | 63.46 | 75.59 | 63.93 | + |
| Txndc9 | 20.12 | 17.84 | 17.72 | 14.58 | 14.15 | 14.61 | 13.55 | 13.94 | 12.57 | 12.42 | 11.71 | 13.66 | + |
| Api5 | 93.42 | 93.08 | 95.82 | 62.86 | 65.65 | 64.35 | 53.94 | 65.36 | 60.89 | 55.08 | 57.8 | 55.7 | + |
| 0610009D07Rik | 90.85 | 95.89 | 92.97 | 56.18 | 56.87 | 59.21 | 56.31 | 58.18 | 59.68 | 46.98 | 47.38 | 48.2 | + |
| Tmem134 | 30.92 | 37.63 | 36.54 | 48.78 | 46.55 | 50.18 | 53.84 | 52.15 | 52.59 | 50.97 | 49.98 | 48.29 | + |
| Gzma | 145.17 | 254.1 | 85.29 | 2625.29 | 2603.07 | 2252.28 | 1102.59 | 557.79 | 736.2 | 2941.32 | 2380.19 | 3445.02 | + |
| Rbm14 | 33.02 | 38.61 | 35.46 | 20 | 20.56 | 22 | 16.16 | 19.32 | 15.1 | 12.65 | 16.92 | 14.71 | + |
| Acaca | 8.87 | 8.81 | 9.37 | 6.34 | 5.63 | 4.93 | 4.76 | 6.11 | 5.87 | 5.25 | 5.08 | 4.38 | + |
| Cisd1 | 149.73 | 155.81 | 140.09 | 92.02 | 86.87 | 95.56 | 128.92 | 121.14 | 124.68 | 77.91 | 95.1 | 91.74 | + |
| Timm8a1 | 88.14 | 84.4 | 82.34 | 34.09 | 38.89 | 37.57 | 27.2 | 41.52 | 34.74 | 22.44 | 21.93 | 21.29 | + |
| 5830454E08Rik | 0.54 | 1.33 | 0.87 | 4.51 | 2.52 | 2.75 | 3.71 | 3.16 | 2.74 | 4 | 4.48 | 5.97 | + |
| Cdc42 | 425.7 | 427.77 | 425.7 | 343.85 | 354 | 371.99 | 384.87 | 376.77 | 386.67 | 343.97 | 360.77 | 352.98 | + |
| Rpe | 35.11 | 38.95 | 38.78 | 29.51 | 29.83 | 33.32 | 33.1 | 34.23 | 34.02 | 30.11 | 30.72 | 29.81 | + |
| Trim30d | 8.55 | 7.01 | 8.36 | 6.31 | 6.46 | 5.76 | 3.23 | 4.08 | 3.91 | 5.05 | 4.95 | 5.92 | + |
| Nle1 | 19.09 | 18.44 | 19.8 | 9.59 | 9.4 | 8.52 | 7.54 | 10 | 9.78 | 6.96 | 7.15 | 8.91 | + |
| Slc25a3 | 668.55 | 675.67 | 661.84 | 540.67 | 544.81 | 553.36 | 524.09 | 568.59 | 537.13 | 496.2 | 518.54 | 492.27 | + |
| Sumo1 | 180.36 | 184.45 | 182.53 | 142.36 | 134.57 | 137.15 | 143.04 | 148.52 | 150.03 | 125.66 | 137.46 | 135.67 | + |
| Dnase1l1 | 5.61 | 4.67 | 5.55 | 9.49 | 9.97 | 7.56 | 8.72 | 9.13 | 8.1 | 8.27 | 7.28 | 10.86 | + |
| Ube2j1 | 26.89 | 28.72 | 28.82 | 22.22 | 22.65 | 22.56 | 21.48 | 24.34 | 20.6 | 20.16 | 21.45 | 17.36 | + |
| Tmem208 | 59.74 | 62.68 | 58.04 | 43.96 | 44.05 | 49.27 | 41.15 | 47.8 | 45.54 | 41.99 | 39.13 | 46.74 | + |
| Pard6a | 10.49 | 9.14 | 9.64 | 13.11 | 14.27 | 13.43 | 16.57 | 16.06 | 15.38 | 14.47 | 16.27 | 15.8 | + |
| Mrps11 | 33.72 | 30.13 | 31.06 | 20.6 | 18.5 | 19.63 | 20.22 | 21.12 | 21.19 | 17.45 | 14.44 | 15.31 | + |
| 9430038I01Rik | 8.41 | 9.35 | 7.86 | 5.05 | 6.09 | 5.75 | 5.51 | 4.7 | 6.8 | 4.35 | 5.04 | 6.13 | + |
| Atp5b | 1252.67 | 1254.97 | 1265.44 | 692.74 | 740.35 | 751.21 | 665.32 | 837.12 | 785.58 | 557.5 | 594.23 | 547.24 | + |
| Osbpl9 | 22.25 | 21.15 | 22.12 | 44.17 | 44.24 | 41.7 | 31.98 | 38.49 | 33.72 | 39.63 | 39.21 | 34.69 | + |
| Mat2a | 212.34 | 227.62 | 221.64 | 86.63 | 93.56 | 91.63 | 51.7 | 70.37 | 66.56 | 58.12 | 46.24 | 52.91 | + |
| Cdc14b | 3.46 | 2.99 | 3.66 | 4.72 | 5.76 | 5.81 | 8.22 | 6.03 | 7.12 | 6.96 | 8.25 | 7.08 | + |
| Sh3yl1 | 2.01 | 1.78 | 1.49 | 3.71 | 4.03 | 4.3 | 4.2 | 2.44 | 2.91 | 4.94 | 3.76 | 3.82 | + |
| Ywhab | 123.77 | 122.92 | 120.54 | 51.36 | 54.89 | 59.13 | 53.36 | 69.42 | 60.03 | 39.65 | 36.49 | 38.55 | + |
| Ifih1 | 4.54 | 4.44 | 3.86 | 3.86 | 3.79 | 4.46 | 2.46 | 3.11 | 2.86 | 3.03 | 3.21 | 2.95 | + |
| Map2k6 | 3.15 | 3.33 | 4.43 | 3.95 | 4.54 | 4.94 | 11.55 | 7.28 | 8.66 | 6.07 | 6.15 | 6.17 | + |
| Galnt6 | 7.96 | 8.52 | 9.03 | 19.94 | 17.7 | 18.91 | 19.31 | 14.71 | 14.91 | 23.7 | 25.13 | 25.24 | + |
| Prpf3 | 24.82 | 23.71 | 22.62 | 10.99 | 13.16 | 12.46 | 9.82 | 12.56 | 11.42 | 13 | 12.07 | 7.45 | + |
| Lgals1 | 1434.73 | 1421.36 | 1437.85 | 826.61 | 895.03 | 904.27 | 1242.39 | 1269.76 | 1243.95 | 848.57 | 876.8 | 896.98 | + |
| Tdp1 | 34.12 | 32.9 | 35.48 | 23.9 | 27 | 24.61 | 19.99 | 21.47 | 21.55 | 16.79 | 19.76 | 16.59 | + |
| Eif4e | 163.1 | 155.95 | 161.45 | 85.01 | 94.09 | 93.27 | 82.84 | 111.3 | 103.99 | 71.62 | 83.82 | 71.15 | + |
| Vps37a | 3.22 | 4.35 | 5.17 | 7 | 8.42 | 6.76 | 7.01 | 9.07 | 8.14 | 8.32 | 7.59 | 6.17 | + |
| Pxk | 5.78 | 7.59 | 7.09 | 15.65 | 12.72 | 11.88 | 15.96 | 14.52 | 13.49 | 18.28 | 17.65 | 18.36 | + |
| Abl1 | 2.93 | 2.84 | 2.96 | 5.21 | 4.52 | 3.99 | 3.81 | 3.92 | 3.56 | 3.95 | 3.53 | 4.55 | + |
| Icos | 32.17 | 36.55 | 36.63 | 149.82 | 150.97 | 153.03 | 55.28 | 38.92 | 47.3 | 115.3 | 122.81 | 137.07 | + |
| Pabpc1 | 185.86 | 200.91 | 203.78 | 267.04 | 274.05 | 273.27 | 304.69 | 305.36 | 303.29 | 283.54 | 335.34 | 282.32 | + |
| Ssr1 | 53.3 | 54 | 54.28 | 37.23 | 38.15 | 39.8 | 34.59 | 36.98 | 38.65 | 31.14 | 35.49 | 32.54 | + |
| Pex2 | 20.14 | 21.23 | 21.33 | 16.21 | 17.08 | 16.82 | 17.55 | 17.57 | 15.5 | 14.37 | 14.65 | 14.43 | + |
| Dld | 68.24 | 68.83 | 69.94 | 44.83 | 48.01 | 49.19 | 43.3 | 49.99 | 48.71 | 37.11 | 40.35 | 34.71 | + |
| Alpk2 | 0.21 | 0.1 | 0.17 | 1.41 | 1.82 | 1.87 | 0.33 | 0.45 | 0.27 | 1.71 | 1.79 | 1.72 | + |
| Eif1 | 371.1 | 421.42 | 413.89 | 507.91 | 503.42 | 507.12 | 554.37 | 558.1 | 541.13 | 545.44 | 597.9 | 545.92 | + |
| Phf19 | 3.58 | 2.74 | 3.16 | 1.18 | 1.85 | 1.65 | 0.89 | 1.3 | 1.03 | 1.12 | 0.94 | 1.11 | + |
| Med28 | 27.3 | 25.78 | 27.14 | 22.57 | 23.67 | 22.17 | 23.29 | 24.26 | 22.76 | 21.44 | 20.71 | 22.2 | + |
| Cnot10 | 32.64 | 30.94 | 32.37 | 24.32 | 23.23 | 22.32 | 20.51 | 21.9 | 23.65 | 20.27 | 19.91 | 18.45 | + |
| Tagln2 | 607.88 | 606.55 | 625.49 | 468.43 | 469.66 | 478.63 | 557.06 | 513.64 | 534.47 | 492.04 | 481.02 | 507.76 | + |
| Wdr75 | 77.52 | 83.4 | 81.18 | 36.9 | 43.16 | 41.1 | 34.85 | 51.66 | 41.95 | 28.56 | 33.36 | 27.63 | + |
| Pdhb | 105.8 | 105.73 | 103.47 | 75.46 | 76.04 | 72.49 | 65.59 | 75.33 | 70.99 | 59.06 | 68.72 | 65.24 | + |
| Tbc1d7 | 26.39 | 29.43 | 25.76 | 16.52 | 15.43 | 14.44 | 25.58 | 21.49 | 23.6 | 17.46 | 18.03 | 18.79 | + |
| Gabarapl2 | 98.28 | 101.86 | 95.87 | 73.2 | 71.52 | 79.45 | 77.65 | 85.11 | 83.27 | 79.15 | 68.67 | 76.07 | + |
| Ccr4 | 12.6 | 11.05 | 15.16 | 9.48 | 8.8 | 7.93 | 5.27 | 6.81 | 7.14 | 2.58 | 2.36 | 2.85 | + |
| Mdh2 | 460.86 | 451.79 | 456.54 | 283.72 | 286.83 | 291.52 | 315.72 | 345.35 | 332.96 | 246.82 | 246.69 | 246.46 | + |
| Epb4.1l2 | 5.94 | 6.56 | 6.47 | 10.23 | 12.88 | 10.72 | 8.34 | 10.33 | 9.93 | 9.68 | 11.25 | 8.24 | + |
| Gale | 66.19 | 57.36 | 58.57 | 56.14 | 55.97 | 58.5 | 62.1 | 58.01 | 57.2 | 34.87 | 42.18 | 43.62 | + |
| Cops6 | 238.56 | 235.82 | 242.3 | 156.68 | 169.84 | 162.26 | 150.37 | 178.94 | 163.46 | 127.43 | 127.38 | 134.27 | + |
| Smtn | 2.43 | 1.69 | 1.57 | 4.3 | 5.01 | 5.01 | 7.61 | 6.27 | 6.26 | 3.72 | 4.36 | 5.36 | + |
| Gbp9 | 4.84 | 3.86 | 4.11 | 9.33 | 9.21 | 9.87 | 7.18 | 6.1 | 7.17 | 9.75 | 7.03 | 8.28 | + |
| Cyp51 | 68.13 | 71.66 | 69.39 | 52.79 | 54.93 | 58.2 | 50.1 | 52.78 | 51.53 | 42.22 | 46.69 | 45.16 | + |
| Raver1 | 24.95 | 28.5 | 27.1 | 17.37 | 19.36 | 18.99 | 14.96 | 15.53 | 16.41 | 15.91 | 12.18 | 15.05 | + |
| Rpsa | 1455.39 | 1486.08 | 1506.05 | 2082.83 | 1979.43 | 2020.24 | 1922.61 | 1894.3 | 1891.35 | 2061.88 | 2217.91 | 2088.08 | + |
| Ogdh | 50.38 | 51.87 | 51.77 | 29.66 | 32.07 | 32.15 | 31.58 | 35.29 | 33.05 | 29.09 | 26.38 | 27.51 | + |
| Zfp944 | 9.44 | 8.54 | 8.83 | 8.85 | 9.42 | 8.21 | 8.9 | 8.86 | 9.23 | 6.2 | 5.54 | 6.48 | + |
| Swap70 | 8.9 | 8.65 | 7.62 | 7.57 | 8.59 | 9.25 | 9.73 | 9.76 | 10.89 | 6.69 | 6.63 | 6.11 | + |
| Itgav | 5.39 | 4.97 | 6.82 | 14.27 | 15.92 | 15.52 | 6.11 | 7.77 | 8.2 | 11.86 | 12.07 | 10.83 | + |
| Araf | 38.5 | 35.84 | 35.82 | 27.16 | 28.26 | 30.76 | 25.49 | 29.84 | 30.3 | 27.82 | 25.58 | 25.09 | + |
| Wdr61 | 98.17 | 102.28 | 100.91 | 51.71 | 56.37 | 57.74 | 43.69 | 55.96 | 50.72 | 42.71 | 44.1 | 37.63 | + |
| Trp53inp1 | 3.83 | 2.9 | 4.21 | 10.86 | 9.97 | 10.35 | 14.17 | 9.85 | 12.27 | 17.29 | 16.15 | 16.37 | + |
| Pdcd6ip | 59.4 | 57.77 | 57.97 | 59.38 | 58.96 | 60.95 | 75.68 | 68.23 | 70.88 | 73.8 | 79.56 | 74.37 | + |
| 0610030E20Rik | 4 | 3.98 | 3.85 | 6.46 | 6.6 | 6.95 | 6.45 | 7.42 | 7.95 | 8.63 | 8.56 | 7.74 | + |
| Nsf | 21.12 | 23.23 | 24.01 | 16.43 | 17.97 | 20.16 | 13.89 | 15.16 | 14.98 | 11.81 | 15.27 | 13.78 | + |
| Ints12 | 8.95 | 8.28 | 8.24 | 12.01 | 12.05 | 11.41 | 16.59 | 15.49 | 15.64 | 14.38 | 18.19 | 18 | + |
| Klhl26 | 6.92 | 5.49 | 6.41 | 9.98 | 10.66 | 10.57 | 10.34 | 10.73 | 10.02 | 11.2 | 11.11 | 10 | + |
| Tm9sf2 | 87.94 | 92.1 | 88.25 | 78.95 | 82.22 | 79.54 | 67.5 | 73.46 | 66.74 | 74.37 | 73.34 | 73.36 | + |
| Cd247 | 101.44 | 99.74 | 98.73 | 83.3 | 86.2 | 81.46 | 114.17 | 103.8 | 108.38 | 124.23 | 114.47 | 126.61 | + |
| C330027C09Rik | 10.33 | 10.32 | 10.39 | 4.63 | 5.37 | 5.5 | 2.75 | 3.06 | 3.82 | 1.99 | 2.85 | 2.07 | + |
| Rap2c | 8.88 | 10.47 | 12.18 | 6.61 | 6.94 | 6.17 | 9.57 | 8.49 | 9.01 | 7.12 | 6.99 | 7.39 | + |
| Taldo1 | 156.07 | 162.25 | 164.65 | 105.9 | 116.27 | 119.7 | 126.51 | 138.23 | 140.48 | 118.79 | 115.09 | 111.96 | + |
| Skap2 | 16.89 | 16.4 | 15.41 | 21.4 | 27.81 | 23.54 | 23.18 | 23.36 | 26.9 | 25.65 | 23.75 | 25.67 | + |
| Riok1 | 25.44 | 24.06 | 24.2 | 17.32 | 16.21 | 18.68 | 18.14 | 18.65 | 19.11 | 16.81 | 14.8 | 17.27 | + |
| Nr1h2 | 29.72 | 30.95 | 29.55 | 41.73 | 38.77 | 43.28 | 44.15 | 39.95 | 39.41 | 43.68 | 41.92 | 41.54 | + |
| Fubp1 | 47.98 | 47.27 | 47.4 | 23.56 | 25.38 | 26.37 | 20.95 | 25.17 | 25.3 | 17.3 | 18.15 | 16.79 | + |
| Tom1 | 10.81 | 8.96 | 11.29 | 13.88 | 13.22 | 13.81 | 22.9 | 20.76 | 17.34 | 16.39 | 16.63 | 19.54 | + |
| Fgr | 1.37 | 1.72 | 1.1 | 0.79 | 0.65 | 0.61 | 0.74 | 0.9 | 0.63 | 0.79 | 0.5 | 0.81 | + |
| Edc4 | 18.26 | 18.53 | 18.93 | 16.18 | 15.44 | 14.84 | 14.87 | 14.16 | 16.29 | 13.55 | 17.17 | 14.96 | + |
| Anxa4 | 19.9 | 16.79 | 16.28 | 46.48 | 39.02 | 42.34 | 40.2 | 27.86 | 32.84 | 33.01 | 33.38 | 33.3 | + |
| Sfi1 | 4.41 | 3.54 | 4.35 | 9.36 | 9.52 | 8.24 | 9.38 | 7.27 | 7.71 | 7.75 | 8.96 | 8.94 | + |
| Usp36 | 9.02 | 8.47 | 8.93 | 5.65 | 5.51 | 4.92 | 4.71 | 5.82 | 6.26 | 5 | 5.63 | 4.8 | + |
| Phlda1 | 3.65 | 2.65 | 3.05 | 0.36 | 0.61 | 0.33 | 0.94 | 1.95 | 1.1 | 0.56 | 0.44 | 0.21 | + |
| Spint2 | 55.02 | 57.04 | 56.36 | 82.49 | 71.98 | 76.48 | 99.55 | 80.09 | 90.38 | 75.59 | 80.87 | 85.25 | + |
| Eif1ad | 42.67 | 49.98 | 42.21 | 22.38 | 24.08 | 23.61 | 19.36 | 24.38 | 21.08 | 17.56 | 16.79 | 16.17 | + |
| Xrn1 | 2.03 | 2.27 | 1.98 | 2.28 | 2.45 | 2.77 | 2.34 | 2.21 | 2.36 | 3.37 | 3.31 | 2.8 | + |
| Entpd1 | 7.6 | 7.76 | 5.32 | 11.73 | 12.55 | 13.51 | 28.15 | 27.45 | 25.8 | 24.58 | 28.14 | 24.53 | + |
| Rnase4 | 1.7 | 0.95 | 2.11 | 4.23 | 5.22 | 6.58 | 5.97 | 4.2 | 4.75 | 5.52 | 3.97 | 5.65 | + |
| Exosc7 | 109.94 | 114.97 | 104.17 | 56.92 | 56.52 | 60.57 | 43.56 | 62.29 | 57.53 | 41.65 | 39.7 | 46.64 | + |
| Gnaz | 0.05 | 0.07 | 0.07 | 0.52 | 0.62 | 0.58 | 0.98 | 0.77 | 1.12 | 0.59 | 0.56 | 0.62 | + |
| Tceb3 | 36.1 | 33.4 | 34.53 | 26.12 | 21.63 | 22.77 | 18.7 | 21.79 | 23.9 | 24.39 | 23.93 | 23.01 | + |
| Grcc10 | 141.19 | 143.16 | 119.58 | 238.99 | 230.95 | 247.31 | 246.38 | 226.56 | 232.2 | 236.67 | 243.7 | 272.34 | + |
| Pdcd5 | 204.08 | 204.11 | 197.24 | 141.1 | 140.14 | 143.46 | 138.83 | 151.37 | 138.74 | 126.58 | 143.87 | 133.78 | + |
| Slamf6 | 4.77 | 7.24 | 5.33 | 4.66 | 3.12 | 3 | 8.09 | 4.98 | 4.5 | 10.33 | 6.79 | 11.16 | + |
| Hsd17b12 | 62.03 | 58.17 | 54.18 | 33.75 | 35.33 | 32.49 | 38.9 | 46.31 | 44.17 | 25.82 | 29.49 | 30.34 | + |
| Nif3l1 | 24.72 | 27.75 | 26.14 | 13.41 | 14.23 | 15.41 | 11.6 | 15.79 | 16.15 | 11.96 | 10.83 | 11.78 | + |
| Omd | 6.84 | 8.41 | 7.61 | 2.48 | 3.67 | 3.08 | 3.57 | 3.4 | 3.41 | 2.28 | 2.98 | 2.81 | + |
| Trappc9 | 13.28 | 12.59 | 12.23 | 16.87 | 15.85 | 15.64 | 18.06 | 17.33 | 17.62 | 17.6 | 18.88 | 17.53 | + |
| Mrpl21 | 33.84 | 35.1 | 29.32 | 22.94 | 24.35 | 20.7 | 21.49 | 22.32 | 23.2 | 18.29 | 18.89 | 21.67 | + |
| Sgip1 | 0.35 | 0.55 | 0.58 | 2.08 | 1.66 | 1.44 | 1.28 | 1.78 | 1.2 | 2.88 | 2.87 | 2.83 | + |
| Exoc4 | 20.39 | 18.13 | 17.2 | 22.35 | 24.72 | 21.8 | 24.04 | 23.79 | 24.96 | 20.82 | 24.85 | 22.86 | + |
| Hdhd2 | 18 | 19.42 | 19 | 13.8 | 12.12 | 12.43 | 13.39 | 15.34 | 15.61 | 12.22 | 11.08 | 12.18 | + |
| Akr7a5 | 19.3 | 20.45 | 21.49 | 14.96 | 14.55 | 15.12 | 17.85 | 19.25 | 17.64 | 12.84 | 16.42 | 16.68 | + |
| Sigmar1 | 46.72 | 50.98 | 50.16 | 22.29 | 23.38 | 21.04 | 20.12 | 23.48 | 25.46 | 18.27 | 15.22 | 17.43 | + |
| Gm5918 | 2.83 | 2.39 | 3.14 | 6.53 | 4.59 | 8.12 | 5.77 | 8.06 | 7.06 | 5.91 | 6.8 | 6.93 | + |
| Asna1 | 30.34 | 29.12 | 30.55 | 14.57 | 15.74 | 15.22 | 17.66 | 18.7 | 16.06 | 13.19 | 13.23 | 15.18 | + |
| Reep5 | 85.25 | 80.98 | 78.96 | 65.78 | 69.15 | 66.74 | 74.88 | 76.48 | 72.83 | 62.12 | 68.15 | 63.02 | + |
| Kpna3 | 32.37 | 34.44 | 36.43 | 26.04 | 25.47 | 24.06 | 23.93 | 22.41 | 25.46 | 22.57 | 26.6 | 22.29 | + |
| Rab43 | 2.82 | 2.15 | 2.49 | 2.78 | 2.77 | 2.37 | 2.6 | 2.99 | 2.23 | 4.44 | 4.19 | 4.08 | + |
| Tcn2 | 7.01 | 4.86 | 5.46 | 15.12 | 14.27 | 13.19 | 18.55 | 14.63 | 13.52 | 22.88 | 17.65 | 22.86 | + |
| Eif1a | 111.77 | 109.38 | 117.7 | 67.92 | 71.69 | 68.29 | 51.88 | 69.71 | 66.56 | 49.73 | 48.89 | 46.88 | + |
| Fxn | 20.41 | 23.81 | 23.33 | 14.16 | 18.32 | 16.94 | 17.87 | 14.75 | 15.42 | 14.12 | 14.87 | 14.44 | + |
| Grpel2 | 8.85 | 8.43 | 8.91 | 7.14 | 6.81 | 6 | 5.51 | 7.14 | 6.25 | 5.53 | 5.82 | 6.98 | + |
| Gm4827 | 3.04 | 4.25 | 6.03 | 8.22 | 8.59 | 8.22 | 23.64 | 20.31 | 20.28 | 7.23 | 12.07 | 9 | + |
| Dyx1c1 | 3.79 | 3.15 | 3.59 | 9.06 | 9.1 | 7.61 | 9.96 | 8.84 | 9.37 | 8.94 | 9.7 | 8.91 | + |
| Pter | 4.23 | 4.5 | 5 | 1.48 | 1.9 | 2.22 | 1.22 | 1.96 | 2.34 | 1.5 | 1.46 | 1.34 | + |
| Maml3 | 0.62 | 0.65 | 0.56 | 2.7 | 1.44 | 1.79 | 1.83 | 1.48 | 1.91 | 2.72 | 2.02 | 2.58 | + |
| Ppat | 52.56 | 53.99 | 55.27 | 22.16 | 23.61 | 23.21 | 16.72 | 25.61 | 20.6 | 16.49 | 17.82 | 15.56 | + |
| Mrps21 | 114.6 | 109.22 | 94.85 | 86.96 | 86.66 | 86.96 | 72.85 | 83.69 | 73.69 | 77.31 | 74.8 | 71.24 | + |
| Slc17a5 | 1.81 | 1.64 | 1.84 | 2.94 | 3.34 | 2.94 | 3.44 | 3.44 | 3.36 | 3.7 | 3.25 | 4.02 | + |
| Khk | 8.38 | 7.19 | 6.97 | 12.39 | 11.85 | 11.99 | 14.7 | 13.46 | 12.92 | 13.83 | 14.79 | 14.81 | + |
| Rnf13 | 26.61 | 27.12 | 20.57 | 32.89 | 31.54 | 32.26 | 45.43 | 39.79 | 41.44 | 38.4 | 39.75 | 35.92 | + |
| Rfc1 | 22.11 | 19.05 | 19.6 | 13.32 | 15.81 | 15.22 | 11.7 | 11.99 | 12.4 | 10.26 | 13.95 | 9.76 | + |
| Samm50 | 102.74 | 109.04 | 97.5 | 58.78 | 60.85 | 63.31 | 59.59 | 72.46 | 69.79 | 43.83 | 44.61 | 47.79 | + |
| Cwc15 | 121.23 | 121.64 | 117.58 | 97.99 | 96.88 | 88.63 | 103.26 | 100.84 | 102.04 | 99.32 | 106.95 | 100.68 | + |
| Uvrag | 4.18 | 4.11 | 5.71 | 8.46 | 8.28 | 7.55 | 9.59 | 11.29 | 11.66 | 13.38 | 11.96 | 10.76 | + |
| Rpl6 | 1458.77 | 1510.33 | 1491.27 | 2201.16 | 2177.21 | 2306.89 | 2118.24 | 2032.32 | 2082.58 | 2150.94 | 2428.07 | 2144.9 | + |
| Hyal2 | 17.74 | 17.13 | 17.34 | 13.37 | 13.17 | 13.09 | 10.67 | 13 | 11.32 | 10.58 | 10.07 | 10.91 | + |
| Nnat | 0.76 | 0.72 | 0.91 | 2.74 | 2.38 | 2.71 | 2.09 | 1.92 | 1.57 | 2.33 | 3.49 | 3.32 | + |
| Fchsd2 | 3.92 | 3.11 | 3.28 | 7.17 | 6.4 | 6.7 | 8.1 | 6.55 | 6.37 | 7.28 | 9.2 | 7.85 | + |
| Bzrap1 | 2.12 | 2.74 | 1.91 | 8.79 | 6.67 | 7.53 | 32.71 | 19.8 | 29.34 | 11.75 | 11.95 | 11.16 | + |
| Foxn3 | 1.53 | 1.43 | 1.57 | 3.24 | 3.56 | 3.05 | 3.57 | 3.62 | 3.06 | 4.25 | 5.83 | 3.62 | + |
| Ccdc14 | 2.89 | 2.96 | 3.16 | 2.23 | 2.23 | 3 | 2.01 | 2.03 | 2.13 | 1.17 | 1.32 | 1.01 | + |
| Papd4 | 52.78 | 53.4 | 55.14 | 64.32 | 61.55 | 64.55 | 67.07 | 61.1 | 59.75 | 78.42 | 84 | 71.46 | + |
| Mrpl49 | 33.42 | 35.13 | 34.48 | 19.32 | 18.74 | 19.29 | 20.06 | 17.88 | 15.24 | 16.07 | 15.17 | 14.8 | + |
| Gins2 | 54.99 | 48.59 | 52.29 | 25.87 | 31.38 | 33.11 | 20.13 | 24.09 | 21.31 | 15.92 | 19.16 | 15.65 | + |

| | | | | | | | | | | | | | |
|---|---|---|---|---|---|---|---|---|---|---|---|---|---|
| Prdx1 | 442.32 | 484.75 | 444.73 | 238.61 | 235.73 | 240.54 | 190.23 | 209.8 | 204.95 | 158.4 | 158.07 | 175.84 | + |
| Usp1 | 6.19 | 7.65 | 6.08 | 4.36 | 3.78 | 4.4 | 3.51 | 3.52 | 3.35 | 3.26 | 3.3 | 4.14 | + |
| Izumo4 | 4.93 | 3.91 | 4.61 | 17.78 | 14.71 | 12.78 | 21.69 | 16.53 | 19.28 | 18.48 | 12.99 | 16.41 | + |
| 3110001D03Rik | 42.14 | 44.97 | 37.45 | 34.05 | 33.39 | 32.77 | 31.28 | 32.54 | 33.22 | 27.66 | 35.17 | 29.94 | + |
| Dnm1 | 0.11 | 0.08 | 0.06 | 0.7 | 0.48 | 0.54 | 1.26 | 0.79 | 1.04 | 1.6 | 1.12 | 1.61 | + |
| Set | 222.95 | 238.86 | 237.32 | 103.84 | 107.87 | 111.57 | 74.69 | 104.23 | 103.81 | 71.98 | 71.23 | 75.96 | + |
| Actl6a | 74.38 | 75.65 | 72.55 | 43.46 | 44.04 | 41.88 | 37.21 | 42.15 | 43.27 | 32.98 | 29.77 | 28.31 | + |
| Flot1 | 54.13 | 48.95 | 55.25 | 129.55 | 135.91 | 130.18 | 111.4 | 88.27 | 95.84 | 99.27 | 105.8 | 111.12 | + |
| Slc25a14 | 7.54 | 7.54 | 9 | 4.38 | 4.96 | 5.23 | 5.11 | 4.7 | 5.73 | 4.87 | 4.46 | 4.07 | + |
| Cse1l | 61.82 | 65.8 | 61.48 | 28.9 | 33.14 | 31.44 | 21.51 | 26.33 | 26.01 | 20.54 | 21.53 | 21.08 | + |
| Nub1 | 25.89 | 25.77 | 26.67 | 41.38 | 38.65 | 38.71 | 40.61 | 44.4 | 40.82 | 45.57 | 54.14 | 45.84 | + |
| Atp6v0a2 | 14.93 | 19.5 | 17.17 | 11.09 | 10.68 | 10.24 | 13.22 | 13.08 | 12.36 | 12.97 | 11.55 | 13.6 | + |
| Mtf1 | 1.67 | 2.04 | 1.96 | 1.21 | 1.14 | 1.34 | 1.2 | 1.47 | 1.17 | 1.36 | 1.32 | 1.15 | + |
| BC031181 | 66.52 | 66.02 | 67.79 | 37.31 | 37.4 | 40.31 | 38.24 | 46.72 | 44.14 | 35.7 | 36.91 | 34.85 | + |
| Btbd19 | 9.33 | 8.41 | 8.59 | 8.29 | 8.82 | 8.35 | 5.67 | 6.21 | 5.71 | 6.53 | 6.04 | 6.23 | + |
| Mtf2 | 16.69 | 14.93 | 15.41 | 7.86 | 9.33 | 8.94 | 9.38 | 10.53 | 11.21 | 8.06 | 7.72 | 8.15 | + |
| Lsm1 | 27.26 | 25.5 | 25.11 | 17.81 | 16.51 | 18.37 | 15 | 18.39 | 19.19 | 15.44 | 15.2 | 16.73 | + |
| Gna13 | 8.12 | 8.74 | 9.28 | 22.23 | 20.75 | 18.95 | 16.3 | 12.93 | 15.14 | 20.88 | 22.07 | 21.75 | + |
| Zfp810 | 3.56 | 3.34 | 3.49 | 6.05 | 5.96 | 5.69 | 5.59 | 5.01 | 4.95 | 5.85 | 5.46 | 4.86 | + |
| Zfp341 | 0.64 | 0.6 | 0.57 | 1.8 | 1.8 | 1.31 | 1.69 | 1.68 | 1.3 | 2.09 | 2.41 | 0.99 | + |
| Ankrd17 | 8.84 | 8.34 | 8.08 | 7.01 | 6.36 | 6.45 | 5.59 | 6.78 | 6.6 | 6.99 | 6.74 | 6.66 | + |
| Sft2d2 | 8.74 | 8.43 | 8.25 | 14.31 | 14.97 | 16.42 | 16.12 | 17.31 | 18.22 | 14.21 | 14.41 | 12.67 | + |
| Ubr2 | 4.28 | 4.53 | 4.4 | 6.45 | 6.09 | 6.75 | 7.5 | 7.98 | 7.15 | 8.14 | 8.83 | 7.13 | + |
| Park7 | 367.12 | 363.43 | 387.75 | 453.13 | 470.63 | 488.73 | 484.8 | 511.33 | 511.41 | 408.47 | 478.46 | 437.11 | + |
| Sub1 | 98.19 | 115.37 | 100.31 | 64.99 | 60.97 | 61.35 | 97.2 | 80.86 | 93.92 | 71.45 | 73.66 | 73.29 | + |
| Mrpl40 | 113.52 | 115.81 | 114.1 | 66.44 | 71.3 | 78.08 | 73.31 | 78.25 | 69.14 | 54.84 | 59.06 | 64.21 | + |
| Rtel1 | 10.68 | 11.27 | 10.12 | 4.33 | 5.13 | 5.11 | 3.69 | 4.35 | 3.94 | 3.68 | 2.93 | 3.98 | + |
| Ubap2l | 69.17 | 71.48 | 69.28 | 54.53 | 55.19 | 56.64 | 50.32 | 51.82 | 47.56 | 51.25 | 52.75 | 52.06 | + |
| Phb | 162.61 | 163.36 | 162.19 | 85.7 | 84.14 | 86.09 | 77.92 | 92.14 | 91.61 | 55.49 | 61.12 | 60.58 | + |
| Rplp1 | 2673.74 | 2637.88 | 2724.76 | 3677.02 | 3747.97 | 3838.76 | 3498.7 | 3455.46 | 3400.2 | 3760.51 | 4103.53 | 3774.47 | + |
| Psma1 | 375.42 | 395.49 | 375.49 | 251.89 | 242.55 | 246.6 | 218.31 | 261.93 | 246.46 | 211.14 | 227.47 | 227.86 | + |
| Rpl38 | 772.97 | 800.48 | 778.99 | 1183.71 | 1232.19 | 1258.49 | 1369.63 | 1170.09 | 1157.39 | 1393.21 | 1478.39 | 1463.79 | + |
| Oat | 51.47 | 46.17 | 46.67 | 30.94 | 35.01 | 33.05 | 34.37 | 38.59 | 36.11 | 25.94 | 32.47 | 29.61 | + |
| Zfml | 4.31 | 4.86 | 4.93 | 3.49 | 3.89 | 3.58 | 3.14 | 3.84 | 2.89 | 3.26 | 3.65 | 2.85 | + |
| Sdccag3 | 17.76 | 24.02 | 25.22 | 11.18 | 14.23 | 13.45 | 11.75 | 16.56 | 13.67 | 17.26 | 14.18 | 13.74 | + |
| Apbb1 | 7.03 | 6.81 | 6.07 | 10.04 | 11.65 | 12.32 | 16.43 | 14.14 | 14.19 | 14.98 | 18.18 | 18.51 | + |
| Pex14 | 19.87 | 21.78 | 21.27 | 16.57 | 15.2 | 14.65 | 14.2 | 14.73 | 14.09 | 14.17 | 12.56 | 14.49 | + |
| Sidt2 | 10.92 | 12.15 | 10.07 | 19.62 | 17.05 | 17.55 | 18.62 | 15.76 | 17 | 19.77 | 19.48 | 20.7 | + |
| Rpa1 | 78.58 | 80.73 | 74.22 | 50.41 | 48.58 | 50.76 | 55.58 | 46.26 | 48.76 | 41.73 | 38.53 | 39.42 | + |
| Lzts2 | 1.5 | 1.95 | 1.92 | 2.23 | 2.89 | 2.02 | 3.95 | 3.12 | 3.68 | 3.08 | 3.08 | 2.9 | + |
| Golga5 | 14.5 | 12.16 | 12.63 | 16.45 | 14.49 | 17.37 | 17.66 | 19.09 | 17.77 | 15.24 | 18.47 | 16.91 | + |
| Gpld1 | 3.66 | 4.4 | 3.83 | 35.66 | 26.72 | 31.81 | 23.72 | 14.43 | 20.54 | 22.63 | 25.21 | 21.88 | + |
| Prim2 | 25.76 | 24.51 | 22.72 | 13.38 | 16.27 | 12.44 | 12.13 | 10.31 | 10.03 | 12.85 | 15.05 | 14.52 | + |
| Gfod1 | 1.34 | 1.51 | 1.54 | 0.71 | 0.66 | 0.62 | 0.98 | 0.91 | 1.1 | 0.76 | 0.99 | 0.57 | + |
| Eif4a3 | 223.52 | 229.25 | 222.87 | 121.01 | 118.27 | 118.75 | 109.56 | 122.16 | 121.6 | 99.2 | 98.03 | 92.62 | + |
| Taf11 | 22.88 | 23.08 | 20.56 | 14.09 | 14.69 | 14.37 | 11.3 | 13.96 | 13.63 | 13.36 | 12.07 | 14.27 | + |
| Rmdn3 | 27.33 | 28.52 | 28.13 | 18.02 | 16.99 | 17.11 | 14.34 | 17.16 | 19.41 | 13.56 | 13.85 | 14.74 | + |
| Chp1 | 70.3 | 61.29 | 70.61 | 114.18 | 103.57 | 109.35 | 128.16 | 113.51 | 114.77 | 138.53 | 152.76 | 139.52 | + |
| Vps45 | 12.44 | 12.16 | 12.91 | 9.41 | 9.32 | 9.28 | 9.37 | 8.88 | 8.5 | 7.8 | 7.74 | 7.66 | + |
| Tpm1 | 25.25 | 23.22 | 25.87 | 17.2 | 18.94 | 15.66 | 16.6 | 17.43 | 17.11 | 15.39 | 15.62 | 17.85 | + |
| Hexa | 25.77 | 26.7 | 22.82 | 33.58 | 31.13 | 32.05 | 33.96 | 34.45 | 33.55 | 34.23 | 37.21 | 37.96 | + |
| Gabbr1 | 0.83 | 1.06 | 0.88 | 3.03 | 3.17 | 3.05 | 2.54 | 2.07 | 2.08 | 3.16 | 3.48 | 3.22 | + |
| Nasp | 62.49 | 65.4 | 64.59 | 26.16 | 29.24 | 24.73 | 19.97 | 25.51 | 20.77 | 17.8 | 18.84 | 16.96 | + |
| Nadk | 73.72 | 75.29 | 75.02 | 45.88 | 46.09 | 47.58 | 44.36 | 53.59 | 46.7 | 38.91 | 33.66 | 38.01 | + |
| Snx5 | 168.51 | 182.72 | 173.17 | 167.55 | 155.81 | 166.37 | 197.35 | 164.9 | 164.75 | 228.01 | 234.97 | 238.53 | + |
| Arpc4 | 147.07 | 140.91 | 136.56 | 101.72 | 103.53 | 100.7 | 124.67 | 117.97 | 122.19 | 117.45 | 108.64 | 122.37 | + |
| Map3k14 | 2.96 | 3.36 | 3.25 | 3.42 | 3.44 | 3.02 | 3.8 | 4.06 | 4.01 | 4.95 | 5.42 | 5.29 | + |
| Sfrs18 | 14.07 | 14.05 | 13.83 | 28.12 | 26.12 | 24.78 | 26.48 | 23.55 | 25.43 | 31.26 | 32.86 | 30.17 | + |
| Sap30bp | 23.1 | 22.51 | 20.97 | 15.79 | 16.97 | 16.42 | 14.74 | 14.87 | 17.2 | 13.85 | 14.45 | 17.96 | + |
| Nptn | 32.08 | 31.69 | 31.7 | 18.2 | 18.09 | 17.07 | 23.91 | 20.84 | 20.54 | 22.36 | 22.5 | 21.4 | + |
| Trp53i11 | 33.73 | 46.04 | 31.57 | 87.09 | 77.32 | 78.31 | 73.27 | 45.97 | 53.78 | 81.04 | 77.43 | 79.44 | + |
| Traf4 | 7.48 | 8.78 | 9.66 | 16.22 | 19.24 | 17.87 | 10.94 | 15 | 15.53 | 13.26 | 14.98 | 11.64 | + |
| Hirip3 | 10.14 | 10.24 | 9.84 | 4.73 | 4.88 | 6.45 | 3.55 | 4.56 | 4.51 | 2.49 | 3.02 | 3.29 | + |
| Gpatch4 | 33.34 | 35.51 | 36.04 | 12.01 | 13.58 | 14.8 | 9.37 | 11.16 | 12.45 | 7.03 | 5.65 | 8.52 | + |
| Ptbp1 | 207.86 | 216.96 | 223.67 | 116.4 | 115.72 | 116.8 | 119.6 | 128 | 125.31 | 108.77 | 105.59 | 114.69 | + |
| Oit3 | 0.95 | 0.79 | 0.44 | 1.77 | 1.7 | 1.29 | 4.13 | 2.24 | 2.75 | 2.74 | 2.09 | 3.65 | + |
| Pelp1 | 7.4 | 7.66 | 7.83 | 4.12 | 3.65 | 4.19 | 3.46 | 3.38 | 3.45 | 2.99 | 3.19 | 3.2 | + |
| Zdhhc6 | 57.06 | 57.87 | 51.63 | 31.62 | 33.85 | 33.8 | 24.85 | 38.07 | 30.01 | 25.35 | 20.27 | 28.64 | + |
| Nampt | 37.97 | 41.25 | 44.33 | 34.26 | 31.03 | 32.88 | 35.36 | 30.61 | 32.12 | 29.4 | 33.18 | 30.86 | + |
| Cd27 | 30.25 | 33.23 | 33.23 | 37.35 | 30.24 | 34.73 | 44.91 | 31.97 | 34.78 | 62.17 | 53.33 | 65.83 | + |
| Sbsn | 0.31 | 0.32 | 0.22 | 3.89 | 3.24 | 5.49 | 8.24 | 3.45 | 4.84 | 8.99 | 11.11 | 12.84 | + |
| Thumpd1 | 58.53 | 59.65 | 55.56 | 39.9 | 36.94 | 37.91 | 32.39 | 40.7 | 36.92 | 36.26 | 31.45 | 32.96 | + |
| R3hcc1 | 13.7 | 12.08 | 11.78 | 8.65 | 9.05 | 9.32 | 10.29 | 8.31 | 8.34 | 8.36 | 7.47 | 9.36 | + |
| Ube2v1 | 178.93 | 176 | 182.61 | 175.4 | 175.51 | 191.5 | 234.09 | 190.02 | 201.54 | 241.38 | 258.12 | 258.37 | + |
| Cdk7 | 9.76 | 10.05 | 10.05 | 5.98 | 5.68 | 6.25 | 5.55 | 5.49 | 6.29 | 5.32 | 4.94 | 5.98 | + |
| Elf1 | 20.21 | 19.62 | 22.14 | 22.78 | 18.99 | 20.54 | 26.65 | 22.66 | 24.3 | 29.68 | 31.48 | 28.15 | + |
| Ankzf1 | 7.37 | 7.38 | 8.52 | 15.59 | 14.34 | 15.02 | 17.1 | 16.43 | 15.58 | 19.29 | 22.14 | 22.09 | + |
| Actr6 | 13.64 | 12.98 | 11.52 | 9.13 | 9.42 | 8.57 | 7.02 | 9 | 7.84 | 7.24 | 7.27 | 6.63 | + |
| Bcl2l11 | 0.28 | 0.64 | 0.37 | 6.17 | 3.49 | 3.24 | 3.1 | 1.09 | 1.69 | 7.03 | 6.25 | 6.94 | + |
| Cog2 | 23.92 | 24.27 | 24.49 | 21.77 | 22.72 | 21.61 | 17.8 | 21.19 | 19.09 | 19.33 | 20.65 | 20.1 | + |
| Klrc1 | 123.64 | 105.7 | 117.36 | 189.21 | 164.83 | 173.6 | 220.46 | 237.63 | 217.29 | 188.38 | 199.78 | 191.87 | + |
| Coro1c | 65.24 | 62.23 | 65.72 | 49.52 | 48.71 | 51.2 | 41.53 | 45.5 | 47.97 | 40.24 | 45.92 | 42.36 | + |
| Sccpdh | 13.93 | 13.41 | 14.66 | 20.46 | 23.98 | 22.56 | 29.99 | 26.91 | 26.32 | 25.07 | 29.73 | 26.24 | + |
| Hmmr | 33.45 | 26.74 | 29.51 | 20.89 | 28.55 | 23.58 | 13.27 | 14.5 | 12.83 | 16.38 | 15.14 | 13.44 | + |
| Hivep2 | 1.26 | 1.16 | 1.17 | 1.78 | 1.67 | 1.69 | 2.24 | 1.99 | 1.97 | 1.86 | 1.89 | 2.19 | + |
| Lpin1 | 4.54 | 4.83 | 5.06 | 8.95 | 8.05 | 9.02 | 11.74 | 9.84 | 9.96 | 9.6 | 11.34 | 10.46 | + |
| Pin1 | 40.69 | 39.38 | 38.63 | 16.48 | 16.87 | 16.96 | 12.87 | 16.44 | 15.94 | 8.27 | 9.16 | 9.47 | + |
| Dock9 | 2.34 | 2.37 | 2.39 | 7.15 | 5.43 | 6.06 | 4.67 | 3.6 | 3.86 | 7.08 | 6.36 | 6.96 | + |
| Ruvbl1 | 137.8 | 142.15 | 148.11 | 50.79 | 61.18 | 61.46 | 43.92 | 64.84 | 58.74 | 36.57 | 33.03 | 34.27 | + |
| Bmpr1a | 0.7 | 0.92 | 0.92 | 1.01 | 1.24 | 1.15 | 1.35 | 1.32 | 1.24 | 2.2 | 2.43 | 2.33 | + |
| Dgcr8 | 12.75 | 12.96 | 11.96 | 7.21 | 6.76 | 6.58 | 3.8 | 6.55 | 5.83 | 4.46 | 3.85 | 4.27 | + |
| Ipcef1 | 2.13 | 1.66 | 1.47 | 2.27 | 1.22 | 1.18 | 2.15 | 1.63 | 1.95 | 3.65 | 2.72 | 4.31 | + |
| Mif4gd | 29.21 | 29.28 | 30.6 | 61.97 | 55.02 | 57.42 | 51.05 | 41.84 | 42.01 | 63.97 | 54.1 | 67.17 | + |
| Slc48a1 | 38.44 | 38.06 | 39.36 | 74.72 | 72.28 | 75.25 | 86.33 | 75.13 | 81.43 | 99.69 | 111.11 | 99.28 | + |
| Tmem43 | 18.65 | 15.73 | 15.88 | 27.56 | 27.29 | 25 | 31.24 | 30.77 | 30.85 | 33.04 | 34.16 | 33.55 | + |
| Sphk2 | 6.87 | 6.54 | 7.08 | 5.84 | 5.43 | 5.12 | 5.32 | 5.35 | 4.96 | 4.56 | 4.77 | 4.41 | + |
| Tmem159 | 14.29 | 12.37 | 14.08 | 19.04 | 19.86 | 20.61 | 29.04 | 31.59 | 26.58 | 31.78 | 30.31 | 30.8 | + |
| Ticam2 | 1.64 | 1.68 | 1.54 | 1.53 | 1.93 | 1.17 | 1.16 | 0.99 | 0.69 | 0.42 | 0.42 | 0.18 | + |
| Pde12 | 25.86 | 23.56 | 23.87 | 19.61 | 18.43 | 20.28 | 16.37 | 18.49 | 18.62 | 19.3 | 17.48 | 20.61 | + |
| Dnaja1 | 126.87 | 126.51 | 131.16 | 65 | 75.5 | 70.41 | 50.18 | 72.92 | 60.59 | 47.74 | 49.14 | 50.29 | + |
| Ganab | 55.04 | 52.2 | 52.27 | 42.26 | 45.89 | 45.61 | 40.82 | 48.45 | 45.8 | 41.48 | 44.27 | 40.74 | + |
| Slc29a1 | 71.49 | 77.59 | 75.72 | 38.64 | 43.13 | 43.36 | 39.56 | 46.62 | 48.5 | 31.53 | 30.9 | 31.43 | + |
| Cenpk | 26.57 | 25.46 | 25.22 | 13.31 | 19.27 | 17.9 | 10.97 | 12.89 | 12.37 | 11.15 | 11.48 | 9.14 | + |
| Ifrd1 | 24.92 | 35.38 | 39.47 | 48.81 | 54.19 | 51.02 | 46.62 | 64.13 | 62.13 | 53.95 | 53.95 | 39.27 | + |
| Polr2b | 42.56 | 41.39 | 39.76 | 27.05 | 26.95 | 24.75 | 24.78 | 25.9 | 25 | 26.04 | 23.8 | 27.16 | + |
| Mut | 11.98 | 10.85 | 11.29 | 7 | 6.17 | 7.11 | 8.18 | 10.31 | 8.31 | 7.18 | 5.52 | 6.95 | + |
| Ndufa5 | 126.34 | 140.84 | 133.35 | 78.29 | 86.08 | 81.54 | 69.4 | 86.54 | 76.71 | 58.65 | 56.13 | 55.97 | + |
| Sra1 | 148.67 | 150.81 | 144.85 | 182.15 | 179.53 | 190.75 | 206.28 | 210.79 | 210.02 | 184.53 | 191.96 | 193.09 | + |
| Edf1 | 194.82 | 203.36 | 199.38 | 165.71 | 157.41 | 159.53 | 175.13 | 158.21 | 166.41 | 155.2 | 153.11 | 165.71 | + |
| Cenph | 22.16 | 25.01 | 19.55 | 12.14 | 13.6 | 13.49 | 10.99 | 11.19 | 10.13 | 6.05 | 8.16 | 9.12 | + |
| Atp5j | 335.54 | 338.29 | 335.62 | 195.57 | 218.98 | 211.86 | 196.09 | 237.19 | 212.62 | 149.35 | 164.47 | 165.17 | + |
| Rab1 | 93.52 | 89.09 | 94.16 | 73.43 | 72.49 | 77.81 | 72.77 | 80.28 | 77.85 | 67.6 | 71.58 | 68.28 | + |
| Trmt10a | 4.9 | 5.57 | 5.14 | 2.84 | 2.8 | 2.74 | 2.92 | 3.27 | 3.32 | 2.44 | 3.21 | 2.21 | + |
| Aftph | 9.38 | 9.2 | 10.15 | 9.74 | 10.39 | 8.83 | 11.12 | 10.82 | 10.44 | 13.23 | 13.91 | 12.63 | + |
| Bnip2 | 27.79 | 27.38 | 26.39 | 18.84 | 18.39 | 19.51 | 16.05 | 17.72 | 18.12 | 15.54 | 14.42 | 15.77 | + |
| Immt | 98.54 | 95.08 | 97.67 | 56.63 | 63.96 | 68.57 | 53.63 | 60.42 | 54.43 | 51.13 | 49.95 | 46.75 | + |
| Qrich1 | 50.01 | 50.56 | 43.51 | 37.85 | 39.05 | 35.26 | 33.27 | 38.77 | 34.97 | 35.02 | 34.83 | 38.06 | + |
| Crebrf | 0.2 | 0.32 | 0.31 | 1.92 | 1.15 | 1.22 | 2.03 | 1.21 | 1.54 | 3.12 | 3.04 | 2.39 | + |
| Fam195b | 16.58 | 19.12 | 19.83 | 40.07 | 43.57 | 38.92 | 51.12 | 41.73 | 42.25 | 56.64 | 56.06 | 51.86 | + |
| Rpl13a | 1274.71 | 1240.23 | 1307.02 | 1874.25 | 1853.02 | 1943.64 | 2015.87 | 1863.74 | 1887.28 | 2054.88 | 2251.64 | 2093.26 | + |
| Rab37 | 6.01 | 7.8 | 4.63 | 16.14 | 10.85 | 10.99 | 24.31 | 12.91 | 19.4 | 20.42 | 15.4 | 21.58 | + |
| Akap1 | 4.19 | 4.28 | 3.18 | 2.43 | 2.23 | 2.49 | 1.98 | 2.71 | 2.87 | 2.6 | 2.47 | 1.94 | + |
| Ppp1r11 | 32.03 | 32.51 | 31.62 | 21.19 | 19.89 | 20.81 | 17.8 | 20.07 | 19.78 | 17.99 | 14.13 | 17.99 | + |
| Neb | 4.2 | 3.96 | 4.23 | 19.77 | 18.26 | 17.56 | 8.28 | 7.13 | 7.6 | 13.11 | 14.21 | 14.58 | + |
| Hdgfrp2 | 46.57 | 42.71 | 46.5 | 35.6 | 37.38 | 39.21 | 36.6 | 35.5 | 34.77 | 33.24 | 33.13 | 34.78 | + |
| 2310015A10Rik | 0.04 | 0.08 | 0.04 | 0.44 | 0.35 | 0.72 | 0.39 | 0.22 | 0.48 | 0.47 | 0.39 | 0.44 | + |
| Gm4951 | 0.78 | 1.01 | 0.62 | 0.89 | 0.43 | 0.85 | 0.07 | 0.04 | 0.04 | 0.21 | 0.05 | 0.22 | + |
| Doc2a | 5.91 | 5.64 | 7.01 | 3.32 | 2.48 | 2.67 | 1.81 | 2.18 | 1.75 | 2.95 | 1.95 | 2.56 | + |
| Ywhae | 226.09 | 242.22 | 233.31 | 172.05 | 179.16 | 188.29 | 164.26 | 179.98 | 179.78 | 137.76 | 144.51 | 144.16 | + |
| Gse1 | 2.18 | 2.38 | 1.99 | 2.46 | 2.42 | 2.49 | 3.24 | 2.94 | 2.85 | 3.46 | 3.84 | 4.12 | + |
| Gatad1 | 20.9 | 20.04 | 20.82 | 29.8 | 29 | 30.2 | 23.92 | 24.72 | 25.49 | 25.6 | 26.52 | 23.13 | + |
| Ctsf | 1.04 | 0.59 | 0.36 | 2.05 | 2.02 | 2.49 | 6.08 | 3.42 | 3.02 | 5.68 | 4.59 | 4.79 | + |
| Akap8 | 31.96 | 35.73 | 35.08 | 23.65 | 22.31 | 21.54 | 18.22 | 20.04 | 20.69 | 20.31 | 22.9 | 22.73 | + |
| Fez2 | 9.95 | 11.48 | 9.59 | 20.21 | 17.02 | 17.32 | 15.67 | 15.74 | 13.25 | 19.94 | 16.13 | 15.2 | + |
| Mapk1ip1l | 49.76 | 50.62 | 53.81 | 31.54 | 31.39 | 30.57 | 27.92 | 31.9 | 31.14 | 30.39 | 28.8 | 28.25 | + |
| Hmgcs1 | 169.76 | 175.48 | 148.65 | 139.22 | 126.78 | 146.11 | 136.69 | 103.01 | 111.8 | 115.38 | 125.44 | 132.07 | + |
| Bptf | 3.07 | 3.42 | 3.4 | 4.98 | 4.87 | 4.67 | 4.76 | 4.64 | 4.9 | 5.11 | 6.09 | 5.43 | + |
| N4bp2l1 | 3.26 | 2.93 | 2.41 | 15.66 | 13.36 | 17.77 | 25.36 | 17.1 | 20.9 | 26.3 | 30.78 | 27.49 | + |
| Ndufaf2 | 33.5 | 30.29 | 33.15 | 22.68 | 21.39 | 27.76 | 18.25 | 24.36 | 22.41 | 19.35 | 20.94 | 21.7 | + |
| Cstf2t | 9.05 | 9.1 | 9.8 | 6.35 | 6.07 | 7.22 | 5.54 | 7.87 | 6.75 | 6.83 | 5.95 | 5.84 | + |
| Eif2s1 | 182.57 | 186.7 | 187.01 | 96.37 | 103.09 | 104.11 | 84.64 | 111.49 | 102.02 | 75.15 | 80.97 | 72.99 | + |
| Wasl | 3.98 | 4.62 | 3.23 | 8.33 | 9.64 | 8.08 | 8.59 | 7.09 | 7.34 | 10.29 | 12.38 | 10.98 | + |
| Uba2 | 140.11 | 146.92 | 146.76 | 72.92 | 80.89 | 78.24 | 68.04 | 78.89 | 75.59 | 64.04 | 63.22 | 65.07 | + |
| Hnrnph1 | 109.59 | 112.52 | 111.97 | 49.45 | 53.17 | 55.82 | 37.6 | 48.21 | 44.73 | 46.77 | 46.6 | 40.06 | + |
| Psmc4 | 145.82 | 157.83 | 146.54 | 69.5 | 74.86 | 71.1 | 56.38 | 73.87 | 69.45 | 57.25 | 47.38 | 53.31 | + |
| Glb1 | 34.4 | 28.85 | 32.31 | 46.08 | 42.07 | 42.17 | 51.06 | 43.72 | 45.69 | 44.78 | 48.72 | 46.3 | + |
| Atic | 18.83 | 18.85 | 21.03 | 11.3 | 12.74 | 10.53 | 9.51 | 13.48 | 12.47 | 8.06 | 9.56 | 8.83 | + |
| Anxa7 | 67.5 | 66.46 | 68.56 | 65.84 | 65.11 | 63.39 | 81.71 | 73.98 | 74.46 | 75.68 | 79.79 | 79.99 | + |
| Kdm2b | 12.09 | 10.39 | 12.91 | 6.5 | 7.61 | 6.48 | 9.05 | 9.72 | 9.62 | 9.63 | 7.75 | 8.53 | + |
| Ufm1 | 11.65 | 11.05 | 11.15 | 7.14 | 6.73 | 7.32 | 7.21 | 7.42 | 7.22 | 6.69 | 6.93 | 6.54 | + |
| Kctd7 | 1.63 | 1.3 | 1.17 | 2.83 | 2.57 | 2.62 | 2.06 | 2.33 | 1.9 | 2.34 | 2.4 | 2.24 | + |

| | | | | | | | | | | | | |
|---|---|---|---|---|---|---|---|---|---|---|---|---|---|
| Rdh12 | 1.94 | 1.53 | 2.23 | 6.95 | 7.35 | 3.47 | 7.11 | 5.82 | 6.76 | 6.13 | 5.15 | 6.73 | + |
| Ccdc117 | 2.35 | 2.02 | 2.19 | 4.87 | 5.46 | 5.68 | 4.93 | 5.97 | 5.74 | 10.09 | 10.46 | 8.49 | + |
| Fasn | 27.79 | 30.28 | 32.8 | 15.78 | 15.26 | 13.88 | 14.33 | 14.59 | 15.76 | 13.24 | 12.57 | 11.07 | + |
| Rps6kb1 | 14.18 | 14.38 | 13.93 | 11.13 | 10.62 | 10.44 | 9 | 9.52 | 9.48 | 9.96 | 10.26 | 9.55 | + |
| Top1mt | 10.69 | 10.62 | 10.67 | 5.26 | 5.9 | 5.65 | 5.73 | 8.34 | 6.71 | 4.41 | 5.56 | 4.87 | + |
| Sorcs2 | 0.12 | 0.13 | 0.07 | 0.28 | 0.35 | 0.12 | 0.7 | 0.89 | 0.98 | 0.55 | 0.76 | 0.17 | + |
| Mgst3 | 16.81 | 14.11 | 14.3 | 36.23 | 34.8 | 36.64 | 35.2 | 28.21 | 29.06 | 35.95 | 31.06 | 32.95 | + |
| Ergic2 | 34.48 | 32.95 | 33.09 | 27.19 | 27.18 | 29.35 | 21.02 | 25.1 | 24.23 | 23.04 | 23.83 | 21.8 | + |
| Snx20 | 0.68 | 2.07 | 3.32 | 11.69 | 22.98 | 14.76 | 23.86 | 32.23 | 38.57 | 40.61 | 37.86 | 23.43 | + |
| Maf1 | 47.57 | 50.33 | 46.12 | 83.47 | 74.54 | 78.57 | 92.37 | 82.83 | 81.87 | 104.21 | 105.46 | 107.56 | + |
| Mettl16 | 31.48 | 31.68 | 34.41 | 10.9 | 11.74 | 11.53 | 7.61 | 12.77 | 12.09 | 6.86 | 7.39 | 7.74 | + |
| Ptplad1 | 59.42 | 56.33 | 56.07 | 46.47 | 52.33 | 51.92 | 33.14 | 40.92 | 39.57 | 46.13 | 45.87 | 47.93 | + |
| Chst15 | 0.27 | 0.16 | 0.04 | 0.74 | 0.32 | 0.7 | 1.4 | 0.98 | 1.04 | 1.02 | 1.05 | 1.65 | + |
| Nt5c3l | 39.29 | 42.69 | 38.17 | 21.84 | 23.49 | 23.96 | 28.09 | 26.67 | 28.2 | 20.85 | 18.79 | 24.8 | + |
| Gdi1 | 48.42 | 43.96 | 48.83 | 70.93 | 67.05 | 64.33 | 57.98 | 62.47 | 58.19 | 61.23 | 63.76 | 64.02 | + |
| Muc2 | 0.15 | 0.06 | 0.12 | 2.03 | 1.16 | 1.52 | 3.92 | 1.61 | 2.12 | 4.54 | 4.57 | 5.11 | + |
| Ncl | 182.4 | 191.25 | 194.25 | 109.45 | 111.11 | 111.9 | 103.51 | 107.94 | 108.9 | 101.99 | 109.07 | 102.16 | + |
| Atp2a2 | 18.72 | 15.39 | 16.63 | 7.18 | 7.55 | 10.03 | 5.68 | 8.3 | 8.05 | 6.65 | 5.23 | 6.31 | + |
| Dctn2 | 82.31 | 81.82 | 77.88 | 94.88 | 88.49 | 94.39 | 106.34 | 98.78 | 99.12 | 90.72 | 96.9 | 97.42 | + |
| Rps14 | 2380.9 | 2455.65 | 2370.64 | 3763.62 | 3693.21 | 3816.24 | 3759.33 | 3448.16 | 3639.8 | 4024.62 | 4311.43 | 4060.25 | + |
| Slc2a9 | 0.68 | 0.71 | 0.43 | 1.88 | 0.92 | 1.43 | 2.86 | 1.91 | 1.88 | 2.11 | 1.52 | 2.11 | + |
| Pomp | 268.31 | 274.74 | 269.68 | 178.67 | 193.14 | 195.29 | 210.35 | 222.11 | 222.14 | 175.14 | 186.09 | 178.34 | + |
| Capn5 | 1.33 | 1.46 | 1.31 | 3.37 | 2.69 | 3.4 | 4.05 | 2.22 | 2.72 | 3.23 | 2.7 | 3.65 | + |
| Pop7 | 41.95 | 48.34 | 41.85 | 26.94 | 25.85 | 26.73 | 27.86 | 27.49 | 26.35 | 22.39 | 22.02 | 21.54 | + |
| Aldoc | 32.23 | 26.46 | 29.93 | 117.39 | 98.12 | 91.36 | 173.25 | 104.42 | 124.92 | 130.15 | 122.11 | 156.39 | + |
| Patz1 | 6.59 | 6.63 | 6.79 | 11.59 | 11.35 | 8.94 | 9.11 | 7.75 | 7.68 | 15.23 | 15.09 | 18.43 | + |
| Ehd3 | 6.06 | 6.94 | 5.71 | 2.38 | 2.03 | 1.73 | 3.63 | 2.39 | 3.65 | 2.11 | 1.55 | 2.4 | + |
| Rrm2 | 269.53 | 279.99 | 260.38 | 61.29 | 72.85 | 68.75 | 44.07 | 48.91 | 45.75 | 25.03 | 30.49 | 28.48 | + |
| Zkscan3 | 9.15 | 8.16 | 8.39 | 17.26 | 15.81 | 14.71 | 10.5 | 11.13 | 13.47 | 14.69 | 14.5 | 16.4 | + |
| Gm15446 | 3.73 | 3.61 | 3.07 | 5.79 | 4.86 | 4.96 | 4.47 | 6.41 | 5.52 | 5.42 | 5.42 | 5.81 | + |
| Pttg1ip | 23.99 | 23.41 | 25.1 | 43.59 | 40.51 | 38.9 | 36.65 | 38.25 | 34.05 | 45.04 | 44.49 | 44.72 | + |
| Mki67ip | 85.67 | 87.71 | 81.01 | 40.65 | 37.65 | 34.87 | 34.93 | 39.49 | 41.13 | 30.48 | 29.06 | 30.31 | + |
| Dus1l | 52.26 | 55.35 | 53.3 | 32.54 | 29.23 | 29.7 | 23.24 | 27.17 | 26.81 | 25.86 | 17.72 | 22.65 | + |
| Rcn3 | 22.16 | 18.9 | 23.7 | 57.85 | 49.71 | 58.03 | 69 | 58.9 | 63.32 | 57.4 | 64.67 | 59.31 | + |
| Dnajc18 | 3.81 | 3.64 | 4.23 | 5.86 | 6.15 | 6.74 | 6.69 | 6.34 | 6.59 | 6.86 | 8.13 | 7.74 | + |
| Rwdd2b | 7.9 | 9.76 | 8.15 | 4.75 | 5.04 | 5.14 | 4.59 | 5.83 | 4.64 | 3.81 | 3.28 | 5.59 | + |
| N6amt2 | 57.23 | 57.63 | 58.27 | 19.56 | 21.8 | 23.8 | 15.94 | 23.68 | 21.31 | 18.75 | 15.04 | 18.33 | + |
| Pdk2 | 0.91 | 0.8 | 0.96 | 3.88 | 3.2 | 5.13 | 8.34 | 6.51 | 5.22 | 6.32 | 7.03 | 7.49 | + |
| Ddx56 | 40.71 | 41.99 | 41.67 | 19.27 | 22.64 | 23.2 | 16.71 | 22.73 | 19.65 | 13.12 | 13.12 | 13.64 | + |
| Tnfsf8 | 12.73 | 12.8 | 13.82 | 11.8 | 11.54 | 13.05 | 3.04 | 3.45 | 3.08 | 1.13 | 1.32 | 1.29 | + |
| Tln1 | 16.47 | 18.34 | 17.97 | 18.88 | 17.12 | 17.75 | 22.13 | 19.34 | 18.74 | 24.83 | 21.23 | 24.27 | + |
| Ippk | 13.12 | 12.79 | 12.26 | 5.79 | 5.91 | 5.93 | 5.25 | 6.63 | 7.08 | 4.91 | 5.19 | 4.3 | + |
| H2-Q1 | 1.16 | 1.25 | 1.71 | 2.52 | 2.62 | 3 | 6.51 | 5.3 | 5.72 | 3.51 | 5.71 | 5.21 | + |
| Gimap7 | 55.77 | 59.09 | 54.97 | 158 | 117.62 | 126.12 | 221.03 | 161.24 | 164.34 | 241.94 | 234.7 | 265.73 | + |
| Dtymk | 92.28 | 97.51 | 98.49 | 58.32 | 61.41 | 60.61 | 46.09 | 54.53 | 50.18 | 40.2 | 44.39 | 44.49 | + |
| Rmnd5a | 3.29 | 3.41 | 3.24 | 5.47 | 4.67 | 4.69 | 4.83 | 4.96 | 4.19 | 5.4 | 5.95 | 4.96 | + |
| Erich1 | 9.29 | 9.19 | 9.97 | 4.42 | 6.55 | 4.62 | 6.38 | 9.09 | 7.69 | 5.35 | 6.17 | 5.97 | + |
| Mmd | 13.02 | 12.49 | 12.48 | 4.92 | 4.88 | 4.7 | 5.66 | 6.91 | 7.2 | 5.96 | 4.86 | 6.15 | + |
| Gns | 2.47 | 2.96 | 2.83 | 5.81 | 5.52 | 5.49 | 5.34 | 7.13 | 5.12 | 5.92 | 6.17 | 5.17 | + |
| Gpaa1 | 33.03 | 33.27 | 28.57 | 21 | 22.42 | 19.89 | 19.67 | 17.19 | 18.89 | 16.18 | 18.1 | 17.5 | + |
| Polrmt | 14.16 | 12.49 | 13.34 | 11.04 | 10.89 | 10.44 | 8.87 | 8.95 | 9.24 | 7.94 | 8.65 | 8.63 | + |
| Mtdh | 36.3 | 38.03 | 35.51 | 22.56 | 21.88 | 21.63 | 21.61 | 24.36 | 24.24 | 19.4 | 20.42 | 18.88 | + |
| Ccdc84 | 6.8 | 9.05 | 7.38 | 7.92 | 6.47 | 8.1 | 9.48 | 8.37 | 7.51 | 12.6 | 14.11 | 13.88 | + |
| Arhgef25 | 4.96 | 5.21 | 4.75 | 8.51 | 10.17 | 9.79 | 11.13 | 10.18 | 10.56 | 11.93 | 14.21 | 13.68 | + |
| Gen1 | 3.37 | 3.03 | 2.93 | 1.25 | 1.79 | 1.36 | 0.67 | 0.86 | 0.95 | 0.66 | 0.68 | 0.47 | + |
| 9030624G23Rik | 0.63 | 0.47 | 0.65 | 1.11 | 0.99 | 1.29 | 1.36 | 1.22 | 1.68 | 1.46 | 1.32 | | + |
| Fastkd5 | 8.45 | 7.52 | 8.29 | 4.94 | 5.13 | 3.22 | 3.27 | 3.38 | 3.77 | 4.13 | 3.61 | 2.73 | + |
| Cdk4 | 266.85 | 282.07 | 274.08 | 162.5 | 157.75 | 163.73 | 142.95 | 145.53 | 143.59 | 127.34 | 115.8 | 140.03 | + |
| Rpl13 | 2212 | 2264.42 | 2263.5 | 3954.24 | 3816.13 | 3972.85 | 3702.96 | 3479.22 | 3530.93 | 4038.97 | 4351.84 | 4135.25 | + |
| Slc25a10 | 6.27 | 5.27 | 5.88 | 3.22 | 3.84 | 3.3 | 2.77 | 2.54 | 2.09 | 2.77 | 2 | 1.92 | + |
| Atg16l2 | 15.41 | 16.43 | 14.82 | 40.54 | 34.59 | 36.77 | 46.63 | 28.58 | 35.17 | 45.67 | 55.14 | 55.53 | + |
| Abce1 | 105.34 | 108.82 | 113.58 | 65.42 | 72.08 | 70.26 | 56.59 | 71.72 | 71.33 | 55.66 | 68.17 | 57.67 | + |
| Ggnbp2 | 14.57 | 18.4 | 21.44 | 37.9 | 38.39 | 34.19 | 36.02 | 35.89 | 35.57 | 44.9 | 44.88 | 36.15 | + |
| Rad23b | 8.75 | 8.69 | 9.55 | 7.15 | 7.01 | 5.32 | 5.53 | 6.69 | 6.66 | 7.26 | 6.58 | 6.49 | + |
| Psmb7 | 184.09 | 187.29 | 186.87 | 89.95 | 89.21 | 88.32 | 93 | 115.24 | 105.14 | 66.42 | 70.54 | 70.14 | + |
| Abcc4 | 3.63 | 4.55 | 4.61 | 2.47 | 2.75 | 2.6 | 2.28 | 2.72 | 2.89 | 2.09 | 1.86 | 2.33 | + |
| Polr2j | 101.12 | 102.87 | 102.78 | 47.67 | 54.65 | 61.6 | 51.97 | 54.03 | 60.61 | 48.99 | 47.96 | 52.53 | + |
| Ephx1 | 4.97 | 3.7 | 4.25 | 6.26 | 6.56 | 7.81 | 7.66 | 8.68 | 9.67 | 7.56 | 6.14 | 7.06 | + |
| Slc4a1ap | 14.84 | 13.51 | 13.98 | 11.7 | 9.95 | 11.41 | 9.98 | 10.68 | 9.82 | 9.32 | 10.32 | 10.13 | + |
| Dph3 | 15.13 | 14.95 | 15.6 | 7.82 | 8.16 | 8.92 | 7.57 | 8.94 | 8.72 | 8.01 | 6.58 | 6.76 | + |
| Stat1 | 75.35 | 86.09 | 77.76 | 74.93 | 71.74 | 72.94 | 91.01 | 60.82 | 59.06 | 108.26 | 88.64 | 118.23 | + |
| Ephb6 | 0.97 | 1.04 | 1.01 | 2.12 | 1.96 | 1.59 | 1.79 | 1.98 | 1.34 | 2.13 | 1.52 | 1.69 | + |
| Surf2 | 40.83 | 43.31 | 45.85 | 25.87 | 30.78 | 28.2 | 27.46 | 28.96 | 31.87 | 27.05 | 25.9 | 24.47 | + |
| Naa35 | 23.13 | 23.8 | 22.88 | 15.02 | 15.78 | 14.69 | 15.48 | 17.08 | 15.02 | 11.74 | 10.65 | 13.82 | + |
| Rps5 | 2338.57 | 2363.48 | 2354.98 | 3789.26 | 3622.06 | 3808.09 | 3568.24 | 3282.98 | 3401.96 | 4119.18 | 4252.12 | 4062.51 | + |
| Peo1 | 7.8 | 7.6 | 7.88 | 6.94 | 5.77 | 6.26 | 4.99 | 5.52 | 5.59 | 4.46 | 6.09 | 5.07 | + |
| Clns1a | 77.72 | 74.09 | 75.16 | 42.45 | 42.51 | 43.76 | 40.17 | 49.21 | 44.99 | 35.02 | 38.78 | 33.43 | + |
| Blmh | 40.94 | 45.73 | 43.46 | 33.12 | 33.31 | 30.44 | 33.43 | 35.57 | 34.82 | 33.64 | 34.43 | 34.69 | + |
| A230046K03Rik | 33.74 | 39.24 | 37.91 | 14.69 | 15.89 | 14.82 | 16.94 | 17.99 | 17.9 | 14.53 | 14.45 | 13.96 | + |
| Tra2a | 55.59 | 54.78 | 55.56 | 19.12 | 22.88 | 21.49 | 15.5 | 21.49 | 20.19 | 14.8 | 16.85 | 13.05 | + |
| Zbtb46 | 1.72 | 2.14 | 1.9 | 4.5 | 5.29 | 4.85 | 2.82 | 6.95 | 4.08 | 4.71 | 4.23 | 2.1 | + |
| Dhx30 | 27.86 | 26.78 | 29.19 | 18.74 | 20.65 | 20.31 | 18.1 | 20.08 | 19.77 | 16.95 | 19.33 | 18.01 | + |
| 15-Sep | 117.81 | 116.63 | 119.56 | 84.54 | 84.53 | 84.93 | 80.11 | 91.49 | 85.71 | 73.62 | 76.74 | 69.46 | + |
| Aqp11 | 0.39 | 0.57 | 0.41 | 0.86 | 1.33 | 1.98 | 2 | 1.24 | 1.38 | 2.12 | 0.84 | 1.48 | + |
| Ccdc111 | 5.5 | 5.74 | 6.06 | 14.2 | 13.35 | 15.36 | 18.05 | 14.36 | 15.83 | 16.55 | 17.92 | 15.24 | + |
| P2rx4 | 9.08 | 6.89 | 8.95 | 16.93 | 15.82 | 18.72 | 23.27 | 21.24 | 20.62 | 16.53 | 16.88 | 18.97 | + |
| Ddx27 | 76.49 | 77.59 | 79.69 | 43.66 | 42.57 | 42.16 | 33.48 | 45.5 | 42.28 | 27.4 | 29.52 | 29.24 | + |
| Il1rap | 3.58 | 2.62 | 2.99 | 1.7 | 2 | 1.45 | 1.92 | 1.99 | 2.35 | 1.55 | 1.55 | 1.66 | + |
| Fcf1 | 193.6 | 198.54 | 192.62 | 124.91 | 131.47 | 132.62 | 119.99 | 152.74 | 133.53 | 104.35 | 107.68 | 105.07 | + |
| Traf3 | 3.5 | 3.84 | 3.67 | 3.36 | 3.29 | 3.94 | 3.66 | 4.62 | 4.26 | 4.04 | 5.27 | 5.21 | + |
| Socs3 | 14.92 | 14.99 | 21.52 | 46.77 | 48.75 | 42.58 | 24.09 | 24.56 | 21.96 | 32.07 | 29.43 | 32.44 | + |
| 4930402H24Rik | 0.76 | 0.63 | 0.65 | 2.71 | 2.9 | 2.93 | 2.49 | 2.08 | 2.06 | 2.56 | 3.4 | 3.33 | + |
| Mapre1 | 41.16 | 39.43 | 40.74 | 23.32 | 26.07 | 25.31 | 23.73 | 25.78 | 25.72 | 22.64 | 21.16 | 21.61 | + |
| Atf7ip | 15.85 | 13.42 | 14.6 | 27.22 | 28.94 | 28.08 | 29.2 | 23.09 | 26.27 | 37.27 | 38.09 | 36.16 | + |
| Mrpl36 | 91.08 | 94.32 | 87.29 | 46.81 | 45.64 | 48.35 | 46.26 | 58.15 | 56.27 | 38.24 | 39.01 | 40.71 | + |
| Tyms | 30.89 | 34.07 | 29.57 | 7.94 | 8.79 | 9.45 | 6.35 | 8.12 | 7.28 | 6.02 | 5.45 | 6.39 | + |
| Manf | 82.6 | 81.94 | 86.82 | 31.2 | 38.49 | 34.36 | 30.74 | 42.87 | 35.09 | 26.17 | 29.59 | 25.84 | + |
| Lemd2 | 3.3 | 4.1 | 3.27 | 8.18 | 8.73 | 7.81 | 9.83 | 8.59 | 7.34 | 10.69 | 12.04 | 9.7 | + |
| Nedd9 | 26.73 | 24.37 | 23.62 | 15.89 | 12.48 | 13.24 | 18.48 | 23.39 | 20.2 | 16.89 | 16.02 | 19.6 | + |
| Dhx33 | 8.52 | 10.47 | 10.47 | 5.35 | 6 | 6.02 | 3.71 | 6.35 | 5.18 | 5.08 | 4.66 | 4.57 | + |
| Prss41 | 0.36 | 0.47 | 0.19 | 1.25 | 1.35 | 1.03 | 1.71 | 1.85 | 1.25 | 1.39 | 1.16 | 1.45 | + |
| Rnaseh1 | 28.09 | 30.56 | 26.88 | 15.27 | 18.72 | 17.68 | 14.12 | 16.97 | 16.39 | 16.28 | 13.41 | 14.64 | + |
| Med13 | 2.42 | 2.26 | 2.46 | 2.77 | 2.68 | 2.59 | 2.17 | 2.64 | 2.32 | 3.06 | 3.39 | 2.78 | + |
| Timm23 | 325.57 | 318.26 | 318.53 | 160.9 | 173.57 | 180.7 | 144.68 | 192.07 | 174.22 | 136.26 | 146.39 | 138.91 | + |
| Zeb1 | 3.01 | 2.71 | 3.38 | 6.71 | 8.26 | 7.62 | 3.72 | 4.68 | 4 | 6.23 | 6.93 | 5.94 | + |
| Ubtd1 | 17.53 | 17.22 | 17.45 | 29.8 | 35.81 | 30.95 | 30.18 | 30.16 | 30.15 | 26.11 | 29.64 | 24.12 | + |
| Hectd3 | 5.29 | 6.31 | 7.1 | 13.02 | 11.14 | 9.46 | 12.26 | 13.15 | 12.68 | 15.17 | 13.32 | 13.28 | + |
| Cst3 | 424.29 | 378.2 | 379.28 | 267.94 | 276.52 | 293.53 | 216.73 | 244.98 | 220.33 | 214.56 | 219.03 | 247 | + |
| Bckdhb | 3.67 | 3.37 | 4.44 | 4.45 | 3.53 | 4.34 | 2.63 | 3.7 | 3.62 | 1.95 | 1.56 | 1.51 | + |
| Gmnn | 107.97 | 103.02 | 106.29 | 59.9 | 66.77 | 63.84 | 45.64 | 52.35 | 51.89 | 44.03 | 40.62 | 36.11 | + |
| Frmd4b | 1.09 | 1.03 | 0.73 | 6.76 | 6.11 | 5.87 | 2.89 | 2.3 | 2.39 | 4.98 | 5.1 | 5.66 | + |
| Slc35c2 | 35.77 | 37.12 | 32.97 | 18.48 | 20.35 | 20.63 | 27.01 | 29.7 | 33.42 | 18.39 | 20.05 | 21.19 | + |
| Lap3 | 74.29 | 79.41 | 77.75 | 34.63 | 41.11 | 40.79 | 38.09 | 44.07 | 44.94 | 27.44 | 28.93 | 30.79 | + |
| Stoml2 | 141.95 | 133.29 | 140.67 | 84.49 | 90.56 | 97.6 | 80.39 | 97.45 | 89.28 | 67.03 | 66.54 | 66.92 | + |
| Ctr9 | 14.87 | 14.97 | 15.52 | 12.66 | 11.16 | 10.59 | 10.17 | 10.18 | 10.58 | 10.86 | 11.47 | 12.94 | + |
| Dph2 | 15.82 | 14.19 | 13.55 | 8.53 | 9.41 | 9.92 | 6.57 | 7.49 | 9.35 | 8.52 | 5.86 | 7.92 | + |
| Txnip | 17.35 | 16.6 | 17.25 | 124.41 | 100.16 | 105.96 | 124.27 | 78.88 | 80.87 | 162.64 | 168.51 | 178.79 | + |
| Dkkl1 | 8.66 | 9.86 | 7.54 | 17.15 | 17.35 | 17.16 | 45.37 | 19.9 | 27.03 | 27.37 | 30.17 | 28.6 | + |
| Abhd4 | 25.55 | 20.37 | 19.7 | 33.94 | 37.8 | 35.13 | 40.9 | 34.35 | 34.09 | 39.65 | 44.05 | 39.34 | + |
| Haus3 | 28.09 | 23.44 | 27.05 | 20.64 | 22.46 | 20.79 | 18.94 | 18.41 | 21.57 | 21.68 | 18.89 | 21.52 | + |
| Ccdc107 | 31.84 | 34.6 | 28.96 | 41 | 45.32 | 46.46 | 46.2 | 45.25 | 45.49 | 44.56 | 43.15 | 46.6 | + |
| Ctu2 | 25.71 | 25.54 | 25.94 | 14.05 | 14.42 | 14.32 | 10.63 | 14.83 | 14.37 | 11.35 | 12.01 | 10.57 | + |
| Bid | 27.73 | 29.35 | 27.58 | 17.92 | 16.24 | 17.54 | 14.07 | 17.21 | 17.13 | 15.12 | 12.71 | 12.83 | + |
| Ppp1ca | 456.69 | 494.24 | 470.6 | 320.95 | 312.46 | 315.03 | 426.01 | 377.58 | 402.81 | 353.61 | 337.07 | 371.22 | + |
| Nemf | 27.39 | 25.05 | 25.6 | 18.33 | 21.1 | 21.55 | 18.31 | 22.1 | 20.11 | 18.94 | 19.98 | 17.77 | + |
| Flna | 25.24 | 23.75 | 23.98 | 17.92 | 16.79 | 16.42 | 15.86 | 18.8 | 16.39 | 16.46 | 14.05 | 14.94 | + |
| Tmem203 | 8.68 | 10.12 | 9.11 | 17.5 | 15.98 | 15.84 | 21.38 | 18.54 | 18.96 | 16.45 | 17.12 | 19.68 | + |
| Susd3 | 38.06 | 38.17 | 33.39 | 74.69 | 68.81 | 72.42 | 85.84 | 66.23 | 75.58 | 92.35 | 81.65 | 95.42 | + |
| 2810403D21Rik | 0.74 | 0.64 | 0.81 | 2.64 | 1.24 | 2.54 | 2.17 | 1.69 | 2.4 | 2.16 | 2.21 | 2.01 | + |
| Pddc1 | 21.7 | 21.84 | 18.34 | 35.34 | 31.06 | 34.87 | 41.53 | 32.24 | 33.87 | 44.63 | 46.35 | 41.24 | + |
| Skp1a | 121.96 | 117.86 | 126.14 | 62.25 | 66.59 | 70.09 | 49.56 | 64.11 | 60.29 | 47.65 | 50.33 | 48.21 | + |
| Ell | 9.27 | 10.26 | 9.57 | 14.01 | 14.49 | 12.65 | 12.39 | 12.12 | 11.9 | 13.15 | 12.94 | 14.09 | + |
| Plbd2 | 7.27 | 9.48 | 12.24 | 18.3 | 18.58 | 17.22 | 16.14 | 19.31 | 17.4 | 20 | 18.45 | 16.02 | + |
| Exo1 | 10.37 | 9.74 | 9.27 | 3.77 | 4.06 | 4.02 | 1.91 | 1.7 | 2.02 | 1.01 | 0.85 | 1.18 | + |
| Mtfp1 | 6.67 | 6.92 | 7.63 | 2.86 | 1.58 | 2.51 | 1.61 | 2.69 | 2.57 | 1.07 | 2.47 | 1.53 | + |
| Plaa | 28.61 | 27.62 | 27.22 | 21.69 | 24.17 | 23.45 | 22.58 | 23.13 | 22.81 | 22.94 | 25.87 | 22.82 | + |
| Mrps35 | 85.24 | 83.17 | 85.11 | 69.84 | 73.05 | 73.16 | 74.91 | 78.15 | 76.61 | 65.36 | 69.83 | 75.42 | + |
| Hp1bp3 | 44.83 | 47.31 | 46.31 | 58.38 | 57.81 | 57.98 | 65.9 | 60.12 | 61.12 | 61.98 | 67.18 | 65.45 | + |
| Abhd10 | 16.23 | 15.55 | 13.76 | 9.56 | 10.29 | 9.82 | 9.35 | 11.16 | 8.76 | 8.43 | 9.08 | 7.9 | + |
| Nfil3 | 14.04 | 19.83 | 22.52 | 51.37 | 54.74 | 51.17 | 56.97 | 70.42 | 74.98 | 85.57 | 96.7 | 65.49 | + |
| Ikzf4 | 11.18 | 10.79 | 10.44 | 13.36 | 12.57 | 11.25 | 5.54 | 7.78 | 7.43 | 2.38 | 3.53 | 2.53 | + |
| Mtss1 | 2.65 | 1.91 | 2.38 | 3.44 | 3.72 | 3.71 | 4 | 3.94 | 4.23 | 3.88 | 4.28 | 3.92 | + |
| Hif3a | 0.4 | 0.59 | 0.45 | 2.15 | 1.87 | 1.44 | 0.72 | 0.67 | 0.71 | 1.13 | 0.9 | 1.1 | + |
| Egln3 | 19.93 | 20.2 | 23.64 | 44.36 | 47.87 | 48.07 | 79.54 | 45.35 | 64.04 | 41.21 | 50.87 | 65.14 | + |
| Zfp775 | 4.38 | 3.76 | 4.98 | 8.03 | 6.54 | 7.7 | 7.88 | 7.18 | 7.47 | 8.25 | 8.35 | 9.2 | + |
| Wdr74 | 87.25 | 83.46 | 85.2 | 36.01 | 42.67 | 41.06 | 33.08 | 49.95 | 42.29 | 29.03 | 26.11 | 31.12 | + |
| Mapk1ip1 | 6.44 | 4.77 | 5.57 | 11.11 | 11.92 | 11.8 | 13.39 | 10.14 | 9.51 | 11.04 | 10.35 | 9.8 | + |
| Cd47 | 286.82 | 313.8 | 279.69 | 387.95 | 391.06 | 388.28 | 467.53 | 459.38 | 466.08 | 384.99 | 407.02 | 385.06 | + |
| Ncbp2 | 36.1 | 41.13 | 39.86 | 17.29 | 17.34 | 19.31 | 19.07 | 21.09 | 18.65 | 15.25 | 11.89 | 12.31 | + |
| Fbxo38 | 18.96 | 15.9 | 17.33 | 28.34 | 26.91 | 25.71 | 28.45 | 25.91 | 25.34 | 30.02 | 28.05 | 28.27 | + |
| Usp16 | 22.96 | 24.06 | 25.82 | 15.89 | 16.29 | 18.14 | 13.44 | 17.76 | 15.76 | 14.46 | 16.33 | 14.74 | + |

| | | | | | | | | | | | | | |
|---|---|---|---|---|---|---|---|---|---|---|---|---|---|
| Uvssa | 2.41 | 2.18 | 2.08 | 3.76 | 4.63 | 4.33 | 3.49 | 3.16 | 3.64 | 4.83 | 3.5 | 4.25 | + |
| Slc37a2 | 4.99 | 2.95 | 4.06 | 1.2 | 1.6 | 1.54 | 1.93 | 2.41 | 1.31 | 2.7 | 1.28 | 1.88 | + |
| Rad17 | 16.18 | 17.13 | 15.35 | 10.54 | 12.73 | 12.17 | 10.06 | 13.78 | 11.58 | 9.39 | 11.84 | 10.98 | + |
| Mcm5 | 105.58 | 111.28 | 109.02 | 46.8 | 52.87 | 43.28 | 33.63 | 36.91 | 33.22 | 21.67 | 24.1 | 23.48 | + |
| Ppm1g | 153.64 | 156.72 | 156.36 | 100.66 | 103.94 | 106.57 | 93.85 | 107.24 | 103.58 | 99.74 | 99.4 | 92.98 | + |
| Zfp324 | 2.92 | 2.85 | 2.27 | 1.5 | 1.19 | 1.59 | 0.83 | 1.24 | 0.99 | 1.12 | 1.14 | 1.16 | + |
| Msh2 | 26.79 | 28.4 | 28.27 | 21.24 | 23.37 | 20.09 | 21.02 | 24.04 | 19.87 | 19.84 | 19.34 | 17.84 | + |
| Tcf7 | 9.93 | 8.28 | 11.45 | 9.54 | 8.42 | 8.72 | 5.37 | 7.39 | 5.57 | 1.62 | 1.39 | 3.32 | + |
| Psmb2 | 397.06 | 420.37 | 391.04 | 234.97 | 247.56 | 262.86 | 238.98 | 284.24 | 264.87 | 213.49 | 203.49 | 235.13 | + |
| Tmem59 | 60.27 | 60.77 | 60.08 | 142.47 | 126.3 | 135.77 | 132.98 | 107.04 | 110.81 | 157.28 | 171.87 | 167.78 | + |
| Ddx49 | 35.64 | 38.99 | 40.32 | 29.46 | 29.02 | 28.86 | 27.24 | 28.77 | 28.68 | 28.11 | 28.66 | 26.29 | + |
| Dhx38 | 16.43 | 16.02 | 14.98 | 10.54 | 10.96 | 7.82 | 11.92 | 11.75 | 11.53 | 7.75 | 9.62 | 8.78 | + |
| Fam102a | 5.18 | 6.16 | 4.35 | 24.54 | 17.04 | 17.74 | 15.82 | 7.26 | 14.29 | 10.51 | 12.2 | 16.73 | + |
| Rala | 9.31 | 13.29 | 13.4 | 20.78 | 21.15 | 20.6 | 28.73 | 27.47 | 27.71 | 27.7 | 31.14 | 26.44 | + |
| Impa2 | 53.55 | 64.01 | 66.94 | 153.86 | 140.9 | 155.44 | 223.09 | 130.79 | 150.76 | 181.14 | 195.66 | 219.96 | + |
| Dek | 56.39 | 86.16 | 82.99 | 41.61 | 49.16 | 39.08 | 30.7 | 32.06 | 30.97 | 31.57 | 27.84 | 26.29 | + |
| Hint1 | 418.39 | 438.68 | 416.65 | 298.45 | 318.3 | 338.61 | 310.1 | 333.29 | 322.17 | 284.17 | 306.06 | 287.29 | + |
| Impa1 | 49.88 | 51.82 | 51.39 | 26.97 | 28.12 | 29.41 | 22.96 | 29.02 | 26.84 | 23.23 | 21.66 | 24.3 | + |
| Farp1 | 0.99 | 0.84 | 1.01 | 1.52 | 1.57 | 2.45 | 1.88 | 1.62 | 1.67 | 1.41 | 1.69 | 1.78 | + |
| Ccbl2 | 14.55 | 10.54 | 11.41 | 2.53 | 2.28 | 1.79 | 1.69 | 4.49 | 4.3 | 1.3 | 1.34 | 1.72 | + |
| Spn | 13.47 | 14.83 | 13.28 | 10.59 | 8.86 | 10.3 | 10.74 | 9.86 | 9.42 | 11.24 | 10.19 | 11.61 | + |
| Adk | 35.82 | 34.48 | 35.02 | 15.39 | 16.92 | 17.31 | 18.27 | 21.72 | 20.68 | 14 | 13.06 | 14.15 | + |
| Ncln | 39.66 | 40.28 | 40.49 | 22.15 | 20.64 | 21.18 | 14.79 | 19.2 | 19.27 | 14.33 | 13.53 | 14.54 | + |
| Mtpn | 92.3 | 94.04 | 97.85 | 64.15 | 64.13 | 61.03 | 69.8 | 71.29 | 76.01 | 63.73 | 56.72 | 57.45 | + |
| Suv420h2 | 10.39 | 9.07 | 8.45 | 17.92 | 18.43 | 17.61 | 21.52 | 15.49 | 16.7 | 18.07 | 21.61 | 19.82 | + |
| Tpt1 | 1122.76 | 1160.49 | 1154.63 | 1991.2 | 1878.74 | 1922.83 | 1845.29 | 1720.01 | 1681.67 | 2127.93 | 2233.71 | 2269.83 | + |
| D230025D16Rik | 8.47 | 8.39 | 8.99 | 12.19 | 12.46 | 11.66 | 11.81 | 11.89 | 11.64 | 11.88 | 10.51 | 12.8 | + |
| Cyp2s1 | 0.87 | 0.84 | 0.89 | 5.35 | 5.95 | 5.57 | 8.73 | 5.51 | 6.02 | 8.25 | 8.25 | 7.74 | + |
| Mns1 | 10.89 | 11.34 | 12.61 | 5.13 | 6.05 | 5.86 | 9.03 | 5.63 | 5.53 | 5.8 | 6.89 | 4.97 | + |
| Gpr174 | 3.84 | 3.81 | 4.34 | 3.46 | 3.16 | 3.55 | 5.78 | 5.13 | 4.76 | 5.78 | 5.51 | 6.6 | + |
| Mpv17 | 53.85 | 57.9 | 52.26 | 38.04 | 39.84 | 40.71 | 42.7 | 38.98 | 41.1 | 37.91 | 39.43 | 36.32 | + |
| Abhd14b | 5.38 | 5.63 | 6.34 | 14.44 | 12.57 | 13.14 | 23.34 | 19.14 | 19.31 | 23.62 | 23.82 | 23.68 | + |
| Phrf1 | 17.26 | 19.78 | 18.86 | 15.21 | 15.47 | 15.3 | 14.72 | 14.32 | 14.36 | 13.07 | 14.48 | 14.43 | + |
| Prpsap2 | 22 | 24.73 | 21.69 | 14.7 | 17.82 | 17.64 | 14.54 | 17.1 | 14.79 | 13.93 | 19.78 | 14.38 | + |
| Bop1 | 100.13 | 99.48 | 98.77 | 36.43 | 42.6 | 42.42 | 28.29 | 47.59 | 45.05 | 31.65 | 28.93 | 28.74 | + |
| Akt1 | 20.85 | 17.01 | 17.79 | 11.33 | 9.42 | 10.84 | 11.82 | 9.76 | 10.29 | 10.68 | 9.26 | 9.71 | + |
| Rps28 | 1190.21 | 1241.09 | 1213.12 | 1788.53 | 1871.77 | 1880.66 | 1764.6 | 1585.09 | 1494.85 | 1823.29 | 1956.59 | 1852.32 | + |
| Rdm1 | 16.43 | 17.38 | 15.32 | 36.49 | 37.47 | 37.11 | 35.07 | 39.48 | 35.23 | 35.76 | 43.11 | 35.18 | + |
| Cpsf3 | 75.26 | 71.1 | 78.52 | 45.06 | 49.25 | 47.55 | 43.57 | 46.78 | 46.96 | 32.36 | 31.34 | 31.86 | + |
| Nt5c2 | 9.91 | 10.47 | 10.49 | 4.83 | 5.42 | 6.45 | 6.28 | 6.95 | 5.3 | 4.89 | 4.74 | 4.39 | + |
| Hadh | 47.66 | 47.8 | 44.95 | 61.07 | 62.11 | 59.14 | 94.48 | 72.97 | 73.76 | 73.98 | 76.57 | 75.87 | + |
| Utp15 | 30.37 | 29.73 | 31.39 | 18.71 | 18.69 | 19.6 | 13.58 | 19.01 | 17.04 | 13.81 | 12.94 | 14.28 | + |
| Pepd | 26.04 | 24.84 | 23.31 | 18.45 | 16.91 | 17.45 | 21.34 | 20.19 | 20.61 | 16.76 | 17.23 | 19.19 | + |
| 2310011J03Rik | 31.93 | 33.39 | 34.38 | 24.87 | 22.12 | 21.87 | 24.7 | 22.69 | 21.96 | 18.83 | 19.58 | 19.98 | + |
| Rftn2 | 0.47 | 0.4 | 0.32 | 1.05 | 0.68 | 0.66 | 0.73 | 1.2 | 0.92 | 1 | 1.16 | 1.05 | + |
| Gipc1 | 50.73 | 41.77 | 47.04 | 33.83 | 36.34 | 36.54 | 34.85 | 36.09 | 33.5 | 30.9 | 28.45 | 30.82 | + |
| Exosc3 | 77.49 | 73.78 | 77.86 | 33.44 | 41.7 | 36.33 | 32.84 | 44.31 | 38.44 | 30.21 | 30.63 | 32.95 | + |
| Amn1 | 8.23 | 6.12 | 5.74 | 9.48 | 9.42 | 9 | 12.65 | 11.84 | 13.65 | 10.66 | 10.26 | 10.1 | + |
| Rcl1 | 57.3 | 64.25 | 60.94 | 34.07 | 34.79 | 34.84 | 27.54 | 36.83 | 33.42 | 27.25 | 29.67 | 24.91 | + |
| Parp6 | 2.59 | 2.84 | 2.66 | 9.14 | 6.15 | 7.37 | 9.07 | 7.19 | 7.21 | 10.6 | 13.83 | 11.68 | + |
| Afg3l1 | 15.73 | 17.49 | 17.9 | 10.47 | 12.06 | 10.64 | 13.03 | 13.2 | 14.15 | 11.42 | 9.83 | 10.76 | + |
| Rps6kc1 | 4.22 | 3.67 | 3.76 | 2.48 | 2.89 | 2.67 | 2.19 | 2.79 | 2.48 | 2.13 | 2.48 | 2.01 | + |
| Gfm1 | 55.4 | 55.32 | 56.73 | 26.71 | 29.4 | 27.61 | 26.94 | 32.4 | 34.25 | 23.67 | 28.72 | 21.17 | + |
| Cyp20a1 | 20.91 | 18.8 | 19.49 | 13.89 | 14.68 | 15.3 | 10.49 | 12.33 | 12.58 | 10.86 | 10.14 | 12.94 | + |
| Dopey2 | 1.3 | 1.25 | 1.37 | 2.42 | 2.68 | 2.43 | 3.08 | 3.01 | 2.58 | 2.46 | 2.62 | 3.08 | + |
| Gm19757 | 2.19 | 2.22 | 2.14 | 3.7 | 3.95 | 4.69 | 5.78 | 4.99 | 5.47 | 5.97 | 7.27 | 5.59 | + |
| Cog7 | 14.86 | 12.63 | 15.61 | 8.39 | 10.27 | 11.01 | 9.95 | 9.72 | 8.7 | 8.47 | 8.5 | 9.46 | + |
| Mrps24 | 64.12 | 71.67 | 65.02 | 43.69 | 48.85 | 46.34 | 53.65 | 52.02 | 47.26 | 43.23 | 46.02 | 42.76 | + |
| Tmem176a | 2.1 | 3.02 | 1.07 | 17.98 | 7.17 | 6.85 | 9.75 | 2.82 | 5.26 | 7.6 | 6.79 | 10.21 | + |
| Dnajc14 | 13.95 | 14.12 | 14.31 | 11.31 | 11.37 | 11.68 | 10.7 | 11.52 | 11.41 | 10.58 | 10.83 | 11.33 | + |
| Safb | 26.47 | 27.35 | 25.88 | 17.87 | 20.36 | 18.15 | 17.39 | 17.31 | 17.04 | 18.33 | 17.97 | 17.93 | + |
| Emc6 | 65.25 | 63.64 | 65.9 | 47.84 | 46.4 | 45.35 | 48.59 | 55.26 | 51.03 | 43.68 | 46.38 | 43.97 | + |
| Aurkaip1 | 77.87 | 83.55 | 75.19 | 63.41 | 61.29 | 67.83 | 67.26 | 67.77 | 67.85 | 61.26 | 64.33 | 58.77 | + |
| Tmem216 | 16.62 | 15.18 | 14.55 | 9.21 | 11.66 | 11.69 | 8.07 | 9.75 | 10.13 | 11.17 | 9.28 | 11.83 | + |
| Papola | 42.98 | 43.1 | 46.04 | 27.33 | 29.69 | 26.86 | 27.64 | 26.24 | 27.74 | 23.58 | 25.41 | 24.65 | + |
| Pold2 | 92.12 | 90.46 | 95.67 | 55.43 | 57.19 | 54.78 | 43.73 | 51 | 48.62 | 36.77 | 37.47 | 35.57 | + |
| Tuba4a | 103.87 | 105.23 | 94.88 | 30.21 | 29.45 | 32.54 | 30.95 | 31.96 | 32.53 | 21.19 | 17.85 | 22.82 | + |
| Ffar2 | 0.12 | 0.05 | 0.21 | 0.55 | 1.12 | 1.36 | 1.23 | 0.49 | 1.54 | 0.85 | 0.97 | 2.69 | + |
| Emc2 | 69.96 | 62.01 | 65.27 | 61.85 | 59.87 | 61.88 | 54.97 | 56.85 | 53.15 | 53.39 | 54.36 | 55.19 | + |
| Pde7a | 7.33 | 6.1 | 5.93 | 3.33 | 3.86 | 4.7 | 3.83 | 4.35 | 4.83 | 4.33 | 4.6 | 4.05 | + |
| Ttc12 | 0.11 | 0.18 | 0.22 | 0.59 | 0.38 | 0.42 | 0.76 | 0.6 | 0.43 | 0.8 | 0.63 | 0.62 | + |
| Kpna6 | 4.61 | 5.13 | 4.78 | 3.73 | 3.57 | 4 | 3.22 | 3.78 | 3.39 | 3.5 | 3.99 | 2.81 | + |
| Zfp207 | 152.91 | 164.55 | 154.9 | 115.56 | 115.53 | 117.3 | 126.71 | 125.8 | 111.57 | 121.67 | 128.96 | 125.34 | + |
| Wdr77 | 46.54 | 48.66 | 49.05 | 22.46 | 23.92 | 24.93 | 17.03 | 25 | 22.4 | 14.8 | 14 | 15.56 | + |
| Rab10 | 13.15 | 14.13 | 14.59 | 18.14 | 15.49 | 16.77 | 15.61 | 14.68 | 14.98 | 21.32 | 21.85 | 20.6 | + |
| Dnajb2 | 4.13 | 5.14 | 7.01 | 10.67 | 9.17 | 10.12 | 7.6 | 9.77 | 10.67 | 8.21 | 9.83 | 9.92 | + |
| Coq10a | 2.23 | 2.09 | 2.24 | 4.61 | 4.01 | 5.49 | 3.74 | 3.99 | 3.57 | 4.34 | 4.68 | 4.39 | + |
| Dpm3 | 22.47 | 25.34 | 23.02 | 34.89 | 36.77 | 31.1 | 44.09 | 37.51 | 40.5 | 43.75 | 38.14 | 44.94 | + |
| Cwf19l1 | 8.08 | 7.05 | 7.41 | 5.36 | 4.97 | 6.09 | 4.57 | 5.4 | 4.4 | 4.55 | 4.78 | 5.83 | + |
| Sun2 | 4.84 | 5.87 | 4.97 | 12.71 | 10.58 | 10.7 | 12.8 | 11.68 | 11.74 | 17.55 | 14.51 | 16.37 | + |
| Dom3z | 13.92 | 15.46 | 14.23 | 9.63 | 10.73 | 9.94 | 8.52 | 11.45 | 8.95 | 9.26 | 10.2 | 11.68 | + |
| Adck3 | 3.78 | 3.69 | 2.61 | 8.99 | 6.47 | 7.24 | 16.17 | 13.38 | 12.2 | 14.13 | 15.43 | 15.63 | + |
| Limk2 | 13.61 | 12.99 | 14 | 27.88 | 22.15 | 22.39 | 25.39 | 21.93 | 22.37 | 29.21 | 33.06 | 28.6 | + |
| Dok2 | 45.83 | 54.73 | 39.49 | 28.23 | 25.27 | 24.29 | 29.72 | 31.61 | 31.3 | 32.23 | 24.01 | 30.99 | + |
| Trappc13 | 30.67 | 26.28 | 28.16 | 15.34 | 15.88 | 16.78 | 13.36 | 15.28 | 15.85 | 12.81 | 15.5 | 12.45 | + |
| Zbtb42 | 0.49 | 0.77 | 0.62 | 2.42 | 1.76 | 1.94 | 2.84 | 2.48 | 2.98 | 3.78 | 3.46 | 3.6 | + |
| L1cam | 0.51 | 0.62 | 0.41 | 0.82 | 1.02 | 0.66 | 0.9 | 1.18 | 1 | 0.86 | 0.98 | 1.19 | + |
| Cenpm | 17.93 | 20.52 | 17.75 | 16.03 | 20.4 | 20.26 | 11.8 | 13.05 | 12.88 | 8.8 | 9.73 | 8.96 | + |
| Tti1 | 9.55 | 8.81 | 10.54 | 6.79 | 7.31 | 6.88 | 4.97 | 6.01 | 5.49 | 5.89 | 4.57 | 5.24 | + |
| Magoh | 110.69 | 103.46 | 95.95 | 39.82 | 46.3 | 48.46 | 39.7 | 44.24 | 40.94 | 36.22 | 36.32 | 36.22 | + |
| Cops2 | 45.07 | 41.52 | 46.06 | 47.62 | 43.01 | 43.44 | 46.75 | 37.99 | 39.78 | 53.68 | 59.64 | 48.91 | + |
| Dlg4 | 1.03 | 0.77 | 1.11 | 3.42 | 2.18 | 2.33 | 3.51 | 2.35 | 2.82 | 3.97 | 3.43 | 3.95 | + |
| Glmn | 19.79 | 17.75 | 19.13 | 15.4 | 13.86 | 11.92 | 15.47 | 17.3 | 14.15 | 14.52 | 14.33 | 15.09 | + |
| C1galt1c1 | 10.46 | 11.89 | 12.5 | 8.55 | 7.75 | 6.98 | 6.25 | 7.91 | 6.94 | 6.61 | 6.3 | 7.15 | + |
| Psmd12 | 150.78 | 158.86 | 155.92 | 78.61 | 85.86 | 88.06 | 70.22 | 90.27 | 80.41 | 67.93 | 66.63 | 66.17 | + |
| Tmem126a | 64 | 66.39 | 66.7 | 45.71 | 47.51 | 42.29 | 44.26 | 55.66 | 47.47 | 40.7 | 39.34 | 37.17 | + |
| Gm16702 | 21.26 | 23.06 | 20.76 | 11.11 | 11.44 | 11.21 | 13.08 | 14.45 | 11.64 | 12.09 | 12.6 | 10.93 | + |
| Lpar5 | 7.48 | 6.19 | 7.58 | 3.52 | 3.91 | 4.63 | 5.22 | 6.41 | 3.4 | 4.67 | 3.69 | 4.59 | + |
| Tmem41a | 16.39 | 16.62 | 15.69 | 9.96 | 10.87 | 10.34 | 6.89 | 10.36 | 11.57 | 9.4 | 11.15 | 9.95 | + |
| Fam132a | 18.84 | 18.15 | 23.08 | 96.68 | 87.75 | 91.09 | 68.38 | 52.42 | 58.97 | 65.74 | 76.09 | 71.96 | + |
| Whsc1l1 | 17.26 | 18.15 | 15.79 | 33.63 | 28.24 | 27.46 | 32.84 | 25.34 | 28.46 | 36.26 | 34.16 | 33.35 | + |
| Fam188a | 18.89 | 20.9 | 17.36 | 10.1 | 11.42 | 11.87 | 9.68 | 10.7 | 9.13 | 11.08 | 10.88 | 11.41 | + |
| Cr1l | 46.44 | 51.61 | 45.22 | 34.27 | 34.14 | 38.49 | 30.64 | 39.89 | 35.71 | 32.07 | 31.56 | 35.72 | + |
| Fam96a | 87.88 | 89.97 | 91.32 | 66.94 | 65.06 | 75.53 | 72.56 | 67.08 | 69.4 | 63.2 | 77.47 | 73.28 | + |
| Crebl2 | 0.11 | 0.18 | 0.15 | 1.15 | 0.62 | 1.18 | 0.29 | 0.44 | 0.83 | 0.45 | 0.78 | 1.11 | + |
| Ddrgk1 | 35.62 | 32.8 | 32.41 | 51.38 | 50.2 | 52.72 | 49.54 | 49.93 | 49.31 | 54.56 | 58.09 | 57.56 | + |
| Ube2e1 | 110.18 | 106.37 | 101.35 | 71.29 | 72.82 | 74.03 | 67.85 | 78.12 | 78.98 | 62.73 | 65.75 | 61.6 | + |
| Eif5 | 90.72 | 94.52 | 88.33 | 60.69 | 61.43 | 59.22 | 51.71 | 61.09 | 59.52 | 53.02 | 51.45 | 51.4 | + |
| Hif1a | 155.07 | 166.11 | 181.59 | 257.02 | 261.18 | 250.8 | 178.83 | 197.4 | 214.68 | 228.11 | 218.74 | 217.7 | + |
| Metap1d | 4.08 | 4.02 | 3.56 | 7.18 | 6.5 | 6.13 | 6.71 | 6.34 | 6.26 | 7.36 | 8.47 | 6.78 | + |
| Hnrnpd | 2.15 | 2.54 | 2.07 | 1.98 | 1.32 | 1.19 | 1.27 | 1.33 | 1.05 | 1.29 | 1.18 | 1.39 | + |
| Becn1 | 66.3 | 58.6 | 62.4 | 46.37 | 46.09 | 43.33 | 47.29 | 49.04 | 48.83 | 41.38 | 38.07 | 43.9 | + |
| Stmn1 | 193.01 | 212.55 | 195.87 | 92.35 | 115.07 | 110.82 | 70.79 | 97.51 | 85.13 | 56.45 | 56.57 | 51.71 | + |
| Farsa | 93.02 | 94.27 | 94.69 | 46.96 | 52.04 | 53.67 | 43.8 | 57.04 | 50.01 | 42.54 | 35.04 | 40.34 | + |
| Mtmr7 | 5.94 | 3.65 | 4.31 | 7.68 | 7.69 | 8.5 | 2.05 | 4.27 | 3.18 | 2.61 | 3.36 | 2.43 | + |
| St8sia1 | 1.17 | 0.75 | 1.14 | 2.22 | 1.94 | 2.19 | 2.33 | 1.81 | 2.08 | 1.82 | 1.73 | 1.48 | + |
| Zfyve1 | 1.76 | 1.65 | 1.96 | 3.84 | 4 | 4.73 | 4.56 | 3.73 | 4.66 | 5.12 | 6.1 | 6.2 | + |
| Gas8 | 7.31 | 6.93 | 6.63 | 4.1 | 3.9 | 2.73 | 3.49 | 3.39 | 2.91 | 3.68 | 2.93 | 3.77 | + |
| Cib1 | 63.09 | 67.08 | 65.24 | 44.1 | 44.45 | 50.36 | 51.42 | 49.26 | 51.02 | 46.27 | 48.16 | 44.28 | + |
| Pigg | 3.42 | 3.61 | 3.37 | 1.99 | 1.78 | 2.18 | 1.4 | 2.33 | 1.58 | 2.1 | 1.69 | 1.42 | + |
| Casp3 | 96.73 | 101.4 | 97.49 | 39.43 | 38.37 | 39.79 | 97.31 | 91.42 | 91.25 | 52.13 | 58.01 | 55.96 | + |
| Rrn3 | 55.96 | 53.01 | 53.14 | 28.28 | 30.45 | 26.6 | 23.64 | 29.75 | 28.95 | 20.02 | 25.03 | 24.15 | + |
| Ube2h | 8.58 | 9.4 | 7.8 | 21.25 | 17.14 | 18.13 | 30.91 | 20.07 | 22.2 | 29.92 | 27.38 | 31.02 | + |
| Ddit4 | 13.96 | 21.24 | 23.12 | 65.57 | 60.33 | 52.35 | 45.12 | 46.12 | 43.46 | 73.94 | 81.55 | 81.66 | + |
| Hspe1 | 514 | 493.28 | 526.85 | 300.39 | 349.78 | 345.68 | 339.47 | 416.03 | 382.32 | 294.87 | 357.01 | 280.57 | + |
| Dut | 112.7 | 132.52 | 121.87 | 59.85 | 64.44 | 61.7 | 56.19 | 60.46 | 62.43 | 52.93 | 57.67 | 55.21 | + |
| Jmjd4 | 5.84 | 4.89 | 4.91 | 3 | 3.13 | 3.09 | 1.78 | 2.45 | 2.36 | 2.25 | 1.56 | 1.93 | + |
| Zswim7 | 27.79 | 29.09 | 28.11 | 30 | 33.54 | 33.33 | 17.04 | 31.32 | 29.39 | 13.15 | 12.46 | 10.58 | + |
| Trmu | 9.88 | 12.1 | 9.78 | 6.36 | 5.75 | 6.46 | 5.38 | 6.72 | 5.68 | 6.51 | 4.37 | 4.78 | + |
| Phf15 | 4.42 | 3.86 | 4.15 | 3.41 | 3.14 | 3.4 | 2.61 | 3.14 | 2.76 | 3.5 | 2.92 | 2.48 | + |
| Gm3604 | 4.78 | 4.24 | 4.56 | 2.57 | 3.16 | 3.44 | 2.69 | 3 | 3.06 | 2.44 | 2.35 | 2.6 | + |
| Scamp4 | 82.68 | 78.75 | 75.61 | 50.19 | 49.5 | 46.25 | 50.7 | 55.02 | 54.7 | 49.94 | 42.65 | 48.99 | + |
| Zfp954 | 10.62 | 11.19 | 10.02 | 18.05 | 16.48 | 16.12 | 21.17 | 17.58 | 18.21 | 22.38 | 24.38 | 24.04 | + |
| Acsl5 | 106.02 | 108.09 | 105.96 | 61.82 | 72.88 | 71.87 | 61.23 | 80.48 | 70.45 | 53.24 | 58.18 | 54.97 | + |
| Clk3 | 49.55 | 49.7 | 50.39 | 64.16 | 62.32 | 61.14 | 74.7 | 72.27 | 71.7 | 70.7 | 76.63 | 68.83 | + |
| Acly | 141.04 | 135.82 | 150.47 | 97.59 | 102.44 | 101.38 | 82.02 | 89.43 | 87.95 | 71.33 | 77.32 | 70.51 | + |
| Tmem220 | 0.7 | 0.51 | 0.47 | 2.48 | 4.52 | 1.61 | 2.81 | 2.56 | 1.73 | 2.54 | 2.36 | 0.9 | + |
| 1110004E09Rik | 34.95 | 40.13 | 43 | 23.69 | 25.87 | 26.32 | 22.78 | 29.52 | 26.62 | 21.89 | 27.82 | 23.15 | + |
| Lta | 231.71 | 225.14 | 221.66 | 34.36 | 37.05 | 40.58 | 36.12 | 58.18 | 63.34 | 17.95 | 17.31 | 11.3 | + |
| Tmem185b | 8.49 | 8.72 | 9.25 | 5.16 | 4.47 | 4.87 | 3.86 | 3.86 | 3.75 | 4.51 | 2.92 | 4.64 | + |
| Stom | 0.83 | 0.74 | 0.75 | 1.68 | 2.11 | 2.24 | 3.94 | 2.88 | 3.11 | 2.91 | 3.33 | 2.92 | + |
| Rras | 1.03 | 0.92 | 0.69 | 3.94 | 3.1 | 4.11 | 4.56 | 3.54 | 3.4 | 3.87 | 3.13 | 3.35 | + |
| Prkra | 10.65 | 9.66 | 10.19 | 21.31 | 22.3 | 24.05 | 25.7 | 22.85 | 19.77 | 25.47 | 27.64 | 29.19 | + |
| Mrpl10 | 59.26 | 59.04 | 57.41 | 45.68 | 44.88 | 44.82 | 47.41 | 46.15 | 50.86 | 44.89 | 44.27 | 42.08 | + |
| Map1s | 23.35 | 26.21 | 24.81 | 18.88 | 19.45 | 20.88 | 18.11 | 20.5 | 19.73 | 19.5 | 18.69 | 17.05 | + |
| Ptprcap | 216.98 | 215.97 | 198 | 147.92 | 144.33 | 139.76 | 149.84 | 156.92 | 152.39 | 147.28 | 117.87 | 151.46 | + |
| Naa50 | 43.88 | 52.37 | 56.68 | 25.65 | 24.19 | 23.79 | 18.41 | 24.83 | 24.21 | 18.11 | 19.33 | 15.92 | + |
| Ddx31 | 14.07 | 13.43 | 14.7 | 7.9 | 8.43 | 9.79 | 7.89 | 9.57 | 9.38 | 7.72 | 7.44 | 7.18 | + |
| Map4k1 | 41.11 | 40.44 | 41.85 | 36.83 | 35.72 | 31.37 | 30.16 | 33.44 | 33.81 | 33.56 | 31.64 | 31.45 | + |
| Ppp2r1b | 36.78 | 34.83 | 36.61 | 18.89 | 18.37 | 18.69 | 16.29 | 21.9 | 21.92 | 15.46 | 11.57 | 13.02 | + |
| Ddx10 | 35.09 | 33.14 | 32.83 | 25.16 | 26.39 | 25.63 | 25.86 | 30.21 | 30.15 | 26.41 | 30.94 | 26.72 | + |

| | | | | | | | | | | | | |
|---|---|---|---|---|---|---|---|---|---|---|---|---|---|
| Stab1 | 0.87 | 0.68 | 0.83 | 1.8 | 1.88 | 1.93 | 2.13 | 2.5 | 2.75 | 1.95 | 2.07 | 1.72 | + |
| Nup43 | 54.48 | 59.05 | 53.72 | 35.09 | 35.86 | 34.67 | 26.33 | 29.19 | 26.49 | 19.69 | 29 | 23.86 | + |
| Zw10 | 17.82 | 18.79 | 17.55 | 11.41 | 14.42 | 14.92 | 11.87 | 13.38 | 10.32 | 13.58 | 14.34 | 12.08 | + |
| Hspb11 | 31.66 | 27.48 | 29.21 | 59.22 | 51.14 | 57.12 | 61.16 | 51.98 | 63.34 | 48.17 | 52.26 | 56.41 | + |
| Ctla2a | 103.3 | 134.23 | 81.89 | 444.74 | 313.57 | 328.53 | 460.75 | 222.21 | 323.77 | 510.52 | 459.88 | 576.85 | + |
| Nit2 | 28.5 | 27.96 | 27.87 | 46.93 | 42.98 | 48.15 | 49.5 | 45.3 | 44.76 | 54.14 | 57.99 | 57.82 | + |
| Cuedc2 | 53.9 | 50.7 | 52.54 | 41.55 | 39.99 | 43.53 | 44.55 | 50.32 | 45.29 | 40.7 | 41.27 | 40.28 | + |
| Lst1 | 14.41 | 11.33 | 9.49 | 25.22 | 18.76 | 24.83 | 44.62 | 23.14 | 24.89 | 23.79 | 21.63 | 28.41 | + |
| Mylpf | 6.35 | 4.01 | 4.8 | 8.54 | 8.02 | 10.22 | 11.04 | 10.89 | 10.37 | 8.71 | 10.78 | 10.23 | + |
| Irf6 | 1.15 | 0.92 | 1.56 | 2.73 | 2.65 | 2.69 | 3.61 | 3.54 | 3.16 | 3.3 | 2.09 | 2.15 | + |
| Lrrfip2 | 2.05 | 2.71 | 3.42 | 5.37 | 4.43 | 4.63 | 6.02 | 6.29 | 7.25 | 6.02 | 6.12 | 5.04 | + |
| Ssr2 | 166.27 | 168.47 | 161.7 | 79.5 | 82.9 | 89.49 | 81.16 | 102.52 | 90.26 | 72.38 | 73.46 | 65.05 | + |
| E130112N10Rik | 1.06 | 0.95 | 0.48 | 1.73 | 2.46 | 3.01 | 3.96 | 3.3 | 3.06 | 2.22 | 2.48 | 1.92 | + |
| Vdac3 | 165.7 | 166.41 | 162.01 | 124.43 | 130.87 | 126.51 | 121.69 | 143.06 | 133.93 | 108.25 | 121.06 | 115.16 | + |
| Pop4 | 38.93 | 35.35 | 33.26 | 18.58 | 23.34 | 23.27 | 21.63 | 23.71 | 21.61 | 18.9 | 17.62 | 20.04 | + |
| Ywhaz | 458.2 | 483.32 | 473.74 | 303.83 | 318.16 | 323.94 | 332.76 | 340.9 | 346.94 | 307.41 | 299.49 | 316.08 | + |
| Ddost | 190.09 | 191.23 | 183.05 | 107.4 | 121.35 | 115.63 | 103.13 | 117.05 | 113.45 | 87.11 | 88.08 | 91.07 | + |
| Atad1 | 48.56 | 52.96 | 49.62 | 33.82 | 32.56 | 34.44 | 32.87 | 34.76 | 36.22 | 33.05 | 29.22 | 31.83 | + |
| 2810417H13Rik | 132.7 | 136.83 | 127.6 | 40.69 | 48.01 | 41.9 | 27.9 | 28.63 | 30.03 | 17.7 | 20.97 | 18.06 | + |
| Rpl21 | 609.69 | 621.99 | 629.85 | 875.81 | 865.79 | 913.81 | 824.94 | 789.48 | 799.14 | 882.25 | 977.84 | 891.68 | + |
| Tubb5 | 891.54 | 836.88 | 799.23 | 880.4 | 846.38 | 899.8 | 1011.08 | 700.45 | 794.57 | 1194.71 | 1359.28 | 1261.7 | + |
| Fam114a2 | 22.87 | 25.22 | 24.92 | 42.59 | 39.2 | 41.4 | 44.43 | 37.66 | 42.04 | 46.93 | 47.94 | 50.88 | + |
| Nsmce2 | 39.64 | 45.35 | 42.79 | 30.35 | 34.03 | 29.73 | 30.24 | 31.28 | 34.87 | 27.62 | 27.16 | 28.72 | + |
| Ciz1 | 13.55 | 13.54 | 14.01 | 23.86 | 20.92 | 20.78 | 22.14 | 19.85 | 20.81 | 23.7 | 24.87 | 26.55 | + |
| Twf1 | 38.83 | 37.28 | 38.11 | 36.35 | 35.94 | 36.1 | 25.77 | 28.86 | 27.94 | 30.48 | 29.99 | 28.39 | + |
| Denr | 67.39 | 68.78 | 65.74 | 37.41 | 40.84 | 40.53 | 33.33 | 42.03 | 42.5 | 30.76 | 28.96 | 29.93 | + |
| Pilra | 0.73 | 0.73 | 0.93 | 0.31 | 0.27 | 0.34 | 0.24 | 0.35 | 0.19 | 0.28 | 0.28 | 0.35 | + |
| Smap2 | 6.15 | 6.17 | 4.22 | 5.53 | 5.35 | 5.2 | 9.34 | 5.94 | 5.82 | 12.04 | 8.21 | 10.97 | + |
| Eno2 | 2.54 | 1.83 | 2.08 | 6.22 | 5.16 | 6.98 | 4.44 | 3.03 | 3.33 | 3.88 | 4.47 | 5.32 | + |
| Adi1 | 50.31 | 50.42 | 51.66 | 37.11 | 39.79 | 37.81 | 32.59 | 37.03 | 37.19 | 35.27 | 35.21 | 34.1 | + |
| Rbm6 | 23.81 | 18.74 | 20.29 | 18.65 | 17.05 | 18.11 | 15.67 | 15.85 | 15.71 | 15.59 | 15.48 | 16.9 | + |
| Slc9a3r1 | 46.16 | 50.6 | 46.45 | 30.65 | 27.13 | 29.03 | 33.14 | 33.52 | 36.27 | 30.42 | 26.33 | 31.94 | + |
| Rpl28 | 959.28 | 1011.21 | 992.19 | 1439.28 | 1465.28 | 1512.43 | 1393.9 | 1439.11 | 1387.51 | 1312.65 | 1499.14 | 1277.87 | + |
| Ppil1 | 137.45 | 134.29 | 143.22 | 67.78 | 77.99 | 84.51 | 70.31 | 78.75 | 70.12 | 56.36 | 56.86 | 55.94 | + |
| Rara | 4.68 | 4.79 | 4.03 | 12.85 | 12.24 | 13.14 | 8.81 | 7.8 | 9.64 | 9.17 | 10.11 | 11.11 | + |
| Gtf3c6 | 26.08 | 26.13 | 26.16 | 15.85 | 17.01 | 17.94 | 14.37 | 15.69 | 15.46 | 11.95 | 11.25 | 12.77 | + |
| Tbrg4 | 51.28 | 47.56 | 48.74 | 25.87 | 25.61 | 27.08 | 22.57 | 28.51 | 24.82 | 20.43 | 21.87 | 19.13 | + |
| Gtf2h4 | 29.27 | 32.47 | 31.54 | 19.12 | 20.63 | 18.12 | 17.96 | 19.41 | 23.25 | 15.18 | 15.44 | 15.22 | + |
| Lmbrd1 | 4.07 | 3.55 | 4.13 | 7.1 | 6.67 | 6.12 | 6.13 | 5.71 | 6.62 | 7.35 | 8.06 | 7.74 | + |
| Fam103a1 | 50.19 | 53.73 | 45.18 | 57.15 | 48.35 | 52.1 | 61.26 | 57.65 | 62.91 | 73.7 | 73.1 | 70.37 | + |
| Abcf1 | 69.02 | 68.63 | 68.98 | 38.64 | 39.64 | 39.42 | 33.39 | 39.2 | 38.66 | 33.21 | 28.19 | 36.04 | + |
| Mpdu1 | 49.53 | 50.22 | 51.21 | 35.83 | 38.15 | 38.55 | 30.88 | 39.69 | 34.91 | 29.6 | 31.6 | 27.7 | + |
| Srp9 | 103.23 | 110.15 | 103.58 | 83.15 | 82.93 | 89.31 | 84.43 | 81.67 | 88.43 | 71.63 | 84.81 | 79.02 | + |
| Atg12 | 14.78 | 12.68 | 12.58 | 30.39 | 26.99 | 30.38 | 37.17 | 30.76 | 32.06 | 34.73 | 40.83 | 40.23 | + |
| Fbxo5 | 27.85 | 29.37 | 25.57 | 14.51 | 18.55 | 15.77 | 9.39 | 10 | 7.55 | 12.46 | 11.25 | 8.74 | + |
| Ndufb6 | 271.9 | 272.07 | 265.59 | 157.73 | 175.79 | 182.37 | 157.22 | 188.51 | 172.54 | 119.71 | 128.92 | 120.9 | + |
| Hemk1 | 35.67 | 35.94 | 37.36 | 46.26 | 50.93 | 47.33 | 25.22 | 39.13 | 37.61 | 16.82 | 20.03 | 17.05 | + |
| Hdac4 | 3.08 | 3.41 | 2.98 | 10.57 | 8.52 | 10.25 | 5.85 | 4.58 | 5.09 | 10.95 | 10.22 | 10.01 | + |
| Med24 | 20.64 | 20.24 | 21.01 | 12.88 | 12.87 | 14.29 | 12.66 | 13.85 | 13.45 | 10.42 | 10.61 | 12.75 | + |
| Got2 | 136.38 | 127.05 | 132.37 | 88.19 | 94.97 | 95.32 | 92.7 | 103.44 | 94.74 | 80.62 | 82.47 | 80.52 | + |
| Pik3ip1 | 4.38 | 5.14 | 4.6 | 79.72 | 56.64 | 65.44 | 58.23 | 23.65 | 30.01 | 122.5 | 118.84 | 134.4 | + |
| Timm10 | 82.89 | 87.77 | 85.46 | 44.11 | 46.22 | 39.26 | 32.96 | 42.86 | 42.37 | 29.81 | 28.5 | 27.3 | + |
| Sirt6 | 16.68 | 17.28 | 15.86 | 24.54 | 22.76 | 23.11 | 36.69 | 26.54 | 28.56 | 25.61 | 29.05 | 28.43 | + |
| Ankrd46 | 23.2 | 19.94 | 25.53 | 38.61 | 38.81 | 36.9 | 25.88 | 30.2 | 29.17 | 32.73 | 30.81 | 34.35 | + |
| Polr3a | 9.18 | 8.7 | 9.39 | 6.73 | 5.68 | 6.56 | 6.45 | 7.3 | 7.93 | 6.47 | 6.16 | 6.77 | + |
| Rpl3 | 2512.47 | 2493.01 | 2499.64 | 4388.98 | 4208.8 | 4479.83 | 4184.24 | 3947.48 | 4000.72 | 4500.24 | 4925.97 | 4372.17 | + |
| Rbm27 | 6.94 | 6.78 | 7.08 | 4.74 | 4.05 | 5 | 4.41 | 5.08 | 4.51 | 4.93 | 4.48 | 4.08 | + |
| Arid5b | 0.9 | 1.4 | 1.01 | 4.3 | 3.26 | 3.31 | 3.57 | 2.45 | 2.4 | 4.28 | 4.86 | 5.19 | + |
| Pfn2 | 6.75 | 5.4 | 5.35 | 15.61 | 13.68 | 14.16 | 17.41 | 14.07 | 17.63 | 14.56 | 19.37 | 18.58 | + |
| Drg2 | 64.19 | 62.77 | 63.39 | 37.04 | 38.96 | 35.78 | 31.77 | 36.76 | 32.26 | 25.28 | 30.47 | 27.46 | + |
| Lrrc33 | 24.86 | 29.06 | 22.45 | 13.69 | 13.63 | 12.15 | 15.47 | 12.83 | 14.34 | 13.06 | 9.79 | 16.04 | + |
| Usmg5 | 417.7 | 426.43 | 418.73 | 252.02 | 264.33 | 265.61 | 276.97 | 290.53 | 283.35 | 200.92 | 233.51 | 215.02 | + |
| Gm19705 | 0.41 | 0.56 | 0.39 | 7.79 | 6.33 | 4.62 | 1.8 | 2.27 | 1.04 | 6.63 | 9.68 | 7.38 | + |
| Gramd1b | 8.05 | 7.65 | 7.36 | 4.4 | 3.88 | 4.16 | 3.53 | 5.03 | 4.48 | 3.94 | 3.81 | 3.22 | + |
| Trak2 | 2.56 | 2.7 | 2.69 | 1.87 | 1.8 | 2.04 | 2.04 | 1.94 | 1.96 | 1.84 | 2.06 | 1.56 | + |
| Ccdc137 | 20.46 | 21.01 | 21.22 | 11.34 | 12.65 | 11.49 | 9.92 | 9.9 | 12.09 | 9.81 | 11.65 | 11.26 | + |
| Cstf1 | 40.9 | 40.12 | 36.55 | 33.86 | 36.8 | 34.13 | 29.1 | 30.69 | 29.35 | 27.5 | 31.3 | 27.46 | + |
| Nxf1 | 48.64 | 54.92 | 58.2 | 37.67 | 38.1 | 35.91 | 32.48 | 33.53 | 37 | 35.21 | 31.87 | 38.36 | + |
| Ube2s | 14.85 | 11.84 | 13.08 | 7.04 | 8.21 | 6.33 | 4.81 | 6.72 | 6.08 | 5.56 | 5.8 | 5.37 | + |
| Psma5 | 295.85 | 296.81 | 293.5 | 130.94 | 127.86 | 125.99 | 139.96 | 142.82 | 148.61 | 120.58 | 121.72 | 121.48 | + |
| Ipo9 | 13.22 | 13.18 | 13.17 | 8.58 | 8.92 | 8.65 | 7.36 | 8.3 | 8.24 | 6.4 | 7.21 | 7.87 | + |
| Tnfrsf4 | 227.86 | 242.84 | 261.45 | 286.8 | 292.5 | 284.42 | 265.08 | 253.62 | 272.48 | 156.81 | 213.39 | 203.4 | + |
| Bsdc1 | 6.42 | 5.56 | 6.2 | 10.1 | 9.7 | 8.54 | 9.68 | 10.67 | 9.85 | 10.65 | 11.03 | 11.73 | + |
| Slc50a1 | 42.89 | 36.38 | 37.35 | 66.59 | 60.31 | 69.09 | 72.09 | 57.08 | 60.12 | 74.96 | 73.22 | 82.9 | + |
| Atp5g3 | 358.56 | 372.06 | 342.25 | 179.2 | 182.4 | 197.76 | 171.21 | 191.06 | 183.85 | 156.69 | 151.86 | 157.31 | + |
| Sf3b3 | 78.47 | 79.46 | 77.24 | 38.84 | 43.29 | 40.81 | 26.83 | 36.23 | 32.47 | 30.9 | 28.08 | 29.05 | + |
| Pusl1 | 33.34 | 33.47 | 34.28 | 18.19 | 15.48 | 15.95 | 14.64 | 21.89 | 18.01 | 14.38 | 12.25 | 12.51 | + |
| Incenp | 32.59 | 29.67 | 29.98 | 27.3 | 31.49 | 29.71 | 22.24 | 19.94 | 19.89 | 21.44 | 24.5 | 23.25 | + |
| Mcts1 | 102.94 | 100.03 | 104.3 | 64.57 | 71.4 | 67.55 | 65.99 | 77.9 | 74.26 | 65.44 | 68.17 | 59.81 | + |
| Rnf215 | 1.33 | 0.78 | 1.12 | 2.15 | 2.74 | 2.11 | 2.23 | 2.94 | 1.55 | 1.68 | 2.51 | 2.63 | + |
| Kdm5b | 2.65 | 3.06 | 3.17 | 6.85 | 5.6 | 5.8 | 8.56 | 7.18 | 9.04 | 8.63 | 9.26 | 8.73 | + |
| Cd68 | 3.34 | 1.56 | 2.2 | 9.34 | 9.1 | 11.77 | 13.06 | 13.11 | 11.18 | 13.22 | 14.49 | 14.41 | + |
| Zbtbd6 | 1.52 | 1.85 | 1.31 | 0.54 | 0.47 | 0.72 | 0.28 | 0.41 | 0.51 | 0.21 | 0.25 | 0.4 | + |
| Atp13a3 | 4.39 | 4.66 | 4.8 | 2.99 | 2.78 | 3.08 | 3.67 | 3.39 | 3.52 | 2.97 | 3.25 | 2.71 | + |
| Mrpl46 | 58.02 | 54.17 | 54.33 | 35.01 | 29.66 | 31.79 | 31.83 | 34.59 | 34.93 | 28.22 | 24.21 | 24.59 | + |
| Aars | 39.29 | 42.2 | 41.49 | 39.96 | 45.75 | 42.69 | 39.01 | 52.84 | 49.64 | 30.31 | 32.07 | 26.94 | + |
| Mib2 | 5.39 | 5.94 | 6.43 | 12.15 | 12.11 | 12.34 | 11.85 | 15.31 | 13.2 | 13.23 | 8.68 | 9.03 | + |
| Rrp8 | 32.35 | 30.23 | 30.45 | 18.76 | 18.92 | 18.51 | 17.08 | 20.05 | 21.04 | 14.58 | 15.64 | 14.29 | + |
| Zbtb24 | 6.21 | 6.04 | 6.42 | 10.45 | 11.09 | 9.26 | 10.05 | 9.38 | 8.81 | 10.48 | 10.06 | 11.85 | + |
| Dcp2 | 1.89 | 2.34 | 2.16 | 1.42 | 1.2 | 1.37 | 1.13 | 1.37 | 1.27 | 1.09 | 1.51 | 1.5 | + |
| Polr1b | 31.67 | 32.3 | 35.57 | 12.85 | 12.05 | 13.9 | 9.04 | 14.07 | 13.88 | 8.44 | 8.36 | 7.72 | + |
| Fyco1 | 1.76 | 1.52 | 1.81 | 3.62 | 3.01 | 2.9 | 3.56 | 3.57 | 3.03 | 4.68 | 5.09 | 4.52 | + |
| Srsf10 | 72.74 | 75.98 | 75.12 | 37.48 | 37.33 | 35.73 | 26.2 | 33.48 | 32.34 | 26.32 | 25.56 | 23.74 | + |
| Cd37 | 67.81 | 66.55 | 65.66 | 132.7 | 130.96 | 137.85 | 159.58 | 122.79 | 141.3 | 173.02 | 169.36 | 196.31 | + |
| Sec23ip | 26.04 | 24.55 | 25.36 | 14.66 | 15.67 | 15.97 | 12.8 | 15.76 | 14.89 | 14.3 | 15.94 | 12.78 | + |
| Pxmp4 | 6.97 | 5.85 | 6.58 | 12.77 | 11.07 | 13 | 17.46 | 10.26 | 10.95 | 13.56 | 10.8 | 13.83 | + |
| Vps28 | 140.04 | 145.5 | 140.87 | 185.83 | 177.61 | 182.52 | 202.41 | 168.37 | 182.94 | 197.22 | 195.34 | 192.99 | - |
| Pex19 | 20.36 | 21.73 | 17.53 | 16.21 | 13.4 | 14.01 | 14.04 | 13.84 | 13.71 | 9.83 | 10.75 | 11.86 | - |
| Katnbl1 | 20.73 | 22.27 | 22.19 | 13.83 | 16.93 | 17.19 | 10.4 | 14 | 13.94 | 12 | 11.51 | 10.7 | - |
| Capzb | 284.27 | 287.78 | 279.45 | 236.07 | 233.09 | 235.62 | 277.45 | 268.88 | 277.21 | 225.78 | 234.35 | 233.15 | - |
| Ptgr1 | 10.12 | 7.1 | 7.48 | 5.77 | 6.2 | 6.37 | 8.71 | 10.3 | 7.85 | 4.98 | 4.22 | 6.19 | - |
| Lnx2 | 9.39 | 7.71 | 8.03 | 10.13 | 11.1 | 10.45 | 9.11 | 9.57 | 8.85 | 11.41 | 10.46 | 13.33 | - |
| Itgb7 | 36.53 | 58.41 | 60.25 | 134.31 | 107.49 | 106.99 | 175 | 101.13 | 130.65 | 209.46 | 149 | 184.84 | - |
| Rnf115 | 8.97 | 9.02 | 10.55 | 13.03 | 11.75 | 13.82 | 13.22 | 13.17 | 12.41 | 17.81 | 16.5 | 15.42 | - |
| Sowahc | 3.49 | 3.06 | 3.92 | 2.31 | 3.51 | 3 | 1.81 | 2.25 | 2.19 | 1.97 | 1.75 | 1.8 | - |
| Nup188 | 13.73 | 13.61 | 15.68 | 9.37 | 8.73 | 9.03 | 7.97 | 9.91 | 9.73 | 7.1 | 8.62 | 5.91 | - |
| 4930565N06Rik | 0 | 0.1 | 0.18 | 0.76 | 0.36 | 0.79 | 0.36 | 0.39 | 0.31 | 0.64 | 0.97 | 0.49 | - |
| Sod2 | 44.35 | 45.09 | 45.03 | 35.88 | 38.13 | 38.5 | 27.31 | 36.04 | 36.94 | 26.73 | 26.23 | 27.34 | - |
| Il17ra | 12.76 | 12.02 | 11.58 | 22.82 | 20.05 | 20.67 | 26.09 | 14.96 | 17.32 | 33.35 | 27.4 | 35.71 | - |
| Sgk1 | 4.63 | 3.82 | 3.74 | 8.42 | 8.69 | 8.52 | 9.32 | 5.6 | 6.99 | 12.95 | 11.45 | 13.08 | - |
| D430042O09Rik | 0.82 | 0.65 | 0.74 | 1.36 | 1.45 | 1.08 | 0.8 | 1.32 | 0.72 | 1.29 | 1.24 | 1.33 | - |
| Zscan22 | 4.62 | 4.55 | 4.71 | 3.4 | 4.26 | 3.75 | 2.11 | 3.09 | 1.85 | 2.1 | 1.98 | 2.2 | - |
| Ptprk | 3.36 | 3.21 | 4.38 | 2.31 | 2.18 | 2.12 | 3.99 | 2.93 | 3.69 | 1.6 | 2.07 | 1.71 | - |
| Kit | 0.17 | 0.34 | 0.06 | 1.48 | 0.4 | 0.9 | 1.83 | 0.49 | 0.85 | 1.21 | 0.49 | 1.44 | - |
| Gata1 | 0.24 | 0.98 | 0.43 | 0.66 | 0.53 | 1.17 | 0.36 | 0.34 | 0.34 | 0.06 | 0 | 0.12 | - |
| Atp7b | 0.11 | 0.11 | 0.09 | 0.93 | 1.09 | 0.85 | 0.7 | 0.15 | 0.38 | 1.2 | 1.32 | 1.44 | - |
| 4931428F04Rik | 0.61 | 1.35 | 0.89 | 1.69 | 1.02 | 1.31 | 2.33 | 1.24 | 2.04 | 2.27 | 2.03 | 3.14 | - |
| Rbbp6 | 6.34 | 6.81 | 7.04 | 8.02 | 7.81 | 8.32 | 6.67 | 7.02 | 6.8 | 8.87 | 8.85 | 7.45 | - |
| Fam117b | 2.33 | 1.91 | 1.87 | 5.76 | 4.2 | 5.25 | 5.5 | 3.73 | 4.68 | 10.02 | 7.74 | 9.69 | - |
| Crabp2 | 77.48 | 47.71 | 68.07 | 16.78 | 13.87 | 20.62 | 7.47 | 27.67 | 18.36 | 5.09 | 2.65 | 4.52 | - |
| Mlf2 | 47.22 | 56.98 | 65.72 | 44.69 | 46.84 | 45.42 | 43.94 | 50.89 | 45.64 | 45.51 | 43.94 | 40.69 | - |
| Agpat3 | 26.15 | 27.56 | 26.77 | 18.16 | 21.69 | 19.42 | 19.58 | 21.98 | 23.8 | 14.13 | 14.03 | 14.36 | - |
| Zfp362 | 1.19 | 1.45 | 0.66 | 1.64 | 2.36 | 2.12 | 1.67 | 1.81 | 1.62 | 2.98 | 2.03 | 2.6 | - |
| Nmral1 | 15.67 | 15.5 | 12.6 | 12.36 | 10.86 | 11.21 | 10.64 | 9.61 | 9.61 | 6.8 | 7.55 | 9.93 | - |
| Vmn1r58 | 12.46 | 12.26 | 12.15 | 21.02 | 18.14 | 17.59 | 15.96 | 17.64 | 13.88 | 21.25 | 21.69 | 19.73 | - |
| Rpl27 | 962.93 | 1018.11 | 978.7 | 1179.44 | 1166.32 | 1217.53 | 1265.91 | 1181.86 | 1238.3 | 1243.2 | 1354.44 | 1278.1 | - |
| Tecr | 211.58 | 218.83 | 220.7 | 214.33 | 214.48 | 233.81 | 206.69 | 214.7 | 213.66 | 179.08 | 193.07 | 179.84 | - |
| Ndufb3 | 144.92 | 155.81 | 142.6 | 108.01 | 108.66 | 122.27 | 121.41 | 126.06 | 118.24 | 101.29 | 92.24 | 96.09 | - |
| Gm5069 | 0.92 | 1.13 | 1.1 | 1.53 | 1.44 | 1.26 | 2.18 | 1.52 | 1.76 | 1.73 | 1.71 | 2.11 | - |
| Bcat1 | 29.41 | 27.68 | 29.41 | 14.62 | 18.83 | 18.3 | 14.96 | 23.61 | 19.92 | 10.91 | 13.88 | 9.22 | - |
| Ankrd49 | 18.48 | 17.62 | 15.48 | 13.71 | 14.54 | 13.85 | 10.62 | 12.75 | 12 | 10.93 | 9.83 | 11.11 | - |
| Mboat7 | 14.37 | 14.34 | 18.72 | 20.01 | 18.14 | 17.22 | 19.97 | 18.8 | 19.32 | 19.05 | 21.21 | 21.92 | - |
| Ppm1h | 4.92 | 5.09 | 3.8 | 6.38 | 3.89 | 4.54 | 6.49 | 4.23 | 4.23 | 7.87 | 5.6 | 7.56 | - |
| Tars2 | 20.47 | 23.59 | 18.38 | 29.27 | 27.13 | 25 | 16.66 | 20.73 | 20.05 | 26.59 | 26.82 | 26.98 | - |
| Gtse1 | 33.36 | 24.45 | 27.5 | 20.58 | 28.32 | 23.48 | 19.63 | 21.91 | 18.11 | 18.78 | 22.01 | 15.32 | - |
| Efhc1 | 0 | 0.11 | 0.02 | 0.14 | 0.82 | 0.4 | 0.42 | 0.45 | 0.27 | 0.83 | 0.47 | 0.68 | - |
| Sfxn2 | 4.48 | 3.98 | 4.13 | 2.17 | 2.43 | 1.85 | 1.04 | 2.87 | 1.72 | 1.44 | 1.48 | 1.44 | - |
| Fam89b | 23.75 | 23.78 | 25.85 | 30.38 | 30.73 | 30.57 | 37.39 | 30.92 | 34.14 | 37.71 | 41.99 | 36.65 | - |
| Rbm43 | 12 | 13.65 | 10.89 | 15.12 | 13.74 | 13.31 | 15.77 | 13.69 | 12 | 15.5 | 16.42 | 16.55 | - |
| Nufip2 | 6.35 | 7.08 | 6.54 | 9.51 | 9.03 | 9.1 | 7.56 | 8.25 | 8.88 | 12.55 | 12.42 | 11.21 | - |
| Nelfa | 28.68 | 31.95 | 30.72 | 23.46 | 24.6 | 27.61 | 22.13 | 25.9 | 24.98 | 23.4 | 21.85 | 19.82 | - |
| Alg1 | 17.25 | 18.37 | 17.08 | 11.58 | 14.8 | 14.26 | 10.95 | 15.48 | 12.99 | 14.21 | 10.01 | 13.57 | - |
| Tmem180 | 26.09 | 25.91 | 25.98 | 20.57 | 17.85 | 19.8 | 20.52 | 26.3 | 24.64 | 12.55 | 17.52 | 12.92 | - |
| Dusp7 | 0.62 | 1.14 | 0.93 | 2.62 | 2.39 | 2.44 | 1.68 | 0.75 | 1.42 | 2.19 | 2.38 | 1.85 | - |
| Zfp760 | 1.62 | 1.8 | 1.89 | 2.59 | 2.64 | 1.95 | 2.65 | 2.61 | 2.94 | 3.8 | 2.59 | 2.67 | - |
| Atp7a | 0.58 | 0.62 | 0.41 | 1.12 | 0.97 | 0.94 | 0.98 | 0.95 | 0.63 | 0.77 | 1.57 | 0.99 | - |
| Asph | 8.02 | 5.65 | 7.55 | 6.08 | 7.69 | 8.03 | 4.58 | 5.86 | 4.95 | 4.26 | 6.16 | 4.18 | - |
| Trafd1 | 23.69 | 21.9 | 19.52 | 32.55 | 29.51 | 32.76 | 31.63 | 24.38 | 25.27 | 44.47 | 39.19 | 53.73 | - |
| Phxr4 | 1.9 | 1.6 | 2.39 | 3.3 | 2.17 | 2.77 | 2.5 | 2.55 | 1.83 | 3.77 | 4.64 | 4.3 | - |
| Sgk3 | 2.52 | 2.73 | 2.04 | 0.85 | 1.49 | 1.48 | 1.83 | 1.43 | 1.52 | 1.01 | 0.75 | 1.36 | - |
| Il2ra | 340.59 | 332.03 | 381.97 | 298.3 | 347.7 | 320.74 | 252.94 | 326.49 | 326.21 | 190.06 | 225.6 | 193.02 | - |

| | | | | | | | | | | | | |
|---|---|---|---|---|---|---|---|---|---|---|---|---|
| Cpt1a | 13.99 | 12.92 | 13.29 | 17.47 | 13.13 | 12.65 | 20.09 | 18.8 | 18.16 | 19.31 | 18.86 | 17.26 | - |
| Arl6ip5 | 51.72 | 43.98 | 43 | 69.29 | 64.29 | 58.11 | 78.11 | 58.73 | 67.7 | 83.51 | 86.86 | 88.96 | - |
| Yes1 | 0.3 | 0.28 | 0.28 | 0.5 | 0.64 | 0.31 | 0.45 | 0.48 | 0.3 | 0.92 | 0.71 | 0.45 | - |
| Pld3 | 44.82 | 46.44 | 41.9 | 103.42 | 80.45 | 85.55 | 99.98 | 65.57 | 68.43 | 126.23 | 106.2 | 132.87 | - |
| Mcm10 | 15.42 | 14.1 | 12.92 | 7 | 7.19 | 7.18 | 2.58 | 5.19 | 3.78 | 2.64 | 2.46 | 2.36 | - |
| Cdca7 | 57.54 | 60.55 | 54.11 | 34.8 | 40 | 38.09 | 22.34 | 31.54 | 33.56 | 20.21 | 18.71 | 17.41 | - |
| Ptpn11 | 3.05 | 2.88 | 3.06 | 2.84 | 2.69 | 3.23 | 2.72 | 2.25 | 2.21 | 1.64 | 1.86 | 2.32 | - |
| Zik1 | 1.13 | 1.28 | 1.65 | 1.18 | 1.66 | 0.94 | 0.39 | 0.85 | 0.65 | 0.78 | 0.53 | 0.81 | - |
| Gm1821 | 321.11 | 311.94 | 297.89 | 346.87 | 391.82 | 381.47 | 388.54 | 350.65 | 374.41 | 405.84 | 454.04 | 407.54 | - |
| Kcna4 | 0.6 | 0.08 | 0.26 | 4.46 | 3.16 | 3.26 | 0.46 | 0.4 | 0.32 | 5.63 | 4.9 | 5.55 | - |
| Ovol2 | 0.19 | 0 | 0 | 0.94 | 1.08 | 0.47 | 0.28 | 0.68 | 0.61 | 2.17 | 0.97 | 2 | - |
| Creb3l2 | 2.93 | 2.74 | 3.65 | 3.73 | 5.98 | 4.13 | 5.7 | 6.55 | 6.03 | 5.88 | 6.69 | 5.68 | - |
| Zfp386 | 16.26 | 18.68 | 15.81 | 22.24 | 17.74 | 21.25 | 27.07 | 18.28 | 19.79 | 25.57 | 26.45 | 32.47 | - |
| AW549877 | 3.22 | 3.45 | 2.85 | 4.13 | 4.51 | 4.54 | 5.15 | 3.75 | 3.84 | 5.21 | 5.09 | 4.97 | - |
| Coro1b | 134.02 | 127.96 | 128.03 | 183.1 | 162.95 | 179.49 | 210.76 | 167.56 | 177.94 | 212.51 | 216.93 | 225.2 | - |
| Uba1 | 206.79 | 205.14 | 203.44 | 174.35 | 184.76 | 186.58 | 193.56 | 198.28 | 190.23 | 164.51 | 177.01 | 165.92 | - |
| Pcsk4 | 2.91 | 3.33 | 2.49 | 5.28 | 3.95 | 3.54 | 6.07 | 3.81 | 4.56 | 5.3 | 6.62 | 6.56 | - |
| Itgb2 | 185.54 | 186.59 | 171.26 | 171.06 | 172.06 | 175.81 | 265.94 | 244.46 | 253.96 | 221.05 | 226.1 | 241.49 | - |
| Ccno | 0.06 | 0.25 | 0.23 | 0.67 | 0.57 | 0.61 | 0.8 | 0.13 | 0.6 | 0.91 | 0.92 | 0.8 | - |
| Pde4d | 1.82 | 1.76 | 2.43 | 3.74 | 3.85 | 2.85 | 4.13 | 3 | 3.19 | 5.23 | 4.61 | 5.04 | - |
| Hlx | 0 | 0.11 | 0.03 | 0.06 | 0.06 | 0 | 1.95 | 0.88 | 0.77 | 1.01 | 0.89 | 1.42 | - |
| Slc25a25 | 5.12 | 5.44 | 4.4 | 3.07 | 3.16 | 3.24 | 1.71 | 2.74 | 2.46 | 1.63 | 1.81 | 1.25 | - |
| Oas1b | 3.01 | 3.06 | 2.76 | 5.74 | 4.85 | 4.47 | 5.2 | 4.8 | 4.34 | 8.04 | 5.82 | 5.13 | - |
| Nkiras2 | 22.8 | 23.39 | 21.31 | 26.59 | 25.74 | 21.6 | 31.22 | 26 | 28.45 | 28.21 | 29.44 | 31.71 | - |
| Ddx26b | 3.67 | 2.81 | 2.71 | 4.51 | 4.6 | 3.6 | 4.35 | 4.32 | 5.1 | 5.39 | 5.82 | 5.48 | - |
| Zfp236 | 1.1 | 1.07 | 0.82 | 1.56 | 1.73 | 1.37 | 1.16 | 1.36 | 1.59 | 1.72 | 2.02 | 1.66 | - |
| Ints1 | 7.7 | 7.31 | 8.1 | 5.65 | 5.76 | 6.09 | 4.56 | 5.66 | 6.36 | 4.57 | 4.53 | 5.25 | - |
| Amdhd2 | 16.06 | 17.54 | 16.3 | 14.69 | 16.64 | 15.74 | 26.63 | 21.16 | 22.41 | 23.16 | 20.35 | 28.96 | - |
| Elmod3 | 5.22 | 6.05 | 6.3 | 7.76 | 7.41 | 6.28 | 7.77 | 6.73 | 5.38 | 10.38 | 6.96 | 9.45 | - |
| Capn3 | 0.72 | 0.58 | 0.31 | 0.75 | 0.33 | 0.75 | 0.07 | 0.2 | 0.31 | 0.09 | 0.14 | 0.04 | - |
| 1110018G07Rik | 8.04 | 6.4 | 8.4 | 8.43 | 8.26 | 8.49 | 11.04 | 14.35 | 12.21 | 9.73 | 10.79 | 9.12 | - |
| Cpne2 | 0.99 | 0.84 | 0.8 | 0.19 | 0.24 | 0.97 | 0.1 | 0.59 | 0.65 | 0.17 | 0.16 | 0.35 | - |
| Zfand1 | 22.86 | 20.89 | 19.68 | 29.05 | 24.64 | 25.4 | 29.8 | 25.54 | 28.51 | 27.07 | 25.57 | 27.83 | - |
| Rell1 | 1.16 | 1.67 | 1.45 | 3.22 | 2.27 | 3.07 | 3.05 | 2.7 | 3.11 | 4.14 | 3.42 | 4.53 | - |
| Irak3 | 0.77 | 0.41 | 0.76 | 1 | 0.48 | 0.37 | 1.94 | 1.15 | 1.43 | 2.18 | 1.64 | 3.49 | - |
| Mpc1 | 121.04 | 115.07 | 117.14 | 87.79 | 99.04 | 103.92 | 73.54 | 95.58 | 86.83 | 62.87 | 65.84 | 63.81 | - |
| Pla1a | 2.34 | 1.59 | 1.4 | 1.95 | 1.14 | 2 | 0.76 | 0.3 | 0.38 | 0.35 | 0.69 | 0.53 | - |
| Plekhj1 | 60.71 | 60.08 | 59.15 | 47.45 | 49.69 | 51.68 | 46.3 | 52.48 | 51.76 | 44.49 | 42.75 | 42.04 | - |
| Prkcz | 3.58 | 3.01 | 2.67 | 4.87 | 5.03 | 4.88 | 5.78 | 5.37 | 5.72 | 6.53 | 6.63 | 7.3 | - |
| Arfgap1 | 21.74 | 20.48 | 21.6 | 19.05 | 19.07 | 15.54 | 14.98 | 17.46 | 16.05 | 14.23 | 12.57 | 13.95 | - |
| R3hdm4 | 30.22 | 33.72 | 30.29 | 43.66 | 40.54 | 44.19 | 57.8 | 50.42 | 49 | 59.93 | 58.36 | 58.47 | - |
| Akip1 | 39.19 | 41.62 | 38.51 | 31.24 | 30.02 | 33.1 | 32.19 | 31.39 | 25.2 | 27.52 | 24.07 | 22.1 | - |
| Rgs16 | 48.36 | 44.24 | 50.15 | 16.32 | 21.31 | 30.26 | 21.53 | 27.95 | 24.44 | 23.4 | 15.36 | 15.5 | - |
| Mettl2 | 12.75 | 13.33 | 13.9 | 9.81 | 8.92 | 10.88 | 8.04 | 9.68 | 9.87 | 7.99 | 7.42 | 7.58 | - |
| Matn1 | 0 | 0 | 0 | 0.09 | 0 | 0.03 | 0.17 | 0.15 | 0.13 | 0.09 | 0.37 | 0.12 | - |
| Tsix | 0.31 | 0.23 | 0.58 | 0.75 | 0.64 | 0.51 | 0.99 | 0.44 | 0.68 | 1.34 | 1.47 | 1.19 | - |
| Rplp2-ps1 | 19.59 | 13.89 | 13.71 | 16.23 | 19.87 | 18.1 | 19.62 | 19.11 | 15.6 | 21.97 | 24.98 | 24.43 | - |
| A330023F24Rik | 0.07 | 0.15 | 0.02 | 0.25 | 0.37 | 0.31 | 0.1 | 0.32 | 0.18 | 0.56 | 0.65 | 0.72 | - |
| Serpinb1a | 1.04 | 2.15 | 0.76 | 3.16 | 1.24 | 0.96 | 11.62 | 1.13 | 5.5 | 12.25 | 5.17 | 12.83 | - |
| Acaa1b | 0.05 | 0 | 0.03 | 1.04 | 1.11 | 1.17 | 0 | 0 | 0 | 2.39 | 2.26 | 2.21 | - |
| Dip2c | 0.45 | 0.33 | 0.51 | 0.54 | 0.5 | 0.54 | 0.47 | 0.41 | 0.74 | 0.67 | 0.89 | 0.84 | - |
| Zfp668 | 5.84 | 6.79 | 5.44 | 5.24 | 4.63 | 5.19 | 3.96 | 4.77 | 4.82 | 4.44 | 3.34 | 4.66 | - |
| Zscan2 | 0.02 | 0.03 | 0.07 | 0.28 | 0.08 | 0.14 | 0.33 | 0.26 | 0.21 | 0.45 | 0.38 | 0.4 | - |
| Pts | 19.13 | 23.38 | 16.72 | 22.73 | 22.22 | 24.38 | 36.6 | 22.93 | 26.89 | 35.25 | 33.74 | 42.62 | - |
| Nfyb | 28.97 | 27.2 | 29.82 | 34.4 | 38.57 | 39.45 | 37.04 | 40.8 | 42.23 | 44.81 | 47.21 | 41.94 | - |
| C030039L03Rik | 0.48 | 0.54 | 0.52 | 1.11 | 0.89 | 0.72 | 1.56 | 0.83 | 0.86 | 1.05 | 1.07 | 1.54 | - |
| Serp1 | 134.77 | 141.74 | 141.98 | 103.17 | 99.68 | 102.95 | 97.44 | 128.2 | 113.55 | 93.89 | 93.93 | 86.71 | - |
| Pdha1 | 96.96 | 97.82 | 101.82 | 98.22 | 102.11 | 103.58 | 83 | 96.76 | 94.92 | 83.73 | 84.1 | 78.53 | - |
| Nploc4 | 10.21 | 8.12 | 9.41 | 7.79 | 7.46 | 7.65 | 6.33 | 7.84 | 7.76 | 5.26 | 6.15 | 6.96 | - |
| 1500032L24Rik | 160.44 | 168.81 | 159.6 | 138.08 | 146.08 | 146.73 | 163.28 | 178.03 | 157.84 | 123.95 | 140.58 | 130.26 | - |
| Jmy | 0.33 | 0.36 | 0.31 | 0.76 | 0.92 | 0.78 | 0.92 | 0.64 | 0.69 | 1.02 | 1.11 | 1.56 | - |
| Lag3 | 84.02 | 69.99 | 72.93 | 160.99 | 160.53 | 172.96 | 237.26 | 180.28 | 187.17 | 303.67 | 304.55 | 348.73 | - |
| Vps26b | 8.66 | 11.76 | 11.6 | 15.63 | 14.6 | 14.07 | 18.37 | 15.47 | 15.78 | 19.07 | 19.78 | 18.7 | - |
| Retsat | 8.28 | 5.98 | 8.37 | 6.07 | 5.93 | 5.99 | 4.51 | 6.28 | 4.57 | 3.65 | 5.54 | 4.61 | - |
| Trim44 | 8.29 | 7.98 | 8.61 | 8.53 | 8.94 | 8.08 | 5.62 | 7.46 | 6.06 | 5.62 | 6.67 | 6.47 | - |
| Pde1a | 0.3 | 0.22 | 0.91 | 0.04 | 0.23 | 0.21 | 0.06 | 0.02 | 0.21 | 0.07 | 0 | 0.03 | - |
| Zfp281 | 5.21 | 6.71 | 8.4 | 13.04 | 13.19 | 12.11 | 7.86 | 8.08 | 7.38 | 11.41 | 11.07 | 11.84 | - |
| Zmym2 | 4.08 | 3.06 | 3.82 | 4.86 | 5.22 | 4.86 | 4.91 | 4.93 | 5.18 | 6.23 | 7.69 | 6.73 | - |
| Enkd1 | 16.72 | 15.23 | 17.4 | 11.33 | 13.1 | 13.38 | 12.42 | 11.77 | 10.59 | 7.35 | 6.71 | 7.72 | - |
| Kcnq2 | 0.45 | 0.26 | 0.16 | 0.79 | 0.33 | 0.63 | 0.48 | 0.76 | 0.37 | 1.06 | 0.98 | 0.83 | - |
| Nek7 | 7.08 | 7.11 | 9.04 | 4.64 | 5.84 | 5.11 | 4.16 | 5.25 | 4.92 | 3.99 | 4.36 | 3.74 | - |
| Acsl3 | 1.32 | 1.83 | 2.47 | 3.12 | 3.68 | 3.21 | 2.39 | 3.2 | 2.89 | 4.47 | 4.22 | 3.16 | - |
| 2410066E13Rik | 0.17 | 0.14 | 0.25 | 0.48 | 0.58 | 1.04 | 0.93 | 0.41 | 0.63 | 1.27 | 0.9 | 1.04 | - |
| Mat2b | 52.28 | 59.47 | 53.27 | 74.99 | 67.43 | 71.53 | 83.89 | 80.09 | 85.67 | 95.95 | 96.7 | 96.23 | - |
| Cdk19 | 1.71 | 1.39 | 1.39 | 2.68 | 2.47 | 2.81 | 3.32 | 1.88 | 2.68 | 3.27 | 3.31 | 2.58 | - |
| Ccng2 | 2.93 | 2.5 | 2.38 | 3.27 | 2.8 | 2.56 | 5 | 3.91 | 5.74 | 5.12 | 6.6 | 4.7 | - |
| 2900008C10Rik | 1.55 | 1.5 | 1.75 | 2.74 | 3.17 | 2.06 | 3.21 | 1.6 | 2.84 | 3.79 | 2.54 | 4.2 | - |
| Acbd5 | 5.15 | 4.85 | 4.91 | 6.26 | 6.24 | 6.35 | 7.17 | 6.21 | 6.37 | 7.24 | 8.23 | 6.01 | - |
| Pole4 | 24.64 | 40.43 | 47.11 | 26.79 | 29 | 26.01 | 20.45 | 28.19 | 28.54 | 22.82 | 20.81 | 18.92 | - |
| Znrf1 | 1.69 | 2.72 | 2.17 | 5.33 | 3.85 | 4.04 | 4.2 | 3.05 | 4.13 | 4.78 | 4.42 | 5.31 | - |
| Pcif1 | 13.28 | 13.89 | 13.12 | 22.28 | 21.28 | 22.13 | 26.46 | 16.71 | 22.79 | 27.69 | 23.83 | 25.62 | - |
| Hax1 | 122.24 | 114.21 | 123.17 | 71.29 | 82.82 | 82.5 | 79.59 | 109.14 | 104.13 | 50.77 | 65.24 | 55.57 | - |
| Galnt11 | 3.64 | 4.97 | 3.73 | 5.4 | 4.53 | 4.97 | 8.14 | 7.17 | 6.31 | 8.17 | 7.1 | 6.66 | - |
| Mthfsd | 9.81 | 12.47 | 12.04 | 8.47 | 7.8 | 9.84 | 6.35 | 7.36 | 8.55 | 5.11 | 3.34 | 6.68 | - |
| Aga | 36.64 | 32.61 | 30.51 | 21.8 | 25.26 | 24.22 | 22.18 | 26.27 | 28.57 | 15.43 | 20.66 | 17.83 | - |
| Ociad1 | 152.7 | 152.16 | 144.95 | 132.78 | 129.46 | 148.47 | 110.47 | 141.8 | 127.39 | 122.8 | 128.28 | 129.59 | - |
| Fam175a | 8.8 | 9.53 | 7.55 | 6.15 | 8.37 | 7.19 | 4.51 | 5.57 | 5.5 | 4.94 | 5.57 | 4.57 | - |
| Fam50a | 25.46 | 22 | 22.17 | 29.59 | 30.64 | 27.95 | 31.08 | 24.46 | 26.82 | 29.62 | 29.72 | 29.51 | - |
| Gypc | 26.73 | 25.25 | 26.09 | 23.38 | 26.46 | 29.08 | 15.75 | 21.46 | 16.87 | 17.93 | 16.27 | 20.52 | - |
| Tstd3 | 31.52 | 28.27 | 31.35 | 31.32 | 31.2 | 35.62 | 43.25 | 38.1 | 38.78 | 41.43 | 41.23 | 40.15 | - |
| Ap2a1 | 11.82 | 10.59 | 13.72 | 10.11 | 10.95 | 9.98 | 10.02 | 11.28 | 9.75 | 8.45 | 8.42 | 8.77 | - |
| Ndst3 | 0 | 0.01 | 0 | 0.05 | 0.12 | 0.11 | 0.14 | 0.19 | 0.14 | 0.33 | 0.5 | 0.38 | - |
| Zfp943 | 2.78 | 2.91 | 2.82 | 4.81 | 4.42 | 4.38 | 3.94 | 3.45 | 2.64 | 5.76 | 4.24 | 4.19 | - |
| Lcorl | 1.16 | 1.36 | 1.64 | 3.16 | 2.53 | 2.45 | 3.27 | 2.46 | 2.55 | 3.35 | 4.81 | 3.05 | - |
| Tomm40l | 17.95 | 16.3 | 17.04 | 13.36 | 14.39 | 13.8 | 12.99 | 15.23 | 14.09 | 12.32 | 9.88 | 12.16 | - |
| Knstrn | 22.05 | 19.77 | 20.81 | 20.07 | 24.33 | 21.37 | 16.42 | 12.81 | 11.61 | 15.06 | 13.38 | 14.35 | - |
| Serhl | 27.43 | 22.25 | 21.9 | 33.45 | 33.76 | 32.76 | 43.41 | 32.84 | 35.61 | 42.05 | 39.91 | 45.11 | - |
| Pex3 | 21.52 | 23.74 | 22.56 | 15.85 | 19.14 | 19.17 | 16.87 | 22.94 | 18.68 | 16.49 | 16.11 | 17.77 | - |
| Dennd2c | 0.65 | 0.73 | 0.38 | 1.45 | 0.63 | 0.82 | 1.01 | 1.45 | 1.23 | 0.94 | 1.16 | 1.36 | - |
| Hist1h2ag | 5.48 | 5.98 | 7.73 | 1.63 | 2.4 | 2.7 | 2.15 | 0.32 | 1.56 | 2.34 | 0.55 | 0.55 | - |
| Rai14 | 4.26 | 3.93 | 3.8 | 0.54 | 0.92 | 0.7 | 0.82 | 2.16 | 1.93 | 0.43 | 0.22 | 0.49 | - |
| Rnf44 | 6.1 | 5.84 | 6.4 | 6.52 | 6.49 | 6.81 | 7.6 | 6.72 | 8.56 | 9.73 | 9.61 | 9.36 | - |
| Rabep2 | 14.91 | 11.99 | 12.44 | 18.87 | 16.42 | 19.82 | 20.02 | 17.33 | 19.27 | 21.7 | 19.37 | 27.36 | - |
| Mrvi1 | 0.11 | 0.11 | 0.1 | 0.05 | 0 | 0 | 0.03 | 0.04 | 0.05 | 0 | 0 | 0 | - |
| Pask | 2.62 | 3.13 | 3.34 | 2.57 | 2.71 | 2.85 | 1.75 | 1.62 | 1.78 | 0.88 | 1.34 | 1.28 | - |
| Rad21 | 12.29 | 12.21 | 12.75 | 10.8 | 12.31 | 12.58 | 9.34 | 10.1 | 10.12 | 8.89 | 8.63 | 8.95 | - |
| Prex1 | 4.53 | 4.79 | 3.67 | 7.57 | 6.02 | 7.17 | 7.45 | 5.47 | 5.73 | 10.85 | 7.78 | 9.96 | - |
| Trpv2 | 47.37 | 50.17 | 43.41 | 66.12 | 62.49 | 61.41 | 73.1 | 59.57 | 62.54 | 75.88 | 72.23 | 81.07 | - |
| Evi2b | 10.43 | 12.52 | 11.16 | 14.42 | 10.65 | 12.31 | 16.49 | 13.17 | 13.93 | 19.53 | 17.01 | 20.76 | - |
| Dbnl | 80.44 | 81 | 81.91 | 65.88 | 62.77 | 65.02 | 61.32 | 58.14 | 56.86 | 57.93 | 45.57 | 65.52 | - |
| Anapc5 | 144.44 | 143.65 | 146.87 | 91.29 | 102.06 | 96.68 | 87.71 | 95.7 | 99.28 | 62.54 | 69.51 | 63.53 | - |
| Slc39a4 | 16.52 | 14.93 | 15.07 | 10.19 | 11.19 | 10.2 | 12.2 | 13.91 | 12.41 | 8.87 | 9.37 | 9.72 | - |
| Rraga | 15.04 | 16.83 | 15.87 | 10.54 | 12.83 | 8.79 | 8.92 | 11.7 | 10.12 | 7.03 | 8.42 | 8.44 | - |
| Gtf2a2 | 74.15 | 75.98 | 78.87 | 72.61 | 75.54 | 73.26 | 45.06 | 59.74 | 54.26 | 48.59 | 56.34 | 54.68 | - |
| Picalm | 10.25 | 10 | 10.22 | 11.85 | 11.29 | 12.7 | 13.83 | 12.45 | 14.21 | 17.53 | 17.24 | 16.95 | - |
| Itgal | 54.09 | 53.7 | 51.93 | 49.59 | 46.83 | 49.42 | 62.23 | 53.13 | 62.64 | 65.67 | 56.94 | 65.17 | - |
| Tmem198b | 2.76 | 2.01 | 3.43 | 4.61 | 4.21 | 4.25 | 3.85 | 3.4 | 4.32 | 4.1 | 4.52 | 5.37 | - |
| Sestd1 | 1.3 | 0.97 | 0.97 | 1.93 | 2.37 | 1.67 | 1.11 | 1.34 | 1.32 | 1.76 | 2.25 | 1.94 | - |
| Stat5b | 13.07 | 12.14 | 13.51 | 14.23 | 14.82 | 12.57 | 13.31 | 16.8 | 16.46 | 16.66 | 17.62 | 16.53 | - |
| Trub1 | 9.28 | 8.98 | 10.05 | 7.4 | 6.87 | 7.99 | 5.15 | 7.21 | 6.39 | 5.54 | 5 | 4.55 | - |
| Babam1 | 50.17 | 53.07 | 47.45 | 38.71 | 44.16 | 38.86 | 39.84 | 42.24 | 41.25 | 32.2 | 28.72 | 33.85 | - |
| Tanc1 | 1.36 | 1.11 | 0.96 | 1.16 | 1.86 | 1.13 | 3.15 | 1.41 | 2.09 | 2.5 | 3.05 | 2.65 | - |
| Terf1 | 9.96 | 10.48 | 9.9 | 14.98 | 16.17 | 15.54 | 15.96 | 11.83 | 12.74 | 15.17 | 17.32 | 13.55 | - |
| Zc3h11a | 15.89 | 16.27 | 15.94 | 20.55 | 18.9 | 20.86 | 18.76 | 16.69 | 19.37 | 19.55 | 23.01 | 18.47 | - |
| Tulp4 | 6 | 4.41 | 4.79 | 9.55 | 8.7 | 8.73 | 7.98 | 8.45 | 9.16 | 17.07 | 14.64 | 17.91 | - |
| Inhba | 138.93 | 91.01 | 146.7 | 24.84 | 56.07 | 44.66 | 15.76 | 49.71 | 35.39 | 17.32 | 19.63 | 25.71 | - |
| Nfkb2 | 56.72 | 56.56 | 58.33 | 36.16 | 40.77 | 36.7 | 32.81 | 43.3 | 40.36 | 27.28 | 28.89 | 30.34 | - |
| Utf1 | 4.63 | 4.76 | 5.69 | 1.2 | 1.29 | 1.37 | 3.81 | 2.84 | 2.87 | 0.89 | 0.85 | 0.8 | - |
| Ptpn9 | 3.1 | 2.96 | 3 | 3.37 | 1.75 | 2.03 | 1.71 | 2.23 | 2.05 | 1.78 | 1.38 | 1.95 | - |
| Entpd7 | 2.41 | 1.95 | 1.56 | 1.07 | 1.27 | 0.85 | 0.97 | 1.56 | 1.08 | 0.77 | 1.04 | 0.95 | - |
| Trim12c | 9.08 | 8.91 | 8.55 | 11.98 | 11.28 | 11.74 | 11.83 | 9.59 | 9.78 | 12.85 | 11.58 | 14.55 | - |
| Gpr55 | 0.36 | 0.82 | 0.32 | 0.22 | 0.34 | 0.1 | 0.11 | 0 | 0.04 | 0.09 | 0.02 | 0.05 | - |
| Cst7 | 166.11 | 162.69 | 158.97 | 301.61 | 288.31 | 291.19 | 296.59 | 239.79 | 243.49 | 385.76 | 377.26 | 402.35 | - |
| 2010003O02Rik | 5.62 | 7.16 | 5.32 | 9.53 | 15.94 | 11.38 | 10.43 | 10.3 | 8.42 | 11.62 | 16.31 | 14.37 | - |
| Hnrpll | 4.98 | 4.74 | 3.85 | 2.72 | 1.9 | 2.95 | 1.79 | 2.45 | 2.36 | 1.8 | 1.38 | 1.22 | - |
| Ncoa2 | 3.21 | 2.96 | 2.94 | 4.82 | 4.11 | 4.59 | 2.98 | 3.76 | 3.72 | 4.97 | 4.61 | 5.12 | - |
| Tbce | 19.48 | 16.91 | 17.31 | 13.8 | 13.79 | 13.08 | 11.64 | 14.82 | 13.44 | 10.35 | 10.54 | 14.77 | - |
| Gtpbp10 | 7.42 | 6.43 | 6.31 | 3.97 | 4.64 | 5.61 | 3.68 | 5.09 | 5.4 | 3.9 | 3.13 | 3.4 | - |
| Tigd2 | 7.09 | 8.53 | 7.77 | 9.17 | 9.99 | 8.77 | 11.33 | 8.69 | 9.43 | 12.37 | 11.13 | 12.86 | - |
| Rrp12 | 22.17 | 21.44 | 23.13 | 10.99 | 11.99 | 12.62 | 8.98 | 13.65 | 12.74 | 7.67 | 6.68 | 6.51 | - |
| Ier5 | 1.75 | 1.55 | 2.05 | 1.51 | 1.51 | 1.64 | 0.53 | 1.4 | 1.21 | 0.84 | 0.55 | 0.78 | - |
| Tnfsf13b | 0.03 | 0.14 | 0.1 | 0.86 | 0 | 0.53 | 0.88 | 0.38 | 1.04 | 1.18 | 0.87 | 2.31 | - |
| Hjurp | 3 | 2.4 | 2.21 | 3.06 | 3.85 | 4.2 | 2.83 | 2.75 | 2.89 | 4.2 | 4.11 | 3.03 | - |
| Ece2 | 33.19 | 37.75 | 30.65 | 22.82 | 25.03 | 24.79 | 19.34 | 28.19 | 24.44 | 16.8 | 18.34 | 16.57 | - |
| Ccdc47 | 38.12 | 42.54 | 40.57 | 32.32 | 37.19 | 37.78 | 31.42 | 31.58 | 31.91 | 30.22 | 28.46 | 30.92 | - |
| Gstt2 | 47.3 | 38.1 | 53.36 | 59.56 | 68.57 | 61.68 | 69.67 | 75.32 | 67.76 | 65.93 | 74.13 | 68.41 | - |
| Psat1 | 261.96 | 252.93 | 257.44 | 149.32 | 186.28 | 175.22 | 139.46 | 211.85 | 191.89 | 87.12 | 107.2 | 91.85 | - |
| Sbds | 44.24 | 46.39 | 50.16 | 60.01 | 59.06 | 61.7 | 59.52 | 57.47 | 55.9 | 68.58 | 77.54 | 69.42 | - |
| Ctnnb1 | 10.76 | 12.69 | 12.54 | 19.25 | 14.79 | 18.69 | 17.06 | 13.41 | 12.74 | 21.63 | 20.66 | 21.45 | - |
| Fnip1 | 1.96 | 1.76 | 1.96 | 2.34 | 2.82 | 2.92 | 2.31 | 2.43 | 2.26 | 2.73 | 3.08 | 2.4 | - |
| Svil | 0.97 | 1.32 | 1.29 | 2.04 | 1.82 | 1.78 | 2.11 | 1.29 | 1.81 | 1.87 | 2.22 | 1.94 | - |

| | | | | | | | | | | | | | |
|---|---|---|---|---|---|---|---|---|---|---|---|---|---|
| 4921525B02Rik | 0.26 | 0.11 | 0.28 | 0.31 | 0.43 | 0.27 | 0.47 | 0.29 | 0.33 | 1.63 | 0.7 | 1.25 | - |
| N4bp2l2 | 8.2 | 7.62 | 8.04 | 9.71 | 9.81 | 10.25 | 9.1 | 9.71 | 9.14 | 11.66 | 12.4 | 10.88 | - |
| Dnaaf2 | 11.94 | 11.89 | 12.34 | 11.76 | 12.09 | 11.39 | 8.47 | 7.88 | 9.64 | 7.35 | 7.87 | 7.79 | - |
| Cd84 | 13.54 | 10.69 | 12.77 | 14.96 | 13.64 | 13.4 | 19.06 | 17.54 | 16.18 | 19.2 | 17.58 | 20.49 | - |
| Tacc1 | 11.91 | 12.81 | 11.3 | 13.39 | 10.95 | 10.3 | 14.89 | 12.85 | 12.71 | 14.96 | 14.04 | 17.24 | - |
| Cbx7 | 4.48 | 5.41 | 5.2 | 9.98 | 10.08 | 10.12 | 9.61 | 8.28 | 9.03 | 14.77 | 12.21 | 14.84 | - |
| Rassf2 | 4.72 | 5.1 | 5.16 | 5.91 | 5.26 | 6.21 | 6.99 | 5.06 | 4.87 | 8.38 | 6.5 | 7.99 | - |
| Ikbke | 6.29 | 7.34 | 5.62 | 11.62 | 9.49 | 9.38 | 10.26 | 9.26 | 9.04 | 11.73 | 9.5 | 11.41 | - |
| Cyth1 | 19.67 | 21.37 | 22.52 | 33.36 | 29.87 | 27.78 | 21.47 | 19.07 | 20.75 | 25.21 | 25.57 | 25.6 | - |
| Chmp6 | 21.39 | 22.18 | 22.58 | 19.44 | 17.12 | 18.92 | 19.06 | 16.19 | 20.52 | 17.48 | 16.38 | 15.77 | - |
| Sh3kbp1 | 24.92 | 25.62 | 24.13 | 34.14 | 25.36 | 30.34 | 48.67 | 33.47 | 35.15 | 46.24 | 41.69 | 46.3 | - |
| Kdsr | 4.76 | 4.48 | 3.97 | 5.71 | 6.35 | 6.01 | 5.68 | 5.91 | 7.1 | 8.02 | 7.62 | 7.21 | - |
| Hdac9 | 0.35 | 0.17 | 0.17 | 0.06 | 0.19 | 0.07 | 0 | 0 | 0.02 | 0.03 | 0 | 0.03 | - |
| Tnni1 | 0.71 | 0.95 | 0.99 | 0.06 | 0.12 | 0.12 | 0.11 | 0 | 0.16 | 0 | 0 | 0 | - |
| Ulk1 | 1.23 | 1.13 | 1.24 | 3.04 | 2.46 | 2.82 | 3.95 | 2.34 | 2.29 | 4.14 | 4.24 | 4.81 | - |
| A630001G21Rik | 6.9 | 7.24 | 5.84 | 5.48 | 4.58 | 5.95 | 4.1 | 4.34 | 5.07 | 4.14 | 2.98 | 4.05 | - |
| Syne3 | 3.99 | 3.94 | 4.47 | 5.6 | 5.47 | 5.73 | 5.55 | 5.3 | 5.26 | 6.17 | 6.55 | 4.58 | - |
| Ifi27l1 | 255.83 | 242.99 | 241.01 | 256.22 | 265.89 | 270.61 | 172.98 | 203.95 | 186.34 | 170.38 | 176.43 | 192.08 | - |
| S100a13 | 46.95 | 44.02 | 41.81 | 47.37 | 40.41 | 48.34 | 67.19 | 54.05 | 56.27 | 56.25 | 54.39 | 58.85 | - |
| Haus2 | 15.36 | 13.85 | 14.03 | 10.04 | 13.41 | 10.8 | 9.26 | 12.86 | 11.9 | 8.71 | 11.27 | 8.71 | - |
| Zfp606 | 1.53 | 2 | 1.99 | 3.2 | 2.55 | 2.59 | 2.14 | 1.81 | 1.63 | 3.83 | 2.99 | 2.96 | - |
| Prkab1 | 14.14 | 15.26 | 14.58 | 15.19 | 17.68 | 16.57 | 18.32 | 19.81 | 17.18 | 22.37 | 19.71 | 20.72 | - |
| Snx2 | 37.43 | 40.41 | 37.04 | 44.3 | 40.61 | 43.3 | 46.87 | 45.17 | 43.58 | 45.44 | 56.45 | 53.67 | - |
| Ttc32 | 6.57 | 6.34 | 6 | 3.1 | 3.68 | 4.21 | 3.39 | 3.35 | 3.65 | 2.86 | 2.34 | 2.08 | - |
| Etv3 | 2.88 | 2.88 | 2.38 | 4.29 | 3.57 | 4.84 | 3.48 | 3.84 | 4.78 | 5.43 | 4.82 | 4.06 | - |
| Trim37 | 10.1 | 11.54 | 11.4 | 7.78 | 8.19 | 9.41 | 7 | 9.99 | 9.48 | 7.69 | 7.61 | 5.74 | - |
| Rnf122 | 0.29 | 0.18 | 0.3 | 1.27 | 0.92 | 0.83 | 1.18 | 0.55 | 0.62 | 1.68 | 1.22 | 1.71 | - |
| Pitpnm1 | 14.74 | 14.97 | 14.24 | 24.98 | 21.92 | 21.78 | 19.55 | 18.18 | 17.38 | 22.5 | 21.56 | 26.86 | - |
| Atxn10 | 53.68 | 50.15 | 46.81 | 67.1 | 60.76 | 65.15 | 67.49 | 57.76 | 58.25 | 77.07 | 77.84 | 75.78 | - |
| Katna1 | 28.85 | 28.32 | 29.97 | 35.91 | 35.06 | 36.89 | 38.2 | 38.58 | 38.72 | 42.96 | 40.69 | 44.23 | - |
| Prdx2 | 440.59 | 387.3 | 421.95 | 343.19 | 364.13 | 362.11 | 373.28 | 378.67 | 345.81 | 308.07 | 322.24 | 327.48 | - |
| Cwc27 | 25.21 | 22.25 | 23.67 | 17.6 | 19.64 | 22.24 | 20.83 | 25.69 | 22.21 | 19.16 | 17.73 | 15.94 | - |
| Ncs1 | 29.94 | 25.33 | 30.07 | 20.46 | 23.27 | 25.38 | 19.48 | 28.73 | 25.44 | 16.54 | 18.42 | 19.49 | - |
| Mcl1 | 6.25 | 7.14 | 7.06 | 10.69 | 10.38 | 10.34 | 9.15 | 9.04 | 9.49 | 15.07 | 14.79 | 13.65 | - |
| Uevld | 3.49 | 2.21 | 2.37 | 4.95 | 4.84 | 5.14 | 3.44 | 3.28 | 4.17 | 4.05 | 5.52 | 3.22 | - |
| Nt5m | 9.4 | 8.51 | 9.4 | 11.57 | 10.56 | 11.57 | 9.85 | 10.7 | 12 | 13.67 | 11.51 | 13.42 | - |
| Cercam | 6.89 | 6.72 | 5.93 | 15.75 | 12.42 | 12.21 | 13.72 | 11.19 | 9.83 | 16.32 | 18.67 | 17.85 | - |
| Narf | 3.07 | 3.4 | 2.95 | 6.89 | 5.77 | 5.69 | 6.79 | 5.12 | 5.89 | 7.29 | 9.3 | 10.04 | - |
| Scp2 | 35.06 | 35.17 | 34.55 | 48.42 | 45.46 | 48.14 | 60.33 | 47.38 | 50.71 | 62.16 | 58.13 | 67.07 | - |
| Pfkfb3 | 6.43 | 7.27 | 5.91 | 9.58 | 7.6 | 8.18 | 12.85 | 9.94 | 11.12 | 11.29 | 16.26 | 13.84 | - |
| Arf4 | 115.72 | 116.29 | 114.24 | 132.85 | 145.82 | 153.67 | 126.71 | 139.26 | 138.87 | 135.55 | 160.54 | 133.67 | - |
| Ivd | 17.15 | 16.44 | 15.82 | 6.51 | 8.52 | 7.25 | 7.02 | 11.45 | 8.64 | 4.98 | 3.73 | 4.74 | - |
| A930013F10Rik | 0.38 | 0.29 | 0.42 | 0.57 | 1.47 | 0.66 | 0.97 | 0.39 | 0.73 | 0.66 | 1.04 | 1.24 | - |
| Plaur | 2.38 | 1.5 | 1.9 | 76.29 | 76.96 | 77.02 | 0.16 | 0.08 | 0.04 | 12.33 | 10.68 | 9.91 | - |
| Upf3b | 11.85 | 10 | 11.42 | 14.33 | 16.21 | 14.4 | 14.22 | 14.55 | 14.19 | 15.16 | 18.73 | 15.49 | - |
| Wdr31 | 4.31 | 3.29 | 4.86 | 2.58 | 1.98 | 2.02 | 3.51 | 3.9 | 3.28 | 2.46 | 1.95 | 1.96 | - |
| Ptgs2 | 0.03 | 0.17 | 0.26 | 0.75 | 0.02 | 0 | 0 | 0 | 0 | 0 | 0 | 0 | - |
| A430107P09Rik | 5.52 | 6.96 | 5.29 | 9.41 | 9.02 | 10.22 | 9.09 | 8.09 | 7.78 | 10.91 | 10.07 | 9.9 | - |
| Ankra2 | 11.9 | 11.2 | 10.94 | 15.06 | 14.96 | 14.29 | 12.98 | 11.53 | 14.53 | 16.06 | 14.12 | 15.91 | - |
| Spata13 | 6.69 | 7.17 | 7.86 | 12.26 | 10.74 | 10.7 | 14.38 | 10.18 | 12.4 | 16.48 | 16.34 | 16.73 | - |
| Slc25a13 | 11.57 | 8.52 | 10.16 | 6.08 | 6.28 | 5.58 | 5.45 | 6.01 | 6.24 | 4.27 | 4.01 | 4.62 | - |
| Hcst | 20.41 | 22.91 | 18.72 | 39.04 | 33.63 | 26.1 | 45.5 | 31.09 | 41.51 | 50.48 | 37.93 | 56.47 | - |
| Cep78 | 1.99 | 1.67 | 1.81 | 1.29 | 1.57 | 1.58 | 1.22 | 1.29 | 1.68 | 1.05 | 0.77 | 0.61 | - |
| Krit1 | 10.19 | 10.42 | 11.18 | 14.41 | 14.1 | 13.11 | 12.3 | 13.2 | 12.78 | 17.37 | 18.18 | 15.33 | - |
| Tmem161a | 19.22 | 19.16 | 19.83 | 16.76 | 16.66 | 16.93 | 16.32 | 17.54 | 17.6 | 15.42 | 13.32 | 13.44 | - |
| Peli1 | 8.24 | 12.31 | 13.61 | 22.22 | 18.88 | 17.99 | 22.97 | 21.27 | 20.68 | 30.28 | 27.97 | 24.06 | - |
| Cyp4v3 | 2.75 | 2.01 | 2.18 | 4.96 | 3.78 | 4.24 | 4.3 | 3.75 | 2.75 | 4.84 | 4.08 | 5.51 | - |
| Pros1 | 6.07 | 7.2 | 5.65 | 5.86 | 4.16 | 3.45 | 3.41 | 3.72 | 4.36 | 2.64 | 2.41 | 2.45 | - |
| Pnrc2 | 33.02 | 31.46 | 32.4 | 48.62 | 49.8 | 49.14 | 52.76 | 50.07 | 43.73 | 62.21 | 63.32 | 63.14 | - |
| Gpr155 | 0.95 | 1.08 | 1.14 | 0.75 | 0.74 | 0.83 | 0.7 | 0.78 | 0.68 | 0.64 | 0.36 | 0.46 | - |
| Ccni | 6.76 | 7.24 | 7.56 | 16.77 | 13.05 | 15.94 | 24.28 | 13.31 | 16.97 | 25.08 | 26.84 | 26.04 | - |
| Spc24 | 56.78 | 58.99 | 57.2 | 35.91 | 41.36 | 36.45 | 20.94 | 28.13 | 25.49 | 18.79 | 23.14 | 15.77 | - |
| Lrrc3b | 0 | 0 | 0 | 0.04 | 0.11 | 0.15 | 0.62 | 0.92 | 0.68 | 1.03 | 1.08 | 1.98 | - |
| Rsad2 | 0.3 | 0.14 | 0.46 | 0.63 | 0.76 | 1.6 | 0.77 | 0.84 | 0.31 | 1.51 | 0.99 | 1.96 | - |
| Alcam | 3.95 | 2.83 | 3.34 | 1.84 | 2.48 | 2.83 | 7.46 | 6.15 | 5.41 | 6.42 | 5.67 | 5.24 | - |
| Vrtn | 0.31 | 0.13 | 0.07 | 0 | 0 | 0 | 0.15 | 0.04 | 0 | 0 | 0 | 0 | - |
| Fars2 | 13.37 | 12.1 | 12.63 | 15.88 | 16.01 | 18.51 | 22.11 | 17.65 | 17.69 | 22.2 | 24.44 | 21.34 | - |
| Gmpr | 0.84 | 0.67 | 0.7 | 0.04 | 0.2 | 0 | 0.04 | 0.23 | 0.21 | 0.07 | 0.08 | 0.04 | - |
| Urb2 | 5.7 | 6.28 | 7.12 | 3.88 | 3.89 | 4.69 | 3.56 | 4.34 | 3.55 | 2.78 | 3.81 | 2.72 | - |
| Mettl1 | 11.07 | 14.04 | 13.17 | 7.84 | 7.88 | 12.15 | 7.71 | 11.99 | 11.49 | 5.16 | 6.6 | 4.61 | - |
| Mfge8 | 12.79 | 11.9 | 8.9 | 25.46 | 18.37 | 19.73 | 23.15 | 15.59 | 15.84 | 25.53 | 28.73 | 31.26 | - |
| Sh3bgrl3 | 386.85 | 409.37 | 359.97 | 557.47 | 502.69 | 506.95 | 579.86 | 432.2 | 493.41 | 501.59 | 466.68 | 558.73 | - |
| Elmo2 | 23.67 | 21.4 | 21.57 | 18.56 | 18.88 | 17.95 | 18.33 | 19.76 | 20.5 | 17.05 | 15.09 | 17.75 | - |
| Dhx34 | 4.93 | 4.26 | 4.31 | 7.78 | 7.44 | 6.86 | 6.05 | 6.94 | 4.86 | 7.92 | 6.39 | 7.42 | - |
| Zfp69 | 0.29 | 0.32 | 0.25 | 0.35 | 0.53 | 0.46 | 0.68 | 0.45 | 0.32 | 1.02 | 1.36 | 0.88 | - |
| Tctex1d1 | 2.1 | 2.19 | 2.47 | 4.42 | 3.83 | 4.01 | 2.77 | 2.47 | 3.45 | 5.34 | 5.8 | 6.15 | - |
| Napa | 85.85 | 84.2 | 85.26 | 72.56 | 76.62 | 71.51 | 70.68 | 76.14 | 72.36 | 61.23 | 65.91 | 68.4 | - |
| Ptges | 0.09 | 0 | 0.02 | 0.14 | 0.09 | 0.19 | 0.28 | 0.11 | 0.02 | 0.72 | 0.3 | 0.59 | - |
| Ttc4 | 27.32 | 26.73 | 28.16 | 25.93 | 25.03 | 24.38 | 21.63 | 21.75 | 21.91 | 22.69 | 20.05 | 22.37 | - |
| Rnf149 | 1.78 | 1.56 | 1.73 | 3.02 | 3.77 | 3.25 | 2.93 | 2.37 | 2.48 | 4.19 | 3.85 | 3.06 | - |
| Zfyve21 | 10.42 | 11.64 | 12.51 | 16.8 | 15.34 | 17.48 | 13.96 | 17.31 | 14.83 | 17.47 | 14.91 | 16.63 | - |
| Mterfd2 | 5.55 | 5.16 | 5.5 | 6.41 | 6.08 | 7.25 | 7.52 | 6.38 | 7.56 | 6.32 | 8.39 | 7.24 | - |
| Gpbp1l1 | 2.69 | 3.32 | 3.49 | 5.06 | 5 | 4.85 | 5.25 | 4.76 | 4.7 | 6.65 | 7.11 | 6.08 | - |
| Mt2 | 148.26 | 149.61 | 147.02 | 243.4 | 261.48 | 283.05 | 439.35 | 224.8 | 282.65 | 463.85 | 564.56 | 528.87 | - |
| Zfp62 | 12.69 | 13.12 | 12.43 | 11.79 | 13.94 | 9.34 | 7.27 | 9.41 | 8.89 | 6.98 | 7.54 | 8.26 | - |
| Synj2 | 1.99 | 1.77 | 1.81 | 1.2 | 1.43 | 1.93 | 1.49 | 1.3 | 1.7 | 0.9 | 0.91 | 0.91 | - |
| Tmem163 | 0.91 | 1.04 | 1.3 | 0.78 | 1.09 | 0.63 | 2.99 | 4.18 | 3.3 | 3.05 | 3.75 | 3 | - |
| Gnptab | 5.89 | 5.35 | 6.12 | 4.28 | 5.14 | 4.67 | 9.02 | 7.13 | 7.63 | 8.41 | 7.28 | 7.57 | - |
| Snapc2 | 20.78 | 20.41 | 20.57 | 23.42 | 21.69 | 23.88 | 25.62 | 23.16 | 26.6 | 27.85 | 28.98 | 28.5 | - |
| St7l | 2.51 | 2.29 | 2.14 | 2.8 | 2.9 | 3.58 | 2.83 | 3.26 | 2.94 | 3.63 | 3.77 | 3.51 | - |
| Rpl10 | 2213.56 | 2250.01 | 2217.72 | 2550.63 | 2565.49 | 2649.33 | 2510.96 | 2440.05 | 2406.99 | 2651.35 | 2918.63 | 2542.68 | - |
| H1f0 | 5.14 | 5.95 | 6.17 | 7.82 | 8.64 | 5.48 | 7.88 | 10.03 | 7.96 | 9.79 | 7.75 | 8.05 | - |
| 1810032O08Rik | 11.64 | 15.12 | 12.92 | 30.86 | 22.92 | 26.1 | 29.07 | 25.21 | 24.18 | 35.73 | 49.55 | 36.89 | - |
| Tdrkh | 8.62 | 7.44 | 9.56 | 6.49 | 8.15 | 8.49 | 4.62 | 5.98 | 5.34 | 3.95 | 3.26 | 2.5 | - |
| Tespa1 | 20.76 | 17.72 | 21.71 | 18.39 | 18.13 | 16.83 | 8.38 | 11.41 | 12.42 | 7.4 | 6.8 | 5.21 | - |
| Slc31a2 | 2.01 | 2.27 | 2.82 | 5.89 | 5.41 | 5 | 5.67 | 3.57 | 4.15 | 7.84 | 6.62 | 7.11 | - |
| Plcl2 | 3.73 | 3.81 | 3.85 | 6.23 | 4.64 | 6.66 | 7.86 | 6.22 | 5.18 | 6.59 | 6.47 | 8.37 | - |
| Fcer1g | 0.09 | 1.59 | 1.23 | 0 | 0 | 0 | 0 | 0.09 | 0.08 | 0 | 0 | 0 | - |
| 1810013L24Rik | 5.69 | 4.51 | 4.72 | 6.66 | 7.72 | 6.87 | 7.94 | 7.05 | 7.72 | 9.07 | 8.31 | 8.79 | - |
| Impdh1 | 31.72 | 39.61 | 35.72 | 37.01 | 32.48 | 34.06 | 50.16 | 43.64 | 45.99 | 44.95 | 42.33 | 42.47 | - |
| Spry4 | 0.34 | 0.29 | 0.27 | 0.07 | 0.12 | 0.03 | 0.35 | 0.22 | 0.2 | 0 | 0.05 | 0.01 | - |
| Egr2 | 8.57 | 7.52 | 9.27 | 2.37 | 2.15 | 2.97 | 0.78 | 3.03 | 1.57 | 0.94 | 0.33 | 0.66 | - |
| Epsti1 | 39.01 | 43.35 | 35.58 | 41.42 | 33.12 | 36.24 | 76.21 | 46.98 | 56.78 | 78.14 | 57.49 | 82.78 | - |
| Zan | 0 | 0 | 0 | 0.17 | 0.08 | 0.18 | 0.01 | 0 | 0 | 0.25 | 0.19 | 0.31 | - |
| Tnfaip3 | 11.45 | 10.76 | 14.51 | 19.33 | 22.06 | 17.96 | 25.41 | 28.33 | 26.88 | 33.14 | 30.13 | 29.11 | - |
| Tmem9 | 22.52 | 23.79 | 22.4 | 30.69 | 26.21 | 27.17 | 32.35 | 25.87 | 28.52 | 30.29 | 30.01 | 30.83 | - |
| Irf8 | 56.04 | 72.5 | 70.43 | 44.52 | 51.33 | 48.21 | 48.78 | 47.04 | 50.12 | 36.81 | 42.5 | 39.02 | - |
| Sec62 | 12.06 | 11.84 | 12.42 | 13.26 | 12.89 | 14.07 | 13.96 | 14.26 | 13.38 | 14.39 | 16.1 | 16.89 | - |
| Abcc1 | 18.69 | 18.8 | 20.89 | 15.31 | 15.49 | 14.41 | 15.41 | 17.63 | 18.99 | 11.3 | 14.08 | 11.6 | - |
| AW209491 | 15.7 | 15.37 | 16.24 | 10.38 | 11.67 | 12.04 | 10.06 | 11.88 | 12.6 | 8.85 | 8.4 | 7.35 | - |
| Ipo7 | 49.86 | 49.62 | 52.45 | 54.37 | 56.71 | 53.14 | 42.5 | 51.65 | 53.07 | 41.84 | 46.65 | 36.86 | - |
| Cxcl10 | 0.99 | 1.27 | 0.34 | 0.57 | 0.4 | 0.41 | 0.05 | 0.21 | 0.15 | 0.05 | 0.11 | 0.1 | - |
| Kifc2 | 0.47 | 0.56 | 0.22 | 1.77 | 1.35 | 0.85 | 1.1 | 0.71 | 1.42 | 2.07 | 2.22 | 2.71 | - |
| Mmachc | 14.87 | 13.38 | 13.31 | 10 | 12.39 | 9.74 | 6 | 9.86 | 10.31 | 6.24 | 7.89 | 6.64 | - |
| Dgkh | 1.42 | 1.42 | 1.46 | 0.66 | 0.25 | 0.58 | 1.42 | 1.26 | 1.5 | 0.58 | 0.67 | 0.7 | - |
| Aip | 55.46 | 56.41 | 57.65 | 53.27 | 54.49 | 48.23 | 55.77 | 54.63 | 56.53 | 43.26 | 46.92 | 49.82 | - |
| Telo2 | 9.92 | 10.34 | 10.12 | 7.98 | 7.4 | 6.91 | 5.27 | 6.6 | 7.07 | 4.51 | 4.76 | 6.41 | - |
| Rab35 | 4.04 | 5.84 | 4.62 | 3.62 | 4.39 | 2.36 | 3.4 | 3.31 | 2.85 | 1.9 | 2.91 | 3.69 | - |
| Rce1 | 38.48 | 39.43 | 36.46 | 33.79 | 32.94 | 34.27 | 31.88 | 35.22 | 34.99 | 29.47 | 28.9 | 30.54 | - |
| Spice1 | 3.59 | 2.97 | 3.95 | 5.37 | 4.73 | 4.22 | 6.88 | 5.1 | 5.02 | 6.19 | 5.03 | 6.5 | - |
| Acox1 | 7.04 | 6.56 | 6.52 | 8.28 | 7.6 | 8.82 | 10.34 | 8.48 | 7.9 | 8.74 | 8.95 | 8.6 | - |
| Zfp960 | 14.7 | 17.46 | 14.07 | 19.84 | 21.97 | 21.79 | 24.96 | 20.48 | 17.22 | 31.22 | 32.25 | 28.79 | - |
| Cd7 | 0 | 0.22 | 0 | 1.76 | 0.06 | 0.25 | 6.85 | 0.06 | 1.34 | 18.26 | 4.98 | 15.69 | - |
| Mef2a | 1.05 | 1.2 | 1.11 | 1.5 | 1.26 | 0.98 | 2.02 | 1.29 | 2.19 | 2.42 | 2.49 | 2.67 | - |
| Ovca2 | 9.28 | 9.15 | 9.36 | 6.85 | 6.77 | 4.99 | 4.45 | 4.98 | 6.96 | 4.38 | 5.86 | 5.58 | - |
| Fxr1 | 63.03 | 58.68 | 62.53 | 66.56 | 68.89 | 70.58 | 70.09 | 76.18 | 65.13 | 70.53 | 89.52 | 66.27 | - |
| Spin4 | 0.34 | 0.45 | 0.53 | 0.17 | 0.31 | 0.37 | 0.15 | 0.21 | 0.23 | 0.15 | 0.1 | 0.06 | - |
| Pik3ap1 | 9.19 | 7.41 | 8.83 | 10.39 | 9.43 | 8.95 | 24.9 | 16.1 | 19.65 | 23.73 | 20.76 | 25.24 | - |
| Selenbp2 | 0.24 | 0.11 | 0.38 | 0.97 | 0.72 | 0.72 | 2.36 | 0.68 | 1.87 | 2.16 | 2.63 | 3.37 | - |
| Tagln3 | 0.79 | 0.2 | 0.88 | 0.16 | 0.11 | 0.11 | 0 | 0.71 | 0.42 | 0.05 | 0 | 0 | - |
| Atg9a | 3.9 | 5.26 | 4.64 | 8.18 | 6.58 | 7.91 | 9.54 | 7.44 | 6.78 | 10.94 | 9.81 | 11.24 | - |
| Rapgef6 | 5.27 | 5.55 | 5.62 | 9.12 | 7.74 | 9.06 | 8.45 | 6.67 | 7.71 | 9.68 | 9.84 | 9.32 | - |
| Tmed1 | 7.46 | 7.99 | 9.43 | 7.2 | 5.94 | 6.92 | 5.64 | 3.8 | 4.84 | 4.83 | 5.01 | 3.83 | - |
| Diablo | 30.29 | 30.85 | 26.63 | 16.69 | 21.82 | 20.39 | 15.03 | 20.11 | 19.89 | 11.51 | 11.25 | 12.54 | - |
| Cdc42ep2 | 0.04 | 0.16 | 0.25 | 1.82 | 0.46 | 1.15 | 2.69 | 0.79 | 1.69 | 1.57 | 2.03 | 1.62 | - |
| Nab1 | 4.69 | 4.32 | 4.01 | 4.5 | 3.91 | 3.76 | 3.34 | 3.71 | 3.22 | 2.95 | 2.52 | 3.48 | - |
| Rpgr | 3.02 | 2.95 | 3.52 | 1.73 | 2.3 | 1.85 | 1.94 | 2.54 | 2.11 | 1.42 | 1.36 | 1.8 | - |
| Syap1 | 1.11 | 1.96 | 1.68 | 2.73 | 2.13 | 2.31 | 2.18 | 2.61 | 2.76 | 3.24 | 3.46 | 2.54 | - |
| Recql4 | 3.91 | 3.52 | 3.61 | 1.78 | 2.62 | 2.57 | 1.07 | 1.78 | 1.91 | 0.47 | 1.22 | 0.71 | - |
| Comtd1 | 10.01 | 10.49 | 11.44 | 8.63 | 9.33 | 7.71 | 6.9 | 9.96 | 10.13 | 6.31 | 7.12 | 7.1 | - |
| Trat1 | 4.34 | 4.34 | 6.01 | 0.18 | 0.82 | 1.31 | 5.88 | 5.15 | 4.47 | 1.67 | 0.9 | 0.93 | - |
| Snapin | 31.14 | 29.75 | 29.07 | 37.91 | 37.29 | 36.62 | 44.01 | 39.07 | 38.01 | 48.41 | 44.35 | 46.98 | - |
| Ypel4 | 0.15 | 0.39 | 0.07 | 1.51 | 1.08 | 1.44 | 2.22 | 1.8 | 1.36 | 1.67 | 1.64 | 2.07 | - |
| Ugt1a5 | 0.5 | 0.19 | 0.4 | 0.05 | 0.04 | 0.05 | 0 | 0.44 | 0.21 | 0 | 0.05 | 0 | - |
| Usp40 | 5.11 | 4.1 | 4.81 | 3.54 | 3.18 | 3.14 | 3.16 | 3.21 | 3.63 | 2.55 | 2.27 | 2.84 | - |
| Prkcq | 24.78 | 30.11 | 22.47 | 38.73 | 32.52 | 34.09 | 53.42 | 39.33 | 39.55 | 53.73 | 56.76 | 59.33 | - |
| Rab9 | 16.48 | 19.73 | 20.27 | 27.54 | 23.19 | 26.87 | 33.16 | 29.2 | 28.16 | 39.74 | 44.57 | 35.91 | - |
| Clptm1l | 11 | 12.52 | 12.32 | 6.78 | 7.09 | 7.51 | 6.13 | 9.07 | 7.17 | 7.05 | 5.39 | 3.95 | - |
| Lcn4 | 1.33 | 1.13 | 0.21 | 0.98 | 1.13 | 0.58 | 11.73 | 1.82 | 5.68 | 5.84 | 3.98 | 6.48 | - |
| Ergic1 | 35.07 | 33.32 | 33.63 | 29.07 | 30.26 | 29.63 | 22.6 | 28 | 24.84 | 20.41 | 23.05 | 20.25 | - |
| Rps11 | 1699.88 | 1791.45 | 1691.84 | 1908.22 | 1817.13 | 1876.37 | 2076.11 | 1915.11 | 2018.1 | 2049.12 | 2127.12 | 2059.24 | - |

| | | | | | | | | | | | | |
|---|---|---|---|---|---|---|---|---|---|---|---|---|
| Gsk3b | 11.51 | 10.25 | 10.53 | 11.8 | 12.74 | 11.77 | 12.42 | 12.03 | 11.99 | 13.26 | 12.44 | 12.44 | - |
| Mis18bp1 | 9.82 | 6.33 | 7.08 | 5.33 | 7.72 | 7.73 | 5.03 | 4.05 | 3.67 | 4.51 | 5.62 | 4.56 | - |
| Cdip1 | 5.97 | 5.19 | 5.12 | 7.56 | 5.89 | 6.7 | 5.9 | 6.56 | 6.52 | 8.24 | 7.19 | 8.14 | - |
| Vbp1 | 147.26 | 145.69 | 146.07 | 127.07 | 132.35 | 134.93 | 114.15 | 136.91 | 125.35 | 107.86 | 121.29 | 106.09 | - |
| Spryd7 | 27.6 | 25.87 | 27.1 | 16.53 | 15.92 | 16.98 | 19.42 | 20.96 | 22.67 | 12.19 | 13.64 | 15.25 | - |
| Ifngr1 | 144.22 | 170.78 | 125.47 | 244.18 | 183.96 | 193.08 | 303.44 | 227.12 | 243.94 | 355.45 | 274.14 | 361.93 | - |
| Vrk2 | 11.53 | 16.16 | 13.54 | 11.83 | 13.67 | 12.73 | 11.23 | 11.43 | 10.34 | 8.94 | 8.4 | 6.84 | - |
| Rec8 | 1.11 | 0.51 | 0.53 | 0.8 | 0.7 | 0.93 | 1.82 | 1.49 | 1.2 | 1.39 | 2.25 | 1.65 | - |
| Sars2 | 14.68 | 14.9 | 14.58 | 9.5 | 11.54 | 12.09 | 9.12 | 10.98 | 10.79 | 10.27 | 8.55 | 8.9 | - |
| Tm9sf1 | 33.69 | 32.91 | 32.3 | 27.08 | 27.89 | 25.85 | 19.76 | 24.63 | 20.51 | 21.53 | 17.76 | 23.62 | - |
| Col20a1 | 0.04 | 0 | 0.1 | 0.57 | 0.49 | 0.62 | 0.47 | 0.29 | 0.32 | 0.67 | 0.75 | 0.21 | - |
| Otub2 | 1.34 | 2.07 | 1.67 | 1.78 | 2.6 | 1.95 | 3.08 | 3.74 | 4.79 | 2.93 | 3.97 | 2.62 | - |
| Styk1 | 0.47 | 0.83 | 0.34 | 0.82 | 0.74 | 0.83 | 2.36 | 2.29 | 2.67 | 1.47 | 1.26 | 2.49 | - |
| Olr1 | 4.18 | 2.56 | 3.96 | 0.03 | 0.09 | 0.02 | 0.34 | 0.67 | 1.32 | 0.1 | 0.02 | 0.11 | - |
| P2ry10 | 14.68 | 15.53 | 16.02 | 29.13 | 22.98 | 24.99 | 31.19 | 21.59 | 25.72 | 39.35 | 30.13 | 39.2 | - |
| Wdr62 | 4.36 | 4.32 | 4.91 | 3.28 | 3.96 | 3.68 | 2.31 | 2.34 | 2.36 | 2.34 | 3.52 | 1.8 | - |
| Zc3hc1 | 31.87 | 30.3 | 29.65 | 24.98 | 26.97 | 25.03 | 25.16 | 30.42 | 24.02 | 20.32 | 25.73 | 20.38 | - |
| Ddx51 | 1.07 | 1.83 | 1.61 | 0.99 | 0.97 | 1.12 | 0.74 | 1.02 | 0.85 | 0.76 | 0.59 | 0.44 | - |
| Akr1a1 | 311.66 | 297.19 | 292.36 | 279.9 | 259.58 | 279.89 | 287.35 | 324.07 | 286.52 | 242.59 | 284.33 | 242.62 | - |
| Ccdc6 | 13.24 | 13.57 | 13.13 | 19.83 | 18.01 | 18.62 | 12.76 | 14.06 | 11.8 | 15.69 | 17.97 | 14.8 | - |
| 2610204G22Rik | 0.76 | 0.3 | 0.45 | 1.35 | 0.58 | 1.64 | 1.12 | 0.6 | 0.72 | 1.47 | 2.22 | 1.66 | - |
| Ncf1 | 23.75 | 24.29 | 24.52 | 48.38 | 38.8 | 36.76 | 32.14 | 27.67 | 27.6 | 37.93 | 28.27 | 42.09 | - |
| Rhog | 87.57 | 83.51 | 83.28 | 157.01 | 129.59 | 140.21 | 184.71 | 135.22 | 145.67 | 212.43 | 219.33 | 235.13 | - |
| Tm4sf5 | 2.46 | 1.13 | 1.44 | 1.1 | 0.77 | 0.85 | 1 | 1.32 | 2.36 | 0.53 | 0.57 | 0.3 | - |
| Dusp11 | 17.78 | 20.72 | 19.73 | 25.71 | 25.58 | 25.53 | 26.54 | 24.88 | 28.2 | 29.33 | 31.66 | 29.58 | - |
| Egfl7 | 12.65 | 14.55 | 13.82 | 9.45 | 9.12 | 9.14 | 11.8 | 12.87 | 10.32 | 6.73 | 5.45 | 5.99 | - |
| Ffar4 | 0.49 | 0.42 | 0.5 | 0.23 | 0 | 0.23 | 0 | 0.08 | 0.27 | 0 | 0.04 | 0.04 | - |
| Tnnt3 | 0 | 0 | 0 | 0.18 | 0.18 | 0.18 | 0.28 | 0 | 0.15 | 0.71 | 0.3 | 0.5 | - |
| Idh1 | 12.46 | 10.73 | 11.42 | 13.65 | 13.18 | 16.08 | 15.73 | 11.16 | 11.04 | 16.56 | 17.43 | 18.87 | - |
| H13 | 39.78 | 42.83 | 40.91 | 50.04 | 45.34 | 44.23 | 55.88 | 49.19 | 50.35 | 55.74 | 55.29 | 60.9 | - |
| Bend6 | 0.77 | 1.42 | 0.94 | 1.76 | 2.59 | 2.72 | 2.88 | 3.28 | 2.65 | 3.77 | 3.46 | 5.73 | - |
| Atpbd4 | 17.33 | 13.58 | 15.5 | 8.88 | 11.49 | 9.78 | 10.43 | 14.26 | 11.94 | 6.49 | 6.16 | 6.26 | - |
| Cntnap1 | 0.01 | 0.01 | 0.03 | 0.07 | 0.05 | 0.05 | 0.18 | 0.18 | 0.13 | 0.2 | 0.21 | 0.15 | - |
| Gmds | 37.77 | 37.57 | 30.11 | 22.18 | 25.88 | 22.26 | 27.67 | 30.81 | 31.12 | 21.14 | 22.04 | 22.4 | - |
| AF529169 | 0.07 | 0.12 | 0.01 | 0.18 | 0.22 | 0.17 | 0.28 | 0.1 | 0.2 | 1.58 | 0.93 | 1.43 | - |
| Acot2 | 5.9 | 6.54 | 5.28 | 4.18 | 3.96 | 3.46 | 3.97 | 4.51 | 4.67 | 3.26 | 3.53 | 2.96 | - |
| Slc9a8 | 7.81 | 8.1 | 6.97 | 7.08 | 6.26 | 6.6 | 4.75 | 6.47 | 6.08 | 5.91 | 3.33 | 4.87 | - |
| Zc3hav1 | 29.66 | 32.94 | 33.7 | 43.86 | 38.14 | 40.02 | 46.35 | 35.93 | 39.9 | 50.79 | 51.61 | 54.01 | - |
| Vrk1 | 27.11 | 27.27 | 26.37 | 20.7 | 20.82 | 20.39 | 15.67 | 16.16 | 15.7 | 11.89 | 11.06 | 11.9 | - |
| Phf8 | 3.01 | 2.83 | 3.63 | 4.53 | 4.39 | 5.71 | 6.33 | 4.47 | 5.54 | 5.96 | 6.81 | 6.57 | - |
| Tprgl | 6.03 | 6.12 | 6.06 | 7.59 | 7.24 | 7.77 | 9.74 | 8.97 | 7.8 | 10.94 | 9.56 | 10.16 | - |
| Ncor1 | 9.18 | 7.89 | 8.89 | 12.92 | 11.93 | 12.83 | 13.07 | 11.59 | 12.86 | 14.84 | 16.16 | 13.92 | - |
| Ska1 | 9.63 | 8.79 | 8.27 | 4.96 | 7.35 | 6.17 | 4.35 | 3.93 | 2.96 | 3.41 | 2.59 | 2.84 | - |
| Pus3 | 12.87 | 13.36 | 12.42 | 11.62 | 8.94 | 9.9 | 9.68 | 9.31 | 10.77 | 6.65 | 6.92 | 7.89 | - |
| Fry | 0.03 | 0.05 | 0.05 | 0.24 | 0.19 | 0.2 | 0.23 | 0.13 | 0.12 | 0.46 | 0.32 | 0.24 | - |
| Aldh9a1 | 64.17 | 57.92 | 61.18 | 42.59 | 46.25 | 49.55 | 46.31 | 53.84 | 48.38 | 36.43 | 39.9 | 40.41 | - |
| Helz | 7.67 | 6.25 | 6.28 | 8.65 | 8.18 | 7.77 | 7.54 | 7.51 | 7.56 | 8.75 | 9.52 | 8.9 | - |
| Chmp5 | 89.46 | 86.79 | 88.76 | 83.46 | 84.35 | 89.69 | 84.98 | 91.3 | 90.37 | 72.59 | 76.91 | 72.63 | - |
| Lins | 3.51 | 4.02 | 3.91 | 3.18 | 2.93 | 3.35 | 2.8 | 3.44 | 3.95 | 2.28 | 2.48 | 2.03 | - |
| Supv3l1 | 16.06 | 19.11 | 20.29 | 13.14 | 14.26 | 14.09 | 11.49 | 15.55 | 13.22 | 11.13 | 11.41 | 9.48 | - |
| Reep4 | 52.46 | 47.48 | 47.76 | 41.33 | 49.04 | 44.07 | 29.03 | 31.97 | 30.17 | 32.6 | 27.42 | 27.09 | - |
| Slc39a6 | 59.6 | 59.96 | 57.42 | 36.61 | 35.42 | 34.86 | 39.08 | 45.25 | 49.33 | 26.89 | 23.94 | 28.77 | - |
| Ndufaf5 | 10.93 | 9.97 | 11.65 | 7.87 | 8.59 | 9.2 | 8.65 | 10.6 | 9.29 | 6.83 | 8.23 | 6.94 | - |
| Frat2 | 0 | 0.03 | 0.05 | 0.09 | 0.09 | 0.07 | 0.19 | 0.08 | 0.15 | 0.42 | 0.57 | 0.37 | - |
| 1200014J11Rik | 11.7 | 11.56 | 11.52 | 10.91 | 10.65 | 11.85 | 10.1 | 10.86 | 10.12 | 8.59 | 8.98 | 9.49 | - |
| Phtf2 | 5.12 | 4.8 | 5.4 | 3.12 | 2.95 | 2.78 | 2.55 | 3.08 | 3.19 | 1.88 | 1.53 | 1.39 | - |
| Mast3 | 4.42 | 4.48 | 4.5 | 7.94 | 6.31 | 6.69 | 6.99 | 5.6 | 5.47 | 8.58 | 8.09 | 9.14 | - |
| Pkp4 | 5.41 | 4.13 | 4.43 | 3.06 | 3.66 | 2.65 | 2.31 | 3.71 | 3.27 | 3.73 | 2.79 | 2.53 | - |
| Pef1 | 38.69 | 37.59 | 36.95 | 40.59 | 38.48 | 39.35 | 45.58 | 44.57 | 48.74 | 48.47 | 44.15 | 46.75 | - |
| Wars | 81.36 | 81 | 86.74 | 43.09 | 51.71 | 47.86 | 45.85 | 78.63 | 65.97 | 31.81 | 29.59 | 34.63 | - |
| Mmgt1 | 7.36 | 8.75 | 8.39 | 7.01 | 7.43 | 6.87 | 5.09 | 7.15 | 5.93 | 6.32 | 6.5 | 5.06 | - |
| Ppme1 | 51.58 | 42.2 | 48.45 | 48.75 | 45.86 | 47.37 | 70.79 | 56 | 59.22 | 52.65 | 58.02 | 63.36 | - |
| Lims1 | 11.49 | 11.46 | 10.3 | 17.57 | 14.83 | 17.3 | 20.7 | 17.27 | 18.55 | 19.25 | 25.29 | 20.9 | - |
| Mrgpre | 2.56 | 2.07 | 2.77 | 3.44 | 3.2 | 3.67 | 5.48 | 4.59 | 4.69 | 7.84 | 5.33 | 7.48 | - |
| Gnai2 | 14.82 | 15.83 | 17.57 | 19.91 | 20.56 | 21.47 | 21.12 | 20.17 | 19.93 | 22.99 | 21.32 | 23.18 | - |
| Uba1y | 0.7 | 0.89 | 0.91 | 1.04 | 1.31 | 1.41 | 1.52 | 1.01 | 1.27 | 1.46 | 1.81 | 1.52 | - |
| Cep192 | 3.46 | 2.61 | 3.67 | 4.44 | 4.39 | 4.63 | 4.71 | 4.09 | 4.83 | 4.86 | 6.62 | 4.86 | - |
| Slu7 | 16.7 | 17.3 | 14.87 | 19.62 | 19.38 | 18.56 | 20.51 | 18.54 | 19.9 | 22.3 | 21.71 | 21.71 | - |
| Hist1h3i | 6.97 | 6.39 | 6.16 | 3.96 | 3.86 | 4.09 | 1.73 | 5.23 | 3.01 | 2.08 | 2.99 | 2.38 | - |
| Phldb1 | 1.33 | 0.9 | 1.58 | 1.34 | 1.12 | 1.17 | 1.18 | 1.38 | 0.79 | 0.85 | 0.44 | 0.6 | - |
| Slc25a45 | 2.87 | 4.03 | 2.89 | 6.31 | 4.43 | 4.94 | 9.91 | 5.7 | 5.06 | 9.74 | 8.98 | 11.42 | - |
| Stat3 | 84.76 | 79.02 | 88.34 | 105.32 | 103.04 | 103.37 | 105.14 | 112.95 | 101.68 | 118.42 | 119.87 | 123.19 | - |
| Coq9 | 25.93 | 29.63 | 28.61 | 39.05 | 38.42 | 40.64 | 43.65 | 36.64 | 37.65 | 47.24 | 51.18 | 46.34 | - |
| Polq | 1.29 | 1.47 | 1.18 | 0.65 | 1.06 | 0.84 | 0.45 | 0.58 | 0.44 | 0.24 | 0.46 | 0.35 | - |
| Tob2 | 9.29 | 8.85 | 10.53 | 6.66 | 7.27 | 6.4 | 6.07 | 7.23 | 7.64 | 4.84 | 3.95 | 4.74 | - |
| Acp5 | 27.91 | 29.46 | 26.35 | 46.35 | 35.89 | 32.25 | 58.62 | 38.54 | 49.22 | 73.04 | 57.97 | 78.88 | - |
| Mtbp | 13.6 | 12.82 | 11.12 | 5.39 | 6.26 | 4.91 | 4.88 | 6.87 | 6.22 | 3.22 | 3.09 | 2.87 | - |
| Pih1d1 | 21.73 | 24.54 | 21.44 | 17.59 | 16.39 | 17.64 | 15.07 | 16 | 16.74 | 11.99 | 13.31 | 14.16 | - |
| Fdx1l | 63.54 | 73.48 | 79.48 | 53.95 | 57.14 | 57.84 | 59.54 | 77.19 | 66.92 | 54.73 | 56.46 | 51.12 | - |
| Cep85 | 14.97 | 12.84 | 14.27 | 15.97 | 18 | 17.6 | 20.5 | 18.69 | 18.63 | 20.45 | 22.93 | 18.55 | - |
| Slc38a8 | 0.07 | 0.07 | 0.06 | 0.1 | 0.1 | 0.27 | 0.26 | 0.25 | 0.51 | 0.41 | 0.61 | 0.15 | - |
| Actr3b | 6.57 | 9.93 | 6.1 | 1.99 | 1.17 | 1.6 | 2.69 | 2.03 | 1.85 | 0.43 | 0.18 | 0.4 | - |
| Jak3 | 62.41 | 66.85 | 70.29 | 139.84 | 115.71 | 121.57 | 136.59 | 103.67 | 119.75 | 163.36 | 142.32 | 183.09 | - |
| Arih2 | 25.57 | 25.82 | 26.09 | 33.77 | 28.85 | 31.36 | 33.56 | 33.93 | 32.75 | 37.19 | 36.92 | 39.19 | - |
| Aim1l | 0.8 | 0.9 | 0.66 | 1.46 | 0.97 | 1.53 | 1.83 | 1.15 | 1.33 | 1.53 | 1.06 | 1.56 | - |
| Exph5 | 0.95 | 0.68 | 0.73 | 0.1 | 0.14 | 0.05 | 0.2 | 0.64 | 0.5 | 0.23 | 0.08 | 0.03 | - |
| Pik3c2a | 1.01 | 1.12 | 1.1 | 2.04 | 1.45 | 1.86 | 2.5 | 1.61 | 2.34 | 2.52 | 2.53 | 2.21 | - |
| Cd8a | 113 | 131.86 | 103.15 | 130.94 | 107.41 | 112.13 | 87.19 | 67.91 | 77.06 | 90.28 | 81.72 | 101.8 | - |
| Adamtsl3 | 0 | 0 | 0 | 0 | 0.02 | 0 | 0.02 | 0 | 0 | 0.04 | 0.08 | 0.06 | - |
| Tmco4 | 6.26 | 4.96 | 6.84 | 7.46 | 9.65 | 9.48 | 6.56 | 8.14 | 7.19 | 9.09 | 7.41 | 8.26 | - |
| Atg13 | 8.32 | 7.15 | 7.73 | 11.78 | 10.96 | 12.12 | 12.3 | 12.31 | 11.47 | 14.41 | 13.23 | 15.97 | - |
| Mettl9 | 6.64 | 9.42 | 9.73 | 14.15 | 14 | 10.36 | 15.03 | 16.91 | 15.83 | 18.42 | 18.9 | 14.04 | - |
| Psph | 97.51 | 98.05 | 103.52 | 52.81 | 59.5 | 55.33 | 55.23 | 81.66 | 80.28 | 36.74 | 38.38 | 38.78 | - |
| Irf5 | 18.55 | 16.56 | 16.57 | 14.88 | 14.12 | 15.53 | 13.03 | 13.64 | 14.24 | 12.37 | 8.97 | 12.28 | - |
| Arih1 | 6.05 | 5.52 | 6.65 | 7.03 | 6.71 | 6.92 | 5.38 | 7.27 | 6.94 | 7.61 | 7.43 | 8.09 | - |
| Slc17a6 | 0.77 | 0.62 | 1.14 | 0.38 | 1.01 | 0.68 | 0.65 | 0.57 | 0.61 | 0.34 | 0.07 | 0.48 | - |
| Kat2a | 8.38 | 11.65 | 12.62 | 9.86 | 8.57 | 7.5 | 7.41 | 10.3 | 7.52 | 8 | 6.59 | 7 | - |
| Maf | 0.02 | 0.03 | 0 | 0.33 | 0.58 | 0.56 | 0.55 | 0.32 | 1.03 | 4.45 | 4.65 | 4.18 | - |
| Ppp1r12b | 1.92 | 2.1 | 2.06 | 3.29 | 2.71 | 2.79 | 2.71 | 3.27 | 3.27 | 3.58 | 4.27 | 4.33 | - |
| Mbd5 | 0.71 | 0.74 | 0.75 | 1 | 0.96 | 1.02 | 0.91 | 1.26 | 0.91 | 1.23 | 0.9 | 1.62 | - |
| Vps41 | 22.4 | 19.33 | 20.43 | 25.48 | 24.5 | 25.45 | 24.55 | 25.19 | 26.46 | 30.74 | 30.11 | 27.74 | - |
| Map1lc3b | 67.36 | 64.71 | 61.74 | 86.5 | 75.77 | 78.35 | 120.29 | 91.98 | 88.66 | 134.56 | 128.55 | 141.81 | - |
| Metap1 | 72.54 | 61.42 | 70.79 | 58.56 | 58.63 | 59.57 | 51.04 | 66.62 | 65.44 | 46.21 | 59.88 | 41.97 | - |
| Spsb2 | 0.97 | 1.04 | 0.83 | 1.66 | 1.11 | 1.51 | 1.46 | 0.75 | 1.46 | 2.13 | 2.82 | 2.44 | - |
| Rictor | 1.37 | 1.59 | 1.55 | 3.55 | 2.84 | 2.77 | 3.1 | 2.19 | 2.56 | 3.99 | 3.81 | 3.61 | - |
| Pigv | 3.69 | 4.52 | 4.18 | 6.24 | 5.15 | 5.1 | 5.63 | 5.31 | 5.73 | 5.41 | 5.89 | 7.26 | - |
| 1700113A16Rik | 2.32 | 2.96 | 3.11 | 4.82 | 3.43 | 4.74 | 4.91 | 4.54 | 3.55 | 6.46 | 5.27 | 6.04 | - |
| Ldhb | 17.04 | 15.79 | 14.75 | 14.6 | 16.74 | 14.58 | 14.74 | 13.39 | 12.93 | 11.17 | 12.13 | 10.81 | - |
| Tmem63a | 3.3 | 2.75 | 2.74 | 4.88 | 5.24 | 4.63 | 9.65 | 7.2 | 5.67 | 9.34 | 8.43 | 9.97 | - |
| Gm6460 | 1.88 | 7.75 | 0.48 | 0 | 0.28 | 0 | 11.17 | 6.93 | 12.79 | 0.25 | 1.05 | 0.23 | - |
| Sv2c | 1.84 | 1.49 | 1.98 | 17.67 | 24.44 | 25.05 | 0.97 | 1.96 | 1.51 | 12.79 | 17.33 | 11.94 | - |
| Rrp1b | 28.78 | 28.44 | 29 | 25.03 | 26.87 | 25.13 | 15.75 | 21.11 | 20.44 | 13.54 | 17.36 | 14.18 | - |
| Pdf | 14.72 | 18.83 | 18.83 | 10.53 | 14.22 | 11.42 | 9.76 | 12.98 | 10.51 | 10.8 | 8.31 | 8.93 | - |
| Gmfb | 21.38 | 21.62 | 21.89 | 21.18 | 21.66 | 18.21 | 16.55 | 18.15 | 18.42 | 14.8 | 16.15 | 13.27 | - |
| Gpx7 | 3.15 | 3.64 | 3.96 | 1.35 | 2.16 | 1.94 | 2.18 | 1.89 | 1.86 | 1.04 | 0.46 | 1.19 | - |
| Trp53bp2 | 3.53 | 4.93 | 4.91 | 3.14 | 3.73 | 3.63 | 3.96 | 3.05 | 3.78 | 2.74 | 3.41 | 2.45 | - |
| Ttll4 | 5.77 | 6.28 | 5.05 | 3.02 | 3.74 | 2.71 | 2.32 | 2.82 | 3.29 | 2.06 | 2.07 | 1.42 | - |
| Cdc25a | 7.98 | 8.58 | 9.21 | 7.85 | 7.11 | 8.77 | 6.95 | 7.55 | 7.8 | 7.08 | 6.25 | 6.25 | - |
| Golim4 | 8.04 | 8.78 | 8.21 | 7.36 | 6.85 | 6.21 | 5.96 | 7.12 | 7.97 | 6.3 | 5.08 | 5.73 | - |
| Lmo2 | 1.2 | 0.96 | 1.07 | 0 | 0.08 | 0.25 | 0.04 | 0.19 | 0.11 | 0 | 0 | 0.04 | - |
| Tmem127 | 12.36 | 13.64 | 12.77 | 19.16 | 19.6 | 19.48 | 14.22 | 15.39 | 14.13 | 19.75 | 16.1 | 19.38 | - |
| Top3b | 14.43 | 14.63 | 14.03 | 10.02 | 13.94 | 12.5 | 10.37 | 11.53 | 12.38 | 11.01 | 8.96 | 9.92 | - |
| Yif1a | 31.28 | 30.89 | 29.33 | 22.52 | 19.61 | 24.99 | 21.98 | 23.8 | 20.3 | 11.95 | 14.53 | 15.52 | - |
| Fam105a | 4.13 | 4.15 | 4.45 | 6.35 | 6.5 | 5.47 | 8.74 | 7.61 | 6.9 | 10.8 | 8.48 | 9.55 | - |
| Mtm1 | 2.76 | 2.72 | 2.62 | 4.44 | 3.52 | 3.34 | 4.85 | 3.1 | 4.89 | 5.36 | 4.27 | 4.71 | - |
| Foxj2 | 1.12 | 1.29 | 0.93 | 1.99 | 1.41 | 1.68 | 1.35 | 1.28 | 1.52 | 2.11 | 2.14 | 1.93 | - |
| Asl | 36.44 | 35.04 | 37.8 | 30.18 | 32.29 | 32.25 | 32.73 | 30.23 | 29.93 | 26.69 | 23.14 | 26.13 | - |
| Timeless | 9.03 | 10.26 | 10.53 | 5.39 | 6.5 | 5.85 | 3.85 | 4.78 | 3.75 | 1.76 | 3.25 | 1.94 | - |
| Mark2 | 7.19 | 7.6 | 7.19 | 8.98 | 7.9 | 8.05 | 8.64 | 5.83 | 7.72 | 10.84 | 8.71 | 11.27 | - |
| Fyb | 20.67 | 21.35 | 19.67 | 47.05 | 41.48 | 45.74 | 57.77 | 39.65 | 44.87 | 73.42 | 66 | 77.58 | - |
| Map3k3 | 2.14 | 1.96 | 2.49 | 3.79 | 3.27 | 3.75 | 3.37 | 2.97 | 3.19 | 4.4 | 3.35 | 4.5 | - |
| Bfar | 15.29 | 15.46 | 15.22 | 19.28 | 19.24 | 17.1 | 16.36 | 17.06 | 17.72 | 19.13 | 19.64 | 22.17 | - |
| Rdh1 | 1.16 | 1.72 | 1.31 | 2.47 | 1.69 | 1.9 | 1.99 | 1.99 | 1.52 | 2.64 | 2.58 | 2.16 | - |
| Cd82 | 152.53 | 153.32 | 141.56 | 124.27 | 149.42 | 134.01 | 124.57 | 150.44 | 138.17 | 99.24 | 108.62 | 100.34 | - |
| Ankle1 | 8.6 | 5.46 | 5.03 | 3.59 | 5.15 | 6.06 | 2.07 | 1.44 | 1.16 | 1.64 | 1.33 | 1.34 | - |
| Ttyh2 | 0.51 | 0.4 | 0.27 | 0.33 | 0.33 | 0.33 | 1.06 | 1.1 | 0.36 | 1.24 | 0.87 | 1.02 | - |
| Commd8 | 14.01 | 13.46 | 13.92 | 15.64 | 16.7 | 16.84 | 16.36 | 13.21 | 15.86 | 18.57 | 17 | 16.23 | - |
| Fam184b | 0.01 | 0.06 | 0.04 | 0.11 | 0.03 | 0.02 | 0.2 | 0.06 | 0.12 | 0.31 | 0.21 | 0.34 | - |
| Cd38 | 4.87 | 4.87 | 3.67 | 1.82 | 2.54 | 1.71 | 10.09 | 4.09 | 6.11 | 2.56 | 1.48 | 2.86 | - |
| Ctss | 49.32 | 48.27 | 52.57 | 95.85 | 90.36 | 92.12 | 66.98 | 61.66 | 58.58 | 100.59 | 100.76 | 108.93 | - |
| Pou6f1 | 0.11 | 0.03 | 0 | 0.66 | 0.63 | 0.74 | 0.84 | 0.13 | 0.62 | 1.13 | 1.02 | 1.23 | - |
| Pcmtd2 | 4.91 | 4.91 | 4.36 | 10.88 | 9.41 | 8.83 | 13.99 | 9.49 | 10.21 | 16.94 | 15.84 | 16.13 | - |
| Rela | 19.23 | 17.01 | 19.03 | 27.13 | 22.51 | 23.56 | 25.95 | 22.46 | 25.19 | 31.12 | 25.03 | 31.01 | - |
| Slc7a5 | 9.07 | 10 | 9.03 | 5.47 | 6.52 | 4.88 | 5.87 | 9.15 | 8.89 | 3.08 | 3.81 | 3.07 | - |
| B2m | 2248.98 | 2326.37 | 2194.87 | 3331.04 | 3250.91 | 3308.03 | 2938.26 | 2589.15 | 2782.9 | 3283.67 | 3343.06 | 3387.17 | - |
| Cox6a1 | 427.9 | 425.44 | 418.03 | 328.23 | 338.12 | 348.98 | 374.87 | 416.97 | 402.59 | 263.39 | 285.61 | 274.16 | - |
| Setd2 | 2.58 | 3.06 | 2.82 | 4.17 | 3.73 | 3.98 | 2.9 | 3.14 | 2.97 | 4.88 | 4.75 | 3.95 | - |
| Cx3cl1 | 0.18 | 0.11 | 0.17 | 0.02 | 0.02 | 0.04 | 0 | 0 | 0.14 | 0 | 0 | 0 | - |
| Prrc2b | 9.06 | 8.72 | 9.04 | 9.84 | 9.94 | 10.41 | 9.02 | 8.6 | 9.41 | 11.47 | 9.62 | 10.86 | - |
| Gpc2 | 0.02 | 0.09 | 0.04 | 0.3 | 0.1 | 0.05 | 0.09 | 0.19 | 0.11 | 0.48 | 0.59 | 0.48 | - |
| Prkab2 | 0.99 | 1.16 | 0.8 | 1.26 | 1.8 | 1.2 | 1.96 | 1.78 | 1.73 | 1.81 | 2.15 | 2.05 | - |

| | | | | | | | | | | | | |
|---|---|---|---|---|---|---|---|---|---|---|---|---|---|
| Echdc2 | 10.16 | 9.75 | 9.02 | 3.39 | 4.2 | 7.16 | 5.72 | 6.83 | 6.61 | 4.68 | 2.88 | 5.4 | - |
| Vamp8 | 87.09 | 87.73 | 81.92 | 69.88 | 75.13 | 75.29 | 78.45 | 77.12 | 71.81 | 72.2 | 63.75 | 67.32 | - |
| Cep89 | 10.59 | 9.63 | 11.23 | 9.55 | 10.48 | 9.47 | 6.83 | 7.48 | 5.97 | 5.42 | 6 | 5.68 | - |
| Sgcb | 3.85 | 4.42 | 3.48 | 3.05 | 2.85 | 2.98 | 2.53 | 2.76 | 2.4 | 2.01 | 1.55 | 2.27 | - |
| Znhit1 | 23.06 | 22.83 | 22.09 | 14.67 | 19.03 | 16.12 | 16.2 | 16.18 | 17.65 | 13.31 | 13.28 | 15.72 | - |
| Sft2d1 | 64.06 | 61.74 | 63.1 | 40.53 | 49.2 | 48.62 | 49.08 | 52.91 | 52.01 | 46.02 | 35.99 | 39.55 | - |
| Rhno1 | 20.58 | 19.47 | 17.53 | 25.93 | 27.54 | 28.46 | 20.06 | 22.38 | 20.69 | 23.87 | 28.51 | 28.71 | - |
| Fam53b | 4.15 | 4.61 | 3.76 | 10.42 | 10.64 | 10.67 | 4.24 | 4.59 | 4.07 | 9.9 | 9.99 | 9.3 | - |
| Aarsd1 | 68.7 | 70.89 | 73.9 | 52.11 | 52.97 | 54.16 | 44.79 | 54.24 | 57.63 | 41.67 | 45.91 | 42.72 | - |
| Map3k9 | 0 | 0.02 | 0 | 0.08 | 0.04 | 0.05 | 0.07 | 0.02 | 0.01 | 0.18 | 0.08 | 0.03 | - |
| Rps6ka4 | 15.51 | 15.83 | 15.44 | 21.35 | 18.47 | 18.93 | 18.43 | 19.32 | 16.83 | 18.66 | 19.6 | 20.68 | - |
| Cir1 | 9.04 | 7.51 | 8.06 | 15.73 | 13.99 | 15.18 | 21.86 | 16.88 | 15.2 | 27.17 | 30.36 | 26.34 | - |
| 4930525D18Rik | 1.04 | 0.93 | 0.98 | 0.07 | 0.07 | 0.14 | 0.25 | 0.5 | 0.29 | 0 | 0.07 | 0 | - |
| Idnk | 13.64 | 15.1 | 16.38 | 20.9 | 16.56 | 17.55 | 22.53 | 19.08 | 18.01 | 25.86 | 22.56 | 25.61 | - |
| Arhgap9 | 27.39 | 39.55 | 37.38 | 59.33 | 55.65 | 51.97 | 71.15 | 63.79 | 57.21 | 77.93 | 63.4 | 68.82 | - |
| Gpr132 | 16.18 | 16.65 | 19.82 | 18.37 | 18.96 | 17.88 | 23.73 | 22.24 | 21.67 | 26.67 | 24.59 | 27.34 | - |
| Rnf38 | 3.21 | 3.21 | 2.91 | 5.67 | 4.7 | 4.77 | 5.12 | 3.75 | 3.74 | 6.92 | 5.81 | 6.02 | - |
| Nipa2 | 20.96 | 26.11 | 24.43 | 19.11 | 21.55 | 17.59 | 15.65 | 19.46 | 18.3 | 17.02 | 15.93 | 13.93 | - |
| Atp5g2 | 363.47 | 403.53 | 373.95 | 447.61 | 462.04 | 499.14 | 486.99 | 458.17 | 468.23 | 491.52 | 562.23 | 505.51 | - |
| Pik3ca | 2.15 | 2.17 | 2.29 | 2.48 | 2.45 | 2.49 | 3.07 | 3.33 | 3.14 | 3.07 | 3.6 | 3.33 | - |
| Srp54b | 46.87 | 42.45 | 43.67 | 32.35 | 34.09 | 33.03 | 25.17 | 34.13 | 30.51 | 25.73 | 24.75 | 25.77 | - |
| Gpt2 | 15.45 | 16.8 | 16.77 | 8.62 | 12.72 | 10.78 | 10.18 | 18.62 | 16.64 | 5.95 | 6.29 | 6.65 | - |
| Rapgef3 | 0.12 | 0.17 | 0.06 | 0.77 | 0.88 | 0.81 | 1.14 | 0.48 | 0.65 | 2.02 | 1.67 | 3.16 | - |
| Mpped2 | 0 | 0 | 0 | 0.25 | 0.17 | 0.15 | 0.02 | 0.02 | 0.02 | 0.41 | 0.29 | 0.14 | - |
| Themis2 | 0.81 | 0.92 | 0.7 | 1.52 | 0.74 | 1.13 | 4.25 | 2.1 | 2.66 | 2.93 | 1.5 | 3.81 | - |
| Tma16 | 30.33 | 30.06 | 33.64 | 19.52 | 22.72 | 21.8 | 15.47 | 32.22 | 25.49 | 15.83 | 17.39 | 15.79 | - |
| Cc2d1b | 18.47 | 22.43 | 21.8 | 15.39 | 15.62 | 13.33 | 17.54 | 16.6 | 17.91 | 13.36 | 11.52 | 14.81 | - |
| Fhl2 | 19.65 | 16.19 | 16.64 | 14.64 | 21.16 | 20.28 | 21.88 | 31.68 | 23.86 | 29.21 | 29.64 | 21.28 | - |
| Rpl34 | 68.26 | 62.38 | 64.17 | 42.61 | 49.23 | 48.88 | 49.87 | 49.33 | 56.28 | 37.29 | 49.71 | 41.85 | - |
| Gatm | 46.74 | 43.55 | 48.66 | 33.57 | 46.33 | 38.63 | 42.45 | 48 | 46.65 | 33.28 | 35.82 | 31.67 | - |
| Nlrc5 | 10.91 | 11.67 | 9.85 | 16.94 | 15.6 | 16.35 | 13.16 | 11.09 | 11.48 | 17.22 | 14.11 | 15.8 | - |
| Col24a1 | 0 | 0 | 0 | 0 | 0 | 0.01 | 0 | 0 | 0.02 | 0.03 | 0.02 | 0.11 | - |
| Dhx58 | 10.6 | 12.42 | 9.11 | 16.48 | 12.36 | 14.75 | 19.3 | 13.03 | 14.39 | 20.35 | 19.25 | 22.42 | - |
| BC029722 | 15.97 | 15.07 | 17.85 | 11.22 | 10 | 12.25 | 13.19 | 11.82 | 14.05 | 9.23 | 10.83 | 10.74 | - |
| Fabp5 | 174.34 | 173.91 | 168.24 | 82.94 | 87.02 | 86.64 | 69.05 | 80.28 | 87.29 | 40.29 | 44.68 | 46.75 | - |
| Card11 | 6.46 | 7.34 | 6.23 | 11.3 | 9.08 | 9.68 | 14.39 | 9.37 | 10.57 | 11.32 | 9.2 | 14.86 | - |
| Fam83d | 11.52 | 7.13 | 8.61 | 8.87 | 13.73 | 10.95 | 4.25 | 4.76 | 3.25 | 4.56 | 5.84 | 3.29 | - |
| Plk4 | 19.7 | 18.68 | 16.53 | 11.51 | 16.05 | 14.49 | 10.67 | 11.82 | 12.57 | 9.32 | 10.1 | 8.86 | - |
| Id2 | 138.95 | 152.03 | 107.55 | 231.59 | 233.09 | 223.64 | 109.24 | 93.41 | 105.98 | 181.12 | 153.62 | 194.21 | - |
| Fut4 | 4.32 | 7.07 | 4.66 | 5.71 | 4.11 | 4.32 | 2.38 | 2.81 | 2.86 | 3.42 | 2.81 | 3.19 | - |
| Aven | 3.21 | 4.54 | 3.34 | 2.81 | 2.49 | 3.09 | 2.45 | 3.04 | 2.92 | 1.45 | 1.74 | 1.47 | - |
| Soat1 | 12.83 | 14.13 | 14.11 | 17.14 | 16.63 | 19.03 | 18.68 | 13.65 | 16.39 | 18.26 | 15.96 | 17.1 | - |
| Kctd12 | 0.13 | 0.16 | 0.07 | 0.32 | 0.2 | 0.23 | 0.35 | 0.32 | 0.26 | 0.94 | 0.49 | 0.99 | - |
| Sp1 | 3.24 | 3.71 | 3.45 | 3.61 | 4.26 | 4.44 | 4.24 | 4 | 3.79 | 4.92 | 5.16 | 4.83 | - |
| Rad18 | 22.17 | 20.29 | 20.27 | 15.53 | 19.24 | 18.84 | 16.45 | 17.76 | 16.86 | 11.97 | 13.39 | 9.66 | - |
| Wdr73 | 16.27 | 13.77 | 14.11 | 8.39 | 10.19 | 9.91 | 6.84 | 8.95 | 9.21 | 7.49 | 6.01 | 7.54 | - |
| Surf1 | 33.83 | 33.31 | 35.93 | 59 | 56.78 | 55.96 | 64.89 | 50.22 | 55.89 | 73.33 | 73.4 | 82.98 | - |
| Syngr2 | 138.08 | 134.13 | 141.34 | 102.23 | 109.08 | 105.23 | 82.24 | 117.93 | 96.73 | 87.39 | 83.19 | 79.54 | - |
| 1600002H07Rik | 6.65 | 6.88 | 7.39 | 5.71 | 5.98 | 8.13 | 4.85 | 4.93 | 5.09 | 3.25 | 4.82 | 4.43 | - |
| Pdk3 | 65.49 | 66.93 | 72.7 | 46.93 | 52.04 | 46.7 | 58.67 | 55.3 | 57.51 | 42.62 | 45.51 | 43.09 | - |
| Palm | 1.22 | 1.01 | 1.61 | 1.66 | 1.45 | 1.95 | 2.8 | 1.73 | 3.41 | 2.83 | 1.96 | 2.83 | - |
| Acpp | 0.41 | 0.21 | 0.11 | 0.5 | 0.26 | 0.12 | 0.72 | 0.33 | 0.83 | 1.29 | 1.24 | 1.37 | - |
| Rdh9 | 0.13 | 0.19 | 0.14 | 0.79 | 0.58 | 0.7 | 0.85 | 0.53 | 0.36 | 1.5 | 1.27 | 1.3 | - |
| Fam195a | 13.57 | 18.47 | 20.54 | 13.63 | 13.69 | 14.3 | 14.51 | 15.91 | 18.28 | 9.81 | 10.2 | 8.12 | - |
| a | 0.91 | 1.01 | 0.86 | 3.29 | 2.44 | 3.13 | 1.49 | 0.34 | 0.47 | 3.06 | 2.67 | 4.41 | - |
| Stbd1 | 0.12 | 0.06 | 0.68 | 0.54 | 0.44 | 0.29 | 1.97 | 1.36 | 1.16 | 3.4 | 3.54 | 2.41 | - |
| Grhl1 | 0.63 | 0.64 | 0.81 | 0.26 | 0.56 | 0.4 | 0.18 | 0.58 | 0.45 | 0.13 | 0.25 | 0.15 | - |
| Emc1 | 10.49 | 10.35 | 11.75 | 8.55 | 10.86 | 8.37 | 5.99 | 6.92 | 7.08 | 7.5 | 6.41 | 6.31 | - |
| Zmiz1 | 6.56 | 6.84 | 6.16 | 8.49 | 7.88 | 7.43 | 9.75 | 7.96 | 8.83 | 9.02 | 8.22 | 9.1 | - |
| Trim8 | 8.79 | 9.75 | 9.51 | 11.66 | 10.15 | 10.92 | 11.83 | 10.64 | 11.04 | 12.44 | 11.77 | 14.69 | - |
| Ptpn5 | 8.76 | 6.59 | 6.64 | 2.4 | 1.28 | 1.79 | 5.26 | 5.61 | 3.9 | 1.74 | 1.26 | 1.47 | - |
| Cxcr4 | 0.19 | 0.55 | 0.24 | 0.77 | 0.83 | 0.98 | 1.14 | 1.17 | 0.92 | 2.11 | 1.12 | 1.78 | - |
| Ska2 | 23.45 | 20.1 | 19.26 | 15.84 | 18.65 | 15.55 | 13.29 | 12 | 11.45 | 11.83 | 11.51 | 9.79 | - |
| Tmem50a | 139.4 | 125.01 | 131.2 | 186 | 165.83 | 187.95 | 207.47 | 179.08 | 189.2 | 225.53 | 232.97 | 253.99 | - |
| Gm5177 | 34.49 | 34.26 | 33.76 | 34.98 | 31.17 | 31.22 | 58.27 | 39.71 | 46.42 | 38.93 | 44.25 | 45.79 | - |
| Tmem2 | 1.47 | 1.42 | 1.64 | 2.55 | 2.54 | 2.91 | 0.55 | 0.93 | 0.61 | 0.72 | 0.82 | 1.19 | - |
| Apitd1 | 9.82 | 8.57 | 9.88 | 4.21 | 6.18 | 4.73 | 2.35 | 3.16 | 4.27 | 2.09 | 1.59 | 1.3 | - |
| 5930412G12Rik | 0.05 | 0 | 0 | 0 | 0.23 | 0.37 | 0.17 | 0.02 | 0.02 | 0.81 | 0.18 | 0.62 | - |
| Nr3c1 | 2.36 | 2.57 | 2.5 | 5.06 | 4.93 | 5.15 | 2.94 | 2.95 | 2.8 | 5.33 | 4.53 | 4.76 | - |
| Rnf121 | 18.22 | 15.52 | 18.6 | 14.31 | 14.76 | 13.73 | 15.4 | 15.47 | 14.06 | 10.48 | 10.81 | 11.54 | - |
| Cenpi | 13.95 | 12.47 | 12.66 | 7.45 | 8.77 | 8.14 | 4.44 | 5.69 | 4.61 | 2.98 | 5.19 | 3.5 | - |
| Fam120a | 30.97 | 29.97 | 32.46 | 25.37 | 24.87 | 26.9 | 23.12 | 24.61 | 26.1 | 23.71 | 21.78 | 22.55 | - |
| Tmem176b | 1.36 | 2.99 | 1.71 | 14.25 | 7.06 | 5.45 | 10.45 | 2.14 | 6.54 | 9.64 | 5.12 | 9.85 | - |
| Arid3b | 6.54 | 5.41 | 5.98 | 8.36 | 6.97 | 6.02 | 6.68 | 6.33 | 7.1 | 8.23 | 8.33 | 8.23 | - |
| Col6a2 | 0.15 | 0.14 | 0.12 | 1.44 | 0.89 | 1.03 | 0.31 | 0.17 | 0.47 | 0.02 | 0 | 0 | - |
| Anapc2 | 33.86 | 38.01 | 38.87 | 51.51 | 47.83 | 50.37 | 45.56 | 45.22 | 45.81 | 55.63 | 54.08 | 49.89 | - |
| Rpusd2 | 3.15 | 3.5 | 3.96 | 2.21 | 3.03 | 2.87 | 2.22 | 2.31 | 2.91 | 2.41 | 2.33 | 2.22 | - |
| Ccndbp1 | 28.48 | 28.84 | 26.69 | 36.95 | 34.23 | 33.92 | 44.96 | 39.08 | 39.27 | 42.66 | 41.13 | 41.42 | - |
| Ubn1 | 3.45 | 3.85 | 3.14 | 4.84 | 4.41 | 4.88 | 5.2 | 4.37 | 4.21 | 5.18 | 5.26 | 5.08 | - |
| Gzme | 10.8 | 15.93 | 3.62 | 0 | 0 | 0 | 0.65 | 0.06 | 0.23 | 0 | 0 | 0 | - |
| Trpm7 | 7.36 | 7.45 | 7.12 | 9.11 | 9.72 | 9.38 | 8.8 | 8.15 | 9.66 | 11.22 | 11.76 | 10.56 | - |
| Gapdh | 5444.19 | 5665.69 | 5739.39 | 5565.36 | 4939.48 | 5164.16 | 9530.92 | 6317.53 | 7662.55 | 6657.56 | 7155.64 | 7278.22 | - |
| Dstyk | 0.51 | 0.47 | 0.51 | 0.64 | 0.92 | 0.69 | 0.6 | 0.7 | 0.59 | 0.89 | 0.69 | 1.25 | - |
| Cwf19l2 | 6.14 | 5.03 | 5.92 | 8.23 | 8.37 | 8.21 | 6.88 | 6.6 | 5.81 | 7.92 | 10.02 | 7.55 | - |
| Pms1 | 9.48 | 10.1 | 8.97 | 7.06 | 5.83 | 6.71 | 5.02 | 5.54 | 6.41 | 5.27 | 4.94 | 3.99 | - |
| Snai3 | 12.45 | 11.24 | 12.56 | 7.4 | 7.46 | 5.86 | 10.2 | 10.12 | 9.05 | 6.34 | 5.62 | 6.44 | - |
| Mrp63 | 27.77 | 28.05 | 27.26 | 19.8 | 22.47 | 21.82 | 22.36 | 29.48 | 24.8 | 18.29 | 18.22 | 17.82 | - |
| Mkrn1 | 4.14 | 5.07 | 5.19 | 6.52 | 5.29 | 6.04 | 8.36 | 6.94 | 8.52 | 8.22 | 8.14 | 8.34 | - |
| Cenpb | 8.51 | 9.27 | 8.18 | 4.97 | 6.78 | 4.85 | 5.1 | 5.11 | 5 | 3.68 | 3.69 | 4.61 | - |
| Pkn2 | 1.81 | 1.76 | 1.89 | 1.91 | 2.21 | 2.06 | 1.98 | 2.04 | 2.44 | 2.49 | 3.21 | 2.34 | - |
| Ptpn13 | 0.38 | 0.46 | 0.65 | 1.2 | 0.92 | 0.82 | 1.36 | 0.9 | 1.19 | 1.51 | 1.09 | 0.94 | - |
| Cyp46a1 | 0 | 0.03 | 0.03 | 0.15 | 0.29 | 0.3 | 0.33 | 0.11 | 0.23 | 0.49 | 0.09 | 0.33 | - |
| Tiparp | 3.32 | 4.36 | 3.76 | 12.2 | 13.25 | 13.29 | 5.85 | 5.42 | 5.11 | 13.43 | 14.94 | 15.32 | - |
| Ccnf | 26.77 | 17.14 | 16.57 | 14.81 | 20.69 | 17.04 | 9.14 | 8.03 | 9.8 | 8.12 | 10.42 | 8.2 | - |
| Zfp217 | 2.58 | 2.34 | 2.61 | 3.54 | 3.34 | 3.26 | 3.41 | 2.91 | 2.77 | 3.2 | 3.71 | 3.59 | - |
| Fig4 | 19.28 | 17.42 | 16.69 | 12.1 | 11.4 | 12.1 | 9.25 | 13.93 | 11.2 | 9.62 | 8.16 | 9.1 | - |
| Gm5801 | 31.85 | 30.78 | 28.8 | 34.58 | 30 | 34.84 | 44.31 | 39.06 | 40.64 | 39.18 | 47.33 | 43.52 | - |
| Itga5 | 3.37 | 2.99 | 3.44 | 2.17 | 2.04 | 2.38 | 2.16 | 2.84 | 2.9 | 1.74 | 1.31 | 1.44 | - |
| Csf2 | 1.66 | 0.56 | 0.63 | 0 | 0 | 0 | 0 | 0 | 0 | 0 | 0 | 0 | - |
| Arhgap12 | 5.57 | 5.23 | 4.98 | 8.5 | 7.53 | 7.4 | 6.94 | 7.06 | 7.09 | 7.71 | 8.91 | 9.04 | - |
| Afg3l2 | 46.26 | 56.34 | 59.62 | 80.03 | 72.55 | 75.51 | 123.44 | 92.82 | 99.94 | 100.98 | 114.37 | 115.99 | - |
| Slc22a15 | 0.24 | 0.45 | 0.19 | 0.53 | 0.39 | 0.29 | 0.67 | 0.19 | 0.47 | 0.79 | 0.6 | 0.75 | - |
| Sec23b | 68.24 | 61.93 | 59.46 | 38.78 | 43.68 | 42.3 | 34.6 | 43.92 | 39.36 | 27.09 | 27.86 | 28.42 | - |
| Dus2l | 23.73 | 21.55 | 23.28 | 17.94 | 18.39 | 20.33 | 17.39 | 21.6 | 19.56 | 13.01 | 15.21 | 17.35 | - |
| Ssc5d | 0.01 | 0 | 0.01 | 0.04 | 0.1 | 0.07 | 0.11 | 0.12 | 0.11 | 0.17 | 0.17 | 0.23 | - |
| Dntt | 0.06 | 0.34 | 0.24 | 0.4 | 0.33 | 0.43 | 3.33 | 0.57 | 3.27 | 2.1 | 2.11 | 2.53 | - |
| Sema6d | 0.28 | 0.38 | 0.39 | 0 | 0.01 | 0 | 0.01 | 0 | 0 | 0 | 0.01 | 0 | - |
| Pus7 | 27.35 | 29.44 | 27.42 | 16.48 | 18.58 | 18.42 | 13.67 | 19.1 | 17.54 | 13.91 | 11.43 | 12.12 | - |
| Smyd5 | 60.48 | 57.71 | 62.6 | 20.51 | 25.16 | 29.26 | 24.57 | 41.72 | 37.16 | 17.42 | 19.55 | 15.75 | - |
| Rassf7 | 10.47 | 9.65 | 11.63 | 8.99 | 9.91 | 8.95 | 6.82 | 10.17 | 8.65 | 6.3 | 8.2 | 6.31 | - |
| Tmem189 | 4.02 | 4.37 | 3.37 | 4.05 | 3.78 | 4.65 | 5 | 4.46 | 6.47 | 6.49 | 6.89 | 5.73 | - |
| Zzef1 | 2.29 | 2.31 | 1.77 | 2.91 | 2.65 | 2.79 | 2.74 | 2.54 | 2.82 | 3.16 | 3.33 | 3.22 | - |
| Atad3a | 25.58 | 33.32 | 39.9 | 18 | 19.87 | 19.3 | 12.64 | 24.23 | 18.99 | 11.8 | 11.67 | 10.63 | - |
| Rab40c | 1.55 | 1.46 | 1.69 | 2.49 | 1.95 | 2.28 | 2.12 | 2.33 | 2 | 3.2 | 2.45 | 2.57 | - |
| Hpcal1 | 19.78 | 19.31 | 19.57 | 40.44 | 33.51 | 36.7 | 69.46 | 37.51 | 46.95 | 63.06 | 60.03 | 67.01 | - |
| Mon2 | 1.48 | 1.27 | 1.6 | 2.3 | 2.05 | 2.58 | 2.11 | 1.98 | 2.15 | 2.69 | 3.08 | 2.28 | - |
| C2cd5 | 9.78 | 9.79 | 9.52 | 7.42 | 6.32 | 8 | 6.73 | 6.34 | 6.47 | 4.16 | 5.48 | 4.71 | - |
| Tex2 | 12.14 | 12.79 | 11.77 | 16.89 | 14.89 | 14.84 | 7.49 | 7.82 | 7.51 | 9.34 | 6.78 | 8.57 | - |
| Ube3b | 5.95 | 7.39 | 6.23 | 8.19 | 8.44 | 8.65 | 9.59 | 7.49 | 7.54 | 9.22 | 8.76 | 9.01 | - |
| Zfp51 | 13.81 | 12.77 | 14.52 | 18.01 | 16.44 | 16.97 | 14.45 | 16.69 | 13.51 | 18.98 | 18.29 | 18.24 | - |
| Abcb1a | 1.13 | 1.61 | 1.78 | 1.64 | 2.22 | 1.26 | 0.7 | 0.75 | 0.93 | 0.61 | 0.22 | 0.91 | - |
| Klrc3 | 0.18 | 0.87 | 0.83 | 2.19 | 1.68 | 2.45 | 1.72 | 1.54 | 1.81 | 2.98 | 3.93 | 2.42 | - |
| Tshz1 | 1.34 | 1.55 | 0.83 | 2.46 | 2.22 | 1.68 | 1.98 | 1.42 | 1.51 | 2.13 | 1.79 | 2.08 | - |
| Fubp3 | 10.44 | 12.03 | 12.22 | 10.95 | 9.39 | 9.31 | 8.58 | 9.88 | 10.07 | 9.07 | 7.71 | 7.89 | - |
| Thtpa | 2.73 | 3.25 | 2.98 | 4.58 | 3.69 | 4.19 | 4.44 | 4.18 | 3.38 | 4.67 | 4.15 | 5.36 | - |
| Tnip1 | 12.6 | 12.66 | 12.66 | 16.64 | 17.13 | 15.01 | 20.11 | 19.44 | 15.47 | 21.04 | 23.11 | 20.58 | - |
| Slc2a8 | 6.06 | 5.67 | 5.71 | 11.38 | 8.42 | 9.15 | 16.64 | 11.08 | 11.87 | 17.93 | 13.37 | 19.53 | - |
| 2810013P06Rik | 1.25 | 2.22 | 2.92 | 2.8 | 3.17 | 1.98 | 3.88 | 2.1 | 2.92 | 5.71 | 4.6 | 5.45 | - |
| Dctn1 | 17.08 | 17.6 | 15.74 | 15.04 | 16.28 | 13.82 | 14.2 | 12.55 | 13.02 | 12.46 | 12.63 | 12.62 | - |
| Paip2b | 4.96 | 4.53 | 3.86 | 3.91 | 5.48 | 4.09 | 3.6 | 3.06 | 2.65 | 2.18 | 3.86 | 2.03 | - |
| Dusp2 | 3.92 | 4.96 | 6.85 | 19.8 | 24.46 | 16.67 | 10.02 | 16.77 | 19.47 | 45.22 | 24.3 | 19.45 | - |
| Arhgap17 | 5.9 | 5.97 | 5.85 | 9.77 | 7.8 | 8.81 | 7.39 | 6.64 | 6.74 | 8.36 | 6.89 | 9.4 | - |
| Fbxw8 | 4.16 | 3.8 | 4.13 | 3.89 | 3.91 | 3.8 | 2.89 | 3.55 | 3.24 | 3.34 | 2.56 | 2.66 | - |
| Uqcrq | 321.39 | 334.69 | 333.71 | 197.98 | 229.29 | 212.01 | 239.39 | 344.87 | 304.51 | 150.54 | 185.88 | 148.46 | - |
| Tpst2 | 33.16 | 35.7 | 31.74 | 62.65 | 44.08 | 42.97 | 63.39 | 35.52 | 45.52 | 106.2 | 80.63 | 107.58 | - |
| Lyn | 1.42 | 1.57 | 1.52 | 2.78 | 1.61 | 1.14 | 2.78 | 3.5 | 2.53 | 4.72 | 3.78 | 4.79 | - |
| 2810025M15Rik | 6.36 | 5.95 | 5.86 | 4.98 | 5.54 | 5.71 | 13.91 | 12.4 | 11.82 | 9.68 | 12.28 | 8.88 | - |
| Cul2 | 49.99 | 44.38 | 46.7 | 33.24 | 35.87 | 34.44 | 27.48 | 41.77 | 36.22 | 27.27 | 26.29 | 26.41 | - |
| Mgat5 | 3.09 | 3.26 | 3.3 | 1.97 | 2.19 | 1.4 | 1.39 | 2.59 | 2.39 | 0.9 | 1.45 | 0.65 | - |
| Galnt14 | 0.3 | 0.31 | 0.33 | 0 | 0 | 0.05 | 0.05 | 0.17 | 0.13 | 0 | 0 | 0 | - |
| Coq3 | 10.07 | 10.92 | 11.48 | 9.07 | 8.77 | 8.71 | 8.85 | 10.54 | 8.77 | 7.38 | 7.07 | 7.81 | - |
| Kdm4c | 7.34 | 8.2 | 7.93 | 8.82 | 9.74 | 9.87 | 9.22 | 8.97 | 10.47 | 9.74 | 13.06 | 9.69 | - |
| Acsl6 | 5.63 | 6.24 | 7.14 | 1.09 | 2.3 | 1.56 | 1.03 | 1.69 | 1.38 | 0.36 | 0.36 | 0.27 | - |
| Arpc2 | 301.61 | 342.23 | 322.45 | 363.9 | 341.4 | 354.54 | 496.2 | 404.09 | 451.45 | 440.57 | 442.39 | 453.44 | - |
| Rgcc | 72.33 | 67.99 | 83.88 | 40.66 | 48.9 | 50.45 | 68.41 | 72.81 | 75.37 | 29.42 | 41.87 | 43.77 | - |
| Bloc1s1 | 29.69 | 31 | 28.71 | 41.94 | 34.62 | 42.67 | 57.05 | 35.88 | 45.55 | 47.32 | 43.81 | 52.51 | - |
| Gbp3 | 12.82 | 14.75 | 13.96 | 19.37 | 19.79 | 16.51 | 23.96 | 20.96 | 22.69 | 29.05 | 28.68 | 29.23 | - |
| Srms | 0.02 | 0 | 0 | 0.17 | 0.17 | 0 | 0.2 | 0.09 | 0.15 | 0.22 | 0.34 | 0.43 | - |
| Mettl7a1 | 0.61 | 1.42 | 0.61 | 2.55 | 2.06 | 1.86 | 2.08 | 1.28 | 2.08 | 2.1 | 2.81 | 2.57 | - |
| Dcaf15 | 6.91 | 7.63 | 6.24 | 12.28 | 11.85 | 12.05 | 16.99 | 12.27 | 13.58 | 18.2 | 21.74 | 18.43 | - |
| Aff3 | 0.83 | 0.61 | 0.89 | 1.31 | 0.84 | 1.11 | 0.4 | 0.4 | 0.61 | 0.23 | 0.05 | 0.54 | - |

| | | | | | | | | | | | | |
|---|---|---|---|---|---|---|---|---|---|---|---|---|
| Fam20b | 5.74 | 5.53 | 6.01 | 3.46 | 4.71 | 4.23 | 3.23 | 4.7 | 3.87 | 3.71 | 3.33 | 2.8 | - |
| Plch1 | 0.28 | 0.07 | 0.09 | 0.01 | 0.08 | 0.05 | 0.03 | 0.02 | 0.03 | 0.01 | 0 | 0 | - |
| Gm16973 | 1.43 | 1.42 | 1.34 | 2.89 | 2.78 | 3.04 | 3.11 | 2.36 | 2.41 | 4.75 | 6.2 | 5.32 | - |
| Fam46c | 1.15 | 1.12 | 1.07 | 1.96 | 3.35 | 3.24 | 5.41 | 2.69 | 3.68 | 8.09 | 7.41 | 6.68 | - |
| Pabpc4 | 22.91 | 20.11 | 21.42 | 14.9 | 15.09 | 16.82 | 11.31 | 16.65 | 15.26 | 10.02 | 10.64 | 10.13 | - |
| Styx | 4.94 | 5.74 | 8.07 | 5.39 | 4.94 | 5.19 | 3.63 | 4.78 | 5.44 | 4.45 | 4.07 | 3.94 | - |
| Zfp781 | 2.45 | 2.06 | 2.56 | 2.98 | 3.76 | 2.76 | 2.17 | 2.56 | 2.32 | 4.84 | 3.2 | 2.82 | - |
| 1810043G02Rik | 4.81 | 3.98 | 4.21 | 6.86 | 6.76 | 5.23 | 7.32 | 5.96 | 5.73 | 6.99 | 8.54 | 8.12 | - |
| Ugt1a9 | 2.88 | 1.88 | 1.83 | 0.13 | 0.24 | 0.11 | 0.3 | 0.83 | 0.35 | 0 | 0 | 0 | - |
| Galnt2 | 4.22 | 4.76 | 4.87 | 5.34 | 5.21 | 4.42 | 5.55 | 5.26 | 6.14 | 7.39 | 5.2 | 7.77 | - |
| Aldh1l1 | 0.43 | 0.28 | 0.07 | 0.39 | 0.65 | 0.14 | 0.65 | 0.38 | 0.77 | 0.96 | 1.14 | 0.95 | - |
| Ssh3 | 3.64 | 3.95 | 2.83 | 8.29 | 6.41 | 7.08 | 11.41 | 6.22 | 8.19 | 10.4 | 10.39 | 11.66 | - |
| Lyrm7 | 3.3 | 2.86 | 2.41 | 1.86 | 1.4 | 1.9 | 1.42 | 1.67 | 1.75 | 0.87 | 0.92 | 1.39 | - |
| Wdr81 | 1.64 | 1.31 | 2.39 | 2.26 | 2.1 | 2.54 | 3.14 | 3.01 | 2.87 | 3.72 | 4.25 | 4.03 | - |
| Ldb1 | 7.28 | 6.96 | 7 | 9.98 | 8.14 | 9.7 | 9.98 | 10.66 | 9.28 | 12.52 | 10.48 | 12.26 | - |
| Cd200 | 133.88 | 110.36 | 135.25 | 28.48 | 37.64 | 36.4 | 44.1 | 92.39 | 82.44 | 23.01 | 25.17 | 22.36 | - |
| Tmem206 | 16.31 | 16.43 | 16.4 | 12.64 | 14.1 | 12.06 | 10.25 | 15.11 | 12.49 | 9.71 | 12.55 | 11.97 | - |
| Aldh7a1 | 19.71 | 18.19 | 18.33 | 12.91 | 14.39 | 15.42 | 15.96 | 18.85 | 17.7 | 9.87 | 10.96 | 13 | - |
| Casz1 | 0.04 | 0.11 | 0.14 | 0.47 | 0.35 | 0.26 | 0.82 | 0.48 | 0.4 | 0.88 | 1.1 | 1.04 | - |
| Kansl3 | 8.29 | 8.66 | 8.55 | 12.8 | 12.5 | 12.23 | 9.09 | 11.44 | 10.43 | 13.69 | 15.05 | 12.65 | - |
| Mcmdc2 | 1.52 | 1.67 | 1.88 | 2.47 | 2.72 | 2.05 | 2.68 | 2.5 | 1.85 | 2.86 | 2.79 | 2.87 | - |
| Gnpnat1 | 49.69 | 49.82 | 53.22 | 32.12 | 35.7 | 34.52 | 33.57 | 53.51 | 47.21 | 24.08 | 25.95 | 24.99 | - |
| Xpo1 | 21.8 | 22.24 | 25.8 | 13.58 | 16 | 15.59 | 13.16 | 16.59 | 15.19 | 11.64 | 11.99 | 10.98 | - |
| Ist1 | 35.59 | 34.43 | 33.43 | 47.82 | 44.86 | 49.18 | 40.25 | 38.05 | 35.67 | 46.48 | 39.39 | 46.35 | - |
| Wdr35 | 2.99 | 2.91 | 3.44 | 2.26 | 2.09 | 2.25 | 1.83 | 2.12 | 2.4 | 1.61 | 1.03 | 1.88 | - |
| Limd2 | 67.21 | 85.91 | 77.46 | 133.92 | 106.14 | 107.29 | 134.98 | 102.9 | 109.53 | 154.1 | 124.32 | 161.57 | - |
| Epm2aip1 | 2.67 | 2.6 | 3.31 | 4.1 | 3.56 | 3.67 | 3.79 | 3.52 | 3.41 | 4.38 | 3.97 | 3.69 | - |
| Pdpr | 1.12 | 1.56 | 1.28 | 1.22 | 1.72 | 1.91 | 2.3 | 1.68 | 1.92 | 2.17 | 2.95 | 2.24 | - |
| Abcg2 | 8.18 | 7.97 | 9.46 | 5.76 | 3.73 | 3.8 | 6.64 | 7.59 | 5.82 | 3.74 | 3.79 | 5.55 | - |
| Slc25a11 | 66.48 | 66.19 | 60.14 | 55.44 | 49.53 | 51.79 | 54.51 | 53.02 | 50.77 | 48.35 | 41.57 | 46.43 | - |
| Dapl1 | 6.27 | 4.56 | 6.88 | 3.41 | 2.61 | 2.78 | 0.2 | 0.61 | 0.47 | 0.1 | 0 | 0.4 | - |
| Aaed1 | 7.75 | 7.12 | 9.02 | 16.58 | 16.24 | 12.93 | 23.76 | 14.32 | 15.68 | 19.07 | 21.45 | 21.59 | - |
| Ptpn22 | 75.76 | 80.9 | 79.37 | 156.13 | 147.62 | 155.34 | 154.18 | 117.48 | 126.79 | 219.63 | 203.37 | 238.38 | - |
| Plac8 | 423.44 | 470.53 | 388.07 | 474.56 | 407.18 | 419.06 | 706.02 | 397.28 | 513.73 | 643.76 | 544.92 | 710.06 | - |
| Mrpl35 | 27.41 | 28.84 | 27.78 | 18.69 | 20.19 | 20.81 | 20.74 | 25.63 | 25.01 | 16.43 | 16.22 | 15.1 | - |
| Mrpl11 | 14.02 | 13.75 | 14.71 | 7.54 | 9.71 | 9.09 | 7.28 | 9.42 | 8.22 | 5.66 | 6.11 | 6.43 | - |
| Nov | 0.77 | 1.03 | 0.48 | 1.76 | 0.42 | 0.89 | 0.34 | 0.7 | 0.43 | 0.1 | 0.05 | 0 | - |
| Fus | 12.14 | 21.74 | 25.18 | 13.81 | 15.61 | 11.13 | 9.37 | 12.58 | 12.22 | 10.3 | 6.18 | 8.38 | - |
| Epn1 | 0.93 | 1.08 | 2.06 | 1.92 | 1.49 | 2.84 | 2 | 2.33 | 1.85 | 3.15 | 2.75 | 2.22 | - |
| Ly6c1 | 1.76 | 4.71 | 3.36 | 8.17 | 6.32 | 8.13 | 3.71 | 1.23 | 3.5 | 10.9 | 9.42 | 10.23 | - |
| Aqp9 | 3.37 | 5.45 | 2.66 | 1.03 | 0.99 | 0.77 | 10.07 | 11.19 | 13.35 | 2.46 | 1.14 | 2.18 | - |
| Cela1 | 3.66 | 3.26 | 4.35 | 6.69 | 8.47 | 8.36 | 5.09 | 5.1 | 5.34 | 7.29 | 13.1 | 8.86 | - |
| Wdr19 | 0.66 | 0.49 | 0.72 | 1.28 | 0.79 | 0.74 | 1.49 | 1.13 | 1.29 | 1.86 | 1.36 | 1.15 | - |
| Rbm5 | 30.75 | 28.38 | 26.96 | 30.71 | 31.99 | 32.74 | 32.2 | 29.62 | 30.76 | 38.16 | 30.9 | 37.63 | - |
| Tbcb | 64.29 | 62.85 | 58.53 | 54.88 | 55.07 | 53.82 | 51.19 | 50.93 | 51.27 | 44.89 | 46.81 | 42.33 | - |
| Mrps28 | 51.87 | 53.76 | 53.69 | 27.31 | 27.93 | 30.45 | 36.45 | 41.24 | 44.4 | 24.09 | 25.3 | 23.85 | - |
| Smpd1 | 22.69 | 22.24 | 20.51 | 39.88 | 32.09 | 35.96 | 54.51 | 35.53 | 40.27 | 45.87 | 52.6 | 65.21 | - |
| Klri2 | 12.13 | 8.76 | 11.73 | 4.77 | 5.03 | 4.56 | 5.99 | 8.84 | 9.01 | 3.67 | 3.1 | 3.31 | - |
| Tjp2 | 1.61 | 1.8 | 1.6 | 0.83 | 1.05 | 1.12 | 0.67 | 0.37 | 0.82 | 0.74 | 0.51 | 0.21 | - |
| Gm17821 | 11.28 | 12.89 | 13.38 | 22.05 | 19.86 | 19.59 | 19.05 | 18.08 | 16.72 | 25.08 | 23.31 | 22.1 | - |
| Eif4b | 187.09 | 189.54 | 192.48 | 171.04 | 172.78 | 168.46 | 167.51 | 179.64 | 178.56 | 158.75 | 175.27 | 157.32 | - |
| Shisa5 | 383.8 | 412.72 | 352.86 | 387.91 | 371.54 | 358.71 | 487.4 | 448.32 | 448.13 | 467.85 | 408.27 | 506.43 | - |
| Abcc5 | 5.5 | 5.49 | 5.56 | 3.36 | 3.85 | 3.86 | 3.47 | 5.01 | 4.79 | 3.12 | 4.8 | 3.48 | - |
| Wdr59 | 4.24 | 4.5 | 4.76 | 4.36 | 4.66 | 4.39 | 3.52 | 3.17 | 3.47 | 2.54 | 3.22 | 3.79 | - |
| Tox | 2.16 | 2.69 | 2.12 | 3.79 | 2.85 | 2.75 | 0.95 | 1.32 | 0.85 | 0.72 | 0.97 | 1.44 | - |
| Mll3 | 1.6 | 1.48 | 1.63 | 2.77 | 2.76 | 2.56 | 2.79 | 2.23 | 2.7 | 3.69 | 3.51 | 3.9 | - |
| Bcl11b | 1.84 | 2.3 | 2.98 | 4.01 | 3.46 | 3.35 | 3.83 | 3.28 | 3.95 | 3.79 | 3.65 | 3.66 | - |
| Ado | 15.83 | 17.46 | 16.12 | 10.72 | 11.96 | 12.37 | 9.48 | 14.42 | 13.85 | 9.61 | 10.25 | 8.97 | - |
| Cad | 25.77 | 24.52 | 28.38 | 11.68 | 13.12 | 12.64 | 9.19 | 13.26 | 13.67 | 7.57 | 7.64 | 6.53 | - |
| Suv39h2 | 1.19 | 0.66 | 1.19 | 0.49 | 0.56 | 0.59 | 0.29 | 0.35 | 0.6 | 0.55 | 0.21 | 0.57 | - |
| Mlkl | 19.35 | 16.96 | 17.04 | 11.85 | 11.04 | 14.46 | 10.87 | 12.35 | 11.82 | 10.99 | 8.55 | 10.97 | - |
| Dnajc17 | 8.63 | 10.33 | 10.24 | 6.21 | 6.69 | 6.58 | 6.73 | 7.02 | 6.39 | 5.28 | 3.73 | 5.72 | - |
| Lsp1 | 45.79 | 56.56 | 35.67 | 127.11 | 98.28 | 104.87 | 118.04 | 45.54 | 61.33 | 126.64 | 87.87 | 152.29 | - |
| Slc9a9 | 0.95 | 0.97 | 1.35 | 1.85 | 1.41 | 1.84 | 3.46 | 2.07 | 2.44 | 3.69 | 2.01 | 3.33 | - |
| E230016M11Rik | 0 | 0 | 0 | 0.14 | 0 | 0.08 | 0.07 | 0.1 | 0.3 | 0.25 | 0.15 | 0.28 | - |
| Pomt2 | 2.53 | 2.4 | 2.19 | 1.36 | 1.62 | 1.84 | 1.65 | 1.9 | 1.7 | 1.28 | 0.86 | 1.33 | - |
| Pja1 | 18.63 | 18.13 | 18.66 | 25.65 | 21.13 | 21 | 21.72 | 21.2 | 20.21 | 26.29 | 19.9 | 26.39 | - |
| Fam174b | 3.11 | 4.89 | 2.73 | 1.86 | 1.68 | 2.26 | 2.36 | 2.54 | 2.2 | 1.3 | 1.24 | 1.61 | - |
| Senp7 | 4.07 | 4.76 | 5.04 | 8.98 | 6.24 | 7.66 | 7.69 | 6.85 | 7.6 | 7.6 | 10.2 | 8.49 | - |
| 1110032F04Rik | 1.9 | 2.31 | 2.33 | 0.86 | 1.09 | 0.84 | 0.85 | 0.45 | 1.1 | 0.53 | 0.33 | 0.4 | - |
| Dph1 | 14.84 | 12.99 | 12.46 | 11.05 | 9.55 | 10.86 | 11.33 | 12.73 | 11.2 | 7.82 | 7.73 | 7.33 | - |
| Arid4a | 1.91 | 2.7 | 1.94 | 1.91 | 2.01 | 2.95 | 3.02 | 2.48 | 2.81 | 3.99 | 3.76 | 3.44 | - |
| Tns1 | 1.8 | 1.75 | 1.69 | 1.44 | 1.32 | 1.35 | 1.31 | 0.88 | 1.37 | 0.85 | 0.6 | 1.24 | - |
| Kank2 | 0.05 | 0.07 | 0.05 | 0.17 | 0.32 | 0.22 | 0.26 | 0.2 | 0.13 | 0.46 | 0.28 | 0.16 | - |
| Qpctl | 6.25 | 5.46 | 6.47 | 4.15 | 4.01 | 3.07 | 3.44 | 4.78 | 3.81 | 3.79 | 2.11 | 4.19 | - |
| A630072M18Rik | 0.32 | 0.36 | 0.33 | 0.68 | 0.68 | 0.51 | 0.85 | 0.92 | 1.05 | 1.59 | 1.44 | 1.36 | - |
| 2900026A02Rik | 1.39 | 1.6 | 1.17 | 1.14 | 1.54 | 1.07 | 4.8 | 2.93 | 3.18 | 4.9 | 2.69 | 5.09 | - |
| Arhgef28 | 0 | 0 | 0 | 0.48 | 0.2 | 0.26 | 0.17 | 0.11 | 0.03 | 0.63 | 0.45 | 0.36 | - |
| Frs2 | 1.38 | 1.14 | 1.49 | 1.67 | 1.94 | 1.96 | 1.23 | 1.73 | 1.46 | 2.33 | 2.61 | 1.48 | - |
| Zfp937 | 1.68 | 2.25 | 2.1 | 3.31 | 3.41 | 3.71 | 2.72 | 2.3 | 2.47 | 3.48 | 3.17 | 3.47 | - |
| Ndufa11 | 27.07 | 26.74 | 24.18 | 20.57 | 22.17 | 22.79 | 22.69 | 24.59 | 23.02 | 17.78 | 18.78 | 20.51 | - |
| Eomes | 2.91 | 3.01 | 2.01 | 0.56 | 0.07 | 0.48 | 0.37 | 0.16 | 0.12 | 0.03 | 0.02 | 0.05 | - |
| 8-Sep | 0.79 | 0.72 | 0.96 | 0.49 | 1.03 | 0.69 | 0.3 | 0.44 | 0.28 | 0.23 | 0.23 | 0.22 | - |
| 2310045N01Rik | 29.4 | 25.99 | 22.71 | 34.02 | 26.95 | 28.91 | 35.34 | 31.62 | 31.59 | 39.66 | 37.8 | 40.54 | - |
| Hdac5 | 6.27 | 6.74 | 5.33 | 12.26 | 12.89 | 11.37 | 16.73 | 10.55 | 11.41 | 15.18 | 17.03 | 18.57 | - |
| Osbpl5 | 3.51 | 3.86 | 1.91 | 4.5 | 3.64 | 3.31 | 9.44 | 3.87 | 5.05 | 10.37 | 7.55 | 11.33 | - |
| Enpp1 | 5.67 | 5.87 | 7.25 | 7.85 | 9.05 | 8.82 | 8.4 | 8.54 | 8.7 | 10.24 | 11.18 | 9.98 | - |
| Fundc2 | 10.14 | 11.13 | 8.9 | 14.83 | 12.39 | 14.59 | 15.56 | 11.83 | 12.03 | 19.49 | 19.79 | 20.84 | - |
| Ttc27 | 17.85 | 18.95 | 17.44 | 11.14 | 10.54 | 11.62 | 9.34 | 12.75 | 11.22 | 6.67 | 6.03 | 6.89 | - |
| Slc46a1 | 1.85 | 1.8 | 1.45 | 0.74 | 0.57 | 0.94 | 0.76 | 0.77 | 0.79 | 0.2 | 0.31 | 0.47 | - |
| Rb1cc1 | 0.84 | 0.78 | 0.9 | 1.32 | 1.43 | 1.57 | 1.13 | 1.48 | 1.85 | 2.16 | 2.06 | 1.44 | - |
| Mrps18a | 75.85 | 76.8 | 71.84 | 47.33 | 45.73 | 44.01 | 45.69 | 51.46 | 47.07 | 28.8 | 33.1 | 27.35 | - |
| Serpine1 | 0.86 | 0.98 | 1.57 | 0.36 | 0.89 | 0.55 | 0.42 | 0.43 | 0.99 | 0.13 | 0.14 | 0.11 | - |
| Ada | 7.65 | 7.73 | 8.45 | 5.16 | 3.56 | 4.76 | 7.33 | 6.11 | 7.54 | 4.47 | 3.48 | 3.18 | - |
| Il1rl1 | 2.42 | 2.3 | 2.75 | 0.97 | 1.32 | 1.27 | 0.63 | 1.18 | 1.18 | 0.71 | 0.67 | 0.71 | - |
| Mrps18c | 110.17 | 109.65 | 110.17 | 87.6 | 91.4 | 100.81 | 81.73 | 91.85 | 78.45 | 63.7 | 80.66 | 80.15 | - |
| Tsc2 | 10.3 | 9.38 | 9.12 | 8.22 | 9.18 | 7.54 | 8.88 | 7.05 | 8.4 | 8.03 | 6.99 | 8.02 | - |
| Csnk1g1 | 11.97 | 10.26 | 10.94 | 12.04 | 12.76 | 12.31 | 13.21 | 13.98 | 15.54 | 14.53 | 15.25 | 13.47 | - |
| Ano10 | 12.69 | 11.78 | 11.67 | 16.44 | 15.51 | 16.83 | 25.13 | 17.18 | 17.03 | 20.34 | 19.51 | 16.67 | - |
| Flt4 | 0 | 0 | 0 | 0 | 0 | 0 | 0.56 | 0.01 | 0.16 | 0.07 | 0.08 | 0.08 | - |
| Srebf2 | 18.45 | 18.63 | 20.07 | 20.72 | 20.3 | 21.92 | 25.76 | 22.94 | 24.52 | 24 | 24.86 | 23.54 | - |
| Hmgn1 | 116.19 | 108.26 | 113.48 | 60.04 | 73.86 | 72.22 | 49.07 | 70.68 | 62.72 | 39.57 | 51.39 | 39.57 | - |
| Pcyox1l | 5.04 | 5.11 | 5.39 | 3.13 | 4.12 | 4.65 | 2.51 | 3.17 | 3.88 | 2.01 | 2.75 | 1.94 | - |
| Eif2ak3 | 2.6 | 2.41 | 2.39 | 5.05 | 3.73 | 5.25 | 4.77 | 3.67 | 3.29 | 8.58 | 7.17 | 9.46 | - |
| C330007P06Rik | 11.95 | 10.19 | 9.83 | 9.67 | 9.05 | 10.75 | 8.93 | 8.57 | 9.88 | 8.08 | 7.78 | 8.51 | - |
| Thada | 3.03 | 2.83 | 2.69 | 8.06 | 6.46 | 6.89 | 7.93 | 4.18 | 4.58 | 14.5 | 13.86 | 17.17 | - |
| Igsf5 | 0.55 | 1.19 | 0.36 | 2.11 | 0.61 | 0.58 | 6.03 | 0.93 | 4.03 | 19.81 | 11.13 | 20.62 | - |
| Hook3 | 1.88 | 2.04 | 2.06 | 2.57 | 2.36 | 2.92 | 2.3 | 2 | 2.33 | 2.67 | 2.59 | 2.45 | - |
| 2410016O06Rik | 1.78 | 2.31 | 2.21 | 1.54 | 1.06 | 1.3 | 1.47 | 1.45 | 1.07 | 1.22 | 0.89 | 0.74 | - |
| Acsbg1 | 4.28 | 5.05 | 3.14 | 7.2 | 7.15 | 4.92 | 8.01 | 5.16 | 5.1 | 8.89 | 6.49 | 9.97 | - |
| Cd99l2 | 4.08 | 3.56 | 3.94 | 8.1 | 6.19 | 6.24 | 5.95 | 5.25 | 3.95 | 6.9 | 6.28 | 7.95 | - |
| Zranb1 | 1.72 | 1.2 | 1.58 | 2.32 | 2.96 | 2.79 | 2.08 | 2.07 | 2.4 | 3.07 | 2.61 | 2.38 | - |
| Higd1a | 82.43 | 85.34 | 87.62 | 58.93 | 63.32 | 60.86 | 89.17 | 61.09 | 72.56 | 58.76 | 62.21 | 70.61 | - |
| Smad7 | 0.59 | 1.01 | 0.9 | 1.65 | 1.69 | 1.57 | 1.98 | 1.16 | 1.45 | 2.28 | 2.23 | 2.22 | - |
| Car13 | 7.35 | 6.71 | 7.69 | 5.14 | 6.9 | 6.75 | 3.3 | 3.17 | 3.36 | 1.54 | 2.98 | 3.13 | - |
| Klra6 | 0 | 0 | 0 | 17.1 | 13.09 | 12.46 | 0 | 0 | 0 | 4.46 | 8.17 | 5.37 | - |
| Seh1l | 33.43 | 33.65 | 36.41 | 35.38 | 35.14 | 38.3 | 41.71 | 35.4 | 38.58 | 38.6 | 47.66 | 42.81 | - |
| Nars | 172.37 | 180.18 | 188.69 | 116.41 | 131.53 | 131.15 | 108.75 | 165.1 | 146.52 | 83.43 | 93.24 | 84.67 | - |
| Pgap1 | 0.29 | 0.47 | 0.37 | 0.51 | 0.67 | 0.52 | 0.99 | 0.76 | 0.64 | 0.77 | 0.86 | 0.71 | - |
| Etnk1 | 6.45 | 6.55 | 6.72 | 9.11 | 6.81 | 8.71 | 7.77 | 7.77 | 7.18 | 9.56 | 8.48 | 7.82 | - |
| D730005E14Rik | 2.04 | 2.68 | 2.66 | 4.88 | 4.57 | 3.99 | 2.27 | 2.71 | 2.87 | 5.8 | 5.09 | 5.45 | - |
| Zfp516 | 0.31 | 0.33 | 0.29 | 0.92 | 0.79 | 0.76 | 0.66 | 0.43 | 0.52 | 1 | 0.78 | 1 | - |
| Abtb2 | 15.53 | 16.4 | 17.65 | 12.95 | 14.49 | 13.32 | 12.86 | 18.44 | 16.43 | 9.8 | 11.06 | 12.36 | - |
| Btbd16 | 0.64 | 0.72 | 0.65 | 1.1 | 0.89 | 0.85 | 1.38 | 1.16 | 1 | 1.34 | 1.49 | 1.5 | - |
| Ica1 | 4.77 | 3.23 | 3.37 | 1.44 | 1.49 | 0.74 | 0.61 | 0.53 | 0.65 | 0.3 | 0.42 | 0.27 | - |
| Rasgrf1 | 0 | 0 | 0.02 | 0.53 | 0.07 | 0.07 | 1.17 | 0.09 | 0.45 | 1.86 | 1.04 | 2.43 | - |
| Piga | 10.76 | 10.59 | 9.48 | 6.43 | 8.34 | 6.6 | 10.65 | 9.94 | 11.39 | 6.86 | 6.88 | 6.31 | - |
| Zfp27 | 3.12 | 3.61 | 4.06 | 2.81 | 3.38 | 4.21 | 3.11 | 3.26 | 2.71 | 2.51 | 1.8 | 2.55 | - |
| 2900076A07Rik | 2.54 | 1.5 | 2.22 | 2.74 | 2.56 | 3.98 | 4.56 | 5.06 | 3.58 | 5.63 | 6.35 | 6.51 | - |
| Ppcdc | 12.08 | 12.38 | 10.21 | 15.99 | 15.06 | 13.69 | 16.15 | 16.57 | 15.61 | 14.84 | 14.73 | 17.77 | - |
| Kazald1 | 0.33 | 0.8 | 1.21 | 1.42 | 1.72 | 1.52 | 2.83 | 1.87 | 1.56 | 3.32 | 2.95 | 3.43 | - |
| Gm2011 | 0.61 | 0.13 | 0 | 1.02 | 1.02 | 0.69 | 0.74 | 0.32 | 0.15 | 1.32 | 1.84 | 1.1 | - |
| Hmgcl | 32.93 | 35.75 | 34.09 | 42.23 | 43.12 | 42.42 | 54.98 | 44.39 | 46.7 | 49 | 58.88 | 51.89 | - |
| Aurkb | 72.3 | 53.43 | 53.14 | 47.17 | 64.56 | 62.96 | 40.86 | 38.72 | 36.91 | 38.53 | 42.74 | 32.93 | - |
| Zkscan14 | 5.01 | 4.26 | 3.36 | 7.59 | 7.09 | 6.92 | 9.49 | 6.82 | 5.92 | 10.77 | 10.14 | 12.13 | - |
| Psme2 | 386.92 | 383.41 | 369.45 | 287.62 | 314.04 | 320.98 | 291.94 | 342.5 | 317.62 | 280.53 | 263.64 | 272.49 | - |
| Rnasek | 96.65 | 80.74 | 84.75 | 109.64 | 102.66 | 108.2 | 130.54 | 115.65 | 111.79 | 136.03 | 143.25 | 141.22 | - |
| Rhoh | 15.22 | 17.45 | 16.05 | 23.58 | 21.06 | 22.59 | 27.89 | 22.29 | 23.95 | 24.3 | 23.69 | 27.97 | - |
| Kif20b | 4.61 | 3.41 | 3.52 | 3.09 | 3.65 | 3.73 | 1.93 | 2.49 | 2.16 | 1.95 | 2.67 | 2.5 | - |
| Tbpl1 | 25.18 | 23.96 | 25.24 | 25.7 | 26.34 | 29.19 | 33.45 | 35.72 | 36.09 | 28.73 | 33.97 | 29.17 | - |
| Noc3l | 8.35 | 9.93 | 8.61 | 3.7 | 6.6 | 4.69 | 3.18 | 5.21 | 4.82 | 2.87 | 4.23 | 3.36 | - |
| 4933434E20Rik | 98 | 96.6 | 92.39 | 87.45 | 86.62 | 89.12 | 76.52 | 88.64 | 88.57 | 69.87 | 79.65 | 75.25 | - |
| Lss | 47.07 | 46.51 | 46.6 | 46.06 | 42.82 | 40.48 | 39.97 | 37.06 | 35.79 | 31.1 | 32.36 | 29.77 | - |
| Kctd8 | 0.02 | 0.02 | 0 | 1.53 | 1.64 | 1.35 | 0.04 | 0 | 0 | 1.96 | 1.4 | 1.18 | - |
| Entpd4 | 16.27 | 21.2 | 21.79 | 29.76 | 24.92 | 25.6 | 49.44 | 37.98 | 43.84 | 46.84 | 37.89 | 36.64 | - |
| Timm21 | 31.28 | 34.03 | 36.24 | 27.97 | 31.51 | 28.28 | 19.88 | 27.38 | 28.44 | 23.67 | 21.14 | 21.44 | - |
| Xpa | 5.62 | 9.75 | 10.98 | 18.26 | 15.36 | 14.88 | 23.16 | 17.86 | 18.26 | 28.51 | 27.1 | 26.45 | - |
| Stard7 | 11.61 | 14.03 | 12.95 | 15.61 | 16.16 | 16.51 | 11.07 | 13.17 | 12.65 | 19.79 | 17.96 | 15.84 | - |
| Srpk1 | 51.3 | 53.32 | 55.98 | 64.67 | 68.08 | 68.02 | 85.89 | 74.77 | 75.84 | 78.9 | 82.89 | 90.45 | - |
| Hspa4l | 13.45 | 13.1 | 16.96 | 17.83 | 20.54 | 17.93 | 21.32 | 22.44 | 21.81 | 22.35 | 20.15 | 21.02 | - |
| Inpp5k | 13.96 | 13.2 | 13.9 | 28.34 | 20.56 | 25.93 | 34.18 | 22.6 | 24.78 | 30.46 | 32.87 | 37.81 | - |

| | | | | | | | | | | | | |
|---|---|---|---|---|---|---|---|---|---|---|---|---|
| Mapkapk5 | 5.74 | 6.35 | 6.27 | 5.09 | 5.26 | 4.45 | 4.21 | 4.52 | 5.2 | 2.85 | 3.52 | 3.89 | - |
| Leprel4 | 0 | 0.03 | 0.17 | 0.98 | 1.14 | 0.72 | 0.71 | 0.63 | 1.32 | 0.74 | 0.86 | 1.11 | - |
| 1810030O07Rik | 27.22 | 25.95 | 25.61 | 27.84 | 30.15 | 30.3 | 33.09 | 30.52 | 31.04 | 31.64 | 31.3 | 35.03 | - |
| D1Ertd622e | 5.98 | 7.4 | 5.53 | 8.17 | 7.68 | 6.38 | 7.7 | 6.76 | 6.63 | 8.04 | 9.23 | 9.28 | - |
| B3gntl1 | 2.87 | 2.02 | 2.57 | 3.47 | 4.17 | 4.39 | 4.25 | 3.77 | 4.42 | 4.47 | 4.6 | 5.86 | - |
| Polm | 9.1 | 8.47 | 9.41 | 12.46 | 11.65 | 12.86 | 12.69 | 11.73 | 12.64 | 13.28 | 12.39 | 13.54 | - |
| Dennd4c | 1.18 | 1.09 | 1.24 | 2.16 | 1.48 | 1.66 | 2.31 | 1.42 | 2.12 | 2.69 | 2.6 | 2.46 | - |
| Slc12a4 | 9.16 | 8.97 | 8.76 | 7.28 | 7.57 | 7.71 | 7.25 | 9.57 | 7.74 | 6.96 | 6.87 | 7.02 | - |
| Armcx3 | 9.41 | 9.21 | 10.46 | 9.11 | 9.05 | 9.97 | 11.9 | 12.15 | 14.27 | 13.67 | 12.89 | 12.31 | - |
| Tars | 114.01 | 122.99 | 117.65 | 72.65 | 81.83 | 83.26 | 68.21 | 102.79 | 92.73 | 56.25 | 63.95 | 58.21 | - |
| Nfatc1 | 9.19 | 10.77 | 10.38 | 12.16 | 11.99 | 11.05 | 12.45 | 8.91 | 10.44 | 15.78 | 12.27 | 12.7 | - |
| Pnpla8 | 3.2 | 5.3 | 6.41 | 9.92 | 8.25 | 7.38 | 7.83 | 9.11 | 11.01 | 8.96 | 9.91 | 7.03 | - |
| Rdh11 | 27.86 | 31.01 | 31.47 | 27.1 | 25.86 | 27.9 | 24.66 | 34.7 | 28.94 | 19.77 | 25.33 | 23.18 | - |
| Mrpl39 | 36.64 | 39.42 | 39.99 | 27.46 | 30.86 | 27.85 | 25.51 | 29.49 | 26.22 | 16.3 | 20.56 | 21.16 | - |
| Glcci1 | 0.51 | 0.61 | 0.83 | 1.31 | 1 | 1.36 | 1.59 | 0.53 | 0.78 | 1.82 | 1.39 | 1.47 | - |
| Lin7b | 0 | 0.08 | 0.15 | 0.18 | 0.17 | 0.62 | 2.65 | 0.25 | 0.97 | 0.96 | 3.1 | 2.17 | - |
| Asxl1 | 9.26 | 9.01 | 8.31 | 11.22 | 11.27 | 12.02 | 14.34 | 12.53 | 12.74 | 15.28 | 13.76 | 16.8 | - |
| Rnaseh2a | 26.89 | 29.47 | 29.63 | 23.69 | 26.17 | 23.21 | 21.96 | 25.18 | 23.12 | 16.78 | 17.85 | 18.25 | - |
| Ceacam1 | 0.47 | 0.47 | 0.1 | 0.14 | 0.12 | 0.04 | 0.49 | 0.53 | 0.37 | 0.02 | 0.07 | 0.05 | - |
| Sapcd2 | 8.01 | 6.81 | 6.32 | 4.92 | 6.62 | 6.42 | 3.65 | 2.58 | 3.78 | 3.54 | 2.35 | 2.72 | - |
| Heatr1 | 7.1 | 7.13 | 7.76 | 3.9 | 3.77 | 3.94 | 3.19 | 4.07 | 3.96 | 2.99 | 2.15 | 2.34 | - |
| Dgkd | 13.64 | 14.35 | 13.83 | 12.11 | 12.28 | 11.74 | 11.65 | 12.05 | 12.64 | 10.94 | 9.52 | 8.09 | - |
| Itpr3 | 2.39 | 2.21 | 2.76 | 4.08 | 4.62 | 4.47 | 4.89 | 3.07 | 4.28 | 6.01 | 6.15 | 5.23 | - |
| Pgk1 | 1192.36 | 1158.63 | 1194.89 | 1489.66 | 1378.82 | 1440.61 | 2255.73 | 1719.2 | 1948.26 | 1748 | 1904.54 | 1860.33 | - |
| Ston1 | 0.15 | 0.29 | 0.25 | 0.53 | 0.59 | 0.75 | 0.39 | 0.28 | 0.13 | 1 | 0.87 | 0.73 | - |
| Lzic | 27.31 | 27.6 | 25.16 | 17.89 | 21.11 | 18.12 | 20.44 | 18.97 | 22.76 | 14 | 16.86 | 15.95 | - |
| Cage1 | 0.12 | 0.35 | 0.36 | 0.69 | 0.61 | 0.42 | 0.68 | 0.66 | 0.88 | 0.92 | 0.91 | 1.07 | - |
| Ufc1 | 72.25 | 70.95 | 70.83 | 56.06 | 54.75 | 66.28 | 69.92 | 78.6 | 72.23 | 59.82 | 60.29 | 58.48 | - |
| Fastkd1 | 3.14 | 2.26 | 3.9 | 2.92 | 2.69 | 2.97 | 1.53 | 2.34 | 2.55 | 1.7 | 2.29 | 1.29 | - |
| Pold1 | 46.07 | 45.23 | 42.81 | 38.49 | 46.25 | 41.96 | 34 | 33.78 | 34.94 | 31.29 | 35.98 | 29.81 | - |
| Zfp830 | 8.12 | 7.86 | 7.7 | 5.19 | 6.59 | 6.24 | 5.38 | 5.38 | 6.11 | 5.51 | 4.56 | 5.06 | - |
| Pea15a | 14.11 | 12.5 | 12.75 | 21.13 | 17.38 | 15.81 | 22.99 | 19.66 | 17.76 | 24.82 | 23.96 | 28.03 | - |
| Tle6 | 1.9 | 1.43 | 1.82 | 3.01 | 3.35 | 2.54 | 6.39 | 3.64 | 5.3 | 4.88 | 5.75 | 5.86 | - |
| Kcnip2 | 0.65 | 0.52 | 0.51 | 1.51 | 1.34 | 1.3 | 1.4 | 1.43 | 1.02 | 2.22 | 1.87 | 1.72 | - |
| Ppie | 79.36 | 79.99 | 80.73 | 59.3 | 59.59 | 64.05 | 57.57 | 72.8 | 68.03 | 48.29 | 52.87 | 51.39 | - |
| Gucy1a3 | 0.27 | 0.26 | 0.3 | 0.04 | 0 | 0.08 | 0.15 | 0.21 | 0.14 | 0 | 0.08 | 0.06 | - |
| Cd200r4 | 1.07 | 0.86 | 1.2 | 1.75 | 1.61 | 1.36 | 2.5 | 0.83 | 1.2 | 2.88 | 2.23 | 2.92 | - |
| Rps9 | 1484.38 | 1508.11 | 1508.11 | 2024.62 | 2002.76 | 2099.47 | 1981.93 | 1836.42 | 1917.53 | 2177.1 | 2297.48 | 2178.25 | - |
| AI462493 | 25.28 | 24.93 | 27.64 | 38.64 | 37.11 | 32.7 | 37.13 | 29.83 | 27.81 | 34.76 | 37.75 | 36.83 | - |
| Lpin2 | 3.07 | 2.88 | 2.93 | 5.16 | 3.44 | 3.39 | 4.04 | 3.59 | 3.47 | 5.19 | 5.88 | 5.93 | - |
| Camk2g | 10.71 | 10.82 | 9.06 | 15.86 | 14.51 | 17.33 | 18.45 | 15.26 | 14.99 | 18.48 | 22.04 | 18.42 | - |
| Phgdh | 557.58 | 580.46 | 565.96 | 313.57 | 364.92 | 358.59 | 319.54 | 452.28 | 410.66 | 226.68 | 229.13 | 212.72 | - |
| Troap | 10.06 | 8.84 | 7.61 | 7.5 | 11.08 | 10.64 | 4.68 | 3.83 | 5.13 | 5.95 | 5.51 | 5.18 | - |
| Klhl25 | 7.79 | 9.56 | 9.29 | 13.67 | 11.16 | 10.26 | 14.6 | 10.41 | 11.91 | 18.02 | 15.07 | 18.5 | - |
| Lars | 64.23 | 64.58 | 67.88 | 35.62 | 40.71 | 38.28 | 29.16 | 48.99 | 45.81 | 23.8 | 22.91 | 20.92 | - |
| Slc2a1 | 30.14 | 28.98 | 28.97 | 16.42 | 17.52 | 15.89 | 19.01 | 18.86 | 20.93 | 8.19 | 12.36 | 16.3 | - |
| Polg2 | 6.86 | 5.8 | 6.03 | 8.65 | 8 | 9.78 | 12.44 | 9.7 | 10.94 | 12.35 | 16.08 | 11.48 | - |
| Dancr | 15.39 | 16.11 | 16.02 | 14.7 | 12.91 | 12.04 | 7.06 | 10.94 | 11.81 | 12.59 | 11.06 | 7.55 | - |
| Gm16845 | 1.36 | 1.35 | 1.66 | 2.93 | 2.13 | 1.35 | 3.53 | 2 | 2.71 | 4.29 | 3.07 | 4.05 | - |
| Zc3h8 | 10.36 | 9.28 | 11.67 | 6.17 | 7.61 | 5.9 | 6.97 | 9.75 | 8.25 | 4.23 | 6.17 | 5.4 | - |
| Gpsm3 | 6.21 | 7.11 | 6.44 | 10.31 | 9.34 | 7.22 | 14.93 | 10.8 | 9.42 | 14.31 | 14.5 | 13.78 | - |
| Lsm11 | 0.61 | 0.57 | 0.57 | 0.99 | 0.99 | 0.9 | 0.9 | 0.83 | 0.69 | 1.15 | 1.47 | 1.06 | - |
| Rbm17 | 69.52 | 88.68 | 104.88 | 68.07 | 72.72 | 67.79 | 71.25 | 83.26 | 81.07 | 61.42 | 66.49 | 54.79 | - |
| Htra3 | 0 | 0 | 0.02 | 0.31 | 0.08 | 0.18 | 0.32 | 0.05 | 0.2 | 0.42 | 0.57 | 1.35 | - |
| Rab31 | 0.84 | 1.87 | 1.03 | 2.6 | 1.8 | 1.85 | 4.96 | 4.92 | 4.45 | 5.46 | 4.73 | 5.65 | - |
| Ubl5 | 30.28 | 30.72 | 30.8 | 37.97 | 34.77 | 37.72 | 45.7 | 39.94 | 40.78 | 45.8 | 46.69 | 46.59 | - |
| Usp53 | 0.93 | 0.75 | 0.69 | 0.99 | 1.18 | 0.81 | 1.07 | 1.17 | 1.31 | 1.51 | 2.53 | 1.56 | - |
| Lipg | 1.04 | 0.92 | 0.96 | 0.42 | 0.54 | 0.89 | 0.59 | 0.56 | 0.64 | 0.23 | 0.31 | 0.29 | - |
| Zfp1 | 6.78 | 5.97 | 6.06 | 10.71 | 8.89 | 9.61 | 7.69 | 7.53 | 6.25 | 10 | 8.47 | 9.33 | - |
| Crlf3 | 44.44 | 55.07 | 58.22 | 39.25 | 39.16 | 36.68 | 40.46 | 45.39 | 42.82 | 33.91 | 33.62 | 39.39 | - |
| Pan3 | 6.81 | 6.97 | 7.36 | 8.34 | 8.69 | 8.27 | 8.82 | 7.32 | 7.84 | 9.69 | 9.68 | 8.62 | - |
| Nol6 | 15.75 | 15.98 | 17.3 | 10.25 | 9.61 | 8.26 | 6.22 | 10.34 | 8.65 | 6.66 | 6.26 | 5.48 | - |
| Def6 | 64.32 | 63.07 | 63.42 | 70.97 | 70.13 | 72.48 | 89.32 | 76.58 | 86.85 | 85.29 | 79.71 | 82.81 | - |
| Ogfod2 | 15.28 | 14.85 | 14.32 | 22.05 | 21.66 | 20.76 | 22.97 | 23.07 | 18.89 | 25.16 | 29.26 | 27.14 | - |
| Wbp4 | 8.12 | 9.46 | 10.27 | 7.25 | 7.45 | 7.69 | 6.14 | 6.86 | 6.25 | 5.08 | 5.57 | 5.63 | - |
| Sco2 | 12.33 | 12.71 | 14.34 | 6.88 | 9.81 | 7.57 | 7.94 | 11.16 | 8.54 | 5.33 | 7.45 | 5.88 | - |
| Tceanc2 | 4.03 | 4.64 | 3.74 | 4.91 | 4.44 | 4.59 | 5.5 | 5.19 | 4.62 | 5.69 | 6.09 | 5.81 | - |
| Lmtk3 | 0.04 | 0.02 | 0.07 | 0.08 | 0.12 | 0.11 | 0.13 | 0.17 | 0.11 | 0.27 | 0.18 | 0.25 | - |
| Yae1d1 | 9.83 | 10.38 | 10.81 | 5.83 | 7.65 | 6.25 | 4.93 | 6.73 | 6.64 | 4.34 | 3.89 | 4.6 | - |
| Usp48 | 4.9 | 6.4 | 6.02 | 10.23 | 8.59 | 9.99 | 11.69 | 10.09 | 9.77 | 14.19 | 13.12 | 12.92 | - |
| AI837181 | 3.99 | 3.46 | 2.54 | 5.53 | 3.75 | 3.5 | 5.37 | 4.57 | 4.28 | 4.6 | 6.4 | 7.33 | - |
| Nek4 | 4.67 | 4.95 | 5.16 | 4.22 | 4.07 | 4.49 | 3.09 | 3.85 | 3.54 | 2.85 | 3.09 | 2.06 | - |
| Rad54b | 7.96 | 8.29 | 7.87 | 4.36 | 5.39 | 4.98 | 3.52 | 4.18 | 3.63 | 2.64 | 2.71 | 2.1 | - |
| Ttc9c | 21.26 | 18.67 | 19.38 | 23.02 | 22.89 | 21.92 | 23.46 | 23.96 | 22.87 | 23.53 | 30.48 | 22.64 | - |
| Dr1 | 22.09 | 23.99 | 23.93 | 19.73 | 19.94 | 18.8 | 16.64 | 19.36 | 20.35 | 18.55 | 16.91 | 17.07 | - |
| Plekhg2 | 2.36 | 1.37 | 1.88 | 4.72 | 4.33 | 5.18 | 6.17 | 3.64 | 3.59 | 6.81 | 4.97 | 7.89 | - |
| Zfp90 | 1.87 | 2.61 | 2.9 | 3.43 | 4.41 | 3.8 | 2.18 | 3.17 | 2.69 | 4.52 | 2.79 | 5.23 | - |
| Lmbr1l | 6.16 | 6.18 | 5.31 | 10.44 | 9.44 | 10.71 | 8.49 | 6.99 | 8.57 | 11.35 | 15.69 | 10.91 | - |
| Ino80d | 1.92 | 2.17 | 2.25 | 2.51 | 2.54 | 2.84 | 2.08 | 2.3 | 2.47 | 2.56 | 3.16 | 2.77 | - |
| Dusp6 | 10.57 | 8.49 | 7.69 | 8.84 | 6.94 | 7.27 | 8.01 | 5.73 | 8.65 | 5.1 | 3.35 | 4.47 | - |
| Fads1 | 16.15 | 17.74 | 17.5 | 15.48 | 15.4 | 15.53 | 13.43 | 13.64 | 13.58 | 11.02 | 13.06 | 12.62 | - |
| Arl6 | 13.51 | 12.56 | 13.44 | 8.66 | 10.73 | 10.02 | 8.18 | 9.65 | 9.47 | 5.85 | 8.53 | 8.32 | - |
| Gbp7 | 8.41 | 9.43 | 8.44 | 10.19 | 9.17 | 10.03 | 12.59 | 10.02 | 10.23 | 15.8 | 11.02 | 13.75 | - |
| Tm4sf19 | 0.45 | 0.32 | 0.65 | 0 | 0 | 0 | 0 | 0 | 0 | 0 | 0 | 0 | - |
| Ovgp1 | 0.2 | 0.28 | 0.28 | 0.93 | 0.69 | 0.73 | 0.32 | 0.95 | 0.49 | 1.25 | 1.34 | 1.1 | - |
| Hscb | 27.91 | 29.09 | 25.29 | 51.72 | 48.58 | 60.36 | 63.04 | 48.29 | 50.34 | 73.78 | 75.8 | 80.73 | - |
| Tomm6 | 50.73 | 54.93 | 57.7 | 31.78 | 31.25 | 34.39 | 36.26 | 39.98 | 37.1 | 23.93 | 25.59 | 20.86 | - |
| Pick1 | 13.33 | 13.82 | 13.94 | 9.14 | 11.93 | 11.48 | 10.6 | 13.51 | 12.21 | 8.39 | 6.85 | 9.29 | - |
| Cbfb | 9.74 | 8.34 | 8.31 | 9.72 | 10.83 | 9.38 | 10.71 | 10.7 | 9.95 | 10.99 | 12.39 | 11.29 | - |
| Hist2h2be | 0.55 | 0.51 | 0.72 | 0.89 | 0.48 | 0.37 | 1.06 | 0.69 | 1.09 | 1.05 | 1.44 | 1.39 | - |
| Melk | 27.19 | 22.96 | 24.05 | 17.87 | 20.95 | 17.88 | 11.52 | 13.83 | 13.44 | 9.01 | 11.45 | 7.91 | - |
| Cstb | 197.61 | 199.43 | 195.16 | 153.32 | 167.26 | 178.5 | 201.21 | 196.18 | 205.51 | 148.24 | 155.2 | 153.78 | - |
| Phf17 | 1.95 | 1.88 | 1.73 | 1.77 | 1.23 | 1.48 | 1.08 | 1.08 | 1.23 | 1.16 | 0.97 | 0.96 | - |
| Mpp2 | 1.03 | 1.15 | 0.42 | 3.96 | 3.07 | 3.25 | 2.92 | 1.15 | 1.72 | 3.26 | 3.62 | 3.38 | - |
| Vgll4 | 1.41 | 1.26 | 1.84 | 4.15 | 3.19 | 3.35 | 3.29 | 2 | 2.9 | 6.24 | 6.53 | 4.15 | - |
| Capns1 | 18.8 | 14.45 | 15.76 | 22.23 | 26.54 | 25.14 | 19.91 | 23.04 | 23.67 | 25.64 | 20.3 | 24.75 | - |
| Moap1 | 1.38 | 2.81 | 1.58 | 0.65 | 0 | 0.81 | 0.42 | 0.86 | 1.89 | 1.15 | 0.65 | 0.76 | - |
| Ankrd37 | 9.73 | 9.98 | 10.47 | 8.11 | 8.91 | 10.11 | 30.96 | 18.52 | 25.41 | 12.28 | 20.45 | 22.7 | - |
| Gm7102 | 8.53 | 7.69 | 7.13 | 11.66 | 10.23 | 10.81 | 10.93 | 10.69 | 10.42 | 13.24 | 14.55 | 12.86 | - |
| Bcl2l1 | 61.96 | 50.53 | 55.77 | 41.05 | 47.44 | 46.56 | 31.36 | 53.11 | 48.17 | 31.63 | 33.04 | 29.43 | - |
| Eif3h | 439.7 | 436.33 | 436.53 | 538.6 | 543.86 | 565.34 | 582.61 | 525.59 | 525.39 | 587.4 | 634.5 | 590.24 | - |
| Rab11b | 20.94 | 19.87 | 19.37 | 17.23 | 19.09 | 18.07 | 16.9 | 17.82 | 16.22 | 16.11 | 14.84 | 15.93 | - |
| Cpd | 0.9 | 0.88 | 0.96 | 0.97 | 1.08 | 1 | 0.2 | 0.12 | 0.39 | 0.34 | 0.26 | 0.28 | - |
| Tmem106b | 4.44 | 4.83 | 4.05 | 7.48 | 5.93 | 7.12 | 5.41 | 6.53 | 6.64 | 8.45 | 8.87 | 8.95 | - |
| Cnpy2 | 39.96 | 41.77 | 41.27 | 36.98 | 35.93 | 38.82 | 32.66 | 34.3 | 32.76 | 27.33 | 28.24 | 27.21 | - |
| Ep300 | 4.47 | 4.48 | 4.36 | 6.77 | 5.59 | 6.32 | 7.67 | 5.91 | 6.99 | 9.45 | 7.54 | 9.09 | - |
| Slc25a33 | 1.89 | 2.64 | 1.93 | 0.71 | 0.63 | 1.16 | 1.1 | 1.25 | 1.03 | 1.37 | 0.72 | 0.34 | - |
| Bin3 | 30.62 | 24.29 | 26.67 | 35.83 | 31.52 | 34.67 | 36.1 | 29.49 | 35.41 | 36.5 | 33.05 | 40.56 | - |
| Gm19557 | 0.33 | 0.46 | 0.47 | 1.37 | 1.45 | 1.32 | 0.65 | 0.79 | 0.76 | 1.95 | 0.87 | 1.75 | - |
| C2cd2 | 0.59 | 0.71 | 0.64 | 1.49 | 1.26 | 0.97 | 1.31 | 0.54 | 0.57 | 1.31 | 1.39 | 1.29 | - |
| Acbd6 | 13.25 | 15.6 | 18.31 | 24.9 | 21.89 | 18.56 | 27.05 | 29.75 | 29.42 | 34.71 | 32.09 | 23.92 | - |
| Arrb2 | 9.21 | 10.99 | 8.81 | 20.37 | 14.96 | 15.35 | 13.74 | 11.75 | 13.57 | 18.28 | 17.44 | 16.38 | - |
| Mt1 | 420.29 | 429.01 | 461.98 | 657.46 | 716.37 | 708 | 882.74 | 487.32 | 573.46 | 1028.03 | 1138 | 1165.56 | - |
| Zap70 | 64.7 | 66.38 | 64.98 | 95.04 | 76.25 | 84.16 | 106.04 | 72.99 | 77.97 | 123.75 | 108.98 | 131.36 | - |
| Trnau1ap | 58.57 | 52.83 | 55.17 | 44.06 | 46.86 | 47.69 | 38.31 | 53.69 | 45.75 | 42.12 | 43.69 | 39.79 | - |
| Rcc2 | 23.79 | 25.95 | 25.31 | 21.38 | 23.94 | 21.6 | 19.33 | 24.21 | 22.37 | 19.34 | 20.48 | 17.27 | - |
| 2700029M09Rik | 85.49 | 90.39 | 89.31 | 58.39 | 59.24 | 65.32 | 47.92 | 59.57 | 56.97 | 30.76 | 34.59 | 35.18 | - |
| Ykt6 | 18.65 | 18.71 | 17.36 | 12.69 | 16.26 | 13.91 | 12.65 | 14.9 | 13.46 | 12.38 | 13.37 | 10.85 | - |
| Mthfd1l | 36.1 | 36.42 | 37.51 | 28.49 | 32.91 | 28.31 | 38.68 | 30.49 | 33.79 | 30.44 | 29.82 | 30.27 | - |
| Nr2c2 | 0.52 | 0.5 | 0.62 | 1.13 | 0.96 | 0.88 | 0.81 | 1.06 | 0.95 | 1.44 | 1.23 | 1.14 | - |
| Gars | 262.13 | 267.65 | 263.73 | 173.08 | 210.9 | 206.04 | 199.95 | 279.94 | 258.11 | 150.62 | 177.87 | 152.35 | - |
| Atp8b4 | 15.57 | 16.86 | 14.52 | 10.46 | 7.95 | 6.4 | 10.18 | 7.24 | 8.72 | 6.62 | 4.37 | 7.48 | - |
| Gprin3 | 6.8 | 8.41 | 8.53 | 5.77 | 6.24 | 5.99 | 8.64 | 9.44 | 10.52 | 5.39 | 5.79 | 4.5 | - |
| Gak | 23.9 | 24.37 | 22.72 | 26.27 | 25.66 | 27.5 | 24.92 | 25.85 | 26.95 | 28.46 | 30.42 | 29.74 | - |
| Coq6 | 20.88 | 21.08 | 17.61 | 16.25 | 14.69 | 12.5 | 13.45 | 14.48 | 14.97 | 9.86 | 12.12 | 6.87 | - |
| Trpc4ap | 10.33 | 10.13 | 11.21 | 17.19 | 16.48 | 17.03 | 19.71 | 17.87 | 17.56 | 22.11 | 26.81 | 22.07 | - |
| Sesn2 | 14.73 | 15.32 | 13.22 | 11.81 | 16.86 | 13.33 | 10.64 | 13.58 | 13.12 | 6.89 | 9.15 | 8.6 | - |
| Pag1 | 1.05 | 1.36 | 1.1 | 2.01 | 1.54 | 1.67 | 1.97 | 1.31 | 1.51 | 2.28 | 2.37 | 2.36 | - |
| Camk2n1 | 0.33 | 0.27 | 0.16 | 0.42 | 0.24 | 0.21 | 1.1 | 0.72 | 0.64 | 0.82 | 0.53 | 0.96 | - |
| Zfp821 | 0.71 | 0.56 | 0.47 | 0.97 | 0.81 | 0.64 | 0.77 | 0.87 | 0.83 | 1.87 | 1.6 | 1.11 | - |
| Tanc2 | 0.71 | 0.51 | 0.45 | 0.45 | 0.48 | 0.36 | 2.37 | 1.16 | 1.53 | 1.79 | 1.26 | 1.98 | - |
| Slc15a2 | 0 | 0.25 | 0.13 | 0.4 | 0.2 | 0 | 0.41 | 0.09 | 0.23 | 0.76 | 0.94 | 1.15 | - |
| Cyhr1 | 7.19 | 7.36 | 7.07 | 10.73 | 9.78 | 10.66 | 13.12 | 8.95 | 10.69 | 14.36 | 14.59 | 15.08 | - |
| Msrb2 | 5.9 | 7.04 | 4.45 | 2.6 | 3.46 | 4.07 | 4.59 | 4.39 | 4.28 | 2.64 | 2.95 | 2.71 | - |
| Tnk2 | 1.93 | 2.48 | 2.58 | 1.68 | 1.97 | 1.13 | 1.47 | 1.47 | 1.14 | 1.77 | 1.19 | 1.32 | - |
| Zmym5 | 3.1 | 2.84 | 2.5 | 3.82 | 4.04 | 4.18 | 4.3 | 3.69 | 3.47 | 5.8 | 6.13 | 5.58 | - |
| Ube2g1 | 15.49 | 16.44 | 15.26 | 16.27 | 18.25 | 19.48 | 15.76 | 16.26 | 17.98 | 18.47 | 19.25 | 20.31 | - |
| Fam5c | 0 | 0.02 | 0.04 | 1.46 | 1.51 | 0.95 | 0 | 0 | 0.02 | 1.25 | 0.99 | 1.14 | - |
| Ndufs2 | 151.25 | 144.71 | 141.96 | 127.54 | 133.48 | 133.39 | 130.28 | 139.6 | 141.62 | 116.23 | 125.01 | 113.99 | - |
| Far2 | 0.67 | 0.64 | 0.5 | 1.09 | 1.28 | 1.04 | 0.86 | 1.14 | 0.84 | 1.24 | 1.19 | 1.12 | - |
| Arsa | 4.5 | 3.58 | 4.43 | 7.04 | 7.49 | 7.4 | 8.49 | 5.77 | 6.44 | 7.62 | 7.06 | 8.95 | - |
| Mtg1 | 35.94 | 33.94 | 33.63 | 20.92 | 23.36 | 23.59 | 18.37 | 28.65 | 25.7 | 15.21 | 16.77 | 14.31 | - |
| 6330403M23Rik | 0.29 | 1.31 | 0.31 | 1.88 | 1.49 | 2.96 | 1.63 | 1.28 | 1.48 | 2.4 | 2.28 | 2.38 | - |
| Afmid | 2.99 | 2.35 | 2.02 | 3.74 | 2.73 | 3.72 | 4.71 | 2.24 | 3.66 | 4.98 | 3.57 | 5.04 | - |
| Grin1 | 0 | 0.03 | 0.03 | 0.38 | 0.41 | 0.35 | 1.24 | 0.68 | 1.34 | 0.87 | 0.58 | 1.13 | - |
| Sat1 | 155.02 | 138.04 | 154.71 | 204.45 | 192.18 | 200.29 | 196.15 | 193.14 | 190.31 | 225.03 | 237.01 | 232.51 | - |
| Usp9x | 5.99 | 6.11 | 6.47 | 8.7 | 8.27 | 9.22 | 7.63 | 7.34 | 7.79 | 9.01 | 10.74 | 9.68 | - |
| Rbpj | 86.97 | 82.03 | 97.27 | 86.82 | 96.82 | 91.4 | 80.51 | 100.12 | 99.86 | 67.83 | 76.86 | 64.03 | - |
| Katnb1 | 9.52 | 8.9 | 9.47 | 6.9 | 7.69 | 7.58 | 6.73 | 6.88 | 6.89 | 7.02 | 6.12 | 5.42 | - |
| Btn1a1 | 0.05 | 0.17 | 0.02 | 0.49 | 0.18 | 0.09 | 0.67 | 0.22 | 0.42 | 0.56 | 0.48 | 0.6 | - |
| Pdcl3 | 48.28 | 44.69 | 47.85 | 34.5 | 36.1 | 40.04 | 36 | 36.07 | 36.95 | 28.27 | 29.48 | 32.49 | - |

| | | | | | | | | | | | | | |
|---|---|---|---|---|---|---|---|---|---|---|---|---|---|
| Lamc1 | 3.13 | 2.72 | 3.05 | 4.64 | 5.08 | 4.48 | 4.07 | 5.39 | 5.04 | 6.68 | 6.04 | 6.55 | - |
| Ddt | 98.15 | 84.94 | 106.09 | 55.21 | 60.68 | 64.71 | 68.98 | 89.21 | 81.14 | 51.01 | 58.7 | 52.72 | - |
| Smg8 | 11.8 | 12.01 | 13.13 | 9.1 | 7.08 | 9.58 | 7.33 | 11.14 | 8.93 | 9.11 | 8.23 | 7.53 | - |
| Zfp213 | 5.01 | 4.18 | 4.65 | 3.05 | 2.89 | 2.77 | 3.48 | 1.91 | 2.29 | 3.04 | 1.71 | 2.52 | - |
| Agtrap | 4.86 | 4.46 | 3.85 | 5.89 | 5.12 | 4.99 | 7.29 | 5.44 | 4.75 | 6.44 | 6.91 | 7.1 | - |
| Nfs1 | 47.41 | 41.92 | 48.8 | 35.87 | 34.51 | 36.34 | 30.42 | 40.6 | 37 | 31.35 | 29.66 | 31.31 | - |
| Itga1 | 0 | 0.02 | 0.01 | 0 | 0 | 0 | 0.33 | 0.06 | 0.06 | 0.12 | 0.03 | 0.23 | - |
| Fam185a | 11.88 | 14.17 | 14 | 11.87 | 11.36 | 10.74 | 8.93 | 13.17 | 10.65 | 7.52 | 8.2 | 10.64 | - |
| 9930111J21Rik1 | 4.01 | 3.38 | 4 | 6.69 | 5.59 | 5.61 | 10.74 | 5.66 | 7.34 | 11.37 | 8.16 | 12.93 | - |
| Mtr | 6.9 | 7.45 | 7.49 | 4.83 | 5.43 | 5.15 | 3.37 | 5.34 | 4.26 | 3.79 | 3.47 | 2.44 | - |
| Cxcr6 | 7.3 | 11.8 | 5.11 | 11.55 | 3.63 | 2.36 | 64.96 | 12.73 | 22.84 | 41.79 | 20 | 47.69 | - |
| Dmwd | 3.62 | 3.93 | 4.06 | 1.66 | 2.45 | 2.02 | 5.09 | 2.87 | 4.47 | 2.54 | 1.86 | 1.86 | - |
| Zfp580 | 1.3 | 2.75 | 2.6 | 5.28 | 2.78 | 3.47 | 4.12 | 3.46 | 3.69 | 5.61 | 3.97 | 4.97 | - |
| 9430020K01Rik | 5.52 | 5.4 | 5.74 | 11.58 | 10.68 | 10.99 | 3.03 | 4.48 | 5.05 | 6.77 | 7.02 | 7.52 | - |
| Zfp426 | 5.57 | 6.49 | 5.74 | 8.57 | 5.92 | 6.93 | 6.53 | 7.05 | 6.38 | 7.55 | 7.83 | 9.81 | - |
| Luzp1 | 1.52 | 1.51 | 1.36 | 0.94 | 1.17 | 1.59 | 0.97 | 1.04 | 1.3 | 0.94 | 0.83 | 0.85 | - |
| Galc | 1.96 | 2.25 | 2.17 | 4.54 | 4.28 | 4.25 | 2.03 | 3.09 | 2.52 | 3.88 | 4.68 | 4.51 | - |
| Brap | 2.88 | 3.93 | 3.95 | 4.91 | 4 | 4.47 | 3.84 | 4.54 | 4.47 | 5.05 | 5.08 | 5.17 | - |
| Qprt | 1.38 | 0.92 | 1.13 | 2.11 | 1.77 | 2.02 | 4.13 | 1.91 | 2.15 | 4.01 | 3.66 | 3.85 | - |
| Fryl | 2.54 | 2.32 | 2.33 | 3.11 | 2.89 | 3.08 | 3.17 | 2.97 | 2.92 | 3.41 | 3.26 | 3.07 | - |
| Exosc5 | 45.63 | 49.16 | 47.93 | 34.7 | 36.75 | 42.17 | 39.02 | 45.66 | 46.83 | 26.08 | 30.01 | 27.17 | - |
| Tbkbp1 | 0.43 | 0.37 | 0.42 | 0.78 | 0.41 | 0.67 | 1.32 | 1.74 | 1.62 | 2.18 | 2.33 | 2.26 | - |
| Fam98b | 3.28 | 2.97 | 4.33 | 1.6 | 2.58 | 2.36 | 1.88 | 3.06 | 2.54 | 1.35 | 2.02 | 2.2 | - |
| Mvk | 16.07 | 16.06 | 16.36 | 11.39 | 13.09 | 14.76 | 14.93 | 12.53 | 11.93 | 10.34 | 12.41 | 12.13 | - |
| A930015D03Rik | 1.8 | 1.41 | 2.81 | 3.09 | 1.09 | 1.67 | 3.45 | 2.29 | 2.87 | 5.69 | 4.78 | 4.01 | - |
| Ddx25 | 1.06 | 0.59 | 0.36 | 0.19 | 0.04 | 0.23 | 0.27 | 0.42 | 0.1 | 0.14 | 0.18 | 0.03 | - |
| 1190002F15Rik | 16.58 | 13.1 | 14.31 | 20.42 | 28.5 | 29.08 | 25.77 | 15.24 | 20.1 | 29.68 | 26.26 | 25.85 | - |
| Lrrc40 | 20.9 | 20.04 | 19.67 | 19.37 | 20.17 | 20.1 | 14.3 | 16.58 | 16.47 | 13.2 | 13.6 | 11.37 | - |
| Vamp5 | 1.04 | 1.16 | 1.12 | 1.66 | 1.76 | 2.07 | 2.19 | 2.21 | 2.42 | 2.9 | 1.43 | 2.89 | - |
| Pgm2l1 | 0.21 | 0.26 | 0.24 | 0.56 | 0.65 | 0.47 | 0.63 | 0.29 | 0.54 | 0.71 | 0.82 | 1.25 | - |
| Paox | 34.51 | 32.25 | 33.62 | 22.2 | 21.24 | 20.49 | 21.5 | 22.11 | 24.68 | 13.44 | 16.82 | 16.06 | - |
| Mul1 | 5.63 | 5.54 | 4.98 | 4.22 | 3.41 | 3.86 | 3.26 | 4.28 | 4 | 2.84 | 2.26 | 2.72 | - |
| Runx1 | 0.24 | 0.25 | 0.31 | 1.07 | 0.75 | 0.75 | 1 | 0.32 | 0.76 | 1.46 | 1.33 | 1.75 | - |
| B3galt6 | 6.11 | 6.32 | 7.24 | 4.05 | 3.21 | 3.14 | 3.04 | 3.95 | 3.85 | 1.98 | 1.71 | 1.76 | - |
| Lrrc28 | 6.98 | 5.63 | 6.28 | 8.06 | 8.34 | 7.21 | 12.27 | 9.08 | 11.19 | 10.85 | 11.34 | 9.8 | - |
| Tomm20 | 84.2 | 93.19 | 88.29 | 61.5 | 62.68 | 65.39 | 61.19 | 71.21 | 70.51 | 51.86 | 55.02 | 48.95 | - |
| Stxbp3a | 11.18 | 11.38 | 11.43 | 13.17 | 15.39 | 17.05 | 11.29 | 12.61 | 12.02 | 15.6 | 13.51 | 15.55 | - |
| Yrdc | 3.09 | 4.42 | 4.2 | 1.95 | 2.33 | 1.81 | 1.96 | 2.14 | 2.59 | 1.77 | 1.25 | 1.74 | - |
| Naa25 | 40.65 | 38.74 | 40.34 | 26.65 | 30.16 | 28.89 | 28 | 36.93 | 33.78 | 24.78 | 24.49 | 25.71 | - |
| Bst1 | 2.22 | 3.11 | 2.18 | 2.16 | 0.91 | 1.36 | 16.26 | 6.81 | 10.86 | 4.26 | 3.3 | 5.72 | - |
| Glul | 6.53 | 6.11 | 6.04 | 11.2 | 10.28 | 11.92 | 11.73 | 10.16 | 9.64 | 14.48 | 15.35 | 15.93 | - |
| L2hgdh | 9.12 | 7.53 | 9.71 | 4.68 | 6.4 | 5.75 | 3.96 | 5.38 | 4.71 | 2.49 | 4.3 | 3.56 | - |
| Mfsd7b | 3.87 | 3.09 | 3.48 | 2.81 | 3.24 | 2.95 | 2.39 | 1.88 | 2.27 | 1.73 | 1.93 | 2.24 | - |
| Vps29 | 129.46 | 128.26 | 127.73 | 78.11 | 93.35 | 90.78 | 83.4 | 101.44 | 102.24 | 65.4 | 72.14 | 76.55 | - |
| Rabl3 | 16.94 | 13.81 | 19.01 | 9.95 | 9.35 | 12.02 | 11.11 | 14.47 | 11.7 | 8.16 | 9.24 | 8.07 | - |
| Clip1 | 6.55 | 7.05 | 8.15 | 10.53 | 9.79 | 8.87 | 12.26 | 11.91 | 11.86 | 12.98 | 13.28 | 10.65 | - |
| Ccdc163 | 9.92 | 9.78 | 9.31 | 14.16 | 16.25 | 15.55 | 13.25 | 11.71 | 14.19 | 15.92 | 19.04 | 17.93 | - |
| Pex16 | 21.83 | 22.31 | 19.47 | 32.45 | 24.36 | 27.89 | 29.42 | 26.23 | 27.15 | 26.19 | 32.95 | 29.6 | - |
| Copg1 | 61.53 | 58.75 | 62.68 | 56.77 | 53.38 | 59.56 | 51.13 | 52.74 | 51.53 | 46.92 | 55.56 | 43.21 | - |
| Bcl7b | 15.56 | 16.63 | 18.5 | 20.14 | 20.53 | 22.32 | 20.66 | 23.09 | 20.8 | 21.43 | 22.2 | 22.42 | - |
| Pgm3 | 11.25 | 9.73 | 10.79 | 9.66 | 10.03 | 9.18 | 7.47 | 9.7 | 9.75 | 6.31 | 6.68 | 7.24 | - |
| Npepl1 | 13.5 | 13.76 | 14.39 | 8.92 | 11.37 | 8.82 | 8.93 | 9.89 | 8.67 | 7.94 | 7.05 | 7.56 | - |
| Zdhhc20 | 19.46 | 20.41 | 19.85 | 22.34 | 21.66 | 22.42 | 25.42 | 23.19 | 23.82 | 24.18 | 25.49 | 24.18 | - |
| BC003331 | 10.39 | 8.78 | 10.73 | 12.04 | 12.48 | 12.47 | 11.76 | 11.87 | 12.36 | 13.83 | 15.38 | 14.3 | - |
| Cyp4f16 | 4.08 | 2.87 | 2.17 | 2.59 | 3.69 | 1.79 | 4.05 | 2.3 | 3.44 | 1.45 | 0.68 | 1.6 | - |
| Neurl3 | 8.47 | 9.1 | 7 | 21.81 | 11.6 | 14.54 | 14.77 | 11.55 | 11.04 | 21.01 | 15.15 | 18.06 | - |
| BC026590 | 8.06 | 7.73 | 8.47 | 6.73 | 6.26 | 6.05 | 5.59 | 5.96 | 6.12 | 5.04 | 4.79 | 5.65 | - |
| Mcm2 | 77.27 | 95.98 | 82.34 | 40.53 | 45.3 | 41.27 | 29.64 | 37.7 | 33.08 | 20.49 | 15.8 | 22.59 | - |
| Lyrm1 | 18.08 | 17.86 | 17.08 | 9.16 | 9.05 | 7.95 | 11.87 | 13.53 | 16.31 | 8.56 | 6.71 | 7.54 | - |
| Nfrkb | 6.56 | 7.08 | 6.86 | 9.06 | 9.86 | 9.16 | 7.18 | 7.89 | 8.31 | 8.72 | 9.4 | 9.94 | - |
| Carhsp1 | 14.53 | 13.32 | 13.04 | 16.08 | 17.51 | 12.49 | 21.54 | 24.68 | 21.56 | 23.32 | 23.64 | 20.68 | - |
| Fam216a | 51.96 | 56.29 | 56.76 | 46.42 | 49.67 | 50.28 | 51.09 | 45.53 | 46.83 | 39.35 | 44.95 | 39.64 | - |
| Id1 | 0.25 | 0 | 0.12 | 0.56 | 0.75 | 0.91 | 0.32 | 0.58 | 0.35 | 0.82 | 0.81 | 1.33 | - |
| Slc19a2 | 3.33 | 3.54 | 4.67 | 5.22 | 5.78 | 4.84 | 5.15 | 6.14 | 6.29 | 6.64 | 7.73 | 5.11 | - |
| Zfp629 | 0.31 | 0.29 | 0.21 | 0.95 | 0.71 | 0.68 | 0.51 | 0.63 | 0.52 | 1.66 | 0.71 | 1.28 | - |
| Baz1b | 12.44 | 11.65 | 13.67 | 10.64 | 10.78 | 10.12 | 8.7 | 10.16 | 11.21 | 10.22 | 9.69 | 9.04 | - |
| Cdh15 | 0.28 | 0.59 | 0.87 | 0.29 | 0.42 | 0.16 | 0.38 | 0.14 | 0.08 | 0.04 | 0.11 | 0.08 | - |
| Rgl2 | 3.55 | 3.56 | 3.4 | 5.44 | 5.07 | 5.12 | 4.54 | 4.08 | 4.42 | 7.37 | 6.08 | 4.94 | - |
| Slx1b | 2.58 | 3.28 | 3.09 | 5.53 | 4.57 | 4.91 | 5.14 | 3.92 | 4.59 | 5.05 | 5.48 | 7 | - |
| Itgam | 0.02 | 0.1 | 0 | 0.84 | 0.35 | 0.12 | 0.73 | 0.18 | 0.18 | 2.31 | 1.65 | 2.17 | - |
| Coa3 | 104.35 | 102.43 | 100.32 | 68.05 | 73.55 | 84.84 | 71.75 | 73.54 | 76.59 | 60.94 | 53.99 | 64.7 | - |
| Parpbp | 9.4 | 7.68 | 7.78 | 7.98 | 10.52 | 8.82 | 5.3 | 4.34 | 4.98 | 3.91 | 5.95 | 4.46 | - |
| Alkbh3 | 30.98 | 28.83 | 30.41 | 25.23 | 31.44 | 25.7 | 23.55 | 26.76 | 25.5 | 19.7 | 18.57 | 21.45 | - |
| 3110056K07Rik | 1.36 | 1.89 | 1.57 | 3.95 | 3.02 | 4.65 | 4.3 | 4.13 | 3.21 | 6.04 | 5.09 | 6.03 | - |
| Tubd1 | 6.94 | 5.8 | 7.15 | 5.15 | 5.89 | 6.08 | 5.11 | 4.22 | 4.32 | 3.38 | 3.57 | 3.01 | - |
| Rps27 | 586.67 | 650.29 | 981.12 | 1607.29 | 1772.87 | 1024.65 | 1152.76 | 1115.82 | 1586.74 | 1640.85 | 2014.27 | 1313.45 | - |
| Ap1m2 | 0.89 | 0.94 | 1.34 | 2.08 | 1.66 | 2.03 | 2.99 | 1.97 | 1.77 | 1.39 | 2.42 | 3.43 | - |
| Zdhhc17 | 0.88 | 0.65 | 0.67 | 1.38 | 1.07 | 1.02 | 1 | 0.65 | 0.97 | 1.5 | 1.27 | 1.75 | - |
| Dnm3 | 0.19 | 0.3 | 0.29 | 0.28 | 0.45 | 0.46 | 1.21 | 0.68 | 0.7 | 1.41 | 0.89 | 1.72 | - |
| Acat1 | 46.83 | 45.25 | 46.27 | 31.14 | 32.87 | 32.16 | 40.26 | 39.98 | 39.22 | 28.03 | 26.86 | 29.96 | - |
| Dhrs3 | 1.3 | 1.53 | 1.39 | 0.93 | 0.84 | 1.37 | 8.46 | 4.54 | 5.66 | 4.07 | 4.88 | 4 | - |
| Chm | 8.43 | 8.42 | 11.6 | 16.29 | 15.41 | 16.47 | 11.98 | 11.94 | 12.12 | 14.35 | 16.09 | 11.62 | - |
| Tpi1 | 928.94 | 971.81 | 992.14 | 832.69 | 836.05 | 857.38 | 1033.33 | 888.1 | 928.29 | 754.9 | 855.99 | 880.58 | - |
| Pqlc1 | 7.57 | 6.22 | 7.31 | 10.78 | 9.54 | 9.75 | 13.32 | 13.95 | 13.82 | 14.8 | 14.51 | 16.93 | - |
| Pkm | 1373.61 | 1381.94 | 1418.39 | 1334.18 | 1264.7 | 1322.9 | 1877.74 | 1372.82 | 1526.38 | 1579.92 | 1671.9 | 1749.28 | - |
| Pprc1 | 15.93 | 18.16 | 19.39 | 13.38 | 13.07 | 13.13 | 9.48 | 12.96 | 11.97 | 9.92 | 10.78 | 8.17 | - |
| Arap1 | 8.74 | 8.07 | 7.21 | 11.14 | 10.68 | 10.59 | 13.7 | 10.98 | 11.85 | 13.97 | 11.61 | 14.54 | - |
| Zfp275 | 4.01 | 3.96 | 3.63 | 4.43 | 4.66 | 5.8 | 3.72 | 3.75 | 4.05 | 4.85 | 5.53 | 5.53 | - |
| Slc29a3 | 6.17 | 5.2 | 5.82 | 4.11 | 4.01 | 4.13 | 3.57 | 5.42 | 4.59 | 3.84 | 4.5 | 3.69 | - |
| Arntl | 2.08 | 2.3 | 2.01 | 2.5 | 3.42 | 3.34 | 3.27 | 2.08 | 2.26 | 4.25 | 4.47 | 4.7 | - |
| BC021614 | 19.62 | 16.72 | 16.58 | 23.69 | 24.32 | 23.9 | 45.62 | 31.15 | 35.13 | 36.7 | 39.44 | 35.4 | - |
| Atp5g1 | 230.82 | 234.5 | 236.97 | 131.53 | 148.19 | 146.87 | 130.12 | 172.45 | 171.29 | 96.82 | 108.3 | 100.08 | - |
| Serinc3 | 12.41 | 13.69 | 12.44 | 24.06 | 20.61 | 19.3 | 11.76 | 10.31 | 11.17 | 22.24 | 16.98 | 21.55 | - |
| Mthfd2 | 150.29 | 150.74 | 151.09 | 100.3 | 113.3 | 109.47 | 116.32 | 170.13 | 154.04 | 76.52 | 78.66 | 68.94 | - |
| Commd10 | 41.69 | 38.14 | 35.77 | 30.96 | 35.21 | 33.61 | 33.43 | 38.45 | 39.64 | 27.11 | 28.8 | 25.69 | - |
| Per2 | 0.72 | 0.71 | 0.71 | 0.1 | 0.3 | 0.44 | 0.33 | 0.51 | 0.48 | 0.07 | 0.25 | 0.05 | - |
| Zfp36 | 3.85 | 5.01 | 4.32 | 9.35 | 7.75 | 9.36 | 10.74 | 7.55 | 7.87 | 13.85 | 13.51 | 14.44 | - |
| Tmem140 | 2.06 | 1.53 | 1.94 | 3.75 | 3.27 | 4.18 | 3.63 | 3.01 | 2.86 | 7.62 | 4.63 | 7.43 | - |
| Gm16515 | 2.59 | 1.82 | 2.41 | 4.36 | 4.46 | 4.35 | 5.21 | 3.3 | 3.48 | 7.03 | 5.35 | 7.18 | - |
| Gimap1 | 80.37 | 82.99 | 79.94 | 157.26 | 128.85 | 136.44 | 182.64 | 125.86 | 140.19 | 221.35 | 202.15 | 240.22 | - |
| Pik3cg | 8.1 | 8.27 | 8.92 | 15.77 | 15.29 | 15.08 | 7.94 | 4.93 | 6.98 | 11.77 | 9.89 | 10.84 | - |
| Fcho2 | 0.95 | 0.85 | 1.1 | 1.9 | 1.38 | 1.51 | 2.16 | 1.07 | 1.73 | 2.21 | 2.29 | 1.93 | - |
| Tbc1d10c | 98.63 | 101 | 86.52 | 119.83 | 103.39 | 104.25 | 146.71 | 109.1 | 127.29 | 149.62 | 126.79 | 159.6 | - |
| Ablim1 | 35.69 | 36.41 | 36.33 | 63.09 | 55.97 | 59.41 | 58.29 | 45.49 | 53.24 | 74.2 | 69 | 71.05 | - |
| Chn2 | 0.02 | 0.04 | 0 | 0.14 | 0.22 | 0.16 | 0.2 | 0.11 | 0.15 | 0.4 | 0.29 | 0.24 | - |
| Dok4 | 0.18 | 0.6 | 0.21 | 1.48 | 0.72 | 0.54 | 2.17 | 1.06 | 1.18 | 2.11 | 1.68 | 1.56 | - |
| Dcaf12 | 31.2 | 29.46 | 30.12 | 37.47 | 32.86 | 33.22 | 39.79 | 33.55 | 35.63 | 35.48 | 33.9 | 35.91 | - |
| Taok1 | 1.88 | 2.19 | 2.38 | 2.67 | 2.95 | 3.12 | 2.56 | 2.29 | 2.78 | 3.08 | 3.55 | 2.69 | - |
| Mettl3 | 16 | 15.73 | 16.01 | 14.96 | 14.45 | 13.69 | 10.92 | 13.86 | 14.18 | 11.94 | 11.09 | 12.03 | - |
| Tlr6 | 0.81 | 0.63 | 0.37 | 1.95 | 1.61 | 0.61 | 3.04 | 1.5 | 1.49 | 1.34 | 1.37 | 2.21 | - |
| Clk1 | 42.42 | 49.04 | 45.01 | 89.88 | 89.02 | 92.16 | 92.16 | 85.27 | 84.58 | 142.72 | 161.13 | 150.1 | - |
| Kcnn4 | 83.96 | 95.53 | 104.7 | 106.49 | 99.7 | 112.15 | 108.6 | 103.6 | 100.99 | 120.7 | 102.07 | 121.36 | - |
| Itga11 | 0.08 | 0 | 0.01 | 0.18 | 0.32 | 0.23 | 0.31 | 0.5 | 0.39 | 0.56 | 1.19 | 0.7 | - |
| Cul4a | 18.8 | 18.8 | 19.69 | 17.04 | 18.19 | 16.21 | 16.05 | 17.72 | 17.29 | 14.8 | 15.52 | 12.67 | - |
| Ccny | 2.3 | 3.65 | 4.33 | 4.28 | 4.58 | 3.84 | 5.09 | 5.1 | 5.3 | 6.04 | 6.01 | 4.88 | - |
| Atf6 | 3.67 | 3.88 | 3.53 | 4.82 | 4.64 | 5 | 4.7 | 5.89 | 4.84 | 6.17 | 5.9 | 5.84 | - |
| Eprs | 75.01 | 74.13 | 82.01 | 64.86 | 74.98 | 67.97 | 57.12 | 82.66 | 78.47 | 46.44 | 53.5 | 44.95 | - |
| Med20 | 15 | 17.57 | 17.86 | 12.98 | 12.4 | 12.82 | 13.77 | 12.27 | 12.53 | 9.79 | 10.03 | 12.32 | - |
| Net1 | 18.44 | 18.38 | 19.81 | 14.01 | 16.96 | 15.29 | 13.1 | 16.99 | 15.16 | 11.01 | 16.07 | 14.24 | - |
| Depdc5 | 3.47 | 3.43 | 3.63 | 2.53 | 3.88 | 2.85 | 2.48 | 2.68 | 2.92 | 2.02 | 1.99 | 2.52 | - |
| Ehmt2 | 15.46 | 15.13 | 15.05 | 16.8 | 19.03 | 20.67 | 21.68 | 18.49 | 18.59 | 23.49 | 22.17 | 22.09 | - |
| Brip1 | 3.75 | 3.29 | 4.27 | 1.94 | 3.51 | 2.57 | 1.95 | 2.39 | 1.88 | 1.89 | 2.31 | 1.65 | - |
| 1810026B05Rik | 1.65 | 1.77 | 2.1 | 3.4 | 3.18 | 2.7 | 3.18 | 3.1 | 3.16 | 3.67 | 4.3 | 4.08 | - |
| Ube2r2 | 4.14 | 3.95 | 3.74 | 5.21 | 6.65 | 4.97 | 4.54 | 4.08 | 4.39 | 6.03 | 4.88 | 5.55 | - |
| F630111L10Rik | 0.28 | 0.26 | 0.36 | 0.57 | 0.47 | 0.46 | 0.64 | 0.38 | 0.51 | 0.62 | 0.64 | 0.51 | - |
| Ugt1a7c | 10.26 | 8.7 | 9.16 | 2.98 | 4.73 | 3.72 | 3.55 | 9.01 | 7.16 | 2.36 | 1.7 | 2.15 | - |
| Il16 | 9.54 | 10.73 | 7.84 | 16.8 | 13.48 | 13.7 | 18.06 | 12.16 | 13.57 | 16.13 | 13.29 | 15.74 | - |
| Atg9b | 3.92 | 4.4 | 4.41 | 4.05 | 3.31 | 3.15 | 8.75 | 5.08 | 5.43 | 5.78 | 6.96 | 8.42 | - |
| Tmsb15l | 2.45 | 2.41 | 2.24 | 5.34 | 5.13 | 3.59 | 6.88 | 4.34 | 5.52 | 5.66 | 9.25 | 9.81 | - |
| Cyfip1 | 12.8 | 10.95 | 12.39 | 8.81 | 10.06 | 9.67 | 7.77 | 9.38 | 10.3 | 6.61 | 5.58 | 6.81 | - |
| Sec31a | 20.47 | 23.63 | 22.91 | 24.98 | 24.3 | 25.22 | 30.2 | 28.13 | 30.05 | 28.55 | 31.28 | 28.45 | - |
| Cep350 | 1.35 | 1.17 | 1.25 | 1.68 | 1.78 | 1.99 | 1.7 | 1.77 | 1.71 | 2.62 | 2.38 | 2.1 | - |
| Ncapg2 | 7.63 | 7.8 | 8.45 | 4.59 | 5.15 | 4.14 | 3.46 | 4.54 | 3.51 | 2.1 | 3.51 | 2.76 | - |
| Foxp1 | 9.95 | 10.28 | 10.42 | 14.04 | 13.5 | 14.33 | 9.13 | 11.43 | 10.28 | 14.48 | 13.64 | 14.53 | - |
| Sdr42e1 | 1.35 | 0.94 | 0.84 | 2.18 | 1.67 | 1.71 | 1.86 | 1.22 | 1.31 | 2.83 | 1.37 | 1.91 | - |
| Bcas3 | 3.87 | 3.94 | 4.25 | 6.49 | 7.72 | 6.69 | 5.69 | 6.6 | 5.84 | 8.04 | 7.83 | 9.12 | - |
| Apobec3 | 146.95 | 142.14 | 139.77 | 187.27 | 195.45 | 187.51 | 149.83 | 144.24 | 136 | 196.17 | 190.93 | 201.58 | - |
| Stat2 | 15.39 | 14.22 | 15.73 | 22.19 | 26.28 | 26.07 | 16.9 | 16.95 | 16.2 | 23.81 | 23.14 | 22.13 | - |
| Tha1 | 1.8 | 1.64 | 0.88 | 5.84 | 4.07 | 4.12 | 4.31 | 3.48 | 3.47 | 8.64 | 5.36 | 9.43 | - |
| Casp8 | 69.75 | 71.52 | 69.75 | 57.06 | 55.09 | 55.98 | 59.69 | 59.89 | 66.83 | 48.91 | 43.15 | 46.51 | - |
| Nudcd1 | 27.12 | 27.58 | 28.37 | 18.96 | 19.7 | 19.97 | 15.16 | 19.46 | 19.6 | 15.3 | 14.64 | 13.96 | - |
| Lrrc57 | 12.26 | 11.56 | 11.61 | 9.4 | 9.93 | 9.82 | 8.83 | 9.5 | 8.93 | 6.54 | 8.62 | 7.85 | - |
| Gtf2ird2 | 2.02 | 2.03 | 2.42 | 4.68 | 4.41 | 3.88 | 2.87 | 2.7 | 3 | 4.32 | 4.37 | 4.37 | - |
| Kif21b | 3.99 | 4.03 | 3.89 | 7.76 | 7.17 | 6.88 | 11.87 | 7.17 | 8.48 | 13.87 | 13.03 | 12.41 | - |
| Ddx58 | 6.47 | 6.32 | 5.62 | 7.6 | 7.17 | 7.06 | 7.87 | 5.8 | 5.99 | 10.16 | 7.38 | 8.3 | - |
| Tmed2 | 237.87 | 285.49 | 313.79 | 230.74 | 233.85 | 234.06 | 226.09 | 277.13 | 263.48 | 242.91 | 247.11 | 196.59 | - |
| Tapbp | 99.04 | 111.68 | 98.21 | 139.61 | 129.58 | 128.18 | 140.04 | 109.33 | 114.38 | 157.43 | 145.93 | 165.09 | - |
| Jam2 | 0.9 | 0.75 | 1.26 | 0.51 | 0.53 | 0.62 | 0.52 | 0.73 | 0.87 | 0.33 | 0.26 | 0.3 | - |
| Ndufb8 | 299.6 | 277.13 | 278.03 | 165.56 | 190.02 | 199.97 | 179.39 | 218.35 | 210.42 | 142.64 | 143.26 | 145.81 | - |
| Acaa1a | 52.37 | 48.93 | 52.53 | 71.65 | 69.57 | 68.33 | 66.41 | 61 | 60.21 | 70.25 | 70.31 | 74.72 | - |
| Tcerg1 | 21.36 | 28.38 | 32.05 | 24.48 | 26.43 | 24.16 | 21.33 | 27.54 | 25.45 | 20.72 | 22.32 | 16.95 | - |

| | | | | | | | | | | | | | |
|---|---|---|---|---|---|---|---|---|---|---|---|---|---|
| Ckap2l | 24.51 | 18.86 | 18.37 | 19.02 | 25.84 | 24.62 | 13.36 | 12.51 | 11.14 | 14.1 | 15.19 | 12.31 | - |
| Dtl | 22.11 | 20.16 | 22.02 | 11.79 | 13.3 | 16.01 | 8.83 | 10.15 | 9.58 | 6.17 | 7.68 | 5.95 | - |
| Zfp868 | 7.14 | 8.53 | 7.86 | 11.09 | 9.23 | 8.78 | 9.78 | 10.11 | 10.62 | 13.41 | 13.62 | 13.89 | - |
| Prtn3 | 0.41 | 0.42 | 0.72 | 0.65 | 0.51 | 0.66 | 3.79 | 2.85 | 3.76 | 1.47 | 3.24 | 2.38 | - |
| Vrk3 | 18.64 | 23.22 | 21.57 | 31.51 | 29.25 | 33.2 | 36.6 | 32.75 | 32.34 | 40.62 | 34.96 | 39.19 | - |
| Ska3 | 15.91 | 13.05 | 13.89 | 8.92 | 13.31 | 13.1 | 6.53 | 8.34 | 7.98 | 8.7 | 7.83 | 8.12 | - |
| Zfp622 | 2.2 | 2.33 | 2.51 | 3.59 | 3.23 | 4.02 | 4.33 | 5.74 | 4.27 | 6.3 | 6.68 | 4.87 | - |
| Zfp488 | 22.81 | 24.37 | 25.92 | 45.02 | 39.86 | 35.65 | 34.98 | 35.37 | 31.98 | 46.01 | 44.51 | 45.49 | - |
| Ufsp1 | 3.11 | 1.64 | 2.39 | 0.76 | 1.13 | 1.85 | 0.52 | 1.42 | 1.29 | 1.15 | 0.25 | 0.41 | - |
| Irgm2 | 6.89 | 9.72 | 6.63 | 9.75 | 8.43 | 7.49 | 9.1 | 7.35 | 6.94 | 13.91 | 9.41 | 11.44 | - |
| Cd63 | 24.83 | 14.24 | 16.42 | 19.97 | 23.12 | 23.62 | 7.41 | 12.22 | 10.8 | 9.04 | 9.19 | 8.27 | - |
| Reps1 | 2.57 | 2.6 | 2.56 | 4.92 | 3.27 | 3.7 | 3.73 | 3.41 | 3.69 | 3.89 | 4.47 | 4.38 | - |
| Ldlr | 23.35 | 25.72 | 25.09 | 23.99 | 21.44 | 22.92 | 18.58 | 20.17 | 20.39 | 19.78 | 16.73 | 17.87 | - |
| Pcyox1 | 7.54 | 7.49 | 9.03 | 10.9 | 9.63 | 11.7 | 13.16 | 10.13 | 9.33 | 13.86 | 11.69 | 12.34 | - |
| Tinf2 | 25.52 | 24.24 | 25.57 | 24.62 | 32.61 | 30.16 | 18.09 | 25.78 | 18.83 | 17.91 | 20.92 | 19.1 | - |
| Thsd7b | 0.01 | 0.02 | 0 | 0.01 | 0.04 | 0.06 | 0 | 0 | 0 | 0.16 | 0.03 | 0.18 | - |
| Nudt13 | 2.69 | 2.47 | 2.19 | 2.33 | 2.93 | 3.36 | 3.42 | 4.28 | 3.67 | 4.31 | 3.86 | 4 | - |
| Prorsd1 | 7.51 | 7.75 | 7.8 | 10.01 | 10.34 | 10.23 | 12.23 | 10.16 | 10.88 | 13.38 | 12.53 | 12.8 | - |
| Naa40 | 28.08 | 27.04 | 28.05 | 19.67 | 20.92 | 22.7 | 18.86 | 23.35 | 23.02 | 16.35 | 17.58 | 21.7 | - |
| Deptor | 0.07 | 0.07 | 0.2 | 0.73 | 0.45 | 0.56 | 0.16 | 0.18 | 0.15 | 0.97 | 0.77 | 1.07 | - |
| Pecr | 18.46 | 17.63 | 16.01 | 11.04 | 16.17 | 17.33 | 13.38 | 16.21 | 16.98 | 10.86 | 9.62 | 11.92 | - |
| 0610007P14Rik | 130.56 | 109.93 | 117.74 | 73.39 | 83.6 | 86.17 | 62.36 | 94.62 | 87.06 | 52.04 | 48.17 | 47.18 | - |
| Hist2h2aa1 | 3 | 1.54 | 2.8 | 2.49 | 3.37 | 5.07 | 7.12 | 4.65 | 5.14 | 8.23 | 6.42 | 8.39 | - |
| Prr5l | 1.62 | 1.25 | 0.85 | 0.97 | 0.87 | 0.66 | 1.76 | 1.26 | 1.02 | 0.64 | 0.47 | 0.75 | - |
| Lgmn | 2.19 | 1.89 | 2.22 | 1.05 | 2.17 | 0.88 | 2.27 | 1.94 | 2.38 | 0.95 | 0.62 | 0.46 | - |
| Wdr95 | 0.07 | 0.19 | 0 | 0.18 | 0.08 | 0.08 | 1.07 | 0.14 | 0.41 | 1.43 | 0.42 | 1.42 | - |
| Dleu7 | 0.94 | 0.96 | 1.87 | 0.52 | 1.12 | 0.81 | 0.91 | 1.15 | 0.81 | 0.39 | 0.19 | 0.04 | - |
| Herc1 | 3.6 | 3.68 | 3.82 | 4.06 | 4.63 | 4.49 | 4.24 | 4.29 | 4.38 | 4.41 | 5.17 | 4.39 | - |
| Rab23 | 3.07 | 3.34 | 2.74 | 2.41 | 2.44 | 2.03 | 1.66 | 1.75 | 1.11 | 1.68 | 1.64 | | - |
| Adat2 | 6.63 | 8.46 | 6.83 | 4.43 | 5.28 | 5.13 | 5.36 | 7.04 | 6.7 | 4.19 | 4.13 | 4.29 | - |
| Hdac11 | 0.21 | 0.12 | 0.07 | 0.23 | 0.55 | 0.61 | 1.06 | 0.57 | 0.22 | 0.99 | 1.09 | 0.6 | - |
| Aldoa | 1506.33 | 1523.44 | 1535.12 | 2514.02 | 2238.78 | 2473.45 | 3843.04 | 2569.82 | 3018.93 | 3519.69 | 3921.94 | 3814.1 | - |
| Snupn | 20.65 | 18.92 | 20.73 | 16.91 | 16.38 | 17.83 | 20.01 | 16.02 | 18.53 | 13.05 | 15.34 | 15.16 | - |
| Amd2 | 47.77 | 66.65 | 55.32 | 31.92 | 1.24 | 29.98 | 13.67 | 13.49 | 31.1 | 28.09 | 22.62 | 9.1 | - |
| Opn3 | 0.8 | 1.12 | 0.85 | 0.4 | 0.4 | 0.48 | 0.37 | 0.24 | 0.25 | 0.33 | 0.29 | 0.13 | - |
| Apol7b | 0.77 | 1.14 | 0.75 | 1.48 | 0.87 | 0.73 | 3.72 | 0.81 | 1.54 | 3.81 | 1.38 | 3.87 | - |
| D630029K05Rik | 0.08 | 0.03 | 0.05 | 0.15 | 0.12 | 0.39 | 0.4 | 0.17 | 0.3 | 0.67 | 0.09 | 0.49 | - |
| Akap7 | 0.2 | 0.45 | 0.61 | 0.87 | 0.56 | 0.74 | 1.51 | 1.94 | 2.16 | 3.53 | 2.39 | 3.05 | - |
| Tmc6 | 17.04 | 16.76 | 15.78 | 23.91 | 20.11 | 21.37 | 30.31 | 24.2 | 27.87 | 29.74 | 29.85 | 29.82 | - |
| Parp10 | 8.04 | 9.54 | 7.36 | 11.01 | 9.91 | 9.32 | 11.92 | 7.4 | 9.88 | 12.17 | 10.22 | 15.6 | - |
| Plcg1 | 7.37 | 8.71 | 10.23 | 17.6 | 15.49 | 13.6 | 18.11 | 13.25 | 16.63 | 21.06 | 18.76 | 20.37 | - |
| Sec14l1 | 7.69 | 6.6 | 7.18 | 11.34 | 9.24 | 9.63 | 7.61 | 8.08 | 6.27 | 9.52 | 11.32 | 8.24 | - |
| Bhlhb9 | 15.2 | 15.52 | 16.53 | 13.79 | 14.06 | 11.83 | 10.52 | 13.17 | 10.84 | 10.34 | 12.68 | 9.87 | - |
| Spns2 | 0.02 | 0.04 | 0 | 0.09 | 0 | 0 | 0.4 | 0.1 | 0.33 | 0.53 | 0.85 | 0.57 | - |
| Cul1 | 19 | 25.52 | 24.99 | 17.24 | 18.29 | 16.1 | 12.3 | 17.49 | 15.92 | 15.31 | 14.11 | 12.05 | - |
| Tnf | 66.18 | 80.41 | 74.58 | 10.2 | 11.54 | 13.87 | 21.59 | 23.09 | 22.68 | 7.28 | 3.12 | 4.06 | - |
| Rpap2 | 17.6 | 16.31 | 17.08 | 19.69 | 19.62 | 20.38 | 23.44 | 21.94 | 21.37 | 21.84 | 28.54 | 25.77 | - |
| Ropn1l | 1.69 | 1.53 | 1.78 | 2.78 | 3.22 | 2.38 | 3.85 | 3.74 | 2.95 | 5.78 | 3.12 | 3.99 | - |
| Med30 | 50.24 | 72.55 | 76.6 | 42.89 | 45.73 | 44.66 | 43.55 | 45.86 | 43.12 | 34.57 | 35.1 | 34.15 | - |
| Lrch1 | 3.92 | 4.82 | 4.85 | 7.4 | 5.86 | 6.51 | 5.51 | 6.08 | 5.39 | 6.72 | 7.32 | 5.93 | - |
| Adpgk | 22.99 | 22.88 | 24.96 | 15.52 | 15.15 | 15.64 | 9.5 | 16.53 | 14.27 | 8.84 | 8.33 | 7.28 | - |
| Ywhag | 47 | 42.8 | 45.9 | 29.42 | 33.47 | 31.33 | 29.95 | 37.47 | 32.91 | 25.92 | 28.14 | 24.23 | - |
| Ulbp1 | 6.36 | 4.92 | 3.35 | 2.93 | 3.99 | 2.64 | 5.43 | 6.18 | 4.47 | 2.19 | 2.12 | 2.2 | - |
| Hpdl | 5.02 | 5.99 | 6.11 | 2.13 | 1.4 | 1.71 | 1.83 | 1.76 | 2.24 | 1.25 | 0.81 | 0.58 | - |
| Tbcd | 25.48 | 20.81 | 22.25 | 18.12 | 16.73 | 17.91 | 19.2 | 20.55 | 21.34 | 16.41 | 16.54 | 16.78 | - |
| Smad2 | 6.93 | 7.24 | 6.82 | 5.74 | 5.97 | 6.74 | 5 | 5.74 | 5.51 | 5.01 | 5 | 5.39 | - |
| Rasgef1a | 0.19 | 0.04 | 0.07 | 0.35 | 0.15 | 0.24 | 0.28 | 0.33 | 0.15 | 0.49 | 0.69 | 0.84 | - |
| Tbc1d25 | 8.62 | 8.01 | 9.72 | 6.43 | 7.39 | 7.18 | 6.99 | 8.3 | 7.93 | 6.71 | 4.69 | 6.49 | - |
| Rhod | 1.45 | 1.17 | 1.05 | 2.13 | 3.59 | 2.81 | 2.96 | 1.93 | 2.09 | 5.99 | 3.68 | 3.45 | - |
| Polr3k | 35.31 | 38.5 | 34.56 | 26.25 | 24.13 | 24.17 | 36.23 | 31.18 | 34.12 | 27.16 | 23.7 | 26.6 | - |
| Csrp2bp | 6.45 | 6.46 | 6.33 | 4.31 | 3.71 | 4.41 | 3.55 | 4.35 | 5.31 | 4.17 | 3.01 | 4.07 | - |
| Dpf1 | 0.15 | 0.64 | 0.31 | 1.05 | 1.07 | 0.95 | 1.46 | 0.64 | 1.08 | 1.73 | 2.08 | 2.73 | - |
| Osm | 57.01 | 61.21 | 61.61 | 33.25 | 32.99 | 31.71 | 17.09 | 23.04 | 28.42 | 8.94 | 9.03 | 10.13 | - |
| Baiap3 | 0.14 | 0.29 | 0 | 1.64 | 0.5 | 0.73 | 3.57 | 0.42 | 0.61 | 5.81 | 2.4 | 6.28 | - |
| Dynll2 | 4.27 | 2.98 | 3.23 | 2.43 | 1.82 | 2.37 | 1.67 | 1.76 | 1.99 | 1.64 | 1.07 | 1.26 | - |
| D630039A03Rik | 5.18 | 5.54 | 4.41 | 8.47 | 4.29 | 4.98 | 4.25 | 2.6 | 3.98 | 1.24 | 2.04 | 2.63 | - |
| Ccdc12 | 60.64 | 54.14 | 59.93 | 45.5 | 50.11 | 46.53 | 56.76 | 53.87 | 56.98 | 46.06 | 46.46 | 49.08 | - |
| 8430410A17Rik | 7.26 | 7.9 | 6.92 | 7.41 | 6.39 | 7.52 | 11.72 | 11.19 | 7.93 | 11.1 | 11.96 | 12 | - |
| BC021767 | 0 | 0 | 0 | 0.06 | 0.11 | 0.11 | 0.47 | 0.18 | 0.19 | 0.15 | 0.08 | 0.31 | - |
| Cd36 | 10.43 | 8.63 | 6.83 | 3.74 | 3.12 | 5.44 | 0.67 | 2.12 | 1.39 | 0.15 | 0.19 | 0.52 | - |
| Fam168b | 7.95 | 7.56 | 7.14 | 10.86 | 8.81 | 9.67 | 10.03 | 9.17 | 8.67 | 11.38 | 11.55 | 12.09 | - |
| 9130221F21Rik | 0.52 | 0.29 | 0.61 | 1.05 | 1.01 | 0.97 | 1.2 | 0.55 | 1.01 | 1.27 | 1.42 | 1.04 | - |
| Rassf1 | 37.56 | 33.52 | 38.68 | 57.67 | 55.79 | 53.97 | 69.03 | 52.09 | 54.07 | 74.04 | 91.01 | 85.57 | - |
| Btbd10 | 23.72 | 21.27 | 19.3 | 13.84 | 14.31 | 13.69 | 15.74 | 19 | 14.82 | 10.87 | 13 | 10.46 | - |
| Gorasp2 | 94.45 | 94.83 | 98.17 | 73.07 | 79.68 | 76.57 | 81.44 | 98.01 | 90.04 | 73.09 | 80.43 | 74.93 | - |
| Mmp15 | 0.01 | 0 | 0 | 0.13 | 0 | 0.04 | 0.08 | 0.1 | 0.13 | 0.17 | 0.14 | 0.19 | - |
| Serpinb5 | 0 | 0 | 0 | 0.2 | 0.05 | 0 | 0.04 | 0.09 | 0.33 | 0.31 | 0.41 | 1.01 | - |
| Dnahc10 | 0.02 | 0.01 | 0 | 0.28 | 0.32 | 0.29 | 0.02 | 0.01 | 0.03 | 0.14 | 0.14 | 0.07 | - |
| Nsun4 | 8.18 | 7.64 | 8.35 | 7.7 | 7.06 | 9.12 | 5.86 | 8.06 | 5.95 | 4.89 | 4.88 | 4.84 | - |
| Hsf2 | 19.56 | 19.15 | 19.05 | 12.86 | 14.87 | 16.53 | 12.09 | 18.67 | 15.58 | 11.62 | 15.3 | 11.82 | - |
| Stxbp2 | 58.64 | 57.37 | 54.81 | 65.76 | 63.38 | 62.92 | 70.11 | 63.89 | 61.85 | 71.63 | 64.4 | 73.28 | - |
| Ptk2b | 7.53 | 8.27 | 7.49 | 13.6 | 10.48 | 10.83 | 15.41 | 11.8 | 12.19 | 18.36 | 15.16 | 17.28 | - |
| Unc5cl | 4.35 | 4.64 | 4.91 | 2.61 | 2.18 | 1.69 | 1.99 | 2.53 | 2.27 | 1.65 | 0.91 | 1.91 | - |
| Pgp | 26.44 | 34.23 | 33.67 | 20.95 | 19.24 | 18.04 | 31.96 | 20.66 | 23.34 | 14.47 | 18.27 | 19.77 | - |
| Trerf1 | 0.65 | 0.85 | 0.78 | 1.39 | 1.11 | 1.15 | 3.76 | 2.45 | 2.68 | 3.47 | 3.01 | 3.23 | - |
| Taf8 | 7.33 | 6.43 | 6.48 | 7.14 | 8.06 | 9.05 | 9.39 | 7.86 | 8.71 | 8.81 | 10.6 | 9.75 | - |
| Fam160b1 | 1.97 | 1.5 | 1.67 | 2.97 | 3.12 | 2.46 | 2.24 | 2.22 | 2.53 | 2.98 | 3.32 | 3.09 | - |
| Baz2a | 3.28 | 3.23 | 3.1 | 5.3 | 4.82 | 4.95 | 5.34 | 4.58 | 5.21 | 6.3 | 6.56 | 6.36 | - |
| Mapk6 | 25.07 | 27.95 | 29.6 | 26.38 | 24.45 | 22.71 | 21.63 | 27.52 | 27.63 | 20.27 | 24.34 | 20.47 | - |
| Map2k4 | 5.61 | 5.59 | 5.8 | 3.85 | 5.39 | 4.35 | 3.66 | 4.39 | 5.72 | 4.2 | 4.12 | 3.75 | - |
| Zcchc4 | 7.95 | 6.62 | 7.84 | 5.09 | 5.45 | 5.8 | 6.69 | 7.97 | 7 | 4.79 | 4.55 | 6.05 | - |
| Ttc33 | 14.05 | 13.31 | 14.86 | 19.15 | 14.51 | 16.4 | 22.58 | 20.44 | 22.63 | 24.13 | 20.26 | 22.8 | - |
| Cd302 | 0.91 | 1.21 | 1.12 | 0.5 | 0.45 | 0.41 | 0.27 | 0.09 | 0.73 | 0.09 | 0.29 | 0.18 | - |
| Etv6 | 4.1 | 3.8 | 4.67 | 4.49 | 3.47 | 3.62 | 3.45 | 3.8 | 3.54 | 2.16 | 2.17 | 3.09 | - |
| Hist1h1b | 5.18 | 8.05 | 4.59 | 1.69 | 3.72 | 3.61 | 2.57 | 2.19 | 3.19 | 1.79 | 2.29 | 2.14 | - |
| 1500009L16Rik | 39.35 | 45.71 | 40.57 | 24.03 | 22.26 | 25.42 | 59.51 | 42.26 | 53.7 | 25.82 | 29.01 | 31.86 | - |
| Igfbp7 | 1.84 | 1.51 | 2.61 | 1.07 | 1.39 | 1.08 | 0.82 | 1.05 | 0.43 | 0.36 | 0.22 | 0.15 | - |
| Ints10 | 45.75 | 44.88 | 44.24 | 25.88 | 32.06 | 28.73 | 31.67 | 33.82 | 31.72 | 24.73 | 21.25 | 22.63 | - |
| Pdzd11 | 60.86 | 59.11 | 55.83 | 41.16 | 52.09 | 47.19 | 36.74 | 43.28 | 39.96 | 26.66 | 28.11 | 29.93 | - |
| Pcmt1 | 48.2 | 43.52 | 43.69 | 36.5 | 39.99 | 40.61 | 38.14 | 42.47 | 39.22 | 32.58 | 39.4 | 35.39 | - |
| Dgkg | 0.05 | 0.03 | 0.07 | 0.1 | 0.26 | 0.36 | 0.52 | 0.17 | 0.3 | 0.72 | 0.72 | 0.93 | - |
| Kctd17 | 1.7 | 1.59 | 1.75 | 0.96 | 0.83 | 0.69 | 0.8 | 0.78 | 1.06 | 0.55 | 0.47 | 0.81 | - |
| Top2a | 82.49 | 67.21 | 62.28 | 57.92 | 70.24 | 65.77 | 41.61 | 37.25 | 42.67 | 37.5 | 40.67 | 27.84 | - |
| Sepsecs | 6.61 | 6.71 | 6.38 | 3.55 | 4.64 | 3.95 | 2.84 | 4.64 | 5.18 | 3.57 | 4.23 | 2.7 | - |
| Usp28 | 2.29 | 1.96 | 1.3 | 4.31 | 3.54 | 3.4 | 5.13 | 2.49 | 3.11 | 7.16 | 6.57 | 7.57 | - |
| Eaf2 | 0.59 | 0.9 | 0.61 | 2.11 | 1.38 | 1.21 | 1.62 | 1.23 | 1 | 2.02 | 3.32 | 2.03 | - |
| Pgap3 | 0.52 | 0.55 | 0.18 | 1.1 | 0.49 | 0.68 | 1.35 | 0.94 | 1.24 | 1.06 | 1.17 | 1.28 | - |
| Hist1h2bk | 2.33 | 4.33 | 1.07 | 0.48 | 2.13 | 1.8 | 0.44 | 0.43 | 0.72 | 0 | 0 | 0.77 | - |
| Anxa11 | 5.74 | 6.83 | 7.07 | 9.29 | 9.96 | 8.15 | 10.27 | 8.04 | 7.5 | 12.09 | 10.07 | 12.42 | - |
| Pex11b | 14.79 | 15.28 | 13.6 | 18.84 | 18.9 | 19.78 | 17.49 | 17.06 | 17.71 | 18.11 | 18.57 | 21.41 | - |
| Fam53a | 29.56 | 27.48 | 29.42 | 30.36 | 32.66 | 33.18 | 30.82 | 36.54 | 31.37 | 36.16 | 36.96 | 40.35 | - |
| Ppan | 89.2 | 85.88 | 89.73 | 41.04 | 39.86 | 40.99 | 30.33 | 48.79 | 47.04 | 27.9 | 24.9 | 22.25 | - |
| Cblb | 14.84 | 15.01 | 14.93 | 26.02 | 21.26 | 22.81 | 23.97 | 22.24 | 22.45 | 37.26 | 35.69 | 37.07 | - |
| Ppp1r12a | 4.11 | 4.46 | 5.01 | 6.42 | 5.35 | 5.38 | 7.34 | 5.9 | 6.58 | 6.62 | 6.99 | 5.91 | - |
| Bnip3 | 172.5 | 170.69 | 161.31 | 209.26 | 192.96 | 195.27 | 370.07 | 245.82 | 307.15 | 217.12 | 257.31 | 300.38 | - |
| Cdkl3 | 3.11 | 2.97 | 3.69 | 4.1 | 5.48 | 4.75 | 3.65 | 4.63 | 4.74 | 6.47 | 5.91 | 5.04 | - |
| Pdk1 | 18.51 | 23.06 | 20.74 | 27.78 | 24.87 | 23.71 | 39.28 | 23.18 | 28.2 | 31.24 | 28.06 | 37.17 | - |
| Abhd2 | 11.02 | 11.56 | 10.37 | 14.65 | 15.25 | 12.68 | 12.81 | 11.67 | 11.07 | 14.14 | 16.85 | 17.62 | - |
| Gpr114 | 19.59 | 21.99 | 17.03 | 51.11 | 35.97 | 37.56 | 32.6 | 22.19 | 27.75 | 47.94 | 32.55 | 46.18 | - |
| Arhgap25 | 6.73 | 7.01 | 7.15 | 12.55 | 9.81 | 10.22 | 15.33 | 11.34 | 12.97 | 19.79 | 16.71 | 20.46 | - |
| Pou2f1 | 0.8 | 0.93 | 0.79 | 1.16 | 1.33 | 1.4 | 1.11 | 1.27 | 1.16 | 1.67 | 2.04 | 1.55 | - |
| Btf3l4 | 17.16 | 16.74 | 16.62 | 14.68 | 14.77 | 15.74 | 13.81 | 14.36 | 14.54 | 13.86 | 12.53 | 14.21 | - |
| Mecr | 24.21 | 21.8 | 22.24 | 14.13 | 13.82 | 13.69 | 11.47 | 15.33 | 12.88 | 8.94 | 8.1 | 10.69 | - |
| Dedd2 | 12.27 | 12.48 | 14.79 | 12.9 | 15.47 | 14.69 | 28.06 | 20.43 | 21.34 | 24.07 | 21.58 | 27.56 | - |
| A430078G23Rik | 0.67 | 0.8 | 1.24 | 1.6 | 1.47 | 1.64 | 1.9 | 1.27 | 1.23 | 2.82 | 1.88 | 3.79 | - |
| Sgms1 | 3.04 | 3.52 | 2.64 | 9.12 | 10.14 | 10.13 | 3.33 | 2.5 | 2.94 | 9.14 | 7.65 | 8.5 | - |
| Clasp2 | 5.55 | 6.04 | 7.32 | 8.21 | 8.24 | 7.56 | 8.32 | 9.38 | 8.33 | 9.1 | 8.34 | 7.44 | - |
| Ubb | 1611.99 | 1571.17 | 1499.98 | 1876.3 | 1874.43 | 1920.63 | 1872.13 | 1854.31 | 1808.19 | 2106.11 | 2196.99 | 2107.59 | - |
| Mrpl30 | 162.4 | 164.05 | 170.18 | 216.88 | 208.59 | 215.87 | 208.09 | 194.14 | 202.33 | 222.02 | 234.39 | 233.86 | - |
| Nlrc3 | 7.75 | 8.02 | 6.86 | 12.63 | 10.17 | 10.27 | 10.43 | 7.54 | 7.68 | 12.86 | 9.12 | 11.56 | - |
| Btbd6 | 2.94 | 3.11 | 4.62 | 6.23 | 5.61 | 5.09 | 4.99 | 4.28 | 4.85 | 6.74 | 4.92 | 5.35 | - |
| Creb3 | 18.17 | 20.82 | 19.27 | 23.27 | 22.87 | 25.23 | 22.47 | 20.46 | 22.94 | 22.46 | 26.33 | 27.78 | - |
| Rnf123 | 9.71 | 11.56 | 10.13 | 13.51 | 12.96 | 14.62 | 16.34 | 14.34 | 13.34 | 17.16 | 14.91 | 16.69 | - |
| Cd164 | 81.78 | 86.62 | 84.87 | 112.13 | 106.17 | 110.69 | 103.81 | 95.59 | 103.78 | 113.37 | 118.35 | 119.04 | - |
| Csrp2 | 6.81 | 7.07 | 6.81 | 10.02 | 7.51 | 9.11 | 13.25 | 7.73 | 9.61 | 10.94 | 9.08 | 15.3 | - |
| Prdm9 | 1.3 | 0.98 | 1.17 | 1.82 | 1.88 | 2.91 | 1.49 | 1.13 | 1.61 | 2.11 | 2.79 | 1.83 | - |
| Rnf219 | 10.82 | 9.68 | 11.07 | 9.25 | 5.4 | 7.16 | 5.16 | 7.26 | 8.14 | 6.83 | 5.85 | 6.89 | - |
| Fam13b | 1.94 | 1.47 | 1.98 | 3.15 | 2.89 | 2.95 | 4.32 | 3.04 | 2.96 | 3.44 | 4.79 | 3.68 | - |
| Nabp1 | 78.15 | 74.63 | 70.23 | 34.56 | 40.94 | 36.4 | 40.86 | 56.34 | 49.82 | 23.79 | 21.85 | 28.98 | - |
| C330024D21Rik | 1.43 | 1.29 | 1.41 | 1.93 | 1.87 | 2.34 | 2.04 | 2.31 | 1.68 | 2.6 | 2.12 | 2.32 | - |
| F8a | 3.58 | 4.3 | 3.41 | 7.11 | 6.84 | 5.35 | 6.37 | 6.93 | 6.27 | 8.87 | 9.21 | 8.61 | - |
| Xpot | 19.39 | 22.99 | 23.41 | 18.2 | 21.91 | 21.22 | 16.32 | 25.43 | 24.04 | 13.43 | 17.78 | 13.85 | - |
| Dlgap1 | 0.03 | 0.01 | 0 | 0.08 | 0.08 | 0.08 | 0.07 | 0.01 | 0.13 | 0.21 | 0.18 | 0.17 | - |
| 0610031J06Rik | 44.22 | 43.88 | 40.35 | 48.4 | 50.77 | 51.74 | 63.08 | 50.52 | 54.9 | 62.9 | 54.09 | 65.34 | - |
| Unc119b | 15.26 | 18.81 | 23.68 | 37.82 | 35.16 | 30.25 | 49.2 | 37.47 | 41.52 | 58.27 | 59.72 | 56.66 | - |
| Rnf128 | 0.19 | 0.63 | 0.12 | 1.47 | 0.5 | 0.63 | 1.09 | 0.34 | 1.21 | 1.23 | 0.92 | 1.32 | - |
| Zfp719 | 3.4 | 3.68 | 3.26 | 2.81 | 2.79 | 2.97 | 2.46 | 2.51 | 1.98 | 2.19 | 2.57 | 1.22 | - |
| Ercc6l | 4.95 | 4.5 | 5.81 | 2.35 | 3.03 | 2.42 | 1.74 | 1.76 | 1.7 | 0.68 | 0.97 | 1.07 | - |
| 1700023L04Rik | 0.17 | 0 | 0 | 2.5 | 1.14 | 2.81 | 0.52 | 0.98 | 0.16 | 3.48 | 3 | 3.5 | - |
| Ehd4 | 11.74 | 12.39 | 12.44 | 10.42 | 10.19 | 10.02 | 8.84 | 9.78 | 10.26 | 8.04 | 9.7 | 8.15 | - |
| Sae1 | 225.39 | 228.43 | 234.45 | 197.11 | 196.22 | 188.42 | 158.97 | 192.06 | 186.1 | 163.2 | 166.67 | 160.85 | - |

| | | | | | | | | | | | | | |
|---|---|---|---|---|---|---|---|---|---|---|---|---|---|
| Tceal1 | 0.21 | 0 | 0.24 | 0.91 | 0.56 | 0.58 | 1.04 | 0.48 | 1.65 | 0.57 | 1.39 | 1.25 | - |
| Fam65b | 2.39 | 2.64 | 2.54 | 7.2 | 4.32 | 5.2 | 12.3 | 5.17 | 9.21 | 10.92 | 9.66 | 10.06 | - |
| Chml | 0.64 | 0.63 | 0.71 | 0.85 | 0.4 | 0.3 | 0.41 | 0.52 | 0.34 | 0.23 | 0.27 | 0.37 | - |
| Pdik1l | 5.01 | 5.37 | 4.82 | 4.43 | 5.23 | 3.6 | 4.02 | 3.94 | 4.09 | 3.62 | 3.57 | 3.17 | - |
| Pnrc1 | 2.73 | 2.69 | 3.93 | 8.14 | 7.33 | 4.97 | 9.64 | 7.29 | 6.52 | 12.01 | 11.49 | 12.55 | - |
| Sh2d3c | 9.8 | 10.04 | 7.94 | 10.32 | 10.16 | 9.65 | 14.71 | 11.27 | 14.21 | 14.88 | 11.45 | 14.67 | - |
| Sptbn1 | 4.28 | 5.25 | 4.64 | 5.3 | 4.84 | 5.04 | 5.96 | 4.68 | 5.41 | 6.75 | 5.7 | 5.8 | - |
| Atf6b | 14.83 | 17.03 | 17.37 | 10.52 | 10.56 | 9.1 | 9.38 | 12.04 | 10.08 | 8.46 | 7.18 | 7.48 | - |
| Slc1a4 | 10.18 | 8.21 | 9.29 | 6.45 | 8.25 | 9.94 | 6.94 | 15.84 | 11.76 | 6.32 | 6.17 | 6.33 | - |
| Brca1 | 2.86 | 3.36 | 3.56 | 2.27 | 3.27 | 2.94 | 1.94 | 2.16 | 2.14 | 1.51 | 2.18 | 1.21 | - |
| Marveld1 | 4.16 | 4.07 | 4.69 | 3.18 | 3.85 | 5.39 | 3.49 | 3.61 | 3.19 | 2.49 | 1.9 | 2.83 | - |
| Kif13b | 0.58 | 0.58 | 0.58 | 1.12 | 1.03 | 0.63 | 0.86 | 0.82 | 0.76 | 1.29 | 1.22 | 1.08 | - |
| Rasa3 | 5.9 | 6.92 | 4.98 | 10.48 | 7.21 | 5.75 | 15.76 | 9.3 | 12.49 | 16 | 12.24 | 16.35 | - |
| Fxyd5 | 344.76 | 339.33 | 320.79 | 588.66 | 469.5 | 509.77 | 886.31 | 498.31 | 643.34 | 774.96 | 710.17 | 890.99 | - |
| Slc25a37 | 5.5 | 5.63 | 6.59 | 7.73 | 8.52 | 8.17 | 11.49 | 11.25 | 9.62 | 12.93 | 13.51 | 12.77 | - |
| Pias1 | 9.86 | 10.32 | 8.99 | 12.54 | 12.18 | 14.21 | 13.27 | 12.16 | 12.46 | 16.36 | 16.18 | 16.49 | - |
| Gapvd1 | 11.36 | 11.37 | 12.65 | 9.04 | 9.87 | 8.34 | 6.92 | 9.37 | 8.95 | 8.61 | 6.68 | 6.69 | - |
| Bzw2 | 104.34 | 109.11 | 106.71 | 70.76 | 66.39 | 69.15 | 53 | 76.72 | 68.53 | 44.8 | 48.26 | 44.27 | - |
| Hist2h3b | 36.33 | 38.34 | 41.56 | 13.22 | 23.67 | 22.65 | 13.52 | 11.16 | 12.56 | 6.24 | 10.65 | 7.07 | - |
| Scel | 0.32 | 0.21 | 0.12 | 0.04 | 0.47 | 0.33 | 1.71 | 1.04 | 1.25 | 1.03 | 0.89 | 1.49 | - |
| Rhbdf2 | 12.06 | 12.13 | 12.3 | 11.6 | 10.6 | 9.24 | 17.62 | 13.87 | 16.2 | 15.39 | 15.39 | 16.09 | - |
| Dynlt1a | 20.68 | 16.93 | 10.29 | 9.94 | 10.42 | 6.98 | 13.92 | 12.71 | 10.87 | 4.15 | 9.38 | 8.75 | - |
| Zfp334 | 0.27 | 0.33 | 0.27 | 0.23 | 0.07 | 0.11 | 0.01 | 0.07 | 0.06 | 0.09 | 0 | 0.02 | - |
| Sppl2a | 16.19 | 15 | 15.39 | 14.07 | 12.9 | 13.75 | 12.43 | 15.63 | 12.92 | 12 | 12.48 | 12.06 | - |
| Ccnh | 55.41 | 53.66 | 53.21 | 44.14 | 43.22 | 45.11 | 43.29 | 48.51 | 47.72 | 36.77 | 41.83 | 40.14 | - |
| Tes | 113.2 | 119.01 | 121.8 | 94.57 | 104.07 | 94.7 | 104.74 | 129.74 | 122.29 | 84.85 | 93.3 | 85.84 | - |
| Zfp934 | 5.03 | 5.19 | 6.32 | 4.03 | 4.9 | 4.53 | 2.89 | 3.07 | 3.22 | 2.76 | 3.01 | 2.55 | - |
| Exoc6 | 17.44 | 16.22 | 18.12 | 26.5 | 26.37 | 24.29 | 32.12 | 23.88 | 25.84 | 29.58 | 32.83 | 25.57 | - |
| Lamp1 | 7.75 | 8.83 | 11.8 | 15.16 | 15.56 | 10.58 | 20.93 | 28.24 | 30.85 | 27.34 | 18.97 | 15.52 | - |
| Prr14 | 18.29 | 19.13 | 17.57 | 23.7 | 22.65 | 24.07 | 22.1 | 23.57 | 23.95 | 29.76 | 27.76 | 27.85 | - |
| Bahcc1 | 0.02 | 0.03 | 0 | 0.04 | 0.05 | 0.04 | 0.1 | 0.11 | 0.05 | 0.08 | 0.14 | 0.19 | - |
| Romo1 | 125.46 | 107.96 | 100.9 | 72.22 | 78.05 | 73.28 | 83.04 | 91.09 | 82.28 | 53.76 | 62.03 | 62.51 | - |
| Ficd | 2.17 | 2.65 | 3.34 | 1.48 | 2.35 | 2.91 | 2.35 | 2.4 | 2.31 | 2.1 | 1.03 | 1.44 | - |
| Slc39a10 | 5.44 | 4.25 | 6.08 | 2.88 | 2.88 | 3.42 | 2.36 | 3.75 | 4.43 | 1.78 | 3.44 | 2.46 | - |
| Rpl19 | 2391.92 | 2460.82 | 2445.98 | 3276.34 | 3173.31 | 3321.19 | 3219.07 | 3009.96 | 3141.33 | 3533.67 | 3809.24 | 3610.95 | - |
| Tbrg3 | 2.46 | 2.6 | 2.54 | 3.61 | 3.15 | 2.9 | 3.44 | 2.75 | 3.08 | 3.48 | 3.93 | 3.64 | - |
| Rps19bp1 | 37.62 | 43.54 | 40.32 | 32.13 | 32.26 | 32.19 | 25.33 | 33.97 | 36.07 | 28.12 | 32.92 | 23.27 | - |
| Ptplb | 4.61 | 6.47 | 5.93 | 6.34 | 6.99 | 6.14 | 7.26 | 7.74 | 7.7 | 8.6 | 8.15 | 6.59 | - |
| Plcb3 | 1.84 | 1.85 | 2.27 | 3.31 | 3.53 | 4.62 | 3.27 | 2.51 | 3.93 | 4.23 | 3.86 | 4.3 | - |
| Abcd4 | 5.31 | 4.36 | 5.95 | 7 | 7.95 | 5.7 | 7.66 | 7.51 | 6.85 | 9.16 | 8.19 | 8.02 | - |
| Ndufc1 | 143.69 | 148.39 | 141.18 | 89.31 | 106.59 | 97.81 | 95.36 | 94.33 | 107.03 | 69.78 | 62.36 | 68.64 | - |
| Emc4 | 141.4 | 134.63 | 128.88 | 67.08 | 71.34 | 71.19 | 59.96 | 85.79 | 79.2 | 46.56 | 43.89 | 47.01 | - |
| Hist1h2ak | 4.39 | 4.52 | 5.11 | 0.52 | 1.99 | 0.42 | 0.39 | 1.82 | 2.19 | 0.38 | 1.71 | 0.64 | - |
| Zfp942 | 4.49 | 4.71 | 5.51 | 7.82 | 5.7 | 5.62 | 4.57 | 5.54 | 4.92 | 6.95 | 6.63 | 6.98 | - |
| Dirc2 | 0.19 | 0.21 | 0.07 | 0.74 | 0.47 | 0.28 | 0.29 | 0.27 | 0.4 | 0.8 | 0.48 | 0.81 | - |
| 2210408I21Rik | 1.77 | 1.32 | 1.45 | 1.31 | 0.61 | 1.29 | 1.34 | 0.97 | 0.72 | 0.86 | 0.54 | 0.51 | - |
| Ppp1r2 | 21.8 | 24.15 | 25.9 | 17.35 | 19.21 | 19.15 | 14.8 | 17.03 | 16.16 | 15.51 | 13.14 | 15.54 | - |
| Chchd6 | 13.49 | 12.24 | 12.82 | 9.39 | 10 | 11.98 | 9.56 | 10.88 | 11.62 | 8.96 | 6.62 | 8.67 | - |
| Mynn | 6.07 | 5.87 | 6.51 | 8.51 | 8.22 | 7.54 | 7.66 | 6.53 | 7.76 | 8.97 | 8.16 | 8.88 | - |
| Rab12 | 1.25 | 1.79 | 2.21 | 3.33 | 3.05 | 2.86 | 2.77 | 2.37 | 2.08 | 2.76 | 4.26 | 4.39 | - |
| Kxd1 | 57.28 | 57.17 | 58.96 | 105.57 | 88.31 | 105.67 | 111.11 | 97.91 | 99.42 | 143.11 | 126.88 | 145.75 | - |
| Nmb | 2.61 | 3.9 | 2.62 | 3.7 | 3.12 | 5.46 | 6.07 | 3.85 | 6.81 | 5.71 | 6.07 | 6.17 | - |
| Il3ra | 3.47 | 3.34 | 3.1 | 8.8 | 7.03 | 5.67 | 4.95 | 4.38 | 4.16 | 6.57 | 5.08 | 6.66 | - |
| Nudt15 | 1.85 | 2.38 | 1.98 | 1.59 | 1.66 | 1.85 | 0.6 | 1.47 | 0.95 | 1.02 | 0.81 | 0.69 | - |
| Rbms2 | 8.96 | 8.09 | 8.56 | 8.98 | 9.59 | 10.46 | 10.08 | 10.68 | 10.1 | 11.61 | 11.19 | 11.37 | - |
| Lpin3 | 0.82 | 0.48 | 0.38 | 0.28 | 0.26 | 0.09 | 0.52 | 0.54 | 0.47 | 0.17 | 0.07 | 0.19 | - |
| Rbm33 | 2.88 | 3.16 | 2.66 | 5.21 | 4.63 | 5.11 | 5.05 | 4.5 | 4.81 | 7.13 | 7.52 | 6.36 | - |
| Sigirr | 32 | 37.75 | 36.26 | 51.36 | 45.61 | 43.17 | 90.76 | 71.27 | 76.73 | 79.72 | 80.03 | 90.05 | - |
| Hddc2 | 47.4 | 48.54 | 48.25 | 40.14 | 41.9 | 38.12 | 44.26 | 42.37 | 45.81 | 39.09 | 34.84 | 39.82 | - |
| Mrrf | 11.42 | 12.08 | 10.02 | 12.89 | 13.31 | 13.12 | 12.67 | 14.75 | 13.62 | 13.08 | 14.96 | 15.7 | - |
| Rassf6 | 0.06 | 0.03 | 0 | 0.03 | 0.03 | 0.03 | 0.7 | 0.12 | 0.08 | 0.61 | 0.47 | 0.5 | - |
| Zfp189 | 2.35 | 2.55 | 2.46 | 3.48 | 2.69 | 4.47 | 3.44 | 2.85 | 4.09 | 3.53 | 4.16 | 3.77 | - |
| Mettl21a | 7.17 | 6.45 | 6.04 | 6.42 | 7.61 | 7.56 | 8.54 | 6.5 | 7.41 | 8.24 | 8.47 | 11.18 | - |
| Mgat2 | 6.26 | 7.83 | 9.07 | 5.63 | 5.97 | 4.31 | 4.5 | 6.31 | 5.81 | 6.3 | 4.36 | 4.25 | - |
| Klhl28 | 2.92 | 2.49 | 2.71 | 3.5 | 3.63 | 4.04 | 4.39 | 4.34 | 3.67 | 4.61 | 5.79 | 4.89 | - |
| Mir17hg | 7.52 | 8.39 | 8.81 | 6.76 | 4.62 | 5.37 | 3.41 | 5.35 | 4.7 | 4.01 | 2.42 | 3.45 | - |
| Cant1 | 9.77 | 8.86 | 12.06 | 12.91 | 12.77 | 14.98 | 13.3 | 12.49 | 11.6 | 16.11 | 12.42 | 15.25 | - |
| LOC100503496 | 0 | 0.25 | 0 | 0.17 | 0.51 | 0.31 | 0.32 | 0.18 | 0.28 | 0.23 | 0.73 | 0.98 | - |
| Adprm | 41.13 | 37.32 | 42.76 | 49.05 | 47.61 | 50.47 | 46.6 | 51.4 | 49.01 | 47.93 | 51.59 | 49.3 | - |
| Pdlim2 | 25.2 | 24.24 | 23.62 | 36.15 | 32.39 | 30.75 | 38.52 | 31.63 | 36.28 | 34.45 | 31.87 | 40.53 | - |
| Gyltl1b | 0.54 | 0.53 | 0.6 | 0.21 | 0.08 | 0.16 | 0.28 | 0.05 | 0.51 | 0.05 | 0 | 0.14 | - |
| Eogt | 7.46 | 6.85 | 8.15 | 5.66 | 6.02 | 6.43 | 5.3 | 6.49 | 5.22 | 5.39 | 4.75 | 4.32 | - |
| Efna1 | 0 | 0.04 | 0 | 0.04 | 0 | 0.09 | 0.51 | 0 | 0.07 | 0.47 | 0.38 | 0.55 | - |
| Bloc1s6 | 19.91 | 18.82 | 19.94 | 22.8 | 19.78 | 24.64 | 23.68 | 22.72 | 24.35 | 23.87 | 26.79 | 24.47 | - |
| Braf | 3.94 | 4.1 | 4.5 | 4.52 | 5.11 | 4.18 | 6.13 | 5.54 | 6.05 | 5.54 | 6.02 | 5.27 | - |
| Ttc37 | 4.55 | 3.83 | 5.07 | 3.74 | 4.08 | 3.57 | 3.81 | 3.47 | 3.7 | 2.55 | 3 | 1.6 | - |
| Mkrn2 | 17.71 | 16.99 | 15.6 | 22.71 | 21.29 | 24.36 | 22.1 | 20.14 | 22.75 | 22.89 | 26.19 | 22.7 | - |
| Gtpbp5 | 14.92 | 13.83 | 15.51 | 8.71 | 13.16 | 9.39 | 9.66 | 11.23 | 10.55 | 8.87 | 9.77 | 9.55 | - |
| Megf8 | 0.48 | 0.37 | 0.47 | 0.31 | 0.47 | 0.27 | 0.25 | 0.42 | 0.32 | 0.16 | 0.29 | 0.18 | - |
| Npw | 2.07 | 2.11 | 2.05 | 0.83 | 0.72 | 0.95 | 0.66 | 1.16 | 1.16 | 0 | 0.41 | 0.38 | - |
| Mef2b | 0.04 | 0.04 | 0 | 1.06 | 1 | 0.97 | 1.8 | 1.79 | 0.57 | 1.8 | 3.15 | 2.77 | - |
| Vps11 | 14.27 | 13.36 | 13.89 | 18.75 | 17.69 | 18.79 | 19.5 | 18.78 | 19.07 | 23.91 | 23.31 | 23.18 | - |
| Dnajc27 | 1.28 | 1.86 | 1.76 | 1.62 | 1.23 | 1.24 | 0.95 | 1.32 | 0.89 | 0.56 | 0.68 | 0.45 | - |
| Rasal3 | 38.32 | 45.29 | 37.15 | 59.97 | 51.83 | 53.98 | 70.01 | 50.27 | 60.1 | 58.89 | 49.95 | 64.15 | - |
| Ubald1 | 7.86 | 8.27 | 7.18 | 10.71 | 9.77 | 9.02 | 16.98 | 13.14 | 11.19 | 21.46 | 13.85 | 20.3 | - |
| Parp3 | 27.34 | 20.25 | 27.09 | 20.97 | 23.69 | 21.69 | 16.91 | 22.4 | 20.84 | 16.75 | 15.96 | 16.59 | - |
| Tg | 20.22 | 16.92 | 23.64 | 17.5 | 22.84 | 21.79 | 16.44 | 30.29 | 24.93 | 16.98 | 15.85 | 16.12 | - |
| Brk1 | 65.27 | 67.49 | 60.8 | 76.02 | 66.75 | 78.16 | 91.21 | 80.64 | 82.26 | 85.99 | 87.39 | 91.7 | - |
| Arv1 | 7.57 | 8.14 | 7.1 | 5.8 | 6.75 | 4.88 | 3.75 | 6 | 6.31 | 5.16 | 3.17 | 3.21 | - |
| Trim41 | 2.57 | 2.1 | 2.59 | 3.95 | 3.8 | 5.13 | 3.95 | 3.4 | 3.53 | 5.01 | 5.28 | 5.2 | - |
| Mylip | 0.28 | 0.29 | 0.4 | 1.44 | 0.52 | 0.73 | 1.43 | 0.22 | 0.56 | 1.94 | 1.45 | 1.7 | - |
| 9530068E07Rik | 65.36 | 70.57 | 67.61 | 55.34 | 61.39 | 65.81 | 59.11 | 64 | 59.81 | 47.62 | 56.44 | 50.67 | - |
| Scarb2 | 24.62 | 26.36 | 22.58 | 26.97 | 24.88 | 31.25 | 38.43 | 29.75 | 28.67 | 33.29 | 34.85 | 37.78 | - |
| Ubiad1 | 15.86 | 14.6 | 15.36 | 6.75 | 8.74 | 8.57 | 6.09 | 8.58 | 8 | 4.22 | 4.42 | 4.37 | - |
| Mtmr9 | 14.49 | 13.22 | 14.57 | 9.46 | 12.05 | 11.71 | 12.41 | 12.27 | 11.85 | 9.17 | 9.21 | 7.88 | - |
| Ctsa | 25.89 | 24.42 | 23.58 | 37.44 | 29.94 | 31.07 | 56.03 | 36.56 | 41.74 | 56.12 | 52.39 | 61.44 | - |
| Zfp780b | 3.71 | 3.84 | 4.1 | 5.69 | 5.5 | 5.54 | 3.95 | 4.46 | 4.02 | 4.89 | 5.76 | 5.49 | - |
| Nudt5 | 93.69 | 96.45 | 97.05 | 50.26 | 64.79 | 55.85 | 42.06 | 61.02 | 52.07 | 32.56 | 32.15 | 31.93 | - |
| Casp1 | 13.24 | 12.31 | 11.26 | 22.35 | 17.78 | 22.55 | 16.14 | 18.94 | 17.2 | 28.85 | 23.67 | 30.67 | - |
| Sipa1 | 8.88 | 13.65 | 15.78 | 24.09 | 18.79 | 17.88 | 21.83 | 21.89 | 25.31 | 31.34 | 26.61 | 28.5 | - |
| Rab8b | 25.61 | 27.37 | 29.17 | 31.18 | 34.87 | 32.28 | 29.06 | 29.92 | 31.49 | 35.62 | 38.88 | 34.68 | - |
| Uxt | 31.8 | 31.73 | 33.43 | 21.09 | 25.93 | 26.11 | 22.35 | 28.2 | 26.54 | 18.13 | 16.65 | 20.9 | - |
| Tmem132a | 0.92 | 0.73 | 1.21 | 0.54 | 0.48 | 0.38 | 0.3 | 0.56 | 0.58 | 0.48 | 0.23 | 0.4 | - |
| Atg2a | 2.46 | 2.84 | 3.06 | 4.66 | 3.89 | 4.33 | 5.18 | 4.27 | 3.78 | 7.63 | 5.64 | 6.26 | - |
| Col5a3 | 0.88 | 0.68 | 1.37 | 0.01 | 0.08 | 0.08 | 0.21 | 0.57 | 0.36 | 0.39 | 0.08 | 0.39 | - |
| Tpk1 | 2.16 | 1.82 | 1.67 | 4.49 | 2.8 | 2.49 | 3.16 | 3.3 | 3.33 | 4.24 | 5.17 | 3.15 | - |
| Atl3 | 10.32 | 10.82 | 11.19 | 12.95 | 13.13 | 13.03 | 11.86 | 13.01 | 12.87 | 11.81 | 13.95 | 13.23 | - |
| Ppdpf | 59.67 | 66.65 | 61.19 | 88.5 | 71.98 | 82.88 | 102.59 | 93.52 | 99.02 | 95.54 | 91.9 | 110.95 | - |
| P2ry1 | 0.15 | 0.06 | 0.13 | 0 | 0.16 | 0.07 | 0 | 0 | 0.04 | 0 | 0 | 0 | - |
| Gbp2 | 15.56 | 14.84 | 13.94 | 28.8 | 27.82 | 29.4 | 42.39 | 29.5 | 33.56 | 52.15 | 43.51 | 54.28 | - |
| Zfp711 | 0.79 | 0.28 | 0.39 | 0.06 | 0.23 | 0.13 | 0 | 0.05 | 0 | 0.04 | 0.09 | 0.03 | - |
| Eml1 | 0.43 | 0.17 | 0.27 | 0.89 | 1.3 | 0.61 | 0.89 | 0.61 | 0.49 | 2.38 | 2.25 | 2.15 | - |
| Plscr1 | 70.42 | 66.39 | 68.66 | 41.81 | 45.45 | 50.05 | 27.14 | 43.75 | 42.02 | 19.48 | 20.74 | 17.34 | - |
| Alpl | 0.75 | 0.79 | 0.75 | 0.23 | 0.02 | 0.18 | 0.09 | 0.05 | 0.06 | 0.02 | 0 | 0.07 | - |
| Tmppe | 4.02 | 4.24 | 4.22 | 3.64 | 2.86 | 3.25 | 2.63 | 2.89 | 2.96 | 3.12 | 2.43 | 2.35 | - |
| Ube2a | 41.47 | 44.23 | 43.62 | 32.93 | 38.98 | 39.02 | 33.15 | 37.21 | 34.34 | 28.48 | 32.73 | 31.47 | - |
| Eid1 | 2.51 | 2.15 | 1.89 | 2.91 | 2.82 | 2.83 | 4.15 | 2.52 | 3.42 | 3.78 | 4.74 | 4.19 | - |
| Ssh2 | 2.34 | 2.27 | 1.94 | 6.09 | 5.32 | 5.18 | 2.93 | 2.48 | 2.89 | 7.45 | 5.09 | 6.54 | - |
| Farsb | 113.56 | 112.91 | 111.89 | 65.46 | 72.38 | 73.58 | 58.72 | 77.94 | 77.41 | 48.95 | 51.91 | 46.1 | - |
| Cbx5 | 15.45 | 14.69 | 13.88 | 11.01 | 12.14 | 11.76 | 8.17 | 9.39 | 9.27 | 7.97 | 7.48 | 7.7 | - |
| 4930486L24Rik | 4.91 | 6.61 | 3.35 | 7.29 | 5.73 | 5.16 | 14.19 | 6.42 | 7.27 | 11.33 | 8.8 | 10.79 | - |
| Smim3 | 30.93 | 28.64 | 30.6 | 36.72 | 42.21 | 40.58 | 31.74 | 33.93 | 32.04 | 35.09 | 42.24 | 42.12 | - |
| Gm10658 | 0.35 | 0.46 | 0.33 | 0.95 | 0.77 | 0.73 | 1.83 | 0.67 | 1.28 | 1.36 | 1.36 | 1.22 | - |
| Fgd2 | 2.74 | 3 | 2.19 | 0.63 | 0.67 | 0.27 | 1.69 | 2.58 | 2.54 | 0.33 | 0.47 | 0.09 | - |
| Gfer | 26.78 | 26.89 | 27.26 | 21.12 | 25.54 | 24.38 | 23.03 | 24.06 | 23.48 | 18 | 21.27 | 17.91 | - |
| Msantd1 | 0.04 | 0.13 | 0.15 | 0.34 | 0.13 | 0.53 | 0.37 | 0.42 | 0.23 | 0.57 | 0.49 | 0.37 | - |
| Usp21 | 7.37 | 8.43 | 9.1 | 13.59 | 14.22 | 13.57 | 13.04 | 12.36 | 12.67 | 14.3 | 18 | 16.23 | - |
| Pfkp | 144.98 | 145.73 | 141.9 | 167.72 | 169.61 | 173.75 | 224.46 | 185.42 | 209.15 | 185.11 | 208.61 | 217.44 | - |
| Camsap2 | 1.68 | 1.55 | 1.72 | 0.65 | 0.87 | 0.95 | 0.61 | 1.61 | 1.09 | 0.72 | 0.94 | 0.75 | - |
| Amotl1 | 0.1 | 0.07 | 0.12 | 0.01 | 0.06 | 0.06 | 0.03 | 0.1 | 0.07 | 0.01 | 0.01 | 0 | - |
| Gabpb1 | 30.96 | 31.94 | 32.48 | 26.23 | 33.64 | 31.64 | 28.05 | 31.93 | 28.12 | 22.8 | 26.47 | 22.56 | - |
| Smg1 | 4.1 | 3.8 | 4 | 5.2 | 4.93 | 5.42 | 4.38 | 3.97 | 4.82 | 4.8 | 5.35 | 4.74 | - |
| Tmem179b | 47.88 | 48.87 | 46.1 | 68.26 | 62.65 | 67.57 | 61.44 | 61.02 | 59.4 | 75.08 | 70.79 | 82.04 | - |
| Ftsjd1 | 3.27 | 4.03 | 3.25 | 2.03 | 2.3 | 1.98 | 1.4 | 2.58 | 2.07 | 2 | 1.5 | 1.5 | - |
| Zc3h7a | 21.46 | 21.19 | 22.67 | 30.67 | 28.76 | 29.76 | 30.85 | 29.33 | 31.04 | 35.94 | 38.21 | 35.27 | - |
| Fbxl5 | 24.56 | 23.41 | 22.56 | 29.02 | 27.93 | 28.08 | 47.13 | 34.69 | 42.31 | 47.33 | 43.07 | 47.46 | - |
| Slc43a3 | 32.15 | 31.59 | 29.74 | 17.88 | 18.53 | 18.76 | 15.44 | 19.4 | 19.08 | 9.52 | 8.41 | 10.97 | - |
| Gpr56 | 0.13 | 0.46 | 0.35 | 2.61 | 2.11 | 1.24 | 2.1 | 0.94 | 1.13 | 4.86 | 3.16 | 5.2 | - |
| Naf1 | 1.33 | 1.63 | 1.55 | 0.87 | 0.88 | 1.19 | 0.58 | 0.68 | 0.9 | 0.52 | 0.64 | 0.71 | - |
| Rnf5 | 57.45 | 50.95 | 52.74 | 47.3 | 54.77 | 54.04 | 50.16 | 48.76 | 48.1 | 42.66 | 43.13 | 45.36 | - |
| 4930422G04Rik | 1.5 | 1.47 | 1.58 | 1.13 | 1.28 | 1 | 1.14 | 1.32 | 0.85 | 0.89 | 1.03 | 0.75 | - |
| Trip10 | 10.35 | 10.68 | 11.18 | 15.22 | 12.69 | 14.94 | 11.6 | 12.35 | 11.39 | 13.24 | 15 | 15.09 | - |
| Ttc39b | 8.43 | 7.64 | 8.68 | 7.51 | 6.58 | 7.44 | 6.32 | 6.98 | 7.18 | 6.85 | 6.6 | 6.75 | - |
| Duoxa1 | 0.42 | 0.23 | 0.14 | 0 | 0 | 0.04 | 0.15 | 0.04 | 0 | 0 | 0 | 0 | - |
| Tap1 | 98.29 | 104.9 | 93.42 | 131.53 | 115.32 | 118.9 | 165.66 | 125.49 | 126.02 | 172.61 | 178.58 | 188.96 | - |
| Tmem173 | 59.61 | 61.85 | 56.81 | 48.87 | 51.58 | 50.56 | 57.18 | 52.05 | 55.35 | 47.07 | 45.2 | 46.48 | - |
| Tet3 | 2.25 | 3 | 2.59 | 3.69 | 2.9 | 3.47 | 3.27 | 3.29 | 3.22 | 3.48 | 3.51 | 3.31 | - |
| Stra6 | 4.66 | 3.07 | 3.75 | 0.17 | 1.04 | 0.47 | 0.88 | 2.48 | 1.5 | 0.19 | 0.14 | 0.08 | - |
| Fbxo7 | 14.17 | 14.78 | 13.82 | 17.39 | 17.67 | 19.07 | 20.47 | 16.82 | 17.75 | 19.69 | 20.82 | 19.52 | - |
| S1pr2 | 9.02 | 9.88 | 7.72 | 9.61 | 10.67 | 10.67 | 17.25 | 12.75 | 13.17 | 11.66 | 10.22 | 13.52 | - |
| Galt | 5.92 | 6.14 | 5.95 | 7.81 | 6.84 | 6.76 | 10.94 | 10.1 | 7.99 | 9.15 | 10.14 | 8.13 | - |

| | | | | | | | | | | | | | |
|---|---|---|---|---|---|---|---|---|---|---|---|---|---|
| Mob2 | 20.89 | 21.66 | 19.42 | 30.77 | 30.27 | 28.86 | 41.56 | 31.1 | 32.72 | 53.47 | 52.53 | 56.3 | - |
| Hs3st6 | 0 | 0.05 | 0 | 0.05 | 0.15 | 0.05 | 0.75 | 0.19 | 0.4 | 0.56 | 0.25 | 0.61 | - |
| Ticrr | 2.59 | 2.84 | 2.98 | 1.77 | 2.76 | 2.53 | 1.29 | 1.73 | 1.29 | 1.53 | 1.66 | 1.25 | - |
| Ing2 | 0.59 | 0.62 | 1.04 | 1.08 | 1.46 | 1.25 | 1.86 | 0.94 | 0.99 | 1.79 | 1.56 | 2.11 | - |
| Kdm1b | 7.43 | 7.28 | 9.21 | 6.11 | 6.36 | 6.2 | 5.47 | 6.06 | 5.65 | 5.4 | 4.92 | 6.35 | - |
| 2810468N07Rik | 1.23 | 2.37 | 1.84 | 2.48 | 3.19 | 2.98 | 3.24 | 2.57 | 3.74 | 3.94 | 2.92 | 5 | - |
| Cdan1 | 0.97 | 1.37 | 0.87 | 1.38 | 2.11 | 1.72 | 1 | 0.96 | 0.92 | 2.01 | 2.04 | 1.59 | - |
| Eif5a | 1586.45 | 1658.83 | 1662.96 | 886.94 | 917.94 | 900.81 | 1028.85 | 1224.26 | 1248.78 | 640.99 | 657.2 | 683.45 | - |
| Optc | 0 | 0 | 0.13 | 0.16 | 0.08 | 0.32 | 0.47 | 0.4 | 0.67 | 0.5 | 0.54 | 0.5 | - |
| Ctso | 3.77 | 3.72 | 4.24 | 7.56 | 6.11 | 8.61 | 7.16 | 5.63 | 5.24 | 6.58 | 7.67 | 8.75 | - |
| Aebp2 | 12 | 10.39 | 12.28 | 19.03 | 19.07 | 19.4 | 13.01 | 11.45 | 12.24 | 17.91 | 16.87 | 15.46 | - |
| Irf4 | 57.39 | 48.57 | 68.34 | 28.88 | 42.21 | 36.53 | 31.37 | 38.97 | 41.46 | 20.56 | 26.15 | 24.9 | - |
| Cd163l1 | 0 | 0.19 | 0.05 | 0.7 | 0.31 | 0.37 | 0.43 | 0.02 | 0.19 | 0.56 | 0.41 | 0.87 | - |
| Insl3 | 0 | 0.63 | 0.74 | 3.08 | 1.51 | 2.06 | 2.87 | 1.23 | 1.16 | 2.05 | 2.73 | 1.96 | - |
| Crebzf | 1.64 | 2.07 | 2.3 | 3.33 | 3.07 | 4.05 | 3.54 | 3.39 | 3.66 | 4.73 | 3.99 | 4.79 | - |
| Ccnk | 7.57 | 7.54 | 7.32 | 7.85 | 9.04 | 8.85 | 10.87 | 9.44 | 8.85 | 10.5 | 12.69 | 10.1 | - |
| Snhg8 | 53.88 | 56.57 | 58.67 | 75.86 | 89.38 | 85.57 | 63.06 | 71.23 | 72.42 | 76.71 | 85.31 | 72.69 | - |
| Srd5a3 | 16.33 | 17.17 | 15.74 | 9.25 | 12.04 | 10.35 | 8.61 | 10.89 | 11.78 | 9.45 | 6.38 | 7.95 | - |
| Cmc1 | 24.47 | 21.65 | 21.89 | 24.47 | 26.12 | 25.23 | 29.1 | 25.47 | 28.84 | 27.9 | 29.8 | 30.21 | - |
| Nupl1 | 18.53 | 17.19 | 19.71 | 12.22 | 16.35 | 13.27 | 14.93 | 14.13 | 12.86 | 12.52 | 10.92 | 11.12 | - |
| Upp1 | 35.37 | 31.7 | 31.43 | 21.59 | 14.42 | 20.37 | 44.42 | 53.14 | 45.38 | 19.68 | 24.82 | 21.46 | - |
| Cfl2 | 7.42 | 7.03 | 7.31 | 10.43 | 11.47 | 12.44 | 10.29 | 11.67 | 10.81 | 14.37 | 15.71 | 14.22 | - |
| Phf13 | 1.75 | 1.8 | 1.87 | 3.37 | 3.15 | 2.81 | 3.23 | 3.55 | 2.76 | 4.48 | 3.27 | 1.9 | - |
| Elf4 | 7.07 | 7.07 | 7.89 | 9.47 | 9.43 | 9.83 | 10.98 | 10.72 | 10.41 | 12.38 | 14.19 | 12.65 | - |
| Slfn9 | 2.73 | 2.37 | 2.93 | 1.82 | 1.79 | 2.17 | 2.2 | 1.67 | 1.58 | 1.94 | 1.78 | 1.1 | - |
| Eif2d | 55.54 | 50.95 | 54.82 | 44.03 | 40.49 | 45.72 | 38.58 | 45.65 | 43.73 | 35.45 | 38.7 | 42.46 | - |
| Pus1 | 35.84 | 41.38 | 40.93 | 26.59 | 29.36 | 26.77 | 21.72 | 31.66 | 29.32 | 21.19 | 24.3 | 19.67 | - |
| Med11 | 47.37 | 43.53 | 47.61 | 26.31 | 31.18 | 31.64 | 26.63 | 36.01 | 28.36 | 28.12 | 20.38 | 23.99 | - |
| Serf1 | 20.06 | 17.3 | 17.9 | 21.14 | 20.56 | 22.85 | 27.9 | 32.71 | 28.69 | 23.7 | 30.24 | 28.09 | - |
| Aacs | 42.86 | 38.97 | 42.26 | 21.07 | 26.03 | 21.58 | 30.52 | 29.12 | 31.16 | 19.96 | 15.9 | 21.95 | - |
| Dnajc22 | 1.02 | 1.27 | 1.04 | 0.39 | 0.47 | 0.35 | 0.67 | 0.68 | 0.81 | 0.39 | 0.34 | 0.28 | - |
| Thumpd3 | 59 | 53.06 | 53.22 | 45.09 | 48.3 | 50.55 | 47.59 | 60.1 | 50.68 | 41.99 | 49.66 | 45.94 | - |
| Klf13 | 23.37 | 24.64 | 23.68 | 37.65 | 30.4 | 32.01 | 32.37 | 25.22 | 25.65 | 52.64 | 37.21 | 54.24 | - |
| Ccdc51 | 9.99 | 9.03 | 7.99 | 7.28 | 8.12 | 6.73 | 7.89 | 5.56 | 7.48 | 6.63 | 4.78 | 6.28 | - |
| Cpne8 | 4.26 | 4.47 | 4.65 | 7.72 | 8.41 | 7.09 | 10.61 | 8.2 | 7.08 | 12.02 | 13.43 | 13.8 | - |
| Miat | 0 | 0 | 0 | 0 | 0 | 0.01 | 0.03 | 0 | 0.01 | 0.07 | 0.04 | 0.11 | - |
| Ets1 | 32.28 | 34.99 | 35.99 | 62.32 | 54.58 | 52.75 | 48.78 | 37.92 | 43.79 | 72.25 | 57.2 | 75.25 | - |
| Tmem222 | 32.05 | 29.33 | 31.41 | 51.48 | 47.55 | 48.3 | 41.02 | 39.51 | 39.64 | 52.14 | 56.35 | 52.23 | - |
| Mast1 | 0.16 | 0.08 | 0.14 | 0.3 | 0.17 | 0.14 | 0.51 | 0.3 | 0.15 | 0.37 | 0.58 | 0.3 | - |
| Lrmp | 35.32 | 37.6 | 32.48 | 60.04 | 47.31 | 49.74 | 29.65 | 19.48 | 25.58 | 44.01 | 41.06 | 49.47 | - |
| 4831440E17Rik | 0.09 | 0.16 | 0.15 | 0.54 | 0.32 | 0.27 | 0.64 | 0.34 | 0.44 | 0.77 | 0.8 | 0.94 | - |
| Smim4 | 19.12 | 19.49 | 19.05 | 24.78 | 19.21 | 23.1 | 30.23 | 23.32 | 24.5 | 30.24 | 32.2 | 29.75 | - |
| Aph1b | 1.09 | 0.67 | 0.79 | 1.03 | 1.05 | 0.97 | 2.49 | 1.88 | 1.54 | 1.51 | 1.54 | 1.9 | - |
| Cdk14 | 0.41 | 0.27 | 0.15 | 0.45 | 0.17 | 0.34 | 0.06 | 0.3 | 0.08 | 0.07 | 0.05 | 0 | - |
| Abcf3 | 16.34 | 15.39 | 15.61 | 15.74 | 13.23 | 13.63 | 12.18 | 11.77 | 12.04 | 12.28 | 10.97 | 12.85 | - |
| Rrp9 | 103.44 | 97.26 | 104.69 | 32.9 | 38.5 | 45.02 | 27.26 | 47.46 | 41.4 | 20.73 | 20.09 | 21.04 | - |
| Tdpoz4 | 1.57 | 0.9 | 1.23 | 2.39 | 5.64 | 3.55 | 0.56 | 1.08 | 0.71 | 0 | 0.44 | 0 | - |
| Phc2 | 3.92 | 3.44 | 3.4 | 7.2 | 5.42 | 4.73 | 4.92 | 3.85 | 4.14 | 7 | 4.57 | 6.39 | - |
| Rab5a | 9.49 | 8.61 | 9.6 | 6.5 | 6.98 | 5.66 | 3.95 | 6.36 | 7.23 | 6.26 | 5.7 | 4.51 | - |
| Blcap | 7.4 | 6.99 | 7.23 | 10.09 | 8.64 | 10.11 | 12.02 | 9.49 | 9.78 | 11.58 | 11.17 | 10.5 | - |
| H2-K1 | 971.89 | 1052.35 | 981.24 | 1690.05 | 1576.41 | 1610.86 | 1429.21 | 1245.08 | 1314.2 | 1797.48 | 1740.94 | 1831.68 | - |
| Arhgap30 | 42.27 | 41.32 | 42.14 | 48.27 | 49.86 | 46.29 | 43.52 | 46.35 | 43.54 | 48.27 | 46.64 | 49.93 | - |
| Furin | 7.99 | 5.6 | 8.06 | 10.57 | 15.64 | 13.72 | 1.85 | 1.82 | 2.07 | 3.72 | 3.52 | 2.91 | - |
| Unkl | 3.81 | 5.32 | 4.71 | 5.61 | 6.15 | 6.6 | 6.69 | 5.78 | 5.56 | 7.88 | 9.02 | 9.21 | - |
| Mesdc2 | 23.9 | 22.53 | 22.8 | 19.23 | 19.63 | 18.27 | 15.44 | 17.47 | 18.51 | 15.46 | 15.77 | 15.96 | - |
| Msra | 5.98 | 6.63 | 7.63 | 4.02 | 5.37 | 6.02 | 4.46 | 2.96 | 4.39 | 4.16 | 3.94 | 4.03 | - |
| Rsbn1l | 2.81 | 2.7 | 3.04 | 4.33 | 3.81 | 3.81 | 4.14 | 4.16 | 4.35 | 5.58 | 5.16 | 5.23 | - |
| Rc3h2 | 1.5 | 1.38 | 1.56 | 2.4 | 2.19 | 2 | 1.84 | 2.08 | 1.88 | 2.48 | 2.7 | 2.59 | - |
| Fbl | 32.26 | 43.48 | 53.23 | 32.67 | 32.42 | 29.67 | 27.77 | 42.39 | 37.68 | 30.16 | 31.43 | 26.1 | - |
| Spata5 | 21.21 | 22.57 | 21.65 | 17.4 | 18.97 | 18.09 | 15.63 | 17.72 | 17.36 | 13.84 | 14.4 | 18.57 | - |
| Ctsd | 229.94 | 212.03 | 168.8 | 213.44 | 174.42 | 193.05 | 430.22 | 328.61 | 335.37 | 385.76 | 381.35 | 407.97 | - |
| Cd3g | 299.9 | 301.33 | 276.35 | 407.52 | 371.28 | 376.34 | 441.35 | 331.91 | 361.46 | 399.79 | 354.03 | 441.03 | - |
| Ace | 0 | 0 | 0 | 0.05 | 0 | 0 | 0.24 | 0 | 0.09 | 0.06 | 0.12 | 0.1 | - |
| Bcat2 | 38.3 | 39.27 | 37.07 | 27.35 | 26.16 | 25.06 | 35.36 | 36.87 | 35.2 | 23 | 26.53 | 25.27 | - |
| Eef2k | 2.72 | 3.24 | 2.68 | 3.3 | 3.79 | 3.54 | 7.21 | 4.44 | 4.96 | 6.66 | 5.56 | 7.13 | - |
| Pyroxd2 | 0.04 | 0.02 | 0 | 0.14 | 0.16 | 0.1 | 0.17 | 0.13 | 0.07 | 0.22 | 0.34 | 0.18 | - |
| Cdkn2aip | 6.72 | 6.8 | 6.96 | 9.01 | 10.29 | 9.05 | 8.62 | 10.96 | 8.43 | 10.33 | 11.78 | 11.87 | - |
| Notch1 | 6.01 | 5.98 | 5.99 | 8.56 | 8.47 | 7.73 | 7.47 | 6.95 | 7.4 | 8.84 | 7.67 | 8.16 | - |
| Map3k15 | 0.11 | 0.04 | 0.05 | 0.25 | 0.18 | 0.19 | 0.08 | 0.06 | 0.42 | 0.17 | 0.35 | 0.41 | - |
| Fpgs | 28.1 | 25.4 | 27.46 | 15.64 | 15.53 | 17.47 | 12.24 | 18.37 | 16.88 | 11.15 | 10.38 | 12.12 | - |
| Zpbp2 | 0.4 | 0.87 | 0.57 | 1.52 | 1.06 | 0.41 | 3.21 | 0.5 | 0.87 | 2.63 | 2.71 | 2.86 | - |
| Pycard | 43.72 | 43.52 | 39.01 | 76.78 | 65.03 | 69.29 | 87.74 | 62.1 | 67.89 | 95.82 | 96.94 | 113.71 | - |
| Itgb3 | 1.41 | 1.45 | 1.02 | 4.27 | 1.26 | 1.66 | 3.7 | 0.97 | 2.16 | 3.71 | 1.48 | 2.62 | - |
| Klhl23 | 2.23 | 1.76 | 2.25 | 0.47 | 0.62 | 0.4 | 0.52 | 0.8 | 0.43 | 0.19 | 0.31 | 0.14 | - |
| Psg17 | 0.44 | 0.73 | 0.08 | 0.08 | 0 | 0 | 0.14 | 0.08 | 0.03 | 0 | 0 | 0 | - |
| Gm9846 | 796.23 | 761.29 | 447.57 | 1388.72 | 1162.24 | 2059.79 | 1727.97 | 1325.96 | 938.78 | 1710.05 | 1603.04 | 2220.36 | - |
| Sypl | 14.38 | 15 | 14.31 | 11.32 | 12.54 | 12.19 | 13.35 | 11.65 | 14.44 | 9.04 | 11.3 | 9.21 | - |
| Padi2 | 10.58 | 12.36 | 9.52 | 17.21 | 14.67 | 13.68 | 18.77 | 15.48 | 14.95 | 16.43 | 14.32 | 17.77 | - |
| Stx11 | 24.11 | 23.72 | 26.71 | 15.34 | 16.45 | 17.88 | 21.6 | 27.47 | 26.69 | 14.92 | 17.04 | 16.53 | - |
| Ide | 28.82 | 28.34 | 30.79 | 21.24 | 22.35 | 22.34 | 19.58 | 27.07 | 22.92 | 18.54 | 19.64 | 17.66 | - |
| Srm | 71.08 | 88.54 | 116.73 | 33.51 | 41.04 | 36.88 | 26.74 | 55.55 | 52 | 22.01 | 24.92 | 16.02 | - |
| D4Wsu53e | 61.93 | 63.86 | 64.42 | 95.81 | 85.52 | 88.73 | 116.33 | 86.96 | 85.84 | 168.32 | 150.53 | 164.26 | - |
| Tmub2 | 14.78 | 13.96 | 11.69 | 22.4 | 21.38 | 25.4 | 23.79 | 18.94 | 19.27 | 27.56 | 27.9 | 26.34 | - |
| Tsen2 | 14.93 | 15.63 | 14.17 | 10.94 | 12.84 | 10.25 | 8.81 | 11.52 | 12.37 | 9.09 | 8.58 | 7.73 | - |
| Tmem120b | 8.99 | 9.44 | 10.73 | 6.66 | 8.56 | 6.75 | 7.14 | 10.96 | 8.11 | 6.27 | 7.02 | 5.23 | - |
| Epc1 | 3.13 | 3.21 | 3.75 | 4.89 | 4.2 | 4.7 | 4.1 | 4.33 | 3.9 | 6.02 | 4.25 | 5.61 | - |
| Tram1 | 19.19 | 21.05 | 21.17 | 27.08 | 22.95 | 21.22 | 25.15 | 23.94 | 24.46 | 34.17 | 30.96 | 30.52 | - |
| Cdk2ap2 | 78.02 | 78.24 | 72.76 | 121.06 | 105.8 | 112 | 108.69 | 99.29 | 100.65 | 134.76 | 141.66 | 141.82 | - |
| Rab3gap2 | 7.2 | 6.08 | 6.38 | 7.43 | 7.79 | 7.98 | 9.53 | 8.55 | 9.3 | 9.47 | 12.55 | 8.64 | - |
| Srpk2 | 5.8 | 5.06 | 4.75 | 11.13 | 8.77 | 9.29 | 11.29 | 8.53 | 9.15 | 13.47 | 15.05 | 12.53 | - |
| Prrx1 | 0.33 | 0.34 | 0.38 | 0.06 | 0 | 0.19 | 0.29 | 0.19 | 0.14 | 0.08 | 0.03 | 0.16 | - |
| Egln1 | 3.11 | 4 | 3.99 | 5.4 | 3.33 | 3.57 | 7.24 | 4.73 | 4.99 | 4.87 | 5.26 | 5.69 | - |
| Lancl2 | 5.04 | 4.52 | 4.15 | 2.61 | 3.5 | 2.13 | 2.07 | 2.87 | 3.32 | 1.71 | 2.21 | 2.29 | - |
| Dnlz | 6.07 | 7.23 | 7.18 | 6.59 | 7.56 | 5.73 | 5.51 | 8.18 | 6.86 | 5.31 | 4.48 | 4.78 | - |
| Tpx2 | 39.07 | 33.65 | 36.37 | 28.58 | 32.15 | 30.56 | 20.05 | 21.42 | 20.05 | 18.11 | 17.4 | 14.99 | - |
| Ftsj2 | 6.05 | 5.73 | 6.48 | 3.9 | 4.8 | 4.46 | 3.42 | 3.13 | 3.46 | 3.29 | 1.79 | 3.48 | - |
| Wdr45 | 7.09 | 8.35 | 7.19 | 15.08 | 14.61 | 11.71 | 17.32 | 13.69 | 14.44 | 21.49 | 19.02 | 19.37 | - |
| Amfr | 8.74 | 9.73 | 11.39 | 11.57 | 10.73 | 12.32 | 11.78 | 15.07 | 14.03 | 14.68 | 16.25 | 12.77 | - |
| Gm11127 | 1.14 | 1.98 | 2.32 | 3.22 | 2.53 | 3.48 | 3.45 | 3.45 | 3.92 | 3.18 | 4.54 | 4.63 | - |
| Eme2 | 3.78 | 3.75 | 3.63 | 2.62 | 1.85 | 3.14 | 1.4 | 1.27 | 2.33 | 0.68 | 1.09 | 1.52 | - |
| Tcf4 | 0.73 | 0.64 | 0.51 | 0.51 | 0.86 | 0.81 | 2.19 | 1.37 | 1.37 | 2.28 | 1.81 | 2.01 | - |
| Hus1 | 5.25 | 5.75 | 5.49 | 3.2 | 3.54 | 3.16 | 2.12 | 3.41 | 3.2 | 2.66 | 2.77 | 1.56 | - |
| Sos2 | 1.87 | 1.83 | 2.02 | 2.7 | 2.95 | 3.2 | 2.34 | 2.88 | 2.46 | 3.15 | 2.86 | 3.08 | - |
| Rtn4ip1 | 13.37 | 9.37 | 10.98 | 7.55 | 10.05 | 8 | 6.67 | 8.95 | 8.67 | 5.14 | 6.04 | 4.08 | - |
| Ect2 | 16.16 | 13.26 | 15.17 | 10.47 | 15.35 | 13.35 | 6.84 | 7.46 | 8.03 | 7.26 | 7.34 | 5.41 | - |
| Epha7 | 0 | 0 | 0.02 | 0.2 | 0.36 | 0.18 | 0 | 0.01 | 0 | 0.51 | 0.22 | 0.45 | - |
| Ccr7 | 200.95 | 168.5 | 210.12 | 269.23 | 287.53 | 281.99 | 99.84 | 140.21 | 135.4 | 111.28 | 96.89 | 143.36 | - |
| Jrk | 1.35 | 1.52 | 1.14 | 0.32 | 0.55 | 0.54 | 0.36 | 0.29 | 0.56 | 0.16 | 0.13 | 0.36 | - |
| Ntmt1 | 60.61 | 61.58 | 60.91 | 43.37 | 52.16 | 49.26 | 29.84 | 42.61 | 32.8 | 21.58 | 23.85 | 23.96 | - |
| Atp2b1 | 4.9 | 5.25 | 5.94 | 7.42 | 7.82 | 7.33 | 5.07 | 4.6 | 5.93 | 7.06 | 7.69 | 6.7 | - |
| Ssbp2 | 4.59 | 4.47 | 4.42 | 8.45 | 6.44 | 7.57 | 4.67 | 3.02 | 3.95 | 7.38 | 5.29 | 6.77 | - |
| A230072C01Rik | 3.97 | 4.1 | 4.47 | 7 | 6.1 | 6.15 | 6.43 | 6 | 4.95 | 7.48 | 7.5 | 6.72 | - |
| Mxd4 | 0.19 | 0.15 | 0 | 0.91 | 1.11 | 0.49 | 1.07 | 0.8 | 0.59 | 0.97 | 1.62 | 0.98 | - |
| Gas2l3 | 3.32 | 2.73 | 2.9 | 2.79 | 3.27 | 3.17 | 2.18 | 1.77 | 1.62 | 1.99 | 2.07 | 1.65 | - |
| Mre11a | 22.73 | 21.5 | 19.83 | 16.36 | 18.6 | 18.77 | 17.38 | 19.5 | 19.82 | 13.84 | 18.79 | 14.36 | - |
| Rnasel | 2.95 | 3.38 | 2.61 | 4.04 | 3.77 | 4.04 | 5.29 | 4.15 | 4.48 | 4.24 | 4.2 | 5.79 | - |
| Rasgrf2 | 0.37 | 0.49 | 0.46 | 0.74 | 0.55 | 0.83 | 0.91 | 0.5 | 0.96 | 0.75 | 1.07 | 0.76 | - |
| Intu | 1.48 | 1.39 | 1.38 | 1.89 | 1.82 | 1.29 | 1.74 | 1.55 | 1.9 | 1.91 | 2.01 | 2.42 | - |
| Sash3 | 97.68 | 99.34 | 96.59 | 119.44 | 112.42 | 110.76 | 195.02 | 130.8 | 149.52 | 160.97 | 149.45 | 180.13 | - |
| Dnajc30 | 18.08 | 16.49 | 16.47 | 13.03 | 15.23 | 13.45 | 10.04 | 10.68 | 13.38 | 8.82 | 7.85 | 12.03 | - |
| Triobp | 8.08 | 6.49 | 8.11 | 9.45 | 10.19 | 7.8 | 14.19 | 11.1 | 11.36 | 9.24 | 11.36 | 10.16 | - |
| Axin2 | 0.04 | 0.01 | 0 | 0.09 | 0.1 | 0.12 | 0.07 | 0.04 | 0.1 | 0.31 | 0.14 | 0.15 | - |
| 2310044G17Rik | 6.96 | 7 | 7.27 | 8.14 | 7.04 | 6.91 | 5.75 | 8.14 | 8.22 | 9 | 10.35 | 8.46 | - |
| Tlcd2 | 6.13 | 6.07 | 6.35 | 4.56 | 4.52 | 6 | 4.23 | 5.69 | 4.77 | 3.02 | 3.3 | 3.8 | - |
| Tmem109 | 119.31 | 116.44 | 119.33 | 80 | 88.06 | 84.18 | 57.61 | 99.03 | 84.16 | 45.42 | 51.88 | 46.31 | - |
| Gtdc2 | 1.13 | 0.37 | 1.1 | 0.4 | 0.58 | 0.73 | 0.29 | 0.32 | 0.36 | 0.17 | 0.21 | 0.22 | - |
| Zfp874b | 0.97 | 0.75 | 1.2 | 1.64 | 1.24 | 1.98 | 2.37 | 1.81 | 1.69 | 2.85 | 2.59 | 3.4 | - |
| Pvt1 | 6.13 | 5.11 | 5.96 | 7.24 | 8.43 | 7.11 | 10.13 | 10.83 | 10.45 | 9.65 | 11.87 | 8.87 | - |
| Prodh | 1.83 | 1.22 | 1.44 | 0.75 | 0.87 | 0.87 | 0.3 | 0.87 | 0.87 | 0.28 | 0.32 | 0.33 | - |
| 5033411D12Rik | 0.1 | 0.1 | 0.03 | 1.87 | 0.87 | 0.74 | 0.3 | 0.41 | 0.16 | 0.56 | 0.79 | 0.43 | - |
| Zfp687 | 1.78 | 2.12 | 2.41 | 3.36 | 3.42 | 2.73 | 2.49 | 2.09 | 2.27 | 3.04 | 3.11 | 3.88 | - |
| Wdr26 | 4.99 | 5.34 | 5.49 | 7.36 | 6.87 | 6.49 | 6.65 | 5.55 | 6.8 | 8.77 | 8.21 | 7.76 | - |
| Elp5 | 68.77 | 61.74 | 66.75 | 38.19 | 41.4 | 42.57 | 34.26 | 42.59 | 41.54 | 25.52 | 29.71 | 27.77 | - |
| Llgl1 | 6.09 | 6.34 | 7.39 | 8.76 | 7.55 | 7.4 | 9 | 6.64 | 6.46 | 9.39 | 8.59 | 9.07 | - |
| Ash1l | 2.25 | 2.09 | 2.3 | 3.63 | 3.33 | 3.32 | 3.48 | 3.27 | 2.9 | 3.89 | 4.44 | 3.76 | - |
| Klhl12 | 13.63 | 10.97 | 11.69 | 8.4 | 8.8 | 8.36 | 7.21 | 10.5 | 8.71 | 8.09 | 6.06 | 8.39 | - |
| Hist1h3d | 1.54 | 1.64 | 1.63 | 0 | 0.34 | 1.6 | 0 | 0 | 0 | 0.28 | 0.07 | 0 | - |
| Rpusd3 | 14.15 | 13.07 | 12.22 | 9.71 | 11.41 | 9.2 | 7.25 | 10.04 | 10.23 | 6.63 | 6.94 | 8.9 | - |
| Mid2 | 0.68 | 0.54 | 0.88 | 0.77 | 1.06 | 0.89 | 1.74 | 1.54 | 1.19 | 1.48 | 1.49 | 1.14 | - |
| Bcl2l15 | 1.08 | 1.34 | 0.89 | 3.53 | 3.66 | 4.56 | 2.35 | 2.11 | 1.83 | 5.83 | 7.92 | 7.52 | - |
| Acot11 | 3.36 | 3.07 | 3.75 | 3.91 | 3.41 | 3.81 | 8.96 | 5.91 | 6.67 | 6.55 | 6.16 | 7.84 | - |
| Helq | 5.67 | 4.8 | 5.3 | 2.97 | 3.4 | 3.29 | 2.31 | 4.72 | 2.69 | 2.44 | 2.89 | 2.86 | - |
| Npc2 | 60.71 | 59.94 | 64.26 | 150.28 | 132.98 | 143.76 | 150.96 | 102.73 | 113.52 | 184.34 | 169.5 | 215.57 | - |
| Apln | 0.07 | 0.15 | 0.1 | 0.36 | 0.24 | 0.24 | 1.02 | 0.37 | 0.17 | 0.58 | 0.67 | 0.31 | - |
| Laptm4b | 3.55 | 1.96 | 2.28 | 4.6 | 3.75 | 4.54 | 5.79 | 3.99 | 4.3 | 5.43 | 5.59 | 4.22 | - |
| Gimap9 | 44.75 | 40.52 | 42.01 | 46.62 | 49.02 | 46.85 | 50.73 | 48.57 | 50.06 | 55.39 | 56.77 | 59.81 | - |
| Fam57a | 5.52 | 5.07 | 6.57 | 3.4 | 3.69 | 4.87 | 4.2 | 4.04 | 3.92 | 3.69 | 2.98 | 2.36 | - |
| Slc30a3 | 0 | 0 | 0 | 0.09 | 0 | 0.1 | 0 | 0.12 | 0.21 | 0.26 | 0.34 | 0.2 | - |
| Fance | 2.62 | 2.58 | 4.94 | 5.51 | 5.16 | 4.1 | 5.48 | 6.05 | 5.31 | 6.45 | 7.15 | 5.91 | - |
| Pten | 1.7 | 2.45 | 1.71 | 2.7 | 1.82 | 2.28 | 2.78 | 1.86 | 1.96 | 3.53 | 3.62 | 3.38 | - |
| Clspn | 3.86 | 4.78 | 5.13 | 2.42 | 2.82 | 2.73 | 1.78 | 1.5 | 1.49 | 1.78 | 0.88 | 0.91 | - |

| | | | | | | | | | | | | | |
|---|---|---|---|---|---|---|---|---|---|---|---|---|---|
| Cs | 253.38 | 244.19 | 248.16 | 184.53 | 204.37 | 195.34 | 202.79 | 222.27 | 213.17 | 174.36 | 197.2 | 162.15 | - |
| Pparg | 6.27 | 5.83 | 5.42 | 6.47 | 5.51 | 4.74 | 2.85 | 4.21 | 3.18 | 3.7 | 2.75 | 3.07 | - |
| Lin7c | 15.51 | 16.56 | 19.2 | 16.78 | 15.23 | 13.94 | 13.19 | 12.74 | 16.9 | 11.76 | 13.03 | 11.59 | - |
| Acot7 | 183.55 | 159.35 | 166.93 | 114.47 | 114.26 | 113.88 | 144.35 | 145.74 | 157.44 | 89 | 102.14 | 99.04 | - |
| Sesn1 | 9.59 | 11.47 | 10.87 | 13.65 | 13.29 | 14.06 | 14.28 | 10.86 | 12.02 | 18.64 | 15.76 | 17.5 | - |
| Zfp395 | 0.22 | 0.58 | 0.22 | 1.15 | 0.97 | 1.03 | 0.61 | 0.39 | 0.66 | 0.74 | 1.1 | 0.93 | - |
| Adcy6 | 1.51 | 1.11 | 1.61 | 1.68 | 2.21 | 2.45 | 2.15 | 1.91 | 1.95 | 2.56 | 2.18 | 2.18 | - |
| Lrp5 | 0.17 | 0.24 | 0.23 | 0.67 | 0.4 | 0.36 | 0.17 | 0.18 | 0.17 | 0.74 | 0.52 | 0.58 | - |
| Grap | 67.41 | 66.05 | 60.51 | 100.63 | 96.5 | 91.55 | 116.06 | 102.86 | 102.49 | 135.93 | 139.26 | 138.3 | - |
| Atxn7l1 | 15.64 | 12.99 | 13.92 | 24.85 | 21.62 | 22.29 | 32.43 | 24.09 | 24.58 | 29.33 | 36.9 | 34.7 | - |
| Thy1 | 141.59 | 151.13 | 142.87 | 112.93 | 109.31 | 104.86 | 137.55 | 139.57 | 139.73 | 111.06 | 100.1 | 123.3 | - |
| Nqo1 | 1.18 | 0.94 | 1.22 | 0.57 | 0.32 | 0.82 | 0.19 | 0.11 | 0.52 | 0.22 | 0.04 | 0.15 | - |
| Ubxn1 | 118.44 | 114.95 | 121.28 | 142.87 | 138.77 | 144.99 | 130.39 | 140.21 | 129.79 | 144.89 | 149.63 | 145.15 | - |
| Hsd17b7 | 7.73 | 7.05 | 8.14 | 4.41 | 4.72 | 5.31 | 4.64 | 5.7 | 6.42 | 2.84 | 3.71 | 2.82 | - |
| Apaf1 | 3.87 | 3.89 | 5.17 | 6.85 | 7.14 | 6.52 | 7.84 | 6.43 | 7.9 | 9.47 | 10.31 | 8.19 | - |
| Ralgapa2 | 1.15 | 1.15 | 1.03 | 2.3 | 2.13 | 2.11 | 2.14 | 1.2 | 1.79 | 3.84 | 3.14 | 4.02 | - |
| Snx29 | 0.77 | 0.59 | 0.79 | 1.04 | 1.59 | 0.57 | 1.55 | 1.8 | 2.13 | 2.46 | 1.5 | 1.86 | - |
| Spag7 | 88.23 | 80.45 | 81.87 | 75.69 | 69.9 | 73.43 | 80.91 | 83.15 | 80.84 | 61.7 | 72.48 | 64.33 | - |
| P2ry14 | 12.27 | 17.7 | 9.07 | 8.05 | 3.67 | 3.27 | 4.44 | 3.83 | 3.8 | 3.03 | 1.95 | 3.42 | - |
| Emp1 | 0.92 | 0.88 | 0.76 | 2.04 | 1.15 | 0.49 | 4.96 | 3.5 | 3.47 | 3.03 | 4.6 | 3.94 | - |
| Mgrn1 | 6.64 | 6.7 | 8.1 | 10.46 | 11.42 | 10.31 | 11.16 | 10.36 | 11.32 | 13.5 | 11.33 | 13.49 | - |
| Fam43a | 0.72 | 0.47 | 0.4 | 0.33 | 0.31 | 1 | 0.06 | 0.12 | 0.19 | 0.15 | 0.1 | 0.06 | - |
| Slamf7 | 8.01 | 10.85 | 8.65 | 11.86 | 10.06 | 11.94 | 22.41 | 12.17 | 20.11 | 14.3 | 14.76 | 20.02 | - |
| Dynlrb1 | 196.65 | 204.5 | 184.75 | 154.97 | 159.97 | 157.21 | 171.64 | 185.11 | 174.25 | 134.88 | 147.52 | 147.97 | - |
| Manba | 9.95 | 8.71 | 10.09 | 9.29 | 10.53 | 10.07 | 12.32 | 11.45 | 10.8 | 12.89 | 12.72 | 12.88 | - |
| Ercc8 | 11.43 | 9.7 | 9.23 | 5.22 | 6.25 | 7.45 | 5.24 | 6.83 | 5.3 | 3.86 | 4.66 | 4.05 | - |
| Rnmtl1 | 10.64 | 15.29 | 13.2 | 8.15 | 6.9 | 7.82 | 5.51 | 7.28 | 5.54 | 4.29 | 3.92 | 3.05 | - |
| Gstm1 | 1.99 | 1.53 | 1.38 | 3.17 | 2.27 | 3.4 | 8.7 | 5.49 | 5.35 | 7.07 | 8.99 | 9.76 | - |
| Mcoln3 | 0.67 | 0.41 | 0.51 | 0.18 | 0.15 | 0.3 | 0 | 0.1 | 0.03 | 0.03 | 0 | 0 | - |
| Mylk | 0.34 | 0.1 | 0.46 | 0.15 | 0.26 | 0.12 | 0.25 | 0.46 | 0.4 | 0.07 | 0.05 | 0.09 | - |
| Snx6 | 58.52 | 55.4 | 56.46 | 51.02 | 54.91 | 56.64 | 49.57 | 56.3 | 55.19 | 46.43 | 46.74 | 47.31 | - |
| Rcn1 | 13.83 | 14.06 | 13.04 | 12.97 | 11.98 | 11.27 | 9.92 | 12.29 | 11.55 | 6.88 | 8.27 | 8.21 | - |
| Prdx6 | 102.74 | 110.31 | 99.98 | 123.61 | 113.02 | 112.38 | 124.11 | 111.86 | 124.39 | 125.56 | 127.97 | 129.38 | - |
| Ndufb2 | 78.67 | 76.92 | 76.06 | 53.06 | 50.63 | 53.26 | 57.09 | 65.7 | 68.03 | 44.62 | 50.93 | 45.1 | - |
| Preb | 21.58 | 19.6 | 21.58 | 18.22 | 16.53 | 16.72 | 15.48 | 16.78 | 16.98 | 14.83 | 13.77 | 15.87 | - |
| Arid1b | 1.66 | 1.22 | 1.47 | 2.17 | 2.19 | 1.9 | 2.09 | 1.59 | 1.98 | 2.32 | 1.92 | 2.04 | - |
| Usp4 | 50.41 | 45.81 | 47.5 | 43.23 | 44.21 | 42.63 | 37.94 | 41.9 | 42.01 | 37.99 | 37.84 | 38.1 | - |
| Eif3e | 330.83 | 339.57 | 319.69 | 417.47 | 410.69 | 420.71 | 415.63 | 397.11 | 406.07 | 459.08 | 494.32 | 464.95 | - |
| Lgi4 | 0.1 | 0.07 | 0.19 | 0.59 | 0.58 | 0.56 | 0.2 | 0.22 | 0.17 | 0.6 | 0.36 | 0.74 | - |
| Pdpn | 17.71 | 10.92 | 15.56 | 3.79 | 5.89 | 4.87 | 9.68 | 21.02 | 16.78 | 1.71 | 4.53 | 4.04 | - |
| Rfc3 | 118.36 | 119.1 | 115.83 | 66.06 | 71.47 | 69.95 | 89.53 | 96.79 | 103.31 | 59.89 | 67.77 | 57.04 | - |
| Dsg2 | 0.01 | 0.02 | 0.02 | 0.08 | 0.05 | 0.09 | 0.23 | 0.05 | 0.08 | 0.25 | 0.04 | 0.19 | - |
| Dclre1a | 4.06 | 4.23 | 4.25 | 1.99 | 2.88 | 2.44 | 2.1 | 2.74 | 2.32 | 1.57 | 1.59 | 1.63 | - |
| Bhlhe41 | 0.1 | 0.11 | 0.12 | 0.01 | 0.14 | 0.1 | 0.64 | 0.33 | 0.26 | 0.66 | 0.78 | 0.87 | - |
| Zfp963 | 1.1 | 0.95 | 1.47 | 3.36 | 2.85 | 1.99 | 2.53 | 1.76 | 1.36 | 3.22 | 2.5 | 4.28 | - |
| Luc7l | 28.43 | 29.14 | 30.86 | 36.49 | 36.71 | 40.01 | 35.58 | 35.98 | 36.34 | 41.93 | 50.09 | 36.47 | - |
| Bard1 | 1.54 | 1.89 | 1.84 | 0.84 | 1.45 | 0.78 | 1 | 0.54 | 0.94 | 0.54 | 0.84 | 0.61 | - |
| Gtf2e1 | 28.4 | 26.45 | 29.36 | 17.71 | 19.17 | 17.18 | 13.9 | 20.28 | 19.14 | 14.91 | 13.42 | 14.04 | - |
| Cnbp | 609.91 | 623.01 | 628.41 | 496.23 | 513.11 | 529.64 | 421.67 | 562.98 | 550.83 | 446.33 | 446.88 | 427.23 | - |
| 1810011H11Rik | 0.12 | 0.43 | 0.11 | 2.36 | 0.21 | 0.35 | 0.12 | 0.32 | 0.07 | 2.68 | 0.84 | 2.31 | - |
| Fgfr1op | 21.22 | 19.27 | 21.27 | 17.67 | 19.6 | 20.46 | 11 | 15.03 | 14.61 | 15.14 | 14.29 | 13.42 | - |
| Gpnmb | 0.03 | 0.18 | 0.03 | 2.21 | 1.18 | 0.91 | 0.86 | 0.19 | 0.27 | 3.72 | 1.87 | 3.41 | - |
| Tnik | 1.56 | 1.49 | 1.11 | 1.65 | 1.28 | 1.59 | 2.67 | 2.08 | 2.2 | 2.12 | 2.31 | 2.82 | - |
| Capn7 | 18.32 | 19.6 | 19.95 | 20.26 | 21.4 | 20.15 | 22.43 | 19.78 | 18.42 | 22.83 | 21.91 | 24.51 | - |
| Htra2 | 63.63 | 56.64 | 60.76 | 42.35 | 49.77 | 52.94 | 31.99 | 48.49 | 41.57 | 26.32 | 26.76 | 24.84 | - |
| Adrb2 | 0.43 | 0.87 | 0.38 | 3.63 | 2.01 | 2.43 | 0.88 | 0.77 | 0.6 | 2.03 | 2.04 | 2.29 | - |
| Evl | 8.12 | 8.44 | 9.59 | 6.44 | 4.6 | 5.3 | 7.93 | 8.94 | 8.35 | 6.08 | 5.47 | 5.59 | - |
| Gpr150 | 0.4 | 0.68 | 0.58 | 0.06 | 0.17 | 0.33 | 0.13 | 0.25 | 0.25 | 0.13 | 0.09 | 0.16 | - |
| Ncaph2 | 75.78 | 74.75 | 72.08 | 72.9 | 70.8 | 67.93 | 72.17 | 64.56 | 64.1 | 62.33 | 63.56 | 62.36 | - |
| Dnm1l | 26.9 | 26.84 | 28.3 | 22.44 | 23.16 | 23.31 | 21.29 | 23.19 | 20.61 | 16.73 | 19.11 | 18.25 | - |
| Ccdc104 | 23.62 | 20.1 | 21.98 | 17.6 | 16.76 | 16.83 | 16.15 | 18.01 | 16.84 | 14.27 | 14.16 | 15.67 | - |
| Stxbp6 | 0.35 | 0.4 | 0.29 | 0.26 | 0.47 | 0.46 | 0.06 | 0.02 | 0.01 | 0.03 | 0.03 | 0.09 | - |
| Ccdc40 | 7.29 | 5.96 | 6.91 | 5.81 | 6.23 | 6.06 | 6.27 | 7.06 | 7.39 | 4.32 | 4.9 | 4.92 | - |
| Ctnnal1 | 0.51 | 0.27 | 0.44 | 0.27 | 0.19 | 0.24 | 0.12 | 0.03 | 0.01 | 0.03 | 0.08 | 0.02 | - |
| 2310047M10Rik | 0.95 | 1.03 | 1.41 | 1.65 | 3.01 | 2.16 | 3.03 | 3.1 | 2.47 | 4.63 | 3.56 | 3.2 | - |
| 5031414D18Rik | 0.02 | 0.04 | 0.02 | 2.46 | 2.64 | 2.72 | 0.69 | 0.38 | 0.32 | 6 | 4.81 | 6.68 | - |
| Abhd15 | 0.7 | 1.27 | 0.35 | 1.44 | 1.08 | 1.12 | 1.47 | 0.85 | 0.88 | 1.76 | 1.44 | 2.2 | - |
| Zfp85-rs1 | 2.91 | 2.37 | 2.89 | 7.72 | 5.74 | 6.17 | 3.12 | 4.2 | 2.92 | 9.29 | 8.26 | 8.05 | - |
| Cisd3 | 71.09 | 57.8 | 65.07 | 51.28 | 57.96 | 59.73 | 54.69 | 57.24 | 60.66 | 46.72 | 43.86 | 49.41 | - |
| Baz2b | 0.65 | 1.03 | 0.89 | 1.22 | 1.13 | 1.06 | 1.89 | 1.48 | 1.41 | 1.15 | 2.09 | 1.52 | - |
| Zfp68 | 10.11 | 9.77 | 9.71 | 13.14 | 13.99 | 13.96 | 12.13 | 12.26 | 13.33 | 12.61 | 14.55 | 15.47 | - |
| Dand5 | 1.17 | 1.47 | 1.5 | 2.79 | 2.34 | 2.91 | 3.57 | 1.97 | 2.8 | 4.2 | 3.37 | 4.75 | - |
| Itk | 83.04 | 85.88 | 83.27 | 75.39 | 70.56 | 73.4 | 68.23 | 94.52 | 89.81 | 55.69 | 44.19 | 53.93 | - |
| Armc5 | 6.39 | 6.41 | 5.99 | 5.42 | 4.35 | 5.18 | 4.23 | 4.74 | 5.06 | 4.04 | 3.55 | 5.19 | - |
| Vps13b | 3.16 | 2.93 | 3.19 | 3.81 | 3.77 | 4.09 | 3.76 | 3.79 | 3.9 | 4.32 | 4.45 | 3.96 | - |
| Zfp867 | 1.39 | 1.09 | 1.55 | 2.02 | 2.84 | 3.1 | 2.22 | 2.12 | 1.57 | 3.07 | 2.56 | 3.02 | - |
| Foxj3 | 6.57 | 7.37 | 7.8 | 8.67 | 8.74 | 8.59 | 8.35 | 9.13 | 10.53 | 8.63 | 10.5 | 9.37 | - |
| Hmga1 | 44.5 | 46.8 | 53.4 | 63.25 | 72.52 | 55.66 | 81.62 | 59.39 | 62.27 | 83.61 | 80.36 | 88.04 | - |
| Vps37b | 13.98 | 20.39 | 17.26 | 36.74 | 25.94 | 22.3 | 36.21 | 28.21 | 27.23 | 35.39 | 32.07 | 34.6 | - |
| 5031434O11Rik | 0.08 | 0.08 | 0 | 1.14 | 0.52 | 0.98 | 2.32 | 0.74 | 1.49 | 2.55 | 1.72 | 1.85 | - |
| Atp5sl | 7.12 | 7.31 | 7.22 | 12.38 | 10.53 | 12.15 | 14.36 | 13.01 | 13.56 | 17.89 | 16.82 | 18.87 | - |
| Cenpf | 3.39 | 2.21 | 2.04 | 2.72 | 3.61 | 3.33 | 1.66 | 1.4 | 1.48 | 2.04 | 1.57 | 1.34 | - |
| Map3k1 | 8.19 | 8.47 | 8.2 | 13.27 | 12.68 | 12.61 | 18.12 | 12.39 | 15.31 | 16.96 | 20.03 | 19.14 | - |
| Eif2c2 | 9.63 | 8.42 | 9.61 | 14.82 | 14.23 | 15.79 | 14.03 | 12.74 | 14.5 | 16.94 | 18.92 | 15.69 | - |
| Fcho1 | 40.75 | 40.61 | 41.05 | 68.35 | 69.58 | 71.34 | 75.53 | 60.02 | 59.28 | 87.3 | 102.67 | 95.53 | - |
| Pkd1 | 1.14 | 1.01 | 1.4 | 1.78 | 1.89 | 1.93 | 1.75 | 1.63 | 1.83 | 2.05 | 2.1 | 2.08 | - |
| Cic | 2.75 | 2.03 | 2.31 | 4.53 | 3.43 | 4.72 | 4.46 | 3.21 | 3.07 | 4.41 | 4.95 | 4.71 | - |
| 11-Sep | 19.3 | 21.83 | 19.3 | 12.12 | 12.46 | 11.24 | 15.46 | 17.35 | 16.95 | 12.03 | 11.32 | 9.24 | - |
| Ube2j2 | 6.62 | 5.64 | 5.4 | 6.13 | 6.82 | 6.57 | 8.35 | 6.83 | 7.55 | 7.71 | 9.33 | 9.36 | - |
| Ccnb2 | 100.1 | 109.23 | 98.05 | 78.3 | 89.13 | 84.83 | 46.5 | 52.75 | 47.12 | 50.08 | 46.19 | 37.16 | - |
| Ralb | 13.99 | 13.8 | 14.92 | 18.05 | 20.1 | 18.85 | 17.24 | 16.61 | 16.87 | 19.75 | 17.13 | 20.98 | - |
| Zfp263 | 20.87 | 19.69 | 20.98 | 31.9 | 30.73 | 31.59 | 26.29 | 23.93 | 26.48 | 32.08 | 31.67 | 32.89 | - |
| 4921511C10Rik | 0.51 | 0.64 | 0.55 | 0.73 | 1.32 | 0.61 | 1.61 | 1.36 | 0.92 | 1.76 | 1.22 | 1.5 | - |
| Dhfr | 10.45 | 9.57 | 9.43 | 4.18 | 6.37 | 5 | 4.1 | 4.04 | 4.01 | 2.85 | 2.16 | 2.78 | - |
| Add1 | 19.9 | 21.97 | 20.98 | 31.48 | 29.15 | 29.2 | 32.07 | 27.17 | 26.86 | 38.59 | 32.97 | 41.55 | - |
| Wdr5b | 3.29 | 2.15 | 2.79 | 2.41 | 1.96 | 3.18 | 1.38 | 1.2 | 0.96 | 1.41 | 1.45 | 1.32 | - |
| Ppp1r18 | 13.1 | 15.28 | 14.04 | 24.5 | 19.33 | 18.77 | 23.64 | 20.29 | 19.08 | 25.12 | 23.21 | 24.26 | - |
| Dcun1d1 | 18.91 | 19.54 | 19.25 | 22.46 | 21.94 | 22.55 | 24.03 | 19.99 | 21.72 | 26.54 | 26.21 | 26.75 | - |
| 9430008C03Rik | 7.6 | 8.77 | 6.9 | 6.6 | 7.65 | 6.62 | 5.12 | 6.73 | 6.9 | 4.96 | 5.64 | 5.2 | - |
| Gm14288 | 33.01 | 32.65 | 33.8 | 45.5 | 42.06 | 45.3 | 54.05 | 46.24 | 50.09 | 48.4 | 62.32 | 60.87 | - |
| Tmem201 | 8.71 | 8.12 | 9.07 | 6.99 | 6.92 | 5.32 | 5.83 | 6.51 | 6.28 | 3.86 | 5.88 | 5.9 | - |
| Setd5 | 8.14 | 8.53 | 8.88 | 8.77 | 9.74 | 10.1 | 10.85 | 10.57 | 10.17 | 10.07 | 11.33 | 10.7 | - |
| Ubash3b | 3.7 | 3.79 | 3.67 | 3.33 | 3.25 | 2.32 | 5.28 | 6.18 | 5.33 | 6.49 | 4.37 | 5.51 | - |
| Gramd3 | 11.15 | 15.01 | 10.84 | 40.7 | 31.44 | 33.76 | 38 | 19.84 | 23.21 | 70.27 | 59.48 | 72.29 | - |
| Fam172a | 12.63 | 12.86 | 12.93 | 10.45 | 11.08 | 11.13 | 10.03 | 10.42 | 10.86 | 8.69 | 8.39 | 8.38 | - |
| BC147527 | 4.03 | 4.84 | 2.02 | 5.94 | 5 | 4.46 | 5.75 | 3.26 | 3.75 | 7.52 | 7.99 | 9.38 | - |
| Glis1 | 0.02 | 0 | 0.04 | 0.07 | 0.06 | 0.02 | 0.02 | 0.14 | 0.04 | 0.18 | 0.21 | 0.44 | - |
| Mettl6 | 21.71 | 21.69 | 21.71 | 17.71 | 18.69 | 19.68 | 15.75 | 20.19 | 21.01 | 14.92 | 17.08 | 16.61 | - |
| Sepw1 | 142.33 | 145.35 | 138.66 | 232.91 | 217.76 | 229.76 | 303.74 | 226.52 | 256.31 | 325.6 | 311.39 | 339.38 | - |
| Spry1 | 3.28 | 3.32 | 3.95 | 1.57 | 2.51 | 3.18 | 0.18 | 0.8 | 1.23 | 0.94 | 0.32 | 0.51 | - |
| Zfp30 | 0.2 | 0.18 | 0.19 | 1 | 0.43 | 0.78 | 0.33 | 0.47 | 0.41 | 0.72 | 0.47 | 0.89 | - |
| Kif7 | 0.35 | 0.41 | 0.55 | 1.16 | 0.44 | 1.24 | 1.08 | 0.91 | 1.07 | 1.1 | 1.2 | 0.85 | - |
| Acpl2 | 2.05 | 3.1 | 2.13 | 5.57 | 5.08 | 3.76 | 6.96 | 3.93 | 6.05 | 5.45 | 6.93 | 6.85 | - |
| Trex1 | 83.21 | 72.95 | 76.87 | 74.73 | 78.74 | 82.3 | 101.32 | 100.68 | 97.58 | 94.66 | 97.09 | 97.33 | - |
| 1810058I24Rik | 37.19 | 41.2 | 36.78 | 41 | 38.41 | 41.83 | 54.37 | 46.11 | 48.84 | 54.79 | 48.78 | 50.6 | - |
| Hipk1 | 6.18 | 6.67 | 6.51 | 7.91 | 7.53 | 8.51 | 6.95 | 6.8 | 7.54 | 8.49 | 8.87 | 7.57 | - |
| Gprc5a | 0.28 | 0.22 | 0.5 | 0.79 | 1.9 | 0.8 | 0.64 | 1.62 | 1.61 | 1.49 | 2.14 | 1.25 | - |
| Aasdhppt | 24.82 | 26.28 | 25.41 | 18.62 | 19.19 | 22 | 17.63 | 21.38 | 19.31 | 15.47 | 17.76 | 16.86 | - |
| Ndufab1 | 177.83 | 182.64 | 178.28 | 105.95 | 122.74 | 115.11 | 103.08 | 139.05 | 125.07 | 82.95 | 99.21 | 83.87 | - |
| Cstad | 0.14 | 0.28 | 0 | 0.93 | 0.31 | 1.17 | 1.55 | 0.65 | 0.59 | 1.68 | 1.74 | 1.98 | - |
| Card10 | 0.02 | 0 | 0.03 | 0 | 0.09 | 0.11 | 0.24 | 0.22 | 0.22 | 0.3 | 0.33 | 0.39 | - |
| Cln6 | 3.74 | 4.85 | 4.77 | 6.54 | 5.08 | 4.68 | 9.22 | 6.48 | 6.38 | 8.73 | 7.26 | 6.39 | - |
| Plod3 | 14.11 | 12.52 | 16.12 | 9.46 | 10.27 | 10.24 | 11.08 | 13.7 | 11.54 | 7.92 | 8.4 | 7.98 | - |
| Foxo3 | 1.72 | 1.77 | 2.6 | 5.47 | 3.76 | 4.04 | 5.51 | 4.51 | 5.48 | 6.47 | 7.84 | 6.89 | - |
| 9930014A18Rik | 0.73 | 0.66 | 0.69 | 0.15 | 0.25 | 0.51 | 0.09 | 0.3 | 0.33 | 0.15 | 0.37 | 0.01 | - |
| Pcdh19 | 0.1 | 0.04 | 0.09 | 0.19 | 0.25 | 0.12 | 0 | 0.01 | 0 | 0.01 | 0 | 0.01 | - |
| Cdca7l | 44.08 | 45.61 | 42.45 | 34.69 | 35.91 | 35.54 | 36.94 | 41.5 | 39.45 | 26.95 | 31.23 | 30.97 | - |
| Tmem237 | 17.04 | 16.29 | 18.51 | 14.85 | 13.78 | 14.32 | 13.12 | 13.9 | 14.1 | 10.18 | 8.64 | 11.3 | - |
| Prickle1 | 1.83 | 1.6 | 2.42 | 2.17 | 1.96 | 1.7 | 1.1 | 1.74 | 1.06 | 1.12 | 1.27 | 0.8 | - |
| Kif11 | 20.23 | 17.16 | 15.42 | 14.2 | 15.73 | 15.06 | 9 | 8.55 | 7.86 | 9.29 | 10.69 | 6.81 | - |
| Fkbp7 | 1.34 | 1.36 | 0.84 | 1.97 | 1.67 | 3.2 | 4.03 | 2.43 | 2.81 | 3.18 | 3.29 | 2.7 | - |
| Nme6 | 19.93 | 19.59 | 20.68 | 10.98 | 11.79 | 13.31 | 8.45 | 12.09 | 10.1 | 7.81 | 7.57 | 4.84 | - |
| Ubp1 | 5.86 | 6.28 | 5.65 | 8.28 | 8.04 | 8.64 | 7.65 | 7.22 | 9.44 | 11.15 | 11.99 | 8.91 | - |
| Chac2 | 7.77 | 9.51 | 9.04 | 5.24 | 4.51 | 3.99 | 6.19 | 4.42 | 5.82 | 3.37 | 3.3 | 3.48 | - |
| Csgalnact1 | 12 | 9.41 | 10.08 | 3.93 | 7.33 | 6.66 | 5.03 | 8.1 | 8.94 | 2.27 | 1.76 | 3.24 | - |
| Sdccag8 | 9.11 | 8.04 | 9.09 | 14.12 | 11.68 | 11.2 | 17.23 | 12.18 | 14.36 | 15.31 | 18.24 | 16.53 | - |
| Fam174a | 4.58 | 8.81 | 9.01 | 12.27 | 10.58 | 9.62 | 12.08 | 15.25 | 13.57 | 14.47 | 11.73 | 9.21 | - |
| Stk39 | 6.47 | 6.86 | 5.71 | 4.03 | 4.04 | 3.73 | 4.28 | 3.94 | 5.2 | 2.96 | 3.93 | 3.46 | - |
| Tm6sf1 | 3.6 | 4.36 | 3.03 | 8.98 | 7.28 | 6.59 | 4.5 | 4.36 | 5.21 | 9.03 | 8.8 | 11.44 | - |
| Ccdc92 | 1.87 | 2.58 | 4.22 | 4.64 | 3.77 | 3.65 | 8.43 | 5.18 | 6.83 | 7.99 | 7.27 | 6.34 | - |
| Ap1s2 | 2.94 | 3.44 | 2.64 | 1.95 | 2.12 | 3.2 | 2.22 | 1.96 | 1.47 | 1.42 | 2.06 | 2.05 | - |
| Gpc1 | 0.4 | 0.43 | 0.12 | 0.68 | 0.44 | 0.27 | 0.87 | 0.33 | 1.01 | 1.29 | 0.78 | 1.22 | - |
| Igj | 0.19 | 1.05 | 0.51 | 0.03 | 0.15 | 0 | 0.55 | 0.28 | 0.54 | 0 | 0.09 | 0.16 | - |
| Rassf4 | 0 | 0 | 0.03 | 0.34 | 0.31 | 0.49 | 0.5 | 0.07 | 0.15 | 0.31 | 0.37 | 0.27 | - |
| Vhl | 9.68 | 9.38 | 8.81 | 12.69 | 11 | 13.39 | 15.53 | 12.99 | 13.48 | 14.23 | 18.74 | 15 | - |
| Chtf18 | 9.63 | 9.35 | 10.29 | 4.72 | 6.69 | 4.74 | 2.98 | 3.57 | 3 | 2.45 | 2.16 | 1.67 | - |
| 0610037L13Rik | 13.91 | 14.52 | 15.2 | 15.2 | 14.09 | 15.25 | 11.63 | 13.29 | 13.97 | 11.68 | 9.5 | 11.34 | - |
| Letm1 | 22.29 | 28.99 | 33.53 | 17.24 | 17.85 | 18.12 | 17.92 | 23.82 | 21.4 | 17.15 | 15.92 | 15 | - |
| Ccbl1 | 3.38 | 3.08 | 2.98 | 4.11 | 3.67 | 2.88 | 4.69 | 5.95 | 4.68 | 5.41 | 5.16 | 5.21 | - |
| Iqgap2 | 2.64 | 3.04 | 2.48 | 4.24 | 4.49 | 3.74 | 2.57 | 1.75 | 2.88 | 4.39 | 5.25 | 4.76 | - |
| Pmf1 | 40.8 | 50.39 | 46.98 | 25.67 | 29.01 | 31.5 | 23.11 | 25.4 | 24.45 | 15.86 | 18.39 | 17.32 | - |

| | | | | | | | | | | | | | |
|---|---|---|---|---|---|---|---|---|---|---|---|---|---|
| Klrb1f | 0.48 | 0.21 | 0.26 | 0.42 | 0.3 | 0.15 | 2.85 | 0.63 | 1.25 | 3.25 | 1.7 | 4.28 | - |
| Plekhm3 | 0.54 | 0.5 | 0.59 | 0.71 | 0.64 | 0.69 | 0.75 | 0.82 | 1.02 | 1.14 | 0.86 | 0.82 | - |
| Fsd1l | 1 | 0.99 | 1.44 | 0.69 | 0.87 | 0.96 | 0.23 | 0.56 | 0.27 | 0.3 | 0.28 | 0.17 | - |
| 1110002L01Rik | 1.42 | 1.76 | 2.27 | 0.45 | 0.81 | 0.27 | 0.45 | 0.21 | 0.66 | 0.25 | 0.17 | 0.44 | - |
| Enah | 0.85 | 1.05 | 0.88 | 2.97 | 2.33 | 2.1 | 0.82 | 0.54 | 0.6 | 1.99 | 1.71 | 2.22 | - |
| Carf | 0.66 | 0.49 | 0.44 | 1.02 | 0.86 | 1.52 | 0.84 | 1.04 | 1.03 | 1.41 | 1.25 | 1.42 | - |
| Krt18 | 0.33 | 0.21 | 0.19 | 0.54 | 0.45 | 1.05 | 4.52 | 1.93 | 2.38 | 2.62 | 3.17 | 3.46 | - |
| Arrdc1 | 37.12 | 36.4 | 36.53 | 48.05 | 45.53 | 47.53 | 45.31 | 42.55 | 44.19 | 46.52 | 44.21 | 47.53 | - |
| Cdyl2 | 2.33 | 1.55 | 1.97 | 1.89 | 1.76 | 1.86 | 3.28 | 3.04 | 3.58 | 2.98 | 3.85 | 3.2 | - |
| 1810034E14Rik | 1.74 | 1.45 | 1.26 | 1.86 | 1.21 | 1.45 | 3.09 | 1.89 | 1.28 | 3.08 | 2.8 | 4.54 | - |
| Ap3s1 | 23.93 | 31.24 | 31.62 | 34.36 | 30.6 | 33.58 | 61.97 | 44.56 | 52.34 | 59.89 | 58.05 | 47.04 | - |
| Cbr1 | 21.28 | 20.56 | 20.48 | 21.28 | 24.48 | 25.91 | 33.06 | 26.73 | 28.44 | 32.86 | 33.88 | 32.69 | - |
| Nod2 | 0.05 | 0.09 | 0.02 | 0.07 | 0.2 | 0.3 | 0.22 | 0.24 | 0.22 | 0.35 | 0.31 | 0.14 | - |
| Actr1b | 39.93 | 42.61 | 43.5 | 32.16 | 34.82 | 34.82 | 24.1 | 28.17 | 29.96 | 26.33 | 23.04 | 21.23 | - |
| Dctd | 23.16 | 22.66 | 21.34 | 14.63 | 14.92 | 13.83 | 10.85 | 17.04 | 16.68 | 9.04 | 8.87 | 8.81 | - |
| H2-Q7 | 136.68 | 131.49 | 123.61 | 159.99 | 156.42 | 156.09 | 176.52 | 147.99 | 153.27 | 179.02 | 168.57 | 174.32 | - |
| Bmf | 0.14 | 0.15 | 0.08 | 0.44 | 0.81 | 0.56 | 0.14 | 0.33 | 0.15 | 0.66 | 0.4 | 0.56 | - |
| Dcun1d2 | 13.66 | 14.1 | 14.87 | 6.81 | 7.39 | 7.48 | 9.09 | 9.05 | 9.16 | 5.37 | 4.75 | 4.72 | - |
| Ifrd2 | 22.37 | 24.33 | 25.37 | 11.01 | 10.69 | 10.98 | 7.57 | 12.28 | 10.05 | 5.81 | 5.1 | 5.18 | - |
| Akr1e1 | 26.39 | 22.73 | 25.48 | 20.22 | 23.03 | 20.16 | 16.9 | 22.53 | 20.48 | 13.89 | 13.35 | 14.84 | - |
| Hmgn3 | 8.46 | 6.91 | 8.01 | 5.58 | 6.99 | 6.42 | 1.43 | 2.26 | 1.64 | 2.07 | 0.91 | 1.17 | - |
| Il22 | 16.13 | 1.21 | 3.61 | 0 | 0.92 | 0.18 | 0.05 | 0 | 0.3 | 0 | 0 | 0 | - |
| Gtf2e2 | 59.85 | 58.24 | 57.52 | 32.42 | 35.71 | 35.7 | 39.82 | 45.87 | 47.62 | 27.17 | 30.07 | 30.71 | - |
| Fam73b | 3.97 | 3.63 | 3.98 | 3.27 | 3.14 | 2.43 | 1.29 | 2.83 | 1.91 | 2.07 | 1.73 | 1.74 | - |
| Pqlc2 | 5.8 | 6.62 | 4.36 | 2.35 | 4.51 | 3.21 | 3.51 | 3.73 | 5.01 | 3.05 | 3.01 | 4.17 | - |
| Mex3b | 0.02 | 0.09 | 0.05 | 0.07 | 0.13 | 0.42 | 0.15 | 0.29 | 0.17 | 0.27 | 0.38 | 0.34 | - |
| Nrep | 0 | 0.03 | 0 | 0.09 | 0 | 0 | 0.13 | 0.05 | 0.02 | 0.13 | 0.49 | 0.21 | - |
| Ncoa4 | 37.37 | 34.79 | 35.92 | 36.29 | 34.89 | 36.25 | 50.14 | 45.19 | 42.95 | 44.31 | 50.01 | 49.98 | - |
| Mrps9 | 64.43 | 64.27 | 65.74 | 55.36 | 52.35 | 54.06 | 54.28 | 58.86 | 56.67 | 42.78 | 45.87 | 44.04 | - |
| B430306N03Rik | 0.08 | 0.56 | 0.36 | 1.69 | 0.84 | 1.13 | 2.62 | 0.87 | 1.78 | 2.75 | 1.67 | 4.05 | - |
| Camk4 | 9.52 | 10.26 | 9.92 | 13.21 | 12.85 | 12.24 | 16.05 | 12.81 | 13.06 | 18.12 | 14.68 | 17.37 | - |
| Tmsb10 | 1221.26 | 1339.73 | 1232.53 | 917.95 | 927.61 | 941.96 | 1542.25 | 1406.85 | 1488.15 | 890.51 | 947.62 | 991.6 | - |
| Spata2l | 0.48 | 0.81 | 0.32 | 1.59 | 1.36 | 2.01 | 2.56 | 1.36 | 0.9 | 2.96 | 2.33 | 2.74 | - |
| Pla2g4e | 0.01 | 0.03 | 0.08 | 0.11 | 0.04 | 0 | 0.46 | 0.43 | 0.25 | 0.09 | 0.33 | 0.37 | - |
| Ly6e | 292.77 | 345.45 | 304.31 | 549.81 | 496.2 | 497.09 | 275.35 | 236.49 | 252.09 | 393.33 | 357.25 | 421.96 | - |
| Trps1 | 1.16 | 1.13 | 1.32 | 1.15 | 1.13 | 0.93 | 0.58 | 1 | 0.63 | 0.84 | 0.69 | 0.65 | - |
| Apex1 | 101.28 | 103.59 | 99.35 | 58.23 | 67.04 | 70.92 | 43.66 | 66.13 | 58.27 | 43.77 | 52.11 | 40.14 | - |
| Flywch1 | 6.87 | 7.91 | 8.96 | 8.87 | 10.13 | 9.15 | 9.34 | 9.69 | 9.71 | 9.99 | 11.3 | 10.71 | - |
| Vamp2 | 4.91 | 5.57 | 5.12 | 8.21 | 8.02 | 7.1 | 8.84 | 7.74 | 6.79 | 10.36 | 8.6 | 9.22 | - |
| Mrpl38 | 36.21 | 36.72 | 38.13 | 28.22 | 23.54 | 27.56 | 24.5 | 28.72 | 25.93 | 18.85 | 25.25 | 18.18 | - |
| Mcin | 0 | 0 | 0 | 0.03 | 0.13 | 0.11 | 0.05 | 0.02 | 0.09 | 0.36 | 0.13 | 0.02 | - |
| Gbe1 | 4.64 | 6.29 | 5.7 | 16.5 | 11.18 | 11.31 | 40.95 | 13.93 | 23.7 | 26.65 | 20.24 | 30.56 | - |
| Dcaf4 | 11.95 | 12.32 | 13.19 | 17.79 | 16.23 | 17.35 | 19.18 | 16.37 | 16.65 | 16.97 | 20.45 | 20.02 | - |
| Rusc1 | 15.42 | 16.88 | 14.55 | 24.88 | 23.04 | 23.54 | 36.43 | 26.11 | 29.1 | 46.49 | 43.45 | 47.13 | - |
| Mas1 | 0.02 | 0.05 | 0.02 | 0.13 | 0.15 | 0.31 | 0.3 | 0.14 | 0.09 | 0.46 | 0.22 | 0.42 | - |
| Rasgrp2 | 0.89 | 2.1 | 0.67 | 2.36 | 0.83 | 1.08 | 3.38 | 1.93 | 2.89 | 3.97 | 1.77 | 3 | - |
| Tk2 | 8.52 | 8.66 | 8.07 | 12.96 | 13.47 | 13.69 | 15.21 | 12.96 | 13.81 | 19.43 | 16.18 | 18 | - |
| Cd33 | 0.03 | 0.17 | 0.13 | 2.3 | 0.67 | 0.72 | 0.19 | 0.05 | 0.06 | 1.78 | 0.52 | 1.29 | - |
| Ckb | 40.6 | 31.89 | 29.78 | 44.43 | 37.4 | 30.42 | 67.38 | 53.14 | 60.4 | 48.46 | 46.43 | 56.12 | - |
| Hist1h1a | 0.31 | 0.78 | 1.01 | 0.51 | 0.33 | 0.94 | 0.46 | 0.39 | 0.14 | 0 | 0 | 0 | - |
| Tusc3 | 32.75 | 33.27 | 37.7 | 27.36 | 28.88 | 29.58 | 27.38 | 28.53 | 28.37 | 28.39 | 25.15 | 24.08 | - |
| Fam64a | 32.4 | 24.29 | 22.72 | 18.49 | 23.5 | 21.23 | 13.05 | 11.22 | 10.03 | 9.72 | 9.16 | 7.67 | - |
| Zcchc18 | 0.52 | 0.73 | 0.22 | 5.91 | 2.45 | 2.3 | 1.01 | 0.78 | 0.6 | 4.43 | 3.37 | 4.92 | - |
| Cdkl2 | 1.97 | 0.9 | 1.8 | 0.75 | 0.72 | 1.02 | 1.04 | 1.29 | 1.6 | 0.83 | 0.86 | 0.78 | - |
| Cab39l | 22.4 | 21.07 | 21.92 | 27.37 | 24.44 | 26.54 | 31.02 | 32.25 | 31.9 | 29.39 | 37.04 | 29.36 | - |
| Pim3 | 36.89 | 42.18 | 44.03 | 16.88 | 19.69 | 18.06 | 15.37 | 19.89 | 22.78 | 11.24 | 9.58 | 9.83 | - |
| Card6 | 2.66 | 2.71 | 2.65 | 6.12 | 4.64 | 4.99 | 6.66 | 3.59 | 5.03 | 11.44 | 9.13 | 11.91 | - |
| 2810403A07Rik | 21.59 | 21.61 | 22.24 | 27.95 | 26.89 | 26.45 | 27.06 | 28.01 | 24.49 | 33.29 | 33.28 | 33.48 | - |
| Bicd2 | 5.09 | 5.34 | 6.15 | 5.45 | 5.61 | 5.02 | 4.17 | 5.46 | 3.95 | 4.42 | 4 | 3.56 | - |
| 1700012B09Rik | 0.25 | 0.25 | 0.71 | 1.2 | 2.62 | 1.86 | 0.34 | 1.54 | 0.7 | 5.34 | 2.16 | 2.61 | - |
| Tuba8 | 2.23 | 3.94 | 0.72 | 0.84 | 0.65 | 1.16 | 0.56 | 0.21 | 0.6 | 0.24 | 0.34 | 0.08 | - |
| Zfp36l2 | 3.3 | 3.74 | 3.2 | 13.3 | 11.47 | 12.74 | 5.26 | 3.61 | 4.82 | 17.26 | 17.74 | 16.63 | - |
| Isca2 | 68.66 | 70.21 | 62.41 | 49.3 | 55.3 | 52.89 | 44.95 | 46.49 | 45.99 | 38.03 | 32.59 | 44.08 | - |
| Plagl2 | 14.67 | 16.47 | 20.93 | 15.96 | 18.31 | 16.59 | 10.31 | 16.41 | 15.15 | 13.25 | 13.22 | 11.18 | - |
| Prickle3 | 8.41 | 9.6 | 9.88 | 17.93 | 15.37 | 16.89 | 17.23 | 14.34 | 16.17 | 21.72 | 19.35 | 21.9 | - |
| Ttf2 | 16.27 | 14.02 | 12.86 | 11.43 | 12.35 | 11.39 | 8.81 | 10.6 | 9.29 | 9.39 | 9.63 | 7.13 | - |
| Zbtb17 | 13.12 | 13.5 | 13.22 | 16.34 | 13.01 | 15.69 | 15.75 | 16.84 | 13.45 | 16.37 | 15.61 | 16.56 | - |
| Zfp935 | 7.72 | 7.55 | 7.79 | 12.56 | 11.54 | 10.84 | 12.51 | 9.85 | 10.73 | 15.54 | 13.41 | 15.77 | - |
| Gpr146 | 3.61 | 5.1 | 3.64 | 11.24 | 8.51 | 6.9 | 8.62 | 7.2 | 7.76 | 9.99 | 8.93 | 12.28 | - |
| Rpusd4 | 16.44 | 17.39 | 17.36 | 21.42 | 19.8 | 20.47 | 24.11 | 25.17 | 22.85 | 23.65 | 26.98 | 24.6 | - |
| 4933430I17Rik | 1.28 | 0.72 | 0.49 | 0.5 | 0.7 | 0.43 | 0.19 | 0.3 | 0 | 0.22 | 0.14 | 0.23 | - |
| Osbpl1a | 0.94 | 0.84 | 0.41 | 1.18 | 1.24 | 1.33 | 2.2 | 1.09 | 1.58 | 1.73 | 1.75 | 1.21 | - |
| Hist1h3c | 4.69 | 4.63 | 5.79 | 1.66 | 0.73 | 2.51 | 1.24 | 0.68 | 2.45 | 1.31 | 1.09 | 0.6 | - |
| Tmem192 | 16.29 | 17.96 | 19.94 | 25.46 | 21.51 | 22.72 | 26.04 | 22.77 | 25.52 | 38.86 | 29.26 | 29.79 | - |
| Dok7 | 0.02 | 0.05 | 0.02 | 0.13 | 0.1 | 0.08 | 0.25 | 0.02 | 0 | 0.35 | 0.45 | 0.19 | - |
| Cd3e | 80.51 | 80.43 | 81.35 | 117.52 | 101.76 | 110.45 | 129.64 | 98.93 | 114.51 | 164.21 | 137.46 | 162.33 | - |
| Plekha2 | 38.87 | 34.43 | 38.34 | 58.58 | 53.37 | 54.63 | 60.25 | 47.01 | 51.09 | 65.31 | 68.9 | 73.14 | - |
| Elmo1 | 61.63 | 55.8 | 57.65 | 30.53 | 31.48 | 28.59 | 38.78 | 55.3 | 52.86 | 27.55 | 26.69 | 28.44 | - |
| 4933433H22Rik | 0 | 0 | 0 | 0 | 0 | 0.14 | 0.03 | 0 | 0.03 | 0.51 | 0.1 | 0.58 | - |
| Rbm15 | 10.86 | 13.26 | 15.07 | 16.14 | 17.71 | 16.63 | 17.36 | 18.68 | 17.21 | 18.58 | 21.54 | 15.23 | - |
| Ptpn23 | 4.32 | 3.72 | 4.79 | 5.6 | 5.31 | 6.44 | 5.65 | 6.18 | 5.49 | 6.05 | 6.03 | 5.04 | - |
| Rexo2 | 87.36 | 102.03 | 112.35 | 58.44 | 66.49 | 61.46 | 60.34 | 85.29 | 79.4 | 49.7 | 41.16 | 45.69 | - |
| Syne1 | 2.29 | 2.09 | 2.85 | 2.84 | 3.92 | 3.63 | 4.74 | 3.98 | 4.04 | 4.18 | 3.78 | 4.67 | - |
| Fam118b | 23.6 | 18.79 | 20.17 | 12.63 | 16.39 | 13.26 | 9.92 | 12.62 | 12.29 | 10.12 | 9.49 | 6.69 | - |
| Fam35a | 3.59 | 3.69 | 3.45 | 1.92 | 3.63 | 3.33 | 2.45 | 3.03 | 4.04 | 2.62 | 2.2 | 2.1 | - |
| Mfsd2a | 15.03 | 15.45 | 19.22 | 9.62 | 14.02 | 15.08 | 7.08 | 10.24 | 12.3 | 6.09 | 6.71 | 6.95 | - |
| Galk1 | 137.49 | 137.64 | 136.62 | 105.45 | 102.31 | 100.05 | 118.16 | 109.13 | 116.34 | 82.77 | 67.61 | 95.15 | - |
| Rabgap1 | 2.32 | 1.92 | 1.94 | 3.1 | 2.55 | 3.47 | 2.94 | 2.76 | 2.97 | 3.48 | 2.76 | 3.51 | - |
| Bax | 151.1 | 160.64 | 162.27 | 88.37 | 104.99 | 101.49 | 134.41 | 135.85 | 141.26 | 85.97 | 97.16 | 94.26 | - |
| Nsdhl | 42.64 | 36.05 | 40.47 | 32.68 | 36.11 | 36.91 | 36.67 | 31.46 | 32.12 | 26.61 | 32.57 | 31.15 | - |
| Psmd5 | 69.21 | 63.17 | 63.53 | 46.71 | 45.21 | 45.07 | 34.2 | 48.28 | 46.42 | 31.19 | 38.39 | 34.14 | - |
| Morc2a | 3.36 | 4.48 | 4.05 | 3.32 | 3.99 | 3.84 | 2.91 | 3.41 | 2.94 | 2.95 | 2.05 | 2.62 | - |
| Il12rb1 | 40.63 | 40.58 | 40.37 | 59.34 | 52.68 | 57.91 | 44.44 | 44.72 | 47.02 | 66.23 | 55.03 | 68.23 | - |
| Fam178a | 3.53 | 3.86 | 4.04 | 4.89 | 4.57 | 4.46 | 4.23 | 3.73 | 4.11 | 4.87 | 6.1 | 5.11 | - |
| Nrm | 21.99 | 21.31 | 21.49 | 33.66 | 35.9 | 34.62 | 28.81 | 25.4 | 25.69 | 38.84 | 33.85 | 36.5 | - |
| Aurka | 57.66 | 40.78 | 40.97 | 27.2 | 39.65 | 35.93 | 20.06 | 20.78 | 19.77 | 18.17 | 16.83 | 16.25 | - |
| Dyrk1b | 0.4 | 0.27 | 0.25 | 1.39 | 0.88 | 0.82 | 1.24 | 0.71 | 0.78 | 1.76 | 1.49 | 1.94 | - |
| Arhgef1 | 15.99 | 23.52 | 24.9 | 37.85 | 33.31 | 32.55 | 49.02 | 38.21 | 46 | 53.95 | 49.8 | 46.83 | - |
| A830080D01Rik | 9.21 | 9.01 | 9.64 | 5.76 | 8.25 | 7.49 | 7.07 | 7.42 | 6.09 | 4.72 | 5.79 | 4.18 | - |
| Cdkn2aipnl | 75.74 | 68.86 | 73.17 | 54.08 | 59.2 | 51.32 | 50.04 | 50.8 | 49.22 | 40.42 | 39.51 | 40.45 | - |
| Zbtb40 | 1.33 | 1.23 | 1.28 | 1.54 | 1.61 | 1.44 | 1.47 | 1.73 | 2.11 | 1.89 | 1.86 | 1.86 | - |
| Tube1 | 14.83 | 10.81 | 11.96 | 10.9 | 11.21 | 10.99 | 8.73 | 7.94 | 8.83 | 6.29 | 5.91 | 6.3 | - |
| Bdp1 | 3.25 | 3.96 | 4.02 | 3.11 | 3.21 | 3.34 | 2.52 | 3.18 | 2.84 | 2.81 | 2.74 | 2.99 | - |
| 6-Mar | 5.77 | 6.19 | 5.88 | 7.61 | 6.61 | 7.34 | 8.91 | 8.42 | 7.92 | 10.37 | 10.94 | 9.03 | - |
| Nktr | 7.31 | 7.78 | 7.79 | 9.83 | 9.16 | 10.45 | 14.43 | 11.61 | 12.88 | 15.22 | 14.97 | 13.79 | - |
| Nt5c | 156.99 | 154.05 | 147.48 | 119.1 | 127.51 | 125.32 | 118.9 | 131.74 | 129.86 | 107.53 | 104.43 | 106.68 | - |
| Leo1 | 20 | 21.92 | 22.13 | 14.39 | 15.63 | 13.96 | 10.43 | 13.58 | 11.36 | 9.21 | 9.05 | 8.78 | - |
| Ctla2b | 22.5 | 36.47 | 17.38 | 80.07 | 57.96 | 54.11 | 101.7 | 37.58 | 59.93 | 122.19 | 93.63 | 148.34 | - |
| 4932438A13Rik | 2.34 | 2.28 | 2.63 | 5.01 | 4.61 | 4.84 | 3.73 | 3.33 | 3.49 | 6.46 | 6.01 | 6.67 | - |
| Lima1 | 0.44 | 0.24 | 0.36 | 0.08 | 0.26 | 0.4 | 0.32 | 0.29 | 0.27 | 0.14 | 0 | 0.03 | - |
| Frmd8 | 44.72 | 42.11 | 42.64 | 71.23 | 54.81 | 58.87 | 79.19 | 49.32 | 57.53 | 81.74 | 68.62 | 88.01 | - |
| Etv5 | 5.02 | 3.98 | 3.61 | 1.5 | 2.69 | 2.39 | 2.56 | 2.86 | 2.72 | 1.15 | 1.05 | 0.87 | - |
| Dph5 | 59.34 | 52.55 | 56.15 | 34.44 | 37.5 | 38.8 | 29.73 | 47.68 | 37.6 | 22.55 | 28.47 | 24.79 | - |
| Dazap2 | 175.08 | 189.53 | 175.82 | 205 | 195.32 | 201.29 | 219.6 | 190.2 | 202.98 | 235.17 | 234.9 | 248.61 | - |
| Mrpl33 | 180.7 | 169.69 | 180.93 | 142.28 | 142 | 158.32 | 179.49 | 172.06 | 172.12 | 128.41 | 134.11 | 138.46 | - |
| Prdm16 | 1.34 | 0.99 | 1.07 | 0.52 | 0.76 | 0.71 | 0.68 | 0.64 | 1.1 | 0.43 | 0.6 | 0.63 | - |
| A530072M11Rik | 0.52 | 0.22 | 0.35 | 1.43 | 0.67 | 0.52 | 0.96 | 1.51 | 1.36 | 0.65 | 1.3 | 1.65 | - |
| Phf7 | 9.08 | 9.06 | 9.21 | 11.75 | 13.22 | 11.98 | 11.3 | 14.84 | 11.65 | 12.23 | 14.54 | 11.97 | - |
| Appl2 | 4.52 | 4.92 | 4.51 | 9.59 | 8.69 | 9.72 | 16.1 | 8.7 | 13.57 | 16.03 | 13.88 | 17.48 | - |
| Snapc4 | 5.91 | 5.12 | 5.52 | 3.38 | 4.08 | 3.55 | 2.73 | 4.46 | 4.09 | 3.42 | 3.46 | 2.82 | - |
| Tnks2 | 4.86 | 5.55 | 5.5 | 4.44 | 4.29 | 5.05 | 4.71 | 4.66 | 4.88 | 4.02 | 4 | 4.18 | - |
| Slc12a6 | 2.43 | 2.35 | 2.68 | 5.22 | 4.29 | 4.69 | 4.55 | 3.86 | 4.78 | 5.09 | 6.52 | 6.14 | - |
| Crim1 | 3.39 | 3.03 | 3.52 | 2.87 | 3.22 | 3.27 | 2.01 | 3.42 | 3.05 | 1.88 | 1.46 | 1.56 | - |
| Ears2 | 8.35 | 7.63 | 7.79 | 6.23 | 8.69 | 6.74 | 7.1 | 10.5 | 7.67 | 5.64 | 4.74 | 5.28 | - |
| Fadd | 5.29 | 4.77 | 5.09 | 4.1 | 4.22 | 3.41 | 2.26 | 2.66 | 2.81 | 4.36 | 2.14 | 2.9 | - |
| 1810037I17Rik | 129.61 | 129.43 | 127.43 | 83.6 | 96 | 96.38 | 131.69 | 115.6 | 135.65 | 64.93 | 75.84 | 71.46 | - |
| Gm14325 | 3.17 | 3.49 | 4.14 | 4.38 | 3.74 | 2.89 | 3.76 | 3.58 | 4.34 | 6.1 | 4.58 | 5.6 | - |
| F2rl2 | 2.87 | 3.56 | 2.28 | 1.51 | 1.22 | 0.8 | 2.99 | 2.43 | 4.79 | 1.09 | 0.7 | 1.59 | - |
| Eri3 | 16.05 | 17.89 | 19.68 | 14.74 | 14.63 | 12.47 | 12.74 | 14.25 | 12.65 | 10.69 | 10.64 | 10.58 | - |
| Midn | 1.8 | 2.09 | 2.51 | 3.27 | 3.08 | 2.92 | 3.63 | 3.28 | 2.85 | 3.13 | 3.31 | 3.82 | - |
| Tom1l1 | 5.38 | 2.91 | 2.76 | 1.39 | 1.17 | 1.42 | 1.61 | 1.01 | 0.68 | 0.25 | 0.54 | 0.54 | - |
| 5830432E09Rik | 1.07 | 1.56 | 0.98 | 1.35 | 1.3 | 1.88 | 2.58 | 1.77 | 2.33 | 3.52 | 2.9 | 1.81 | - |
| Dennd3 | 7.83 | 6.62 | 8.73 | 10.97 | 12.94 | 14.12 | 13.61 | 17.91 | 14.9 | 16.16 | 19.92 | 15.19 | - |
| Mettl4 | 10.99 | 11.2 | 12.01 | 10.35 | 9.23 | 11.96 | 6.17 | 10.62 | 11.2 | 7.48 | 7.85 | 6.17 | - |
| Mnda | 6.13 | 9.35 | 9.43 | 16.04 | 16.82 | 19.02 | 13.33 | 2.92 | 5.03 | 22.79 | 16.16 | 23.8 | - |
| Ak1 | 6.7 | 3.34 | 4.65 | 2.86 | 3.98 | 2.79 | 3.19 | 3 | 1.71 | 2.28 | 3.09 | 2.43 | - |
| Nt5dc2 | 3.64 | 2.87 | 3.33 | 1.86 | 2.64 | 4.19 | 3.02 | 2.71 | 1.99 | 1.83 | 1.24 | 1.88 | - |
| Tmem154 | 5.91 | 5.61 | 4.76 | 5.5 | 5.68 | 5.86 | 3.03 | 3.57 | 4.16 | 3.06 | 2.86 | 3.38 | - |
| Idh3b | 130.89 | 131.01 | 129.38 | 149.56 | 139.42 | 145.67 | 155.17 | 142.34 | 150.36 | 160.29 | 184.64 | 168.6 | - |
| Helb | 3.51 | 3.18 | 3.14 | 4.15 | 5.96 | 5.78 | 6.91 | 4.19 | 4.66 | 6.19 | 6.73 | 7.64 | - |
| Rpain | 21.04 | 20.9 | 20.47 | 32.49 | 28.03 | 28.89 | 27.13 | 23.6 | 20.46 | 28.22 | 26.84 | 27.14 | - |
| Grtp1 | 5.14 | 4.71 | 5.57 | 1.45 | 0.67 | 1.7 | 3.82 | 4.26 | 3.19 | 0.61 | 1.41 | 0.75 | - |
| Plekha5 | 2.49 | 2.27 | 2.76 | 4.59 | 4.12 | 3.71 | 4.17 | 3.84 | 3.57 | 4.95 | 4.55 | 5.63 | - |
| Pnpla2 | 20.41 | 19.49 | 20.58 | 25.15 | 21.68 | 19.72 | 26.44 | 24.9 | 25.39 | 30.39 | 29.68 | 29.96 | - |
| Gpi1 | 293.61 | 299.64 | 295.65 | 501.96 | 427.93 | 450.84 | 686.18 | 451.04 | 529.49 | 663.68 | 693.79 | 723.91 | - |
| 9630033F20Rik | 3.09 | 2.82 | 2.84 | 1.18 | 1.49 | 2.01 | 0.79 | 1.85 | 1.63 | 0.38 | 0.81 | 0.88 | - |
| Fgfr1 | 0.02 | 0.01 | 0.03 | 0.33 | 0.39 | 0.07 | 0.15 | 0.07 | 0.09 | 0.46 | 0.22 | 0.51 | - |
| Sys1 | 12.11 | 19.12 | 20.98 | 21.85 | 16.89 | 19.99 | 26.09 | 21.78 | 25.03 | 27.95 | 24.95 | 26.71 | - |
| Minos1 | 55.61 | 58.7 | 55.71 | 42.66 | 41.57 | 44.7 | 49.66 | 48.84 | 50.93 | 34.15 | 42.15 | 38.45 | - |
| Rps2 | 919.45 | 1525.18 | 1908.63 | 2543.17 | 2479.66 | 2246.81 | 3000.76 | 3132.2 | 3222.1 | 3210.55 | 3300.82 | 2367.66 | - |
| Fbxo17 | 9.28 | 7.44 | 7 | 5.44 | 5.55 | 5.3 | 4.66 | 5.2 | 4.55 | 4.13 | 3.62 | 2.94 | - |

| | | | | | | | | | | | | |
|---|---|---|---|---|---|---|---|---|---|---|---|---|
| Cass4 | 0.38 | 0.81 | 1.1 | 1.07 | 0.42 | 0.97 | 2.66 | 0.89 | 2.5 | 2.57 | 1.09 | 3.04 | - |
| Dennd1b | 2.33 | 2.57 | 2.32 | 2.64 | 2.5 | 1.76 | 3.66 | 3.49 | 2.81 | 2.94 | 4.4 | 3.23 | - |
| Gpr157 | 0.97 | 1.04 | 0.84 | 1.51 | 1.29 | 1.29 | 1.25 | 1.12 | 1.44 | 2.03 | 1.61 | 1.31 | - |
| Mdh1 | 143.68 | 136.17 | 140.87 | 125.76 | 127.01 | 126.14 | 123.53 | 126.81 | 128.41 | 107.08 | 116.08 | 108.79 | - |
| Pola1 | 13.98 | 14.33 | 14.37 | 7.59 | 7.48 | 6.69 | 4.16 | 5.73 | 6.36 | 3.63 | 3.52 | 3.31 | - |
| Thap11 | 5.16 | 5.64 | 7.14 | 10.43 | 8.68 | 8.08 | 9.55 | 10.93 | 11 | 16.04 | 12.14 | 12.23 | - |
| Klhl36 | 3.38 | 4.37 | 3.33 | 4.81 | 3.73 | 5.8 | 5.66 | 4.5 | 5.21 | 5.07 | 4.89 | 7.78 | - |
| Ptdss2 | 12.3 | 12.46 | 11.25 | 15.89 | 15.2 | 16.41 | 20.8 | 17.73 | 19.14 | 23.41 | 21.99 | 20.33 | - |
| Ccdc102a | 3.47 | 6.41 | 6.36 | 12.56 | 10.84 | 10.76 | 7.83 | 7.38 | 10.69 | 10.58 | 8.99 | 10.16 | - |
| Eif2s2 | 224.04 | 235.83 | 229.09 | 191.73 | 205.74 | 208.01 | 206.89 | 236.29 | 230.36 | 187.06 | 199.79 | 183.26 | - |
| Zfp524 | 10.69 | 8.37 | 10.57 | 18.42 | 19.43 | 17.83 | 26.94 | 16.69 | 19.42 | 30.9 | 35.53 | 40.74 | - |
| Nudcd2 | 83.98 | 80.15 | 80.95 | 52.64 | 62.03 | 62.25 | 55.65 | 73.24 | 66.06 | 50.39 | 54.84 | 48.02 | - |
| 2700060E02Rik | 160.28 | 160.03 | 152.98 | 187.06 | 181.74 | 202.96 | 215.92 | 205.91 | 200.76 | 208.29 | 239.78 | 212.3 | - |
| Nfatc3 | 4.01 | 4.68 | 5.36 | 7.09 | 7.3 | 7.08 | 11 | 7.93 | 8.14 | 13.65 | 11.78 | 11.26 | - |
| Ndufaf7 | 22.8 | 21.12 | 22.8 | 18.32 | 15.61 | 15.84 | 15.47 | 17.86 | 16.99 | 12.51 | 13.93 | 16.18 | - |
| Clic4 | 45.83 | 48.46 | 48.52 | 32.13 | 42.87 | 40.26 | 38.38 | 52.07 | 46.39 | 31.27 | 33.94 | 32.11 | - |
| Prep | 96.79 | 91.92 | 96.43 | 57.17 | 65.43 | 60.75 | 98.23 | 95.97 | 98.55 | 61.39 | 68.04 | 66.83 | - |
| Prmt7 | 73.63 | 73.33 | 75.29 | 52.55 | 51.73 | 54.77 | 36.56 | 56.34 | 47.74 | 28.68 | 36.77 | 30.18 | - |
| Slc25a20 | 28.14 | 27.9 | 28.93 | 23.68 | 22.34 | 20.65 | 26.05 | 28.76 | 25.84 | 21.78 | 19.01 | 20.04 | - |
| Ghdc | 9.8 | 8.73 | 8.09 | 11.13 | 9.75 | 11.4 | 12.82 | 11.75 | 12.42 | 11.62 | 14.86 | 13.15 | - |
| Slc28a2 | 9.98 | 8.92 | 7.35 | 11.93 | 9.73 | 10.99 | 16.23 | 12.34 | 14.83 | 11.87 | 10.54 | 14.48 | - |
| Pigf | 42.14 | 37.46 | 36.9 | 28.18 | 34.36 | 35.85 | 37.22 | 36.2 | 32.93 | 28.59 | 33.65 | 28.24 | - |
| Zdhhc21 | 2.13 | 2.76 | 2.63 | 3.05 | 2.57 | 3.51 | 2.75 | 2.92 | 3.85 | 3.59 | 3.47 | 3.29 | - |
| Timd2 | 3.05 | 1.28 | 2.36 | 1.23 | 1.63 | 2.13 | 0.7 | 2.04 | 1.52 | 0.78 | 0.92 | 1.38 | - |
| Stim1 | 9.62 | 10.37 | 9.17 | 12.01 | 11.19 | 12.03 | 12.63 | 11.39 | 12.76 | 11.94 | 13.05 | 14.15 | - |
| Ston2 | 0.42 | 0.32 | 0.45 | 0.52 | 0.54 | 0.54 | 1.61 | 0.93 | 1.18 | 1.52 | 1.48 | 1.23 | - |
| Tex264 | 14.66 | 16.82 | 14.47 | 19.98 | 17.96 | 15.81 | 21.98 | 18.34 | 15.54 | 20.84 | 20.42 | 21.26 | - |
| Zcchc7 | 17.32 | 18.01 | 17.69 | 22.75 | 20.14 | 22.72 | 21.57 | 22.25 | 21.03 | 23.17 | 28.44 | 26.38 | - |
| Lman2l | 11.8 | 8.93 | 10.05 | 8.55 | 10.32 | 10.63 | 6.82 | 9.42 | 7.98 | 5.96 | 6.7 | 6.53 | - |
| Zfp874a | 3.19 | 3.2 | 4.51 | 4.54 | 5.86 | 5.69 | 5.91 | 5.09 | 5.58 | 6.86 | 9.32 | 7.34 | - |
| Gm3414 | 0.18 | 0.31 | 0.17 | 0.51 | 0.3 | 0.35 | 0.4 | 0.24 | 0.53 | 0.81 | 0.64 | 0.5 | - |
| Reep1 | 2.85 | 2.36 | 2.33 | 2.46 | 2.04 | 1.59 | 9.53 | 4.79 | 6.43 | 4.8 | 4.75 | 5.51 | - |
| Palb2 | 3.58 | 4.29 | 4.1 | 1.77 | 3.13 | 2.69 | 1.76 | 2.99 | 2.1 | 1.95 | 1.32 | 1.02 | - |
| St3gal2 | 2.35 | 2.67 | 2.39 | 1.26 | 1.58 | 1.43 | 0.86 | 0.85 | 1.2 | 0.63 | 0.51 | 1.11 | - |
| Atox1 | 78.2 | 79.71 | 79.73 | 56.71 | 60.29 | 67.85 | 69.59 | 66.65 | 68.96 | 59.56 | 50.51 | 59.35 | - |
| 2210039B01Rik | 1.28 | 1.37 | 1.37 | 2.26 | 2.75 | 2.84 | 2.6 | 2.04 | 2.55 | 2.4 | 4.83 | 3.95 | - |
| 6-Sep | 39.21 | 38.41 | 37.73 | 57.68 | 48.88 | 54.9 | 66.56 | 46.22 | 52.43 | 64.04 | 61.22 | 67.98 | - |
| Exosc6 | 2.08 | 2.03 | 2.06 | 0.7 | 1.04 | 1.16 | 0.46 | 1.02 | 0.89 | 0.41 | 0.49 | 0.82 | - |
| Ctps2 | 11.08 | 10.84 | 10.42 | 15.78 | 14.02 | 15.48 | 17.57 | 13.26 | 13.52 | 16.6 | 17.18 | 17.71 | - |
| Crtc2 | 6.36 | 7.57 | 7.98 | 9.39 | 9.71 | 7.52 | 6.25 | 7.89 | 8.18 | 10.86 | 9.59 | 9.82 | - |
| Smurf2 | 0.84 | 1.08 | 0.94 | 3.01 | 2.22 | 2.85 | 1.29 | 0.95 | 1.1 | 3.28 | 3.24 | 2.94 | - |
| Ttc30a1 | 1.7 | 2.08 | 1.39 | 0.9 | 1.71 | 1.13 | 0.4 | 0.83 | 0.4 | 0.15 | 0.22 | 0.15 | - |
| Sar1a | 92.35 | 92.29 | 92.72 | 87.4 | 83.42 | 89.8 | 75.52 | 89.3 | 88.09 | 73.12 | 79.13 | 72.71 | - |
| Nbn | 13.22 | 15.11 | 12.61 | 11.62 | 11.85 | 11.96 | 9.44 | 12.8 | 9.46 | 9 | 8.84 | 8.01 | - |
| Spryd4 | 11.08 | 10.86 | 13.48 | 7.98 | 9.24 | 9.68 | 8.36 | 11.69 | 9.22 | 7.26 | 8.61 | 9.3 | - |
| 4930506M07Rik | 0.52 | 0.2 | 0.52 | 0.09 | 0.18 | 0.05 | 0.03 | 0.09 | 0.14 | 0.06 | 0 | 0 | - |
| Sfmbt1 | 1.89 | 2.32 | 1.71 | 1.96 | 1.66 | 1.3 | 0.97 | 1.42 | 1.16 | 1.42 | 1.23 | 1.23 | - |
| Mansc1 | 2.31 | 2.33 | 2.78 | 2.02 | 2.13 | 1.58 | 1.72 | 2.55 | 1.69 | 1.06 | 0.93 | 0.78 | - |
| Sult2b1 | 3.32 | 3.18 | 2.52 | 8.01 | 3.96 | 5 | 8.09 | 3.52 | 4.3 | 15.01 | 7.32 | 13.7 | - |
| Pion | 1.34 | 1.59 | 1.52 | 4.86 | 2.75 | 2.85 | 6.77 | 1.52 | 2.94 | 5.83 | 3.85 | 7.08 | - |
| Ubxn7 | 4.71 | 4.4 | 4.53 | 5.56 | 5.15 | 5.45 | 5.36 | 5.29 | 5.32 | 5.67 | 6.11 | 5.55 | - |
| Dse | 2.31 | 3.31 | 2.83 | 2.84 | 1.55 | 2.43 | 1.82 | 1.81 | 1.7 | 1.29 | 1.05 | 1 | - |
| Syce2 | 69.22 | 72.17 | 77.93 | 43.38 | 49.03 | 46.53 | 51.96 | 52.66 | 53.01 | 32.02 | 33.42 | 30.08 | - |
| 2310008H04Rik | 11.87 | 11.13 | 14.92 | 18.77 | 15.81 | 15.12 | 18.72 | 15.88 | 17.36 | 16.32 | 15.42 | 17.98 | - |
| Mki67 | 11.55 | 11.17 | 10.92 | 8.46 | 10.8 | 10.21 | 5.76 | 6.09 | 5.95 | 5.89 | 5.99 | 5.2 | - |
| Gjc2 | 0 | 0 | 0 | 0.03 | 0.03 | 0.06 | 0.03 | 0 | 0 | 0.13 | 0.31 | 0.08 | - |
| Xpc | 4.21 | 5.24 | 4.81 | 6.34 | 6.66 | 6.93 | 8.34 | 6.94 | 8.08 | 7.56 | 8.01 | 9.51 | - |
| Ethe1 | 10.16 | 10.7 | 11.91 | 13.27 | 13.92 | 14.91 | 14.64 | 13.61 | 14.14 | 15.03 | 13.17 | 16.47 | - |
| Taf1a | 5.51 | 5.53 | 5.66 | 4.21 | 4.4 | 3.69 | 3.71 | 4 | 3.27 | 2.39 | 2.54 | 3.86 | - |
| Cyp27a1 | 0.85 | 0.65 | 0.74 | 0.23 | 0.1 | 0.37 | 1.25 | 0.87 | 0.2 | 0.03 | 0 | 0.3 | - |
| Abcg1 | 2.83 | 3.37 | 2.64 | 7.96 | 4.91 | 6.21 | 11.15 | 6 | 8.56 | 11.75 | 11.42 | 11.49 | - |
| Plxnb2 | 5.62 | 4.65 | 5.59 | 4.37 | 5.6 | 4.68 | 5.03 | 4.57 | 4.81 | 2.75 | 3.45 | 3.52 | - |
| Flt3l | 11.05 | 10.66 | 8.95 | 18.4 | 15.97 | 14.89 | 16.95 | 16.34 | 16.27 | 24.21 | 18.4 | 25.58 | - |
| Sytl3 | 44.79 | 44.88 | 49.85 | 64.02 | 65.33 | 62.04 | 54.38 | 59.03 | 63.05 | 92.7 | 86.4 | 89.29 | - |
| Tmco3 | 11.14 | 11.27 | 10.69 | 6.58 | 6.89 | 6.42 | 8.09 | 8.59 | 9.45 | 5.53 | 6.61 | 5.43 | - |
| Zc3h14 | 10.86 | 10.36 | 11.09 | 11.89 | 12.86 | 12.54 | 12.04 | 12.37 | 13.84 | 13.97 | 15.51 | 12.56 | - |
| Rab11fip1 | 2.14 | 2.93 | 3.31 | 4.95 | 5.51 | 5.86 | 5.64 | 5.16 | 4.67 | 6.05 | 8.23 | 5.9 | - |
| Tmc4 | 0.73 | 0.92 | 0.92 | 2.1 | 2.66 | 1.52 | 1.74 | 0.75 | 1.26 | 2.02 | 2.26 | 1.4 | - |
| Myl12a | 256.44 | 240.77 | 236.87 | 155.1 | 164.62 | 168.4 | 185.75 | 222.59 | 205.91 | 149.68 | 143.87 | 148.37 | - |
| Cdc14a | 2.99 | 3.67 | 2.8 | 2.23 | 3.73 | 3.1 | 2.45 | 2.44 | 2.62 | 2.47 | 1.98 | 2.03 | - |
| Stx16 | 8.94 | 7.81 | 8.84 | 16.06 | 12.82 | 12.2 | 13.19 | 12.38 | 12.32 | 16.54 | 15.69 | 17.58 | - |
| Tmem170b | 9.15 | 8.74 | 10.71 | 10.84 | 8.21 | 9.94 | 9.85 | 8.94 | 10.19 | 12.68 | 11.39 | 11.27 | - |
| E2f3 | 5.85 | 4.71 | 4.94 | 3.26 | 3.73 | 3.19 | 2.37 | 3.03 | 2.58 | 1.77 | 2.09 | 1.98 | - |
| Sdc4 | 6.85 | 5.75 | 7.78 | 15.42 | 13.42 | 14.93 | 37.79 | 19.34 | 25.11 | 29.81 | 37.29 | 37.35 | - |
| Ints6 | 2.41 | 2.78 | 3.29 | 3.8 | 3.84 | 3.34 | 4.15 | 4.24 | 3.86 | 3.65 | 4.73 | 3.71 | - |
| Add3 | 9.27 | 10.09 | 8.93 | 12.6 | 12.14 | 10.03 | 15.5 | 11.45 | 12.07 | 13.37 | 12.26 | 16.4 | - |
| Sema3f | 0.07 | 0 | 0.02 | 0.39 | 0.33 | 0.23 | 0.07 | 0.31 | 0.11 | 0.2 | 0.53 | 0.36 | - |
| Setx | 5.86 | 5.46 | 6.64 | 6.98 | 6.51 | 6.83 | 8.17 | 6.99 | 8.22 | 8.1 | 8.3 | 8.77 | - |
| Zfp949 | 1.82 | 1.48 | 1.85 | 3.24 | 3.43 | 4.09 | 4.01 | 2.89 | 2.81 | 5.21 | 6.03 | 4.4 | - |
| Pik3c3 | 13.45 | 13.39 | 12.55 | 17.01 | 15.33 | 15.97 | 18.8 | 15.58 | 15.93 | 17.81 | 18.64 | 19.06 | - |
| Rcn2 | 27.71 | 29.83 | 26.24 | 20.8 | 22.87 | 22.45 | 21.42 | 21.63 | 23.77 | 16.76 | 18.05 | 20.7 | - |
| Slc16a3 | 66.91 | 73.56 | 77.12 | 52.64 | 47.85 | 49.22 | 65.27 | 46.29 | 56.37 | 40.04 | 46.4 | 59.41 | - |
| Rbbp8 | 10.53 | 11.01 | 12.96 | 10.16 | 12.4 | 10.97 | 10.59 | 12.86 | 11.84 | 8.72 | 9.2 | 8.96 | - |
| B130006D01Rik | 0.2 | 0.49 | 0.17 | 0.57 | 0.74 | 0.7 | 0.7 | 0.48 | 0.44 | 0.84 | 0.73 | 0.76 | - |
| Gm16039 | 10.43 | 9.38 | 9.34 | 12.37 | 12.5 | 13.62 | 13.36 | 12.41 | 10.4 | 14.59 | 16.31 | 14.29 | - |
| Apc2 | 0.08 | 0.07 | 0.09 | 0.17 | 0.25 | 0.2 | 0.35 | 0.25 | 0.39 | 0.38 | 0.12 | 0.39 | - |
| Mndal | 7.27 | 10.31 | 5.13 | 17.1 | 11.77 | 11.84 | 10.91 | 3.38 | 5.64 | 18.2 | 11.03 | 17.65 | - |
| Srgap3 | 3.14 | 2.43 | 2.97 | 4.09 | 4.51 | 5.57 | 5.08 | 5.7 | 5.3 | 5.99 | 7.23 | 7.01 | - |
| Rgp1 | 2.12 | 2.04 | 2.14 | 3.84 | 2.86 | 3.36 | 3.65 | 4.1 | 3.47 | 5.11 | 5.16 | 5.2 | - |
| Dnase2a | 3.13 | 2.76 | 2.05 | 4.8 | 3.97 | 5.16 | 4.72 | 2.34 | 3.52 | 4.72 | 4.76 | 5.62 | - |
| Pex26 | 0.5 | 0.51 | 0.57 | 0.99 | 0.92 | 0.74 | 0.56 | 0.88 | 0.74 | 1.11 | 0.99 | 1.14 | - |
| Bach1 | 0.21 | 0.31 | 0.2 | 0.39 | 0.39 | 0.4 | 0.68 | 0.41 | 0.37 | 0.86 | 0.74 | 0.54 | - |
| Ar | 1.87 | 0.97 | 1.32 | 2.19 | 2.27 | 2.17 | 2.66 | 1.53 | 1.61 | 3.34 | 3.39 | 3.4 | - |
| Dcdc2c | 11.52 | 11.14 | 11.68 | 16.5 | 13.21 | 16.39 | 17.2 | 13.69 | 13.31 | 16.96 | 17.7 | 17.47 | - |
| Lonp2 | 26.63 | 24.51 | 23.99 | 33.92 | 33.44 | 31.53 | 35.13 | 30.87 | 33.48 | 35.38 | 41.2 | 42.01 | - |
| Grk6 | 6.81 | 8.56 | 6.96 | 9.21 | 7.16 | 7.57 | 9.76 | 6.89 | 9.22 | 11.46 | 8.92 | 10.66 | - |
| Sfrp2 | 0.17 | 0.61 | 0.22 | 3.85 | 1.87 | 2.49 | 0.23 | 0.44 | 0.59 | 1.49 | 0.74 | 1.3 | - |
| Tmem184b | 25.52 | 21.68 | 29.42 | 24.74 | 26.75 | 25.53 | 42.38 | 38.1 | 41.35 | 32.46 | 39.07 | 34.9 | - |
| Nrbp2 | 0.23 | 0.77 | 0.32 | 1.02 | 1 | 1.86 | 1.41 | 1.39 | 1.44 | 1.33 | 1.56 | 2.36 | - |
| Enthd1 | 1.62 | 2.07 | 2 | 1.02 | 1.39 | 1 | 1.17 | 1.22 | 1.25 | 0.68 | 0.47 | 0.82 | - |
| Pigx | 63.75 | 68.2 | 59.21 | 69.27 | 62.23 | 71.01 | 97.81 | 69.26 | 79.76 | 83.24 | 83.33 | 84.44 | - |
| Ap1g2 | 37.2 | 33.93 | 36.03 | 38 | 36.28 | 34.38 | 30.22 | 39.33 | 36.63 | 28.01 | 27.73 | 29.87 | - |
| Lmnb1 | 27.21 | 40.33 | 36.73 | 25.4 | 34.22 | 22.74 | 27.41 | 24.12 | 24.02 | 23.47 | 22.73 | 22.07 | - |
| Fam199x | 1.3 | 1.34 | 1.23 | 1.79 | 1.78 | 1.61 | 2.08 | 2.01 | 1.94 | 2.04 | 2.55 | 1.47 | - |
| Lig3 | 9.77 | 9.06 | 9.16 | 7.29 | 6.36 | 6.39 | 4.56 | 5.5 | 6.94 | 5.41 | 4.45 | 5 | - |
| Zfp511 | 22.78 | 22.69 | 21.63 | 22.47 | 21.05 | 22.84 | 30.89 | 25.1 | 27.22 | 26.11 | 31.32 | 30.5 | - |
| Ostf1 | 319.92 | 285.83 | 322.88 | 450.53 | 461.56 | 474.22 | 336.7 | 369.5 | 362.13 | 431.29 | 415 | 448.88 | - |
| Inip | 15.61 | 15.01 | 14.15 | 12.34 | 11.81 | 14.37 | 12.58 | 14.16 | 13.34 | 11.99 | 12.84 | 11.18 | - |
| Agpat5 | 17.84 | 18.98 | 16.41 | 14.51 | 15.8 | 16.59 | 12.56 | 12.38 | 13.72 | 9.99 | 13.77 | 11.32 | - |
| Bahd1 | 2.57 | 2.08 | 2.08 | 3.41 | 2.97 | 2.77 | 2.93 | 3.09 | 2.81 | 3.33 | 3.2 | 4.15 | - |
| Kdelc1 | 19.07 | 17.58 | 18.47 | 14.09 | 16.38 | 16.82 | 12.41 | 16.23 | 15.06 | 10.09 | 10.42 | 9.08 | - |
| Cd28 | 19.54 | 18.14 | 18.29 | 31.98 | 33.89 | 35.83 | 48.93 | 37.2 | 39.84 | 61.19 | 60.85 | 56.07 | - |
| Rnf180 | 0.42 | 0.32 | 0.26 | 0.05 | 0.18 | 0.22 | 0 | 0.16 | 0.1 | 0.03 | 0.05 | 0 | - |
| Lamp2 | 43.07 | 37.73 | 45.36 | 61.99 | 62.85 | 68.02 | 44.64 | 37.81 | 39.27 | 58.02 | 63.59 | 54.67 | - |
| H2-D1 | 1104.49 | 1086.82 | 1045.01 | 1838.53 | 1672.35 | 1715.77 | 1537.76 | 1380 | 1427.11 | 1929.91 | 1876.45 | 1968.77 | - |
| Meaf6 | 14.79 | 16.62 | 13.69 | 12.19 | 13.11 | 13.9 | 10.89 | 12.07 | 13.05 | 9.14 | 10.8 | 10.53 | - |
| Mxd3 | 9.27 | 7.68 | 5.68 | 5.57 | 7.14 | 7.15 | 5.74 | 4.04 | 4.15 | 4.96 | 5 | 3.79 | - |
| Ubl7 | 46.32 | 45.72 | 48.31 | 55.9 | 54.41 | 55.59 | 74.96 | 63.83 | 65.42 | 71.01 | 70.13 | 74.15 | - |
| Irf1 | 24.02 | 27.85 | 23.09 | 38.3 | 28.94 | 27.47 | 54.74 | 27.92 | 30.67 | 71.44 | 57.43 | 64.91 | - |
| Ski | 2.67 | 2.62 | 2.27 | 3.4 | 3.21 | 2.81 | 2.93 | 2.92 | 2.73 | 3.42 | 3.63 | 3.42 | - |
| Aph1a | 20.15 | 20.15 | 21.04 | 22.77 | 20.26 | 19.84 | 24.69 | 19.92 | 20.18 | 25.5 | 23.72 | 27.23 | - |
| Ppm1f | 9.19 | 9.47 | 9.91 | 8.37 | 9.05 | 6.15 | 5.84 | 7.43 | 6.85 | 7.25 | 8.33 | 5.7 | - |
| Aebp1 | 1.69 | 1.62 | 2.18 | 1.31 | 1.77 | 1.46 | 0.16 | 0.59 | 0.58 | 0.13 | 0.29 | 0.21 | - |
| Stard3 | 10.6 | 10.11 | 10.74 | 13.21 | 11.51 | 12.83 | 13.49 | 14.24 | 13.89 | 15.68 | 15.24 | 12.29 | - |
| Hexb | 25.62 | 23.24 | 24.03 | 36.19 | 32.34 | 34.11 | 48.93 | 31.08 | 39.46 | 41.1 | 39.94 | 43.88 | - |
| Ankrd44 | 10.04 | 9.16 | 8.37 | 10.6 | 10.01 | 9.72 | 11.69 | 9.03 | 11.18 | 11.83 | 11.78 | 11.96 | - |
| Xlr4c | 20.46 | 22.15 | 19.89 | 18.15 | 15.98 | 15.9 | 15.64 | 13.14 | 16.6 | 11.82 | 15.33 | 11.86 | - |
| Enpp4 | 4.25 | 3.92 | 5.21 | 2.01 | 2.88 | 1.88 | 1.94 | 2.6 | 3.27 | 0.97 | 0.68 | 0.95 | - |
| Smcr7l | 8.99 | 10.1 | 9.5 | 7.95 | 6.68 | 6.63 | 7.08 | 6.36 | 6.93 | 5.18 | 5.53 | 5.24 | - |
| Cox20 | 32.88 | 35.59 | 31.86 | 38.23 | 40.41 | 42.83 | 59.52 | 47.33 | 45.99 | 59.6 | 67.82 | 52.22 | - |
| Fam72a | 3.66 | 3.07 | 2.47 | 1.8 | 2.79 | 1.27 | 1.64 | 1.4 | 2.17 | 1.74 | 1.35 | 1.56 | - |
| Endod1 | 6 | 8.36 | 6.63 | 4 | 3.61 | 4.67 | 6.74 | 5.32 | 6.49 | 5.29 | 3.7 | 4.22 | - |
| Gaa | 23.28 | 20.65 | 22.49 | 22.43 | 18.93 | 21.38 | 17 | 19.07 | 19.22 | 18.42 | 18.87 | 17.63 | - |
| Ddx20 | 27.43 | 25.19 | 23.44 | 14.25 | 19.39 | 19.28 | 16.71 | 15.3 | 14.83 | 11.63 | 13.05 | 12.78 | - |
| 9030624J02Rik | 16.4 | 14.59 | 16.48 | 12.17 | 13.75 | 13.67 | 15.13 | 13.96 | 15.18 | 12.51 | 11.61 | 13.2 | - |
| Zfp869 | 11.4 | 10.97 | 11.76 | 18.05 | 15.65 | 14.88 | 15.95 | 13.46 | 13.93 | 17.35 | 15.68 | 14.98 | - |
| Hgsnat | 2.56 | 3.69 | 3.23 | 5.03 | 4.87 | 3.93 | 4.89 | 5.39 | 5.29 | 6.75 | 5.24 | 6.46 | - |
| Brdt | 0.22 | 0.22 | 0.21 | 0.2 | 0.2 | 0.37 | 0.35 | 0.26 | 0.44 | 0.68 | 0.51 | 0.65 | - |
| Raph1 | 1.43 | 1.78 | 1.59 | 1.71 | 1.89 | 2.79 | 1.95 | 2.15 | 1.79 | 2.68 | 2.92 | 2.66 | - |
| Cenpj | 3.91 | 3.67 | 3.57 | 2.84 | 3.03 | 2.78 | 2.77 | 2.53 | 3.32 | 2.9 | 2.15 | 2.77 | - |
| Yipf3 | 41.35 | 41.23 | 41.21 | 59.17 | 55.06 | 57.35 | 64.52 | 55.33 | 60.77 | 73.01 | 62.27 | 72.51 | - |
| Dnm2 | 39.47 | 37.82 | 40.69 | 44.93 | 49 | 45 | 52.38 | 50.51 | 47.01 | 53.63 | 57.01 | 52.38 | - |
| Otx1 | 0.16 | 0.1 | 0.19 | 0 | 0.22 | 0.06 | 0 | 0 | 0 | 0 | 0 | 0 | - |
| Il6 | 2.05 | 1.28 | 3.68 | 0.06 | 0.17 | 0.12 | 0.05 | 0 | 0 | 0 | 0 | 0 | - |
| 4930511M06Rik | 8.71 | 9.31 | 9.66 | 14.97 | 12.96 | 14.23 | 12.97 | 13.48 | 12.4 | 17.31 | 16.11 | 16.68 | - |
| Thra | 0.43 | 0.34 | 0.53 | 1.15 | 0.78 | 1.28 | 1.33 | 1.02 | 0.96 | 1.45 | 2.22 | 1.05 | - |
| Cers4 | 5.09 | 5.45 | 6.25 | 8.45 | 6.81 | 8.36 | 9.43 | 7.17 | 8.51 | 10.86 | 10.77 | 12.46 | - |
| 5830418P13Rik | 0.47 | 0.59 | 0.55 | 0.84 | 0.57 | 0.85 | 3.06 | 1.81 | 2.7 | 2.13 | 1.46 | 3.12 | - |
| Ptpmt1 | 18.46 | 20.27 | 19.65 | 16.84 | 15.5 | 17.99 | 14.34 | 18.89 | 17.5 | 12.2 | 13.16 | 15.45 | - |
| Tada3 | 18.4 | 17.59 | 18.69 | 15.69 | 16.21 | 17.52 | 15.5 | 15.63 | 12.94 | 12.22 | 14.67 | 14.87 | - |
| Pram1 | 0.3 | 0.28 | 0.05 | 0.22 | 0.36 | 0.25 | 0.28 | 0.31 | 0.35 | 0.81 | 0.54 | 0.96 | - |

| | | | | | | | | | | | | | |
|---|---|---|---|---|---|---|---|---|---|---|---|---|---|
| Wdr89 | 8.06 | 8.49 | 9.15 | 4.75 | 5.09 | 4.77 | 2.99 | 5.07 | 5.03 | 1.8 | 2.07 | 2.56 | - |
| Pias2 | 3.82 | 4.08 | 4.45 | 7.39 | 8.65 | 8.2 | 4.19 | 5.58 | 4.2 | 7.93 | 7.82 | 5.92 | - |
| Bckdk | 34.88 | 37.7 | 37.47 | 22.27 | 24.3 | 24.83 | 21.16 | 23.58 | 19.63 | 16.85 | 12.86 | 18.01 | - |
| Ibtk | 6.56 | 6.12 | 7.72 | 3.84 | 4.92 | 5.23 | 4.06 | 5.49 | 5.45 | 3.53 | 3.69 | 3.48 | - |
| Gab3 | 2.23 | 3.28 | 1.86 | 4.17 | 3.59 | 4.14 | 6.52 | 4.59 | 4 | 8.07 | 5.8 | 7.87 | - |
| Nsl1 | 10.9 | 11.85 | 11.53 | 10.26 | 12.67 | 11.79 | 7.42 | 8.61 | 7.7 | 7.28 | 8.06 | 8.1 | - |
| Apoe | 4.65 | 3.12 | 4.49 | 2.68 | 2.84 | 3.42 | 2.16 | 2.96 | 2.27 | 1.89 | 1.28 | 2.86 | - |
| Mina | 72.03 | 72.24 | 71.41 | 49.57 | 54.15 | 52.96 | 56.95 | 65.82 | 60.39 | 51.05 | 42.75 | 44.77 | - |
| Orc1 | 17.29 | 17.56 | 19.8 | 7.49 | 6.84 | 7.79 | 2.92 | 6.4 | 5.5 | 1.42 | 2.09 | 2.47 | - |
| Kntc1 | 2.7 | 2.32 | 3.12 | 1.68 | 2.17 | 1.65 | 0.84 | 1.05 | 1.39 | 0.91 | 0.6 | 0.7 | - |
| Rrad | 3.95 | 7.17 | 7.62 | 10.67 | 9.47 | 8.38 | 11.75 | 9.43 | 11.24 | 14.71 | 11.82 | 12.87 | - |
| Paqr7 | 7.24 | 6.73 | 7.74 | 17.64 | 15.95 | 17.03 | 25.6 | 15.83 | 16.43 | 30.39 | 28.54 | 33.07 | - |
| Hivep1 | 4.89 | 4.27 | 5.02 | 4.24 | 5.22 | 6.05 | 6.83 | 6.43 | 6.61 | 6.2 | 6.4 | 6.67 | - |
| Cnn2 | 143.81 | 165.18 | 140.75 | 226.33 | 186.39 | 197.36 | 316.07 | 226.07 | 244.83 | 315.18 | 273.15 | 344.24 | - |
| Wisp1 | 0.47 | 0.79 | 0.92 | 0.67 | 0.7 | 0.6 | 0 | 0.23 | 0.03 | 0.26 | 0.09 | 0.15 | - |
| Bend5 | 0.39 | 0.4 | 0.51 | 2.13 | 1.55 | 1.85 | 2.85 | 0.81 | 1.08 | 1.9 | 2.7 | 3.47 | - |
| Rnf2 | 12.44 | 11.9 | 12.82 | 18.01 | 14.53 | 15.52 | 13.23 | 14.43 | 12.51 | 14.74 | 17.36 | 15.48 | - |
| Il27ra | 20.32 | 22.02 | 20.95 | 26.92 | 23.74 | 25.65 | 29.49 | 23.32 | 25.78 | 28.9 | 28.54 | 30.69 | - |
| Tspan6 | 14.76 | 10.53 | 12.82 | 10.68 | 15.9 | 14.59 | 7.04 | 7.46 | 7.76 | 8.76 | 8.46 | 8.21 | - |
| Zfp784 | 0.53 | 0.54 | 0.53 | 1.8 | 1.6 | 1.1 | 1.98 | 1.45 | 0.95 | 0.83 | 1.23 | 3.04 | - |
| Rad1 | 26.76 | 23.72 | 26.59 | 20.06 | 20.87 | 21.36 | 16.49 | 18.96 | 19.37 | 15.11 | 16.81 | 17.15 | - |
| Edem2 | 91.03 | 78.42 | 90.98 | 41.38 | 43.8 | 45.97 | 60.12 | 71.9 | 71.02 | 41.22 | 38.22 | 40.29 | - |
| H2-K2 | 7.03 | 5.19 | 5.47 | 15.68 | 11.85 | 14.72 | 18.46 | 10.86 | 12.83 | 21.49 | 25.34 | 28.06 | - |
| Srr | 2.81 | 2.71 | 2.97 | 5.69 | 5.09 | 4.48 | 4.55 | 5.51 | 3.82 | 5.46 | 4.05 | 7.45 | - |
| Slc37a4 | 18.1 | 20.9 | 17.5 | 37.89 | 31.2 | 33.67 | 49.94 | 34.79 | 38.42 | 48.1 | 56.44 | 54.73 | - |
| Shd | 0.04 | 0 | 0.07 | 0.33 | 0.16 | 0.17 | 0.3 | 0.46 | 0.21 | 0.04 | 1.37 | 0.45 | - |
| Dyrk3 | 1.66 | 2.3 | 2.17 | 5.89 | 3.28 | 3.4 | 3.8 | 3.23 | 4.3 | 5.27 | 5.18 | 3.62 | - |
| Csda | 17.46 | 20.99 | 18.7 | 16.98 | 14.32 | 15.69 | 19.02 | 22.1 | 22.16 | 13.84 | 15.06 | 13.88 | - |
| Tsr1 | 49.86 | 52.96 | 54.28 | 31.95 | 33.01 | 34.71 | 21.04 | 36.96 | 35.05 | 26.1 | 24.21 | 25.66 | - |
| Cnnm4 | 5.92 | 7.39 | 6.65 | 7.76 | 6.87 | 6.71 | 5.48 | 5.08 | 6.11 | 5.19 | 4.88 | 4.76 | - |
| Fn3k | 0.06 | 0 | 0 | 0.63 | 0.31 | 1.14 | 1.6 | 0.18 | 0.85 | 2.5 | 1.23 | 1.95 | - |
| Guk1 | 106.7 | 105.7 | 94.7 | 73.08 | 68.04 | 77.27 | 79.52 | 95.93 | 88.56 | 66.96 | 58.99 | 60.58 | - |
| Tmem55b | 2.38 | 2.92 | 2.23 | 3.64 | 2.74 | 4.44 | 2.67 | 3.44 | 3.89 | 4.34 | 3.87 | 4.88 | - |
| Taf2 | 3.29 | 4.03 | 4.82 | 3.5 | 3.29 | 2.85 | 3.24 | 3.24 | 3.5 | 2.96 | 2.91 | 2.15 | - |
| Slc1a5 | 112.63 | 118.3 | 114.15 | 88.99 | 98.47 | 93.29 | 80.55 | 95.09 | 100.51 | 78.9 | 73.72 | 78.56 | - |
| Dtx3l | 12.52 | 13.97 | 14.05 | 21.14 | 17.72 | 18.94 | 22.15 | 14.73 | 17.23 | 31.72 | 27.98 | 32.74 | - |
| Gm15800 | 2.19 | 2.24 | 1.9 | 4.05 | 3.88 | 3.81 | 4.67 | 3.35 | 3.7 | 6.6 | 6.47 | 6.43 | - |
| BC002163 | 127.87 | 131.63 | 128.28 | 85.01 | 79.86 | 91.56 | 67.71 | 114.86 | 96.77 | 69.5 | 66.25 | 61.45 | - |
| Lrrc58 | 10.11 | 10.81 | 11.61 | 8.98 | 10.05 | 9.85 | 6.51 | 7.69 | 7.97 | 7.52 | 7.75 | 7.23 | - |
| Parvg | 69.12 | 79.94 | 73.41 | 57.27 | 56.93 | 59.4 | 70.78 | 69.07 | 74.08 | 51.09 | 51.79 | 53.58 | - |
| Wdr11 | 9.4 | 9.57 | 10.23 | 6.97 | 6.82 | 7.33 | 8.02 | 7.47 | 8.69 | 7.33 | 5.74 | 5.78 | - |
| Uimc1 | 17.51 | 17.69 | 17.8 | 19.11 | 19.57 | 21.23 | 20.86 | 21.08 | 21.38 | 25.17 | 24.9 | 23.03 | - |
| Pglyrp1 | 5.62 | 9.23 | 2.78 | 27.66 | 12.1 | 12.74 | 90.04 | 15.18 | 48.76 | 72.08 | 62.17 | 93.02 | - |
| Bora | 16 | 11.95 | 12.54 | 9.21 | 10.8 | 10.54 | 9.12 | 7.71 | 8.14 | 8.31 | 8.28 | 6.64 | - |
| Ppp1r15b | 11.48 | 12.24 | 12.47 | 10.67 | 9.23 | 9.51 | 9.59 | 10.17 | 9.98 | 8.38 | 8.68 | 7.97 | - |
| 4930455F23Rik | 7.87 | 6.32 | 7.61 | 6.9 | 9.25 | 8.95 | 11.88 | 9.85 | 10.78 | 10.95 | 10.04 | 10.36 | - |
| Dgkz | 10.85 | 11.71 | 11.64 | 14.59 | 13.35 | 14.16 | 18.84 | 15.74 | 14.64 | 16.39 | 19.77 | 20.79 | - |
| H2-T9 | 127.05 | 118.67 | 117.85 | 189.61 | 177.28 | 184.14 | 244.38 | 186.85 | 196.82 | 258.81 | 295.81 | 276.54 | - |
| Arhgap15 | 33.99 | 29.35 | 31.63 | 73.73 | 62.13 | 65.69 | 74.62 | 54.23 | 60.41 | 98.1 | 87.59 | 97.43 | - |
| Wdr96 | 0.04 | 0.05 | 0.03 | 0.04 | 0.17 | 0.05 | 0.02 | 0.07 | 0.07 | 0.13 | 0.25 | 0.18 | - |
| Ctnnd1 | 2.41 | 1.18 | 2.32 | 0.48 | 0.69 | 0.66 | 0.46 | 1.03 | 0.88 | 0.51 | 0.42 | 0.26 | - |
| Fam212b | 1.19 | 1.09 | 1.36 | 0.88 | 1.6 | 0.8 | 1.18 | 0.66 | 0.76 | 0.49 | 0.83 | 0.65 | - |
| Itfg3 | 6.59 | 7.06 | 7.11 | 16.86 | 12.48 | 14.15 | 26.41 | 14.83 | 18.3 | 22.43 | 26.57 | 28.21 | - |
| AI413582 | 59.69 | 56.45 | 53.65 | 80.47 | 74.67 | 70.18 | 99.32 | 87.23 | 84.9 | 101.58 | 102.14 | 107.5 | - |
| Lrp10 | 16.93 | 19.57 | 18.58 | 23.98 | 24.21 | 21.47 | 26.39 | 26.13 | 23.96 | 33.45 | 27.56 | 28.19 | - |
| Phf3 | 3.16 | 3.16 | 3.86 | 4.17 | 4.57 | 4.13 | 4.76 | 4.45 | 4.57 | 5.14 | 5.31 | 4.58 | - |
| Mll1 | 3.48 | 3.57 | 3.41 | 5.22 | 5.2 | 5 | 4.59 | 4.57 | 4.27 | 6.24 | 6.38 | 5.94 | - |
| Gm8369 | 25.75 | 27.73 | 28.06 | 20.34 | 17.72 | 19.31 | 22.02 | 19.87 | 19.89 | 17.09 | 11.17 | 18.63 | - |
| Serpinb6b | 309.36 | 208.12 | 249.5 | 851.87 | 781.48 | 834.01 | 276.93 | 399.22 | 338.73 | 869.06 | 888.07 | 913.96 | - |
| Acsl1 | 5.93 | 6.33 | 5.02 | 3.74 | 3.98 | 4.39 | 3.93 | 4.48 | 5.27 | 3.4 | 3.33 | 2.46 | - |
| Ebi3 | 33.98 | 29.55 | 25.44 | 40.53 | 36.41 | 34.75 | 22.57 | 18.26 | 19.29 | 18.17 | 21.16 | 18.8 | - |
| Ppp3cb | 3.54 | 4.27 | 4.01 | 5.15 | 4.58 | 4.3 | 5.17 | 4.24 | 4.67 | 6.19 | 4.65 | 6.34 | - |
| Ccdc22 | 12.58 | 12.55 | 12.79 | 10.42 | 11.63 | 7.49 | 10.72 | 10.8 | 10.8 | 10.62 | 7.65 | 7.95 | - |
| Agap2 | 0.94 | 0.96 | 0.86 | 1.4 | 0.98 | 0.96 | 1.24 | 0.91 | 1.13 | 1.48 | 1.29 | 1.81 | - |
| Slc24a6 | 2.95 | 2.01 | 2.25 | 3.86 | 3.22 | 3.4 | 5.25 | 4.05 | 5.19 | 5.15 | 5.7 | 4.4 | - |
| Prkacb | 14.39 | 16.58 | 15.87 | 26.71 | 19.55 | 23.77 | 33.08 | 20.53 | 26.79 | 38.07 | 32.41 | 33.02 | - |
| Ap2b1 | 19.25 | 19.18 | 18.85 | 16.73 | 16.14 | 17.53 | 15.54 | 19.88 | 17.44 | 12.53 | 15.8 | 13.82 | - |
| B3galt5 | 0.01 | 0.11 | 0.06 | 0.21 | 0.1 | 0.21 | 0 | 0 | 0.03 | 0.54 | 0.2 | 0.25 | - |
| Tceb2 | 270.95 | 272.61 | 273.74 | 203.29 | 223.15 | 232.48 | 233.25 | 262.66 | 236.67 | 191.44 | 202.12 | 195.05 | - |
| Pced1b | 5.81 | 6.69 | 7.08 | 8.36 | 7.32 | 6.99 | 8.15 | 7.48 | 7.85 | 8.81 | 8.85 | 9.38 | - |
| Kremen2 | 0.47 | 0.4 | 0.48 | 1.1 | 0.83 | 0.71 | 2.15 | 1.73 | 1.2 | 1.61 | 1.07 | 2.08 | - |
| Faah | 13.33 | 13.05 | 12.61 | 17.14 | 14.43 | 16.39 | 18.9 | 13.5 | 15.2 | 18.15 | 14.34 | 19.38 | - |
| Armcx6 | 3.65 | 2.98 | 3.99 | 4.69 | 3.28 | 3.45 | 10.4 | 7.83 | 9.55 | 8.41 | 11.65 | 9.92 | - |
| Zyx | 26.93 | 26.92 | 25.25 | 40.67 | 32.7 | 33.51 | 49.05 | 35.75 | 35.76 | 57.91 | 46.46 | 56.65 | - |
| Nkd2 | 0.16 | 0.25 | 0.1 | 0.08 | 0.08 | 0.36 | 0.14 | 0.23 | 0.2 | 0.02 | 0 | 0 | - |
| Ncoa7 | 2.21 | 2.49 | 2.98 | 2.7 | 3.33 | 2.71 | 0.81 | 1.15 | 1.2 | 0.96 | 1.19 | 1.44 | - |
| Ugt1a6b | 6.57 | 3.53 | 5.91 | 0 | 1.35 | 0.73 | 0 | 0.37 | 0.9 | 0 | 0 | 0.05 | - |
| Nr2c1 | 2.13 | 3.22 | 2.87 | 2.91 | 2.57 | 2.8 | 1.81 | 2.06 | 2.13 | 1.93 | 1.21 | 1.88 | - |
| Papd7 | 3.16 | 3.58 | 3.98 | 2.5 | 2.31 | 2.51 | 2.06 | 1.82 | 2.01 | 2.02 | 1.79 | 0.92 | - |
| Rreb1 | 1.65 | 0.97 | 1.17 | 1.86 | 1.96 | 2.33 | 2.24 | 1.67 | 1.74 | 2.93 | 2.65 | 2.44 | - |
| Birc3 | 20.42 | 21.14 | 21.87 | 22.61 | 25.99 | 24.5 | 26.92 | 26.86 | 26.64 | 25.42 | 26.33 | 27.25 | - |
| Morc3 | 14.52 | 13.98 | 15.72 | 12.02 | 13.5 | 12.48 | 13.13 | 10.41 | 11.67 | 12.16 | 11.43 | 10.72 | - |
| Ier3 | 44.06 | 36.25 | 43.35 | 33.05 | 37.8 | 33.7 | 28.63 | 23.92 | 25.79 | 20.15 | 18.83 | 19.83 | - |
| Smad3 | 2.91 | 3.06 | 2.79 | 3.87 | 3.63 | 3.36 | 5.59 | 5.86 | 5.07 | 5.33 | 5.09 | 5.87 | - |
| Cyba | 285.09 | 309.33 | 275.07 | 247.9 | 254.06 | 253.75 | 286.42 | 265.5 | 284.65 | 207.33 | 204.46 | 220.78 | - |
| Ddc | 3.64 | 4.16 | 4.67 | 4.52 | 4.6 | 4.5 | 2.38 | 2.16 | 3.24 | 2.11 | 1.45 | 2.12 | - |
| Gpt | 1.34 | 1.1 | 2.39 | 3.07 | 3.96 | 3.32 | 4.6 | 2.58 | 3.31 | 4.77 | 3.8 | 3.03 | - |
| Lum | 53.53 | 29.42 | 44.06 | 20.68 | 26.68 | 30.36 | 6.56 | 16.93 | 8.35 | 10.28 | 11.17 | 9.5 | - |
| Naa16 | 14.74 | 15.71 | 15.74 | 14.7 | 15.97 | 13.29 | 12.15 | 13.91 | 13.38 | 11.34 | 12.75 | 12.48 | - |
| Trak1 | 4 | 3.37 | 3.98 | 4.01 | 4.6 | 4.51 | 5.02 | 5.94 | 5.72 | 6.63 | 6 | 5.1 | - |
| Asb1 | 1.92 | 1.28 | 1.38 | 0.95 | 0.9 | 0.86 | 0.98 | 1.29 | 1.39 | 1 | 0.83 | 0.75 | - |
| Zfp652 | 0.96 | 1.53 | 0.87 | 2.56 | 2.32 | 2.27 | 2.33 | 2.2 | 1.97 | 3.99 | 3.33 | 3.07 | - |
| Megf9 | 0.26 | 0.42 | 0.67 | 1.18 | 0.82 | 0.87 | 1.33 | 0.65 | 0.77 | 1.77 | 0.95 | 0.73 | - |
| Ap3m1 | 26.66 | 26.27 | 24.57 | 23.37 | 22.32 | 23.99 | 22.71 | 25.35 | 25.46 | 20.94 | 20.22 | 20.07 | - |
| Hrc | 0.87 | 0.92 | 0.53 | 0.21 | 0.13 | 0.16 | 0.78 | 0.8 | 0.55 | 0.19 | 0.29 | 0.15 | - |
| Spint1 | 1.46 | 1.43 | 1.95 | 0.81 | 1.1 | 0.87 | 1.16 | 1.73 | 1.16 | 0.48 | 0.68 | 0.56 | - |
| Tnfrsf1a | 7.34 | 7.7 | 7.58 | 9.56 | 9.76 | 8.84 | 12.64 | 9.81 | 10.84 | 11.02 | 8.72 | 10.97 | - |
| Zfp429 | 4.65 | 5.17 | 4.75 | 8.47 | 9.66 | 9.23 | 8.34 | 7.09 | 7.5 | 10.21 | 13.52 | 12.03 | - |
| Cttn | 0.88 | 0.79 | 0.89 | 0.87 | 0.74 | 0.74 | 0.52 | 0.83 | 0.69 | 0.17 | 0.46 | 0.35 | - |
| Tcf3 | 6.33 | 6.31 | 7.68 | 5.02 | 5.92 | 4.05 | 5.63 | 5.27 | 4.42 | 5.2 | 4.49 | 5.05 | - |
| Ndufv2 | 129.11 | 137.52 | 133.81 | 117.81 | 126.24 | 121.39 | 115.78 | 117.66 | 118.02 | 104.3 | 110.78 | 109.29 | - |
| Zfp871 | 4.36 | 4.47 | 4.66 | 6.72 | 5.98 | 5.75 | 5.38 | 5.4 | 5.67 | 7.31 | 7.79 | 6.7 | - |
| Ddx17 | 23.15 | 21.42 | 22.02 | 32.16 | 31.74 | 30.76 | 31.74 | 29.58 | 29.71 | 35.13 | 32.41 | 36.1 | - |
| Gstp1 | 235.14 | 248.04 | 232.42 | 379.54 | 354.95 | 374.53 | 222.77 | 217.94 | 225.06 | 309.39 | 278.98 | 337.82 | - |
| Relt | 11.79 | 14.69 | 11.69 | 18.96 | 17.59 | 17.36 | 16.22 | 17.64 | 18.32 | 18.46 | 16.63 | 20.12 | - |
| Ube2q1 | 4.37 | 4.09 | 5.04 | 3.41 | 3.48 | 3.45 | 3.1 | 3.07 | 3.22 | 2.69 | 2.86 | 2.79 | - |
| Lrrc8a | 3.45 | 3.69 | 3.57 | 6.48 | 5.83 | 6.07 | 6.41 | 4.21 | 4.79 | 5.92 | 5.21 | 6.76 | - |
| Osgin1 | 2.04 | 2.4 | 1.16 | 2.51 | 1.78 | 1.72 | 9.03 | 5.12 | 6.48 | 7.5 | 5.94 | 8.82 | - |
| Otud1 | 0.02 | 0 | 0.02 | 0.12 | 0.15 | 0.1 | 0.2 | 0.16 | 0.11 | 0.29 | 0.22 | 0.21 | - |
| Ms4a4b | 249.02 | 314.01 | 225.31 | 172.29 | 153.08 | 165.77 | 184.31 | 116.05 | 126.46 | 78.5 | 51.95 | 102.05 | - |
| Ankrd54 | 13.86 | 16.62 | 18.6 | 25.58 | 26.33 | 20.88 | 29.64 | 24.39 | 25.48 | 27.32 | 29.31 | 27.35 | - |
| Slc46a3 | 2.43 | 1.99 | 2.72 | 4.02 | 2.94 | 2.69 | 4.44 | 2.98 | 3.32 | 4.18 | 3.97 | 3.69 | - |
| Trim7 | 0.8 | 0.43 | 0.8 | 0.94 | 1.65 | 1.33 | 1.67 | 1.77 | 1.65 | 1.72 | 1.63 | 2.13 | - |
| Slc52a2 | 10.55 | 11.48 | 13.5 | 8.47 | 10.57 | 9.16 | 7.28 | 10.03 | 8.88 | 6.29 | 6.35 | 7.39 | - |
| Ywhaq | 158.46 | 165.25 | 161.39 | 115.62 | 120.72 | 124.02 | 77.91 | 116.94 | 109.59 | 64.21 | 77.87 | 74.8 | - |
| 2700049A03Rik | 2.4 | 2.34 | 2.25 | 3.58 | 4.19 | 3.61 | 3.31 | 2.51 | 3.16 | 3.72 | 3.66 | 3.57 | - |
| Upb1 | 4.22 | 3.45 | 4.33 | 1.15 | 1.48 | 1.24 | 0.84 | 2.01 | 1.83 | 0.92 | 0.69 | 0.88 | - |
| Ubox5 | 7.9 | 8.38 | 8.07 | 6.91 | 6.43 | 5.48 | 4.89 | 7.04 | 6.19 | 6.16 | 4.94 | 6.3 | - |
| Slc11a2 | 13.76 | 11.16 | 12.43 | 22.17 | 18.23 | 20.09 | 15.78 | 13.7 | 14.15 | 23.31 | 18.7 | 19.41 | - |
| Fgd3 | 0.95 | 1.37 | 1.44 | 1.61 | 1.43 | 2.05 | 2.55 | 2.04 | 2.47 | 3.24 | 3.49 | 3.84 | - |
| Bcar1 | 1.23 | 1.03 | 1.04 | 2.23 | 3.06 | 2.6 | 2.93 | 1.38 | 2.19 | 2.16 | 3.18 | 3.44 | - |
| Ahcyl2 | 6.22 | 5.14 | 6.04 | 5.01 | 4.33 | 4.41 | 5.04 | 6.52 | 6.25 | 4.18 | 4.14 | 4.57 | - |
| Dus4l | 7.62 | 8.4 | 8.64 | 4.11 | 5.86 | 2.96 | 2.37 | 5.68 | 4.67 | 2.69 | 3.87 | 2.38 | - |
| Tatdn1 | 12.42 | 10.14 | 9.54 | 13.46 | 15.35 | 13.77 | 15.39 | 14.58 | 12.85 | 14.87 | 15.57 | 15.28 | - |
| Usp20 | 4.48 | 3.56 | 3.81 | 5.69 | 5.42 | 4.4 | 5.63 | 4.97 | 6.01 | 5.02 | 5.99 | 6.2 | - |
| Dysf | 0.01 | 0 | 0.01 | 0.18 | 0.32 | 0.37 | 0.84 | 0.23 | 0.65 | 0.43 | 0.86 | 0.59 | - |
| Ret | 0 | 0 | 0 | 0 | 0 | 0.01 | 0.03 | 0 | 0.02 | 0.07 | 0.03 | 0.16 | - |
| 1700010I14Rik | 2.19 | 1.84 | 2.55 | 3.13 | 3.79 | 3.12 | 4.1 | 3.66 | 4 | 4.45 | 4.4 | 5.59 | - |
| Tnfaip1 | 17 | 14.64 | 16.29 | 13.75 | 12.24 | 15.63 | 13.37 | 14.23 | 12.05 | 12.61 | 12.52 | 9.53 | - |
| Yaf2 | 13.04 | 13.87 | 14.18 | 7.82 | 10.9 | 10.9 | 8.66 | 14.47 | 12.01 | 9.66 | 10.47 | 7.05 | - |
| Qsox2 | 2.73 | 3.07 | 4 | 2.12 | 1.78 | 1.59 | 1.13 | 2.83 | 1.84 | 1.07 | 0.83 | 0.21 | - |
| Srl | 0.52 | 0.46 | 0.6 | 0.15 | 0.23 | 0.32 | 0.17 | 0.29 | 0.33 | 0.21 | 0.16 | 0.19 | - |
| 7-Mar | 19.63 | 19.13 | 20.55 | 30.43 | 30.33 | 27.37 | 35.15 | 28.54 | 29.24 | 35.97 | 38.9 | 31.06 | - |
| Trmt61a | 7.23 | 7.15 | 7.13 | 4.03 | 2.86 | 3.58 | 2.11 | 4.1 | 3.01 | 1.57 | 2.06 | 1.48 | - |
| Gm14308 | 5.1 | 5.43 | 4.79 | 8.68 | 5.84 | 7.07 | 9.89 | 8.51 | 6.62 | 8.13 | 12.07 | 10.54 | - |
| Arl14ep | 61.13 | 55.8 | 60.97 | 102.63 | 122.14 | 118.87 | 55.69 | 75.14 | 74.09 | 110.49 | 131.34 | 115.51 | - |
| Foxm1 | 4.8 | 4.07 | 3.42 | 2.91 | 5.63 | 5.57 | 2.62 | 2.23 | 1.88 | 2.23 | 2.37 | 2.06 | - |
| Il6st | 1.15 | 1.05 | 0.82 | 2.02 | 1.98 | 1.54 | 1.88 | 1.11 | 1.56 | 1.94 | 2.19 | 2.14 | - |
| Syn1 | 0.31 | 0.88 | 0.68 | 0.28 | 0.12 | 0.18 | 0.2 | 1.05 | 0.76 | 0.11 | 0.16 | 0.07 | - |
| Itpkb | 0.87 | 0.71 | 0.73 | 1.31 | 1.19 | 1.31 | 1.28 | 1.09 | 1.09 | 1.47 | 1.45 | 1.66 | - |
| Calu | 63.27 | 61.4 | 67.64 | 45.44 | 55.35 | 54.44 | 44.16 | 59.49 | 52.16 | 41.04 | 47.44 | 36.63 | - |
| Casp2 | 10.38 | 10.6 | 12.48 | 13.82 | 14.67 | 12.04 | 11.55 | 12.08 | 11.88 | 16.6 | 13.08 | 17.25 | - |
| Phip | 5.39 | 4.83 | 4.81 | 5.79 | 6.56 | 6.44 | 6.11 | 5.49 | 5.85 | 6.69 | 8.38 | 7.26 | - |
| Baat1 | 0.58 | 0.53 | 0.19 | 0.29 | 0.3 | 0.22 | 0.06 | 0.08 | 0.12 | 0.13 | 0 | 0.12 | - |
| Eif2a | 71.06 | 72.27 | 72.65 | 92.36 | 82.52 | 88.23 | 111.64 | 91.43 | 95.72 | 118.62 | 134.58 | 124.59 | - |
| Ipo5 | 91.03 | 83.54 | 87.03 | 66.86 | 71.74 | 69.66 | 62.51 | 71.44 | 69.61 | 56.25 | 60.76 | 48.09 | - |
| Pink1 | 0.48 | 0.74 | 0.46 | 1.82 | 1.4 | 1.78 | 3.42 | 1.56 | 1.27 | 2.68 | 3.49 | 3.19 | - |
| Med9 | 12.26 | 11.61 | 11.54 | 10.24 | 11.72 | 9.67 | 9.12 | 8.85 | 9.54 | 7.23 | 6.54 | 8.57 | - |
| Kansl1 | 5.19 | 5.27 | 5.07 | 7.02 | 5.98 | 6.83 | 6.66 | 5.63 | 6.45 | 7.29 | 6.48 | 6.61 | - |
| Tmc8 | 4.43 | 4.41 | 4.03 | 5.11 | 5.54 | 4.69 | 7.89 | 5.26 | 6.17 | 6.43 | 6.63 | 6.69 | - |
| Apeh | 35.19 | 33.44 | 41.2 | 31.32 | 29.37 | 27.62 | 30.77 | 25.93 | 30.05 | 26.18 | 26.45 | 27.63 | - |

| | | | | | | | | | | | | | |
|---|---|---|---|---|---|---|---|---|---|---|---|---|---|
| Zfp839 | 0.83 | 0.97 | 1.1 | 2.39 | 1.44 | 1.4 | 2.08 | 1.01 | 1.16 | 1.64 | 2.1 | 2.08 | - |
| Tfpi | 5.58 | 5.45 | 6.34 | 4.66 | 6.36 | 4.78 | 3.58 | 5.54 | 5 | 2.78 | 4.3 | 3.1 | - |
| Irgc1 | 0.06 | 0 | 0.03 | 3.94 | 7.3 | 6.2 | 0 | 0 | 0 | 1 | 0.77 | 0.38 | - |
| Ergic3 | 92.21 | 81.61 | 84.34 | 103.78 | 93.85 | 105.82 | 94.42 | 85.95 | 93.9 | 106.37 | 107.96 | 105.81 | - |
| Rad51 | 47.47 | 44.8 | 44 | 20.54 | 31.56 | 25.71 | 13.14 | 19.14 | 16.69 | 10.73 | 12.46 | 10 | - |
| Abca3 | 4.48 | 4.61 | 5.14 | 7.19 | 6.61 | 5.99 | 6.18 | 5.25 | 6.63 | 5.74 | 6.71 | 6.66 | - |
| Rangrf | 36.53 | 39.94 | 41.95 | 25.86 | 29.4 | 26.99 | 32.5 | 41.72 | 38.11 | 25.17 | 28.25 | 28.14 | - |
| Porcn | 0.06 | 0 | 0 | 0.54 | 0.17 | 0.17 | 1.35 | 1.04 | 0.49 | 1.6 | 0.86 | 0.71 | - |
| Hey1 | 0.03 | 0 | 0 | 0.03 | 0.06 | 0 | 0.16 | 0.11 | 0.1 | 0.49 | 0.11 | 0.58 | - |
| Tbl2 | 9.93 | 8.25 | 8.93 | 7.07 | 7.73 | 8.31 | 7.79 | 7.71 | 7.61 | 6.21 | 7.04 | 5.74 | - |
| Pak6 | 9.42 | 9.29 | 8.91 | 6.43 | 6.78 | 6.75 | 7.06 | 8.23 | 8.56 | 5.85 | 5.14 | 5.41 | - |
| Vps13c | 0.6 | 0.49 | 0.41 | 0.9 | 0.99 | 0.94 | 0.91 | 0.6 | 0.81 | 0.95 | 1.19 | 0.76 | - |
| A630007B06Rik | 1 | 0.82 | 1.35 | 1.31 | 1.41 | 1.27 | 1.57 | 1.86 | 1.83 | 2.72 | 1.57 | 2.13 | - |
| Eng | 12.86 | 10.67 | 11.14 | 10.8 | 9.47 | 9.99 | 12.85 | 13.13 | 11.65 | 9.89 | 7.61 | 7.7 | - |
| Tbcel | 0.49 | 0.71 | 0.53 | 1.18 | 0.67 | 0.9 | 1.09 | 1.31 | 1.22 | 1.31 | 1.54 | 1.37 | - |
| Abhd11 | 29.13 | 24.97 | 29.11 | 21.32 | 21.78 | 23.05 | 20.48 | 22.32 | 20.63 | 17.93 | 16.36 | 19.16 | - |
| Dhodh | 21.33 | 19.93 | 20.01 | 8.78 | 9.96 | 11.45 | 5.95 | 10.44 | 10.21 | 4.37 | 5.08 | 5.5 | - |
| Srbd1 | 9.27 | 9.03 | 9.65 | 7.92 | 9.11 | 7.8 | 7.13 | 7.05 | 7.87 | 7.67 | 6.26 | 6.19 | - |
| Piwil4 | 1.17 | 1.94 | 1.11 | 1.05 | 1.07 | 0.56 | 0.59 | 0.4 | 0.43 | 0.26 | 0.23 | 0.42 | - |
| Ehbp1l1 | 17.64 | 16.79 | 17.87 | 21.06 | 24.54 | 23.37 | 23.25 | 22.11 | 22.05 | 23.88 | 23.68 | 23.85 | - |
| Dcun1d5 | 10.08 | 10.61 | 9.54 | 7.79 | 6.76 | 5.84 | 4.79 | 8.95 | 5 | 5.19 | 4.41 | 7.1 | - |
| Atp8a2 | 0.02 | 0 | 0.01 | 0.03 | 0 | 0.03 | 1.12 | 0.18 | 0.25 | 0.78 | 0.5 | 1.38 | - |
| Smpdl3a | 15.68 | 14.59 | 15.43 | 26.51 | 21.11 | 27.63 | 43.03 | 34.07 | 31.33 | 42.26 | 36.41 | 48.49 | - |
| Psen1 | 22.19 | 23.52 | 22 | 26.48 | 25.25 | 25.59 | 28.4 | 28.29 | 26.91 | 27.87 | 30.4 | 30.05 | - |
| 9430083A17Rik | 0.02 | 0.04 | 0.05 | 0.21 | 0.06 | 0.32 | 0.02 | 0.13 | 0.07 | 0.29 | 0.3 | 0.24 | - |
| 3230401D17Rik | 21.38 | 22.98 | 22.15 | 26.14 | 26.52 | 23.97 | 27.96 | 26.28 | 24.87 | 35.74 | 31.88 | 30.59 | - |
| Tmub1 | 7.33 | 6.66 | 7 | 10.39 | 11.26 | 11.98 | 11.06 | 10.23 | 10.73 | 16.21 | 12.61 | 11.83 | - |
| Tmem110 | 1.3 | 1.73 | 1.52 | 1.98 | 1.25 | 2 | 3.41 | 2.62 | 2.93 | 2.75 | 2.51 | 3.12 | - |
| Gna11 | 7.57 | 6.89 | 5.62 | 8.04 | 7.64 | 6.57 | 7.58 | 6.36 | 8.26 | 9.21 | 8.35 | 9.22 | - |
| Nedd1 | 19.37 | 17.47 | 19.76 | 11.4 | 14.65 | 11.65 | 9.79 | 11.34 | 12.77 | 9.05 | 8.95 | 10.62 | - |
| Smap1 | 3.64 | 4.02 | 3.67 | 3.82 | 4.57 | 5.7 | 6.76 | 6.81 | 5.38 | 6.09 | 6.74 | 6.54 | - |
| Anxa2 | 738.51 | 721.65 | 700.83 | 883.33 | 744.77 | 797.93 | 1230.87 | 881.65 | 1015.19 | 863.61 | 815.18 | 958.1 | - |
| Vac14 | 4.63 | 5.09 | 4.99 | 2.46 | 3.13 | 2.61 | 2.29 | 3.1 | 3.49 | 1.93 | 2.03 | 2.3 | - |
| Rars2 | 33.33 | 32.23 | 32.89 | 19.35 | 21.63 | 23.3 | 17.36 | 18.94 | 21.15 | 15.56 | 14.26 | 13.25 | - |
| Gm4759 | 1.35 | 1.72 | 1.26 | 3.03 | 1.52 | 2.04 | 2.4 | 1.88 | 1.98 | 3.53 | 2.29 | 4 | - |
| Vamp1 | 1.72 | 2.12 | 2.07 | 3.97 | 3.29 | 4.06 | 4.08 | 3.56 | 3.71 | 4.84 | 4.9 | 6.22 | - |
| Eral1 | 22.57 | 23.17 | 21.85 | 17.7 | 15.95 | 15.8 | 14.98 | 16.91 | 16.38 | 15.91 | 13.53 | 11.59 | - |
| Ano6 | 6.82 | 5.98 | 7.43 | 7.78 | 7.86 | 7.3 | 8.75 | 7.95 | 8.03 | 9.18 | 10.47 | 10.39 | - |
| Nf2 | 16.23 | 14.75 | 17.21 | 13.6 | 13.15 | 13.02 | 13.76 | 14.5 | 13.4 | 12.47 | 11.86 | 13.61 | - |
| Fam102b | 0.56 | 0.88 | 0.81 | 1.13 | 0.94 | 0.89 | 1.71 | 0.98 | 1.63 | 2.05 | 1.68 | 1.42 | - |
| Os9 | 21.16 | 20.87 | 19.94 | 29.81 | 27.87 | 26.63 | 29.29 | 23.02 | 21.63 | 39.79 | 34.25 | 43.16 | - |
| Fndc9 | 1.14 | 1.11 | 1.05 | 1.59 | 1.41 | 1.32 | 2.08 | 1.64 | 2.2 | 2.38 | 2.23 | 1.88 | - |
| Mycbp2 | 3.72 | 4.06 | 3.84 | 5.15 | 5.05 | 5.18 | 5.93 | 4.89 | 5.12 | 7.04 | 6.47 | 6.44 | - |
| Fam179b | 3.23 | 2.92 | 3.35 | 4.75 | 4.17 | 4.71 | 4.95 | 4.45 | 5.37 | 5.08 | 6.43 | 5.04 | - |
| Ndufa2 | 233.75 | 219.74 | 221.69 | 182.13 | 185.96 | 195.05 | 193.65 | 206.29 | 209.36 | 167.94 | 169.58 | 166.83 | - |
| Adipor1 | 39.38 | 44.36 | 48.16 | 67.76 | 62.17 | 63.76 | 80.63 | 76.2 | 78.23 | 98.07 | 90.59 | 74.39 | - |
| 5830418K08Rik | 3.79 | 3.21 | 3.82 | 2.73 | 3.02 | 3.2 | 2.06 | 2.45 | 2.39 | 1.89 | 2.4 | 1.42 | - |
| Gzmk | 0.07 | 0.88 | 0.07 | 0 | 0 | 0 | 4.72 | 0.67 | 1.56 | 4.84 | 2.26 | 7.86 | - |
| Acsf3 | 6.03 | 4.53 | 6.1 | 2.86 | 4.27 | 2.98 | 3.18 | 4.18 | 3.31 | 1.97 | 1.59 | 2.48 | - |
| Naca | 714.74 | 747.96 | 735.46 | 863.4 | 847.86 | 904.85 | 832.88 | 781.16 | 822.54 | 932.96 | 989.44 | 911.41 | - |
| Mrps14 | 65.81 | 64.21 | 63.87 | 45.69 | 43.98 | 43.85 | 39.61 | 46.66 | 49.54 | 33.63 | 34.23 | 32.98 | - |
| Dcaf8 | 23.58 | 23.99 | 25.53 | 30.5 | 30.54 | 30.02 | 31.75 | 27.57 | 26.91 | 30.37 | 33.49 | 36.38 | - |
| Plk1s1 | 5.26 | 4.55 | 3.42 | 9.4 | 7.84 | 8.57 | 9.14 | 5.76 | 6.46 | 8.04 | 10.92 | 9.57 | - |
| Ndc80 | 19.31 | 18.61 | 16.9 | 16.84 | 20.85 | 17.28 | 13.12 | 11.43 | 10.58 | 12.01 | 12.98 | 12.35 | - |
| Gimap5 | 45.53 | 38.74 | 41.93 | 40.9 | 37.03 | 37.97 | 33.29 | 32.38 | 34.08 | 30.99 | 31.08 | 35.66 | - |
| Pccb | 3.54 | 4.04 | 3.99 | 2.21 | 3.02 | 2.23 | 3.61 | 2.31 | 3.09 | 1.57 | 2.31 | 1.76 | - |
| Zfp354c | 0.31 | 0.16 | 0.38 | 0.79 | 0.45 | 0.34 | 0.82 | 0.62 | 0.94 | 0.54 | 0.77 | 0.71 | - |
| Deaf1 | 4.8 | 5.68 | 7.45 | 9.01 | 7.5 | 8.99 | 8.24 | 9.19 | 6.02 | 8.9 | 9.65 | 7.78 | - |
| Kif4 | 8.18 | 6.58 | 5.82 | 5.03 | 5.8 | 6.88 | 3.6 | 3.04 | 2.88 | 3.27 | 3.47 | 3.47 | - |
| Actbl2 | 0.15 | 0.3 | 0.14 | 0 | 0 | 0.05 | 0 | 0.02 | 0.12 | 0 | 0 | 0 | - |
| Nck1 | 22.53 | 18.16 | 19.26 | 20.41 | 18.52 | 21.8 | 22.64 | 21.45 | 25.12 | 26.96 | 25.74 | 24.05 | - |
| Arpc3 | 133.86 | 123.89 | 104.43 | 115.17 | 128.15 | 100.59 | 120.52 | 96.03 | 101.16 | 81.37 | 107.13 | 91.57 | - |
| Lrch3 | 4.11 | 4.53 | 4.98 | 7.36 | 6.57 | 6.02 | 6.85 | 6.5 | 6.4 | 6.01 | 8.34 | 7.1 | - |
| Rsrc1 | 13.11 | 13.59 | 13.25 | 10.25 | 10.91 | 12.33 | 11.63 | 12.77 | 11.95 | 9.73 | 9.41 | 9.49 | - |
| Kdm5a | 4.84 | 4.74 | 5.01 | 7.4 | 5.98 | 6.27 | 7.02 | 5.65 | 5.76 | 8.02 | 8.87 | 8.32 | - |
| Calhm2 | 7.66 | 8.35 | 6.76 | 10.7 | 6.46 | 9.03 | 10.99 | 9.34 | 8.82 | 9.24 | 11.07 | 13.22 | - |
| Nsmf | 3.56 | 3.78 | 3.91 | 4.98 | 5.33 | 4.48 | 5.12 | 4.47 | 3.94 | 5.28 | 6.13 | 4.8 | - |
| Med12l | 0.27 | 0.45 | 0.24 | 0.62 | 0.7 | 0.62 | 1.07 | 0.52 | 0.7 | 0.91 | 1.02 | 0.82 | - |
| Sertad2 | 2.36 | 2.72 | 2.21 | 3.34 | 3.6 | 2.77 | 3.6 | 3.21 | 3.03 | 4.46 | 4.41 | 4.21 | - |
| Med27 | 33.44 | 38.73 | 35.99 | 22.16 | 26.33 | 25.57 | 22.02 | 26.79 | 26.83 | 12.68 | 15.11 | 15.95 | - |
| Fbrs | 5.36 | 6.68 | 6.89 | 8.91 | 8.36 | 8.44 | 8.22 | 7.91 | 6.87 | 10.29 | 9.89 | 9.93 | - |
| Cth | 61.77 | 56.57 | 63.17 | 24.43 | 38.06 | 31.87 | 29.45 | 66.52 | 52.35 | 14.99 | 16.82 | 16.04 | - |
| Tcea1 | 84.49 | 93.46 | 94.12 | 82.78 | 85.07 | 82.85 | 72.28 | 86.35 | 85.77 | 74.65 | 80.62 | 74.17 | - |
| Jak1 | 34.22 | 34.58 | 35.19 | 38.62 | 33.83 | 37.15 | 44.6 | 34.24 | 37.11 | 45.6 | 40.32 | 44.93 | - |
| Efcab7 | 5.29 | 4.19 | 4.43 | 3.37 | 3.33 | 2.07 | 1.99 | 2.58 | 3.01 | 2.42 | 2.47 | 1.52 | - |
| Bcap29 | 122.09 | 128.14 | 133 | 69.54 | 83.99 | 74.57 | 67.62 | 98.06 | 86.29 | 58.28 | 56.93 | 56.68 | - |
| Zbtb2 | 2.81 | 2.99 | 3.69 | 6.82 | 6.07 | 7.13 | 5.72 | 4.39 | 4.51 | 7.64 | 7.89 | 7.41 | - |
| A930024E05Rik | 0.75 | 0.46 | 0.63 | 1.07 | 0.96 | 1.18 | 0.61 | 1.02 | 0.99 | 2 | 1.89 | 2.25 | - |
| Havcr2 | 12.45 | 15.14 | 7.46 | 16.59 | 16.86 | 19.23 | 12.93 | 8.61 | 9.15 | 22.5 | 19.94 | 22.58 | - |
| Klhl18 | 4.3 | 4.08 | 3.17 | 3.38 | 3.56 | 3.48 | 3.48 | 3.22 | 2.97 | 2.54 | 2.78 | 2.56 | - |
| Scd1 | 26.47 | 26.12 | 27.55 | 24.09 | 25.02 | 24.33 | 24.68 | 21.21 | 22.77 | 21.76 | 24.66 | 20.29 | - |
| Oxnad1 | 13.48 | 11.54 | 11.56 | 11.16 | 12.26 | 11.6 | 8.22 | 9.03 | 8.9 | 6.99 | 7.05 | 6.76 | - |
| Snx14 | 9.72 | 11.54 | 10.19 | 15.52 | 13.48 | 15.39 | 22.88 | 16.63 | 17.02 | 22.64 | 19.53 | 22.38 | - |
| Polg | 3.2 | 4.63 | 5.73 | 7.39 | 8.51 | 7.85 | 5.42 | 5.41 | 5.8 | 7.75 | 6.29 | 7.14 | - |
| Klhdc5 | 2.48 | 2.16 | 2.6 | 3.25 | 2.75 | 2.76 | 2.82 | 2.64 | 2.47 | 3.82 | 3.5 | 2.73 | - |
| Top2b | 5.91 | 5.32 | 4.94 | 6.73 | 6.03 | 5.68 | 6.41 | 6.43 | 7.25 | 7.66 | 7.36 | 7.53 | - |
| Acot9 | 47.68 | 48.15 | 47.9 | 57.86 | 53.02 | 56.19 | 56.87 | 48.68 | 43.29 | 66.7 | 59 | 72.58 | - |
| Ptpla | 11.6 | 12.87 | 12.02 | 7.2 | 6.41 | 8.06 | 7.72 | 7.03 | 8.72 | 5.62 | 4.45 | 6.38 | - |
| Acy1 | 22.74 | 20.28 | 24.72 | 9.63 | 14.88 | 15.73 | 11.11 | 15.01 | 15.31 | 8.24 | 7.34 | 7.91 | - |
| Rcor3 | 2.22 | 2.99 | 2.41 | 4.96 | 4.82 | 5.04 | 2.52 | 3.65 | 4.01 | 6.42 | 3.98 | 5.42 | - |
| Rnf138 | 18.66 | 20.95 | 18.49 | 34.72 | 34.03 | 33.48 | 37.97 | 31.42 | 33.76 | 49.01 | 51.26 | 46.76 | - |
| Nanos1 | 0.52 | 0.49 | 0.41 | 0.4 | 0.75 | 0.97 | 0.69 | 0.42 | 0.59 | 1.02 | 0.82 | 1.14 | - |
| Lclat1 | 3.64 | 3.2 | 3.21 | 2.53 | 2.71 | 3.13 | 2.77 | 3.24 | 3.37 | 2.07 | 2.13 | 2.34 | - |
| Yars2 | 6.79 | 5.14 | 5.73 | 2.23 | 2.84 | 3.58 | 2 | 4.08 | 2.91 | 3.18 | 2.06 | 1.89 | - |
| Ercc5 | 6.76 | 5.13 | 6.97 | 11.27 | 11.44 | 13.42 | 13.14 | 11.28 | 11.23 | 14.18 | 19.07 | 11.5 | - |
| Dcps | 57.22 | 54.97 | 51.88 | 76.81 | 72.36 | 67.03 | 57.13 | 61.64 | 62.09 | 73.21 | 73.09 | 81.69 | - |
| Mphosph8 | 18.25 | 18.88 | 15.67 | 14.09 | 17.26 | 16.48 | 14.26 | 13.86 | 16.97 | 12.62 | 12.7 | 11.23 | - |
| Cul7 | 3.66 | 3.04 | 3.44 | 3.52 | 4.31 | 3.98 | 3.17 | 2.42 | 2.03 | 2.5 | 1.97 | 1.7 | - |
| Kbtbd8 | 1.2 | 1.56 | 1.61 | 0.38 | 0.77 | 0.62 | 0.45 | 0.66 | 0.63 | 0.51 | 0.26 | 0.27 | - |
| H2-Q2 | 1.59 | 1.67 | 1.56 | 2.8 | 2.23 | 2.97 | 7.18 | 4.87 | 6.17 | 5.95 | 6.75 | 6.89 | - |
| Mid1 | 18.79 | 19.98 | 19.21 | 17.58 | 16.92 | 18.55 | 16.26 | 16.87 | 17.67 | 15.22 | 15.94 | 15.67 | - |
| Etv4 | 1 | 0.77 | 1.23 | 0.24 | 0.51 | 0.27 | 0.66 | 0.78 | 0.6 | 0.24 | 0.24 | 0.27 | - |
| H2-Q9 | 53.68 | 61.09 | 49.77 | 162.53 | 118.56 | 117.44 | 172.27 | 120.79 | 137.67 | 242.85 | 235.59 | 247.17 | - |
| Ccdc85b | 2.17 | 3.31 | 3.6 | 4.07 | 3.44 | 3.48 | 3.87 | 4.85 | 4.44 | 5.26 | 4.31 | 4.42 | - |
| Cyld | 10.15 | 10.61 | 10.58 | 11.68 | 11.02 | 10.1 | 12.81 | 11.98 | 12.92 | 12.88 | 14.27 | 10.94 | - |
| Vars | 51.28 | 67.91 | 79.72 | 107.23 | 98.05 | 91.37 | 99.31 | 93.3 | 94.89 | 111.08 | 116.76 | 103.16 | - |
| Sec24c | 49.98 | 47.03 | 49.09 | 64.7 | 58.45 | 58.06 | 59.97 | 54.48 | 55.51 | 66.08 | 73.24 | 64.2 | - |
| Ppic | 7.46 | 6.02 | 5.89 | 2.02 | 1.6 | 3.42 | 2.87 | 1.78 | 2.09 | 1.6 | 0.48 | 1.21 | - |
| Syf2 | 43.32 | 48.79 | 52.41 | 56.27 | 59.06 | 63.12 | 65.71 | 70.93 | 70.27 | 70.93 | 74.27 | 74.57 | - |
| St3gal6 | 1.65 | 2.13 | 1.43 | 6.47 | 4.07 | 5.69 | 4.5 | 1.73 | 1.35 | 3.7 | 3.17 | 4.16 | - |
| Tmem161b | 6.78 | 5.89 | 6.3 | 4.87 | 4.3 | 4.55 | 4.26 | 4.77 | 4.23 | 4.54 | 4.59 | 2.88 | - |
| Dtx1 | 0.06 | 0 | 0 | 0.09 | 0.03 | 0 | 0.19 | 0.27 | 0.33 | 0.26 | 0.38 | 0.28 | - |
| Ankhd1 | 2.46 | 2.76 | 2.68 | 3.69 | 3.28 | 3.19 | 3.25 | 3.2 | 3.36 | 3.5 | 3.73 | 3.54 | - |
| Rwdd4a | 21.83 | 23.26 | 23.9 | 14.56 | 14.91 | 15.01 | 10.76 | 15.98 | 16.87 | 11.76 | 11.13 | 10.05 | - |
| Ass1 | 111.45 | 118.19 | 103.06 | 170.21 | 137.88 | 145.98 | 161.61 | 112.53 | 131.54 | 124.65 | 129.89 | 146.49 | - |
| Pstpip1 | 48.36 | 45.52 | 38.17 | 63.78 | 54.92 | 56.54 | 117.05 | 75.38 | 92.8 | 107.52 | 101.23 | 129.21 | - |
| Zkscan6 | 9.3 | 8.92 | 8.5 | 13.92 | 12.18 | 13.55 | 14.23 | 11.84 | 10.91 | 13.05 | 14.24 | 15.37 | - |
| Gm129 | 7.41 | 5.92 | 5.58 | 5.12 | 5.73 | 6.86 | 8.08 | 9.14 | 8.81 | 8.82 | 9.81 | 9.21 | - |
| Smarcc1 | 4 | 4.39 | 3.79 | 2.7 | 2.8 | 2.83 | 2.78 | 3.34 | 3.18 | 1.97 | 3.17 | 2.23 | - |
| Gm11435 | 0 | 0 | 0 | 0 | 0.1 | 0.03 | 0.09 | 0.03 | 0.03 | 0.64 | 0.46 | 0.28 | - |
| Lime1 | 45.61 | 45.45 | 44.63 | 54.86 | 44.87 | 51.01 | 94.52 | 66.95 | 81.34 | 85.4 | 69.08 | 98.35 | - |
| Akt2 | 35.1 | 35.36 | 31.2 | 39.78 | 35.77 | 37.15 | 40.45 | 33.87 | 35.2 | 45.96 | 39.81 | 48.17 | - |
| Smim15 | 36.91 | 42.92 | 40.29 | 67.58 | 61.09 | 60.47 | 49.16 | 36.51 | 48.41 | 76.56 | 72.1 | 78.64 | - |
| R74862 | 0.15 | 0.18 | 0.2 | 0.5 | 0.46 | 0.87 | 0.36 | 0.31 | 0.28 | 0.51 | 0.75 | 0.91 | - |
| Uba3 | 51.57 | 49.63 | 48.87 | 40.52 | 41.02 | 42.17 | 40.19 | 45.53 | 45.82 | 33.64 | 41.31 | 36.03 | - |
| Slc6a6 | 23.38 | 21.05 | 21.86 | 15.42 | 15.66 | 15.15 | 12.09 | 16.56 | 15.35 | 10.91 | 12.09 | 11.6 | - |
| E330033B04Rik | 0.7 | 0.88 | 0.97 | 1.99 | 1.37 | 1.93 | 1.48 | 0.93 | 1.18 | 2.09 | 1.96 | 2.44 | - |
| Atp10a | 7.6 | 8.42 | 8.43 | 9.43 | 8.38 | 8.94 | 3.55 | 4.99 | 4.51 | 6.73 | 5.31 | 5.11 | - |
| Prkd2 | 13.17 | 13.77 | 12.15 | 19.87 | 18.08 | 20.26 | 24.82 | 19.65 | 19.97 | 26.12 | 28.61 | 25.92 | - |
| Ntrk3 | 1.37 | 1.4 | 1.21 | 0.54 | 0.4 | 0.55 | 0.22 | 0.44 | 0.2 | 0 | 0.13 | 0.06 | - |
| Tmed3 | 93.87 | 85.75 | 81.32 | 93.55 | 95.3 | 103.56 | 103.17 | 107.4 | 96.85 | 103.76 | 116.89 | 101.78 | - |
| Slc2a3 | 29.92 | 30.57 | 24.2 | 36.38 | 30.3 | 28.49 | 133.72 | 57.75 | 92.48 | 47.96 | 50.91 | 66.16 | - |
| Gm2382 | 23.33 | 26.99 | 20.22 | 37.05 | 33.58 | 32.34 | 44.2 | 29.69 | 36.79 | 46.56 | 52.29 | 50.78 | - |
| Rapsn | 2.5 | 2.73 | 1.9 | 1.37 | 1.43 | 1.74 | 1.88 | 1.81 | 1.49 | 1.24 | 0.53 | 1 | - |
| Pcf11 | 3.89 | 3.85 | 3.5 | 5.54 | 6.01 | 6.13 | 5.54 | 6.27 | 5.66 | 7.73 | 8.9 | 7.44 | - |
| Nek2 | 28.48 | 23.2 | 26.32 | 18.63 | 23.8 | 18.15 | 12.1 | 15.74 | 11.56 | 12.74 | 13.33 | 8.8 | - |
| Atg4c | 1.64 | 2.2 | 1.95 | 4.03 | 3.75 | 3.08 | 2.75 | 2.81 | 2.71 | 4.26 | 2.4 | 3.31 | - |
| Cd2 | 272.14 | 268.19 | 275.42 | 325.96 | 310.04 | 307.24 | 211 | 207.12 | 222.96 | 216 | 193.56 | 228.69 | - |
| Atg2b | 2.03 | 2.12 | 2.3 | 2.59 | 3.19 | 3.44 | 3.27 | 3.1 | 2.93 | 3.49 | 3.76 | 3.85 | - |
| Dym | 28.83 | 25.79 | 24.74 | 51.59 | 43.05 | 49.86 | 52.5 | 44.03 | 42.88 | 62.62 | 73.23 | 68.82 | - |
| Nxpe3 | 1.2 | 0.98 | 1.26 | 1.67 | 1.73 | 1.99 | 2.1 | 1.63 | 1.85 | 1.97 | 2.42 | 2.53 | - |
| Abr | 4.04 | 4.05 | 3.57 | 8.53 | 7.48 | 8.78 | 8.56 | 5.76 | 6.47 | 10.98 | 10.77 | 13.31 | - |
| Clk4 | 20.94 | 20.88 | 18.33 | 25.42 | 27.37 | 29.12 | 29.86 | 28.69 | 29.52 | 36.8 | 42.87 | 36.99 | - |
| Ankrd23 | 7.54 | 7.68 | 6.39 | 12.42 | 11.34 | 10.17 | 10.23 | 8.36 | 9.39 | 11.8 | 11.42 | 12.69 | - |
| Utp11l | 109.79 | 107.27 | 109.39 | 87.54 | 97.4 | 105.19 | 87.8 | 102.92 | 96.88 | 76.47 | 91.36 | 80.02 | - |
| Shisa2 | 0.38 | 0.22 | 0.29 | 0.44 | 0.24 | 0.12 | 0.88 | 0.49 | 0.77 | 1 | 0.98 | 0.9 | - |
| Brd4 | 5 | 5.81 | 5.83 | 6.38 | 7.22 | 6.74 | 5.74 | 6.78 | 5.51 | 6.81 | 7.97 | 7.1 | - |
| Enam | 0.43 | 0.43 | 0.43 | 0.08 | 0.08 | 0.14 | 0 | 0.05 | 0.05 | 0.02 | 0.03 | 0.01 | - |
| Gpd2 | 18.19 | 17.28 | 18.48 | 12.89 | 16.41 | 15.93 | 12.37 | 18.5 | 16.45 | 13.07 | 14.15 | 12.31 | - |
| Rrs1 | 27.61 | 40.44 | 47.32 | 23.49 | 24.38 | 18.03 | 16.33 | 21.62 | 21.8 | 17.7 | 11.86 | 13.34 | - |
| Stoml1 | 16.2 | 15.67 | 15.26 | 24.72 | 19.57 | 22.07 | 28.91 | 24.35 | 22.37 | 24.19 | 23.37 | 29.54 | - |

| | | | | | | | | | | | | |
|---|---|---|---|---|---|---|---|---|---|---|---|---|
| Ripk3 | 30.82 | 33 | 29.27 | 33 | 27.19 | 27.91 | 46.12 | 36.07 | 35.53 | 37.85 | 41.13 | 40.4 | - |
| Zfp169 | 1.8 | 2.06 | 1.55 | 2.25 | 2.58 | 1.87 | 2.03 | 2.27 | 2.56 | 2.62 | 2.93 | 3.53 | - |
| Tnfrsf8 | 79.45 | 81.06 | 93.04 | 43.85 | 57.75 | 49.97 | 70.18 | 81.74 | 79.16 | 47.8 | 49.19 | 53.29 | - |
| Saal1 | 9.71 | 9.09 | 9.75 | 8.31 | 8.62 | 9.23 | 7.76 | 7.28 | 9.01 | 6.87 | 6.45 | 6.55 | - |
| Lrrcc1 | 5 | 4.31 | 3.93 | 7.39 | 6.2 | 6.07 | 5.3 | 5.47 | 6.95 | 8.22 | 6.25 | 7.56 | - |
| Nfkbib | 54.75 | 57.26 | 58.6 | 31.75 | 37.09 | 39.94 | 29.9 | 45.5 | 38.54 | 27.56 | 30.07 | 27.53 | - |
| Zfp420 | 0.99 | 1.1 | 0.94 | 1.55 | 1.97 | 1.56 | 1.22 | 1.97 | 1.4 | 1.58 | 2.07 | 1.69 | - |
| Higd2a | 76.73 | 82.77 | 82.56 | 78.15 | 80.04 | 88.12 | 107.81 | 98.78 | 107.55 | 99.2 | 106.07 | 114.8 | - |
| Phf12 | 2.95 | 2.84 | 2.7 | 4.01 | 3.35 | 3.24 | 3.48 | 3.3 | 3.15 | 3.86 | 4.84 | 3.78 | - |
| 5830416P10Rik | 0.15 | 0.26 | 0.58 | 0.98 | 0.97 | 1.08 | 1.39 | 0.77 | 1.37 | 2.02 | 2.28 | 2.02 | - |
| Mgea5 | 8.16 | 9.06 | 9.6 | 12.32 | 11.98 | 12.22 | 12.91 | 11.68 | 12.86 | 15.62 | 15.74 | 13.71 | - |
| Camkk2 | 0.78 | 1.08 | 0.92 | 0.34 | 0.54 | 0.49 | 0.42 | 0.65 | 0.85 | 0.51 | 0.43 | 0.3 | - |
| Gdi2 | 213.86 | 213.17 | 219.72 | 276.03 | 261.7 | 278.52 | 322.49 | 268.56 | 285.62 | 310.51 | 352.68 | 313.71 | - |
| Sdf2 | 44.39 | 40.1 | 40.93 | 72.92 | 63.89 | 74.75 | 79.35 | 68.4 | 60.54 | 91.95 | 94.1 | 87.62 | - |
| Adck2 | 4.52 | 3.05 | 3.25 | 5.77 | 4.21 | 6.18 | 6.17 | 5.36 | 6.11 | 7.19 | 5.94 | 6.3 | - |
| Pde2a | 0.14 | 0.33 | 0.3 | 2.75 | 0.74 | 0.4 | 2.95 | 0.36 | 1.24 | 8.24 | 3.04 | 7.21 | - |
| Depdc1a | 6.63 | 6.42 | 6.09 | 4.54 | 4.9 | 3.58 | 2.84 | 2.94 | 3.41 | 3.22 | 3.96 | 1.93 | - |
| Calm3 | 54.06 | 52.13 | 55.02 | 41.74 | 45.49 | 45.26 | 42.66 | 43.95 | 46.57 | 33.29 | 32.16 | 30.41 | - |
| Cd9 | 27.83 | 23.14 | 26.63 | 17.39 | 19.71 | 16.86 | 16.15 | 17.64 | 16.29 | 14.25 | 9.25 | 12.28 | - |
| Nlgn2 | 2.7 | 1.83 | 2.73 | 4.12 | 3.71 | 4.12 | 4.58 | 3.97 | 4.04 | 3.61 | 5.42 | 4.26 | - |
| Dixdc1 | 0.26 | 0.3 | 0.06 | 0.04 | 0.13 | 0.06 | 1.49 | 0.52 | 0.7 | 0.49 | 0.55 | 0.89 | - |
| Trim14 | 2.27 | 2.36 | 2.38 | 5.03 | 5.23 | 4.5 | 4.53 | 4.27 | 4.83 | 7.51 | 4.52 | 5.03 | - |
| Agk | 19.28 | 16.4 | 16.45 | 12.39 | 12.63 | 12.78 | 12.35 | 13.07 | 11.28 | 10.46 | 8.89 | 9.25 | - |
| Opa1 | 11.89 | 10.92 | 13.24 | 9.23 | 11.07 | 11.15 | 10.05 | 9.74 | 10.7 | 9.14 | 8.73 | 8.59 | - |
| Pigk | 20.23 | 18.37 | 19.8 | 13.5 | 13.56 | 14.5 | 12.27 | 14.5 | 14.92 | 10.81 | 12.28 | 9.85 | - |
| BC022687 | 0.83 | 1.24 | 1.41 | 1.91 | 2.21 | 1.47 | 1.42 | 2.6 | 2.13 | 2.12 | 2.54 | 2.44 | - |
| Wdfy4 | 0.23 | 0.34 | 0.07 | 0.09 | 0.01 | 0.05 | 0.09 | 0.15 | 0.07 | 0 | 0.03 | 0 | - |
| Sh3gl1 | 34.61 | 35.39 | 34.28 | 62.15 | 54.09 | 56.82 | 55.37 | 42.1 | 43.88 | 63.61 | 66.65 | 69.81 | - |
| Sdcbp2 | 17.87 | 13.34 | 16.42 | 13.8 | 19.14 | 15.86 | 27.22 | 25.78 | 30.14 | 27.47 | 24.96 | 23.51 | - |
| 2310035C23Rik | 1.54 | 2.08 | 2.34 | 2.97 | 2.01 | 2.6 | 2.76 | 2.43 | 2.88 | 3.69 | 3.32 | 2.44 | - |
| Btf3 | 739.79 | 783.63 | 761.9 | 919.83 | 905.34 | 963 | 992.73 | 906.66 | 976.64 | 997.5 | 1085.91 | 983.88 | - |
| Tmem30a | 27.04 | 39.52 | 41.8 | 41.69 | 43.61 | 35.82 | 57.36 | 62.37 | 68.05 | 64.2 | 57.62 | 47.5 | - |
| Fkbp1a | 337.47 | 340.31 | 333.03 | 299.28 | 288.8 | 294.87 | 243.24 | 289.9 | 306.49 | 196.33 | 192.6 | 191.27 | - |
| Pfkfb1 | 2.95 | 3.48 | 2.98 | 2.2 | 2.91 | 1.78 | 1.84 | 2.35 | 2.03 | 1.46 | 1.15 | 0.94 | - |
| Gm14718 | 2.72 | 4.7 | 5.29 | 1.09 | 0.33 | 1.27 | 0.46 | 1.01 | 1.21 | 0.08 | 0.49 | 0.23 | - |
| Eif3k | 255.9 | 266.03 | 247.95 | 328.09 | 314.33 | 318.19 | 335.33 | 301.11 | 305.39 | 349.01 | 365.14 | 352.43 | - |
| Phpt1 | 34.35 | 39.32 | 40.23 | 31 | 29.47 | 28.66 | 30.36 | 26.39 | 26.95 | 22.47 | 21.46 | 27.13 | - |
| Slc12a9 | 1.83 | 1.97 | 1.4 | 3.15 | 2.17 | 1.98 | 1.82 | 2.55 | 2.2 | 2.67 | 4 | 2.78 | - |
| Slc41a1 | 3.49 | 5.01 | 4.33 | 5.75 | 3.86 | 5.41 | 5.02 | 4.85 | 4.73 | 5.14 | 6.3 | 7.12 | - |
| Sepp1 | 26.42 | 22.69 | 21.46 | 36.94 | 28.24 | 33.63 | 68.68 | 38.69 | 42.1 | 67.12 | 45.73 | 72.53 | - |
| Esm1 | 1 | 2.32 | 0.46 | 2.7 | 2.17 | 2.02 | 18.7 | 7.46 | 9.93 | 9.08 | 4.86 | 8.8 | - |
| Slc2a6 | 15.29 | 12.65 | 14.04 | 4.12 | 3.81 | 4.22 | 3.67 | 7.61 | 6.55 | 1.48 | 1.13 | 1.24 | - |
| Hrsp12 | 37.17 | 32.14 | 35.41 | 31.86 | 29.08 | 31.37 | 28.15 | 30.25 | 29.04 | 23.6 | 21.95 | 22.21 | - |
| Rab27a | 22.83 | 25.09 | 21 | 28.51 | 27.8 | 22.78 | 38.32 | 31.32 | 33.16 | 28.89 | 29.2 | 29.79 | - |
| Atp2a3 | 17.24 | 18.38 | 17.34 | 19.16 | 15.24 | 16.11 | 26.9 | 23 | 22.93 | 24.86 | 20.09 | 25.52 | - |
| Uba7 | 9.58 | 9.24 | 9.39 | 15.11 | 12.02 | 14.86 | 15.75 | 11.12 | 11.72 | 16.17 | 12.06 | 17.05 | - |
| Epb4.1 | 8.85 | 8.03 | 8.7 | 14.8 | 12.16 | 11.01 | 10.08 | 9.52 | 10.15 | 14.18 | 11.17 | 12.99 | - |
| Gltscr1 | 1.27 | 1.31 | 1.23 | 2.77 | 2.54 | 2.64 | 3.04 | 2.1 | 2.11 | 3.68 | 3.46 | 4.14 | - |
| Mgst2 | 36.83 | 40.7 | 33.32 | 64.38 | 49.07 | 55.44 | 62.63 | 43.48 | 50.74 | 57.22 | 52 | 62.04 | - |
| Zswim8 | 2.9 | 3.16 | 3.32 | 4.92 | 4.23 | 3.27 | 3.37 | 3.63 | 3.61 | 4.74 | 4.19 | 4.73 | - |
| Cdkn1a | 164.29 | 156.5 | 157.31 | 97.6 | 127.81 | 115.77 | 153.18 | 177.78 | 189.79 | 94.71 | 109.35 | 104.2 | - |
| Prkch | 21.28 | 23.7 | 24.61 | 34.47 | 31.68 | 26.57 | 34.37 | 34.27 | 33.43 | 50.12 | 38.37 | 39.11 | - |
| A630089N07Rik | 34.77 | 38.24 | 40.93 | 59.46 | 53.94 | 52.7 | 51.79 | 47.06 | 44.28 | 65.24 | 61.12 | 56.15 | - |
| Sco1 | 9.3 | 8.95 | 9.02 | 5.13 | 6.39 | 6.56 | 5.93 | 7.57 | 7.42 | 4.03 | 4.28 | 5.28 | - |
| Stk3 | 5.12 | 6.18 | 5.5 | 2.28 | 4.18 | 3.33 | 3.55 | 5.71 | 5.42 | 2.06 | 3.41 | 2.26 | - |
| Plekhf1 | 1.9 | 1.6 | 1.25 | 3.77 | 2.52 | 4.01 | 7.41 | 5.01 | 5.78 | 8.26 | 8.62 | 9.94 | - |
| Srp72 | 60.7 | 54.9 | 56.46 | 60.31 | 54.54 | 63.56 | 84.9 | 68.69 | 73.91 | 77.83 | 88.29 | 71.1 | - |
| Wrn | 2.18 | 2.16 | 2.32 | 3.15 | 2.66 | 3.15 | 3.07 | 2.57 | 2.76 | 3.65 | 3.07 | 3.53 | - |
| Mtfr1l | 30.81 | 29.41 | 31.67 | 39.95 | 34.58 | 42.56 | 45 | 36.62 | 42.5 | 44.16 | 48.38 | 45.04 | - |
| Nid1 | 1.53 | 0.9 | 0.95 | 0.6 | 0.77 | 0.95 | 0.29 | 0.64 | 0.61 | 0.32 | 0.33 | 0.51 | - |
| Plod2 | 3.98 | 4 | 4.47 | 5.4 | 5.52 | 4.47 | 7.8 | 4.5 | 6.36 | 6.18 | 6.95 | 4.97 | - |
| Nr4a3 | 2.79 | 4.05 | 4.22 | 0.94 | 2.02 | 1.56 | 0.49 | 1.87 | 1.22 | 0.82 | 0.62 | 0.52 | - |
| Casp9 | 2.49 | 2.39 | 2.89 | 5.34 | 5.54 | 4.88 | 5 | 3.71 | 3.93 | 4.92 | 7.06 | 5.67 | - |
| Ngrn | 23.33 | 26.44 | 27.08 | 32.43 | 31.13 | 30.67 | 33.59 | 32.79 | 34.31 | 36.76 | 35.5 | 32.81 | - |
| Rpp38 | 16.95 | 18.41 | 18.91 | 9.17 | 9.97 | 9.44 | 10.24 | 12.47 | 12.49 | 5.63 | 6.05 | 9.26 | - |
| Sipa1l1 | 8.56 | 8.52 | 8.53 | 16.94 | 16.79 | 16.11 | 10.3 | 9.6 | 10.07 | 17.46 | 17.8 | 19.24 | - |
| Mccc1 | 10.77 | 10.33 | 10.76 | 7.64 | 7.1 | 6.06 | 7.64 | 7.2 | 9.26 | 5.34 | 7.09 | 6.75 | - |
| Hist3h2a | 1.49 | 0.78 | 1.2 | 2.48 | 1.29 | 2.02 | 2.23 | 1.6 | 1.92 | 3.01 | 1.95 | 3.16 | - |
| Tnrc6c | 1.32 | 0.95 | 1.12 | 2.13 | 1.91 | 2.29 | 1.63 | 1.19 | 1.55 | 2.24 | 2.36 | 2.56 | - |
| Lamtor1 | 71.34 | 67.56 | 67.52 | 58.09 | 59.92 | 55.87 | 67.71 | 59.72 | 60.57 | 55.9 | 52.48 | 57.22 | - |
| Pde1b | 0.89 | 1.1 | 1.17 | 1.5 | 1.61 | 1.55 | 2.49 | 1.77 | 1.67 | 2.52 | 2.61 | 2.95 | - |
| Shmt1 | 75.87 | 76.85 | 72.9 | 20.94 | 25.97 | 27.15 | 13.74 | 24.52 | 22.6 | 11.69 | 9.58 | 7.93 | - |
| Chd8 | 2.78 | 2.45 | 3.12 | 3.16 | 2.82 | 3.33 | 3.25 | 3.27 | 3.12 | 3.68 | 3.77 | 3.58 | - |
| Tgoln1 | 16.91 | 17.44 | 18.72 | 30.14 | 27.79 | 25.88 | 31.81 | 25.51 | 30.3 | 35.86 | 41.57 | 39.91 | - |
| Ube2b | 76.4 | 69.12 | 68.38 | 90.51 | 85.76 | 93.13 | 95.8 | 82.01 | 82.93 | 99.07 | 97.44 | 93.68 | - |
| Cand2 | 3.01 | 3.16 | 3.11 | 3.22 | 2.82 | 3.17 | 2.65 | 1.68 | 2.43 | 1.85 | 1.96 | 2.52 | - |
| Prpf19 | 3.27 | 2.9 | 3 | 1.98 | 2.66 | 2.44 | 1.61 | 2.65 | 1.81 | 1.91 | 2.06 | 1.39 | - |
| Lef1 | 44.16 | 46.63 | 45 | 73.48 | 72.69 | 68.23 | 72.83 | 61.44 | 60.08 | 86.12 | 77.99 | 89.33 | - |
| Lrp11 | 0.32 | 0.29 | 0.46 | 0.04 | 0.04 | 0.08 | 0.1 | 0.36 | 0.19 | 0 | 0 | 0 | - |
| Ftl1 | 1245.14 | 1206.94 | 1252.36 | 963.87 | 1077.97 | 1142.14 | 1216.37 | 1362.22 | 1260.53 | 971.36 | 1123.14 | 986.95 | - |
| Maoa | 5.54 | 4.37 | 6 | 3.85 | 3.91 | 4.44 | 2.97 | 4.4 | 3.37 | 2.97 | 2.72 | 2.9 | - |
| Acer2 | 0.76 | 0.42 | 0.55 | 1.98 | 1.34 | 1.17 | 1.15 | 0.96 | 0.86 | 2.83 | 2.33 | 2.44 | - |
| Coprs | 2.7 | 3.56 | 4.04 | 3.47 | 4.83 | 4.45 | 5.88 | 7.1 | 4.82 | 6.85 | 8.38 | 5.17 | - |
| Tapt1 | 2.34 | 2.14 | 2.78 | 3.05 | 2.45 | 3.02 | 3.13 | 2.93 | 3.24 | 3.95 | 4.01 | 3.42 | - |
| Il7r | 45.37 | 41.4 | 44.33 | 127.4 | 106.34 | 109.26 | 104.68 | 84.05 | 91.97 | 174.56 | 129.32 | 178.58 | - |
| Flcn | 10.17 | 9.21 | 9.18 | 16.48 | 13.88 | 14.07 | 16.44 | 13.28 | 14.33 | 19.5 | 21.24 | 19.29 | - |
| E2f8 | 6.14 | 6.29 | 5.58 | 2.57 | 2.99 | 2.09 | 0.96 | 1.78 | 1.6 | 0.95 | 0.79 | 0.78 | - |
| 2700038G22Rik | 9.79 | 12.6 | 12.67 | 6.06 | 9.56 | 8.72 | 6.4 | 8.06 | 8.35 | 6.58 | 5.99 | 7.19 | - |
| Kif24 | 1.98 | 1.74 | 1.46 | 1.05 | 2.2 | 1.96 | 1.28 | 1.08 | 1.2 | 0.95 | 1.17 | 1.18 | - |
| Wdr1 | 377.87 | 355.72 | 388.42 | 305.41 | 318.01 | 298.63 | 259.67 | 304.46 | 309.73 | 217.81 | 216.48 | 231.97 | - |
| Sla2 | 24.25 | 27.01 | 20.5 | 40 | 27.39 | 29.9 | 38.53 | 28.4 | 33.12 | 41.42 | 37.02 | 43.43 | - |
| Commd1 | 78.44 | 78.46 | 68.67 | 49.84 | 52.2 | 51.45 | 49.39 | 57.3 | 54.18 | 34.77 | 43.4 | 39.53 | - |
| Hopx | 2.58 | 3.11 | 4.11 | 2.75 | 1.98 | 4.04 | 7.28 | 8.93 | 9.66 | 7.3 | 5.19 | 7.82 | - |
| Csgalnact2 | 1.62 | 1.32 | 1.2 | 1.78 | 1.74 | 1.79 | 1.13 | 2.36 | 2.42 | 2.25 | 2.51 | 2.13 | - |
| Rbl2 | 18.05 | 15.1 | 17.95 | 19.37 | 21.73 | 20.83 | 25.19 | 28.14 | 31.11 | 34.54 | 34.42 | 33.85 | - |
| Iqgap3 | 2.02 | 1.69 | 1.94 | 1.64 | 1.59 | 1.52 | 0.93 | 1.15 | 1.21 | 1.16 | 0.99 | 1.03 | - |
| Zfp229 | 0.31 | 0.36 | 0.62 | 0.1 | 0.11 | 0.2 | 0.2 | 0.18 | 0.17 | 0.04 | 0.2 | 0.13 | - |
| Ramp1 | 0.68 | 0.71 | 0.31 | 4.4 | 2.6 | 2.18 | 5.79 | 1.06 | 3.2 | 6.29 | 4.62 | 5.38 | - |
| Mgarp | 16.01 | 12.95 | 17.25 | 32.63 | 25.73 | 28.6 | 58.59 | 26.33 | 39.92 | 33.97 | 37.49 | 48.07 | - |
| Gm12216 | 1.44 | 0.56 | 1.27 | 2.19 | 1.57 | 0.94 | 3.54 | 1.43 | 1.12 | 3.18 | 3.58 | 4.5 | - |
| Tspan14 | 10.84 | 11.99 | 9.87 | 21.52 | 18 | 22.25 | 18.54 | 13.25 | 14.08 | 24.54 | 20.81 | 24.82 | - |
| Mrps10 | 107.56 | 101.27 | 103.73 | 52.87 | 50.43 | 56.4 | 49.54 | 66.52 | 65.39 | 34.16 | 34.38 | 31.65 | - |
| 1810055G02Rik | 17.47 | 17.01 | 18.13 | 10.95 | 12.44 | 11.39 | 9.59 | 13.05 | 9.53 | 9.85 | 6.39 | 7.78 | - |
| Zfp574 | 6.55 | 7.08 | 7.9 | 8.82 | 8.57 | 8.62 | 7.59 | 7.19 | 7.85 | 10.13 | 8.33 | 9.13 | - |
| Rab11fip4 | 0.73 | 0.78 | 0.33 | 0.46 | 0.67 | 0.49 | 1.24 | 1.7 | 0.94 | 0.98 | 1.35 | 2 | - |
| Nprl3 | 14.89 | 15.87 | 14.13 | 10.71 | 12.93 | 12.88 | 11.38 | 14.33 | 11.01 | 11.84 | 11.72 | 9.19 | - |
| St14 | 7.5 | 7.26 | 6.3 | 5.37 | 4.74 | 5.32 | 7.48 | 7.8 | 8.11 | 4.46 | 4.37 | 5.01 | - |
| Elk4 | 1.54 | 1.25 | 2.4 | 3.36 | 3.21 | 3.01 | 2.82 | 2.4 | 2.63 | 3.51 | 2.65 | 3.09 | - |
| Inpp5d | 14.18 | 14.62 | 12.95 | 22.44 | 17.56 | 18.84 | 14.7 | 11.85 | 13.15 | 21.89 | 18.31 | 23.12 | - |
| 4833420G17Rik | 15.02 | 17.08 | 13.34 | 23.25 | 22.97 | 22.04 | 25.08 | 19.62 | 22.12 | 30.81 | 30.21 | 36.15 | - |
| Trmt2b | 21.26 | 20.99 | 22.08 | 24.38 | 26.41 | 27.88 | 23.42 | 25.98 | 23.74 | 25.47 | 30.63 | 24.72 | - |
| Tmem131 | 2.22 | 1.67 | 2.33 | 2.96 | 2.9 | 3.41 | 3.49 | 2.9 | 3.35 | 3.93 | 4.24 | 3.77 | - |
| Zfp53 | 11.19 | 9.43 | 10.72 | 15.02 | 12.25 | 13.85 | 12.32 | 12.13 | 14.04 | 17.87 | 18.06 | 20.52 | - |
| B930003M22Rik | 0 | 0 | 0.22 | 0.83 | 0.2 | 0.37 | 0.7 | 0.48 | 0.39 | 1.08 | 0.91 | 0.33 | - |
| Pcdhgb2 | 0.09 | 0.13 | 0.16 | 0 | 0.08 | 0.06 | 0 | 0.08 | 0 | 0 | 0 | 0 | - |
| Depdc7 | 0.6 | 0.78 | 0.57 | 0.07 | 0.25 | 0.19 | 0.37 | 0.2 | 0.34 | 0.23 | 0.04 | 0 | - |
| Swi5 | 304.27 | 303.75 | 282.49 | 222.07 | 242.67 | 234.08 | 208.16 | 233.56 | 216.48 | 158.21 | 194.48 | 162.08 | - |
| Tnfsf11 | 5.75 | 6.77 | 9.7 | 1.78 | 1.87 | 1.88 | 1.65 | 5.31 | 5.09 | 0.46 | 0.36 | 1.08 | - |
| Sgol1 | 16.67 | 14.33 | 12.58 | 11.03 | 13.94 | 11.89 | 6.5 | 7.1 | 6.17 | 6.58 | 6.47 | 5.27 | - |
| Gbp10 | 4 | 3.06 | 3.16 | 6 | 4.34 | 6.16 | 4.48 | 2.98 | 3.18 | 6.95 | 5.68 | 8.41 | - |
| Wwp1 | 8.13 | 8.8 | 9.43 | 6.16 | 6.73 | 5.18 | 7.61 | 11.14 | 10.57 | 5.95 | 6.76 | 5.19 | - |
| Kif1a | 0.03 | 0.06 | 0 | 0.61 | 0.21 | 0.43 | 2.35 | 0.56 | 0.7 | 1.3 | 1.32 | 1.4 | - |
| Ankrd11 | 4.52 | 4.7 | 4.22 | 6.6 | 4.93 | 6.68 | 6.16 | 4.9 | 5.71 | 7.61 | 6.71 | 7.39 | - |
| Csrnp1 | 9.89 | 11.33 | 10.07 | 11.18 | 12.61 | 12.64 | 7.47 | 9.8 | 8.19 | 6.51 | 9.58 | 6.06 | - |
| Elovl4 | 0.79 | 1.14 | 0.92 | 0.52 | 0.27 | 0.04 | 0.28 | 0.6 | 0.3 | 0.28 | 0.22 | 0.15 | - |
| Prss30 | 1.2 | 1.58 | 1.33 | 1.25 | 1.55 | 1.82 | 7.2 | 6.4 | 7.01 | 6.53 | 4.24 | 7.3 | - |
| Ercc6l2 | 2.3 | 2.98 | 2.76 | 2.42 | 2.29 | 2 | 2.06 | 2.25 | 2.26 | 1.74 | 2.16 | 1.24 | - |
| Zfp410 | 17.43 | 16.77 | 17.08 | 14.28 | 13.26 | 15.32 | 12.11 | 13.94 | 13.87 | 10.71 | 11.34 | 12.12 | - |
| Egln2 | 15.58 | 18.09 | 20.12 | 25.35 | 22.81 | 23.91 | 24.46 | 20.54 | 21.27 | 27.08 | 26.7 | 30.91 | - |
| Slc29a2 | 4.52 | 4.16 | 4.5 | 2.1 | 3.28 | 2.63 | 1.84 | 3.04 | 3.28 | 2.61 | 2.39 | 1.26 | - |
| Ifitm2 | 528.48 | 589.22 | 458.35 | 309.36 | 280.2 | 291.33 | 706.56 | 544.19 | 654.87 | 355.95 | 346.77 | 422.92 | - |
| Tm2d1 | 37.11 | 33.46 | 36.52 | 33.71 | 34.06 | 32.53 | 25.29 | 29.07 | 33.52 | 20.54 | 23.3 | 22.98 | - |
| Lama3 | 0.2 | 0.18 | 0.2 | 0.02 | 0.01 | 0 | 0 | 0.02 | 0.01 | 0 | 0 | 0 | - |
| Chtf8 | 19.85 | 21.74 | 24.77 | 16.35 | 18.76 | 16.97 | 17.51 | 18.11 | 19.14 | 15.89 | 14.26 | 17.1 | - |
| Iars | 80.82 | 75.83 | 75.83 | 50.66 | 61.03 | 55.26 | 54.59 | 71.47 | 69.08 | 45.36 | 44.04 | 39.79 | - |
| Ptpre | 1.43 | 2.12 | 1.51 | 1.55 | 1.71 | 1.31 | 2.63 | 2.4 | 2.78 | 3.45 | 3.23 | 2.31 | - |
| Kdm3b | 2.82 | 2.65 | 2.39 | 3.87 | 4.08 | 3.71 | 4.21 | 3.85 | 3.49 | 5.67 | 6.83 | 4.85 | - |
| 2310067B10Rik | 4.15 | 4.69 | 4.24 | 7.59 | 8.47 | 6.38 | 7.53 | 7.09 | 7.69 | 10.03 | 9.21 | 9.23 | - |
| Col11a2 | 0.44 | 0.26 | 0.27 | 0.62 | 0.61 | 0.65 | 0.53 | 0.57 | 0.53 | 1.38 | 1.01 | 1.15 | - |
| Apol7e | 0.77 | 1.14 | 0.75 | 1.48 | 0.87 | 0.73 | 3.72 | 0.81 | 1.47 | 3.81 | 1.38 | 3.87 | - |
| D17Wsu104e | 131 | 126.89 | 130.82 | 96.28 | 109.67 | 106.24 | 96.3 | 119.95 | 121.58 | 70.8 | 77.7 | 70.54 | - |
| Tgif1 | 15.02 | 15.07 | 16.84 | 15.66 | 18.76 | 15.78 | 18.58 | 17.15 | 19.03 | 22.01 | 21.94 | 20.3 | - |
| Man2c1 | 17.47 | 15.77 | 17.36 | 11.89 | 12.64 | 12.3 | 11.08 | 14.17 | 13.39 | 9.86 | 8.3 | 9.88 | - |
| Tefm | 16.55 | 15.51 | 12.27 | 10.12 | 8.83 | 10.36 | 8.89 | 10.04 | 10.92 | 7.07 | 7.09 | 8.55 | - |
| Etfdh | 33.79 | 28.16 | 29.49 | 32.03 | 35.19 | 32.92 | 35.62 | 39.94 | 32.53 | 36.43 | 41.12 | 35.35 | - |
| Snx32 | 2.46 | 1.59 | 2.16 | 2.95 | 3.67 | 3.05 | 3.76 | 4.31 | 2.93 | 4.02 | 3.29 | 4.57 | - |
| Gm13157 | 23.26 | 24.52 | 24.01 | 27.81 | 25.32 | 27.09 | 26.07 | 29.49 | 30.1 | 29.12 | 32.31 | 28.49 | - |
| Tgfbr3 | 2.15 | 2.01 | 2.69 | 2.65 | 2.89 | 3.24 | 5.06 | 4.52 | 4.41 | 4.83 | 4.68 | 4.27 | - |
| S100a11 | 198.45 | 172.23 | 170.14 | 201.53 | 179.09 | 192.46 | 282.01 | 222.7 | 223.96 | 238.92 | 209.84 | 263.85 | - |
| Anxa6 | 204.94 | 197.27 | 193.54 | 164.58 | 160.19 | 161.84 | 211.39 | 193.84 | 204.84 | 171.24 | 153.88 | 175.85 | - |
| Dlgap5 | 21.88 | 21.81 | 21.19 | 17.35 | 20.62 | 18.33 | 11.98 | 12.18 | 11.14 | 10.61 | 11.66 | 8.33 | - |

| | | | | | | | | | | | | | |
|---|---|---|---|---|---|---|---|---|---|---|---|---|---|
| Kiss1r | 0.04 | 0.11 | 0.19 | 0.32 | 0.12 | 0.3 | 0.25 | 0.2 | 0.25 | 0.63 | 0.35 | 0.42 | - |
| Pdcd2 | 15.65 | 22.97 | 26.21 | 30.36 | 28.09 | 31.2 | 32.15 | 34.03 | 37.57 | 40.69 | 40.67 | 38.89 | - |
| 1110001J03Rik | 24.73 | 33.65 | 31.51 | 17.2 | 22.51 | 20.53 | 17.72 | 26.24 | 20.07 | 15.96 | 19.57 | 15.55 | - |
| Ica1l | 0.14 | 0.1 | 0.15 | 0.74 | 0.56 | 0.58 | 0.39 | 0.16 | 0.18 | 1.03 | 1.33 | 1.33 | - |
| Btbd2 | 1.91 | 1.9 | 1.7 | 3.53 | 2.62 | 2.55 | 3.48 | 2.56 | 2.3 | 3.16 | 2.69 | 3.79 | - |
| Ppip5k2 | 5.69 | 5.54 | 6.83 | 5.26 | 4.73 | 3.92 | 4.01 | 4.58 | 5 | 3.6 | 4.07 | 4.7 | - |
| Rassf5 | 42.56 | 45.74 | 43.06 | 56.07 | 45.72 | 47.53 | 47.2 | 39.05 | 44.55 | 63.49 | 51.65 | 57.89 | - |
| Rspry1 | 9.61 | 8.86 | 9.76 | 10.95 | 11.1 | 10.12 | 11.84 | 10.97 | 10.71 | 12.01 | 13.11 | 10.93 | - |
| Ndst1 | 2.83 | 2.93 | 2.48 | 5.33 | 4.94 | 5.5 | 5.05 | 3.73 | 4.6 | 9.74 | 8.64 | 8.92 | - |
| Efemp2 | 3.67 | 3.22 | 3.93 | 7.19 | 5.78 | 7.58 | 6.02 | 5.17 | 5.69 | 7.21 | 6.02 | 9.26 | - |
| Bysl | 29.8 | 27.58 | 28.95 | 9.68 | 11.6 | 13.31 | 10.09 | 15.56 | 14.13 | 6.72 | 6.42 | 7.28 | - |
| Lypd6b | 0.47 | 0.2 | 0.45 | 1.16 | 0.55 | 1.28 | 2.79 | 0.82 | 1.5 | 1.93 | 0.65 | 1.72 | - |
| Fbxo25 | 9.12 | 8.84 | 8.08 | 9.51 | 11.47 | 10.39 | 12.53 | 10.51 | 10.5 | 11.77 | 13.78 | 11.65 | - |
| Wnt4 | 0.06 | 0.06 | 0.16 | 0.12 | 0.32 | 0.34 | 0.15 | 0.27 | 0.19 | 0.36 | 0.44 | 0.39 | - |
| Coro2a | 15.44 | 17.02 | 16.17 | 12.86 | 11.78 | 11 | 6.89 | 8.75 | 8.51 | 4.41 | 4.18 | 4.76 | - |
| Gcm2 | 0.61 | 0.81 | 0.6 | 0.88 | 0.11 | 0.02 | 4.61 | 1.59 | 3.5 | 1.49 | 0.86 | 2.12 | - |
| Kremen1 | 1.69 | 1.64 | 1.73 | 2.5 | 2.32 | 2.35 | 3.37 | 2.06 | 2.37 | 2.63 | 2.81 | 3.64 | - |
| Bcl2l14 | 0.9 | 0.47 | 0.54 | 0.37 | 0.14 | 0.37 | 0.54 | 0.58 | 0.89 | 0.1 | 0.08 | 0.16 | - |
| Lgals9 | 104.76 | 104.3 | 108.6 | 100.84 | 92.99 | 103.4 | 100.94 | 92.07 | 96.71 | 86.63 | 77.9 | 98.87 | - |
| Cd320 | 25.38 | 22.13 | 19.41 | 14.39 | 16.76 | 15.07 | 18.29 | 21.97 | 17.8 | 12.59 | 12.3 | 12.94 | - |
| Slc43a2 | 1.34 | 1.47 | 0.92 | 1.68 | 1.72 | 1.5 | 2.87 | 1.81 | 2.32 | 2.16 | 1.69 | 1.9 | - |
| Ndufs8 | 127.48 | 130.81 | 136.03 | 91.33 | 102.85 | 97.9 | 76.72 | 95.37 | 87.4 | 65.6 | 71.53 | 69.34 | - |
| Cacna1s | 0.23 | 0.23 | 0.11 | 0.91 | 0.66 | 0.53 | 1.18 | 0.62 | 0.59 | 1.98 | 2.68 | 2.74 | - |
| Smarcb1 | 68.03 | 71.93 | 68.85 | 50.61 | 48.73 | 52.77 | 41.57 | 48.88 | 45.73 | 35.08 | 31.09 | 37.47 | - |
| Ankrd50 | 0.64 | 0.45 | 0.43 | 0.72 | 0.43 | 0.74 | 0.89 | 0.66 | 0.87 | 1.63 | 0.67 | 1.13 | - |
| Zfp945 | 0.75 | 0.74 | 0.51 | 1.34 | 0.87 | 0.95 | 1.87 | 1.17 | 1.32 | 2.42 | 2.07 | 2.77 | - |
| Adam8 | 53.71 | 48.48 | 42.37 | 49.98 | 45.08 | 51.29 | 117.1 | 76.18 | 90.05 | 97.24 | 89.82 | 102.67 | - |
| Itm2b | 200.21 | 202.68 | 188.78 | 283.83 | 263.08 | 273.62 | 473.08 | 308.38 | 352.18 | 380.6 | 387.58 | 414.27 | - |
| Gad2 | 0.48 | 0.6 | 0.33 | 0.09 | 0.09 | 0.1 | 0.11 | 0.17 | 0.14 | 0.06 | 0.03 | 0.09 | - |
| Faf1 | 2.75 | 3.63 | 3.65 | 4.74 | 4.57 | 4.49 | 5.51 | 6.22 | 6.61 | 5.55 | 6.98 | 4.62 | - |
| Fermt3 | 186.4 | 174.33 | 181.63 | 201.85 | 193.06 | 196.84 | 207.24 | 190.86 | 191.29 | 197.96 | 201.87 | 215.97 | - |
| Tmem17 | 1.08 | 1.44 | 0.75 | 1.43 | 0.91 | 1.12 | 0.34 | 0.26 | 0.51 | 0.17 | 0.4 | 0.08 | - |
| Abi3 | 2.35 | 2.38 | 1.95 | 5.9 | 5.22 | 5.83 | 4.43 | 2.74 | 2.66 | 5.73 | 4.34 | 5.05 | - |
| Bcl9 | 0.96 | 1.12 | 1.05 | 1.9 | 1.34 | 1.52 | 1.7 | 1.53 | 1.57 | 1.83 | 1.87 | 1.72 | - |
| Gm9159 | 1.58 | 1.65 | 1.58 | 2.75 | 2.72 | 3.19 | 2.5 | 2.49 | 2.07 | 2.83 | 3.27 | 3.24 | - |
| Oxr1 | 7.26 | 7.89 | 7.48 | 12.96 | 12.12 | 13.3 | 15.43 | 10.49 | 13.53 | 20.27 | 16.34 | 16.18 | - |
| Msantd4 | 12.47 | 13.37 | 13.49 | 10.25 | 11.45 | 10.51 | 9.2 | 12.28 | 9.68 | 8.76 | 8.02 | 10.64 | - |
| Mtmr4 | 3.91 | 3.76 | 4.06 | 2.78 | 2.93 | 2.39 | 2.7 | 3.79 | 3.23 | 1.93 | 2.79 | 2.16 | - |
| Hdac7 | 11.75 | 11.83 | 12.52 | 24.55 | 21.92 | 24.44 | 42.26 | 22.14 | 26.52 | 34.38 | 39.22 | 38.58 | - |
| Canx | 222.64 | 220.84 | 222.77 | 155.51 | 167.01 | 169.56 | 142.42 | 173.9 | 171.73 | 125.43 | 140.3 | 116.96 | - |
| Coq5 | 30.98 | 29.4 | 28.88 | 22.68 | 24.61 | 27.09 | 18.98 | 25.3 | 25.28 | 16.05 | 17.03 | 18.12 | - |
| Gimap8 | 8.84 | 9.48 | 9.34 | 13.28 | 11.92 | 11.73 | 11.18 | 9.69 | 10.59 | 13.14 | 10.93 | 14.25 | - |
| Emb | 143.61 | 140.32 | 133.92 | 275.53 | 217.07 | 232.1 | 280.8 | 166.47 | 191.17 | 391.55 | 305.38 | 412.22 | - |
| Rnf6 | 15.75 | 15.06 | 15.9 | 17.12 | 14.59 | 17.71 | 18 | 16.93 | 16.44 | 19.89 | 19.39 | 19.06 | - |
| Bicd1 | 1.24 | 0.56 | 0.69 | 0.18 | 0.14 | 0.28 | 0.24 | 0.27 | 0.11 | 0.14 | 0.07 | 0.15 | - |
| Runx3 | 23.87 | 29.13 | 28.86 | 52.89 | 42.41 | 46.06 | 77.04 | 52.63 | 62.11 | 75.34 | 76.56 | 78.09 | - |
| Ube2l6 | 12.92 | 12.18 | 11.94 | 14.43 | 11.96 | 12.75 | 22.42 | 17.24 | 16.89 | 22.07 | 19.66 | 18.39 | - |
| Insrr | 0.36 | 0.6 | 0.47 | 0.61 | 0.53 | 0.62 | 1.28 | 1.08 | 1.35 | 1.2 | 1.22 | 1.54 | - |
| Zc3h3 | 3.71 | 2.95 | 3.13 | 4.18 | 4.44 | 3.31 | 4.73 | 3.29 | 3.95 | 4.8 | 4.87 | 4.99 | - |
| Zfp286 | 1.73 | 1.74 | 2.07 | 1.22 | 1.41 | 0.98 | 1.22 | 1 | 1.29 | 1.05 | 0.45 | 1 | - |
| Zfpl1 | 18.09 | 19.76 | 19.5 | 29.6 | 24.64 | 25.48 | 33.53 | 24.89 | 27.84 | 27.49 | 32.71 | 28.16 | - |
| Cd53 | 395.5 | 398.61 | 384.92 | 473.19 | 489.08 | 482.09 | 619.29 | 572.87 | 604.82 | 589.78 | 531.52 | 610.3 | - |
| Wipi2 | 9.97 | 11.47 | 15.41 | 17.95 | 15.52 | 15.59 | 16.5 | 18.85 | 17.57 | 19.04 | 19.56 | 16.63 | - |
| Chd1l | 17.3 | 17.3 | 16.14 | 10.23 | 12.15 | 10.48 | 9.29 | 10.51 | 11.13 | 6.56 | 8.68 | 8.51 | - |
| Shprh | 2.27 | 2.21 | 2.75 | 2.52 | 2.63 | 3.54 | 2.51 | 2.38 | 3.11 | 3.46 | 3.69 | 2.98 | - |
| Arl6ip6 | 20.44 | 20.56 | 23.78 | 13.99 | 15.62 | 15.43 | 10.15 | 12.03 | 13.97 | 11.1 | 10.24 | 9.96 | - |
| Usp31 | 1.54 | 1.34 | 1.35 | 0.97 | 0.95 | 1.07 | 0.85 | 1.16 | 1.25 | 0.85 | 0.63 | 1.1 | - |
| Eci2 | 54.8 | 51.01 | 54.57 | 69.33 | 70.57 | 64.03 | 82.15 | 66.55 | 75.08 | 70.76 | 77.61 | 75.09 | - |
| Hist2h3c2 | 21.36 | 23.35 | 18.36 | 9.55 | 15.33 | 11.73 | 6.22 | 6.52 | 6.09 | 4.44 | 4.72 | 4.91 | - |
| Rnf19b | 13.18 | 13.51 | 13.22 | 12.73 | 13.16 | 13.79 | 5.88 | 9.65 | 8.92 | 6.53 | 6.09 | 6.56 | - |
| Oaf | 0.38 | 0.56 | 0.75 | 0.13 | 0.21 | 0.13 | 0.05 | 0.22 | 0.31 | 0.19 | 0.13 | 0 | - |
| Gpr68 | 24.02 | 24.34 | 25.26 | 29.13 | 24.03 | 26.22 | 41.01 | 30.02 | 32.31 | 39.71 | 35.07 | 39.89 | - |
| Mll5 | 7.76 | 7.09 | 7.74 | 13.73 | 12.76 | 14.36 | 12.01 | 9.58 | 10.1 | 15.81 | 16.71 | 16.51 | - |
| Ranbp1 | 269.3 | 333.33 | 325.13 | 111.27 | 127.7 | 128.52 | 94.7 | 137.18 | 125.81 | 62.38 | 55.55 | 60.08 | - |
| Tnfsf9 | 22.51 | 21.71 | 22.77 | 12.13 | 16.09 | 13.79 | 25.56 | 20.82 | 23.29 | 13.21 | 13.13 | 10.07 | - |
| Il3 | 29.65 | 15.22 | 24.84 | 0.22 | 0.51 | 0.75 | 0.41 | 0 | 1.01 | 0.07 | 0 | 0 | - |
| Lipt2 | 4.69 | 5.44 | 6.03 | 3.45 | 4.77 | 3.63 | 1.73 | 3.35 | 4.67 | 2.65 | 2.51 | 2.63 | - |
| 4632434I11Rik | 9.71 | 8.4 | 9.82 | 6.38 | 9.37 | 6.63 | 5.6 | 6.43 | 5.4 | 5.09 | 6.47 | 5 | - |
| Uqcc | 21.32 | 20.52 | 21.7 | 17.06 | 14.58 | 15.61 | 12.54 | 15.76 | 16.48 | 11.62 | 13.37 | 10.28 | - |
| Tgfbr2 | 6.58 | 9.97 | 8.6 | 15.62 | 12 | 13.14 | 19.4 | 14.04 | 17.1 | 23.37 | 15.06 | 19.13 | - |
| Mlxip | 3.51 | 3.93 | 3.92 | 7.1 | 5.77 | 5.93 | 5.62 | 5.37 | 5.48 | 10.44 | 9 | 8.76 | - |
| Myh11 | 0.25 | 0.03 | 0.09 | 0.05 | 0 | 0.04 | 0.01 | 0 | 0 | 0 | 0.01 | 0 | - |
| Tubgcp2 | 21.7 | 20.75 | 19.14 | 18.25 | 21.43 | 16.41 | 18.31 | 14.83 | 17.91 | 14.64 | 14.68 | 15.19 | - |
| Sugp1 | 22.57 | 22.47 | 25.79 | 25.12 | 26.15 | 24.24 | 28.22 | 24.04 | 25.95 | 29.27 | 27.32 | 30.72 | - |
| Jun | 12.84 | 12.5 | 14 | 5.39 | 6.99 | 6.52 | 16.06 | 18.04 | 15.84 | 6.73 | 8.24 | 6.64 | - |
| Gbp5 | 2.23 | 1.66 | 2.85 | 3.28 | 3.24 | 3.93 | 3.37 | 3.29 | 2.49 | 3.96 | 3.47 | 3.34 | - |
| 0610010F05Rik | 5.48 | 5.98 | 6.84 | 2.75 | 4.63 | 3.2 | 2.29 | 3.84 | 2.67 | 2.47 | 1.73 | 2.12 | - |
| Usf2 | 2.93 | 4.16 | 3.4 | 5.44 | 5.63 | 5.43 | 5.97 | 4.78 | 5.52 | 7.52 | 7.39 | 9.46 | - |
| Zfp952 | 5.31 | 6.36 | 5.3 | 7.13 | 7.41 | 6.85 | 7.18 | 6.8 | 6.74 | 7.85 | 7.2 | 8.25 | - |
| Nckap1l | 42.15 | 41.56 | 38.82 | 49.4 | 47.36 | 46.01 | 50.73 | 49.52 | 48.76 | 52.92 | 53.63 | 56.64 | - |
| Foxo1 | 2.61 | 2.71 | 2.42 | 4.67 | 4.72 | 3.61 | 3.59 | 3.83 | 3.85 | 4.03 | 3.67 | 5.32 | - |
| Dolpp1 | 12.25 | 11.57 | 10.93 | 9.33 | 8.86 | 8.92 | 9.14 | 10.24 | 10.67 | 8.88 | 7.77 | 7.29 | - |
| Cyb561d1 | 1.89 | 1.95 | 2.23 | 2.9 | 2.29 | 2.34 | 3.2 | 3.13 | 2.97 | 3.38 | 2.43 | 3.75 | - |
| Dusp4 | 13.74 | 12.68 | 13.41 | 7.73 | 11.08 | 10.56 | 5.94 | 11.41 | 8.62 | 6.41 | 6.15 | 4.2 | - |
| Fbxo32 | 1.01 | 1.28 | 1.26 | 1.44 | 1.61 | 1.07 | 2.4 | 1.86 | 1.89 | 2.71 | 2.91 | 2.94 | - |
| Prrg1 | 1.15 | 1.27 | 1.92 | 1.19 | 1.11 | 1.18 | 0.92 | 0.86 | 1.2 | 0.74 | 0.87 | 0.76 | - |
| Arhgap19 | 4.85 | 4.69 | 5.26 | 3.65 | 4.25 | 4.57 | 3.14 | 2.47 | 2.75 | 2.24 | 2.64 | 1.35 | - |
| Tln2 | 0.14 | 0.07 | 0.14 | 0.03 | 0.02 | 0.12 | 0.11 | 0.06 | 0.09 | 0.03 | 0.02 | 0.04 | - |
| Scml4 | 0.64 | 0.87 | 0.64 | 2.85 | 2.45 | 2.58 | 3.05 | 1.22 | 1.46 | 4.06 | 3.69 | 4.32 | - |
| Bbs9 | 3.28 | 3.79 | 3.53 | 5.38 | 4.5 | 4.3 | 4.5 | 4.67 | 4.88 | 4.81 | 5.3 | 6.82 | - |
| Lrig2 | 1.93 | 1.72 | 1.48 | 2.16 | 3.27 | 2.95 | 2.66 | 3.24 | 2.2 | 3.95 | 3.53 | 3.36 | - |
| Ak2 | 169.52 | 171.1 | 170.52 | 108.7 | 105.03 | 115.02 | 82.82 | 102.11 | 101.86 | 64.49 | 73.62 | 65.57 | - |
| Dctn3 | 128.09 | 127.91 | 123.44 | 99.97 | 110.93 | 101.85 | 108.25 | 122.14 | 109.89 | 94.08 | 93.28 | 98.99 | - |
| Herc3 | 2.91 | 3.08 | 3.32 | 4.38 | 5.03 | 3.95 | 5.75 | 4.24 | 4.76 | 5.42 | 5.68 | 6.47 | - |
| Cept1 | 37.89 | 35.05 | 33.32 | 44.73 | 43.86 | 46.04 | 46.27 | 49.46 | 44.17 | 47.38 | 52.99 | 52.21 | - |
| Nkain1 | 1.79 | 1.44 | 1.76 | 0.27 | 1.03 | 1.02 | 0.63 | 1.19 | 1.11 | 0.34 | 0.53 | 0.54 | - |
| Hsd17b4 | 24.88 | 25.76 | 24.87 | 21.7 | 18.74 | 20.83 | 18.34 | 23.98 | 21.96 | 19.94 | 18.13 | 17.82 | - |
| Psmd11 | 106.94 | 103.45 | 106.98 | 78.79 | 82.54 | 86.09 | 72.44 | 94.23 | 86.48 | 70.87 | 76.69 | 74.47 | - |
| Ermp1 | 2.87 | 3.15 | 3.46 | 3.9 | 3.61 | 2.7 | 4.35 | 4.69 | 4.21 | 5.13 | 4.78 | 4.66 | - |
| Glo1 | 77.61 | 79.81 | 76.08 | 68.25 | 67.89 | 66.17 | 72.64 | 77.39 | 77.57 | 62.32 | 65.16 | 63.14 | - |
| Myeov2 | 146.38 | 142.2 | 140.83 | 170.25 | 148 | 160.61 | 166.93 | 164.48 | 161.34 | 167.73 | 166.28 | 180.45 | - |
| Rad54l | 16.82 | 15.7 | 15.1 | 6.46 | 8.71 | 7.05 | 4.83 | 6.02 | 4.3 | 2.23 | 2.9 | 3.2 | - |
| Nek6 | 83.45 | 65.54 | 77.09 | 39.32 | 46.45 | 45.59 | 27.15 | 49.45 | 46.7 | 21.81 | 22.45 | 17.39 | - |
| Stk24 | 7.01 | 8.88 | 7.29 | 11.29 | 9.65 | 10.4 | 10.25 | 8.9 | 9.48 | 13.27 | 12.01 | 11.01 | - |
| Ppp4r2 | 4.5 | 5.34 | 5.94 | 3.96 | 3.86 | 4.01 | 2.73 | 3.66 | 3.94 | 2.86 | 3.98 | 3.23 | - |
| Oip5 | 19.26 | 15.12 | 15.87 | 12.26 | 15.82 | 12.9 | 10.02 | 10.7 | 9.32 | 8.02 | 7.42 | 8.46 | - |
| Il20rb | 0.62 | 0.61 | 1.05 | 0.23 | 0.2 | 0.31 | 0.51 | 0.12 | 0.13 | 0.05 | 0.02 | 0.12 | - |
| Serpinb1b | 1.03 | 0.71 | 0.79 | 5.7 | 4.35 | 2.71 | 0.87 | 1.33 | 1.5 | 4.54 | 3.44 | 4.03 | - |
| Snrk | 3.73 | 4.18 | 3.32 | 5.95 | 5.65 | 6.25 | 7.76 | 6.35 | 6.96 | 9.25 | 7.07 | 8.37 | - |
| Miip | 29.63 | 19 | 23.34 | 41.49 | 42.14 | 40.99 | 31.41 | 28.38 | 27.14 | 39.54 | 39.57 | 41.34 | - |
| Tceal3 | 0.15 | 0.26 | 0 | 2.44 | 1.62 | 1.77 | 0.41 | 0.63 | 0.29 | 1.28 | 0.72 | 1.23 | - |
| Nfkbia | 135.04 | 120.87 | 137.86 | 190.59 | 217.69 | 194.47 | 136.2 | 132.17 | 120.81 | 178.09 | 180.95 | 171.2 | - |
| Sec13 | 213.27 | 206.25 | 212.62 | 124.03 | 135.68 | 137.35 | 118.11 | 159.33 | 150.35 | 98.5 | 107.93 | 101.63 | - |
| Hbegf | 8.32 | 8.62 | 8.99 | 2.82 | 3.92 | 3.52 | 1.74 | 3.96 | 4.22 | 0.65 | 0.99 | 0.97 | - |
| Fnbp4 | 16.36 | 17.15 | 16.89 | 24.43 | 23.8 | 22.35 | 21.94 | 20.6 | 20.75 | 30.73 | 32.01 | 27.21 | - |
| Ms4a4c | 13.02 | 12.67 | 10.01 | 3.4 | 3.96 | 4.18 | 6.16 | 2.86 | 2.5 | 1.77 | 0.63 | 2.2 | - |
| Zfp783 | 2.41 | 2.4 | 2.07 | 2.34 | 2.16 | 1.58 | 0.74 | 1.72 | 1.56 | 1.17 | 0.64 | 0.98 | - |
| Cd52 | 373.29 | 389.97 | 278.15 | 505.27 | 450.49 | 443.18 | 730.93 | 483.04 | 545.46 | 614.94 | 575.42 | 705.32 | - |
| Macf1 | 5.5 | 5.21 | 5.15 | 7.33 | 5.91 | 6.29 | 7.94 | 5.87 | 6.6 | 8.82 | 8.38 | 8.94 | - |
| Lyst | 1.19 | 1.13 | 0.8 | 1.27 | 1 | 0.91 | 2.03 | 1.17 | 1.62 | 2.26 | 1.97 | 2.02 | - |
| D430020J02Rik | 1.97 | 2.6 | 3.08 | 1.38 | 1.64 | 1.68 | 0.77 | 1.1 | 1.13 | 0.63 | 0.47 | 0.28 | - |
| Smyd2 | 54.49 | 60.4 | 62.72 | 33.79 | 38.48 | 38.38 | 31.19 | 44.72 | 40.58 | 24.13 | 22.19 | 24.1 | - |
| Zfp277 | 21.78 | 25.01 | 22.15 | 39.12 | 33.62 | 34.91 | 41.93 | 30.51 | 35.09 | 45.01 | 44.76 | 46.15 | - |
| Lztfl1 | 6.03 | 5.05 | 6.43 | 7.24 | 7.61 | 6.24 | 10.33 | 9.26 | 10.6 | 10.72 | 10.23 | 10.34 | - |
| Prmt10 | 1.91 | 1.44 | 1.5 | 2.68 | 1.89 | 2.5 | 1.88 | 1.77 | 1.68 | 3.02 | 2.73 | 3.46 | - |
| Gpsm1 | 1.85 | 1.63 | 1.93 | 3.25 | 2.68 | 2.69 | 3.93 | 3.02 | 2.7 | 3.79 | 3.62 | 3.65 | - |
| Hapln4 | 0.81 | 0.76 | 1.07 | 0.48 | 1.01 | 0.36 | 0.8 | 0.92 | 0.78 | 0.43 | 0.12 | 0.32 | - |
| Capn11 | 0.04 | 0.02 | 0.02 | 0.07 | 0.1 | 0.12 | 0.09 | 0.02 | 0.02 | 0.21 | 0.22 | 0.49 | - |
| Tmtc4 | 4.04 | 3.57 | 4 | 2.37 | 2.76 | 2.88 | 1.78 | 2.43 | 2.26 | 0.98 | 0.87 | 2.04 | - |
| Zzz3 | 5.91 | 5.21 | 5.32 | 4.87 | 4.98 | 4.02 | 3.54 | 3.98 | 4.01 | 3.73 | 2.67 | 3.56 | - |
| Tmem181b-ps | 11.21 | 11.06 | 10.42 | 12.73 | 12.56 | 12.78 | 14.25 | 13.54 | 13.07 | 16.03 | 14.54 | 15.03 | - |
| Rfc4 | 52.19 | 54.13 | 49.6 | 31.55 | 38.79 | 36.71 | 26.61 | 29.02 | 28.17 | 18.04 | 22.88 | 23.12 | - |
| Jakmip1 | 5.3 | 7.5 | 5.63 | 14.25 | 11.77 | 11.24 | 21.23 | 10.11 | 13.58 | 19.86 | 16.04 | 25.75 | - |
| Mdm1 | 4.32 | 4.38 | 3.66 | 2.45 | 3.18 | 3.1 | 2.41 | 3.2 | 2.85 | 2.25 | 1.83 | 2.13 | - |
| H2-Q10 | 104.18 | 83.9 | 90.6 | 76.75 | 86.48 | 78.63 | 155.65 | 169.16 | 161.6 | 133.93 | 139.11 | 132.43 | - |
| Slc31a1 | 17.8 | 20.4 | 16.66 | 12.8 | 14.63 | 15.46 | 13.66 | 13.4 | 11.96 | 11.45 | 10.57 | 12.57 | - |
| P2ry4 | 0 | 0 | 0 | 0.01 | 0.01 | 0.11 | 0 | 0.01 | 0.02 | 0.13 | 0.07 | 0.12 | - |
| Tnks1bp1 | 0.31 | 0.29 | 0.34 | 0.68 | 0.69 | 0.44 | 0.97 | 0.43 | 0.71 | 0.84 | 0.5 | 0.85 | - |
| Plekha4 | 0.02 | 0.13 | 0.16 | 0.4 | 0.35 | 0.26 | 0.43 | 0.44 | 0.26 | 0.39 | 0.56 | 0.56 | - |
| Whamm | 0.78 | 0.76 | 0.79 | 1.52 | 1.18 | 1.43 | 2.36 | 1.28 | 1.28 | 1.55 | 2.07 | 2.22 | - |
| Spink2 | 11.57 | 9.41 | 9.01 | 3.41 | 3.98 | 6.15 | 11.27 | 12.38 | 12.97 | 4.21 | 4.73 | 5.56 | - |
| Stag2 | 9.9 | 7.64 | 8.84 | 12.31 | 12.81 | 13.38 | 11.16 | 11.05 | 11.84 | 14.13 | 14.47 | 12.19 | - |
| Dhrs13 | 9.85 | 12.31 | 9.7 | 7.94 | 7.48 | 5.93 | 5.67 | 7.37 | 7.84 | 4.99 | 5.58 | 5.92 | - |
| Harbi1 | 5.04 | 5.21 | 6.04 | 7.82 | 8.46 | 9.15 | 7.72 | 7.89 | 8.42 | 9.45 | 8.51 | 9.29 | - |
| Fancl | 13.36 | 12.36 | 12.91 | 11.52 | 10.89 | 12.91 | 9.92 | 12.1 | 11.11 | 9.6 | 8.22 | 9.38 | - |
| Spred1 | 0.93 | 0.56 | 1.1 | 0.59 | 0.65 | 0.79 | 0.42 | 0.94 | 0.9 | 0.32 | 0.35 | 0.27 | - |
| Gm4858 | 1.03 | 0.38 | 0.58 | 1.18 | 2.36 | 1.61 | 0.21 | 0.45 | 0.2 | 0 | 0.1 | 0.06 | - |
| Fkbp15 | 8.59 | 10.02 | 10.03 | 11.84 | 12.1 | 11.63 | 12.86 | 11.49 | 11.71 | 12.19 | 14.11 | 13.53 | - |
| Herpud2 | 2.68 | 2.97 | 2.54 | 5.18 | 4.64 | 4.09 | 3.81 | 4.97 | 3.93 | 6.32 | 5.39 | 4.2 | - |
| Npl | 2.16 | 1.7 | 1.85 | 1.13 | 1.45 | 1.32 | 0.98 | 1.17 | 1.68 | 0.65 | 0.35 | 1.03 | - |
| Nus1 | 5.81 | 7.25 | 6.33 | 4.45 | 4.73 | 4.44 | 4.41 | 4.91 | 4.84 | 4.52 | 3.54 | 3.55 | - |

| | | | | | | | | | | | | |
|---|---|---|---|---|---|---|---|---|---|---|---|---|
| Fkrp | 8.28 | 7.53 | 7.15 | 5.07 | 5.76 | 3.89 | 5.39 | 5.13 | 4.76 | 4.61 | 2.68 | 4.41 | - |
| Alas1 | 22.29 | 21.04 | 24.23 | 16.8 | 17.3 | 16.47 | 16.42 | 16.76 | 15.11 | 11.98 | 12.93 | 13.42 | - |
| Ghr | 0 | 0 | 0 | 0 | 0.03 | 0 | 0.03 | 0.16 | 0 | 0.19 | 0.11 | 0.43 | - |
| Mrpl52 | 85.36 | 84.74 | 91.99 | 109.46 | 115.15 | 109.74 | 102.8 | 113.91 | 113.44 | 108.58 | 108.3 | 117.25 | - |
| Smad4 | 2.84 | 3.17 | 2.67 | 5.22 | 4.26 | 5.15 | 3.34 | 4.24 | 2.92 | 5.88 | 6.33 | 5.04 | - |
| C77080 | 0.29 | 0.28 | 0.5 | 0.06 | 0.06 | 0.22 | 0.18 | 0.26 | 0.22 | 0.09 | 0.05 | 0.13 | - |
| Papss2 | 1.17 | 0.98 | 1.53 | 0.44 | 0.47 | 0.72 | 0.13 | 0.44 | 0.24 | 0.08 | 0.24 | 0.44 | - |
| Shmt2 | 187.62 | 195.94 | 198.32 | 106.25 | 137.59 | 117.48 | 106.61 | 166.35 | 149.88 | 65.78 | 72.3 | 69.71 | - |
| Cdk1 | 72.15 | 55.22 | 48.76 | 31.69 | 44.37 | 38.72 | 21.85 | 26.09 | 22.66 | 21.56 | 19.59 | 17.43 | - |
| Mcm8 | 8.1 | 7.44 | 9.21 | 4.59 | 5.88 | 4.64 | 2.57 | 4.28 | 3.43 | 2.79 | 1.75 | 2.98 | - |
| Cyr61 | 2.59 | 0.89 | 1.87 | 0 | 1.91 | 1.19 | 0.09 | 0.9 | 0.26 | 0.25 | 0.06 | 0.03 | - |
| Jak2 | 2.85 | 3.08 | 4.52 | 4.07 | 4.91 | 4.36 | 4.69 | 5.76 | 5.12 | 5.92 | 5.72 | 5.94 | - |
| Uros | 6.87 | 6.36 | 6.12 | 3.63 | 3.65 | 4.08 | 2.86 | 2.27 | 3.47 | 1.93 | 2.08 | 2.67 | - |
| Taf6 | 31.51 | 30.53 | 31.12 | 27.36 | 32.98 | 27.83 | 25.05 | 27.29 | 24.88 | 20.45 | 24.72 | 25.44 | - |
| Zfp617 | 8.41 | 8.55 | 7.84 | 11.74 | 9.73 | 9.16 | 9.29 | 8.59 | 10.22 | 11.12 | 9.81 | 11.32 | - |
| Akap5 | 0.17 | 0.21 | 0.29 | 0.46 | 0.21 | 0.44 | 0.3 | 0.33 | 0.51 | 1.05 | 0.48 | 0.61 | - |
| P4ha1 | 39.35 | 36.72 | 37.4 | 39.84 | 39.43 | 45.65 | 62.48 | 52.84 | 58.71 | 47.21 | 61.93 | 56.34 | - |
| Gm4926 | 0 | 0 | 0 | 0.59 | 0.23 | 0.39 | 1.03 | 0.73 | 0.63 | 0.49 | 0.23 | 0.36 | - |
| Srrm2 | 20.97 | 20.47 | 22.56 | 29 | 27.47 | 28.65 | 31.59 | 30.62 | 31.22 | 37.42 | 35.34 | 34.86 | - |
| Nfatc2 | 3.87 | 4.93 | 4.04 | 6.23 | 6.13 | 5.07 | 5.5 | 5.21 | 6.26 | 6.26 | 5.63 | 6.16 | - |
| Klhl6 | 31.2 | 34.51 | 29.56 | 78.05 | 58.91 | 69.18 | 116.38 | 76.07 | 77.72 | 135.88 | 133.88 | 144.04 | - |
| Snhg10 | 0.97 | 2.97 | 2.25 | 4.18 | 4.01 | 3.01 | 3.04 | 3.77 | 2.94 | 6.49 | 6.17 | 4.14 | - |
| Frat1 | 1.14 | 1.54 | 1.47 | 2.76 | 2.63 | 2.12 | 2.9 | 1.89 | 2.79 | 3.52 | 3.47 | 4.65 | - |
| Hmgxb3 | 7.3 | 9.22 | 9.64 | 7.49 | 7.75 | 7.66 | 5.19 | 8.03 | 7.58 | 5.34 | 5.88 | 5.68 | - |
| Chsy1 | 20.53 | 22.68 | 19.62 | 20.63 | 17 | 20.33 | 14.64 | 14.28 | 16.05 | 14.77 | 15.29 | 16.59 | - |
| Tmem191c | 2.31 | 0.87 | 1.08 | 4.4 | 2.93 | 3.08 | 8.35 | 3.38 | 6.78 | 8.29 | 8.19 | 7.57 | - |
| Cacnb4 | 0.07 | 0.02 | 0.04 | 0.14 | 0.09 | 0.06 | 0.09 | 0.08 | 0.07 | 0.34 | 0.25 | 0.22 | - |
| Fbln5 | 0.06 | 0.1 | 0.17 | 0.08 | 0 | 0 | 0.03 | 0.11 | 0.03 | 0 | 0 | 0 | - |
| Armc7 | 5.2 | 5.36 | 6.68 | 8.12 | 7.16 | 7.06 | 9.35 | 6.19 | 7.47 | 9.74 | 11.51 | 13.17 | - |
| Sos1 | 1.33 | 1.44 | 1.4 | 1.57 | 1.79 | 1.82 | 1.9 | 1.61 | 1.69 | 1.93 | 2.32 | 1.91 | - |
| Mxd1 | 2.88 | 2.91 | 3.2 | 19.17 | 13.82 | 14.51 | 12.36 | 7.83 | 7.94 | 27.97 | 21.73 | 32.8 | - |
| Il23a | 42.64 | 40.56 | 32.12 | 2.56 | 6.89 | 8.77 | 1.18 | 10.29 | 6.59 | 4.17 | 1.13 | 0.21 | - |
| Gm15421 | 5.4 | 5.7 | 5.46 | 9.28 | 9.18 | 10.35 | 7.02 | 6.36 | 8.7 | 9.76 | 9.96 | 10.3 | - |
| Ell2 | 4.97 | 5.09 | 4.86 | 3.98 | 3.75 | 3.08 | 2.5 | 3.46 | 2.46 | 2.67 | 2.9 | 1.95 | - |
| Zfp97 | 11.49 | 11.34 | 10.2 | 19.64 | 16.19 | 16.96 | 21.15 | 15.42 | 14.55 | 28.17 | 23.79 | 28.01 | - |
| Copb1 | 64.8 | 62.59 | 61.7 | 65.21 | 73.2 | 71.52 | 66.34 | 72.94 | 65.89 | 76.44 | 85.65 | 72.45 | - |
| 1110059E24Rik | 47.41 | 47.8 | 47.7 | 49.48 | 45.81 | 43.45 | 41.02 | 39.48 | 41.26 | 34.05 | 38.66 | 31.76 | - |
| S1pr4 | 13.14 | 16.13 | 13.59 | 16.85 | 15.32 | 14.95 | 32.38 | 17.69 | 17.86 | 27.2 | 24.28 | 35.22 | - |
| Cnn3 | 46.41 | 43.66 | 49.93 | 39.26 | 43.72 | 44.1 | 41.14 | 53.9 | 48.45 | 33.59 | 39.73 | 34.29 | - |
| Crtc3 | 0.62 | 1.02 | 0.56 | 1.7 | 1.81 | 1.43 | 0.86 | 1.12 | 1.3 | 1.75 | 1.35 | 1.39 | - |
| Nisch | 23.01 | 22.99 | 26.33 | 34.88 | 33.6 | 35.85 | 33.9 | 29.84 | 31.61 | 40.47 | 44.84 | 42.32 | - |
| Acrbp | 1.09 | 0.83 | 0.77 | 3.92 | 2.97 | 3.75 | 4.06 | 2.99 | 2.47 | 5.68 | 6.17 | 8.24 | - |
| A930004D18Rik | 0.51 | 0.48 | 1.02 | 0.51 | 0.54 | 0.33 | 0.09 | 0.25 | 0.17 | 0.15 | 0.14 | 0.27 | - |
| Osbpl7 | 5.68 | 5.28 | 5.26 | 8.01 | 6.89 | 8.53 | 7.25 | 8.01 | 7.81 | 8.82 | 10.91 | 9.21 | - |
| St3gal1 | 12.36 | 9.72 | 13.16 | 13.39 | 15.07 | 15.59 | 20 | 23.24 | 21.59 | 24.14 | 27.97 | 23.31 | - |
| Agps | 4.89 | 4.93 | 5.63 | 4.88 | 5.31 | 4.36 | 3.4 | 4.18 | 4.35 | 3.67 | 4.63 | 3.59 | - |
| B4galt3 | 21.5 | 21.97 | 22.6 | 26.16 | 23.12 | 28.41 | 28.7 | 29.97 | 28.11 | 28.46 | 28.78 | 29.03 | - |
| Nipsnap3b | 45.59 | 48.72 | 50.08 | 42.19 | 39.73 | 41 | 43.86 | 40.07 | 43.35 | 37.43 | 35.59 | 38.1 | - |
| Tmem234 | 50.4 | 50.08 | 48.87 | 60.48 | 53.48 | 57.52 | 70.6 | 55.33 | 61.73 | 70.82 | 70.92 | 77.9 | - |
| Cd226 | 11.88 | 12.39 | 11.97 | 17.11 | 14.84 | 16.45 | 4.67 | 4.02 | 4.33 | 3.34 | 4.17 | 5.13 | - |
| Emilin2 | 15.2 | 22.75 | 16.19 | 10.1 | 7.17 | 7.88 | 34.38 | 25.24 | 37.95 | 13.09 | 11.77 | 12.73 | - |
| Mapk8ip1 | 0.18 | 0.1 | 0.13 | 0.52 | 0.23 | 0.52 | 0.72 | 0.42 | 0.22 | 0.53 | 0.4 | 0.71 | - |
| Itgb3bp | 2.86 | 3.1 | 2.49 | 3.01 | 2.87 | 2.79 | 1.57 | 1.39 | 1.41 | 1.63 | 1.42 | 1.79 | - |
| Ccdc114 | 0.05 | 0.07 | 0.11 | 0.31 | 0.46 | 0.31 | 0.75 | 0.31 | 0.22 | 0.37 | 0.53 | 0.78 | - |
| Kctd2 | 1.07 | 0.76 | 0.73 | 2.58 | 1.56 | 1.84 | 1.89 | 1.83 | 1.06 | 2.2 | 2.65 | 2.42 | - |
| Iah1 | 51.91 | 51.61 | 56.65 | 34.52 | 36.91 | 36.96 | 40.28 | 57.82 | 48.66 | 28.71 | 30.05 | 24.76 | - |
| Dapp1 | 6.99 | 5.48 | 5.52 | 9.27 | 9.76 | 9.66 | 10.79 | 8.47 | 7.89 | 10.67 | 12.42 | 10.6 | - |
| Larp1 | 11.39 | 10.82 | 11.11 | 7.09 | 6.72 | 7.45 | 5.98 | 7.81 | 8.12 | 4.55 | 5.42 | 5.08 | - |
| Rps17 | 1834.43 | 1885.81 | 1919.93 | 1734.63 | 1855.82 | 1889.77 | 1594.94 | 1758.79 | 1650.59 | 1409.8 | 1650.95 | 1403.5 | - |
| Zscan21 | 9.45 | 8.7 | 8.77 | 13.91 | 11.79 | 12.96 | 11.15 | 10.27 | 11.71 | 13.38 | 13.08 | 12.59 | - |
| Abhd8 | 30.81 | 29.05 | 30.04 | 48.65 | 42.7 | 43.82 | 54.31 | 40.83 | 40.04 | 53.06 | 55.39 | 53.5 | - |
| Ube2d1 | 5.07 | 6.91 | 6.73 | 9.01 | 7.2 | 5.82 | 9.08 | 6.85 | 7.21 | 9.27 | 10.29 | 8.62 | - |
| AI467606 | 27.53 | 28.31 | 26.7 | 37.29 | 35.41 | 34.91 | 53.15 | 35.89 | 39.72 | 53.53 | 41.61 | 60.68 | - |
| Suz12 | 7 | 7.06 | 7.16 | 6.37 | 7.19 | 6.12 | 5.8 | 5.35 | 5.57 | 5.28 | 5.77 | 4.65 | - |
| Blm | 8.55 | 7.03 | 8.4 | 5.96 | 6.12 | 6 | 5.1 | 4.32 | 5.39 | 4.34 | 3.46 | 3.7 | - |
| Panx1 | 4.43 | 6.14 | 7.66 | 10.51 | 10.77 | 10.88 | 11.8 | 12.77 | 11.86 | 17.34 | 14.04 | 14.11 | - |
| Nup62 | 74.78 | 79.98 | 75.47 | 53 | 64.41 | 56.24 | 49.01 | 60.98 | 53.79 | 40.74 | 44 | 39.61 | - |
| Psmg1 | 64.22 | 70.77 | 65.32 | 40.4 | 43.85 | 46.49 | 36 | 47.94 | 42 | 27.01 | 30.58 | 28.21 | - |
| Zfp512 | 6.48 | 7.26 | 6.03 | 13.99 | 12.7 | 11.45 | 10.98 | 9.97 | 12.25 | 16 | 13.37 | 15.77 | - |
| Fosl1 | 3.55 | 1.53 | 3.06 | 0.81 | 1.43 | 0.86 | 0.32 | 0.51 | 1.09 | 0.47 | 0.56 | 0.21 | - |
| Polr3d | 23.93 | 21.97 | 25.96 | 11.98 | 13.18 | 14.24 | 8.72 | 15.08 | 13.88 | 8.55 | 9.56 | 8.46 | - |
| Gspt2 | 2.61 | 2.59 | 2.21 | 1.27 | 1.28 | 1.25 | 0.99 | 1.25 | 1.63 | 0.8 | 0.47 | 0.46 | - |
| Nfkb1 | 21.53 | 23.03 | 22.37 | 21.43 | 24.07 | 21.79 | 13.33 | 14.79 | 15.15 | 13.07 | 13.56 | 13.56 | - |
| Grk4 | 16.84 | 19 | 19.98 | 31.2 | 30.13 | 27.04 | 29.05 | 24.49 | 23.94 | 34.82 | 31.37 | 29.86 | - |
| Ptgr2 | 12.92 | 13.03 | 13.26 | 11.26 | 11.98 | 10.38 | 10.04 | 9.13 | 11.19 | 8.64 | 8.86 | 9.57 | - |
| Mettl23 | 61.21 | 58.47 | 59.66 | 76.8 | 70.47 | 70.44 | 97.68 | 75.8 | 85.41 | 96.95 | 91.16 | 96.8 | - |
| Carm1 | 7.08 | 8.07 | 7.24 | 5.69 | 5.08 | 3.92 | 4.12 | 4.56 | 5.08 | 4.58 | 3.41 | 3.74 | - |
| Cd109 | 0.09 | 0.07 | 0.09 | 0.05 | 0.07 | 0.01 | 0.27 | 0.31 | 0.23 | 0.29 | 0.26 | 0.28 | - |
| Zfp202 | 2.22 | 3.82 | 3.85 | 2.26 | 2.62 | 1.76 | 2.25 | 1.81 | 1.8 | 1.72 | 1.59 | 2.12 | - |
| Carkd | 42.94 | 45 | 41.81 | 39.22 | 40.25 | 42.66 | 33.62 | 40.93 | 42.45 | 33.17 | 33.73 | 32.3 | - |
| Git2 | 7.39 | 9.21 | 8.83 | 13.72 | 10.96 | 11.76 | 13.44 | 11.49 | 11.4 | 14.92 | 14.56 | 18.02 | - |
| Rgs9 | 0.05 | 0.01 | 0.03 | 0.22 | 0.3 | 0.32 | 0.26 | 0.25 | 0.22 | 0.17 | 0.27 | 0.46 | - |
| Dnajc10 | 26.19 | 24.19 | 25.88 | 19.78 | 24.13 | 20.91 | 16.75 | 23.41 | 22.72 | 17.13 | 16.4 | 18.07 | - |
| Ndufb4 | 136.31 | 143.68 | 141.12 | 101.35 | 112.2 | 105.91 | 97.97 | 117.86 | 110.37 | 76.72 | 85.91 | 93.53 | - |
| Rps6kl1 | 2.12 | 1.08 | 1.6 | 0.85 | 0.44 | 0.63 | 1.22 | 1.34 | 1.23 | 0.23 | 0.36 | 0.52 | - |
| Tgfb2 | 0.33 | 0.33 | 0.31 | 0.03 | 0.05 | 0.01 | 0.11 | 0.11 | 0.11 | 0 | 0.07 | 0.01 | - |
| Rhbdd2 | 7.06 | 6.09 | 7.87 | 6.44 | 7.03 | 6.09 | 5.63 | 6.5 | 5.99 | 3.99 | 3.77 | 4.81 | - |
| Hhat | 3.42 | 3.22 | 3.51 | 13.73 | 11.87 | 14.71 | 2.47 | 5.51 | 3.94 | 10.59 | 13.91 | 13.81 | - |
| Klf6 | 20.79 | 18.33 | 22.35 | 24.35 | 26.08 | 24.79 | 28.67 | 32.17 | 31.48 | 29.99 | 30.09 | 31.19 | - |
| Psmb6 | 298.47 | 302.29 | 291.21 | 193.46 | 201.66 | 196.28 | 176.28 | 236.81 | 201.95 | 142.3 | 156.51 | 138.77 | - |
| Esyt2 | 4.55 | 3.99 | 4.33 | 4.61 | 5.62 | 6.04 | 6.09 | 4.98 | 6.04 | 7.44 | 7.13 | 6.82 | - |
| Blzf1 | 6.34 | 6.17 | 6.3 | 5.9 | 4.23 | 4.48 | 4.32 | 5.16 | 4.75 | 2.93 | 4.02 | 3.46 | - |
| Ifit1 | 0.88 | 0.54 | 0.51 | 1.86 | 0.78 | 2.82 | 0.71 | 3.98 | 0.75 | 3.16 | 1.39 | 3.05 | - |
| Kctd14 | 3.21 | 2.06 | 1.65 | 0.79 | 1.37 | 1.6 | 1.42 | 1.9 | 2.05 | 1.34 | 1.11 | 1.03 | - |
| Pja2 | 7.76 | 5.69 | 7.69 | 9.6 | 11.53 | 10.75 | 9.09 | 9.87 | 9.96 | 11.47 | 11.17 | 9.96 | - |
| E2f7 | 1.35 | 1.13 | 0.78 | 0.9 | 0.73 | 0.44 | 0.21 | 0.28 | 0.51 | 0.34 | 0.19 | 0.41 | - |
| Rnaset2a | 88.15 | 108.48 | 118.41 | 149.87 | 144.12 | 138.31 | 154.89 | 148.06 | 152.82 | 154.57 | 136.17 | 154.26 | - |
| Wdr70 | 16.76 | 16.47 | 20.22 | 14.25 | 13.94 | 14.97 | 14.61 | 14.68 | 14.34 | 12.27 | 13.44 | 14.37 | - |
| Ankmy2 | 7.42 | 7.65 | 6.55 | 11.5 | 8.8 | 8.19 | 8.62 | 7.88 | 7.57 | 10.08 | 9.2 | 10.78 | - |
| Zfp13 | 0 | 0.05 | 0.03 | 0.62 | 0.38 | 0.75 | 0.51 | 0.52 | 0.4 | 0.59 | 0.49 | 0.38 | - |
| Plp2 | 216.34 | 225.33 | 220.76 | 338.09 | 291.89 | 302.61 | 297.76 | 256.48 | 276.77 | 335.33 | 305.32 | 336.71 | - |
| Tnfaip8l2 | 12.47 | 10.49 | 12.72 | 18.98 | 16.25 | 18.98 | 25.86 | 19.53 | 19.27 | 25.39 | 17.69 | 22.76 | - |
| Zfp319 | 2.72 | 2.39 | 2.62 | 3.84 | 3.83 | 3.82 | 4.28 | 3.05 | 3.81 | 5.06 | 3.53 | 5.14 | - |
| Slc30a9 | 4.54 | 6.36 | 7.53 | 9.84 | 9.7 | 9.22 | 10.97 | 11.04 | 9.45 | 11.12 | 10.26 | 11.2 | - |
| Ift140 | 3.48 | 2.88 | 4.09 | 5.59 | 5.62 | 5.54 | 3.99 | 3.84 | 4.29 | 5.28 | 5.66 | 3.97 | - |
| Acp6 | 13.43 | 13.19 | 13.49 | 26.87 | 21.87 | 21.9 | 27.15 | 19.8 | 19.98 | 28.44 | 24.88 | 27.34 | - |
| Abcd1 | 6.26 | 7.89 | 5.48 | 8.7 | 7.47 | 6.53 | 9.26 | 6.37 | 7.19 | 9.05 | 7.79 | 11.27 | - |
| Ndufaf6 | 22.94 | 28.93 | 32.08 | 20.47 | 22.19 | 21.97 | 22.72 | 33.17 | 27.21 | 18.06 | 22.6 | 18.47 | - |
| Cxcr1 | 0.3 | 0.1 | 0.1 | 0.45 | 0.28 | 1.19 | 1.58 | 0.37 | 1.15 | 1.53 | 0.55 | 0.77 | - |
| Zfp799 | 1.51 | 1.64 | 1.47 | 2.71 | 1.85 | 2.07 | 2.15 | 1.52 | 2.26 | 2.53 | 2.18 | 2.56 | - |
| Arf5 | 18.05 | 31.37 | 28.63 | 31.97 | 30.31 | 25.22 | 38.93 | 33.55 | 39.45 | 39.29 | 36.66 | 34.47 | - |
| Atr | 1.94 | 1.95 | 1.69 | 2.97 | 2.88 | 2.64 | 4.09 | 2.66 | 3.36 | 3.59 | 3.96 | 3.57 | - |
| Pot1b | 2.57 | 2.39 | 3.07 | 3.43 | 3.69 | 3.58 | 3.81 | 2.93 | 3.33 | 4.98 | 5.08 | 4.14 | - |
| Arhgef12 | 0.6 | 0.64 | 0.55 | 0.75 | 0.59 | 0.53 | 0.89 | 0.69 | 0.86 | 1.14 | 1 | 1.47 | - |
| Irf9 | 23.79 | 25.57 | 25.36 | 32.17 | 35.28 | 33.94 | 29.6 | 28.11 | 26.92 | 40.18 | 36.09 | 47.38 | - |
| Tmem45a | 0.28 | 0.63 | 0.52 | 1.41 | 1.25 | 1.44 | 2.59 | 0.89 | 1.59 | 2.78 | 2.03 | 3.74 | - |
| Zfp458 | 0.56 | 0.49 | 0.36 | 0.09 | 0.11 | 0.04 | 0 | 0.22 | 0.34 | 0 | 0 | 0 | - |
| Nat2 | 5.74 | 6.06 | 5 | 7.55 | 8.73 | 7.99 | 12.26 | 7.14 | 9.14 | 16.21 | 13.55 | 14.53 | - |
| Irak2 | 4.37 | 3.49 | 3.62 | 9.96 | 8.53 | 8.79 | 9.98 | 7.59 | 9.35 | 16.39 | 14.88 | 17.6 | - |
| Klra7 | 0.16 | 0.05 | 0.35 | 9.76 | 12.14 | 10.2 | 1.71 | 4.69 | 4.84 | 30.43 | 27.38 | 20.52 | - |
| Lamtor4 | 28.51 | 28.93 | 28.05 | 29.76 | 27.05 | 30.11 | 37.87 | 32.08 | 37.38 | 38.88 | 37.67 | 40.75 | - |
| Snrnp27 | 77.09 | 75.31 | 74.15 | 63.36 | 64.29 | 73.54 | 58.84 | 67.38 | 66.21 | 57.21 | 62.29 | 57.57 | - |
| Oaz1-ps | 524.05 | 529.88 | 545.66 | 435.48 | 448.33 | 458.42 | 488.57 | 507.59 | 491 | 441.39 | 403 | 447.65 | - |
| Zranb3 | 2.14 | 2.69 | 1.74 | 2.43 | 2.7 | 1.69 | 1.04 | 1.57 | 1.45 | 0.55 | 0.99 | 0.99 | - |
| Alg2 | 9.03 | 9.89 | 9.82 | 8.4 | 9.26 | 9.89 | 6.81 | 10.3 | 8.17 | 4.42 | 6.23 | 6.32 | - |
| Pgam1 | 916.2 | 923.27 | 956.96 | 761.18 | 739.94 | 768.77 | 1030.72 | 855.73 | 970.8 | 711.11 | 777.31 | 837.71 | - |
| Chmp1a | 52.87 | 48 | 52.28 | 55.71 | 53.22 | 58.63 | 74.28 | 59.3 | 65.09 | 77.81 | 76.14 | 79.04 | - |
| Fitm2 | 4.05 | 3.61 | 3.59 | 2.85 | 2.11 | 2.76 | 2.03 | 2.61 | 1.89 | 1.64 | 1.2 | 1.25 | - |
| Chchd1 | 161.29 | 160.97 | 151.39 | 100.28 | 99.88 | 112.65 | 104.2 | 127.54 | 123.02 | 81.92 | 89.5 | 79.15 | - |
| Tpgs1 | 23.59 | 24.95 | 22.54 | 29.1 | 26.42 | 29.42 | 33.04 | 30.51 | 29.09 | 35.76 | 33.16 | 39.89 | - |
| Fbxl15 | 4.7 | 5.56 | 4.55 | 2.84 | 2.8 | 2.53 | 4.4 | 4 | 3.8 | 2.62 | 2.96 | 2.8 | - |
| Rbm7 | 82.48 | 76.35 | 72.62 | 84.22 | 80.24 | 88.15 | 94.31 | 79.35 | 84.73 | 98.19 | 106.18 | 92.83 | - |
| 0610009B22Rik | 19.25 | 19.25 | 17.27 | 12.56 | 15.47 | 14.92 | 13.44 | 14.41 | 12.82 | 10.22 | 11.52 | 13.7 | - |
| Atp1a3 | 0.32 | 0.33 | 0.37 | 0.36 | 0.29 | 0.62 | 1.45 | 0.77 | 0.48 | 0.96 | 0.78 | 1.1 | - |
| Vps13a | 3.8 | 3.86 | 3.85 | 5.21 | 4.46 | 4.86 | 4.96 | 4.48 | 4.97 | 5.42 | 5.82 | 5.29 | - |
| Neil3 | 7.17 | 8.55 | 7.68 | 6.06 | 7.04 | 8.15 | 3.57 | 4.41 | 3.67 | 2.31 | 3.64 | 4.22 | - |
| Stard10 | 2.43 | 2.04 | 1.94 | 2.9 | 2.66 | 3.41 | 4.37 | 4.56 | 4.75 | 6.21 | 7.15 | 5.63 | - |
| Arnt | 9.29 | 9.08 | 9.6 | 12.59 | 11.71 | 10.61 | 10.54 | 12.36 | 12.16 | 12.7 | 13.48 | 12.44 | - |
| Slfn1 | 8.02 | 10.25 | 7.63 | 17.7 | 15.37 | 14.29 | 9.44 | 4.9 | 5.78 | 20.05 | 14.56 | 21.23 | - |
| Jmjd1c | 2.27 | 2.31 | 2.52 | 4.2 | 3.54 | 3.53 | 3.54 | 2.51 | 3.4 | 4.25 | 5.27 | 4.43 | - |
| Clcf1 | 1.74 | 1.49 | 0.86 | 2.4 | 1.6 | 2.73 | 3.79 | 3.09 | 2.95 | 3.5 | 3.39 | 5 | - |
| Cpne3 | 9.36 | 9.07 | 10.75 | 8.23 | 8.86 | 8.04 | 7.19 | 7.47 | 8.34 | 8.63 | 7.42 | 7.23 | - |
| Ndrg3 | 35.42 | 32.92 | 33.79 | 46.89 | 43.53 | 45.36 | 51.04 | 47.52 | 45.04 | 51.36 | 53.61 | 56.3 | - |
| Nr4a1 | 18.03 | 22.56 | 24.29 | 13.12 | 24.18 | 14.4 | 7.03 | 8.11 | 8.39 | 9.87 | 6.99 | 6.43 | - |
| Dip2b | 2.43 | 2.36 | 2.64 | 3.67 | 3.13 | 3.09 | 3.06 | 3.17 | 3.46 | 4.17 | 4 | 4.13 | - |
| Chkb | 17.17 | 17.53 | 17.19 | 23.49 | 23.86 | 22.24 | 26.4 | 22.63 | 21.57 | 25.93 | 26.04 | 26.33 | - |
| Alkbh5 | 14.56 | 18.29 | 20.94 | 22.1 | 23.41 | 22.09 | 27.09 | 24.72 | 25.26 | 25.03 | 28.21 | 23.68 | - |
| Tbx6 | 0.1 | 0 | 0.16 | 1 | 0.73 | 0.48 | 0.47 | 0.62 | 0.63 | 1.4 | 1.8 | 0.91 | - |
| Gba | 32.98 | 30.02 | 28.79 | 33.87 | 34.53 | 36.04 | 42.74 | 38.63 | 33.3 | 40.55 | 36.84 | 39.73 | - |
| Gcnt2 | 1.81 | 2.75 | 1.4 | 1.45 | 1 | 0.89 | 3.1 | 3.01 | 2.83 | 0.43 | 0.68 | 1.18 | - |
| Fxyd7 | 0.08 | 0.26 | 0.16 | 8.2 | 2.21 | 2.16 | 19.49 | 0.61 | 5.29 | 23.1 | 11.54 | 24.48 | - |

| | | | | | | | | | | | | |
|---|---|---|---|---|---|---|---|---|---|---|---|---|
| Capn2 | 49.43 | 46.49 | 43.74 | 34.33 | 38.22 | 34.67 | 51.58 | 47.82 | 48.63 | 33.64 | 33.67 | 33.99 | - |
| Arid4b | 5.25 | 5.5 | 5.34 | 7.47 | 6.89 | 7.47 | 6.67 | 6.55 | 7.3 | 8.39 | 9.97 | 8.8 | - |
| Mtif2 | 11.53 | 9.83 | 10.22 | 8.5 | 9.44 | 9.35 | 7.63 | 7.82 | 8.54 | 8.62 | 6.19 | 7.2 | - |
| Sdhb | 280.39 | 285.11 | 275.9 | 197.74 | 217.77 | 210.81 | 190.82 | 238.36 | 225.52 | 157.63 | 167.03 | 153.42 | - |
| Gnl1 | 12.92 | 15.03 | 16.22 | 13.06 | 9.87 | 12.71 | 12.69 | 12.26 | 9.58 | 11.05 | 9.7 | 10.48 | - |
| Plin3 | 25.44 | 23.19 | 23.2 | 36.86 | 35.47 | 35.63 | 44.59 | 34.54 | 36.78 | 47.04 | 51.21 | 48.3 | - |
| Cep128 | 2.28 | 2.12 | 2.4 | 1.75 | 2.17 | 1.87 | 0.94 | 1.52 | 1.44 | 0.77 | 1.14 | 1.19 | - |
| Scin | 1.05 | 1.15 | 1.4 | 0.36 | 0.63 | 0.56 | 0.25 | 0.32 | 0.35 | 0.05 | 0 | 0.04 | - |
| H2-Ab1 | 25.7 | 30.14 | 27.78 | 76.71 | 50.39 | 56.34 | 189.44 | 64.4 | 102.45 | 158.05 | 167.87 | 177.27 | - |
| Mettl21d | 32.3 | 33.26 | 35.85 | 27.02 | 32.58 | 33.9 | 54.33 | 44.15 | 47.23 | 42.34 | 50.35 | 44.84 | - |
| Sdad1 | 10.58 | 13.08 | 11.14 | 8.11 | 8.97 | 9.53 | 8.4 | 8.95 | 10.94 | 6.87 | 6.38 | 7.54 | - |
| Prpf31 | 56.09 | 54.74 | 57.01 | 20.76 | 25.91 | 25.39 | 13.72 | 21.83 | 21.88 | 9.61 | 7.47 | 9.65 | - |
| Pla2g16 | 12.22 | 11.51 | 11.47 | 18.29 | 17.12 | 16.74 | 19.46 | 17.76 | 19.2 | 22.2 | 20.53 | 22.72 | - |
| Camk2b | 0.16 | 0.17 | 0.12 | 0.38 | 0.12 | 0.32 | 0.63 | 0.31 | 0.43 | 0.97 | 0.45 | 0.73 | - |
| Rprd2 | 3.93 | 3.5 | 4.2 | 4.35 | 5.15 | 4.88 | 4.47 | 4.81 | 3.96 | 5.97 | 5.33 | 6.44 | - |
| Tgds | 18.1 | 21.19 | 19.51 | 12.19 | 15.49 | 15.37 | 15.74 | 15.91 | 16.01 | 10.73 | 10.61 | 11.3 | - |
| Bbc3 | 0.55 | 0.74 | 0.67 | 2.2 | 1.55 | 2.1 | 1.7 | 1.31 | 1.06 | 4.9 | 4.03 | 6.04 | - |
| Lnpep | 14.63 | 13.83 | 14.97 | 24.41 | 23.89 | 24.06 | 30.53 | 22.44 | 25.39 | 32.91 | 34 | 31.25 | - |
| Trub2 | 11.58 | 11.67 | 11.85 | 6.09 | 8.05 | 8.49 | 5.2 | 9.22 | 7.88 | 5.74 | 5.77 | 6.14 | - |
| Chchd7 | 20.01 | 22.11 | 20.63 | 30.29 | 24.6 | 30.66 | 32.43 | 26 | 26.72 | 29.21 | 32.25 | 32.71 | - |
| Acbd4 | 3.38 | 4.71 | 2.92 | 5.02 | 4.93 | 4.15 | 8.36 | 6.61 | 6.28 | 6.81 | 6.38 | 7.97 | - |
| Pde3b | 6.09 | 6.52 | 5.9 | 12.05 | 9.24 | 11.1 | 11.25 | 8.75 | 9.27 | 15.54 | 13.73 | 13.78 | - |
| Ccnd2 | 131.87 | 114.87 | 121.71 | 87.38 | 106.26 | 97.64 | 69.6 | 104.11 | 98.46 | 54.29 | 51.97 | 54.47 | - |
| Ccl5 | 31.61 | 11.62 | 14.66 | 26.07 | 26.5 | 27.26 | 100.97 | 86.96 | 105.32 | 249.72 | 184.99 | 248.85 | - |
| Npcd | 0 | 0.06 | 0.06 | 0.82 | 0.5 | 0.8 | 0.41 | 0.83 | 0.72 | 0.46 | 0.8 | 1.51 | - |
| Brox | 15.74 | 15.71 | 15.74 | 23.21 | 23.2 | 20.35 | 25.84 | 22.24 | 25.19 | 24.04 | 28.58 | 27.58 | - |
| Abcb9 | 14.71 | 17.89 | 14 | 10.35 | 9.61 | 9.91 | 15.62 | 15.54 | 13.86 | 10.23 | 10.29 | 9.23 | - |
| Stra13 | 49.16 | 46.66 | 44.36 | 41.08 | 33.47 | 43.42 | 36.59 | 32.6 | 33.07 | 27.15 | 35.76 | 37.34 | - |
| Hist2h4 | 5.98 | 4.75 | 5.28 | 2.36 | 4.31 | 4.95 | 4.79 | 4.22 | 2.16 | 1.15 | 2.06 | 1.16 | - |
| Lrr1 | 6.58 | 5.74 | 7.02 | 2.19 | 3.47 | 2.12 | 2.57 | 1.13 | 1.14 | 1.07 | 0.7 | 0.42 | - |
| Pls1 | 0.43 | 0.3 | 0.42 | 0.84 | 0.95 | 0.48 | 2.37 | 2.42 | 0.93 | 4.01 | 4.53 | 2.6 | - |
| Rab19 | 11.06 | 11.5 | 10.27 | 18.52 | 14.89 | 15.69 | 25.23 | 13.61 | 16.9 | 20.92 | 18.1 | 25.7 | - |
| Slc12a7 | 4.25 | 3.3 | 4.42 | 5.79 | 5.33 | 6.12 | 8.08 | 5.9 | 6.53 | 8.29 | 7.12 | 7.68 | - |
| AI836003 | 13.67 | 10.57 | 11.63 | 12.54 | 16.67 | 13.96 | 3.55 | 7.64 | 6.06 | 4.91 | 5.05 | 4.43 | - |
| Evi5 | 0.75 | 0.55 | 0.8 | 1.04 | 0.86 | 1.19 | 1.16 | 1.74 | 1.55 | 1.45 | 1.49 | 1.24 | - |
| Rad9a | 28.68 | 25.2 | 27.62 | 25.56 | 23.65 | 25.45 | 21 | 22.1 | 23.35 | 23.83 | 20.68 | 21.13 | - |
| Ppp1r12c | 4.23 | 6.41 | 5.66 | 9.88 | 7.55 | 7.1 | 8.79 | 7.17 | 9.14 | 10.34 | 7.76 | 8.5 | - |
| Wdtc1 | 2.2 | 1.95 | 1.97 | 3.75 | 3.01 | 3.43 | 3.78 | 3.93 | 3.43 | 5.67 | 4.77 | 5 | - |
| 9530082P21Rik | 3.62 | 3.26 | 3.61 | 4.87 | 4.99 | 4.54 | 4.85 | 4.47 | 4.93 | 5.54 | 4.86 | 4.66 | - |
| Espl1 | 6.27 | 5.34 | 5.67 | 4.17 | 4.96 | 4.17 | 1.99 | 2.29 | 2.36 | 2.69 | 2.39 | 1.17 | - |
| Nol3 | 0.81 | 0.38 | 0.63 | 1.27 | 1.03 | 0.61 | 0.87 | 1.33 | 1.09 | 1.33 | 1.48 | 1.13 | - |
| Gar1 | 7 | 8.38 | 8.68 | 3.83 | 3.68 | 3.11 | 3.3 | 4.62 | 4.81 | 3.47 | 2.96 | 1.5 | - |
| Desi1 | 12.14 | 18.02 | 22 | 22.82 | 24.68 | 20.22 | 31.61 | 33.67 | 29.38 | 31.54 | 33.26 | 25.38 | - |
| Tnfsf4 | 3.8 | 3.44 | 3.66 | 2.28 | 2.68 | 2.77 | 1.4 | 3.25 | 1.7 | 2.12 | 1.19 | 1.83 | - |
| Selm | 1.9 | 2.47 | 1.49 | 3.99 | 2.79 | 3.95 | 23.33 | 9.63 | 14.46 | 16.2 | 14.21 | 21.22 | - |
| Polr3e | 17.86 | 17 | 16.37 | 11.23 | 12.57 | 12.95 | 9.45 | 13.27 | 10.5 | 8.95 | 9.18 | 7.55 | - |
| Rttn | 1.08 | 1.19 | 0.98 | 0.83 | 1.08 | 0.76 | 0.99 | 0.72 | 0.69 | 0.71 | 0.47 | 0.42 | - |
| Tnip3 | 8.22 | 6.53 | 5.07 | 1.04 | 1.77 | 1.42 | 1.77 | 2.35 | 1.75 | 0.38 | 0.39 | 0.47 | - |
| Agap1 | 0 | 0.01 | 0.01 | 0.03 | 0.07 | 0.07 | 0.13 | 0.07 | 0.14 | 0.04 | 0.06 | 0.11 | - |
| Cpm | 1.89 | 2.07 | 2.58 | 2.37 | 2.06 | 1.94 | 6.85 | 5.75 | 6.29 | 4.61 | 3.56 | 4.23 | - |
| Uck1 | 29.09 | 28.29 | 28.41 | 33.89 | 33.66 | 36.9 | 33.06 | 32.72 | 34.9 | 39.01 | 45.24 | 37.79 | - |
| Otud7b | 3.73 | 3.54 | 3.43 | 3.24 | 3.62 | 3.49 | 3.21 | 3.8 | 3.54 | 2.52 | 3.2 | 2.22 | - |
| Hes6 | 22.28 | 21.53 | 20.04 | 29.35 | 25.87 | 27.8 | 36.37 | 27.54 | 30.66 | 36.3 | 32.59 | 34.97 | - |
| 2810029C07Rik | 0.32 | 0.06 | 0.11 | 0.71 | 0.35 | 0.2 | 0.32 | 0.66 | 0.55 | 1.14 | 0.35 | 0.83 | - |
| Sgsm3 | 9.48 | 8.89 | 7.23 | 6.33 | 5.73 | 6.06 | 5.22 | 5.98 | 6.05 | 2.96 | 4.01 | 4.31 | - |
| Atf3 | 55.49 | 45.99 | 55.62 | 41.58 | 53.81 | 54.42 | 19.89 | 38.28 | 34.26 | 17.89 | 23.83 | 18.88 | - |
| Ap5m1 | 12.14 | 11.81 | 10.95 | 15.51 | 14.42 | 15.74 | 14.35 | 13.86 | 13.38 | 16.74 | 16.89 | 16.93 | - |
| Ppp1r15a | 9.38 | 9.73 | 10.39 | 10 | 12.39 | 13.23 | 13.5 | 16.26 | 12.67 | 12.98 | 15.22 | 13.79 | - |
| Fam46a | 5.91 | 6.74 | 7.02 | 2.36 | 2.45 | 1.74 | 1.81 | 2.07 | 2.28 | 0.56 | 0.92 | 0.65 | - |
| Siglec5 | 3.31 | 3.4 | 3.16 | 1.32 | 0.74 | 0.42 | 2.86 | 4.29 | 4.14 | 1.05 | 1.06 | 1.67 | - |
| Actr10 | 34.26 | 33.57 | 32.72 | 36.49 | 34.33 | 33.79 | 60.02 | 45 | 50.29 | 55 | 61.07 | 62.02 | - |
| Sap25 | 1.63 | 0.19 | 1.86 | 3.96 | 3.5 | 3.5 | 3.43 | 3.3 | 3.29 | 4.65 | 6.1 | 5.58 | - |
| Npepps | 9.6 | 9.08 | 10.67 | 12.91 | 14.15 | 13.77 | 15.41 | 12.16 | 12.12 | 16.21 | 18.15 | 15.43 | - |
| Tbck | 3.58 | 2.14 | 2.6 | 3.7 | 4.58 | 5.27 | 5.19 | 3.87 | 4.15 | 4.23 | 5.68 | 4.08 | - |
| Hist1h2ab | 9.35 | 8.9 | 8.34 | 5.11 | 6.79 | 3.17 | 3.04 | 4.4 | 4.44 | 2.72 | 3.4 | 1.76 | - |
| Hltf | 7.54 | 7.69 | 7.93 | 9.54 | 11.07 | 9.36 | 9.52 | 9.76 | 10.85 | 9.78 | 10.88 | 9.38 | - |
| Selplg | 180.07 | 191.43 | 180.04 | 327.39 | 274.4 | 291.04 | 365.63 | 258.07 | 293.44 | 349.77 | 280.17 | 373.31 | - |
| Kat6a | 2.11 | 1.81 | 1.56 | 2.79 | 3.03 | 2.59 | 2.52 | 2.01 | 2.2 | 3.28 | 3.32 | 2.68 | - |
| Ugt1a6a | 2.49 | 1.23 | 2.56 | 0.3 | 0.66 | 0.19 | 0.3 | 1.01 | 0.27 | 0.14 | 0.1 | 0.18 | - |
| Gnb2l1 | 1181.55 | 1174.82 | 1216.8 | 1415.46 | 1420.85 | 1461.37 | 1328.38 | 1388.04 | 1345.48 | 1374.37 | 1485.8 | 1350.91 | - |
| Il2 | 20.91 | 11 | 15.96 | 0 | 0.27 | 0.07 | 0 | 0 | 2.11 | 0 | 0 | 0 | - |
| Prkcd | 12.45 | 16.84 | 17.26 | 25.39 | 22.77 | 22.75 | 29.2 | 24.66 | 24.57 | 30.88 | 32.28 | 25.84 | - |
| Mast2 | 3.55 | 3.24 | 3.39 | 2.82 | 3.05 | 2.49 | 2.24 | 2.5 | 2.69 | 2.08 | 1.97 | 2.04 | - |
| Rnf125 | 12.75 | 13.13 | 15.97 | 17.91 | 15.25 | 13.82 | 17.53 | 16.25 | 20.55 | 28.19 | 19.68 | 24.15 | - |
| Ppp1r9b | 1.5 | 2.04 | 2.21 | 3.71 | 3.46 | 2.59 | 2.73 | 1.95 | 2.33 | 4.08 | 3.09 | 3.58 | - |
| Cldn12 | 3.68 | 4.13 | 4.1 | 2.54 | 3.13 | 3.6 | 2.46 | 4.16 | 3.81 | 2.28 | 1.59 | 2.39 | - |
| Slc25a28 | 1.79 | 2.35 | 1.97 | 3.75 | 3.5 | 4.03 | 4.01 | 2.91 | 2.9 | 4.4 | 4.7 | 3.12 | - |
| 6720489N17Rik | 7.6 | 4.47 | 6.3 | 5.59 | 5.03 | 3.89 | 3.38 | 4.26 | 3.42 | 2.45 | 2.52 | 3.94 | - |
| Rgs18 | 0.89 | 0.94 | 0.55 | 0.27 | 0.07 | 0.07 | 0.03 | 0.03 | 0 | 0 | 0.2 | 0 | - |
| Surf6 | 22.7 | 24.04 | 24.9 | 17.57 | 19.91 | 16.09 | 18.19 | 22.35 | 20.87 | 15.01 | 15.41 | 16.54 | - |
| Slc25a46 | 27.2 | 23.39 | 23.99 | 21.72 | 23.39 | 20.72 | 18.25 | 19.86 | 20.79 | 19.57 | 18.98 | 17.32 | - |
| Hist1h3a | 1.54 | 1.78 | 1.08 | 0.17 | 0.36 | 0.56 | 0.7 | 0 | 0.33 | 0.41 | 0.18 | 0 | - |
| Mplkip | 1.77 | 1.76 | 2.2 | 2.81 | 2.97 | 2.86 | 3.58 | 2.92 | 2.95 | 4.59 | 4.35 | 4.22 | - |
| Abca1 | 0.04 | 0.05 | 0.01 | 0.11 | 0.21 | 0.1 | 0.25 | 0.27 | 0.17 | 0.49 | 0.26 | 0.28 | - |
| Il33 | 0.89 | 0.91 | 1.25 | 1.63 | 1.47 | 1.85 | 1.87 | 2.69 | 1.24 | 2.13 | 1.88 | 3.25 | - |
| 9430091E24Rik | 0.45 | 0.4 | 0.28 | 1.05 | 0.45 | 0.56 | 0.68 | 0.51 | 0.42 | 1.61 | 0.55 | 1.36 | - |
| Plec | 6.52 | 5.72 | 6.79 | 6.34 | 6.94 | 6.68 | 6.78 | 6.15 | 6.76 | 8.52 | 6.82 | 8.26 | - |
| Rsph3a | 21.08 | 19.38 | 24.41 | 17.94 | 20.12 | 19.29 | 21.6 | 18.27 | 18.41 | 14.13 | 16.26 | 18.87 | - |
| Tmem175 | 11.66 | 11.3 | 10.84 | 9.51 | 10.96 | 8.32 | 7.44 | 9.48 | 8.31 | 7.46 | 5.94 | 8.6 | - |
| 2410002F23Rik | 31.84 | 36.49 | 33.5 | 22.32 | 26.17 | 28.25 | 16.35 | 23.27 | 26.12 | 18.24 | 14.49 | 18.56 | - |
| Uap1 | 7.23 | 9.97 | 8.46 | 4.97 | 5.93 | 5.56 | 5.88 | 6.25 | 7.54 | 4.71 | 5.75 | 4.01 | - |
| Bcl6 | 0.07 | 0.19 | 0.21 | 0.87 | 0.86 | 1.05 | 2.92 | 1.07 | 0.52 | 5.12 | 4.45 | 5.11 | - |
| Rp9 | 25.57 | 34.68 | 43.41 | 60.05 | 56.18 | 54.89 | 66.23 | 54.85 | 59.95 | 66.37 | 65.6 | 61.45 | - |
| Zmynd11 | 5.47 | 4.95 | 5.84 | 8.86 | 8.85 | 7.83 | 7.85 | 6.66 | 6.95 | 12.52 | 11.03 | 10.61 | - |
| Slc38a9 | 1.31 | 1.1 | 1.03 | 2.94 | 1.99 | 2.36 | 3.18 | 1.92 | 2.74 | 3.49 | 3.47 | 3.44 | - |
| Fam189b | 7.32 | 7.8 | 9.39 | 12.09 | 9.31 | 8.09 | 13.47 | 9.2 | 8.09 | 12.98 | 11.93 | 12.62 | - |
| Tmcc3 | 0.06 | 0 | 0.09 | 0.15 | 0.15 | 0.09 | 0.37 | 0.16 | 0.23 | 0.49 | 0.83 | 0.31 | - |
| Zfp655 | 9.74 | 10.86 | 12.01 | 14.32 | 14.99 | 14.84 | 15.31 | 16.22 | 15.56 | 16.54 | 14.78 | 17.3 | - |
| Bola1 | 32.73 | 34.16 | 33.6 | 29.13 | 28.09 | 27.89 | 27.51 | 29.33 | 30.32 | 22.21 | 22.16 | 21.9 | - |
| Dbf4 | 5.82 | 5.88 | 7.1 | 4.7 | 5.2 | 4.99 | 5.71 | 5.24 | 4.37 | 3.37 | 4.87 | 4.08 | - |
| Traf3ip3 | 32.72 | 35.71 | 34.95 | 41.79 | 36.24 | 37.77 | 49.88 | 35.16 | 37.09 | 52.36 | 46.59 | 55.59 | - |
| Elp2 | 96.29 | 92.19 | 94.69 | 64.88 | 65.39 | 71.7 | 65.94 | 80.45 | 79.38 | 54.81 | 55.36 | 52.77 | - |
| Zfp120 | 7.46 | 7.66 | 7.8 | 17.61 | 15.96 | 16.46 | 13.5 | 11.48 | 12.02 | 21.41 | 22.66 | 21.47 | - |
| Vps16 | 46.04 | 43.58 | 46.47 | 40.35 | 40.31 | 39.74 | 36.3 | 38.95 | 39.12 | 36.7 | 33.97 | 35.6 | - |
| Fam222b | 5.63 | 5.59 | 5.22 | 5.1 | 5.5 | 5.2 | 3.18 | 3.55 | 4.23 | 3.55 | 3.9 | 3.95 | - |
| Mutyh | 9.2 | 7.56 | 8.9 | 9.05 | 7.44 | 10.16 | 5.57 | 8.16 | 7.1 | 5.17 | 6.46 | 6.08 | - |
| Cdk13 | 1.09 | 0.78 | 0.86 | 1.43 | 1.18 | 1.51 | 1.44 | 1.2 | 1.71 | 1.63 | 1.92 | 1.67 | - |
| Piwil2 | 1.35 | 1.49 | 1.63 | 1.29 | 1.75 | 1.52 | 3.12 | 3.23 | 3.33 | 2.34 | 2.56 | 2.67 | - |
| Il1rl2 | 1.43 | 0.72 | 1.33 | 0.62 | 2.08 | 1.34 | 0.44 | 0.87 | 0.63 | 0.59 | 0.25 | 0.3 | - |
| Ifnar1 | 14.56 | 15.05 | 15.34 | 22.1 | 18.61 | 18.73 | 22.21 | 18.7 | 17.08 | 21.68 | 21.48 | 24.78 | - |
| Nelfb | 54.9 | 54.54 | 56.47 | 60.88 | 57.95 | 61.79 | 66.89 | 59.78 | 59.81 | 67.07 | 69.16 | 73.44 | - |
| Gm4013 | 1.1 | 0.98 | 0.91 | 1.98 | 2.7 | 3.53 | 5.39 | 3.25 | 4.26 | 5.36 | 6.37 | 4.71 | - |
| Mfi2 | 0 | 0 | 0 | 0.06 | 0.11 | 0.09 | 0.14 | 0 | 0.03 | 0.06 | 0.19 | 0.03 | - |
| Wash | 2.94 | 4.6 | 4.03 | 6 | 4.5 | 5.52 | 6 | 5.38 | 4.26 | 6.97 | 9.16 | 7.95 | - |
| Sfn | 1.88 | 2.13 | 1.98 | 1.14 | 1.12 | 0.95 | 1.71 | 1.71 | 1.1 | 0.92 | 0.96 | 0.71 | - |
| Cd83 | 8.7 | 7.79 | 9.28 | 9.37 | 13.04 | 14.92 | 1.29 | 2.35 | 2.15 | 2.37 | 2.7 | 1.69 | - |
| Cnih | 113.95 | 117.09 | 107.91 | 78.66 | 81.86 | 81.36 | 75.08 | 91.31 | 88.54 | 67.42 | 66.01 | 65.52 | - |
| Supt3 | 14.36 | 15.89 | 15.46 | 17.73 | 24.16 | 25.33 | 28.21 | 26.41 | 27.13 | 27.65 | 34.45 | 27.87 | - |
| Spin2 | 2.71 | 2.85 | 1.86 | 2.47 | 1.34 | 2.88 | 2.02 | 1.03 | 1.77 | 1.31 | 0.94 | 0.84 | - |
| Terf2ip | 10.06 | 8.11 | 9.64 | 12.84 | 10.39 | 12.54 | 11.71 | 12.92 | 11.44 | 14 | 14.14 | 14.72 | - |
| Bach2 | 2.29 | 1.74 | 2.17 | 2.32 | 2.11 | 2.34 | 3.48 | 3.2 | 3.32 | 3.38 | 2.51 | 4.03 | - |
| Rps25 | 1264.87 | 1292.72 | 1278.4 | 1483.19 | 1552.92 | 1600.1 | 1517.17 | 1531.81 | 1491.81 | 1498.25 | 1707.5 | 1475.22 | - |
| Mdn1 | 4.97 | 5.06 | 5.28 | 3.55 | 3.16 | 3.63 | 2.38 | 3.16 | 3.01 | 2.2 | 2.14 | 2.13 | - |
| Pqlc3 | 39.48 | 37.72 | 37.83 | 51.63 | 46.04 | 47.86 | 57.52 | 50.21 | 52.29 | 52.34 | 47.89 | 55.18 | - |
| Pear1 | 2.79 | 3.2 | 2.58 | 3.87 | 3.4 | 3.65 | 10.13 | 5.4 | 7.17 | 6.53 | 6.67 | 10.03 | - |
| Aif1l | 0.1 | 0.16 | 0.16 | 0.21 | 0.49 | 0.48 | 0.41 | 0.24 | 0.38 | 1.23 | 1.11 | 0.55 | - |
| Chl1 | 0.46 | 0.23 | 0.1 | 0 | 0.04 | 0 | 0.1 | 0.16 | 0.03 | 0.04 | 0 | 0.01 | - |
| Rbm12b2 | 3.98 | 4.2 | 4.42 | 3.41 | 3.97 | 4.42 | 3.46 | 3.49 | 4.16 | 2.41 | 2.45 | 2.75 | - |
| Iqcc | 9.37 | 8.72 | 10.77 | 6.99 | 7.31 | 7.38 | 7.3 | 8.7 | 7.69 | 6.06 | 5.21 | 6.32 | - |
| Pspc1 | 26.73 | 26.16 | 23.45 | 22.01 | 23.6 | 23.55 | 17.49 | 16.96 | 18.77 | 16.98 | 15.34 | 15.29 | - |
| Nmt2 | 27.46 | 28.55 | 27.13 | 26.67 | 29.82 | 28.66 | 21.16 | 29.67 | 25.73 | 21.36 | 21.31 | 17.54 | - |
| Trim27 | 6.04 | 6.62 | 8.28 | 8.73 | 6.89 | 7.59 | 7.66 | 7.7 | 9.87 | 10.74 | 9.82 | 8.82 | - |
| Anapc16 | 22.23 | 22.15 | 22.7 | 32.96 | 32.35 | 33.88 | 24.56 | 31.77 | 25.23 | 34.05 | 38.21 | 35.48 | - |
| Lif | 44.69 | 49.3 | 60.56 | 12.44 | 15.05 | 13.69 | 4.81 | 8.92 | 10.41 | 1.26 | 1.21 | 0.36 | - |
| Gtpbp4 | 126.23 | 125 | 130.51 | 75.42 | 80.88 | 84.51 | 64.69 | 98.99 | 82.73 | 54.26 | 61.51 | 54.3 | - |
| Eri2 | 5.84 | 5.9 | 5.3 | 5.31 | 5.35 | 6.54 | 4.64 | 5.28 | 5.37 | 4.81 | 2.9 | 4.1 | - |
| Rad50 | 10.88 | 9.24 | 10.11 | 8.86 | 11.45 | 10.88 | 7.3 | 8.43 | 8.64 | 6.73 | 8.12 | 7.33 | - |
| Tspyl1 | 26.28 | 27.39 | 27.8 | 34.85 | 30.43 | 30.69 | 32.96 | 30.41 | 30.02 | 37.21 | 34.6 | 38.02 | - |
| Fancb | 4.62 | 5.98 | 3.64 | 2.05 | 2.6 | 2.44 | 1.65 | 2.68 | 2.01 | 1.41 | 1.27 | 0.72 | - |
| Secisbp2l | 1.87 | 2.19 | 2.2 | 2.79 | 2.75 | 2.45 | 2.94 | 2.88 | 2.72 | 2.94 | 2.99 | 2.76 | - |
| Arhgap26 | 2.25 | 2.58 | 2.15 | 6.53 | 5.29 | 6.1 | 5.81 | 3.31 | 4.1 | 9.82 | 10.21 | 10.36 | - |
| Gnpat | 57.16 | 51.48 | 56.95 | 46.64 | 47.09 | 43.28 | 49.09 | 46.88 | 54.47 | 41.98 | 43.99 | 44.18 | - |
| Prkcb | 2.85 | 2.43 | 2.69 | 3.05 | 3.04 | 3.21 | 2.93 | 3.47 | 2.89 | 3.78 | 3.88 | 4.28 | - |
| Yeats4 | 37.01 | 45.03 | 41.61 | 30.35 | 32.9 | 38.45 | 32.65 | 37.17 | 34.18 | 28.03 | 31.78 | 30.58 | - |
| Rbm20 | 0 | 0.01 | 0 | 0.22 | 0.09 | 0.23 | 0.04 | 0.02 | 0.07 | 0.1 | 0 | 0.21 | - |
| BC005624 | 29.32 | 29.48 | 28.29 | 27.51 | 24.68 | 23.87 | 21.47 | 24.58 | 22.5 | 20.85 | 20.51 | 23.13 | - |
| Sirt2 | 42.53 | 43.64 | 39.19 | 48.13 | 46.54 | 47.06 | 53.21 | 43.54 | 46.89 | 57.25 | 55.48 | 62.06 | - |
| Aen | 54.55 | 50.69 | 51.83 | 36.54 | 44.13 | 42.19 | 36.94 | 48.63 | 45.34 | 34.46 | 37.29 | 30.41 | - |
| Atraid | 29.97 | 26.86 | 23.73 | 45.53 | 41.77 | 49.97 | 51.01 | 34.65 | 44.44 | 55.12 | 57.33 | 68.74 | - |
| Rnf181 | 69.27 | 62.06 | 61.26 | 84.18 | 89.91 | 84.23 | 98.47 | 78.97 | 81.04 | 87.11 | 96.52 | 94.86 | - |
| Cdc7 | 18.93 | 17.25 | 16.76 | 13.62 | 16.76 | 14.47 | 13.08 | 13.45 | 13.96 | 11.59 | 12.68 | 9.68 | - |

| | | | | | | | | | | | | |
|---|---|---|---|---|---|---|---|---|---|---|---|---|
| Gemin6 | 40.26 | 39.63 | 41.47 | 15.49 | 20.23 | 17.44 | 10.74 | 17.53 | 15.17 | 7.64 | 7.49 | 9.24 | - |
| Wipf2 | 1.77 | 1.64 | 1.84 | 1.88 | 2.57 | 1.9 | 2.45 | 1.96 | 2.46 | 2.58 | 2.54 | 2.56 | - |
| Dsn1 | 13.25 | 12.99 | 13.21 | 8.89 | 12.66 | 10.75 | 6.62 | 6.76 | 6.27 | 5.21 | 5.94 | 5.02 | - |
| Smoc1 | 0.2 | 0.3 | 0.17 | 0 | 0.05 | 0.02 | 0 | 0.12 | 0.17 | 0.05 | 0 | 0 | - |
| Tada2a | 17.29 | 17.98 | 17.11 | 9.16 | 11.9 | 12.01 | 10.31 | 14.25 | 11.7 | 11.39 | 9.39 | 8.43 | - |
| Ltn1 | 7.15 | 7.07 | 7.7 | 4.7 | 5.74 | 5.17 | 5.04 | 3.95 | 6.01 | 5.04 | 4.38 | 3.95 | - |
| Pigp | 48.23 | 35 | 41.31 | 35.46 | 39.92 | 33.61 | 29.76 | 34.64 | 36.34 | 25.73 | 26.76 | 30.68 | - |
| Lat2 | 9.33 | 12.64 | 6.27 | 6.35 | 7.74 | 6.41 | 38.15 | 25.84 | 33.46 | 24 | 23.46 | 27.7 | - |
| Gys1 | 9.57 | 9.16 | 10.07 | 10.05 | 12.3 | 9.33 | 19.17 | 12.57 | 15.03 | 13.11 | 11.08 | 13.06 | - |
| Dapk1 | 0.09 | 0.1 | 0.03 | 0.1 | 0.11 | 0.16 | 0.96 | 0.32 | 0.6 | 0.5 | 0.49 | 0.22 | - |
| R3hdm1 | 16.87 | 16.35 | 17.11 | 24.18 | 23.39 | 21.94 | 27.18 | 19.37 | 23.79 | 22.13 | 32.39 | 23.95 | - |
| Apbb2 | 0.43 | 0.37 | 0.32 | 0.59 | 0.91 | 0.55 | 1.18 | 0.96 | 1.03 | 2.03 | 1.49 | 1.85 | - |
| Cbwd1 | 14.04 | 12.82 | 12.69 | 5.71 | 6.6 | 7.2 | 6.11 | 8.9 | 7.69 | 3.91 | 4.72 | 4.77 | - |
| Arl5c | 2.16 | 3.06 | 1.59 | 5.39 | 2.92 | 2.29 | 7.78 | 5.49 | 4.4 | 5.79 | 4.86 | 5.72 | - |
| Rab33a | 3.17 | 2.69 | 3.11 | 2.73 | 4.75 | 3.74 | 9.04 | 6.04 | 8.82 | 6.23 | 5.89 | 5.69 | - |
| Zfp740 | 4.91 | 4.46 | 4.28 | 7.77 | 6.07 | 7.51 | 7.6 | 6.26 | 7.04 | 9.45 | 7.79 | 8.16 | - |
| H2-Eb1 | 1.68 | 1.08 | 1.23 | 3.18 | 4.34 | 6.53 | 2.51 | 2.74 | 1.8 | 3.98 | 3.51 | 5.18 | - |
| Fhod1 | 0.41 | 0.66 | 0.56 | 0.78 | 0.73 | 0.81 | 1.04 | 0.78 | 0.82 | 1.32 | 1.52 | 1.06 | - |
| Phf23 | 13.07 | 15.02 | 16.27 | 13.72 | 13.67 | 12.32 | 10.99 | 9.86 | 10.71 | 10.65 | 11.41 | 10.82 | - |
| Pgpep1 | 6.75 | 5.8 | 7.2 | 9.06 | 9.99 | 9.77 | 9.78 | 9.36 | 9.27 | 10.85 | 10.9 | 9.15 | - |
| 1190007I07Rik | 22.31 | 23.33 | 23.03 | 14.42 | 16.02 | 15.36 | 16.89 | 20.38 | 16.31 | 9.69 | 12.01 | 9.97 | - |
| Zfp143 | 16.23 | 16.38 | 15.7 | 20.14 | 20.07 | 20.09 | 20.71 | 20.44 | 19.77 | 23.13 | 24.59 | 25.46 | - |
| Glrx | 84.64 | 114.15 | 69.42 | 134.5 | 113.56 | 116.25 | 282.83 | 128.58 | 169.51 | 247.52 | 218.24 | 279.98 | - |
| H2afx | 41.91 | 33.07 | 34.34 | 29.45 | 39.55 | 32.88 | 29.61 | 25.42 | 27.3 | 21.79 | 21.59 | 22.73 | - |
| 2-Mar | 33.16 | 34.3 | 32.16 | 47.23 | 43.76 | 45.45 | 54.58 | 50.62 | 50.05 | 61.66 | 59.6 | 62.39 | - |
| Per1 | 5.52 | 4.51 | 5.24 | 12.34 | 12.35 | 12.25 | 8.97 | 8.34 | 8.34 | 11.74 | 16.57 | 16.65 | - |
| Spc25 | 44.86 | 32.51 | 31.74 | 20.76 | 31.28 | 28.11 | 14.06 | 17.82 | 20.51 | 14.46 | 16.7 | 11.92 | - |
| Sertad4 | 0 | 0 | 0 | 0.06 | 0.06 | 0.04 | 0 | 0 | 0 | 0.09 | 0.15 | 0.14 | - |
| Trabd | 33.18 | 33.46 | 33.61 | 33.96 | 35.16 | 36.32 | 43.51 | 42.12 | 38.53 | 37.31 | 44.04 | 38.97 | - |
| Kmo | 4.46 | 4.59 | 3.42 | 0.18 | 0.1 | 0.26 | 0.18 | 0.19 | 0.24 | 0 | 0 | 0 | - |
| Siae | 3.09 | 2.97 | 2.87 | 5.17 | 4.78 | 5.7 | 4.2 | 4.17 | 3.75 | 5.63 | 4.92 | 6.79 | - |
| E130317F20Rik | 0.29 | 0.44 | 0.32 | 1.02 | 0.68 | 0.56 | 0.64 | 0.67 | 0.84 | 0.89 | 0.86 | 0.96 | - |
| Col7a1 | 0.04 | 0.07 | 0.1 | 0.12 | 0.18 | 0.2 | 0.16 | 0.1 | 0.09 | 0.21 | 0.33 | 0.31 | - |
| Inpp4b | 8.66 | 8.72 | 6.92 | 8.33 | 5.53 | 6.68 | 5.79 | 6.01 | 7.38 | 4.18 | 3.41 | 4.17 | - |
| Taok3 | 12.97 | 12.8 | 13.17 | 13.5 | 12.98 | 14.75 | 17.29 | 14.44 | 14.67 | 17.31 | 16.72 | 17.4 | - |
| Cd1d1 | 3.78 | 3.78 | 3.51 | 5.97 | 5.37 | 6.33 | 7.85 | 5.57 | 5.37 | 9.93 | 6.66 | 10.42 | - |
| Fbxo44 | 3.86 | 3.79 | 3.62 | 6.14 | 5.07 | 6.27 | 7.17 | 5.42 | 3.93 | 6.51 | 6.78 | 7.62 | - |
| 5730507C01Rik | 2.46 | 4.72 | 5.09 | 6.67 | 6.96 | 5.22 | 6.55 | 5.17 | 5.94 | 9.78 | 8.75 | 7.77 | - |
| Pcsk1 | 0.29 | 0.09 | 0.64 | 0.25 | 0.25 | 0.03 | 0.09 | 0 | 0.17 | 0 | 0 | 0 | - |
| Gm14305 | 1.83 | 0.85 | 0.81 | 1.63 | 2.83 | 3.71 | 2.77 | 1.99 | 2.77 | 3.56 | 4.01 | 5.34 | - |
| Pecam1 | 4.61 | 5.31 | 4.84 | 3.31 | 3.44 | 2.48 | 11.18 | 8.24 | 9.08 | 7.19 | 5.61 | 9.21 | - |
| Slc16a1 | 21.36 | 28.6 | 28.32 | 13.71 | 15.72 | 14.65 | 11.35 | 19.38 | 18.72 | 10.4 | 11.94 | 11.36 | - |
| Ccl27a | 2.88 | 3.12 | 1.24 | 3.61 | 5.15 | 3.26 | 8.21 | 3.8 | 4.28 | 5.88 | 7.05 | 6.28 | - |
| Pcnxl3 | 3.8 | 3.69 | 3.54 | 5.77 | 5.19 | 4.6 | 4.82 | 4.58 | 4.31 | 5.3 | 4.28 | 4.86 | - |
| Alg3 | 33.21 | 39.03 | 37.37 | 18.89 | 23.04 | 22.06 | 20.65 | 25.88 | 24.96 | 15.78 | 14.19 | 14 | - |
| BC028528 | 1.02 | 0.39 | 1.09 | 1.13 | 1.54 | 1.14 | 6.56 | 7.55 | 4.8 | 2.81 | 5.24 | 4.9 | - |
| Kcnq1ot1 | 4.21 | 4.55 | 4.79 | 6.4 | 5.83 | 5.61 | 5.41 | 5.33 | 5.12 | 6.78 | 6.56 | 6.13 | - |
| D130043K22Rik | 0.15 | 0.02 | 0.09 | 0 | 0 | 0.01 | 0.1 | 0.06 | 0.08 | 0 | 0 | 0 | - |
| Fdps | 393.77 | 411.84 | 434.09 | 325.77 | 343.04 | 361.23 | 382.86 | 343.33 | 359.25 | 261.01 | 300.04 | 266.44 | - |
| Rheb | 25.26 | 33.71 | 30.18 | 24.59 | 27.58 | 28.66 | 21.33 | 26.76 | 24.35 | 21.65 | 20.57 | 21.87 | - |
| Abi2 | 6.93 | 5.99 | 7.43 | 5.38 | 5.36 | 6.04 | 5.22 | 5.3 | 4.94 | 4.52 | 5.52 | 4.44 | - |
| Fam120b | 13.15 | 12.17 | 12.05 | 16.02 | 13.49 | 14.23 | 12.82 | 12.88 | 13.27 | 16.33 | 12.91 | 17.72 | - |
| Gldc | 3.88 | 2.97 | 4.09 | 10.52 | 11.95 | 14.09 | 2.68 | 3.48 | 3.36 | 7.97 | 6.48 | 5.2 | - |
| Gphn | 8.5 | 8.31 | 8.54 | 11.22 | 11.05 | 10.22 | 8.85 | 11.36 | 10.54 | 9.97 | 11.84 | 11.53 | - |
| Lrch4 | 22.02 | 18.65 | 20.07 | 29.53 | 25.91 | 27.96 | 33.09 | 28.1 | 28.63 | 32.43 | 32.69 | 34.98 | - |
| Rif1 | 3.79 | 3.92 | 3.5 | 3.11 | 3.1 | 3.04 | 2.93 | 2.63 | 3.22 | 2.07 | 2.6 | 2.2 | - |
| Fam60a | 97.03 | 94.46 | 99.64 | 64.75 | 66 | 70.28 | 70.44 | 89.59 | 86.79 | 52.74 | 60.27 | 57.49 | - |
| Lpxn | 88.34 | 88.5 | 91.86 | 114.9 | 106.9 | 109.69 | 142.23 | 111.91 | 124.46 | 129.04 | 137.64 | 144.5 | - |
| Faf2 | 18.32 | 17.44 | 18.25 | 17.4 | 18.11 | 18.97 | 15.02 | 15.82 | 15.6 | 14.65 | 15.53 | 13.74 | - |
| Lgals6 | 0.78 | 0.34 | 0.3 | 1.69 | 0.77 | 0.76 | 0.77 | 0.46 | 1.23 | 2.37 | 2.25 | 0.83 | - |
| Ubxn4 | 28.82 | 31.45 | 31.51 | 38.25 | 40.5 | 38.75 | 32.36 | 39.6 | 35.56 | 42.14 | 47.27 | 42.01 | - |
| 4933406I18Rik | 0.31 | 0.06 | 0.06 | 0.61 | 0.53 | 0.27 | 0.62 | 0.13 | 0.29 | 1.1 | 1.25 | 1.3 | - |
| Ramp2 | 0.85 | 0.13 | 0.5 | 0.07 | 0 | 0 | 0.4 | 0.34 | 0.06 | 0 | 0 | 0 | - |
| Setd3 | 22.37 | 24.81 | 24.64 | 27.33 | 26.92 | 25.39 | 30.3 | 30.78 | 32.54 | 30.1 | 28.24 | 31.48 | - |
| Impdh2 | 274.23 | 297.97 | 304.8 | 262.44 | 265.41 | 286.7 | 247.71 | 254.92 | 245.05 | 227.77 | 265.45 | 218.66 | - |
| Dis3 | 12.07 | 17.74 | 17.57 | 8.6 | 8.59 | 7.05 | 4.22 | 7.55 | 7.25 | 5.92 | 5.02 | 3.7 | - |
| Hist1h3f | 1.69 | 2.37 | 0.82 | 0.75 | 0.36 | 1.16 | 0.82 | 0.49 | 0.42 | 0.08 | 0.66 | 0.06 | - |
| Egfr | 0.88 | 0.76 | 0.75 | 0.19 | 0.39 | 0.22 | 0.61 | 0.69 | 0.48 | 0.04 | 0.2 | 0.05 | - |
| Ahctf1 | 5.24 | 4.33 | 4.82 | 3.89 | 2.98 | 3.62 | 3.11 | 3.81 | 3.69 | 3.4 | 3.03 | 2.62 | - |
| Prpf38a | 54.08 | 55.81 | 51.72 | 38.82 | 39.72 | 42.4 | 30.83 | 33.55 | 32.16 | 26.4 | 29.42 | 21.55 | - |
| Disp1 | 2 | 1.93 | 1.97 | 6.35 | 4.27 | 5.32 | 3.12 | 3.12 | 3.18 | 9.22 | 8.82 | 8.08 | - |
| 4930509E16Rik | 3.53 | 4.06 | 4.76 | 2.39 | 2.57 | 1.77 | 1.88 | 1.49 | 1.71 | 0.62 | 1.35 | 1.06 | - |
| Hs1bp3 | 3.34 | 2.5 | 4.61 | 6.09 | 5.47 | 5.07 | 5.42 | 5.98 | 6.69 | 5.89 | 5.92 | 7 | - |
| Itpk1 | 18.11 | 18.6 | 14.96 | 28.24 | 29.33 | 27.1 | 22.56 | 18.75 | 20.99 | 37.79 | 35.5 | 38.76 | - |
| Lrrc8d | 7.29 | 6.51 | 7.86 | 6.92 | 7.27 | 6.51 | 3.36 | 4.47 | 4.35 | 3.95 | 2.26 | 2.91 | - |
| Ganc | 1.9 | 2.19 | 2.46 | 3.62 | 3.46 | 2.96 | 2.66 | 2.95 | 2.52 | 3.33 | 2.9 | 4.26 | - |
| Smox | 16.23 | 13.78 | 19.65 | 17.12 | 21.59 | 19.02 | 22.38 | 23.86 | 19.94 | 23.87 | 26.22 | 26.68 | - |
| Pacsin2 | 10.87 | 13.36 | 12.79 | 6.67 | 8.4 | 6.82 | 7.5 | 10.04 | 8.76 | 7.22 | 5.47 | 5.11 | - |
| Ifitm1 | 388.29 | 501.69 | 341.84 | 151.92 | 201.68 | 206.44 | 1212.54 | 1211.55 | 1250.65 | 595.02 | 612.86 | 626.09 | - |
| Eif3c | 263.72 | 272.91 | 264.86 | 240.14 | 270.83 | 259.95 | 232.85 | 289.85 | 268.89 | 201.75 | 229.34 | 206.88 | - |
| Mtcp1 | 6.2 | 4.47 | 5.18 | 7.14 | 7.23 | 6.11 | 7.12 | 6.8 | 6.55 | 9.42 | 8.03 | 8.6 | - |
| Nlrp6 | 0.22 | 0.47 | 0.37 | 0 | 0.1 | 0.02 | 0.02 | 0.25 | 0.08 | 0 | 0.08 | 0 | - |
| Snx15 | 13 | 10.8 | 12.42 | 10.25 | 11.59 | 10.3 | 7.91 | 10.86 | 8.6 | 9.21 | 7.96 | 8 | - |
| Zrsr1 | 0.52 | 0.39 | 0.43 | 0.86 | 0.94 | 0.81 | 0.74 | 0.47 | 0.84 | 0.99 | 1.07 | 0.81 | - |
| Myc | 264.63 | 265.8 | 281.75 | 161.1 | 157.74 | 155.38 | 164.62 | 228.14 | 208.04 | 137.6 | 104.93 | 114.15 | - |
| Btg2 | 6.04 | 6.83 | 6.09 | 8.89 | 7.51 | 7.94 | 13.4 | 7.79 | 9.43 | 19.04 | 11.63 | 19.35 | - |
| 4933439C10Rik | 0.28 | 0.34 | 0.26 | 0.68 | 0.7 | 0.48 | 0.95 | 0.63 | 0.75 | 1.01 | 1.26 | 1.01 | - |
| Mnt | 1.61 | 1.48 | 1.44 | 2.72 | 2.55 | 2.48 | 2.68 | 2.37 | 2.68 | 2.94 | 3.71 | 4.01 | - |
| Scpep1 | 20.44 | 24.89 | 22.37 | 35.2 | 35.37 | 40.65 | 54.3 | 37.02 | 39.73 | 55.75 | 56.55 | 57.98 | - |
| Trip4 | 7.28 | 6.32 | 7.63 | 9.11 | 8.13 | 9.29 | 9.41 | 8.6 | 8.9 | 8.21 | 9.24 | 9.74 | - |
| Med31 | 10.74 | 11.02 | 11.77 | 8.15 | 9.68 | 9 | 8.18 | 10.92 | 9.47 | 7.14 | 8.75 | 7.34 | - |
| Cdkn3 | 19.29 | 18.9 | 17.38 | 15.24 | 20.21 | 19.13 | 10.31 | 9.29 | 8.84 | 7.7 | 10.71 | 8.01 | - |
| Ptpn6 | 121.78 | 120.63 | 120.98 | 101.41 | 98.09 | 108.28 | 108.56 | 102.6 | 97.23 | 80.49 | 83.24 | 90.61 | - |
| Hn1l | 29.55 | 31.28 | 28.25 | 17.87 | 19.52 | 19.8 | 11.69 | 13.41 | 12.6 | 8.03 | 8.92 | 8.01 | - |
| Il13 | 5.35 | 2.51 | 5.25 | 0 | 0 | 0.05 | 0 | 0.24 | 0 | 0 | 0 | 0 | - |
| Gdpgp1 | 4.67 | 3.77 | 4.11 | 3.1 | 3.14 | 2.43 | 2.93 | 2.66 | 2.45 | 2.44 | 2.01 | 3.4 | - |
| Kcnk5 | 12.11 | 12.5 | 10.69 | 7.37 | 7.96 | 6.33 | 8.66 | 9.74 | 10.27 | 4.88 | 4.69 | 7.01 | - |
| Pkd1l3 | 1.67 | 1.38 | 1.81 | 0.71 | 1.28 | 0.99 | 0.84 | 1.11 | 1.19 | 0.66 | 0.47 | 0.77 | - |
| Oraov1 | 9.44 | 7.99 | 8.64 | 12.68 | 12.91 | 13.38 | 13.85 | 11.78 | 12.46 | 13.9 | 16.14 | 16.3 | - |
| 1110008P14Rik | 32.5 | 30.47 | 31.85 | 18.37 | 23.37 | 24.36 | 24.4 | 28.61 | 29.15 | 18.2 | 17.35 | 18.69 | - |
| Pisd | 57.47 | 53.97 | 55.93 | 49.29 | 51.29 | 55.13 | 48.24 | 64.18 | 56.43 | 38.06 | 42.49 | 41.57 | - |
| Tmem194b | 2.92 | 3.65 | 3.48 | 5.26 | 4.82 | 6.32 | 7.69 | 5.79 | 5.8 | 7.77 | 7.55 | 6.14 | - |
| Fancd2 | 5.22 | 4.88 | 4.57 | 2.82 | 3.63 | 2.84 | 1.18 | 2.21 | 1.9 | 0.69 | 1.6 | 1.34 | - |
| Srp54a | 6.67 | 6.54 | 5.26 | 3.99 | 5.39 | 6.01 | 4.23 | 4.77 | 5.2 | 3.94 | 3.61 | 4.18 | - |
| Esco1 | 8.87 | 9.79 | 10.48 | 7.1 | 8.87 | 7.04 | 7.06 | 8.43 | 8.15 | 7.8 | 7.24 | 7.77 | - |
| H2-Q8 | 36.62 | 27.18 | 46.13 | 76.08 | 58.45 | 75.47 | 51.27 | 48.81 | 68.64 | 67.42 | 82.6 | 65.77 | - |
| Vapb | 0.7 | 0.76 | 0.55 | 0.84 | 0.93 | 0.75 | 0.79 | 0.75 | 0.71 | 1.01 | 1.21 | 1.26 | - |
| Slc2a4rg-ps | 5.16 | 5.32 | 5.88 | 6.94 | 5.16 | 5.19 | 12.97 | 7.83 | 9.24 | 11.77 | 9.65 | 12.23 | - |
| Adcy7 | 10.76 | 10.75 | 11.68 | 12.48 | 10.37 | 10.17 | 16.03 | 12.44 | 12.94 | 18.07 | 13.03 | 16.39 | - |
| Matk | 2.43 | 2.08 | 2.2 | 17.2 | 7.13 | 8.05 | 15.81 | 4.7 | 9 | 20.55 | 18.41 | 28.02 | - |
| Procr | 17.76 | 14.28 | 18.91 | 4.2 | 6.18 | 7.04 | 23.8 | 26.56 | 26.43 | 5.78 | 7.66 | 5.29 | - |
| Sema4d | 66.65 | 71.92 | 67.46 | 82.17 | 79.03 | 79.93 | 115.32 | 94.62 | 109.72 | 102.76 | 105.15 | 103.04 | - |
| Psmc3ip | 25.55 | 23.82 | 21.28 | 9.88 | 10.21 | 12.07 | 8.22 | 10.51 | 7.62 | 4.12 | 4.82 | 4.39 | - |
| Atxn1 | 3.66 | 3.17 | 3.44 | 3.15 | 3.69 | 4 | 3.79 | 4.29 | 3.7 | 4.59 | 4.39 | 4.27 | - |
| Arhgdig | 14.99 | 11.45 | 15.61 | 11.15 | 15.73 | 15.39 | 6.68 | 11.33 | 11.33 | 6.76 | 5.26 | 5.19 | - |
| As3mt | 11.02 | 8.27 | 8.89 | 5.2 | 6.49 | 4.08 | 6.4 | 9.59 | 8.64 | 5.55 | 6.67 | 4.64 | - |
| 4930427A07Rik | 10.11 | 7.72 | 8.33 | 5.64 | 5.66 | 5.27 | 3.83 | 3.86 | 3.73 | 3.1 | 2.77 | 2.41 | - |
| Cdc25b | 15.69 | 10.69 | 10.05 | 21.67 | 25.69 | 22.18 | 15.61 | 12.84 | 11.27 | 19.97 | 19.4 | 14.71 | - |
| Zfp592 | 4.43 | 4.56 | 5.22 | 8.39 | 8.5 | 8.54 | 7.89 | 6.76 | 6.89 | 11.11 | 9.42 | 7.97 | - |
| Klra18 | 0.12 | 0 | 0.51 | 0.2 | 0.37 | 0.77 | 0.09 | 0.79 | 0.22 | 0.83 | 0.78 | 1.84 | - |
| Dopey1 | 1.38 | 1.61 | 1.38 | 2.09 | 1.95 | 2.15 | 1.73 | 1.89 | 1.77 | 2.24 | 2.65 | 2.58 | - |
| Crem | 66.25 | 57.13 | 68.64 | 61.96 | 67.8 | 66.11 | 39.02 | 51.68 | 50.91 | 36.69 | 32.31 | 36.46 | - |
| Zfp947 | 2.83 | 2.71 | 2.96 | 1.78 | 2.19 | 2.99 | 1.83 | 1.89 | 1.81 | 1.71 | 1.43 | 1.65 | - |
| Crcp | 34.56 | 29.82 | 31.37 | 34.96 | 30.87 | 35.49 | 41.98 | 41.38 | 40.55 | 40.08 | 45.08 | 38.04 | - |
| Lrp8 | 3.26 | 2.95 | 2.91 | 2.29 | 2.55 | 2.53 | 1.69 | 1.69 | 2.02 | 1.22 | 1.81 | 1.35 | - |
| Cnrip1 | 1.3 | 2 | 2.18 | 0.66 | 0.61 | 0.41 | 0.52 | 0.69 | 0.52 | 0.07 | 0.08 | 0.3 | - |
| Tpp1 | 13.48 | 10.66 | 11.57 | 18.32 | 17.33 | 17.47 | 14.73 | 14.42 | 15.45 | 16.5 | 19.27 | 18.35 | - |
| Pyroxd1 | 7.92 | 7.63 | 6.46 | 4.89 | 6.37 | 5.56 | 5.14 | 7.28 | 6.67 | 4.5 | 4.3 | 5.41 | - |
| Hmha1 | 34.53 | 40.53 | 34.03 | 82.55 | 71.74 | 72.36 | 114.11 | 69.12 | 75.28 | 126.94 | 107.42 | 137.61 | - |
| Mll2 | 11.2 | 11.13 | 11.11 | 17.97 | 16.19 | 17.07 | 16.62 | 14.63 | 15.16 | 20.58 | 20.33 | 21.84 | - |
| Galnt3 | 3.7 | 3.7 | 4.2 | 1.39 | 1.25 | 1.35 | 2.65 | 4.64 | 4.05 | 1.79 | 1.36 | 1.39 | - |
| Herc4 | 13.51 | 13.66 | 11.88 | 16.11 | 14.02 | 15.59 | 14.76 | 13.74 | 15.28 | 19.62 | 18.07 | 18.28 | - |
| Aak1 | 1.73 | 1.88 | 1.44 | 2.9 | 2.36 | 2.45 | 2.89 | 2.03 | 2.28 | 3.02 | 2.52 | 3.32 | - |
| Cnot8 | 52.24 | 50.67 | 51.73 | 47.26 | 45.57 | 42.69 | 45.04 | 46.29 | 50.83 | 40.73 | 42.67 | 43.99 | - |
| 4930470H14Rik | 19.62 | 17.73 | 19.26 | 31.48 | 30.09 | 28.46 | 26.43 | 25.73 | 23.78 | 34.37 | 29.54 | 31.51 | - |
| Tcf25 | 105.08 | 104.87 | 103.32 | 124.96 | 120.66 | 118.35 | 167.91 | 132.44 | 143.54 | 154.32 | 162.65 | 154.48 | - |
| Spag5 | 13.14 | 11.13 | 10.79 | 7.9 | 11.56 | 10.73 | 6.37 | 6.3 | 7.47 | 5.58 | 6.69 | 6.26 | - |
| Tsc22d4 | 18.78 | 21.56 | 20.71 | 29.47 | 24.15 | 23.94 | 32.24 | 25.74 | 26.25 | 33.05 | 33.53 | 37.23 | - |
| Brca2 | 1.31 | 1.34 | 1.54 | 1.32 | 1.24 | 1.02 | 0.81 | 1.03 | 0.81 | 0.62 | 0.64 | 0.71 | - |
| Zfp787 | 0.18 | 0.25 | 0.65 | 0.93 | 1.08 | 0.6 | 0.61 | 1.05 | 0.65 | 1.57 | 1.17 | 0.91 | - |
| Ecsit | 32.97 | 33.25 | 30.67 | 19.99 | 21.52 | 20.32 | 20.82 | 23.82 | 24.51 | 17.41 | 16.17 | 14.23 | - |
| Yars | 66.5 | 70.87 | 71.89 | 54.74 | 60.09 | 61.93 | 53.31 | 75 | 70.69 | 40.24 | 47.32 | 37.32 | - |
| AW112010 | 930.5 | 945.72 | 839.41 | 1104.69 | 1126.75 | 1152.91 | 1572.84 | 1160.98 | 1201.69 | 1651.77 | 1477.39 | 1893.22 | - |
| Ppm1d | 0.89 | 1.47 | 1.01 | 1.4 | 1.14 | 1.28 | 1.69 | 1.5 | 1.41 | 2.12 | 2.6 | 1.57 | - |
| Syt5 | 0.78 | 0.63 | 0.4 | 1.26 | 1.14 | 0.98 | 2.27 | 0.84 | 1.13 | 1.05 | 1.94 | 2.27 | - |
| Tmem181c-ps | 2.55 | 2.51 | 2.66 | 3.22 | 3.32 | 3.35 | 4.05 | 4.05 | 4.08 | 5.1 | 4.1 | 4.36 | - |
| Cdnf | 0.17 | 0.12 | 0.24 | 0.59 | 0.21 | 0.67 | 0.84 | 0.45 | 0.5 | 0.47 | 0.7 | 0.69 | - |
| Lin37 | 20.25 | 17.67 | 17.34 | 26.03 | 25.83 | 27.59 | 29.88 | 24.23 | 26.07 | 31.81 | 30.75 | 30.38 | - |
| Snx27 | 6.91 | 6.41 | 7.21 | 5.44 | 5.48 | 5.82 | 4.73 | 6.2 | 5.84 | 4.99 | 5.89 | 4.42 | - |
| Pacs2 | 1.01 | 0.93 | 1.34 | 2.16 | 2.2 | 2.26 | 2.8 | 1.92 | 2.2 | 4.44 | 3.04 | 4.5 | - |
| Arpc1b | 493.25 | 502.89 | 492.75 | 433.96 | 422.13 | 428.97 | 487.12 | 499.48 | 501.69 | 427.27 | 389.38 | 442.39 | - |

| | | | | | | | | | | | | |
|---|---|---|---|---|---|---|---|---|---|---|---|---|
| Bace1 | 1.4 | 1.28 | 1.36 | 1.88 | 2.24 | 2.57 | 2.3 | 1.87 | 1.73 | 3.44 | 2.36 | 2.39 | - |
| Ypel2 | 0.12 | 0 | 0.02 | 0.71 | 0.43 | 0.52 | 1.14 | 0.58 | 0.54 | 0.63 | 1.3 | 1.18 | - |
| Pnpla7 | 2.61 | 2.5 | 2.24 | 5.05 | 3.92 | 4 | 5.77 | 3.28 | 5.34 | 8.13 | 6.94 | 7.86 | - |
| Prmt3 | 23.71 | 30.6 | 38.37 | 23.94 | 27.22 | 26.53 | 19.81 | 29.3 | 27.21 | 19.42 | 17.83 | 16.6 | - |
| Vps54 | 4.99 | 4.8 | 6.21 | 10.44 | 10.41 | 12.05 | 9.39 | 7.8 | 9.22 | 13.41 | 12.31 | 11.55 | - |
| Kif18a | 10.28 | 7.39 | 8.03 | 8.39 | 11.22 | 9.9 | 5.87 | 5.46 | 4.7 | 6.06 | 6.24 | 4.71 | - |
| Gid4 | 5.23 | 5.89 | 5.86 | 10.16 | 9.65 | 8.44 | 8.99 | 7.96 | 9.99 | 12.85 | 10.64 | 10.83 | - |
| Eif4e3 | 24.76 | 29.92 | 26.07 | 27.01 | 25.54 | 27.31 | 49.9 | 38.26 | 42.14 | 41.51 | 50.08 | 47.64 | - |
| Ptbp3 | 11.7 | 13.38 | 11.86 | 13.67 | 12.45 | 13.33 | 17.39 | 15.68 | 17.73 | 16.63 | 18.57 | 17.87 | - |
| Slc35b2 | 23.74 | 31 | 31.23 | 44.17 | 39.56 | 39.51 | 36.45 | 35.27 | 36.51 | 42.49 | 43 | 42.06 | - |
| Papss1 | 9.81 | 10.52 | 10.28 | 12.25 | 11.06 | 11.55 | 16.73 | 11.97 | 16.38 | 12.92 | 18.8 | 14.72 | - |
| Med18 | 13.96 | 13.44 | 11.7 | 9.11 | 9.41 | 10.98 | 7.68 | 9.44 | 9.85 | 5.62 | 6.35 | 7.69 | - |
| Zfp36l1 | 4.65 | 4.41 | 4.46 | 7.58 | 7.26 | 6.07 | 5.49 | 3.94 | 4.53 | 6.13 | 5.89 | 6.41 | - |
| Slc9a1 | 4.82 | 4.3 | 5.13 | 6.75 | 6.94 | 6.4 | 7.3 | 5.61 | 7.56 | 8.23 | 10.02 | 9.34 | - |
| Fmnl1 | 13.16 | 17.98 | 20.37 | 28.89 | 26.7 | 24.23 | 27.56 | 28.77 | 24.34 | 36.66 | 32.34 | 35.71 | - |
| Gm20605 | 3.47 | 3.27 | 2.59 | 5.26 | 3.91 | 3.19 | 4.08 | 2.68 | 2.72 | 6.32 | 5.77 | 5.35 | - |
| Egr3 | 0.81 | 1.15 | 2.21 | 0.22 | 0.4 | 0.32 | 0.49 | 0.5 | 0.45 | 0.12 | 0 | 0.04 | - |
| Galm | 9.13 | 7.57 | 10.21 | 8.89 | 8.04 | 6.79 | 4.38 | 6.15 | 5.71 | 4.46 | 3.79 | 4.27 | - |
| Rab6b | 0.87 | 0.49 | 0.53 | 1.43 | 1.7 | 1.74 | 1.23 | 1.19 | 1.57 | 2.03 | 2.73 | 2.3 | - |
| Sytl2 | 1.55 | 1.84 | 0.99 | 2.91 | 1.58 | 1.7 | 5.46 | 2.02 | 2.78 | 8.26 | 5.52 | 9.83 | - |
| Fkbp11 | 23.43 | 19.9 | 18.66 | 6.76 | 6.36 | 8.32 | 10.07 | 12.61 | 16.84 | 5.83 | 4.44 | 5.33 | - |
| Mrpl19 | 17.05 | 18.09 | 19.63 | 11.8 | 13.34 | 12.03 | 13.8 | 15.97 | 15.45 | 10.4 | 10.71 | 9.66 | - |
| Phf20 | 4.95 | 5.46 | 5.35 | 7.21 | 6.79 | 6.55 | 6.69 | 6.02 | 6.98 | 6.81 | 8.31 | 7.91 | - |
| Abca2 | 8.19 | 8.45 | 7.94 | 8.86 | 9.15 | 9.01 | 11.17 | 8.89 | 9.03 | 9.9 | 10.28 | 10.15 | - |
| Zfp87 | 7.58 | 7.95 | 9.98 | 11.3 | 11.91 | 10.81 | 9.11 | 9.45 | 9.85 | 11.01 | 13.9 | 12.07 | - |
| Ntan1 | 58.78 | 63.79 | 57 | 61.84 | 61.97 | 60.12 | 72.83 | 65.85 | 72.41 | 72.8 | 75.32 | 82.74 | - |
| Mocs2 | 11.15 | 10.41 | 10.45 | 14.2 | 14.3 | 12.81 | 14.16 | 18.62 | 15.73 | 19.43 | 19.9 | 19.71 | - |
| Cxxc1 | 20.36 | 18.9 | 20.29 | 23.59 | 25.79 | 24.41 | 24.55 | 24.17 | 23.64 | 22.89 | 25.99 | 28.38 | - |
| Lrrc25 | 0 | 0.14 | 0.34 | 0.94 | 0.54 | 1 | 1.4 | 0.69 | 1.18 | 0.67 | 1.16 | 1.31 | - |
| Pi4kb | 2.04 | 2.29 | 2.03 | 2.75 | 2.7 | 3.22 | 3.39 | 2.07 | 1.98 | 3.49 | 2.81 | 3.14 | - |
| Tead2 | 0.22 | 0.25 | 0.2 | 0.36 | 0.47 | 0.3 | 0.57 | 0.5 | 0.46 | 0.78 | 0.67 | 0.93 | - |
| Slc1a2 | 10.85 | 7.49 | 7.85 | 0.3 | 0.74 | 0.43 | 1.2 | 5.61 | 3.04 | 0.02 | 0.18 | 0.36 | - |
| Ppap2a | 5.24 | 4.9 | 4.59 | 2.71 | 4.06 | 4.54 | 2.46 | 1.8 | 2.05 | 0.61 | 1.27 | 2.29 | - |
| Atpif1 | 310.87 | 309.19 | 304.29 | 199.82 | 209.04 | 208 | 199.26 | 252.52 | 228.43 | 133.84 | 162.13 | 143.09 | - |
| Vps13d | 1.02 | 0.81 | 0.9 | 1.46 | 1.38 | 1.39 | 1.25 | 0.99 | 1.35 | 1.58 | 1.44 | 1.5 | - |
| Plekho2 | 12.28 | 11.36 | 13.08 | 15.39 | 14.37 | 12.88 | 18.86 | 16.66 | 14.92 | 20.16 | 17.24 | 20.47 | - |
| Ttyh3 | 0.25 | 0.19 | 0.21 | 0.5 | 0.15 | 0.48 | 0.52 | 0.23 | 0.24 | 1.04 | 0.32 | 0.83 | - |
| Fkbp5 | 10.33 | 10.71 | 9.9 | 36.95 | 32.18 | 36.19 | 11.35 | 8.62 | 10.42 | 36.38 | 35.17 | 37.46 | - |
| Klf4 | 0.04 | 0 | 0.04 | 0.04 | 0.25 | 0.1 | 0.15 | 0.17 | 0.07 | 0.23 | 0.36 | 0.34 | - |
| Il17a | 3.22 | 2.88 | 1.06 | 0 | 0 | 0.05 | 0 | 0 | 0 | 0 | 0 | 0 | - |
| Tbc1d2b | 3.32 | 3.41 | 3.03 | 4.55 | 3.92 | 3.97 | 4.39 | 4.21 | 4.33 | 6.02 | 5.05 | 5.5 | - |
| Pip5k1c | 7.63 | 8.66 | 9.95 | 9.49 | 8.09 | 7.08 | 6.48 | 5.95 | 6.53 | 5.46 | 5.32 | 4.73 | - |
| Suclg2 | 55.34 | 51.48 | 52 | 38.46 | 39.66 | 39.72 | 29.37 | 39.08 | 35.74 | 26.45 | 28.5 | 27.39 | - |
| Ttll3 | 1.2 | 1.44 | 1.35 | 2.51 | 1.96 | 1.49 | 2.06 | 1.75 | 2.61 | 3.03 | 1.98 | 2.75 | - |
| Erbb2ip | 2.15 | 2.85 | 2.67 | 3.35 | 2.96 | 2.93 | 2.8 | 2.22 | 2.85 | 3.45 | 3.4 | 3.54 | - |
| Ppox | 10.28 | 8.8 | 8.79 | 16.59 | 15.98 | 18.7 | 17.99 | 12.61 | 13.08 | 19.47 | 18.22 | 19.86 | - |
| Cyp2d22 | 0.06 | 0.17 | 0.02 | 0.32 | 0.56 | 0.51 | 1.77 | 1.46 | 1.55 | 1.47 | 2.72 | 2.21 | - |
| Mid1ip1 | 46.66 | 45.81 | 43.86 | 33.44 | 33.82 | 37.18 | 36.52 | 42.55 | 42.19 | 27.12 | 29.91 | 30.85 | - |
| Usp47 | 10.57 | 10.29 | 11.44 | 13.61 | 11.82 | 11.36 | 13.01 | 12.83 | 13.49 | 17.42 | 15.62 | 14.49 | - |
| Prokr1 | 0.2 | 0.09 | 0.11 | 0.06 | 0.02 | 0.06 | 0.01 | 0.09 | 0.12 | 0 | 0 | 0.01 | - |
| Gm16907 | 2.55 | 2.86 | 2.86 | 5.25 | 4.91 | 4.8 | 4.55 | 4.43 | 3.44 | 5.93 | 5.54 | 6.89 | - |
| Tmem51 | 0.91 | 0.56 | 1.41 | 5.09 | 2.5 | 3.13 | 5.95 | 2.57 | 3.95 | 5.49 | 4.18 | 7.6 | - |
| Pkmyt1 | 20.87 | 23.87 | 23.48 | 13.51 | 19.57 | 14.85 | 10.89 | 12.21 | 10.53 | 10.51 | 12.06 | 8.28 | - |
| Smpd2 | 6.2 | 6.54 | 5.02 | 8.52 | 7.75 | 7.51 | 7.71 | 7.44 | 6.6 | 10.53 | 10.01 | 11.26 | - |
| 2210417K05Rik | 1.33 | 1.04 | 0.78 | 1.93 | 3.47 | 1.37 | 3.4 | 2.32 | 2.89 | 3.29 | 3.43 | 3.1 | - |
| Xlr4a | 32.23 | 30.28 | 28.08 | 19.1 | 19.94 | 24.54 | 21.16 | 24.51 | 25.51 | 16.79 | 17.6 | 16.73 | - |
| Sult5a1 | 0 | 0 | 0 | 0.05 | 0.22 | 0 | 0.05 | 0 | 0.23 | 0.59 | 0.43 | 0.15 | - |
| Il11ra1 | 1.07 | 1.06 | 0.98 | 2.05 | 1.42 | 1.19 | 2.66 | 2.08 | 2.25 | 2.66 | 3.94 | 3.14 | - |
| Mars | 92.4 | 92.64 | 95.06 | 71.57 | 88.77 | 83.75 | 73.61 | 95.71 | 91.83 | 56.37 | 70.66 | 58 | - |
| Stk17b | 46.14 | 50.05 | 44.17 | 87.03 | 80.13 | 79.73 | 66.87 | 51.22 | 58.74 | 94.24 | 74.4 | 90.1 | - |
| Kif15 | 11.89 | 10.8 | 11.35 | 6.28 | 7.95 | 6.38 | 6.44 | 5.99 | 7.56 | 4.05 | 4.97 | 3.47 | - |
| Usp22 | 15.78 | 16.61 | 15.77 | 20.66 | 21.28 | 22.28 | 22.08 | 19.83 | 22.43 | 24.38 | 23.57 | 22.2 | - |
| Txnl4b | 14.02 | 13.17 | 13.79 | 14.6 | 13.45 | 11.08 | 11.58 | 12.07 | 11.95 | 9.97 | 8.78 | 10.78 | - |
| Hspa9 | 347.62 | 350.21 | 356.73 | 233.34 | 276.86 | 262.17 | 215.78 | 319.31 | 288.45 | 152.66 | 183.48 | 162.8 | - |
| Kdm3a | 10.73 | 11.36 | 10.36 | 10.33 | 9.42 | 9.75 | 20.39 | 15.74 | 16.93 | 13.32 | 16.3 | 14.51 | - |
| Ipo4 | 47.37 | 41.83 | 45.68 | 21.48 | 26.16 | 26.37 | 18.16 | 26.43 | 26.38 | 15.68 | 15.55 | 13.74 | - |
| Scn9a | 0.12 | 0.03 | 0.03 | 0.01 | 0.13 | 0.11 | 0.01 | 0.02 | 0.01 | 0 | 0.01 | 0 | - |
| Tmem242 | 18.78 | 19.54 | 21.27 | 24.73 | 23.79 | 24.04 | 26.92 | 23.77 | 25.36 | 26.74 | 29.63 | 24.81 | - |
| Eefsec | 18.62 | 17.42 | 17.46 | 12.67 | 12.91 | 15.4 | 10.62 | 15.39 | 10.84 | 8.97 | 10.9 | 10.19 | - |
| Junb | 336.31 | 321.4 | 382.94 | 254.84 | 316.17 | 275.18 | 218.72 | 275.43 | 267.06 | 186.85 | 193.21 | 216.37 | - |
| Afap1l1 | 0.15 | 0.03 | 0.02 | 0.19 | 0.06 | 0.07 | 0.78 | 0.03 | 0.19 | 1.18 | 0.49 | 1.69 | - |
| Sp110 | 47.47 | 43.9 | 41.1 | 37.52 | 40.17 | 37.95 | 27.05 | 33.37 | 31.37 | 27.65 | 24.36 | 30.4 | - |
| Plxnc1 | 3.21 | 2.97 | 3.84 | 2.14 | 2.85 | 2.58 | 2.25 | 2.74 | 2.83 | 2.08 | 2.39 | 2.68 | - |
| D630041G03Rik | 0 | 0 | 0 | 0.16 | 0.12 | 0.02 | 0.05 | 0.03 | 0.12 | 0.23 | 0.08 | 0.19 | - |
| Xdh | 5.17 | 5.1 | 4.59 | 4.15 | 3.5 | 3.72 | 6.25 | 4.26 | 4.09 | 3.49 | 2.46 | 3.52 | - |
| Zfp82 | 6.73 | 5.5 | 6.48 | 9.56 | 10.05 | 8.1 | 7.46 | 8.8 | 8.99 | 9.97 | 9.38 | 9.6 | - |
| Crebbp | 3.32 | 2.98 | 3.26 | 4.33 | 4.51 | 4.79 | 4.59 | 4.35 | 3.79 | 4.76 | 4.91 | 4.93 | - |
| Echdc1 | 12.97 | 10.98 | 13.57 | 10.66 | 12.72 | 12.35 | 10.78 | 10.09 | 10.78 | 8.8 | 7.28 | 8.57 | - |
| Adipor2 | 4.94 | 5.55 | 5.13 | 6.87 | 7.46 | 6.95 | 7.87 | 6.7 | 6.48 | 8.36 | 7.29 | 8.46 | - |
| Rps6ka3 | 2.44 | 2.57 | 2.7 | 5.03 | 4.78 | 5.36 | 3.49 | 4.99 | 3.62 | 5.94 | 4.63 | 5.24 | - |
| A430093F15Rik | 6.68 | 5.53 | 5.7 | 7.29 | 7.08 | 7.24 | 10.18 | 9.59 | 8.55 | 9.98 | 9.21 | 10.68 | - |
| Alyref | 4.39 | 4.83 | 5.11 | 2.12 | 2.54 | 2.7 | 1.73 | 2.91 | 1.85 | 1.36 | 1.2 | 1.02 | - |
| Acat2 | 102.52 | 104.19 | 110.48 | 109.54 | 109.56 | 108.26 | 86.34 | 75.33 | 88.23 | 65.54 | 80.5 | 68.48 | - |
| Plk2 | 38.2 | 33.99 | 46.63 | 22.72 | 33.94 | 29.19 | 14.89 | 28.05 | 29.42 | 12.58 | 15.34 | 9.8 | - |
| Scfd1 | 46.16 | 46.58 | 44.58 | 33.28 | 34.33 | 36.77 | 24.84 | 35.75 | 34.49 | 27.33 | 28.03 | 29.14 | - |
| Gimap4 | 146.47 | 156.51 | 136.53 | 280.78 | 221.57 | 230.66 | 248.56 | 197.52 | 211.26 | 339.59 | 276.11 | 353.79 | - |
| Leprot | 24.07 | 25.76 | 28.14 | 33.31 | 34.58 | 31.28 | 29.67 | 32.25 | 29.99 | 33.94 | 32.15 | 31.55 | - |
| Rrbp1 | 8.06 | 10.43 | 9.2 | 9.58 | 9.42 | 9.93 | 13.08 | 11.61 | 13.23 | 14.79 | 13.86 | 14.99 | - |
| Mctp2 | 3.17 | 3.67 | 3.7 | 4.26 | 4.6 | 4.14 | 9.67 | 8.4 | 8.33 | 7.94 | 8.69 | 8.63 | - |
| Zscan26 | 11.68 | 11.15 | 10.41 | 16.02 | 14.18 | 14.6 | 12 | 11.3 | 11.11 | 13.59 | 13.55 | 14.89 | - |
| Golph3l | 8.3 | 10.71 | 8.91 | 15.42 | 12.29 | 15 | 11.87 | 11.2 | 10.54 | 14.21 | 14.58 | 13.78 | - |
| Smad1 | 0.54 | 0.4 | 0.6 | 0.98 | 0.64 | 1.17 | 0.62 | 0.81 | 0.9 | 1.28 | 1.03 | 1.68 | - |
| Rap1gds1 | 20.45 | 20.35 | 21.68 | 16.99 | 15.81 | 13.84 | 18.98 | 18.85 | 19.67 | 15.22 | 15.93 | 12.04 | - |
| Gzmd | 14.84 | 21.63 | 8.83 | 0 | 0 | 0 | 0.71 | 0 | 0.23 | 0 | 0.86 | 0 | - |
| Dgka | 58.43 | 55.5 | 58.95 | 108.17 | 92.57 | 99.52 | 132.06 | 94.28 | 101.32 | 146.94 | 116.13 | 138.11 | - |
| Sp2 | 3.73 | 3.93 | 3.53 | 3.04 | 2.88 | 3.26 | 2.27 | 2.9 | 2.26 | 2.15 | 2.34 | 2.46 | - |
| Rbm47 | 0.38 | 0.37 | 0.23 | 0.49 | 0.4 | 0.26 | 1.4 | 0.53 | 0.62 | 1.82 | 1.26 | 2.02 | - |
| AA414768 | 1.23 | 1.36 | 1.52 | 0.45 | 0.49 | 0.88 | 1.21 | 0.77 | 0.77 | 0.38 | 0.71 | 0.28 | - |
| Arhgef18 | 0.48 | 0.4 | 0.31 | 1.25 | 0.69 | 0.93 | 1.27 | 0.75 | 0.85 | 2.44 | 1.26 | 2.41 | - |
| Rapgef4 | 0.49 | 0.57 | 0.95 | 1.3 | 1.07 | 1.33 | 1.82 | 1.33 | 1.13 | 1.23 | 1.41 | 2.2 | - |
| Skil | 7.7 | 7.68 | 8.16 | 7.18 | 8.2 | 7.58 | 4.4 | 6.16 | 6.07 | 6.07 | 5.78 | 5.28 | - |
| Cbx4 | 2.03 | 2.38 | 2.14 | 3.62 | 3.75 | 3.29 | 2.8 | 2.74 | 2.67 | 3.8 | 3.87 | 4.3 | - |
| Syt11 | 4.18 | 5.18 | 4.24 | 6.74 | 6.02 | 6.43 | 6.56 | 5.72 | 4.81 | 7.45 | 7.33 | 8.31 | - |
| Cltc | 32.8 | 30.28 | 36.8 | 23.18 | 26.02 | 25.97 | 21.08 | 24.83 | 25.82 | 22.52 | 23.77 | 18.64 | - |
| Tcam1 | 0.99 | 0.74 | 0.43 | 0.44 | 0.46 | 0.15 | 0.21 | 0.2 | 0.58 | 0.21 | 0 | 0.02 | - |
| Sec22c | 0.71 | 0.79 | 0.67 | 1.57 | 1.49 | 0.81 | 1.51 | 1.33 | 1.29 | 1.85 | 1.44 | 1.49 | - |
| Ubxn2a | 8.93 | 9.53 | 11.3 | 5.52 | 6.27 | 5.79 | 4.18 | 7.84 | 7.93 | 5.97 | 5.23 | 5.15 | - |
| Zfp365 | 2.11 | 2.2 | 2.04 | 1.65 | 1.48 | 1.12 | 1.74 | 0.66 | 1.01 | 1.14 | 0.91 | 1.19 | - |
| Acp2 | 5.11 | 3.89 | 4.65 | 3.61 | 3.5 | 3.01 | 2.08 | 3.62 | 3.04 | 2.7 | 1.82 | 2.87 | - |
| Lyrm4 | 48.18 | 46.13 | 50.36 | 31.12 | 33.33 | 30.22 | 26.25 | 38.82 | 32.33 | 22.88 | 22.78 | 20.77 | - |
| Jmjd7 | 9.7 | 10.35 | 8.13 | 16.13 | 13.95 | 14.62 | 16.58 | 15.8 | 17.77 | 15.64 | 23.71 | 14.92 | - |
| Dars2 | 11.89 | 11.54 | 13.11 | 8.94 | 8.89 | 8.85 | 7.01 | 6.58 | 5.77 | 5.7 | 4.59 | 4.98 | - |
| Clgn | 1.08 | 0.66 | 0.52 | 0.39 | 0.03 | 0.08 | 0.33 | 0.23 | 0.28 | 0.2 | 0.11 | 0.2 | - |
| Pfdn1 | 111.61 | 115.69 | 118.55 | 80.94 | 83.76 | 89.88 | 96.2 | 100.78 | 100.83 | 67.93 | 80.86 | 65.83 | - |
| Pitpnc1 | 6.39 | 6.84 | 6.21 | 14.13 | 12.9 | 12.58 | 19.27 | 14.98 | 16.22 | 25.59 | 23.67 | 28.26 | - |
| Kif9 | 1.41 | 0.85 | 0.6 | 2.24 | 1.77 | 1.74 | 1.63 | 1.74 | 1.55 | 2.17 | 1.78 | 2.29 | - |
| Pitpnm2 | 1.48 | 1.91 | 1.89 | 1.4 | 1.41 | 1.6 | 1.51 | 1.8 | 1.51 | 1.03 | 1.03 | 1.38 | - |
| Pnp2 | 6.68 | 5.54 | 4.87 | 1.27 | 2.51 | 2.38 | 2.27 | 2.93 | 2.08 | 0.77 | 0.74 | 0.45 | - |
| Dhx57 | 3.94 | 3.67 | 3.66 | 5.45 | 5 | 4.62 | 6.93 | 4.66 | 4.45 | 6.13 | 6.79 | 5.85 | - |
| Penk | 156.4 | 138.83 | 149.68 | 110.6 | 209.72 | 230.76 | 17.62 | 26.7 | 19.3 | 9.4 | 24.97 | 29.72 | - |
| Acadm | 25.74 | 26.3 | 25.83 | 32.24 | 31.34 | 30.77 | 33.75 | 31.44 | 28.62 | 33.01 | 33.21 | 34.94 | - |
| Gcc2 | 1.52 | 1.83 | 1.57 | 1.51 | 1.83 | 2.16 | 1.82 | 1.89 | 1.8 | 2.42 | 2.31 | 2.71 | - |
| Camkv | 0.32 | 0.44 | 0.2 | 0.02 | 0.08 | 0.23 | 0.04 | 0.08 | 0.23 | 0.08 | 0 | 0 | - |
| Lsr | 3.94 | 3.34 | 2.63 | 11.16 | 7.01 | 8.69 | 6.54 | 2.86 | 3.78 | 13.41 | 7.97 | 11.3 | - |
| Ccnt2 | 9.63 | 9.61 | 9.06 | 13.35 | 12.59 | 11.42 | 11.43 | 14.2 | 10.63 | 13.5 | 15.09 | 13.52 | - |
| Mms19 | 19.46 | 16.57 | 17.46 | 23.99 | 22.11 | 24.22 | 33.74 | 23.7 | 27.67 | 36.03 | 40.29 | 33.41 | - |
| Mettl5 | 34.12 | 27.75 | 27.42 | 25.49 | 25.91 | 23.49 | 17.45 | 26.74 | 20.23 | 16.6 | 20.12 | 17.21 | - |
| Pcbp2 | 48.05 | 51.31 | 50.08 | 58.34 | 56.92 | 60.61 | 57.24 | 52.74 | 54.43 | 57.6 | 61.64 | 56.55 | - |
| Ighmbp2 | 5.03 | 4.88 | 5.33 | 3.55 | 4.28 | 3.75 | 3.62 | 4.12 | 3.72 | 3.07 | 3.03 | 3.07 | - |
| Pdcd1 | 67.29 | 60 | 62.19 | 140.55 | 108.51 | 123.25 | 30.87 | 40.93 | 37 | 100.42 | 71.33 | 97.88 | - |
| Zscan29 | 1.39 | 1.41 | 1.54 | 2.48 | 2.26 | 2.22 | 2 | 1.59 | 1.83 | 2.66 | 1.89 | 1.87 | - |
| Gm6313 | 1.21 | 1.19 | 0.98 | 1.91 | 1.39 | 2.11 | 1.77 | 1.38 | 1.9 | 2.29 | 2.31 | 2.27 | - |
| Tmx4 | 1.98 | 2.17 | 1.95 | 3.62 | 1.92 | 2.13 | 3.66 | 2.08 | 3.39 | 3.81 | 5 | 3.85 | - |
| Dscr3 | 24.54 | 25.3 | 23.62 | 30.72 | 27.77 | 29.88 | 37.78 | 30.7 | 31.23 | 31.99 | 32.21 | 37.56 | - |
| Podnl1 | 3.9 | 4.14 | 4.19 | 4.07 | 5.64 | 4.44 | 9.34 | 7.09 | 8.22 | 12.73 | 10.34 | 15.58 | - |
| Cnppd1 | 12.45 | 13.63 | 12.95 | 23.85 | 21.78 | 20.64 | 24.72 | 20.98 | 21.28 | 29.25 | 27.93 | 30.97 | - |
| Ydjc | 44.37 | 43.77 | 45.92 | 24.2 | 28.56 | 26.62 | 23.95 | 33.77 | 29.3 | 19.03 | 17.36 | 18.08 | - |
| Cirbp | 8.73 | 10.28 | 9.86 | 19.17 | 16.4 | 14.37 | 14.37 | 9.96 | 11.06 | 19.74 | 12.51 | 17.99 | - |
| Rad23a | 48.19 | 53.12 | 52.32 | 39.75 | 37.71 | 37.72 | 34.23 | 40.48 | 43.96 | 28.08 | 28.72 | 29.83 | - |
| Capza2 | 387.75 | 381.22 | 397.28 | 330.54 | 346.1 | 354.98 | 288.15 | 350.19 | 338 | 287.14 | 290.75 | 300.92 | - |
| Lysmd1 | 3.95 | 3.73 | 3.02 | 4.42 | 3.93 | 4.7 | 6.22 | 5.68 | 4.54 | 4.53 | 5.67 | 6.6 | - |
| Zfp9 | 2.57 | 2.39 | 2.66 | 2.79 | 2.54 | 2.93 | 3.02 | 5.23 | 3.83 | 2.97 | 4.9 | 3.55 | - |
| Ubac2 | 46.47 | 48.05 | 49.66 | 89.42 | 69.14 | 81.67 | 93.98 | 78.07 | 77.14 | 133.59 | 137.9 | 139.37 | - |
| BC016579 | 0.49 | 0.42 | 0.37 | 0 | 0 | 0.03 | 0.34 | 0.03 | 0.91 | 0 | 0.03 | 0 | - |
| Lacc1 | 0.5 | 0.43 | 0.8 | 0.21 | 0.05 | 0.26 | 0.31 | 0.32 | 0.07 | 0.17 | 0.18 | 0.12 | - |
| Accs | 1.66 | 2.01 | 1.65 | 3.67 | 3.19 | 3.42 | 4.46 | 2.7 | 3.98 | 5.24 | 4.72 | 5.31 | - |
| Taf1d | 45.61 | 46.38 | 49.23 | 83.72 | 79.13 | 84.04 | 64.77 | 64.01 | 67.13 | 99.32 | 114 | 99.01 | - |
| Litaf | 16.8 | 18.68 | 14.49 | 19.24 | 14.79 | 15 | 35.62 | 25.23 | 27.51 | 25.35 | 21.57 | 22.47 | - |
| Abcd2 | 0.08 | 0.06 | 0.03 | 0.11 | 0.16 | 0.1 | 0.08 | 0.03 | 0.06 | 0.36 | 0.3 | 0.19 | - |
| C230081A13Rik | 4.13 | 3.55 | 3.31 | 5.52 | 4.78 | 5.36 | 5.27 | 4.64 | 5.38 | 5.13 | 5.85 | 5.92 | - |
| Mysm1 | 11.1 | 9.86 | 9.84 | 12.54 | 11.87 | 12.48 | 12.36 | 12.84 | 12.07 | 15.72 | 15.76 | 14.44 | - |

| | | | | | | | | | | | | |
|---|---|---|---|---|---|---|---|---|---|---|---|---|
| Bmp2 | 0 | 0 | 0 | 0 | 0.02 | 0.08 | 0 | 0.26 | 0.21 | 0.18 | 0.27 | 0.09 | - |
| Lingo3 | 0.28 | 0.19 | 0.46 | 0.22 | 0.24 | 0.09 | 0.02 | 0.24 | 0.03 | 0.05 | 0.02 | 0.08 | - |
| Lad1 | 4.37 | 3.68 | 4.99 | 0.59 | 0.31 | 0.6 | 0.12 | 0.31 | 0.02 | 0 | 0.04 | 0.12 | - |
| Tbrg1 | 133.06 | 122.74 | 128.24 | 130.45 | 141.5 | 145.78 | 160.29 | 170.3 | 162.73 | 154.99 | 166.56 | 160.16 | - |
| Ntf5 | 0.06 | 0.3 | 0.44 | 1.34 | 0.54 | 0.56 | 1.41 | 1.14 | 0.66 | 1.15 | 1.41 | 1.15 | - |
| Gca | 2.19 | 2.37 | 1.64 | 1.15 | 1.39 | 1.63 | 1.01 | 1.16 | 1.45 | 0.63 | 1.05 | 0.7 | - |
| Orc4 | 31.05 | 29.28 | 30.91 | 24.34 | 24.8 | 24.61 | 21.33 | 25.88 | 25.36 | 21.13 | 20.79 | 17.79 | - |
| Gm17296 | 0.83 | 0.9 | 1.23 | 0.55 | 0.44 | 0.62 | 0.27 | 0.39 | 0.54 | 0.27 | 0.3 | 0.19 | - |
| Fmo5 | 0.22 | 0.19 | 0.25 | 0.35 | 0.69 | 0.49 | 1.5 | 0.77 | 0.65 | 1.86 | 1.1 | 1.65 | - |
| Prss55 | 0 | 0 | 0 | 0 | 0 | 0 | 0.45 | 0 | 0.63 | 0.39 | 0.3 | 0.51 | - |
| Cct4 | 419.56 | 401.05 | 421.87 | 347.46 | 350.96 | 365.99 | 326.67 | 389.2 | 361.86 | 297.24 | 331.56 | 287.82 | - |
| Smg6 | 2.86 | 2.35 | 2.7 | 3.2 | 3.38 | 3.56 | 4.17 | 4.11 | 4.68 | 5.18 | 6.08 | 5.1 | - |
| Bub1 | 19.9 | 16.55 | 15.75 | 11.38 | 15.36 | 11.57 | 7.08 | 7.86 | 6.65 | 7.22 | 6.8 | 5.64 | - |
| Ndufv1 | 97.59 | 100.61 | 98.54 | 70.36 | 80.24 | 71.73 | 65.94 | 76.18 | 71.95 | 52.77 | 54.42 | 55.91 | - |
| Ccng1 | 182.18 | 158.65 | 172.42 | 126.12 | 141.78 | 140.14 | 139.5 | 155.99 | 153.25 | 121.01 | 149.11 | 119.58 | - |
| Plag1 | 0.37 | 0.29 | 0.4 | 0.97 | 0.71 | 0.76 | 1.09 | 0.8 | 1.18 | 1.88 | 1.26 | 1.26 | - |
| Rac1 | 20.44 | 24.85 | 27.89 | 29.8 | 37.9 | 28.7 | 36.72 | 50.53 | 55.98 | 57.71 | 49.52 | 32.7 | - |
| Ints2 | 4.74 | 4.23 | 4.96 | 3.05 | 3.45 | 2.37 | 2.87 | 3.39 | 3.15 | 2.03 | 2.41 | 2.26 | - |
| Ccl1 | 34.93 | 25.46 | 39.82 | 0 | 1.73 | 0.13 | 0 | 0.12 | 0 | 0 | 0 | 0 | - |
| Atf7 | 3.96 | 3.56 | 4.17 | 6.17 | 5.38 | 6.65 | 6.68 | 4.97 | 5.98 | 7.96 | 8.54 | 9 | - |
| Aff4 | 1.94 | 1.65 | 1.56 | 2.75 | 2.05 | 2.35 | 1.96 | 1.88 | 1.93 | 2.56 | 2.59 | 2.32 | - |
| 4930429F24Rik | 0.04 | 0.08 | 0.21 | 0.08 | 0 | 0 | 0.87 | 0.39 | 0.14 | 0.68 | 0.77 | 1.02 | - |
| Ccdc88c | 2.26 | 2.33 | 2.83 | 4.88 | 4.44 | 4.75 | 4.25 | 3.51 | 4.32 | 5.78 | 6.58 | 6.15 | - |
| Got1l1 | 0.21 | 0.13 | 0.24 | 0.37 | 0.83 | 0.8 | 1.44 | 1.21 | 0.47 | 1.31 | 1.68 | 0.85 | - |
| Mlf1ip | 3.92 | 4.6 | 4.25 | 1.7 | 2.37 | 2.19 | 2.56 | 1.75 | 2.09 | 1.83 | 1.58 | 1.07 | - |
| Zwint | 24.7 | 23.75 | 25.71 | 20.22 | 21.51 | 22.62 | 19.73 | 21.29 | 20.41 | 14.39 | 17.43 | 14.6 | - |
| Ift52 | 23.78 | 24.93 | 23.37 | 29.82 | 28.54 | 27.77 | 34.04 | 30.64 | 29.4 | 35.09 | 34.27 | 35.85 | - |
| Ccdc25 | 29.05 | 29.03 | 27.66 | 19.94 | 19.62 | 22.26 | 20.36 | 21.38 | 24.06 | 15.84 | 17.91 | 19.27 | - |
| Cnp | 38.16 | 44.63 | 41.91 | 63.2 | 50 | 55.07 | 61.87 | 51.64 | 56.05 | 55.66 | 49.62 | 64.68 | - |
| Pex7 | 15.3 | 15.91 | 19.44 | 13.64 | 14.56 | 13.22 | 17.53 | 16.83 | 14.13 | 9.26 | 12.51 | 10.76 | - |
| Ercc1 | 24.68 | 26.67 | 28.53 | 20.64 | 21.17 | 23.59 | 21.87 | 19.58 | 20.98 | 20.05 | 17.97 | 20.9 | - |
| Slc26a11 | 1.89 | 1.1 | 1.31 | 2.49 | 1.86 | 2.3 | 4.01 | 1.94 | 3.35 | 4.46 | 4.3 | 5.87 | - |
| Numa1 | 8.03 | 9 | 8.34 | 12.34 | 11.16 | 11.02 | 13.79 | 10.15 | 10.31 | 12.55 | 13.48 | 13.26 | - |
| Sdcbp | 68.26 | 78.13 | 68.12 | 72.64 | 68.16 | 70.8 | 95.53 | 77.12 | 87.16 | 90.06 | 84.63 | 102.93 | - |
| Wdr90 | 3.85 | 3.16 | 3.15 | 2.16 | 2.4 | 2.54 | 1.55 | 1.78 | 1.85 | 1.28 | 1.14 | 1.33 | - |
| Slc43a1 | 1.23 | 1.42 | 1.62 | 0.67 | 1.32 | 1.3 | 0.61 | 1.17 | 0.79 | 0.19 | 0.73 | 0.42 | - |
| Arhgap4 | 13.47 | 14.32 | 10.43 | 17.7 | 14.16 | 16.08 | 18.29 | 13.65 | 14.71 | 20.55 | 14.62 | 22.44 | - |
| Dexi | 0.29 | 0.42 | 0.78 | 1.27 | 1.03 | 1.24 | 1.57 | 0.59 | 0.81 | 1.32 | 1.86 | 1.87 | - |
| Ppp1r13l | 0.24 | 0.48 | 0.6 | 0.99 | 0.78 | 1 | 1.67 | 0.9 | 0.97 | 1.24 | 2.99 | 1.38 | - |
| Maml1 | 0.78 | 0.64 | 0.65 | 1.3 | 0.99 | 1.13 | 1.15 | 1.43 | 1.23 | 1.75 | 1.74 | 1.33 | - |
| Fam129a | 12.42 | 9.71 | 10.5 | 6.13 | 7.38 | 7.66 | 12.5 | 18.34 | 14.64 | 6.93 | 6.4 | 7.38 | - |
| Myh7b | 0.01 | 0.04 | 0.05 | 0.05 | 0.07 | 0.09 | 0.32 | 0.16 | 0.15 | 0.23 | 0.05 | 0.36 | - |
| Trpa1 | 0.39 | 0.44 | 0.41 | 0.04 | 0.12 | 0.06 | 0.01 | 0.01 | 0.25 | 0.05 | 0.03 | 0.08 | - |
| Ldlrap1 | 3.78 | 3.88 | 3.54 | 3.62 | 4.46 | 4.35 | 4.97 | 4.05 | 5.28 | 5.44 | 5.98 | 5.42 | - |
| Colec12 | 1.51 | 1.85 | 1.74 | 2.54 | 2.06 | 1.68 | 2.25 | 1.93 | 1.67 | 2.8 | 3.11 | 2.73 | - |
| Bcl2l12 | 11.04 | 10.93 | 12.67 | 10.4 | 10.85 | 10.07 | 7.48 | 8.01 | 10.91 | 7.34 | 7.05 | 8.62 | - |
| Ssx2ip | 2.08 | 2.67 | 2.36 | 1.16 | 1 | 1.17 | 0.8 | 1.55 | 1.22 | 0.73 | 0.29 | 0.64 | - |
| Dusp3 | 16.99 | 17.53 | 17.55 | 14.45 | 15.06 | 15.1 | 17.52 | 19.7 | 16.98 | 14.5 | 13.76 | 14.25 | - |
| Dennd5a | 8.78 | 7.83 | 7.76 | 14.56 | 11.6 | 11.38 | 1.99 | 2.66 | 1.35 | 2.07 | 2.23 | 1.95 | - |
| Msl1 | 2.69 | 2.96 | 2.72 | 5.49 | 4.21 | 4.48 | 5.13 | 4.54 | 4.85 | 6.9 | 6.88 | 5.57 | - |
| Uchl1 | 25.88 | 17.71 | 20.88 | 13.94 | 14.53 | 13.01 | 25.75 | 35.68 | 35.69 | 12.82 | 9.73 | 15.28 | - |
| Rgs14 | 14.37 | 13.34 | 13.2 | 16.23 | 14.76 | 13.64 | 19.1 | 13.81 | 16.53 | 17.51 | 18.21 | 16.78 | - |
| Ppp3cc | 12.07 | 11.65 | 11.37 | 14.92 | 12.82 | 16.79 | 18.41 | 17.85 | 16.94 | 18.12 | 19.05 | 17.64 | - |
| Lhx6 | 0.05 | 0.02 | 0 | 0.1 | 0.18 | 0.12 | 0.16 | 0.06 | 0.15 | 0.2 | 0.35 | 0.18 | - |
| Ms4a6b | 120.55 | 127.53 | 125.28 | 98.01 | 79.28 | 82.38 | 135.46 | 112.03 | 117 | 96.25 | 76.09 | 116.69 | - |
| Ctla4 | 96.8 | 87.3 | 97.12 | 179.22 | 205.4 | 192.93 | 69.05 | 63.78 | 60.52 | 131.28 | 127.17 | 129.69 | - |
| 2410089E03Rik | 0.73 | 0.84 | 0.94 | 1.27 | 1.32 | 0.99 | 1.16 | 0.91 | 0.91 | 1.26 | 1.26 | 1.41 | - |
| 2010315B03Rik | 8.32 | 7.71 | 7.64 | 11.05 | 11.08 | 9.81 | 11.06 | 9.3 | 10.63 | 10.81 | 12.28 | 12.59 | - |
| Hemgn | 4.05 | 6.6 | 4.01 | 0.38 | 0.24 | 0.3 | 0.23 | 0.79 | 0.88 | 0 | 0.13 | 0.37 | - |
| Slc7a1 | 11.73 | 11.79 | 12.02 | 7.73 | 8.3 | 8.64 | 7.09 | 11.8 | 11.26 | 6.37 | 6.29 | 4.85 | - |
| Atg4d | 22.36 | 21.06 | 22.73 | 16.37 | 16.7 | 15.92 | 14.83 | 17.35 | 16.15 | 12.89 | 12.38 | 12.9 | - |
| Alg6 | 7.23 | 6.36 | 6.16 | 4.91 | 4.53 | 3.46 | 2.62 | 4.06 | 4.17 | 2.28 | 2.59 | 2.31 | - |
| Slc35f5 | 3.43 | 3.49 | 3.37 | 5.99 | 8.12 | 5.75 | 6.28 | 5 | 4.67 | 7.41 | 7.71 | 7.68 | - |
| Lipo1 | 10.18 | 7.95 | 8.23 | 4.86 | 8.6 | 7.66 | 5.34 | 6.86 | 5.86 | 4.31 | 6.02 | 4.17 | - |
| Atg14 | 2.1 | 2.06 | 1.36 | 4.27 | 4.14 | 3.73 | 6.14 | 3.9 | 4.24 | 5.98 | 7.91 | 7.5 | - |
| Nxpe4 | 0.04 | 0 | 0.02 | 0.1 | 0 | 0.04 | 0.2 | 0.02 | 0.05 | 0.63 | 0.14 | 0.36 | - |
| Endou | 0 | 0.07 | 0.04 | 0.26 | 0.49 | 0.03 | 0.76 | 0.07 | 0.25 | 1.21 | 1.03 | 0.86 | - |
| Fam149b | 2.39 | 2.74 | 2.75 | 3.46 | 5.58 | 5.09 | 5.58 | 5.16 | 6.08 | 6.46 | 7.08 | 6.7 | - |
| Zfp71-rs1 | 5.67 | 6.71 | 5.43 | 9.62 | 8.7 | 7.77 | 7.04 | 7.19 | 6.44 | 8.13 | 9.39 | 8.9 | - |
| Tubb2a | 22.84 | 18.39 | 19.65 | 15.24 | 14.87 | 14.39 | 17.1 | 16.35 | 17.44 | 11.26 | 11.59 | 12.62 | - |
| Chfr | 21.48 | 22.59 | 21.43 | 23.31 | 23.27 | 21.13 | 24.14 | 25.32 | 25.56 | 27.32 | 25.93 | 28.61 | - |
| Rufy2 | 1.64 | 1.23 | 1.67 | 2.69 | 2.12 | 2.09 | 1.95 | 1.93 | 1.56 | 2.74 | 3.03 | 2.94 | - |
| Gpr107 | 8.03 | 8.06 | 8.28 | 10.95 | 9.75 | 9.79 | 10.95 | 10.16 | 10.09 | 10.99 | 11 | 11.41 | - |
| Ube2c | 266.84 | 175.09 | 165.63 | 179.78 | 238.7 | 202.66 | 127.03 | 107.14 | 101.76 | 125.29 | 125 | 105.42 | - |
| Thyn1 | 59.36 | 57.73 | 60.88 | 41.45 | 43.88 | 43.35 | 36.13 | 48.47 | 42.85 | 29.44 | 31.43 | 28.25 | - |
| Alad | 15.83 | 14.84 | 14.38 | 13.53 | 12.51 | 13.4 | 12.97 | 12.92 | 11.96 | 10.09 | 10.52 | 10.89 | - |
| Spdl1 | 14.78 | 13.65 | 13.57 | 7.93 | 10.42 | 9.45 | 6.3 | 8.49 | 7.4 | 3.58 | 7 | 5.1 | - |
| Mbd4 | 3.08 | 3.5 | 3.01 | 3.87 | 5 | 4.45 | 4.55 | 3.95 | 3.59 | 4.51 | 4.25 | 5.04 | - |
| Wdr82 | 20.97 | 20.59 | 17.84 | 19.41 | 20.39 | 16.18 | 17.12 | 14.18 | 14.59 | 16.64 | 15.79 | 16.25 | - |
| Fam78a | 12.18 | 12.91 | 11.09 | 17.87 | 20.61 | 18.35 | 24.83 | 24.67 | 24.4 | 30.68 | 33.52 | 34.02 | - |
| Gpank1 | 17.68 | 18.32 | 16.79 | 25.43 | 21.63 | 22.4 | 31.72 | 22.29 | 27.02 | 34.56 | 32.89 | 35.46 | - |
| Sdhc | 61.64 | 58.34 | 55.86 | 50.66 | 54.43 | 55.6 | 56.69 | 59.1 | 58.61 | 43.11 | 50.55 | 47.18 | - |
| Npm3 | 13.91 | 11.8 | 10.67 | 8.47 | 10.34 | 6.84 | 7.27 | 8.49 | 9.68 | 6.6 | 7.68 | 5.63 | - |
| Pofut2 | 16.52 | 16.1 | 13.84 | 11.71 | 11.31 | 11.32 | 14.19 | 14.56 | 14.29 | 10.08 | 11.73 | 9.76 | - |
| Itgb1 | 12.14 | 14.23 | 13.24 | 11.42 | 10.31 | 10.51 | 4.27 | 6.04 | 6.14 | 5.57 | 4.35 | 4.3 | - |
| Il18 | 5.27 | 3.7 | 4.56 | 4.38 | 4.54 | 5.23 | 10.75 | 7.32 | 8.24 | 11.16 | 8.9 | 9.39 | - |
| Cnot3 | 10.44 | 11.36 | 12.26 | 15.24 | 14.5 | 13.44 | 14.76 | 14.71 | 14.85 | 15.72 | 14.05 | 14.64 | - |
| Rgs11 | 10.88 | 8.74 | 9.49 | 25.75 | 19.08 | 19.04 | 27.32 | 19.94 | 22.35 | 35.22 | 29.62 | 40.09 | - |
| Ppp2r5a | 2.71 | 3.52 | 2.98 | 4.54 | 3.54 | 3.64 | 4.99 | 3.17 | 4.53 | 5.81 | 4.97 | 5.61 | - |
| Cd70 | 20.48 | 19.59 | 16.89 | 3.38 | 5.71 | 6.06 | 2.93 | 8.09 | 6.12 | 2.79 | 1.61 | 2.46 | - |
| Bcorl1 | 1.15 | 1.13 | 1.27 | 1.54 | 1.89 | 1.78 | 2.25 | 1.29 | 1.77 | 1.56 | 2.4 | 2 | - |
| Gm4285 | 0.25 | 0.39 | 0.47 | 0.96 | 0.63 | 0.47 | 0.67 | 0.61 | 0.3 | 0.96 | 1.69 | 1.34 | - |
| Mpzl2 | 1.35 | 1.81 | 0.66 | 5.97 | 4.54 | 5.79 | 3.22 | 1.92 | 2.65 | 5.39 | 7.36 | 5.07 | - |
| Tet1 | 0.11 | 0.07 | 0.19 | 0.19 | 0.28 | 0.32 | 0.55 | 0.28 | 0.26 | 0.4 | 0.54 | 0.62 | - |
| Diap3 | 2.55 | 2.18 | 2.32 | 2.01 | 2.09 | 2.1 | 1.06 | 1.64 | 1.88 | 0.83 | 1.3 | 1.25 | - |
| Cramp1l | 1.2 | 1.48 | 1.44 | 1.8 | 1.51 | 1.74 | 1.61 | 1.3 | 1.57 | 2.03 | 2.43 | 1.86 | - |
| Zfp287 | 1.19 | 1.58 | 1.18 | 1.9 | 2.19 | 1.9 | 2.2 | 1.78 | 1.57 | 2.55 | 1.75 | 2.49 | - |
| Ankrd13b | 3.13 | 2.9 | 3.45 | 2.39 | 2.38 | 2.54 | 2.1 | 2.2 | 2.27 | 1.77 | 2.24 | 1.94 | - |
| Irf2bpl | 3.11 | 3.66 | 3.85 | 7.07 | 5.15 | 4.97 | 4.77 | 6.13 | 5.83 | 6.62 | 6.59 | 6.62 | - |
| Mtx2 | 65.56 | 74.38 | 79.41 | 57.53 | 64.01 | 58.71 | 43.95 | 61.09 | 61.63 | 43.69 | 47.96 | 38.78 | - |
| Ddx60 | 0.25 | 0.12 | 0.2 | 0.66 | 0.57 | 0.55 | 0.34 | 0.23 | 0.04 | 0.53 | 0.64 | 1.11 | - |
| Paqr4 | 3.41 | 3.31 | 3.35 | 2.17 | 3.48 | 3.01 | 3.56 | 2.46 | 3.01 | 1.84 | 2.11 | 2.42 | - |
| Slc35d3 | 4.3 | 3.72 | 3.87 | 3.87 | 4.39 | 3.78 | 2.86 | 4.53 | 3.39 | 2.28 | 2.71 | 2.59 | - |
| Fam210a | 4.68 | 4.56 | 4.77 | 3.27 | 3.13 | 3.51 | 3.14 | 3.45 | 3.36 | 2.44 | 2.48 | 2.14 | - |
| Snrnp35 | 9.04 | 5.4 | 9.06 | 4.28 | 5.95 | 5.02 | 5.26 | 4.68 | 4.72 | 3.61 | 4.52 | 3.85 | - |
| 2410131K14Rik | 6.71 | 6.86 | 7.07 | 3.15 | 4.51 | 4.59 | 4.62 | 6.53 | 5.91 | 4.69 | 3.16 | 3.61 | - |
| Hps1 | 10.55 | 11.53 | 9.4 | 16.27 | 16.07 | 14.94 | 18.74 | 14.29 | 17.81 | 17.97 | 18.93 | 20.57 | - |
| Hmgxb4 | 6.83 | 7.3 | 6.62 | 5.77 | 6.17 | 4.46 | 5.37 | 5.49 | 6.82 | 5.64 | 4.36 | 4.26 | - |
| Pigu | 32.47 | 37.64 | 33.09 | 24.54 | 23.26 | 25.61 | 24.31 | 21.96 | 27.64 | 20.08 | 19.01 | 19.77 | - |
| Tmem87b | 1.46 | 2.11 | 2.02 | 2.42 | 2.19 | 2.65 | 3.21 | 3.47 | 2.16 | 2.93 | 2.87 | 2.57 | - |
| Zfp831 | 0.09 | 0.05 | 0.1 | 0.22 | 0.06 | 0.05 | 0.17 | 0.14 | 0.19 | 0.29 | 0.21 | 0.21 | - |
| Cep55 | 27.57 | 24.61 | 23.94 | 19.88 | 27.95 | 23.05 | 16.58 | 14.91 | 14.67 | 16.01 | 16.14 | 15.67 | - |
| Aqp3 | 0.03 | 0.26 | 0.15 | 0.32 | 0.67 | 0.33 | 0.26 | 0.17 | 0.12 | 1.3 | 0.45 | 1.28 | - |
| Tnrc6b | 1.12 | 1.02 | 1.31 | 1.76 | 1.56 | 1.88 | 1.85 | 1.91 | 1.88 | 2.61 | 2.17 | 2.44 | - |
| Fra10ac1 | 24.54 | 23.01 | 22.01 | 16.91 | 16.46 | 17.65 | 18.17 | 25.45 | 20.53 | 15.55 | 17.02 | 12.41 | - |
| Tagap1 | 21.65 | 18.46 | 19.69 | 35.07 | 35.68 | 35.4 | 32.97 | 25.51 | 25.64 | 39.42 | 34.95 | 35.23 | - |
| Slc35a3 | 2.78 | 2.79 | 2.99 | 2.18 | 1.78 | 1.65 | 2.07 | 1.74 | 2.49 | 1.82 | 1.44 | 1.25 | - |
| Cdca8 | 91.99 | 83.53 | 80.78 | 68.49 | 81.8 | 77.11 | 45.34 | 54.45 | 46 | 44.36 | 53.49 | 39.56 | - |
| Sbf1 | 4.66 | 5.92 | 5.52 | 7.58 | 7 | 7.13 | 8.5 | 6.65 | 6.45 | 9.78 | 7.78 | 7.56 | - |
| Ola1 | 83.57 | 78.67 | 79.41 | 54.71 | 53.99 | 58.51 | 53.57 | 62.8 | 65.98 | 44.32 | 54.9 | 45.58 | - |
| Atm | 1.64 | 1.98 | 1.82 | 2.73 | 2.3 | 2.51 | 1.8 | 2.22 | 2.42 | 2.33 | 2.7 | 2.32 | - |
| Oaz2 | 14.95 | 18.96 | 15.69 | 26.56 | 24.88 | 20.85 | 24.43 | 22.87 | 25.06 | 27.39 | 26.37 | 27.79 | - |
| Lynx1 | 0.19 | 0.09 | 0.11 | 0.3 | 0.3 | 0.44 | 0.28 | 0.09 | 0.18 | 0.67 | 0.39 | 0.42 | - |
| Specc1l | 5.48 | 3.9 | 6 | 5.79 | 6.13 | 5.66 | 6.55 | 6.49 | 5.73 | 8.53 | 8.08 | 8.5 | - |
| Id3 | 9.82 | 10.99 | 9.19 | 12.07 | 14.28 | 15.06 | 3.56 | 4.01 | 4.28 | 6 | 5.17 | 4.17 | - |
| Rnf111 | 2.9 | 2.98 | 3.85 | 5.64 | 3.98 | 4.17 | 5.03 | 4.28 | 4.42 | 6.22 | 5.35 | 5.45 | - |
| Gas5 | 133.62 | 130.06 | 129.84 | 200.65 | 206.14 | 218.2 | 138.52 | 166.37 | 151.33 | 197.41 | 214.14 | 174.14 | - |
| Insl6 | 47.83 | 38.26 | 38.91 | 55.26 | 56.65 | 58.94 | 53.39 | 67.56 | 56.01 | 67.95 | 65.09 | 65.93 | - |
| Dhx40 | 7.13 | 7.34 | 5.84 | 10.66 | 8.6 | 8.33 | 11.51 | 10.27 | 10.51 | 14.13 | 15.84 | 15.84 | - |
| Ptprn | 0.87 | 0.46 | 0.86 | 0 | 0.12 | 0.14 | 0.1 | 0.2 | 0.44 | 0.1 | 0.02 | 0.11 | - |
| Ascc2 | 27.26 | 25.43 | 24.51 | 24.65 | 22.77 | 25.35 | 20.21 | 27.81 | 26.02 | 16.87 | 18.79 | 15.74 | - |
| Bub1b | 36.05 | 31.81 | 30.65 | 26.24 | 29.94 | 26.92 | 16.71 | 18.87 | 14.62 | 13.95 | 19.99 | 14.97 | - |
| Zfp54 | 8.81 | 8.04 | 7.62 | 9.94 | 9.09 | 13.39 | 10.1 | 10.13 | 9.54 | 10.45 | 11.91 | 11.52 | - |
| B3gnt9 | 1.45 | 3.2 | 1.55 | 5.28 | 5.07 | 4.36 | 6.31 | 4.71 | 6.21 | 7.12 | 5.8 | 8.62 | - |
| Tor1b | 10.89 | 10.78 | 11.02 | 8.29 | 10.04 | 8.98 | 7.8 | 9.15 | 7.38 | 6.96 | 6.72 | 8.4 | - |
| Ceacam15 | 0.16 | 0.46 | 0.35 | 3.59 | 3.95 | 3.48 | 0.21 | 0.59 | 0.12 | 2.06 | 1.02 | 2.28 | - |
| Fbxo18 | 18.13 | 16.18 | 15.51 | 19.87 | 21.37 | 20.82 | 20.6 | 19.15 | 21.71 | 21.17 | 19.98 | 22.08 | - |
| Poglut1 | 24.66 | 23.1 | 21.89 | 20.12 | 22.49 | 21.66 | 17.3 | 23.06 | 18.46 | 17.78 | 15.43 | 18.44 | - |
| Spsb1 | 0.15 | 0.15 | 0.11 | 1.26 | 0.78 | 0.75 | 0.55 | 0.13 | 0.54 | 1.55 | 1.51 | 0.83 | - |
| Rcbtb1 | 5.92 | 5.67 | 5.82 | 8.32 | 7.87 | 7.37 | 8.2 | 8.8 | 8.11 | 9.82 | 8.64 | 7.55 | - |
| Itpr2 | 0.87 | 0.65 | 0.72 | 1.52 | 1.12 | 1.11 | 1.62 | 1.26 | 1.38 | 1.69 | 1.66 | 1.67 | - |
| Ndufa3 | 180.58 | 189.41 | 171.96 | 166.65 | 178.68 | 167.2 | 176.28 | 147.42 | 153.73 | 139.82 | 137.04 | 147.16 | - |
| Grn | 26.79 | 27.28 | 28.13 | 38.85 | 33.78 | 36.17 | 44.95 | 36.36 | 35.83 | 38.67 | 41.4 | 41.29 | - |
| 5730508B09Rik | 0.53 | 0.54 | 0.4 | 1.09 | 0.82 | 0.99 | 1.61 | 1.36 | 1.71 | 4.14 | 2.42 | 2.73 | - |
| Inpp1 | 7.63 | 8.27 | 8.05 | 12.93 | 8.8 | 12.84 | 17.73 | 13.61 | 11.61 | 20.32 | 17.27 | 20.34 | - |
| Gmpr2 | 56.9 | 51.78 | 53.05 | 38.56 | 43.84 | 40.89 | 34.27 | 41.01 | 38.61 | 30.78 | 31.65 | 34.27 | - |
| Ndufa12 | 249.43 | 254.77 | 241.53 | 137.3 | 159.46 | 158.46 | 132.04 | 180.59 | 151.95 | 102.32 | 113.04 | 106.3 | - |
| Lxn | 36.86 | 36.45 | 35.15 | 26.28 | 25.67 | 27.21 | 38.36 | 36.9 | 37.14 | 24.05 | 24.07 | 31.51 | - |
| Zranb2 | 62.78 | 62.12 | 65.11 | 42.62 | 49.15 | 41.26 | 37.3 | 44.71 | 40.34 | 31.42 | 28.72 | 32.36 | - |
| Sp100 | 32.06 | 34.39 | 31.09 | 44.52 | 38.8 | 41.04 | 54.48 | 39.85 | 43.45 | 72 | 59 | 73.53 | - |
| Mcrs1 | 38.26 | 37.01 | 35.61 | 28.31 | 29.53 | 29.84 | 24.47 | 33.75 | 31.2 | 27.25 | 25.37 | 26.91 | - |

| | | | | | | | | | | | | | |
|---|---|---|---|---|---|---|---|---|---|---|---|---|---|
| Igfbp2 | 17.03 | 19.73 | 29.95 | 1.33 | 1.13 | 1.25 | 1.6 | 3.85 | 0.97 | 0 | 0.14 | 0.96 | - |
| Ispd | 0.47 | 0.26 | 0.34 | 0.21 | 0.14 | 0.05 | 0 | 0.11 | 0.04 | 0.02 | 0 | 0 | - |
| Pcyt1a | 11.68 | 10.98 | 10.82 | 5.76 | 5.55 | 5.84 | 5.9 | 9.86 | 7.67 | 5.45 | 5.03 | 4.2 | - |
| Mrps15 | 88.74 | 91.72 | 78.84 | 60.83 | 71.54 | 66.83 | 71.73 | 72.76 | 71.5 | 50.86 | 63.6 | 62.81 | - |
| Ern1 | 0.78 | 0.76 | 1.2 | 1.18 | 1.29 | 0.96 | 1.39 | 1.65 | 1.31 | 1.57 | 1.84 | 1.42 | - |
| Nelfe | 55.97 | 60.1 | 59.86 | 44.52 | 46.22 | 46.4 | 44.98 | 44.74 | 42.11 | 33.84 | 34.34 | 35.93 | - |
| Sp3 | 7.65 | 11.13 | 11.25 | 15.12 | 12.51 | 11.71 | 13.02 | 11.28 | 13.53 | 14.81 | 17.51 | 14.61 | - |
| Snx11 | 7.74 | 6.75 | 6.89 | 4.46 | 4.74 | 5.31 | 4.39 | 6.17 | 5.92 | 4.37 | 3.65 | 4.49 | - |
| Ahnak | 20.62 | 20.28 | 18.88 | 27.26 | 25.1 | 26.14 | 36.41 | 30.77 | 33.01 | 42.44 | 39.59 | 45.39 | - |
| Csnk1g2 | 5.5 | 5.45 | 5.31 | 6.56 | 6.56 | 6.23 | 7.21 | 5.58 | 5.93 | 7.15 | 7.62 | 9.21 | - |
| Trim46 | 3.55 | 3.58 | 2.84 | 7.5 | 6.01 | 7.89 | 8.68 | 4.75 | 5.36 | 9.26 | 9.42 | 8.12 | - |
| Crip1 | 184.68 | 193.92 | 191.41 | 215.2 | 215.34 | 218.59 | 283.6 | 210.79 | 234.46 | 255.32 | 234.61 | 274.16 | - |
| Tenc1 | 0.18 | 0.11 | 0.14 | 0.34 | 0.12 | 0.08 | 0.7 | 0.4 | 0.58 | 0.22 | 0.69 | 1.04 | - |
| Dnmt3a | 3.87 | 3.65 | 3.16 | 2.64 | 2.4 | 1.94 | 2.02 | 2.41 | 2.55 | 2.28 | 1.5 | 2.02 | - |
| Chit1 | 0.77 | 0.17 | 0.57 | 0.63 | 0.33 | 0.45 | 3.16 | 3.63 | 4.01 | 2.07 | 3.71 | 2.53 | - |
| E2f2 | 4.25 | 4.43 | 4.04 | 5.58 | 4.97 | 4.58 | 6.14 | 4.8 | 4.34 | 6.11 | 5.62 | 6.8 | - |
| Ctsb | 61.19 | 55.45 | 58.64 | 89.45 | 77.45 | 85.25 | 89.75 | 81.85 | 85.17 | 99.66 | 106.61 | 104.17 | - |
| Ctsw | 36.51 | 43.36 | 26.58 | 104.73 | 71.91 | 72.42 | 151.5 | 77.13 | 101.38 | 172.22 | 145.02 | 211.64 | - |
