## Supplementary material for "An endogenous glucocorticoid-cytokine signaling circuit promotes CD8^+^ T cell dysfunction in the tumor microenvironment": Table S2

| Exhaustion_up | Exhaustion_down |
|---|---|
| Chst12 | Dnajc7 |
| Timp1 | Fas |
| Cysltr2 | Samhd1 |
| Alg8 | Map4k2 |
| Gsg2 | Urb1 |
| Aplp1 | Pus7l |
| Kif22 | Man1c1 |
| Ccdc109b | Amz2 |
| Nup107 | Lrp12 |
| Crat | Rftn1 |
| Ifng | Il4ra |
| Trib3 | Klf9 |
| Rabgef1 | Idua |
| 5330426P16Rik | Clasp1 |
| Ccl4 | Snhg7 |
| Gpr160 | Psap |
| Serpine2 | Bcl3 |
| Rpa2 | Zscan12 |
| Arsb | Pcca |
| Enpp2 | Gpd1l |
| Nsmaf | Tmem50b |
| Cdca3 | Tcp11l2 |
| Cenpn | Dhcr24 |
| Cdc27 | Rarg |
| Slc35b1 | Zer1 |
| Il1r2 | Idh2 |
| Shcbp1 | Atp2a1 |
| Dclk1 | Nfia |
| Kpna2 | Vdr |
| Esco2 | Mtap |
| Ppp1r3b | Gramd1a |
| Eno3 | Cmah |
| Ero1l | Npc1 |
| Dennd4a | Gpr18 |
| Sepn1 | Nefh |
| Mtmr1 | Rbm38 |
| Abcb1b | Macrod1 |
| Birc5 | Zc3h12d |
| Magohb | Gimap6 |
| Samsn1 | Cd24a |
| Zfp692 | Slfn5 |
| Gcsh | Slco3a1 |
| Ccl9 | Bdh1 |
| Zwilch | Noc4l |
| Spp1 | Ddr1 |
| Zc3h12a | Ctsl |
| Tmbim4 | Btla |
| Tmem171 | Epcam |
| Casp4 | Gramd4 |
| Tnfrsf18 | Cryl1 |
| Ube2i | Mboat1 |
| Mthfs | Igfbp4 |
| Plekhb2 | Bphl |
| Sdf4 | Atp10d |
| Ccl3 | Rere |
| Serpinb6a | Afap1 |
| Anapc4 | Hs3st3b1 |
| Ifitm3 | St6gal1 |
| Ndfip2 | Ppa1 |
| Sec23a | Ung |
| Stard4 | Mbp |
| Fasl | Rpp40 |
| Ccnb1 | Arhgef3 |
| Nek8 | Satb1 |
| Myadm | Cxcr3 |
| Tmem48 | Exosc1 |
| Cenpt | Sf3a2 |
| Gm11110 | Acss2 |
| Ttk | Iigp1 |
| Arhgap18 | 2610035D17Rik |
| Slc7a3 | Als2cl |
| Chac1 | Rnf157 |
| Armc1 | Jhdm1d |
| Cars | Tpd52 |
| Mlf1 | Kti12 |
| Gab2 | Ralgps2 |
| Cenpa | Prnp |
| Tmem218 | Itgae |
| Nrp1 | Atp1b3 |
| Sephs1 | Ecm1 |
| Prdm1 | Sema4f |
| Sh3bgrl | Ddx54 |
| Filip1 | Pde4b |
| Agl | Aldh6a1 |
| Qsox1 | Socs1 |
| Plek | Tlr1 |
| Adam9 | Camkk1 |
| Igf2r | Tspan13 |
| Pkp2 | Vmac |
| Npnt | Med4 |
| Kif20a | Sorl1 |
| Cep170 | Taf5l |
| Idh3a | Tecpr1 |
| Pmaip1 | Fads2 |
| Wbp5 | Kbtbd11 |
| Prf1 | Hagh |
| Lgals3 | AB124611 |
| Gdpd5 | Sesn3 |
| Ccdc50 | Serinc5 |
| Lpgat1 | Fosl2 |
| Tyw1 | Vav2 |
| Zbtb32 | Tax1bp3 |
| BC068157 | Map4k4 |
| Arhgef9 | Gatsl3 |
| Nrn1 | Rras2 |
| 2310001H17Rik | 3110043O21Rik |
| Tacc3 | Lcmt2 |
| Spry2 | Rgs10 |
| Slco4a1 | Me2 |

| | |
|---|---|
| Htatip2 | Ifnar2 |
| Nr4a2 | Itga7 |
| Trappc4 | Clpb |
| Abhd14a | Mppe1 |
| Nupr1 | Rbm19 |
| AA467197 | Ccpg1 |
| Polk | Dzip1 |
| Anxa3 | Pdlim5 |
| Cnnm2 | Ctdsp2 |
| Ndrg1 | Prcp |
| Lrp1 | Cdt1 |
| Slc16a13 | Zcchc11 |
| Sh2d2a | Ptcra |
| Dnajc24 | Ifngr2 |
| Gins1 | Med8 |
| Cox17 | Insr |
| Zfp52 | Slc16a6 |
| Gata3 | 1810043H04Rik |
| Dynlt3 | Stk38 |
| Gtf2ird1 | Ascc1 |
| Vldlr | Zdhhc13 |
| Dctn4 | Arid5a |
| Syngr3 | Ddx18 |
| Asns | 2610002M06Rik |
| Uhrf2 | Tfam |
| Ino80c | Phc1 |
| Ppap2c | Parp9 |
| Pggt1b | Sqrdl |
| Cd93 | Gtf2i |
| Ttc39c | Lmo4 |
| Tubb6 | Gpx1 |
| Dab2 | Tmem141 |
| Zdhhc5 | Trim62 |
| Usp46 | Icam2 |
| Mrps6 | Irf7 |
| Kctd11 | Utrn |
| Gnb5 | Cd40lg |
| B4galt5 | Smc6 |
| Gzmc | Zfp260 |
| Tk1 | Dhx37 |
| Rhoq | Trib2 |
| Gja1 | Ift80 |
| 3110082I17Rik | Klhdc1 |
| Osbpl3 | Nop2 |
| Tm9sf3 | Umps |
| Smc2 | Pim2 |
| Cenpp | Igf1r |
| Copz2 | Osbpl9 |
| Klf10 | Cdc14b |
| Degs1 | Ccr4 |
| Nbeal1 | Tom1 |
| Cdca5 | Fgr |
| 4932415G12Rik | Slamf6 |
| Adam15 | Sgip1 |
| Mpi | Sigmar1 |
| Dapk2 | Uvrag |
| Plk1 | Sdccag3 |
| Hip1r | Traf4 |
| Amigo1 | Lpin1 |
| Optn | Ipcef1 |
| 1700017B05Rik | Rab37 |
| Ctsc | Top1mt |
| Ltbp3 | Chst15 |
| Mrps36 | Ehd3 |
| Atf4 | Zkscan3 |
| Rab5b | Rcn3 |
| Acadl | Tnfsf8 |
| Pld2 | Abcc4 |
| Ptrf | Ephx1 |
| Nqo2 | Peo1 |
| Scyl2 | A230046K03Rik |
| Myo1e | P2rx4 |
| Tmem39a | Socs3 |
| Nuf2 | Zeb1 |
| Rnaseh2c | Frmd4b |
| Spag9 | Skp1a |
| Atad5 | Egln3 |
| Ncaph | Mapk1ip1 |
| 2610318N02Rik | Tcf7 |
| Traip | Adk |
| Gem | Cyp2s1 |
| Elk3 | Pepd |
| Egr1 | Rcl1 |
| Adssl1 | Pde7a |
| 1110007C09Rik | Tmem41a |
| Icos | St8sia1 |
| Tbc1d7 | Hspb11 |
| Itgav | Irf6 |
| C330027C09Rik | Pik3ip1 |
| Anxa4 | Polr3a |
| Entpd1 | Lrrc33 |
| Omd | Nxf1 |
| Pter | Zbtb24 |
| Ccdc14 | Itgb7 |
| Apbb1 | Fam117b |
| Gpld1 | Zfp362 |
| Gabbr1 | Nufip2 |
| Oit3 | Sgk3 |
| Hmmr | Arl6ip5 |
| Tmem159 | Yes1 |
| Cenph | Capn3 |
| Ndufaf2 | Cpne2 |
| Kdm2b | Dip2c |
| Ptplad1 | Serp1 |
| Aldoc | Jmy |
| Mmd | Zfp281 |
| Atg16l2 | Zmym2 |
| Ephb6 | 2410066E13Rik |
| Aqp11 | Znrf1 |
| Nedd9 | Gypc |
| Dkkl1 | Tanc1 |

| | |
|---|---|
| Abhd4 | Ikbke |
| Pddc1 | Cyth1 |
| Exo1 | Ulk1 |
| Slc37a2 | Ttc32 |
| Impa2 | Etv3 |
| Farp1 | Rnf122 |
| Gpr174 | Plaur |
| Rps6kc1 | Gpr155 |
| Cyp20a1 | Rsad2 |
| Tmem126a | Tctex1d1 |
| Fam188a | Rnf149 |
| Hif1a | Tdrkh |
| Mtmr7 | Egr2 |
| Casp3 | Cxcl10 |
| Dut | Mmachc |
| Stab1 | Rab35 |
| Ctla2a | Cd7 |
| E130112N10Rik | Rapgef6 |
| 2810417H13Rik | Trat1 |
| Pilra | Prkcq |
| Eno2 | Ncf1 |
| Tnfrsf4 | Tm4sf5 |
| Ptgr1 | Vrk1 |
| Ptprk | Pus3 |
| Kit | Frat2 |
| Crabp2 | Acp5 |
| Tecr | Ldhb |
| Bcat1 | Commd8 |
| Tmem180 | Pou6f1 |
| Zfp760 | Fam53b |
| Il2ra | Gpr132 |
| Ptpn11 | Dhx58 |
| Kcna4 | Card11 |
| Ovol2 | Kctd12 |
| Creb3l2 | Acpp |
| Pla1a | Cxcr4 |
| Rgs16 | Dus2l |
| Lag3 | Dntt |
| Rai14 | Hpcal1 |
| Pask | 2810013P06Rik |
| Sestd1 | Tpst2 |
| Tulp4 | Aff3 |
| Utf1 | Dapl1 |
| Cst7 | Aqp9 |
| Hnrpll | Wdr59 |
| Tnfsf13b | Mll3 |
| Cercam | Ado |
| Ptgs2 | Slc9a9 |
| Ankra2 | Pja1 |
| Slc25a13 | Hmgn1 |
| Cyp4v3 | Eif2ak3 |
| Pros1 | Etnk1 |
| Alcam | Abtb2 |
| Fars2 | 2900076A07Rik |
| Elmo2 | Srpk1 |
| Mt2 | D1Ertd622e |
| Fcer1g | Glcci1 |
| Spin4 | Ceacam1 |
| Rpgr | Heatr1 |
| Usp40 | Itpr3 |
| Styk1 | Gucy1a3 |
| Ccdc6 | Usp53 |
| Bend6 | Zfp1 |
| AF529169 | Phf17 |
| Ppme1 | Brap |
| Mrgpre | BC026590 |
| Mtbp | Pqlc1 |
| Exph5 | Pprc1 |
| Slc17a6 | Arntl |
| Ankle1 | Tlr6 |
| Cd38 | Kcnn4 |
| Vamp8 | Net1 |
| Nipa2 | Foxp1 |
| Gpt2 | Tha1 |
| Fhl2 | Kif21b |
| Gatm | Vrk3 |
| Id2 | Irgm2 |
| Nr3c1 | Herc1 |
| Cenpi | Unc5cl |
| Gzme | Cd302 |
| Itga5 | Usp28 |
| Csf2 | Pgap3 |
| Slc22a15 | Gpr114 |
| Sema6d | Dedd2 |
| Slc2a8 | Sgms1 |
| Mettl7a1 | Mrpl30 |
| Cd200 | Btbd6 |
| Tjp2 | Fam65b |
| Tox | Rasa3 |
| Suv39h2 | Bzw2 |
| Mlkl | Dirc2 |
| 2900026A02Rik | Nudt15 |
| Il1rl1 | Dnajc27 |
| Higd1a | Parp3 |
| Car13 | Mylip |
| Ica1 | Gbp2 |
| Kazald1 | Alpl |
| Kif20b | Ssh2 |
| Nfatc1 | Phc2 |
| Cd200r4 | Furin |
| Rab31 | Eef2k |
| Rad54b | Notch1 |
| Dusp6 | Wdr45 |
| Arl6 | Tcf4 |
| Bcl2l1 | Ccr7 |
| Mt1 | Ssbp2 |
| Camk2n1 | Axin2 |
| Fam5c | 2310044G17Rik |
| Sat1 | Npc2 |
| Rbpj | Laptm4b |
| Lamc1 | Gimap9 |

| | |
|---|---|
| Agtrap | P2ry14 |
| Itga1 | Fam43a |
| Cxcr6 | Mcoln3 |
| Galc | Dclre1a |
| 1190002F15Rik | Fgfr1op |
| Bst1 | Tnik |
| L2hgdh | Adrb2 |
| Lyrm1 | Evl |
| Carhsp1 | Pkd1 |
| Tpi1 | 4921511C10Rik |
| Mthfd2 | Gramd3 |
| Pik3cg | Prickle1 |
| Fcho2 | Gpc1 |
| Chn2 | Ccbl1 |
| Ncapg2 | Iqgap2 |
| Apobec3 | Klrb1f |
| Jam2 | Cdyl2 |
| Dtl | B430306N03Rik |
| Aldoa | Rasgrp2 |
| Tnf | Tk2 |
| Ulbp1 | Card6 |
| D630039A03Rik | Gpr146 |
| Mapk6 | Dph5 |
| 1500009L16Rik | Crim1 |
| Igfbp7 | Fgfr1 |
| Kctd17 | Fbxo17 |
| Pex11b | Ctps2 |
| Bnip3 | Nbn |
| Xpot | Dse |
| Unc119b | Xpc |
| Rnf128 | Add3 |
| Ehd4 | Pex26 |
| Rhbdf2 | Bach1 |
| Ptplb | Sfrp2 |
| Rab12 | Irf1 |
| Tg | Enpp4 |
| Casp1 | Thra |
| Tpk1 | Gab3 |
| Plscr1 | Bend5 |
| Ftsjd1 | Tmem55b |
| Slc43a3 | Dtx3l |
| Gpr56 | Arhgap15 |
| 4930422G04Rik | Itfg3 |
| Trip10 | Gm8369 |
| Irf4 | Acsl1 |
| Upp1 | Faah |
| Ctsd | Armcx6 |
| Padi2 | Smad3 |
| Stx11 | Zfp652 |
| Ide | Lrrc8a |
| Tmem120b | Slc11a2 |
| Prrx1 | Dus4l |
| Tpx2 | Il6st |
| Ect2 | Hey1 |
| Bcl2l15 | Atp8a2 |
| Fam57a | Smpdl3a |
| Acot7 | 3230401D17Rik |
| Mylk | Mycbp2 |
| Pdpn | Zfp354c |
| Rfc3 | Fbrs |
| Bard1 | Klhdc5 |
| 1810011H11Rik | Ptpla |
| Cisd3 | Kbtbd8 |
| Cenpf | St3gal6 |
| Ccnb2 | Dtx1 |
| Gm14288 | Cd2 |
| Ubash3b | Dym |
| Glis1 | Shisa2 |
| Cstad | Camkk2 |
| Fkbp7 | Pde2a |
| Stk39 | Slc41a1 |
| Ap1s2 | Sepp1 |
| Krt18 | Atp2a3 |
| Camk4 | Mgst2 |
| Spata2l | Sipa1l1 |
| Trps1 | Chd8 |
| Flywch1 | Lef1 |
| Gbe1 | Tapt1 |
| Fam64a | Il7r |
| 1700012B09Rik | Ramp1 |
| Etv5 | Elk4 |
| Dennd3 | Zfp53 |
| Nt5dc2 | Csrnp1 |
| Sar1a | Tgfbr3 |
| Sult2b1 | Lypd6b |
| Sytl3 | Kremen1 |
| Cdc14a | Bcl9 |
| Tmem170b | Emb |
| E2f3 | Arl6ip6 |
| Srgap3 | Rnf19b |
| Zfp511 | Foxo1 |
| Ostf1 | Fbxo32 |
| Fam72a | Scml4 |
| Endod1 | Ms4a4c |
| Raph1 | Jakmip1 |
| Nsl1 | Jak2 |
| Apoe | P4ha1 |
| Hivep1 | Klhl6 |
| Wisp1 | Frat1 |
| Tspan6 | Hmgxb3 |
| Csda | Sos1 |
| Pglyrp1 | S1pr4 |
| Serpinb6b | Crtc3 |
| Ppp3cb | Nisch |
| B3galt5 | Ifit1 |
| Ier3 | Tnfaip8l2 |
| Ddx17 | Fitm2 |
| Osgin1 | Atp1a3 |
| Foxm1 | Slfn1 |
| Calu | Jmjd1c |
| Rad51 | Cpne3 |

| | |
|---|---|
| Anxa2 | Gcnt2 |
| Ndc80 | Slc12a7 |
| Kif4 | Abca1 |
| Med12l | Bcl6 |
| Efcab7 | Fam189b |
| Havcr2 | Il1rl2 |
| Lclat1 | Bach2 |
| Atp10a | Mdn1 |
| Slc2a3 | Lif |
| Rapsn | Prkcb |
| Gpd2 | Kmo |
| Depdc1a | Pcnxl3 |
| Calm3 | Zrsr1 |
| Sdcbp2 | Myc |
| Fkbp1a | Scpep1 |
| Rab27a | As3mt |
| Plekhf1 | Zfp592 |
| Nid1 | Dopey1 |
| Plod2 | Egr3 |
| E2f8 | Zfp87 |
| Kif24 | Smad1 |
| Iqgap3 | Dgka |
| St14 | Arhgef18 |
| Tnfsf11 | Rapgef4 |
| Elovl4 | Zfp365 |
| Ifitm2 | Pitpnc1 |
| S100a11 | Gcc2 |
| Adam8 | Lysmd1 |
| Tnfsf9 | Ubac2 |
| 0610010F05Rik | Ldlrap1 |
| Dusp4 | Hemgn |
| Ermp1 | Tubb2a |
| Hbegf | Ddx60 |
| Npl | Aqp3 |
| Cyr61 | Id3 |
| Armc7 | Spsb1 |
| Emilin2 | Itpr2 |
| Ndufb4 | Grn |
| Hhat | 5730508B09Rik |
| E2f7 | |
| Slc30a9 | |
| Klra7 | |
| Neil3 | |
| Abcb9 | |
| Rab19 | |
| AI836003 | |
| Tnfsf4 | |
| Selm | |
| Siglec5 | |
| Actr10 | |
| Il2 | |
| Cldn12 | |
| Il33 | |
| Tmem175 | |
| Tmcc3 | |
| Piwil2 | |
| Pear1 | |
| Nmt2 | |
| Lat2 | |
| Phf23 | |
| Abi2 | |
| Gldc | |
| Lpxn | |
| Cdkn3 | |
| Fancd2 | |
| Atxn1 | |
| Cnrip1 | |
| Galnt3 | |
| Spag5 | |
| Brca2 | |
| Kif18a | |
| Sytl2 | |
| Plekho2 | |
| Tbc1d2b | |
| Mid1ip1 | |
| A430093F15Rik | |
| Gzmd | |
| Penk | |
| Pdcd1 | |
| Litaf | |
| Bub1 | |
| Ccng1 | |
| Ccl1 | |
| 4930429F24Rik | |
| Dusp3 | |
| Ctla4 | |
| Alg6 | |
| Slc35f5 | |
| Ube2c | |
| Mpzl2 | |
| Diap3 | |
| Paqr4 | |
| Cep55 | |
| Cdca8 | |
| Oaz2 | |
| Bub1b | |
| Lxn | |
| Pcyt1a | |
| Ern1 | |
